# DNA-encoded library (DEL)-enabled discovery of proximity-inducing small molecules

**DOI:** 10.1101/2022.10.13.512184

**Authors:** Jeremy W. Mason, Liam Hudson, Matthias V. Westphal, Antonin Tutter, Gregory Michaud, Wei Shu, Xiaolei Ma, Connor W. Coley, Paul A. Clemons, Simone Bonazzi, Frédéric Berst, Frédéric J. Zécri, Karin Briner, Stuart L. Schreiber

## Abstract

Molecular glues and bifunctional compounds that induce protein–protein associations provide a powerful and general mechanism to modulate cell circuitry. We sought to develop a platform for the direct discovery of compounds able to induce association of any two pre-selected proteins, using the first bromodomain of BRD4 and the VHL–elongin C–elongin B (VCB) complex as a test system. Leveraging the screening power of DNA-encoded libraries (DELs), we synthesized ∼one million DNA-encoded compounds that possess a VHL-targeting fragment, a variety of connectors, and a diversity element generated by split-and-pool combinatorial chemistry. By screening our DEL against BRD4^BD1^ in the presence and absence of VCB, we could identify VHL-bound molecules that simultaneously bind BRD4. For highly barcode-enriched library members, ternary complex formation leading to BRD4 degradation was confirmed in cells. Furthermore, a ternary complex crystal structure was obtained for the most enriched library member. Our work provides a foundation for adapting DEL screening to the discovery of proximity-inducing small molecules.

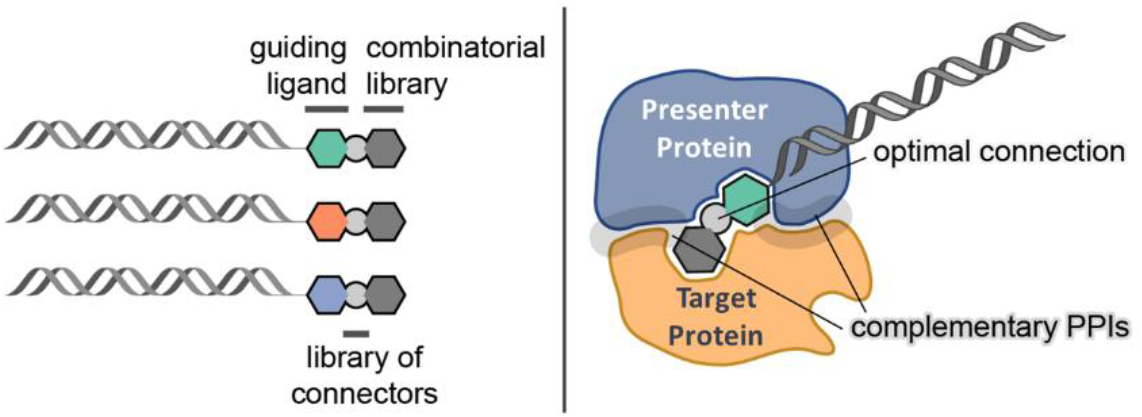

## Introduction

The discovery of small molecules that can modulate specific protein–protein interactions (PPIs) has long been of interest to the biomedical field and, although challenging, provides broad therapeutic opportunities.^1^ While the discovery of PPI inhibitors has become an established paradigm in drug discovery, the identification of compounds that induce novel (neo)-protein associations has been gaining momentum. The therapeutic relevance of such compounds, referred to here as chemical inducers of proximity (CIPs), was established with the realization that the drugs cyclosporin (sandimmune),^2^ FK506 (tacrolimus),^2^ and later rapamycin (sirolimus)^3,4^ work by such a mechanism. Initially, the perceived biophysical demands of these molecular glues were considered too great to be replicated by simple synthetic compounds (*vs* requiring natural selection over eons). However, the development of the first bifunctional compounds, which target genetic fusion proteins, provided a work-around to CIP identification and demonstrated the breadth of cellular processes that can be modulated by CIPs, especially by inducing association of neo-substrates with cellular enzymes.^5^ The slow but steady identification of simpler synthetic molecules that induce neo-associations of native proteins, such as tool compounds synstab A^6^ and indisulam,^7,8^ and drugs thalidomide and lenalidomide (Revlimid),^9,10^ have ignited widespread interest in the development of proximity-inducing compounds.

Notable areas of interest include the development of CIPs that promote targeted protein degradation of native proteins. For example, heterobifunctional degraders known as proteolysis-targeting chimeras (PROTACs), recruit proteins to an E3 ubiquitin ligase complex for targeted degradation and have gained broad interest in drug discovery.^11,12^ Lysosome-targeting chimeras have been introduced for recruiting an extracellular protein of interest (POI) to cell-surface receptors that traffic cargo to the lysosome for degradation.^13-15^ Mirroring the use of CIPs with genetic fusion proteins, additional emerging applications include targeting proteins to kinases^16^ or phosphatases,^17,18^ recruiting deubiquitinases to a target protein for stabilization,^19^ and recruiting histone acetyltransferases for targeted protein acetylation. ^20^

While the potential applications of CIPs continue to expand, the available tools for identifying them remain limited. As such, we set out to develop a platform that could accelerate the discovery of CIPs for any two pre-selected proteins, which we herein label as targets and presenters. We hoped to develop a platform where both target and presenter proteins would be screened simultaneously to identify bifunctional compounds that induce optimal ternary complex formation in a single step, rather than requiring independent efforts to optimize separate binding and linking events. The PROTAC concept was selected for our initial implementation, and we targeted a library of bifunctional compounds that could in principle recruit any desired protein (the target) to the E3 ubiquitin ligase VHL (the presenter) for ubiquitination and subsequent degradation. The transcriptional and epigenetic regulator BRD4 was chosen as the target for our proof-of-concept study; however, this library can be used to screen other targets of interest for recruitment to VHL.

DNA-encoded libraries (DELs), which are used to identify binders to target proteins,^21^ have not yet been adapted to discover compounds that induce ternary complex formation. We reasoned that a DNA headpiece could be functionalized with a VHL-binding fragment and a library of connectors, and the connectors could then be functionalized using the split-and-pool synthesis strategy with DNA barcoding. Next, we could use affinity screens with and without the VHL–elongin C–elongin B (VCB) complex to identify compounds that bind the target protein in the presence of VHL. This approach was especially appealing since it enables identification of CIPs that induce optimal ternary complex assembly in a single process. Here we present our first implementation of the “CIP-DEL” approach and report the successful validation of the platform through the identification of novel protein degraders.

## Results

### Library design and synthesis

The established BRD4 degrader **MZ1** was used as a template for our initial library design and the available ternary complex structure (PDB: 5T35) was used to select an attachment vector for the requisite DNA barcodes (**Fig. 1a and 1b**).^22,23^ The thiazole-methyl group of the VHL-targeting moiety of **MZ1** resides within a small channel directed towards solvent. For our library, the VHL-targeting portion of **MZ1** was synthesized with an alkyne functionality appended to the thiazole-methyl for copper(I)-catalyzed azide-alkyne cycloaddition (CuAAC) coupling (click reaction) to the DNA headpiece. This modified VHL ligand was functionalized with a panel of connectors having varied lengths and compositions. We focused primarily on shorter heterocyclic connectors to bias our screens towards CIPs that induce a close association between VHL and the POI, although traditional alkyl and PEG-based linkers were also included (**Fig. 1d; Supplementary Fig. S2**). To ensure that our VHL ligand–connector combinations did not disrupt VHL binding, a critical feature for our screens, we tested 22 ligand–connector combinations in a TR-FRET assay for VCB binding after the Fmoc-protected amines were converted to acetamides. We identified a collection of 15 connectors that displaced a HIF1α derived peptide from the VCB complex with EC50 values of ∼1 µM (range: 0.75 – 1.51 µM). These connectors were selected for library inclusion, coupled to the DNA headpiece, and then encoded by the addition of a unique DNA barcode.

**Figure 1.**
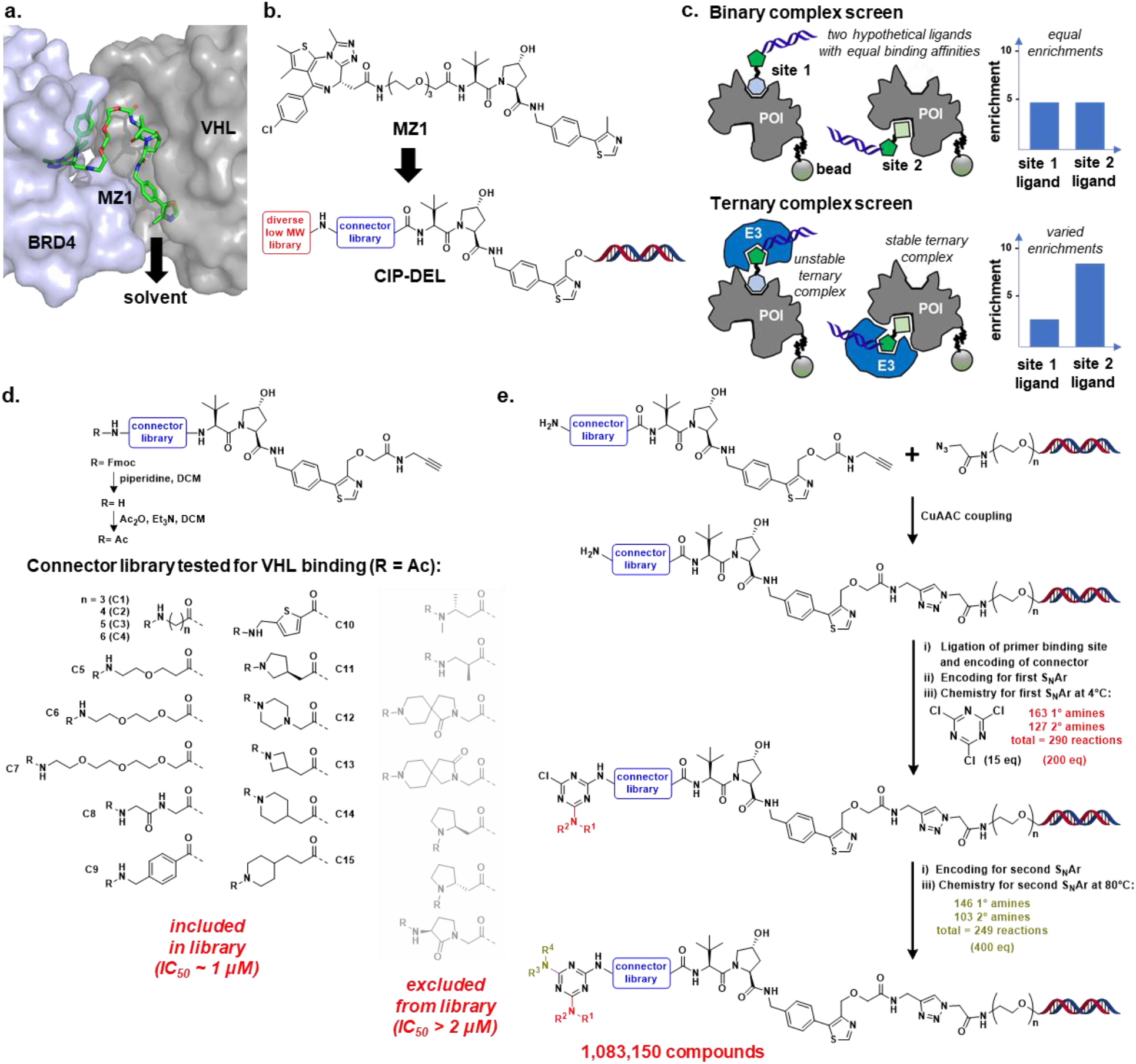
Library design and synthesis. (**a**) Using the ternary complex crystal structure of **MZ1** with BRD4^BD2^ and VHL (PDB: 5T35), the thiazole-methyl group of **MZ1** was selected as the site for DNA attachment as it projects towards a solvent exposed channel, potentially minimizing disruption of the ternary complex. (**b**) **MZ1** was used as a template for CIP-DEL library design. The VHL ligand was functionalized with a connector library and appended to DNA. Each connector terminated with an amine that was functionalized using split-and-pool DEL synthesis methods. (**c**) Affinity screening strategy for identifying complex-inducing library members. Parallel screens with the protein of interest +/-the E3 ubiquitin ligase are performed and enrichment signals that remain high in both screens suggest library members that are capable of stable ternary complex induction. (**d**) The VHL ligand was modified with an alkyne to enable DNA attachment and functionalized with 22 Fmoc-amino acid connectors. The Fmoc groups were replaced by acetyl groups, and each compound was assayed for binding to VHL. We selected 15 connectors that retained binding to VHL to include in the CIP-DEL library. (**e**) Overview of the CIP-DEL library synthesis. Full synthetic details are provided in the supplementary information.

Each connector terminated with a primary or secondary amine that was used to construct a split- and-pool library of bifunctional compounds. Numerous DEL library designs could be used for functionalizing the connector amines; here, we selected the well-established triazine-library concept for our initial experiments given the large diversity that can be generated with sequential SNAr reactions on the triazine core.^24^ Each connector amine was directly functionalized with the triazine scaffold, and two additional rounds of S Nar were performed to complete the triazine substitution (**Fig. 1e**). To select amines for inclusion in our library, we performed validation experiments with a set of 264 primary amines and 169 secondary amines. Building blocks that gave a minimum of 75% conversion to product based on UPLC-MS analysis were selected for inclusion. This resulted in the inclusion of 290 amines (163 primary and 127 secondary) for the second triazine substitution and 249 amines (146 primary and 103 secondary) for the final substitution. Therefore, with 15 connectors (cycle 1), 290 amines included in the second substitution (cycle 2), and 249 amines included in the third substitution (cycle 3), the final DEL library contains 1,083,150 potential CIPs for targeting VHL. Detailed experimental protocols for library production are provided in the **Supplementary Information** and **Extended Data Fig. 7**.

Since our library was designed using **MZ1** as a template, a DNA-tagged derivative of this degrader was synthesized (**MZ1-DEL**; **Supplementary Fig. S18 – S20**). This control compound was synthesized separately from the library, encoded with a unique barcode, and spiked into the library at a theoretical equimolar ratio to each library member.

### Affinity Screens and Sequencing Analysis

For CIP-DEL screening, we targeted the first bromodomain (BD1) of BRD4 and screened the library using three parallel conditions: (i) a “beads-only” screen to control for matrix binders; (ii) a “BRD4 only” screen (denoted BRD4 (-) VHL) to identify binders to BRD4^BD1^; and (iii) a “BRD4 with VHL” screen (denoted BRD4 (+) VHL) that included excess VCB complex (∼85 equivalents of VCB complex compared to CIP-DEL library; ∼43 equivalents of VCB compared to BRD4^BD1^) to identify library members that bind BRD4 in the presence of VCB. Our expectation was that BRD4 ligands identified in the BRD4 (-) VHL screen would show reduced enrichments in the screen with VCB if the library compound was unable to induce a stable ternary complex (*e*.*g*., due to steric or electrostatic clashes between presenter and target proteins: **Fig. 1c**). In contrast, BRD4 ligand–connector combinations that induced a stable ternary complex would yield high enrichment levels in both (+/-) VHL screens. In the ideal situation, induction of neo-PPIs would lead to increases in experimental enrichment values in the presence of VCB, suggesting cooperative ternary complex formation.

Following the affinity-based screens, library members retained in each experimental condition were separated from the beads by heating and the output DNA was PCR-amplified and subjected to next-generation sequencing (NGS). Analysis of our sequencing results showed that we obtained roughly equal sequencing depth for each of the three screening conditions (**Extended Data Fig. 1**). However, in the BRD4 (+) VHL screen, we observed a marked increase in the number of high-count barcodes. To investigate the count distribution further, we looked at barcode counts for each individual building block across each screening condition (**Extended Data Fig. 2**). For each connector included in the library (denoted BB1), we saw roughly even distribution of barcode counts in the beads-only and BRD4 (-) VHL screens. Remarkably, in the screen with VHL present, we found a dramatic shift in NGS reads towards library members that possessed connector **C15** suggesting that this connector was a key component for successful ternary complex induction. Intriguingly, the second most prevalent connector was **C14**, which is a one methylene truncated analog of **C15** suggestive of a structure–activity relationship (SAR).

We next looked for preferred cycle-2 and -3 building blocks (denoted BB2 and BB3, respectively). In the BRD4 (+) VHL screen, the highest counts in cycle 2 came from **BB2–109** (**Extended Data Fig. 2**). Interestingly, the cycle-3 building block with the highest counts in the BRD4 (+) VHL screen was **BB3-114**, which has the same structure as **BB2-109** (**Extended Data Fig. 2**). Since a common set of amine-based building blocks was profiled for cycle-2 and -3 triazine substitutions, and amines that provided at least 75% UPLC-MS product yield were included in the library, some amines were included in both cycles 2 and 3. That **BB2-109** / **BB3-114** was the most enriched building block within each substitution cycle strongly suggests that this building block is a favored chemotype.

To refine our screening analysis further, we calculated enrichment values for each library member by comparing the barcode counts from the BRD4 +/-VHL screens to the counts from the beads-only screen. We modeled the NGS counts as a Poisson sampling of the barcodes and estimated an enrichment ratio, and the upper- and lower-bounds of the 95% confidence interval (see the **Supplementary Information** for details). For all subsequent analyses, we used the lower bound (lb) of the 95% confidence interval as our enrichment metric since this provided a more conservative estimate of enrichment. Cube plots were generated from the enrichment values to visualize SAR trends (**Fig. 2a – c**). We looked for library members with three different output profiles: (i) high-confidence enrichment in the BRD4 (-) VHL screen (**Fig. 2a**); (ii) high-confidence enrichment in the BRD4 (+) VHL screen (**Fig. 2b**); and (iii) molecules that showed enrichment in both screens (**Fig. 2c**). Points in **Fig. 2a** are sized by BRD4 (-) VHL enrichment (range: 0 to 120) and points in **Fig. 2b and c** are sized by BRD4 (+) VHL enrichment (range: 0 to 389). The **MZ1-DEL** control compound is shown at the origin of each plot (1, 1, 1; red).

**Figure 2.**
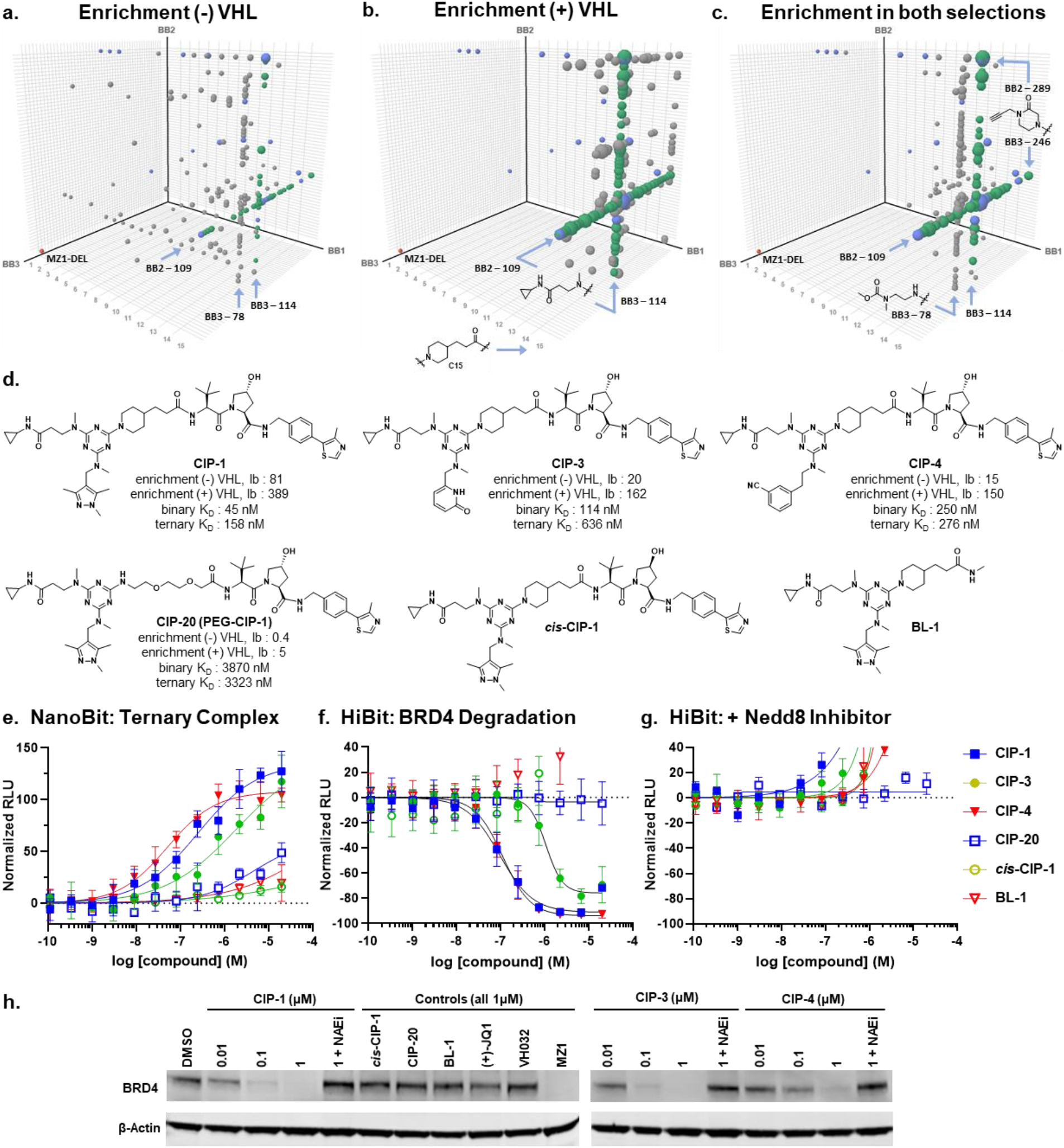
CIP-DEL screening analysis and evaluation of compounds synthesized off-DNA. (**a -c**) Cube plots of enriched features in (**a**) the BRD4 (-) VHL screen (enrichment cutoff ≥ 7), (**b**) the BRD4 (+) VHL screen (enrichment cutoff ≥ 80), and (**c**) both the BRD4 (-) VHL and BRD4 (+) VHL screens (enrichment cutoff ≥ 5 in both). Points are sized by (-) VHL enrichment values in (**a**) and (+) VHL enrichment values in (**b**) and (**c**). Coloring: red = **MZ1-DEL**; grey = meets enrichment cutoff; green = contains **C15** and **BB2-109** or **BB3-114**; blue = synthesized off-DNA. Compounds selected for synthesis are shown in all plots and may not meet indicated enrichment cutoffs. (**d**) Structures of select library members synthesized off-DNA with binary and ternary KD values from SPR (see **Table 1** for additional details). (**e**) Ternary complex induction by select CIP compounds using the NanoBiT assay. (**f**) Compound induced BRD4^BD1^ degradation measured by the HiBiT assay. (**g**) BRD4^BD1^ degradation with CIP compounds is reversed in the presence of a Nedd8 inhibitor. Data in **e** -**g** represent the mean ± s.d. of n = 2 independent experiments. (**h**) Western blot analysis of endogenous BRD4 degradation with CIP compounds and controls. Uncropped gels are provided in the supplementary information.

In the BRD4 (-) VHL screen, we found weaker overall enrichments compared to the BRD4 (+) VHL screen and greater dispersion of the compounds meeting the enrichment cutoff ≥ 7 (cutoff values were selected to give ∼200 plotted data points). However, several line features were present and many of the enriched features contained **C15** and **BB2-109** / **BB3-114** (shown in green). When VCB complex was included in the screen, the preference for compounds containing the **C15** connector paired with **BB2-109** or **BB3-114** became strongly apparent (**Fig. 2b**; green). Strong enrichment values were seen across both **BB2-109** and **BB3-114** line features, suggesting that a variety of amines may be tolerated at the final diversity element. Given the strong enrichments for **C15** and **BB2-109** / **BB3-114**, we focused our off-DNA follow-up efforts on compounds containing these features. To confirm that our screening results were predictive of off-DNA binding affinities, we synthesized compounds containing **BB2-109** / **BB3-114** but with connectors that gave weaker enrichments in our screens. Library members selected for off-DNA synthesis are colored blue in **Fig. 2a – c**.

**Table 1.**
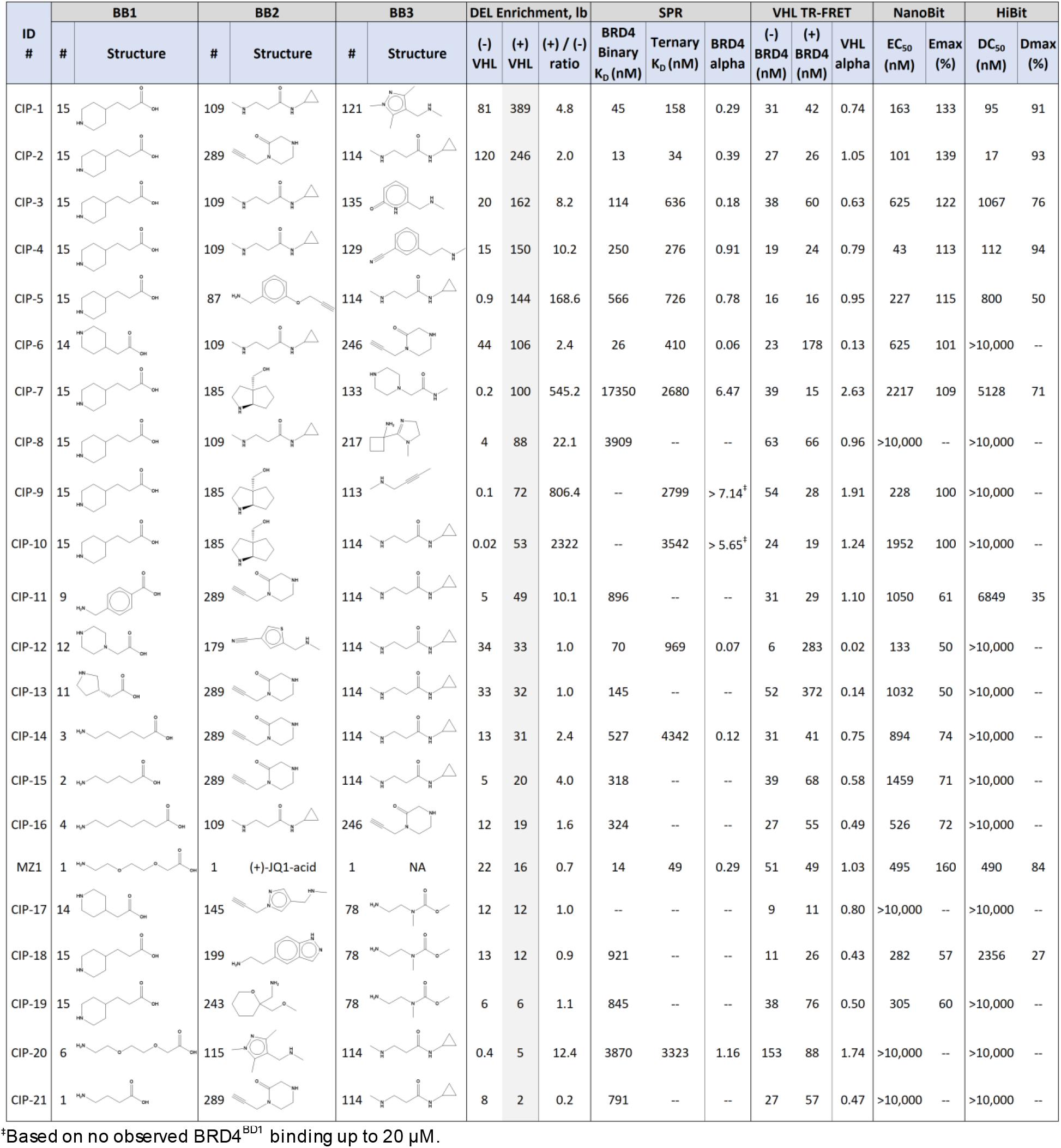
Profiling of compounds selected for off-DNA synthesis. On-DNA enrichment values corresponding to each DNA-free compound listed in the table are provided for the screens (-/+) VHL, along with a calculated ratio between the two screens. Binary KD values for BRD4^BD1^ were obtained by SPR. Ternary KD values were obtained by immobilizing BRD4^BD1^ to the SPR chip and flowing a titration of pre-incubated mixtures of compound and VCB complex over the chip. All compounds were tested up to 20 µM with BRD4 (binary kinetics) and missing values indicate that the data obtained could not be fitted to a 1:1 Langmuir or steady-state binding model. KD values are averages of all experimentally determined values (for detailed kinetic and steady-state parameters, see **Extended Data Table 1**). A TR-FRET peptide displacement assay was used to obtain EC50 values for binding to VHL in the absence or presence of BRD4^BD1^ (n = 2 independent experiments). Cellular ternary complex induction was measured using the NanoBiT assay and protein degradation of BRD4^BD1^ was assessed using the HiBiT assay (n = 2 experiments each). Standard deviations for these data can be found in **Table S1**.

As an additional query of our screening data, we plotted library members that met a nominal enrichment value in both BRD4 (+/-) VHL screens (cutoff ≥ 5; **Fig. 2c**). We observed similar trends to those seen when the BRD4 (+) VHL screen was analyzed independently (**Fig. 2b**) with some minor variations. Notably, we saw the appearance of an additional vertical line feature for **BB3–78**. However, this line feature displayed much lower enrichment values compared to **BB2-109** / **BB3-114** and as such, was deemed a second tier series.

### SPR Validation of Off-DNA Library Members

A total of 21 library members were selected for off-DNA synthesis and evaluation (denoted **CIP-1** – **21**) in addition to the **MZ1** control. BRD4 binding affinities for the compounds were first measured by surface plasmon resonance (SPR), wherein biotinylated BRD4^BD1^ was immobilized to the SPR chip. The control compound **MZ1** gave a binding affinity of 14 nM. In comparison, **CIP-1**, which corresponds to the most highly enriched barcode from the BRD4 (+) VHL screen, had a binary KD of 45 nM (**Fig. 2d**; **Table 1**). Next, we sought to determine a ternary KD by immobilizing BRD4^BD1^ on the SPR chip, adding a pre-incubated mixture of compound and VCB to the flow cell, and measuring the binding affinity of the complex. A ternary KD of 49 nM was obtained for **MZ1** and a ternary KD of 158 nM was measured for **CIP-1**. From the binary and ternary KD values, the cooperativity factor α (defined as α = KDbinary / KDternary)^25^ was calculated to be 0.29 for **CIP-1**, indicating negative cooperativity despite remaining a potent CIP.

In the BRD4 (-) VHL screen, the most highly enriched barcode corresponded to **CIP-2**, which was also the fourth most highly enriched barcode in the BRD4 (+) VHL screen (**Table 1**). **CIP-2** contains the highly enriched **C15** and **BB3-114**. The final triazine substituent was **BB2-289**, a structure present in both cycles 2 and 3 that showed strong enrichments in both positions (**Fig. 2c**). **CIP-2** was evaluated by SPR and displayed a binary KD of 13 nM and a ternary KD of 34 nM, matching the potencies of **MZ1** with no off-DNA optimization.

Two additional representatives that contain the **C15** / **BB2-109** pairing and gave strong enrichment values in both screens are **CIP-3** and **CIP-4** (**Fig. 2d**). SPR analysis for **CIP-3** and **CIP-4** gave binary KD values of 114 nM and 250 nM, and ternary KD values of 636 nM and 276 nM, respectively. Interestingly, the weaker BRD4^BD1^ ligand resulted in a stronger ternary KD, indicating enhanced cooperativity.

For additional CIP-DEL validation, we synthesized **CIP-20** which contains the same cycle 2 and 3 building blocks as **CIP-1**, but with a connector that did not show strong enrichment in our screening (**Fig. 2d**; **Table 1**). We chose connector **C6** as this is the structure used in **MZ1**. Despite containing the same cycle 2 and 3 building blocks as **CIP-1, CIP-20** displayed substantially weaker binary and ternary KD values, as suggested by our screening data. This indicates that the **C15** connector is indeed an important contributor to both binary and ternary binding affinity for this series. The full collection of 21 library members selected for off-DNA synthesis and profiling can be found in **Table 1** and detailed SPR results are provided in **Extended Data Table 1**.

### Confirmation of VHL Binding by TR-FRET

Each VHL ligand–connector pairing included in the CIP-DEL library had shown comparable binding to VCB when acetylated at the terminal amine of the connector (**Fig. 1d**). To verify that the addition of the triazine component did not disrupt binding, we tested our off-DNA validation compounds in the TR-FRET assay for VCB binding. All compounds were found to retain binding to VCB with an average EC50 of 35 nM (range: 6 nM to 153 nM; **Table 1**). The TR-FRET assay was also performed with the addition of BRD4^BD1^ to look for cooperative complex induction. Most compounds showed slight negative cooperativity in the presence of BRD4^BD1^. Notably, **CIP-12**, which contains a basic amine in the connector, displayed strong negative cooperativity (α = 0.02). Interestingly, **CIP-7, -9**, and **-10** displayed slight positive cooperativity for VCB binding in the TR-FRET assay (α = 2.63, 1.91, and 1.24, respectively) and positive cooperativity for BRD4^BD1^ binding in our SPR analysis (α = 6.47, > 7.14, and > 5.65, respectively). For **CIP-9** and **-10**, we could not determine binary KD values by SPR when titrating the compounds up to 20 µM; however, measurable ternary KD values were obtained indicating a minimum cooperativity of 7.14 and 5.65 for **CIP-9** and **-10**, respectively. Common to **CIP-7, -9**, and **-10** is **BB2-185**, which contains a constrained hydroxymethyl group. We speculate that this functionality may be a key contributor to the positive cooperativity.

### Ternary Complex Induction and Protein Degradation

To complement our SPR and TR-FRET datasets, we assessed ternary complex induction in cells using the high-sensitivity NanoBiT split-luciferase assay, wherein the luciferase was split between BRD4^BD1^ and VHL.^26^ **CIP-1, -3**, and **-4** induced robust complex assembly with EC50 values of 163 nM, 625 nM, and 43 nM, respectively (**Fig. 2e)**. Importantly, **CIP-20** induced minimal complex assembly, further validating our screening results. To confirm that the NanoBiT signal was dependent on VHL binding, we tested the *cis*-hydroxyproline derivative of **CIP-1** (***cis*-CIP-1**). As expected, ***cis*-CIP-1** produced negligible ternary complex induction. A truncated analog of **CIP-1** (denoted **BL-1**), which comprises only the BRD4^BD1^ ligand with the **C15** connector, was synthesized and tested in the NanoBiT assay to confirm further that complex assembly required the VHL-targeting motif. Negligible activity was observed with **BL-1**, confirming that the complete bifunctional compound is necessary for complex induction. The complete set of NanoBiT EC50 and Emax values can be found in **Table 1**.

Although our CIP-DEL screens were not coupled to protein degradation, the discovery of CIPs targeting BRD4 to VHL could result in functional degraders. To establish this, we profiled our DNA-free compounds in a HiBiT assay using cells transfected with a HiBiT-tagged BRD4^BD1^ plasmid.^27^ We found that **CIP-1** and **CIP-4** produced potent BRD4^BD1^ degradation with DC50 values of 95 nM and 112 nM and Dmax values of 91% and 94% (**Fig. 2f**). Interestingly, **CIP-3** was found to be ∼10x less potent than **CIP-4** even though **CIP-3** was a more potent BRD4^BD1^ ligand, reinforcing the idea that binary affinity to the POI is less consequential for degradation than cooperativity and ternary KD. **CIP-2** provided a DC50 of 17 nM and a Dmax of 93%, representing the most potent degrader we identified. Considering that **CIP-2** came directly from our screen with no ligand or connector optimization, this provides compelling validation of our CIP-DEL approach.

Consistent with our NanoBiT data, ***cis*-CIP-1** and **BL-1** induced no protein degradation in the HiBiT assay, confirming the importance of VHL binding for the biological effect (**Fig. 2f**). Treating cells with **CIP-20** produced no BRD4^BD1^ degradation, suggesting that enrichment in the CIP-DEL screen was predictive of functional degraders. To confirm that the degradation observed with **CIP-1, -3**, and **-4** was cullin-RING ligase dependent, the HiBiT assay was repeated in the presence of the NEDDylation inhibitor (NAEi) MLN 4924,^28^ which fully reversed degradation (**Fig. 2g**). In fact, addition of the NAEi generated a dose-dependent increase in the HiBiT curves. A similar result was observed for ***cis*-CIP-1** and **BL-1** in the absence of NAEi. These data suggest that BRD4^BD1^ binding promotes ligand-induced stabilization of the protein, a phenomenon recently exploited in the development of a HiBiT-based CETSA assay.^29^

Since an isolated protein domain was used in the HiBiT assay, we performed Western blot analysis of HEK293 cells treated with CIPs to assess endogenous BRD4 degradation (**Fig. 2h**). A dose-dependent degradation was observed for **CIP-1, -3**, and **-4**, consistent with the HiBiT results; however, the relative potencies for **CIP-3** and **-4** appear to be different when examining endogenous BRD4 degradation. Co-treatment of cells with active CIPs and a NAEi completely reversed the observed degradation. As expected, no degradation was observed for **CIP-20, *cis*-CIP-1**, or **BL-1** at 1 µM, whereas **MZ1** induced complete BRD4 removal at this concentration. The BRD4-binding element **(+)-JQ1**^30^ and the VHL-binding element **VH032**^31^ showed no degradation as monovalent ligands. Finally, cell viability was measured to verify that the observed protein degradation was not due to compound toxicity. After 16 h of compound treatment, no toxicity was observed for the tested CIPs up to 20 µM (**Supplementary Fig. S18**).

### CIP-DEL Correlations with Off-DNA Validation Data

Correlations between on-DNA screening results and off-DNA validation data were examined to assess the predictive capacity of the CIP-DEL platform (**Fig. 3**). The most direct comparisons can be drawn between on-DNA enrichment values and off-DNA binding affinities from our SPR data. Comparing BRD4 (-) VHL enrichments versus binary KD (**Fig. 3a**), and BRD4 (+) VHL enrichments versus ternary KD (**Fig. 3b**), we found that higher enrichment values were generally predictive of more potent binders in each of the screens. To assess the ability of CIP-DEL screening to identify compounds that promote ternary complex formation and protein degradation in cells, we compared BRD4 (+) VHL enrichment values to our NanoBiT and HiBiT results (**Fig. 3c** and **3d**). BRD4 (+) VHL enrichments correlated well with NanoBiT Emax and HiBiT Dmax values, which represent important degrader attributes (**Fig. 3d**). HiBiT DC50 values showed reasonable correlation with BRD4 (+) VHL enrichments, but poor association was seen between NanoBiT EC50 data and enrichment values (**Fig. 3c**). Similar trends were observed when the NanoBiT and HiBiT data was compared to ternary KD values from SPR (**Fig. 3e** and **3f**), indicating that barcode enrichment and ternary complex binding by SPR were similarly predictive of performance in the NanoBiT and HiBiT assays. HiBiT data and NanoBiT data showed good inter-assay correlations (**Fig. 3g** and **h**).

**Figure 3.**
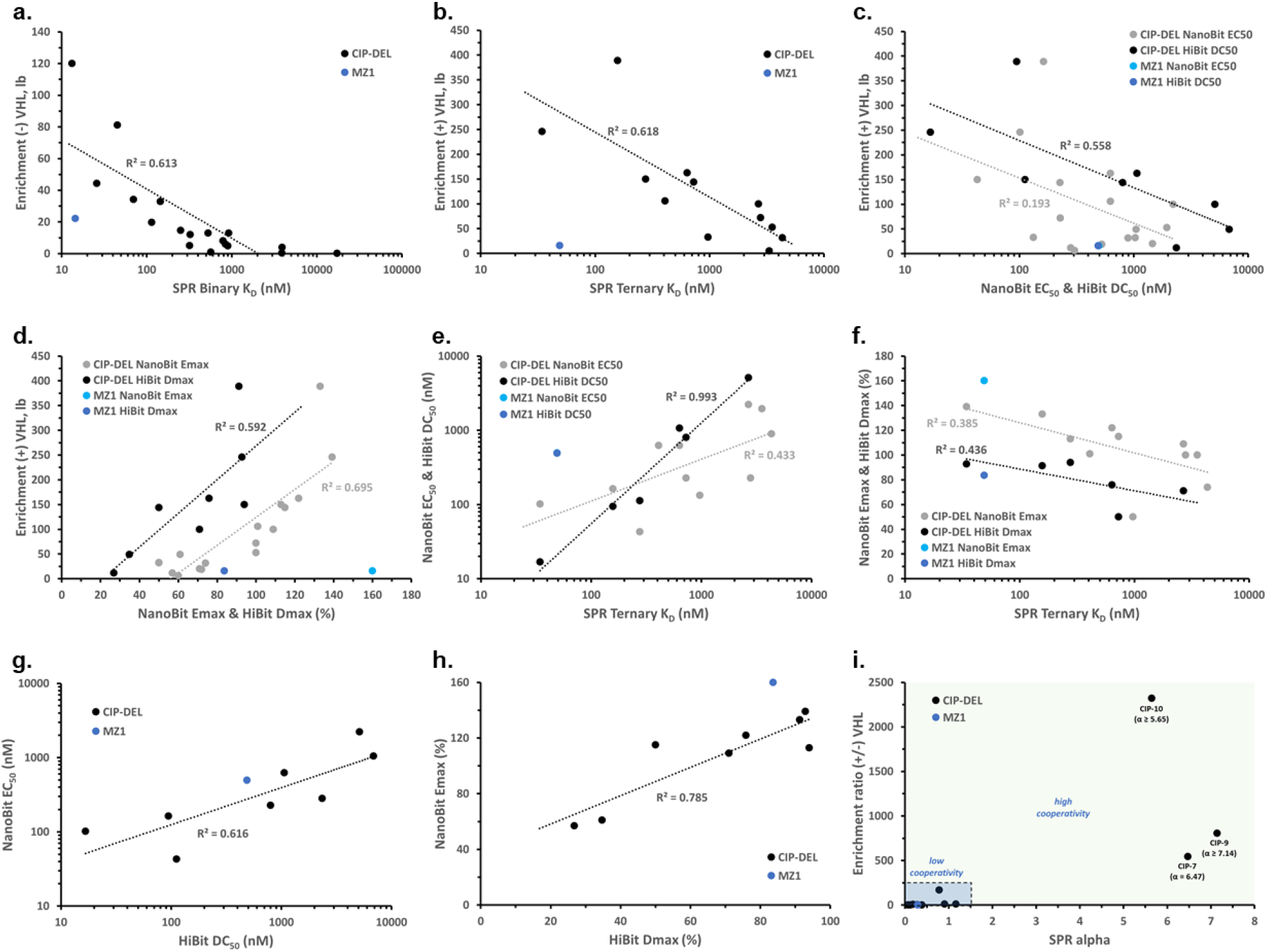
Correlations between CIP-DEL screening results and off-DNA validation data. (**a**) Correlation between BRD4 (-) VHL enrichment values (lower-bound of 95% confidence interval) and binary KD values obtain by SPR. (**b**) Relationship between BRD4 (+) VHL enrichments and ternary KD values obtained by SPR. (**c**) BRD4 (+) VHL enrichment values compared to NanoBiTEC50 and HiBiT DC50 values. (**d**) BRD4 (+) VHL enrichments compared to NanoBiT Emax and HiBiT Dmax results. (**e**) Relationship between NanoBiT EC50 and HiBiT DC50 values, and the ternary KD values obtained from SPR. (**f**) Correlation of NanoBiT Emax and HiBiTDmax with the ternary KD results obtained by SPR. (**g**) Association between the NanoBiT EC50 and HiBiT DC50 results obtained for each active off-DNA compound. (**h**) Correlation between NanoBiT Emax and HiBiT Dmax values for active DNA-free compounds. (**i**) Comparison of CIP-DEL enrichment ratios (+/-) VHL and the alpha values calculated from binary and ternary SPR KD values. In all panels, data points for **MZ1** were excluded from the line fitting due to potential ternary complex disruptions caused by attachment of the DNA tag (see text).

Lastly, we hoped to determine if our screening system was predictive of CIP cooperativity. We calculated a ratio of the enrichment values from the screens +/-VHL and compared these ratios to the α values obtained by SPR (**Fig. 3i**). While the largest enrichment ratios from our screen did correspond to positively cooperative compounds, we identified few cooperative compounds and additional data will be needed to validate these trends.

### Ternary Complex Crystal Structure

To understand better the striking preferences observed for **C15** and **BB2-109** / **BB3-114**, we solved the X-ray crystal structure of the BRD4^BD1^–**CIP-1**–VHL–Elongin B–Elongin C complex (PDB: XXXX, **Fig. 4; Extended Data Fig. 3**). In contrast to the “folds on itself” conformation of PEG-based degraders such as **MZ1**, the **C15** connector adopts a linear, extended conformation minimizing the entropic penalty incurred during complex induction. Assembly of the complex generates a short channel stretching between the two proteins that is occupied by the **C15** connector and surrounded primarily by hydrophobic residues from both VHL and BRD4^BD1^ (**Extended Data Fig. 3c -f**). The connector structure generates an effective balance of close protein associations without inducing steric clashes and results in 216 Å^2^ of solvent accessible surface area that is buried by the interface (**Extended Data Fig. 4**). The complex is further stabilized by ligand-induced PPIs. Most notably, Asp145 of BRD4^BD1^ forms two hydrogen bonds with VHL, one to Tyr112 and the second to His110 (**Fig. 4b**). All these factors combine to explain the strong preference observed for the **C15** connector.

**Figure 4.**
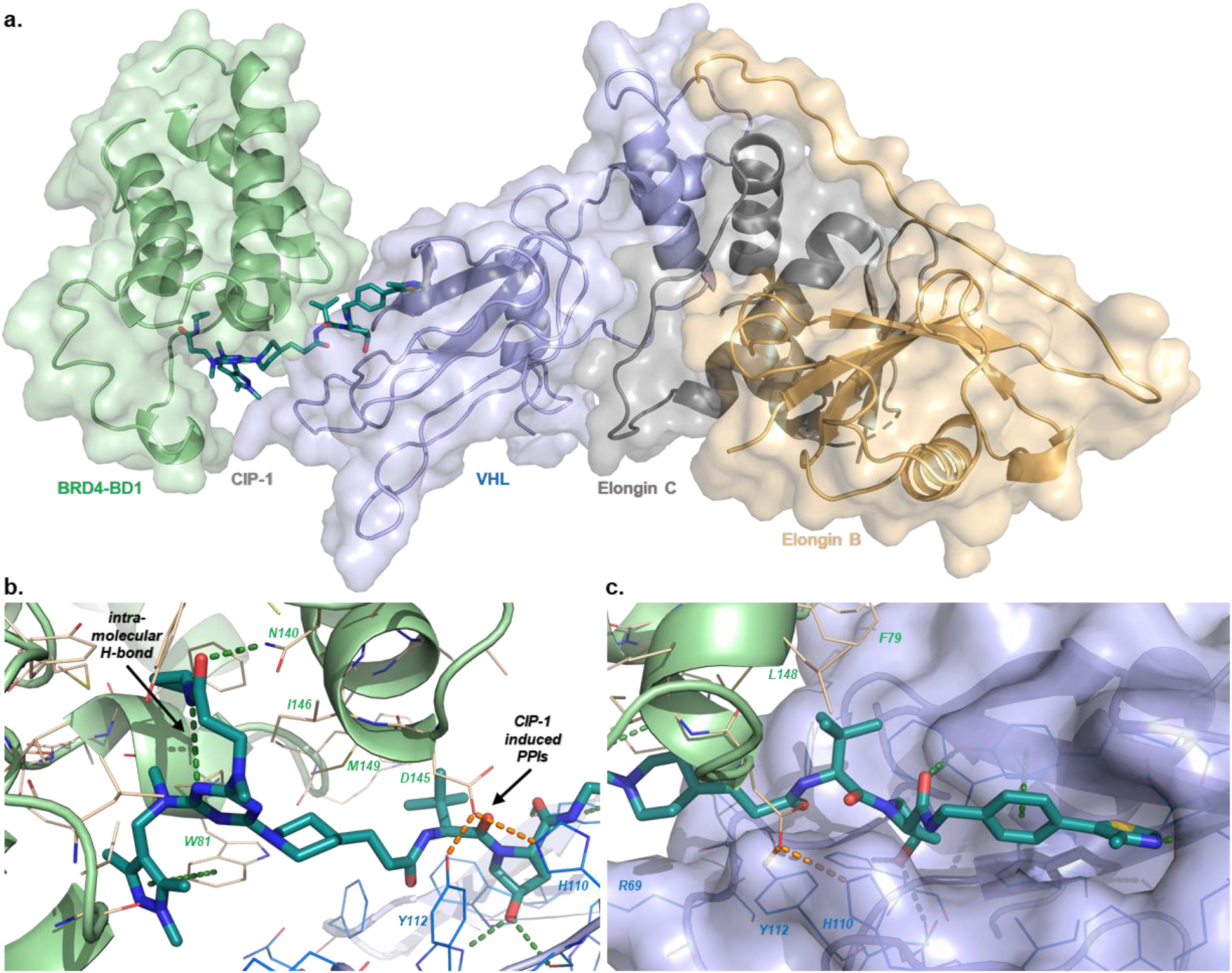
Ternary complex crystal structure of CIP-1 with BRD4 and VHL-Elongin C-Elongin B. (**a**) Overall architecture of complex assembly. Protein components are shown as ribbons with semi-transparent surfaces and labeled by protein color. **CIP-1** is shown as sticks. (**b**) Key interactions of **CIP-1** with BRD4^BD1^. An amide carbonyl from **CIP-1** forms an H-bond with Asn140 of BRD4^BD1^ and the pyrazole substituent of **CIP-1** forms a pi-stacking interaction with Trp81. An intramolecular H-bond between an amide NH and a triazine nitrogen of **CIP-1** reinforces the bound conformation of the BRD4 binding component. The ternary complex is stabilized by two H-bonds formed between Asp145 of BRD4^BD1^ and Tyr112 and His110 of VHL. (**c**) Key interactions of **CIP-1** with VHL. The VHL binding component of **CIP-1** maintains equivalent interactions with the E3 ligase as the parent **VH032** ligand.

The VHL-targeting component of **CIP-1** maintains equivalent interactions with the E3 ligase as the parent ligand **VH032** (**Fig. 4c**).^31^ In addition, the *tert*-butyl group of the VHL ligand interacts with hydrophobic residues Phe79 and Met149 of BRD4^BD1^, further stabilizing the complex. The triazine component of **CIP-1** is positioned within the acetyl-lysine binding pocket of BRD4^BD1^ and forms several key interactions with the WPF shelf residues of the protein. Phe83 of the WPF shelf, along with Met107 and Ile146, form a hydrophobic pocket occupied by the cyclopropyl group of the ligand. A hydrogen bond is formed between the cyclopropylamide carbonyl and Asn140 of BRD4^BD1^ (**Fig. 4b**). Interestingly, **CIP-1** forms an intramolecular hydrogen bond between the cyclopropylamide N-H and a triazine nitrogen, which stabilizes the bound conformation of the ligand. Combined, these interactions provide a rationale for the strong enrichments observed for **BB2-109** / **BB3-114**. The pyrazole substituent of the triazine forms a π-stacking interaction with Trp81 of the WPF shelf and hydrophobic interactions with Leu92 (**Fig. 4b**).

Unexpectedly, the orientation of BRD4^BD1^ relative to VHL in the **CIP-1** complex was dramatically different than the orientation of BRD4^BD2^ and VHL in the **MZ1** structure (**Extended Data Fig. 4**).^23^ We compared the orientation of additional VHL-based degraders,^32,33^ and found substantial variability between protein orientations. These differences highlight the strong influence that the bifunctional small molecule imposes on ternary complex geometry.

## Discussion

The identification of proximity-inducing small molecules can be a tedious, multistage process entailing optimization of target- and presenter-protein binders, identification of exit vectors, and optimization of a connector. Adding to the complexity, each of the components are highly interdependent – requiring optimization of a complex multi-element ensemble. Our CIP-DEL approach streamlines this process by directly screening both target and presenter proteins using a large library of DNA-barcoded heterobifunctional compounds with varying connectors.

We show here that a library of DNA-barcoded bifunctional compounds can be readily synthesized and screened, and the sequencing outputs can be used to identify functional CIPs. As a testament to the success of this approach, we identified a 17 nM degrader of BRD4^BD1^ directly from our library. Notably, this included the discovery of a novel BRD4 ligand and the identification of a preferred connector to the E3 ligase with no off-DNA optimization. Another benefit of the CIP-DEL approach is that it is not restricted to active-site ligands since the affinity-based screening interrogates all accessible protein pockets.

For our inaugural CIP-DEL library, we modified a VHL-targeting ligand to incorporate a vector for DNA barcode attachment. The native substrate of VHL is the hydroxyproline-containing HIF-1α peptide, which spans an extended groove on the protein surface and exits at both N- and C-termini.^34^ The two accessible exit vectors from this binding site enabled the design of a presenter ligand with a DNA barcode at one end, and a connector and split-and-pool library at the other. However, other E3 ligands, such as thalidomide, do not provide a second exit vector for DNA barcode attachment. For these ligands, a modified bespoke library design might be adapted where other components, such as the connectors, provide the requisite barcode attachment sites.^35^ Nevertheless, we anticipate that the high-level library design and screening principles identified in this study should be broadly applicable to diverse CIP-DEL implementations.

As a positive control for our screening experiments, we generated a DNA-barcoded derivative of **MZ1**. The on-DNA compound **MZ1-DEL** appeared to be an outlier when compared to the off-DNA potencies for **MZ1** (*e*.*g*., **Fig. 3b**). In fact, at the outset of our analysis we were surprised to find that **MZ1-DEL** was not a highly enriched library member given the known potency of **MZ1**. The DNA attachment point used for the CIP-DEL library was designed to minimize ternary complex disruption but was added to the VHL ligand near the PPI interface induced by **MZ1** (**Fig. 1a**). Our data suggests that this attachment vector interferes with ternary complex formation for this degrader and as a result, **MZ1** data points were excluded from the correlations shown in **Fig. 3**. Interestingly, our ternary complex structure with **CIP-1** places BRD4^BD1^ further from the barcode attachment point, implying that attachment vectors to the presenter protein ligand may influence ternary complex orientations.

The field of small molecule-induced proximity induction, while launched just over 30 years ago with the first reports of molecular glues and bifunctional compounds,^1^ is now in its growth stage and is expanding in many new directions. Preselecting targets and presenter proteins with robust discovery of proximity-inducing small molecules will enable many neo-protein and neo-substrate-related activities. The approach described here, although illustrated thus far through a single proof-of-concept protein degrader screen, is appealing in its modularity and potential generality. Many opportunities now exist to extend this concept to DELs whose members have simple, compact structures, and to novel screening conditions and data analyses to discover molecular glues having highly cooperative binding bidirectionally.

## Supporting information

Spectral Data

## Acknowledgements

John Capece, Jennifer Poirier, Philip Michaels, Carmelina Rakiec, Thomas Dice and Ritesh Tichkule are gratefully acknowledged for excellent technical and analytical support. Drs. Bruce Hua, Christopher Ge **r**y, Wenyu Wang, Shuang Liu, Andrew Reidenbach, Shubhroz Gill, Christian Gampe, and Nichola Smith are gratefully acknowledged for their support, guidance, and valuable feedback during the preparation of this manuscript. We also thank Dustin Dovala and Michael Romanowski for supplying the protein reagents used in SPR. The research was supported in part by the National Institute of General Medical Sciences (R35GM127045 awarded to S.L.S.) and by the NIBR Scholar’s Program

## Author contributions

J.W.M. conceived the reported project, designed and synthesized the CIP-DEL library, performed the screening, analyzed the data, and produced screening hits off-DNA. L.H. and M.V.W. contributed to the design, screening, and data analysis and produced off-DNA screening hits. A.T. conducted the NanoBiT, HiBiT, CTG, and TR-FRET experiments. G.M. generated and analyzed all SPR data. W.S. and X.M. ran crystallization screens and solved the ternary complex structure. C.W.C. and P.A.C. developed the CIP-DEL data analysis pipeline. S.B., F.B., F.J.Z., K.B., and S.L.S. provided context for the framing of the original goals, supervision, guidance, operational support, assisted in the interpretation of experimental outcomes, and made recommendations for a subset of the reported experiments. J.W.M. and S.L.S. wrote the manuscript and all authors read and edited the paper.

## Competing interests

The authors declare the following competing financial interests: S.L.S. is a shareholder and serves on the Board of Directors of Jnana Therapeutics and Kojin Therapeutics; is a shareholder and advises Kisbee Therapeutics, Belharra Therapeutics, Magnet Biomedicine, Exo Therapeutics, and Eikonizo Therapeutics; advises Vividian Therapeutics, Eisai Co., Ltd., Ono Pharma Foundation, F-Prime Capital Partners, and the Genomics Institute of the Novartis Research Foundation; and is a Novartis Faculty Scholar.

## Methods and Supplementary Information

Detailed methods and supplementary information are available online at https://doi.org/xxx

**Extended Data Figure 1.**
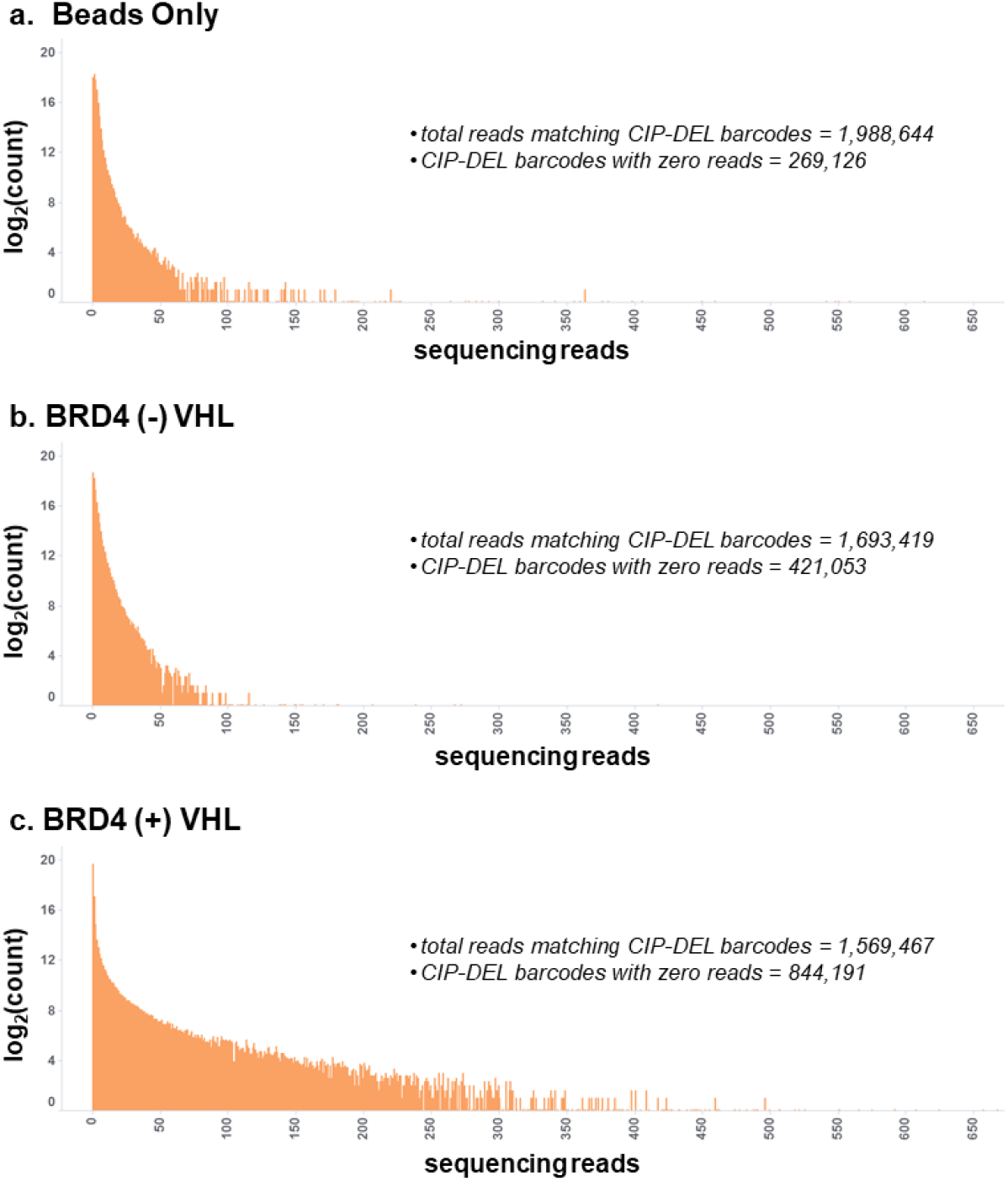
Distribution of barcode counts for each CIP-DEL library member across each screening conditions. (**a**) Barcode counts for the beads only screen. (**b**) Barcode counts for the BRD4 (-) VHL screen. (**c**) Barcode counts for the BRD4 (+) VHL screen.

**Extended Data Figure 2.**
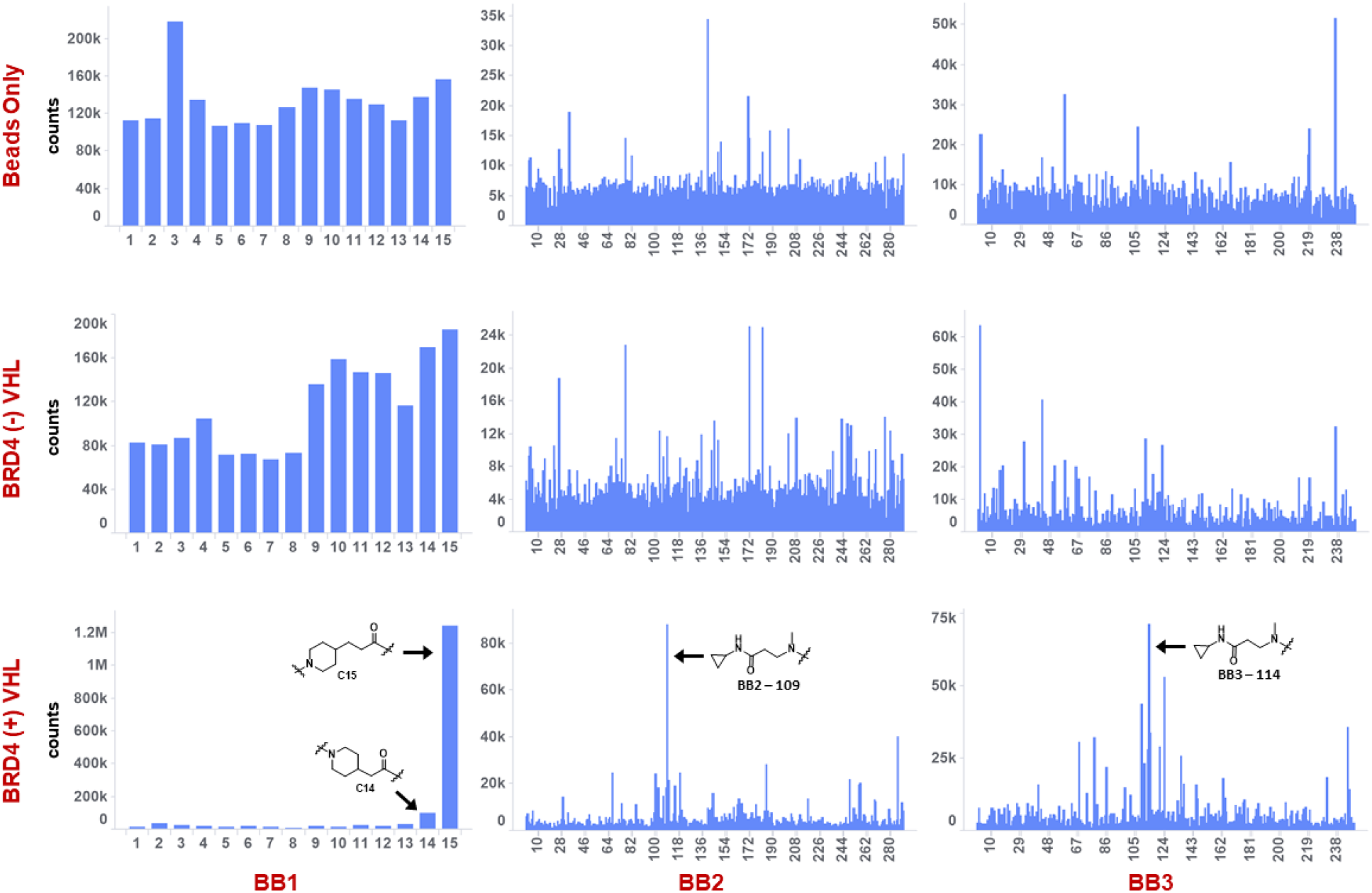
Individual building block barcode counts for each screening condition.

**Extended Data Table 1.**
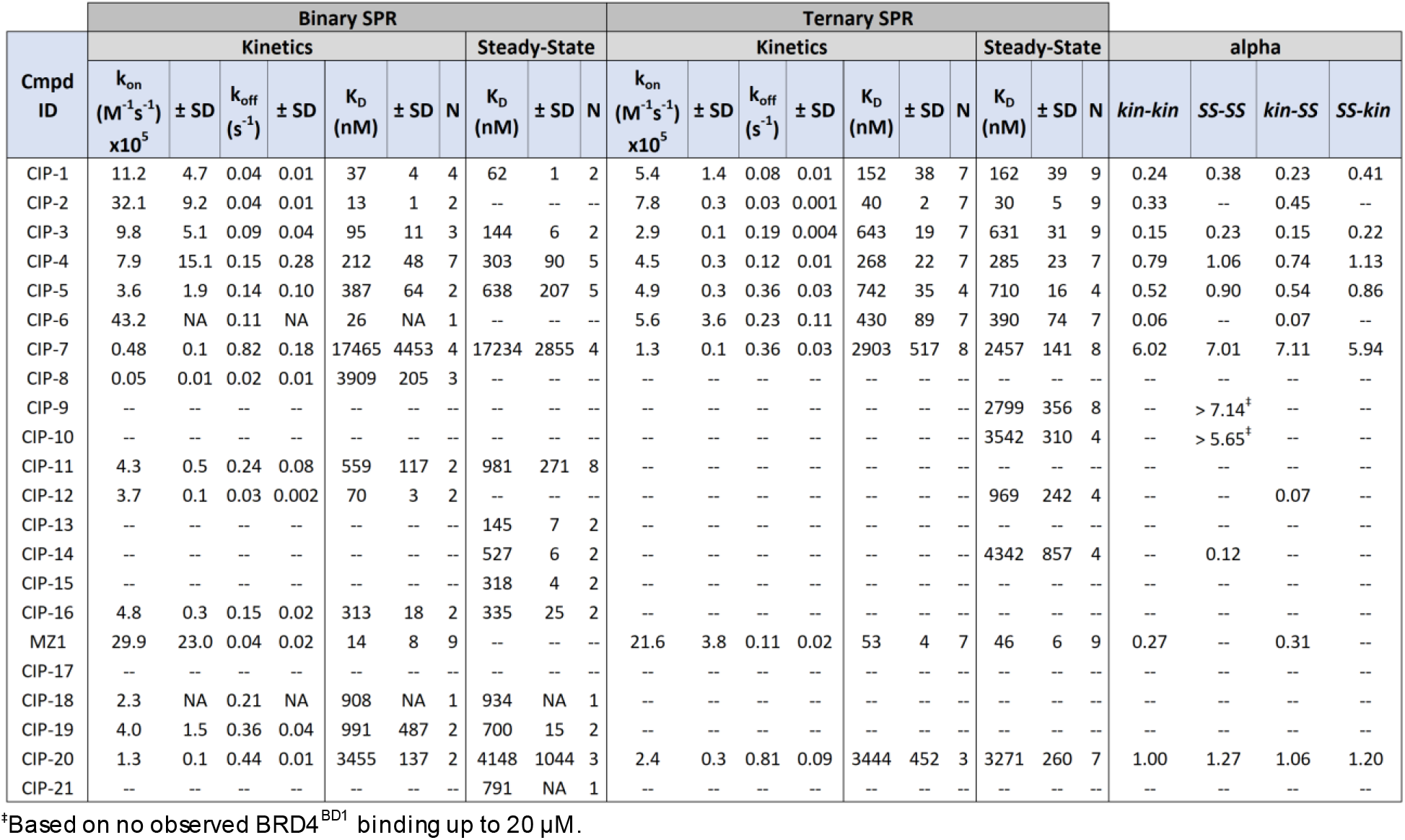
Detailed SPR data for CIP-DEL compounds synthesized off-DNA. Binary KD values were obtained with BRD4^BD1^ immobilized on the SPR chip. Ternary KD values were obtained with BRD4^BD1^ immobilized on the SPR chip and injection of a titration of pre-incubated solutions of CIP compound and VCB complex in to the flowcell. All experiments were performed in single-cycle kinetics mode. Data fitting to a 1:1 Langmuir binding model was performed where possible to provide kinetic parameters; steady-state affinity fitting was also performed where possible. Average kinetic and steady-state values obtained from binary and ternary complex experiments are provided ± s.d. for the indicated number of experiments (N). All compounds were tested up to 20 µM with BRD4 (binary kinetics) and missing values indicate that the data obtained couldnot be fitted to the indicated binding model. Cooperativity values (α) were calculated from each binary and ternary KD value obtained from kinetic (*kin*) and steady-state (*SS*) data.

**Extended Data Table 2.**
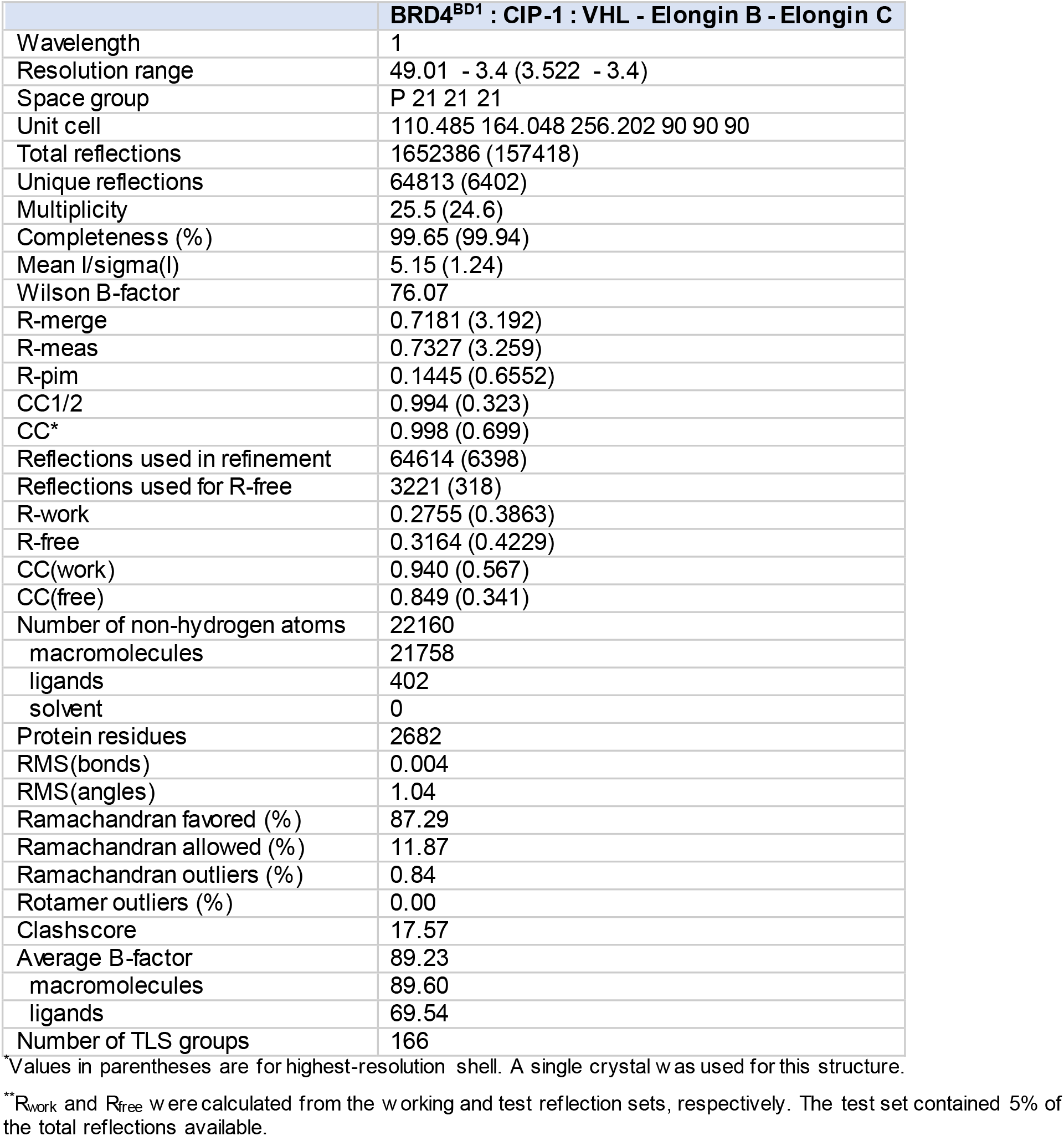
Data collection and refinement statistics.

|  | **BRD4<sup>BD1</sup> : CIP-1 : VHL - Elongin B - Elongin C** |
| --- | --- |
| Wavelength | 1 |
| Resolution range | 49.01 - 3.4 (3.522 - 3.4) |
| Space group | P 21 21 21 |
| Unit cell | 110.485 164.048 256.202 90 90 90 |
| Total reflections | 1652386 (157418) |
| Unique reflections | 64813 (6402) |
| Multiplicity | 25.5 (24.6) |
| Completeness (%) | 99.65 (99.94) |
| Mean I/sigma(I) | 5.15 (1.24) |
| Wilson B-factor | 76.07 |
| R-merge | 0.7181 (3.192) |
| R-meas | 0.7327 (3.259) |
| R-pim | 0.1445 (0.6552) |
| CC1/2 | 0.994 (0.323) |
| CC* | 0.998 (0.699) |
| Reflections used in refinement | 64614 (6398) |
| Reflections used for R-free | 3221 (318) |
| R-work | 0.2755 (0.3863) |
| R-free | 0.3164 (0.4229) |
| CC(work) | 0.940 (0.567) |
| CC(free) | 0.849 (0.341) |
| Number of non-hydrogen atoms | 22160 |
| macromolecules | 21758 |
| ligands | 402 |
| solvent | 0 |
| Protein residues | 2682 |
| RMS(bonds) | 0.004 |
| RMS(angles) | 1.04 |
| Ramachandran favored (%) | 87.29 |
| Ramachandran allowed (%) | 11.87 |
| Ramachandran outliers (%) | 0.84 |
| Rotamer outliers (%) | 0.00 |
| Clashscore | 17.57 |
| Average B-factor | 89.23 |
| macromolecules | 89.60 |
| ligands | 69.54 |
| Number of TLS groups | 166 |

\*Values in parentheses are for highest-resolution shell. A single crystal was used for this structure.
\*\*R<sub>work</sub> and R<sub>free</sub> were calculated from the working and test reflection sets, respectively. The test set contained 5% of the total reflections available.

**Extended Data Figure 3.**
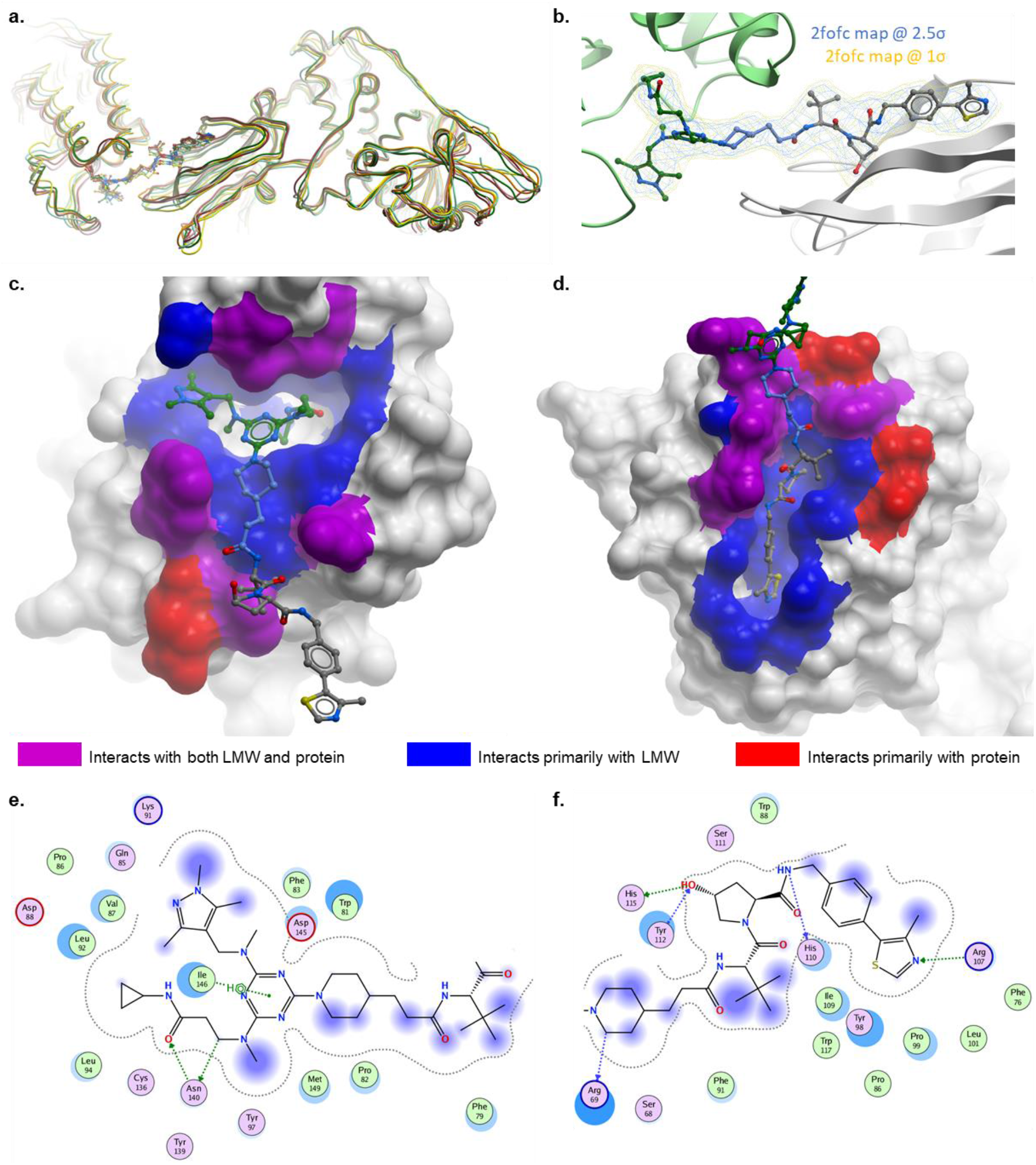
CIP-1 ternary complex structural analysis. (**a**) Six copies of the ternary complex are present in each asymmetric unit. An overlay of the six ternary complexes shows the structures are highly similar with RMS deviations of 1.06 – 1.37 Å. (**b**) The electron density for **CIP-1** is well defined in 4 of the 6 complexes of the asymmetric unit. *2Fo – Fc* maps contoured at 2.5σ and 1σ are shown for **CIP-1** in a ternary complex with well-defined density. (**c**) BRD4^BD1^ protein surface with residues colored based on interactions with **CIP-1** (blue), VHL (red), or both (purple). **CIP-1** is shown in sticks. (**d**) VHL protein surface with residues colored based on interactions with **CIP-1** (blue), BRD4^BD1^ (red), or both (purple). (**e**) 2D diagram of **CIP-1** interactions with BRD4^BD1^. (**f**) 2D diagram of **CIP-1** interactions with VHL.

**Extended Data Figure 4.**
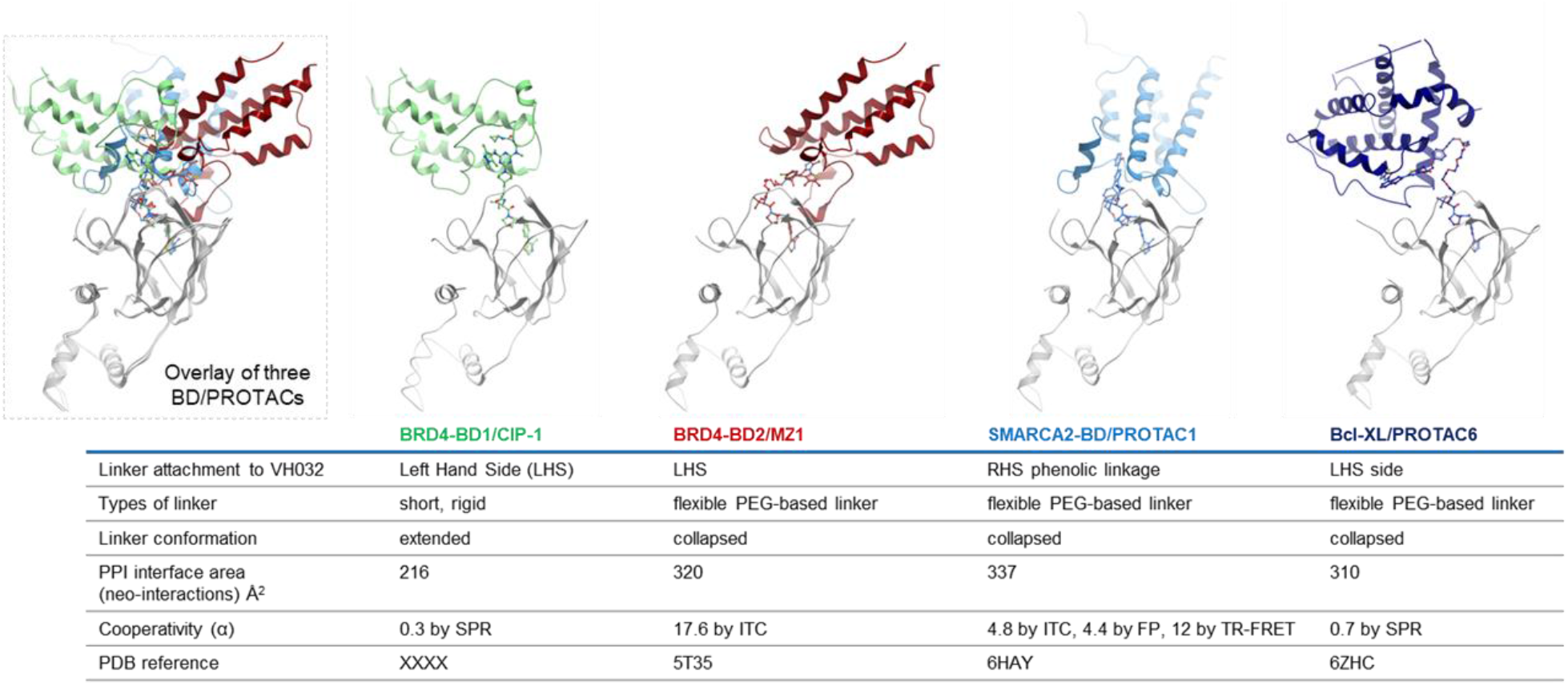
Comparison of ternary complex assemblies for four VHL-targeting bifunctional degraders. The first three structures show bromodomain targets (BRD4^BD1^, BRD4^BD2^, and SMARCA4^BD^) and the fourth complex is a non-bromodomain targeting degrader (Bcl-XL). VHL proteins are aligned at the bottom of each structure and the target protein orientation is shown at the top. Key details for each complex are provided in the table. Inset: the three bromodomain containing complexes are overlaid with VHL aligned at the bottom of the structures, highlighting the diverse target protein orientations.

**Extended Data Figure 5.**
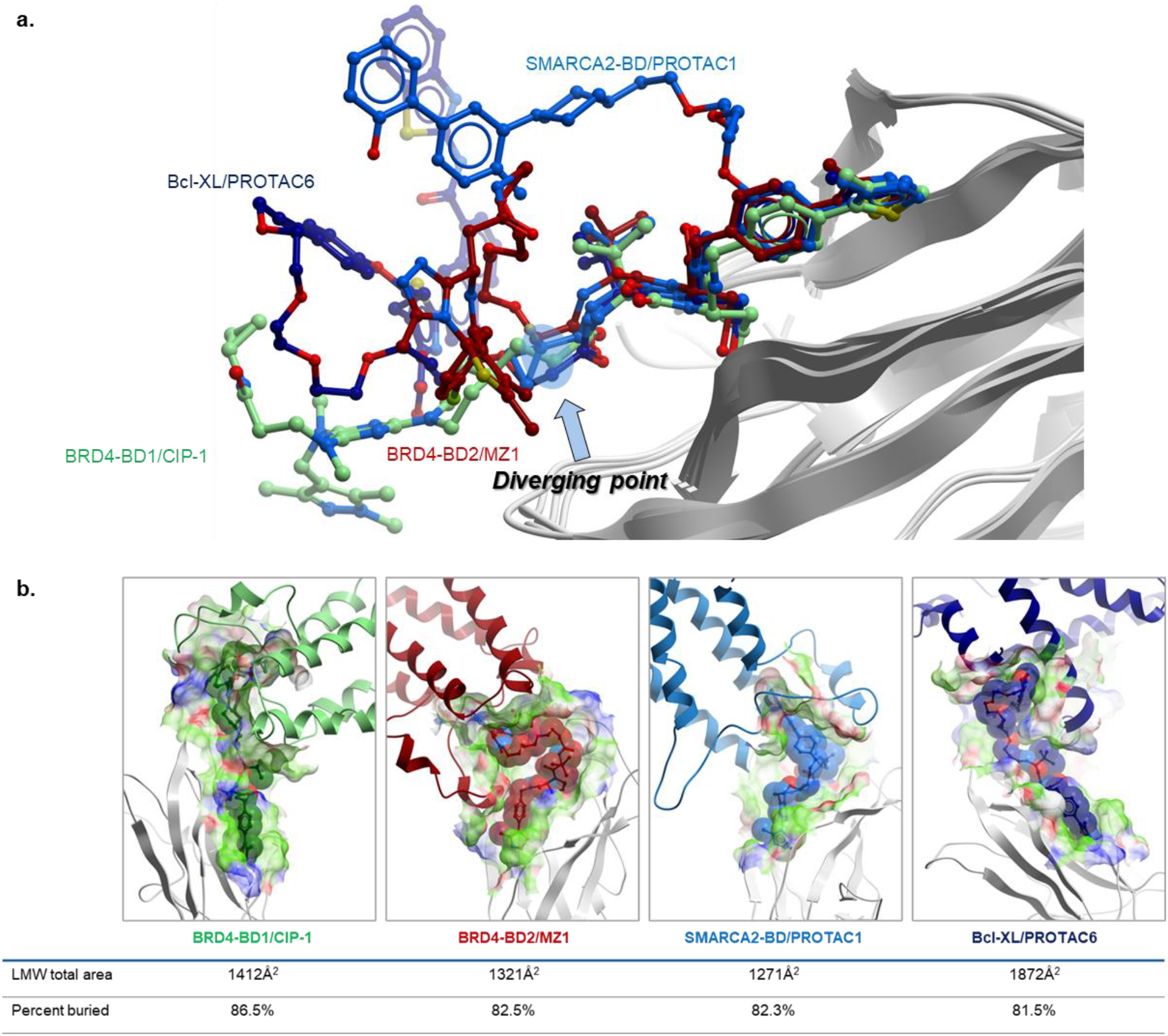
Comparison of the small molecule components of VHL PROTAC ternary complex structures. (**a**) Overlaying the VHL binding moieties of ternary complex assemblies, the diverse projection vectors of the connectors and target protein ligands are shown. The initial point of divergence for the bifunctional molecules is the carbon atom following the terminal amide of the VH032 ligand. (**b**) Comparison of the buried surface area for four VHL-targeting CIPs. The same ternary complex structures from **Extended Data Fig. 4** were used for comparison.

**Extended Data Figure 6.**
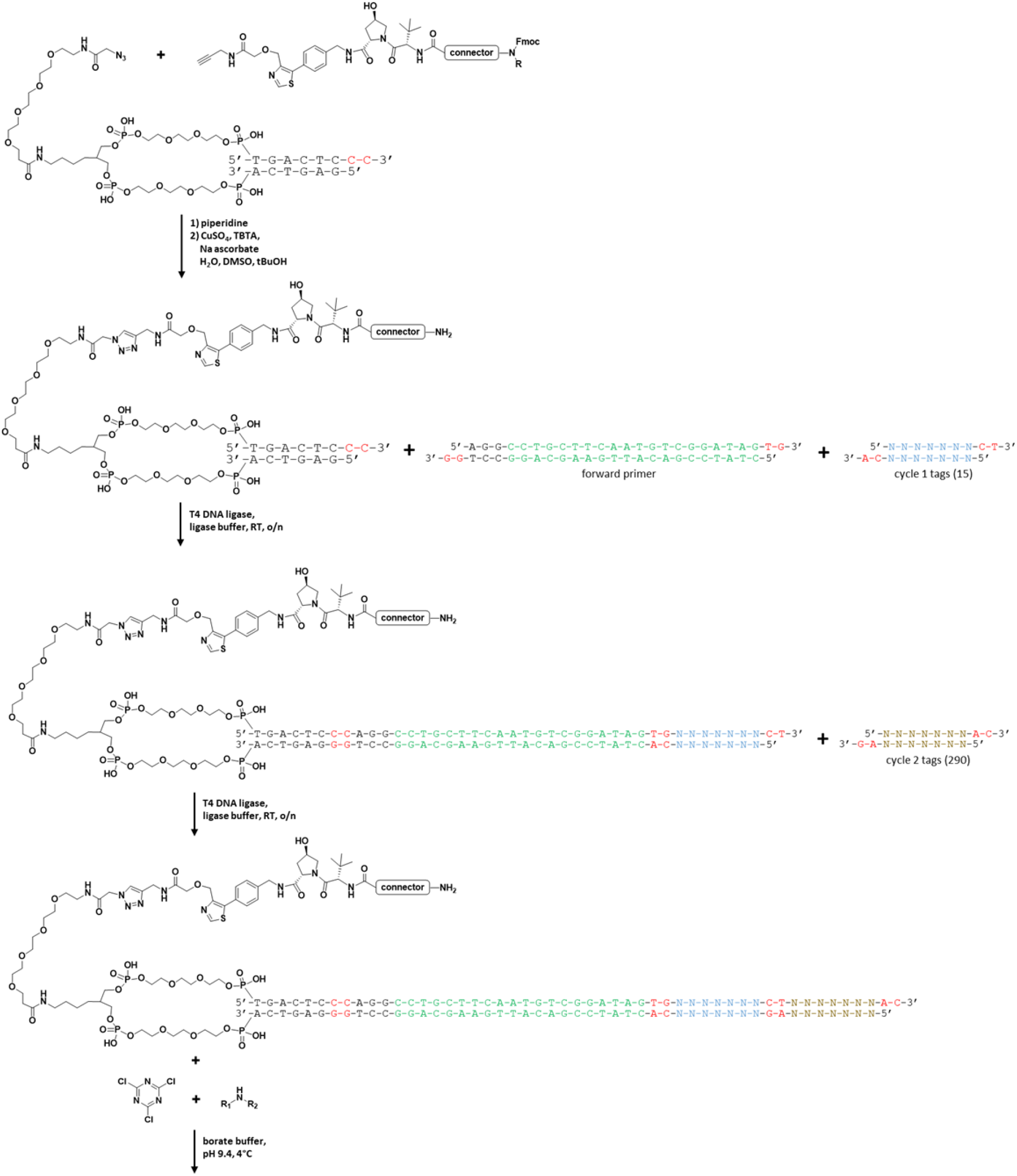
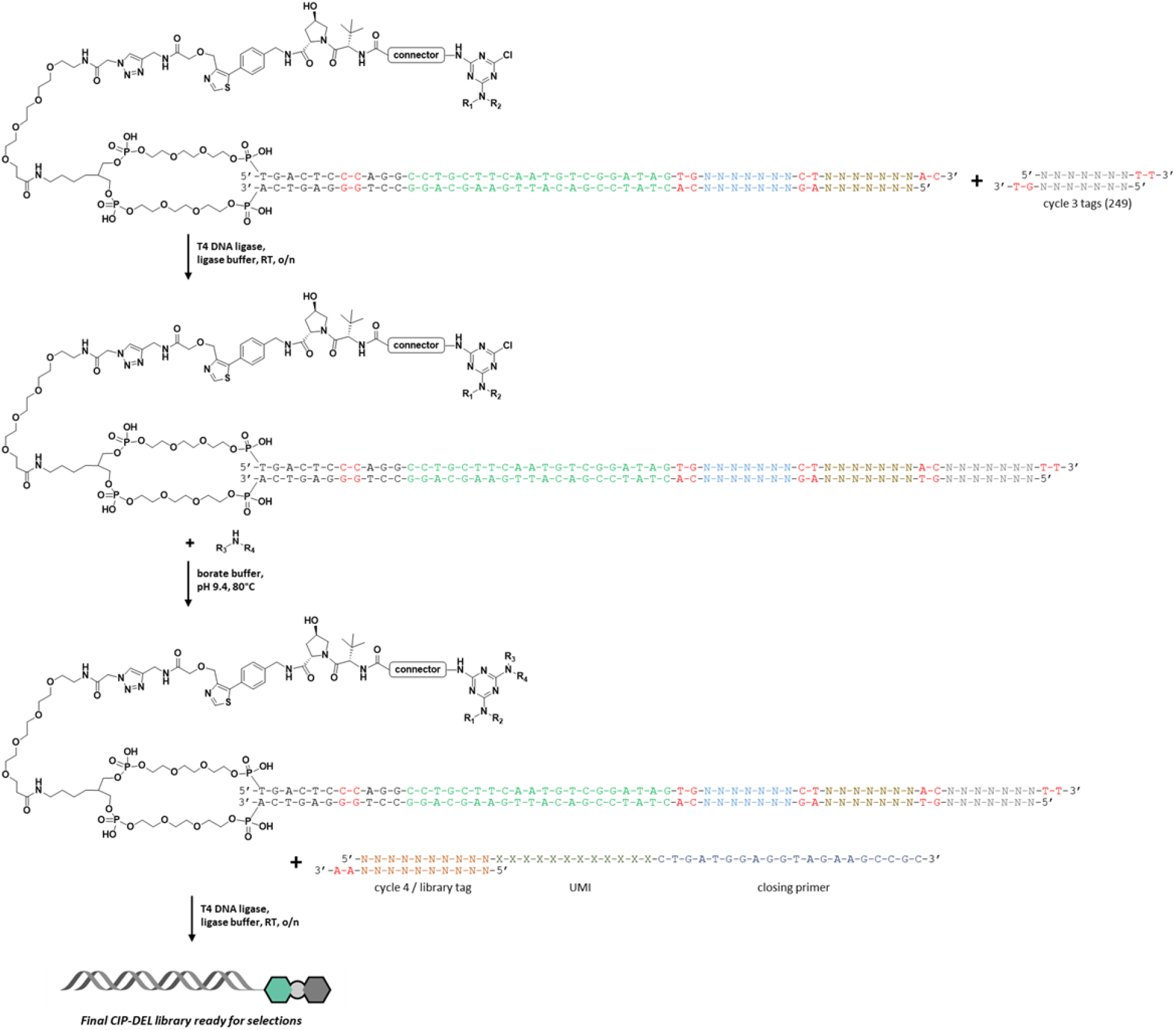
Detailed synthetic sequence for CIP-DEL library production.

