## Supplementary Information for "DNA-encoded library (DEL)-enabled discovery of proximity-inducing small molecules"

### Table of Contents


Experimental Procedures: 3

1. General chemistry information. 3
2. General CIP-DEL library information. 3
3. General experimental procedures. 4
4. Synthesis of modified VHL ligands for incorporation in CIP-DEL. 6
5. Synthesis of modified VHL ligand – connector conjugates for VHL TR-FRET binding assay. 16
6. Synthesis of CIP-DEL screening hits and controls for validation experiments. 21
7. Building block validation for the CIP-DEL library. 32
8. CIP-DEL library production. 34
9. Synthesis of MZ1-DEL positive control. 53
10. CIP-DEL selections. 57
11. NGS data processing and analysis. 60
12. VHL TR-FRET binding assay. 62
13. SPR experimental details. 62
14. NanoBit cellular ternary complex complementation assay. 63
15. HiBit cellular protein degradation assay. 65
16. Cell-Titer Glo cell viability assay. 68
17. Western blotting. 68
18. X-ray crystallography. 69
19. References 70

#### Experimental Procedures:

##### 1. General chemistry information.

Anhydrous solvents were purchased from Sigma Aldrich and used as received. *N*,*N*-Diisopropylethylamine (DIPEA), triethylamine ($Et_3N$), piperidine, sodium hydride (NaH; 60% in mineral oil), trifluoroacetic acid (TFA), and cyanuric chloride were purchased from Sigma Aldrich. 4-(4,6-dimethoxy-1,3,5-triazin-2-yl)-5-methyl-morpholinium chloride (DMT-MM), HATU, and acetic anhydride ($Ac_2O$) were purchased from Oakwood. Palladium(II) acetate was purchased from Strem Chemicals. Other chemicals were purchased from various suppliers and used as received.

Thin layer chromatography (TLC) was performed using Agela 60Å F254 aluminum-backed plates and visualized with UV light and/or staining with a solution of $KMnO_4$. Column chromatography was performed using Teledyne ISCO CombiFlash systems with RediSep Rf columns.

Ultra-performance liquid chromatography – mass spectrometry (UPLC-MS) was performed using Waters ACQUITY systems equipped with ACQUITY UPLC BEH C18 1.7 µM, 2.1 x 30 mm columns kept at 50°C. UV signals were monitored with ACQUITY UPLC PDA detectors (210 – 400 nm) and light scattering signals were recorded with ACQUITY ELSD detectors. Low-resolution mass spectra (LRMS) were obtained with a Waters SQD-2 or QDa detector with a scan time of 0.3 sec. Acidic gradients consisted of water (solvent A) and acetonitrile (solvent B) mobile phases both modified with 0.1% formic acid and basic gradients consisted of the same solvents modified with 5 mM ammonium hydroxide. Linear mobile phase gradients ran from 2% to 98% solvent B over a runtime of 2 mins and at a flowrate of 1.0 mL/min. High-resolution mass spectra (HRMS) were obtained with a Waters Xevo G2 QToF detector using positive mode electron spray ionization (ESI) and scan times of 0.2 sec.

Preparative high-performance liquid chromatography (prep-HPLC) was performed using a Waters system equipped with Waters 515 pumps, a Waters 2545 binary gradient module, a Waters ACQUITY QDa detector, a Waters 2998 PDA detector, and a Waters 2767 sample manager. Separations were performed with a Waters XBridge C18 OBD, 5 µM, 30 mm x 50 mm column using focused gradients of water (solvent A) and acetonitrile (solvent B) modified with 0.1% formic acid or 5 mM ammonium hydroxide (75 mL/min flowrate). Fraction collection was triggered using the UV signal and a user-defined target mass.

NMR Spectra were recorded on a Bruker AV-III spectrometer equipped with a BBO-F cryoprobe with 1H-NMR recorded at 400 MHz and 13C-NMR at 100 MHz. Chemical shifts are expressed in ppm and referenced to residual solvent peaks according to literature values[34] with multiplicities reported as: s = singlet, d = doublet, t = triplet, q = quartet, m = multiplet, and br = broad signal. Coupling constants are reported in Hertz (Hz).

##### 2. General CIP-DEL library information.

The starting DNA construct AOP-headpiece (AOP-HP) was purchased from LGC Biosearch Technologies with anion-exchange purification. Amine building blocks for CIP-DEL library generation were purchased from Enamine, split into multiple barcoded Matrix tubes (ThermoFisher), and stored neat at 4°C until use. Prior to use, amine building blocks were dissolved in 1:1 acetonitrile : water to generate 200 mM stock solutions. Other CIP-DEL library buffers and chemicals were purchased from varying vendors and used as received. Reactions involving DNA-conjugates were performed in polypropylene vessels (Eppendorf low-bind tubes®) or 96-well polypropylene PCR plates (Eppendorf twin.tec®). Oligo tags and primer DNA was purchased from LGC Biosearch Technologies with desalting and ordered as normalized solutions in water. All DNA was ordered as 5'-phosphorylated oligos.

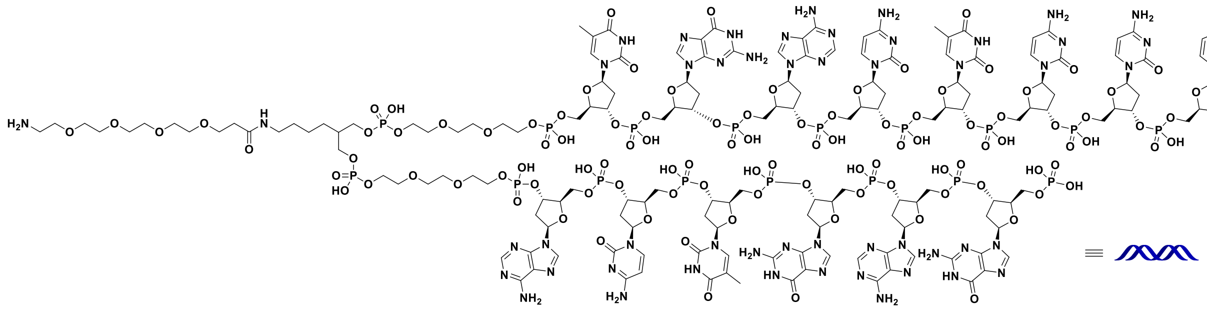

**Figure S1.** Chemical structure of AOP-HP.

UPLC-MS analysis of DNA conjugates was performed using ion-pairing chromatography with gradients of 250 mM hexafluoroisopropanol and 8 mM triethylamine in water (solvent A) and methanol (solvent B). Samples were resolved with a Waters ACQUITY system equipped with an Oligonucleotide BEH C18 1.7 µM, 2.1 x 50 mm column heated to 50°C. UV signals were recorded with an ACQUITY TUV detector at 260 nm and mass spectra were obtained with a Waters SQD2 detector in negative ion mode with a scan time of 0.2 sec. A scanning range of 500 – 3000 Da was recorded and MassLynx MaxEnt1 deconvolution was used to obtain parent ion masses. The system was run at a flowrate of 0.5 mL/min and gradients were selected based on conjugate polarity (primarily DNA length) with a runtime of 6 mins.

Prep-HPLC purification of DNA conjugates were performed on an Agilent Infinity system equipped with Infinity 1260 pump system (G5611A), Infinity 1260 autosampler (G1330B), Infinity 1290 thermostatted column heater (G5667A), Infinity 1260 diode array detector (G4212B), and Infinity 1260 fraction collector (G5664A). Mobile phase gradients of 50 mM triethylammonium acetate in water (solvent A) and methanol (solvent B) were used to resolve injected species on a Waters XBridge BEH C18 OBD, 130 Å 5 µm, 10 x 150 mm column kept at 50°C. The system was operated at a flowrate of 5 mL/min and gradients were selected based on conjugate polarity with a runtime of 20 mins. Fraction collection was triggered by UV signal at 260 nm.

Electrophoresis of DNA conjugates was performed using an Invitrogen E-Gel Power Snap system with commercial E-Gel EX 4% agarose gels. Aqueous solutions of DNA were quantified spectrophotometrically using a Thermo Scientific NanoDrop One system or a Qubit 4 Fluorometric Quantification system with the dsDNA Broad Range assay kit per the manufacturer's instructions. SDS-PAGE protein separations were performed using the Thermo Fisher XCell SureLock Mini-Cell electrophoresis system with NuPAGE 4-12% Bis-Tris precast gels. The Thermo Fisher SeeBlue Plus2 pre-stained protein ladder (P/N: LC5625) was used for protein sizing. For Western blotting, proteins were transferred to nitrocellulose membranes using the BioRad Trans-Blot Turbo Transfer system and the MagicMark XP protein ladder (P/N: LC5602) was used was protein sizing.

#### 3. General experimental procedures.

***GP1** – General procedure for HATU-mediated amide couplings with acid pre-activation:*

Carboxylic acid (1.1 equiv) was dissolved in DMF (0.10 M solution) and HATU (1.1 equiv) and DIPEA (3.0 equiv) were added. After stirring at room temperature for 10 mins, the amine (1.0 equiv) was added and stirring was continued until UPLC-MS analysis showed complete consumption of starting material. The reaction was diluted with $CH_2Cl_2$, added to a solution of saturated aqueous $NH_4Cl$, and the organic phase was isolated. The aqueous layer was further extracted with $CH_2Cl_2$ (3x) and the combined organics were dried with $MgSO_4$, filtered, and concentrated. The crude material was purified by flash chromatography on silica or preparative reverse-phase HPLC to provide pure product, unless specified otherwise.

***GP2*** *– General procedure for HATU-mediated amide couplings without acid pre-activation:*

To a solution of amine (1.0 equiv), DIPEA (3.5 equiv), and carboxylic acid (1.1 equiv) in DMF (0.10 M solution) was added HATU (1.1 equiv) at 0°C and the solution was allowed to warm to room temperature. The solution was stirred until UPLC-MS analysis showed complete consumption of starting material and the reaction was diluted with $CH_2Cl_2$ and saturated aqueous $NH_4Cl$. The organic layer was isolated, and the aqueous layer was further extracted with $CH_2Cl_2$ (3x). The combined organics were dried with $MgSO_4$, filtered, and concentrated. The crude material was purified by flash chromatography on silica or preparative reverse-phase HPLC to provide pure product, unless specified otherwise.

***GP3*** *– General procedure for Boc deprotection with TFA:*

*N*-Boc protected amine (1.0 equiv) was dissolved in $CH_2Cl_2$ (0.25 M solution) and TFA (15 equiv) was added. The reaction was stirred at room temperature until UPLC-MS analysis showed complete consumption of starting material. The reaction was diluted with $CH_2Cl_2$ and concentrated to afford crude product as the TFA salt.

***GP4*** *– General procedure for conversion of Fmoc-protected amines to acetylated amines:*

*N*-Fmoc protected amine (1.0 equiv) was dissolved in $CH_2Cl_2$ (0.05 M solution) and piperidine (15.0 equiv) was added at room temperature. Stirring was continued until UPLC-MS analysis showed complete consumption of starting material. The reaction was diluted with $CH_2Cl_2$ and concentrated. Dilution with $CH_2Cl_2$ and concentration was repeated (2x) to ensure complete removal of piperidine. The resulting material was dissolved in $CH_2Cl_2$ (0.5 M solution) and $Et_3N$ (3.3 equiv) and $Ac_2O$ (1.8 equiv) were added at room temperature. The reaction was monitored by UPLC-MS and additional portions of $Ac_2O$ were added as needed until complete consumption of starting material was observed. The reaction mixture was partitioned between $CH_2Cl_2$ and saturated aqueous $NH_4Cl$. The organic phase was isolated, and the aqueous phase was further extracted with $CH_2Cl_2$ (3x). The combined organics were dried with $MgSO_4$, filtered, and concentrated. The crude material was purified by reverse-phase HPLC to provide pure product, unless specified otherwise.

***GP5*** *– General procedure for the synthesis of di-substituted triazines from cyanuric chloride:*

Fresh 80 mM solutions of amine #1 (typically VHL-ligand / connector conjugates, 1.0 equiv) and cyanuric chloride (1.0 equiv) were prepared in 1:1 acetonitrile : 150 mM borate buffer, pH 9.4. The solution of cyanuric chloride was cooled to 0°C and the solution of amine #1 was added dropwise over 1 min. After stirring at 0°C for 30 mins, a solution of amine #2 (1.0 equiv) in $CH_3CN$ (0.1 M) was added to the reaction, followed by $K_2CO_3$ (5.0 equiv). The reaction was allowed to warm to room temperature and monitored by UPLC-MS for complete consumption of starting material. Once complete, the reaction was diluted with $CH_2Cl_2$ and saturated aqueous $NH_4Cl$ and the organic phase was isolated. The aqueous phase was further extracted with DCM (3x) and the combined organics were dried with $MgSO_4$, filtered, and concentrated under reduced pressure. The crude material was purified by flash chromatography on silica ($CH_2Cl_2$ – MeOH gradient) to provide pure product, unless stated otherwise.

***GP6*** *– General procedure for the third $S_NAr$ substitution of triazines with an amine:*

The di-substituted triazine with one remaining chloride (1.0 equiv) was dissolved in DMF (0.05 M solution) and amine #3 (4 equiv) and $K_2CO_3$ (8 equiv) were added. The solution was heated to 50°C and monitored by UPLC-MS until complete consumption of the di-substituted triazine was observed. The reaction mixture was then partitioned between $CH_2Cl_2$ and saturated aqueous $NH_4Cl$. The organic phase was isolated, and the aqueous phase was further extracted with DCM (3x). The combined organics were dried with $MgSO_4$, filtered, and concentrated. The crude material was purified by reverse-phase HPLC to provide pure product, unless stated otherwise.

#### 4. Synthesis of modified VHL ligands for incorporation in CIP-DEL.

**Scheme S1.** Synthesis of alkyne-functionalized amine **S6**.

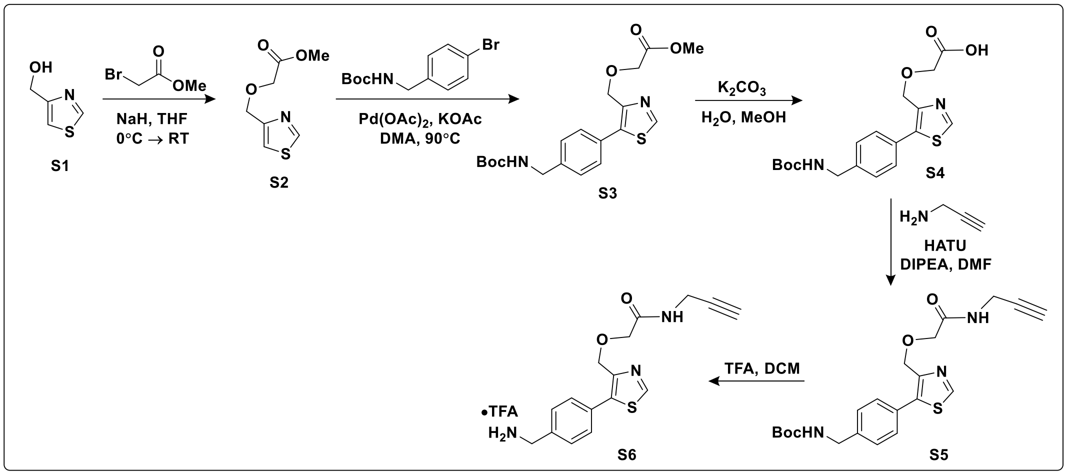


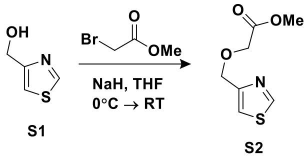


***Synthesis of S2:*** To a round bottom flask was added 4-hydroxymethylthiazole (**S1**) (2.0 g, 17.4 mmol, 1.0 equiv) and THF (22 mL). The solution was cooled to 0°C and NaH (60% in mineral oil, 0.903 g, 22.6 mmol, 1.3 equiv) was added. After the initial bubbling had ceased (~5 mins), methyl bromoacetate (1.81 mL, 19.1 mmol, 1.1 equiv) was added and the solution was allowed to warm to room temperature. After 1.5 h, UPLC-MS analysis showed a single peak with product mass. The reaction mixture was cooled to 0°C, diluted with $CH_2Cl_2$, and quenched with the dropwise addition of MeOH (4 mL). A solution of saturated $NH_4Cl$ was added and the organic phase was isolated. The aqueous phase was further extracted with three portions of $CH_2Cl_2$ and the combined organics were dried with $MgSO_4$, filtered, and concentrated under reduced pressure. The crude product was purified by flash chromatography on silica (heptanes – ethyl acetate gradient) to provide **S2** (2.40 g, 12.8 mmol, 74% yield) as a clear light-orange oil.

**$^1$H NMR** (400 MHz, $CDCl_3$) δ = 8.65 (d, *J* = 2.4 Hz, 1H), 7.20 (d, *J* = 2.4 Hz, 1H), 4.62 (s, 2H), 4.03 (s, 2H), 3.55 (s, 3H). **$^{13}$C NMR** (101 MHz, $CDCl_3$) δ = 170.1, 153.2, 153.0, 116.3, 68.3, 67.2, 51.4. **LRMS** (ESI+) req. for $C_7H_{10}NO_3S$ $[M+H]^+$ 188.0, found 188.2.

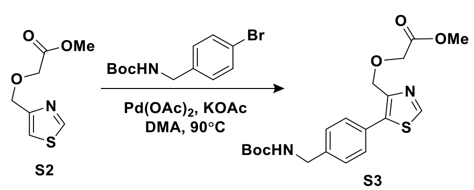


***Synthesis of S3:*** To a reaction vial was added **S2** (2.40 g, 12.8 mmol, 1.8 equiv), DMA (18 mL), tert-butyl (4-bromobenzyl)carbamate (2.0 g, 6.99 mmol, 1.0 equiv), potassium acetate (1.37 g, 14.0 mmol, 2 equiv), and palladium(II) acetate (0.016 g, 0.070 mmol, 0.01 equiv). The vial was closed with a pressure release cap and the resulting mixture was heated to 90°C in an oil bath overnight. The black slurry was cooled to room temperature, filtered through a pad of Celite®, and partitioned between $Et_2O$ and saturated aqueous

$NH_4Cl$. The organic layer was isolated and the aqueous layer was further extracted with $Et_2O$ (3x). The combined extracts were dried with $MgSO_4$, filtered, and concentrated. The crude material was purified by flash chromatography on silica (heptanes – ethyl acetate gradient) to obtain a thick orange oil that contained a mixture of product **S3** and thiazole starting material **S2**. The mixture was further purified on a 50 g reverse-phase ISCO C18 column and eluted with a gradient of water and acetonitrile, both modified with 0.1% formic acid (5 – 95% gradient), to provide product **S3** (1.15 g, 2.93 mmol, 42% yield) and thiazole starting material **S2** (0.439 g, 2.34 mmol).

**$^1$H NMR** (400 MHz, $CDCl_3$) δ = 8.55 (s, 1H), 7.35 (d, *J* = 7.9 Hz, 2H), 7.18 (d, *J* = 8.1 Hz, 2H), 5.62 (t, *J* = 6.4 Hz, 1H), 4.53 (s, 2H), 4.16 (d, *J* = 5.9 Hz, 2H), 4.06 (s, 2H), 3.53 (s, 3H), 1.29 (s, 9H). **$^{13}$C NMR** (101 MHz, $CDCl_3$) δ = 170.2, 155.8, 150.8, 147.0, 139.8, 137.7, 129.2, 128.9, 127.3, 78.8, 66.9, 66.0, 51.3, 43.6, 28.0. **HRMS** (ESI+) req. for $C_{19}H_{25}N_2O_5S$ $[M+H]^+$ 393.1479, found 393.1486.

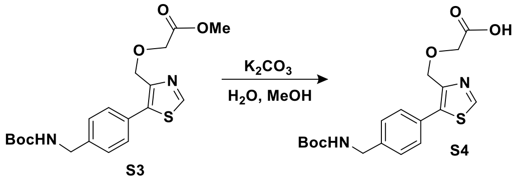


***Synthesis of S4:*** To a reaction vial containing **S3** (1.47 g, 3.74 mmol, 1.0 equiv) was added MeOH (6 mL), water (6 mL), and $K_2CO_3$ (1.55 g, 11.2 mmol, 3 equiv). After stirring at room temperature for 2 h, UPLC-MS analysis indicated complete consumption of starting material. The reaction was diluted with water and acidified with 0.2 M HCl (pH ~2). EtOAc was added and the organic phase was isolated. The aqueous phase was further extracted with EtOAc (3x) and the combined organics were dried with $MgSO_4$, filtered, and concentrated to give a white foam that could be broken up into a fine powder of **S4** (1.29 g, 3.40 mmol, 91% yield).

**$^1$H NMR** (400 MHz, MeOD) δ = 8.90 (s, 1H), 7.51 (d, *J* = 8.3 Hz, 2H), 7.35 (d, *J* = 8.0 Hz, 2H), 4.65 (s, 2H), 4.26 (s, 2H), 4.17 (s, 2H), 1.45 (s, 9H). **HRMS** (ESI+) req. for $C_{18}H_{23}N_2O_5S$ $[M+H]^+$ 379.1322, found 379.1325.

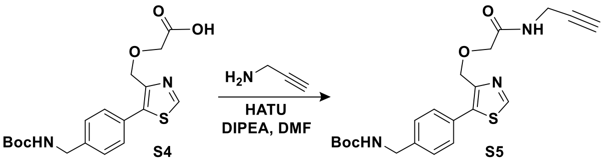


***Synthesis of S5:*** To a round bottom flask was added acid **S4** (1.09 g, 2.87 mmol, 1.0 equiv), DMF (10 mL), DIPEA (1.75 mL, 10.0 mmol, 3.5 equiv), and HATU (1.20 g, 3.15 mmol, 1.1 equiv) at 0°C. After stirring for 15 mins, propargyl amine (0.331 mL, 5.16 mmol, 1.8 equiv) was added and stirring was continued until UPLC-MS analysis showed complete consumption of starting material. The reaction was diluted with EtOAc and saturated aqueous $NH_4Cl$, then warmed to room temperature. The organic phase was isolated and the aqueous phase was further extracted with EtOAc (3x). The combined organic extracts were dried with $MgSO_4$, filtered, concentrated, and purified by flash chromatography on silica (heptanes – ethyl acetate gradient) to afford an impure product mixture. This material was further purified on a 50 g reverse-phase ISCO C18 column and eluted with a gradient of water and acetonitrile, both modified with 0.1% formic acid (5 – 95% gradient), to provide product **S5** (1.15 g, 2.77 mmol, 82% yield).

**$^{1}$H NMR** (400 MHz, $CDCl_3$) δ = 8.61 (s, 1H), 7.36 (t, *J* = 5.5 Hz, 1H), 7.22 (dd, *J* = 10.0, 8.8 Hz, 4H), 5.76 (t, *J* = 6.3 Hz, 1H), 4.49 (s, 2H), 4.17 (d, *J* = 6.2 Hz, 2H), 3.95 – 3.78 (m, 4H), 2.15 (t, *J* = 2.9 Hz, 1H), 1.29 (s, 9H). **HRMS** (ESI+) req. for $C_{21}H_{26}N_3O_4S$ $[M+H]^+$ 416.1639, found 416.1644.

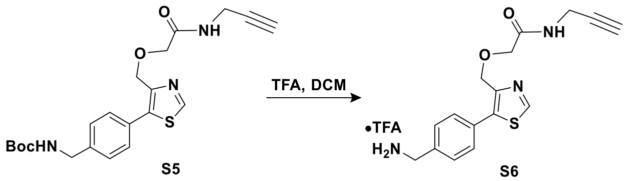


***Synthesis of S6:*** Starting material **S5** (1.15 g, 2.77 mmol, 1.0 equiv) was deprotected according to **GP3** to afford product **S6** (1.19 g, 2.77 mmol, quantitative yield).

**$^{1}$H NMR** (400 MHz, MeOD) δ = 9.09 (s, 1H), 7.61 (d, *J* = 8.1 Hz, 2H), 7.55 (d, *J* = 8.2 Hz, 2H), 6.19 (s, 3H), 4.67 (s, 2H), 4.23 (s, 2H), 4.03 (s, 2H), 3.99 (d, *J* = 2.9 Hz, 2H), 2.52 (t, *J* = 2.6 Hz, 1H). **$^{13}$C NMR** (101 MHz, MeOD) δ = 171.8, 154.9, 148.0, 138.4, 135.4, 132.0, 131.1, 130.9, 80.4, 72.3, 70.4, 67.1, 49.4, 49.2, 49.0, 48.8, 48.6, 43.9, 29.1. **HRMS** (ESI+) req. for $C_{16}H_{18}N_3O_2S$ $[M+H]^+$ 316.1114, found 316.1122.

**Scheme S2.** Synthesis of alkyne functionalized VHL ligand **S12**.

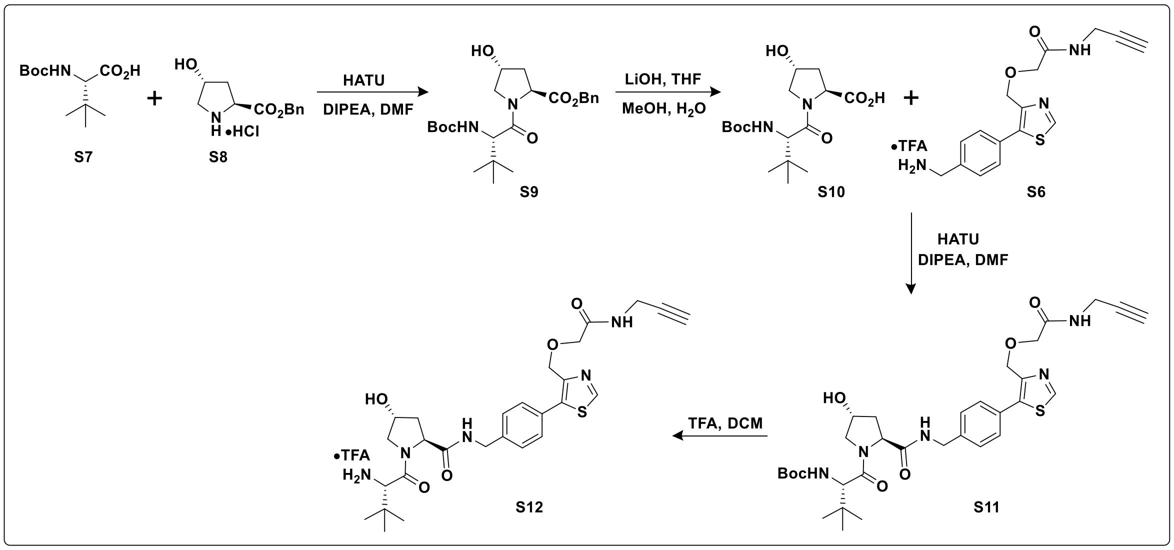


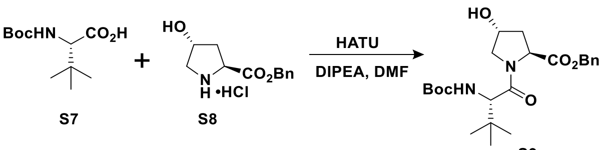


***Synthesis of S9:*** A solution of H-Hyp-OBzl·HCl (**S8**) (2.50 g, 9.70 mmol, 1.0 equiv), Boc-Tle-OH (**S7**) (2.36 g, 10.2 mmol, 1.05 equiv), and DIPEA (3.73 ml, 21.3 mmol, 2.2 equiv) in DMF (24.3 mL) was cooled to 0°C. HATU (3.87 g, 10.2 mmol, 1.05 equiv) was added and the reaction was monitored by UPLC-MS analysis. Additional DIPEA (0.5 mL), Boc-Tle-OH (**S7**) (1.0 g), and HATU (1.0 g) were added portionwise to give complete consumption of starting material **S8**. The reaction was diluted with $Et_2O$ and water, and warmed to room temperature. The organic phase was isolated and the aqueous phase was further extracted with $Et_2O$ (3x). The combined organics were dried with $MgSO_4$, filtered, and concentrated under reduce pressure. The crude material was purified by flash chromatography on silica (heptanes – ethyl acetate gradient) to afford **S9** (3.94 g, 9.07 mmol, 93% yield).

**$^1$H NMR** (400 MHz, $CDCl_3$) δ = 7.34 – 7.19 (m, 5H), 5.40 (d, *J* = 9.4 Hz, 1H), 5.15 – 5.02 (m, 2H), 4.64 (t, *J* = 8.4 Hz, 1H), 4.40 (s, 1H), 4.15 (d, *J* = 9.4 Hz, 1H), 3.85 (d, *J* = 11.2 Hz, 2H), 3.62 (dd, *J* = 11.2, 4.0 Hz, 1H), 2.26 (dd, *J* = 13.6, 8.1 Hz, 1H), 1.90 (ddd, *J* = 13.3, 9.1, 4.4 Hz, 1H), 1.33 (s, 9H), 0.92 (s, 9H). **$^{13}$C NMR** (101 MHz, $CDCl_3$) δ = 171.6, 170.8, 155.5, 135.2, 128.1, 127.9, 127.9, 79.2, 69.4, 66.4, 58.3, 57.8, 56.0, 37.1, 35.6, 28.0, 25.9. **HRMS** (ESI+) req. for $C_{23}H_{34}N_2NaO_6$ $[M+Na]^+$ 457.2309, found 457.2318.

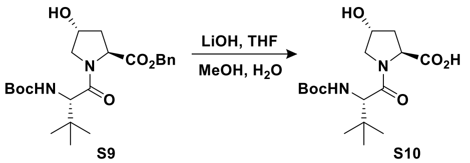


***Synthesis of S10:*** To a round bottom flask containing **S9** (3.94 g, 9.07 mmol, 1.0 equiv) was added THF (10 mL), methanol (5 mL), water (5 mL), and $LiOH{\cdot}H_2O$ (0.684 g, 16.3 mmol, 1.8 equiv). After stirring for 1 h at room temperature, UPLC-MS analysis showed complete consumption of starting material **S9** and the reaction was diluted with 0.2 M HCl (to pH ~2) and EtOAc. The organic phase was isolated and the aqueous phase was further extracted with EtOAc (3x). The combined organics were dried with $Na_2SO_4$, filtered, and concentrated. The crude material was purified by flash chromatography on silica (heptanes – ethyl acetate gradient) to give **S10** as a white foam that could be broken up into a powder (2.76 g, 8.01 mmol, 88% yield).

**$^1$H NMR** (400 MHz, MeOD) δ = 4.61 – 4.40 (m, 2H), 4.31 (s, 1H), 3.93 – 3.66 (m, 2H), 2.31 (dd, *J* = 13.6, 8.0 Hz, 1H), 2.06 (ddd, *J* = 13.2, 8.7, 4.2 Hz, 1H), 1.43 (s, 9H), 1.03 (s, 9H). **$^{13}$C NMR** (101 MHz, MeOD) δ = 174.9, 172.5, 157.4, 80.5, 70.7, 59.8, 59.1, 57.4, 38.3, 36.7, 28.7, 26.8. **HRMS** (ESI+) req. for $C_{16}H_{26}N_2NaO_6$ $[M+Na]^+$ 367.1840, found 367.1845.

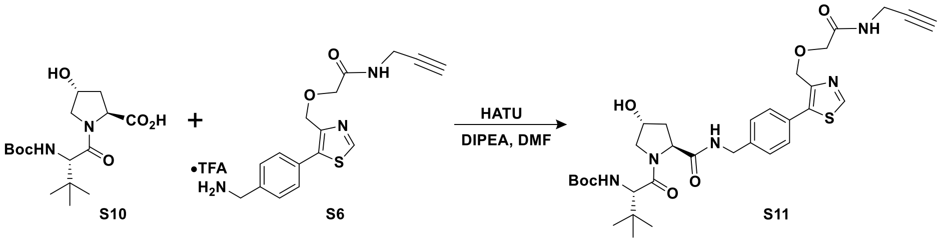


***Synthesis of S11:*** A solution of **S6** (1.19 g, 2.77 mmol, 1 equiv) in DMF (6 mL) was cooled to 0°C. DIPEA (2.90 mL, 16.6 mmol, 6 equiv), **S10** (1.05 g, 3.05 mmol, 1.1 equiv), and HATU (1.16 g, 3.05 mmol, 1.1 equiv) were added and the solution was allowed to warm to room temperature overnight. Analysis by UPLC-MS showed remaining starting material **S6** and additional **S10** (0.191 g, 0.554 mmol, 0.2 equiv) and HATU (0.211 g, 0.554 mmol, 0.2 equiv) were added. Subsequent UPLC-MS analysis showed complete consumption of starting material and the reaction was cooled to 0°C and diluted with EtOAc and saturate aqueous $NH_4Cl$. The organic phase was isolated and the aqueous phase was further extracted with EtOAc (3x). The combined organics were dried with $MgSO_4$, filtered, and concentrated. The crude material was purified by flash chromatography on silica ($CH_2Cl_2$ – MeOH gradient) to provide a large fraction of pure product and a small amount of impure material which was re-purified in the same way. The combined product **S11** (1.49 g, 2.33 mmol, 84% yield) was isolated as a white foam.

**$^1$H NMR** (400 MHz, MeOD) δ = 8.99 (s, 1H), 8.68 (t, *J* = 6.0 Hz, 1H), 7.48 (s, 4H), 6.37 (d, *J* = 9.3 Hz, 1H), 4.67 (s, 2H), 4.63 – 4.47 (m, 3H), 4.40 – 4.27 (m, 2H), 4.03 (s, 2H), 4.00 (d, *J* = 2.4 Hz, 2H), 3.89 (d, *J* = 11.2 Hz, 1H), 3.80 (dd, *J* = 11.2, 3.9 Hz, 1H), 2.59 (t, *J* = 2.5 Hz, 1H), 2.23 (dd, *J* = 13.2, 7.8 Hz, 1H), 2.09 (ddd, *J* = 13.3, 9.0, 4.5 Hz, 1H), 1.44 (s, 10H), 1.01 (s, 10H). **$^{13}$C NMR** (101 MHz, MeOD) δ = 174.4, 172.8,

171.7, 157.7, 153.9, 148.5, 141.0, 139.0, 130.6, 130.3, 129.0, 80.5, 72.3, 71.0, 70.4, 67.4, 60.8, 60.3, 58.0, 43.6, 38.8, 36.8, 29.0, 28.7, 26.9. **HRMS** (ESI+) req. for $C_{32}H_{44}N_5O_7$ $[M+H]^+$ 642.2956, found 642.2979.

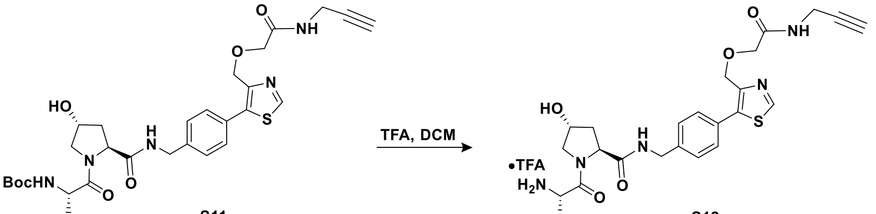


S11 S12

***Synthesis of S12:*** Boc-protected **S11** (1.49 g, 2.33 mmol, 1.0 equiv) was deprotected according to **GP3** to provide **S12** (1.53 g, 2.33 mmol, quantitative yield).

**$^1$H NMR** (400 MHz, MeOD) δ = 9.16 (s, 1H), 7.47 (d, *J* = 8.0 Hz, 2H), 7.41 (d, *J* = 8.2 Hz, 2H), 4.75 – 4.62 (m, 3H), 4.55 (d, *J* = 15.5 Hz, 2H), 4.36 (d, *J* = 15.6 Hz, 1H), 4.05 (d, *J* = 13.6 Hz, 3H), 3.98 (d, *J* = 2.8 Hz, 2H), 3.86 (d, *J* = 11.2 Hz, 1H), 3.71 (dd, *J* = 11.1, 3.5 Hz, 1H), 2.51 (t, *J* = 2.6 Hz, 1H), 2.33 (dd, *J* = 13.2, 7.5 Hz, 1H), 2.10 (ddd, *J* = 13.4, 9.6, 4.1 Hz, 1H), 1.36 (dd, *J* = 6.8, 4.8 Hz, 1H), 1.13 (s, 9H). **$^{13}$C NMR** (101 MHz, MeOD) δ = 174.0, 171.8, 168.7, 161.3, 160.9, 160.5, 160.2, 154.9, 147.0, 141.3, 139.3, 130.6, 129.7, 129.1, 121.1, 118.2, 115.4, 112.5, 80.3, 72.3, 71.2, 70.5, 67.0, 61.0, 60.4, 58.0, 49.6, 49.4, 49.2, 49.0, 48.8, 48.6, 48.4, 43.7, 38.9, 35.7, 29.1, 26.7. **HRMS** (ESI+) req. for $C_{27}H_{36}N_5O_5S$ $[M+H]^+$ 542.2432, found 542.2443.

**Scheme S3.** Synthesis of alkyne-functionalized VHL ligands with Fmoc-protected amino acid connectors.

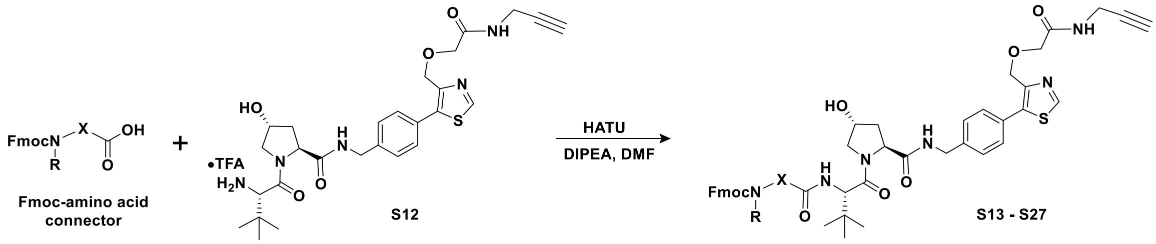


***Synthesis of S13 – S27:*** Coupling of Fmoc-amino acid connectors was performed according to **GP2** and products were purified by reverse-phase HPLC using gradients of solvent A (water) and solvent B (acetonitrile) both modified with 0.1% formic acid to generate the following compounds:

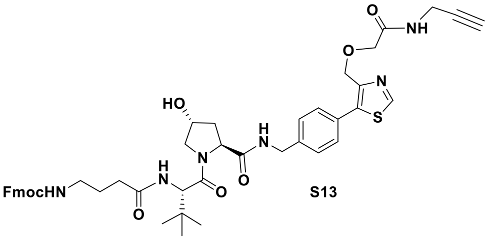


S13

**$^1$H NMR** (400 MHz, MeOD) δ = 8.97 (s, 1H), 7.78 (d, *J* = 7.5 Hz, 2H), 7.63 (d, *J* = 7.5 Hz, 2H), 7.52 – 7.42 (m, 4H), 7.37 (t, *J* = 7.4 Hz, 2H), 7.29 (t, *J* = 7.3 Hz, 2H), 4.68 – 4.53 (m, 5H), 4.50 (s, 1H), 4.40 – 4.30 (m, 3H), 4.18 (t, *J* = 6.7 Hz, 1H), 4.01 (s, 2H), 3.99 (d, *J* = 2.6 Hz, 2H), 3.91 (d, *J* = 11.0 Hz, 1H), 3.80 (dd, *J* = 11.0, 3.9 Hz, 1H), 3.12 (t, *J* = 7.0 Hz, 2H), 2.58 (t, *J* = 2.7 Hz, 1H), 2.33 – 2.16 (m, 3H), 2.08 (ddd, *J* = 13.3, 9.1, 4.5 Hz, 1H), 1.76 (p, *J* = 7.6 Hz, 2H), 1.04 (s, 10H). **$^{13}$C NMR** (101 MHz, MeOD) δ = 175.4, 174.5, 172.4, 171.9, 158.9, 154.0, 148.6, 145.4, 142.6, 141.2, 139.1, 130.7, 130.4, 129.1, 128.8, 128.1, 126.1,

120.9, 80.5, 72.2, 71.1, 70.5, 67.5, 67.5, 60.8, 59.2, 58.0, 43.7, 41.1, 38.9, 36.5, 33.8, 29.0, 27.3, 27.1. **HRMS** (ESI+) req. for $C_{46}H_{53}N_6O_8S$ $[M+H]^+$ 849.3640, found 849.3651.

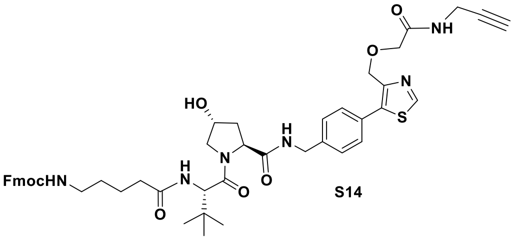


S14

**[1]H NMR** (400 MHz, MeOD) δ = 8.97 (s, 1H), 7.84 – 7.73 (m, 3H), 7.66 – 7.56 (m, 2H), 7.46 (dd, *J* = 12.2, 8.1 Hz, 4H), 7.37 (t, *J* = 7.5 Hz, 2H), 7.29 (t, *J* = 7.4 Hz, 2H), 4.64 (s, 2H), 4.63 – 4.45 (m, 4H), 4.34 (dd, *J* = 11.3, 4.4 Hz, 3H), 4.18 (t, *J* = 6.8 Hz, 1H), 4.02 (s, 2H), 4.00 – 3.96 (m, 2H), 3.91 (d, *J* = 11.0 Hz, 1H), 3.80 (dd, *J* = 11.0, 3.9 Hz, 1H), 3.12 (dd, *J* = 6.6, 5.8 Hz, 2H), 2.59 (t, *J* = 2.6 Hz, 1H), 2.36 – 2.14 (m, 3H), 2.08 (ddd, *J* = 13.3, 9.1, 4.5 Hz, 1H), 1.62 (p, *J* = 7.6 Hz, 2H), 1.50 (p, *J* = 7.0 Hz, 2H), 1.03 (s, 10H). **[13]C NMR** (101 MHz, MeOD) δ = 175.8, 174.5, 172.3, 171.9, 158.9, 154.0, 148.6, 145.4, 142.6, 141.1, 139.1, 130.7, 130.4, 129.1, 128.8, 128.1, 126.2, 120.9, 80.5, 72.2, 71.1, 70.5, 67.6, 67.5, 60.8, 59.0, 58.0, 43.7, 41.2, 38.9, 36.5, 36.2, 30.4, 29.0, 27.1, 24.1. **HRMS** (ESI+) req. for $C_{47}H_{55}N_6O_8S$ $[M+H]^+$ 863.3797, found 863.3824.

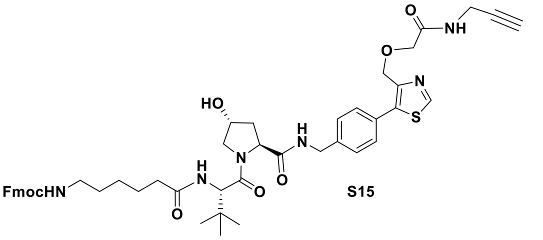


S15

**[1]H NMR** (400 MHz, MeOD) δ = 8.97 (s, 1H), 7.78 (d, *J* = 7.3 Hz, 3H), 7.63 (d, *J* = 7.4 Hz, 2H), 7.46 (dd, *J* = 12.7, 8.2 Hz, 4H), 7.38 (t, *J* = 7.5 Hz, 2H), 7.29 (t, *J* = 7.5 Hz, 2H), 4.69 – 4.60 (m, 3H), 4.56 (dd, *J* = 15.4, 5.9 Hz, 2H), 4.49 (s, 1H), 4.38 – 4.28 (m, 3H), 4.17 (t, *J* = 6.8 Hz, 1H), 4.02 (s, 2H), 3.99 (d, *J* = 2.4 Hz, 2H), 3.91 (d, *J* = 10.9 Hz, 1H), 3.79 (dd, *J* = 10.9, 3.9 Hz, 1H), 3.10 (dd, *J* = 7.0, 6.1 Hz, 2H), 2.58 (t, *J* = 2.6 Hz, 1H), 2.38 – 2.16 (m, 4H), 2.07 (ddd, *J* = 13.3, 9.1, 4.5 Hz, 1H), 1.62 (p, *J* = 7.3 Hz, 2H), 1.49 (p, *J* = 7.3 Hz, 2H), 1.33 (p, *J* = 8.5 Hz, 2H), 1.03 (s, 10H). **[13]C NMR** (101 MHz, MeOD) δ = 176.0, 174.5, 172.4, 171.9, 158.9, 154.0, 148.6, 145.4, 142.6, 141.1, 139.1, 130.7, 130.4, 129.1, 128.8, 128.1, 126.2, 120.9, 80.5, 72.2, 71.1, 70.5, 67.5, 67.5, 60.8, 59.0, 58.0, 43.7, 41.6, 38.9, 36.5, 36.5, 30.6, 29.0, 27.4, 27.0, 26.7. **HRMS** (ESI+) req. for $C_{48}H_{57}N_6O_8S$ $[M+H]^+$ 877.3953, found 877.3964.

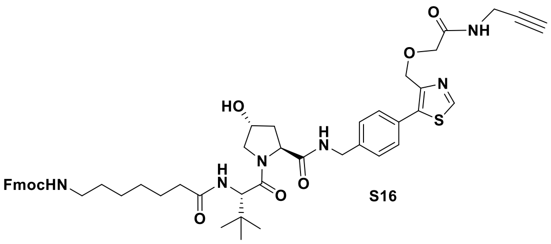


S16

**[1]H NMR** (400 MHz, MeOD) δ = 8.97 (s, 1H), 7.78 (dd, *J* = 8.4, 4.4 Hz, 3H), 7.62 (d, *J* = 7.4 Hz, 2H), 7.45 (dd, *J* = 12.7, 8.3 Hz, 4H), 7.37 (t, *J* = 7.5 Hz, 2H), 7.29 (t, *J* = 7.4 Hz, 2H), 4.63 (s, 3H), 4.60 – 4.52 (m,

2H), 4.49 (s, 1H), 4.37 – 4.29 (m, 3H), 4.17 (t, *J* = 6.8 Hz, 1H), 4.01 (s, 2H), 3.99 (d, *J* = 2.4 Hz, 2H), 3.91 (d, *J* = 11.0 Hz, 1H), 3.80 (dd, *J* = 11.1, 3.9 Hz, 1H), 3.09 (q, *J* = 6.5 Hz, 2H), 2.58 (t, *J* = 2.5 Hz, 1H), 2.33 – 2.17 (m, 3H), 2.08 (ddd, *J* = 13.3, 9.2, 4.5 Hz, 1H), 1.61 (p, *J* = 7.2 Hz, 2H), 1.48 (p, *J* = 6.8 Hz, 2H), 1.39 – 1.28 (m, 4H), 1.03 (s, 9H). **$^{13}$C NMR** (101 MHz, MeOD) δ = 176.1, 174.5, 172.3, 171.8, 158.9, 153.9, 148.6, 145.4, 142.6, 141.1, 139.1, 130.7, 130.4, 129.1, 128.8, 128.1, 126.2, 120.9, 80.5, 72.2, 71.1, 70.5, 67.5, 67.5, 60.8, 59.0, 58.0, 54.8, 43.7, 41.7, 38.9, 36.5, 36.5, 30.7, 29.9, 29.0, 27.5, 27.0, 27.0, 26.9. **HRMS** (ESI+) req. for $C_{49}H_{59}N_6O_8S$ $[M+H]^+$ 891.4110, found 891.4141.

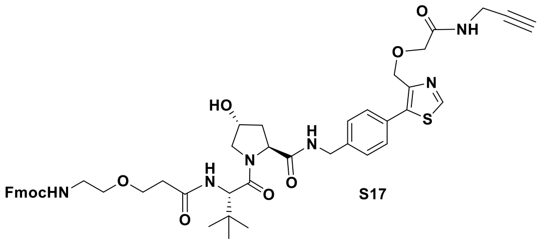


S17

**$^1$H NMR** (400 MHz, MeOD) δ = 8.93 (s, 1H), 7.77 (d, *J* = 7.5 Hz, 2H), 7.62 (d, *J* = 7.6 Hz, 2H), 7.49 – 7.39 (m, 4H), 7.37 (t, *J* = 7.4 Hz, 2H), 7.28 (t, *J* = 7.4 Hz, 2H), 4.70 – 4.45 (m, 6H), 4.35 – 4.26 (m, 2H), 4.15 (t, *J* = 6.7 Hz, 1H), 4.02 – 3.95 (m, 4H), 3.89 (d, *J* = 11.0 Hz, 1H), 3.79 (dd, *J* = 10.9, 3.8 Hz, 1H), 3.75 – 3.62 (m, 2H), 3.50 (t, *J* = 5.5 Hz, 2H), 3.29 (d, *J* = 5.3 Hz, 1H), 2.58 (t, *J* = 2.5 Hz, 1H), 2.56 – 2.43 (m, 2H), 2.25 – 2.14 (m, 1H), 2.14 – 2.01 (m, 1H), 1.03 (s, 9H). **$^{13}$C NMR** (101 MHz, MeOD) δ = 174.4, 173.9, 172.2, 171.8, 158.8, 153.9, 148.5, 145.3, 142.6, 141.1, 139.0, 130.6, 130.3, 129.1, 128.8, 128.2, 126.2, 121.0, 80.5, 72.2, 71.1, 70.8, 70.5, 68.0, 67.7, 67.5, 60.8, 59.0, 58.0, 54.8, 43.7, 41.8, 38.9, 37.4, 36.8, 29.0, 27.0. **HRMS** (ESI+) req. for $C_{47}H_{55}N_6O_9S$ $[M+H]^+$ 879.3746, found 879.3840.

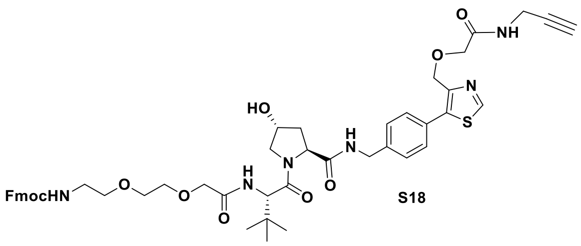


S18

**$^1$H NMR** (400 MHz, MeOD) δ = 8.94 (s, 1H), 7.78 (d, *J* = 7.6 Hz, 2H), 7.61 (dd, *J* = 7.7, 3.5 Hz, 2H), 7.43 (s, 3H), 7.37 (t, *J* = 7.4 Hz, 3H), 7.28 (t, *J* = 7.3 Hz, 2H), 4.76 – 4.47 (m, 6H), 4.42 – 4.23 (m, 3H), 4.15 (t, *J* = 6.9 Hz, 1H), 4.06 – 3.94 (m, 6H), 3.83 (q, *J* = 11.0 Hz, 2H), 3.70 (t, *J* = 4.2 Hz, 2H), 3.64 (t, *J* = 5.7 Hz, 2H), 3.54 (t, *J* = 5.5 Hz, 2H), 3.35 (s, 1H), 2.58 (s, 1H), 2.29 – 2.03 (m, 2H), 1.04 (s, 10H). **$^{13}$C NMR** (101 MHz, MeOD) δ = 174.3, 172.2, 171.9, 171.7, 158.8, 153.9, 148.6, 145.3, 142.6, 141.0, 139.0, 130.7, 129.1, 128.8, 128.2, 126.2, 121.0, 80.5, 72.4, 72.2, 71.2, 71.2, 71.0, 71.0, 70.5, 67.6, 67.5, 60.8, 58.1, 43.7, 41.8, 38.9, 37.3, 29.0, 27.0. **HRMS** (ESI+) req. for $C_{48}H_{57}N_6O_{10}S$ $[M+H]^+$ 909.3851, found 909.3843.

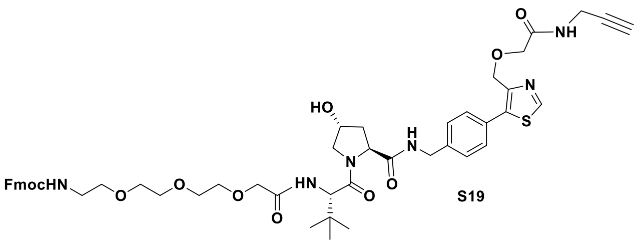


S19

**$^{1}$H NMR** (400 MHz, MeOD) δ = 8.94 (s, 1H), 7.77 (d, *J* = 7.5 Hz, 2H), 7.62 (d, *J* = 7.4 Hz, 2H), 7.44 (dd, *J* = 10.1, 7.6 Hz, 4H), 7.37 (t, *J* = 7.4 Hz, 2H), 7.29 (t, *J* = 7.5 Hz, 2H), 4.72 – 4.61 (m, 3H), 4.61 – 4.51 (m, 2H), 4.49 (s, 1H), 4.37 – 4.28 (m, 3H), 4.16 (t, *J* = 6.9 Hz, 1H), 4.00 (h, *J* = 6.1 Hz, 6H), 3.87 (d, *J* = 11.1 Hz, 1H), 3.79 (dd, *J* = 11.0, 3.8 Hz, 1H), 3.69 – 3.56 (m, 8H), 3.48 (t, *J* = 5.5 Hz, 2H), 3.25 (t, *J* = 5.5 Hz, 2H), 2.59 (t, *J* = 2.6 Hz, 1H), 2.27 – 2.16 (m, 1H), 2.08 (ddd, *J* = 13.3, 9.2, 4.4 Hz, 1H), 1.03 (s, 9H). **$^{13}$C NMR** (101 MHz, MeOD) δ = 174.4, 172.1, 171.8, 171.6, 158.8, 153.9, 148.6, 145.3, 142.6, 141.1, 139.0, 130.7, 130.4, 129.1, 128.8, 128.2, 126.2, 121.0, 80.5, 72.2, 71.6, 71.5, 71.4, 71.1, 71.0, 70.9, 70.5, 67.6, 67.5, 60.8, 58.1, 43.7, 41.8, 38.9, 37.1, 29.0, 27.0. **HRMS** (ESI+) req. for $C_{50}H_{61}N_6O_{11}S$ $[M+H]^+$ 953.4114, found 953.4130.

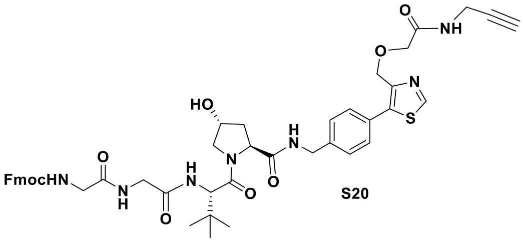


S20

**$^{1}$H NMR** (400 MHz, MeOD) δ = 8.92 (s, 1H), 7.76 (d, *J* = 7.4 Hz, 2H), 7.62 (dd, *J* = 7.7, 2.5 Hz, 2H), 7.46 – 7.39 (m, 4H), 7.35 (t, *J* = 7.4 Hz, 2H), 7.27 (t, *J* = 7.3 Hz, 2H), 4.66 – 4.45 (m, 7H), 4.38 – 4.28 (m, 3H), 4.17 (t, *J* = 6.9 Hz, 1H), 4.02 – 3.96 (m, 5H), 3.96 – 3.73 (m, 7H), 2.58 (t, *J* = 2.5 Hz, 1H), 2.21 (dd, *J* = 13.0, 7.5 Hz, 1H), 2.07 (ddd, *J* = 13.3, 9.0, 4.5 Hz, 1H), 1.03 (s, 10H). **$^{13}$C NMR** (101 MHz, MeOD) δ = 174.4, 172.8, 172.0, 171.8, 171.2, 159.0, 153.9, 148.5, 145.3, 145.2, 142.6, 141.1, 139.0, 130.6, 130.3, 129.1, 128.8, 128.2, 126.2, 120.9, 80.5, 72.2, 71.1, 70.5, 68.1, 67.5, 60.8, 59.1, 58.0, 45.1, 43.6, 43.4, 38.9, 36.8, 29.0, 27.0. **HRMS** (ESI+) req. for $C_{46}H_{52}N_7O_9S$ $[M+H]^+$ 878.3542, found 878.3572.

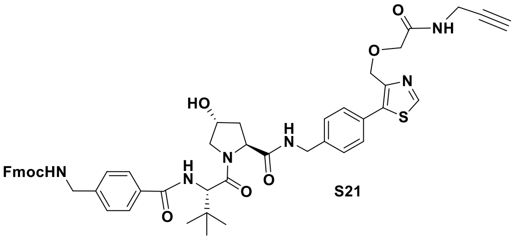


S21

**$^{1}$H NMR** (400 MHz, MeOD) δ = 8.95 (s, 1H), 7.81 – 7.73 (m, 4H), 7.62 (d, *J* = 7.2 Hz, 2H), 7.54 – 7.41 (m, 4H), 7.32 (ddt, *J* = 19.3, 14.7, 7.4 Hz, 6H), 4.68 – 4.55 (m, 4H), 4.52 (s, 1H), 4.49 – 4.37 (m, 3H), 4.36 – 4.25 (m, 3H), 4.17 (t, *J* = 6.6 Hz, 1H), 4.08 – 3.93 (m, 5H), 3.85 (dd, *J* = 11.1, 3.8 Hz, 1H), 2.57 (s, 1H), 2.24 (dd, *J* = 13.3, 7.6 Hz, 1H), 2.10 (ddd, *J* = 13.4, 9.4, 4.4 Hz, 1H), 1.10 (s, 9H). **$^{13}$C NMR** (101 MHz, MeOD) δ = 174.4, 172.3, 171.8, 169.6, 159.0, 153.9, 148.5, 145.3, 144.9, 142.6, 141.1, 139.0, 133.9, 131.0, 130.7, 130.4, 129.1, 128.8, 128.7, 128.3, 128.1, 126.1, 120.9, 80.5, 72.2, 71.1, 70.5, 67.6, 67.5, 60.9, 59.3, 58.2, 45.0, 43.7, 38.9, 37.3, 29.0, 27.1. **HRMS** (ESI+) req. for $C_{50}H_{53}N_6O_8S$ $[M+H]^+$ 897.3640, found 897.3644.

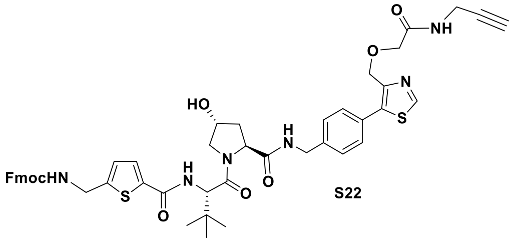


S22

**$^{1}$H NMR** (400 MHz, MeOD) δ = 8.95 (s, 1H), 8.70 (t, *J* = 6.5 Hz, 1H), 7.76 (d, *J* = 7.8 Hz, 2H), 7.61 (d, *J* = 7.9 Hz, 2H), 7.47 (dd, *J* = 12.2, 8.4 Hz, 4H), 7.35 (t, *J* = 7.7 Hz, 2H), 7.27 (t, *J* = 7.4 Hz, 2H), 6.93 (d, *J* = 3.9 Hz, 1H), 4.68 – 4.55 (m, 4H), 4.53 – 4.41 (m, 3H), 4.40 – 4.28 (m, 3H), 4.17 (t, *J* = 7.1 Hz, 1H), 4.05 – 3.90 (m, 5H), 3.82 (dd, *J* = 11.3, 2.8 Hz, 1H), 2.57 (t, *J* = 2.5 Hz, 1H), 2.23 (dd, *J* = 13.4, 7.5 Hz, 1H), 2.09 (ddd, *J* = 13.6, 9.4, 4.4 Hz, 1H), 1.08 (s, 9H). **$^{13}$C NMR** (101 MHz, MeOD) δ = 174.4, 172.2, 171.8, 163.9, 158.7, 153.9, 150.4, 148.6, 145.2, 142.6, 141.1, 139.0, 138.0, 130.7, 130.4, 130.2, 129.1, 128.8, 128.2, 126.9, 126.2, 120.9, 80.5, 72.2, 71.1, 70.5, 67.8, 67.5, 60.9, 59.2, 58.2, 43.7, 40.6, 38.9, 37.3, 29.0, 27.1. **HRMS** (ESI+) req. for $C_{48}H_{51}N_6O_8S$ $[M+H]^+$ 903.3204, found 903.3207.

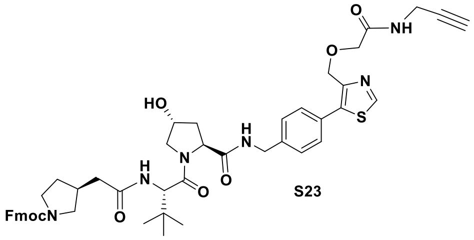


S23

**$^{1}$H NMR** (400 MHz, $CDCl_3$) δ = 8.90 (s, 1H), 7.73 (d, *J* = 7.5 Hz, 2H), 7.68 (t, *J* = 6.1 Hz, 1H), 7.59 (t, *J* = 8.7 Hz, 1H), 7.51 (t, *J* = 6.5 Hz, 2H), 7.36 (q, *J* = 8.3, 7.8 Hz, 4H), 7.32 – 7.21 (m, 5H), 6.55 (dd, *J* = 25.3, 8.8 Hz, 1H), 4.78 (t, *J* = 8.3 Hz, 1H), 4.69 – 4.45 (m, 6H), 4.34 – 4.09 (m, 4H), 4.08 – 4.00 (m, 5H), 3.76 – 3.60 (m, 2H), 3.50 (t, *J* = 8.3 Hz, 1H), 3.34 (qd, *J* = 11.8, 10.4, 7.2 Hz, 1H), 3.21 – 3.09 (m, 1H), 2.69 – 2.44 (m, 1H), 2.42 – 2.30 (m, 2H), 2.28 – 2.17 (m, 2H), 2.13 – 2.02 (m, 1H), 1.67 – 1.52 (m, 1H), 0.96 (s, 9H). **$^{13}$C NMR** (101 MHz, $CDCl_3$) δ = 171.8, 171.6, 171.0, 169.4, 155.0, 152.2, 146.2, 144.1, 141.4, 140.3, 137.9, 129.6, 128.3, 127.8, 127.1, 125.1, 120.1, 116.5, 79.6, 71.5, 70.6, 70.2, 70.1, 67.3, 66.5, 59.0, 57.5, 57.3, 52.0, 51.0, 47.3, 45.7, 45.3, 43.1, 39.5, 37.4, 36.6, 36.1, 35.9, 35.5, 35.2, 32.0, 30.8, 28.6, 26.6, 26.5. **HRMS** (ESI+) req. for $C_{48}H_{55}N_6O_8S$ $[M+H]^+$ 875.3797, found 875.3829.

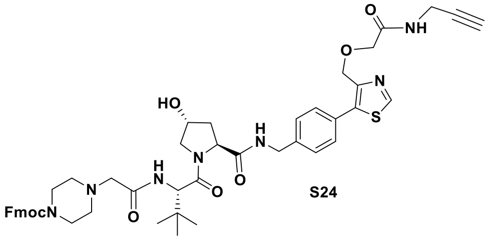


S24

**$^{1}$H NMR** (400 MHz, MeOD) δ = 8.93 (s, 1H), 7.78 (d, *J* = 7.6 Hz, 2H), 7.57 (d, *J* = 7.4 Hz, 2H), 7.49 (dd, *J* = 13.3, 8.3 Hz, 4H), 7.37 (t, *J* = 7.5 Hz, 2H), 7.29 (t, *J* = 7.5 Hz, 2H), 4.63 (s, 3H), 4.60 – 4.53 (m, 2H), 4.53 – 4.45 (m, 3H), 4.35 (d, *J* = 15.6 Hz, 1H), 4.19 (t, *J* = 5.9 Hz, 1H), 4.03 – 3.93 (m, 4H), 3.88 (d, *J* = 10.9 Hz, 1H), 3.79 (dd, *J* = 11.0, 3.7 Hz, 1H), 3.52 – 3.31 (m, 4H), 3.13 – 2.92 (m, 2H), 2.58 (t, *J* = 2.5 Hz, 1H), 2.52 – 2.28 (m, 4H), 2.24 (dd, *J* = 13.3, 7.6 Hz, 1H), 2.09 (ddt, *J* = 13.3, 9.3, 4.3 Hz, 1H), 1.04 (s, 9H). **$^{13}$C NMR** (101 MHz, MeOD) δ = 174.3, 172.0, 171.8, 171.8, 156.7, 153.9, 148.6, 145.3, 145.2, 142.7, 141.1, 139.0, 130.7, 130.4, 129.2, 128.8, 128.2, 126.0, 121.0, 80.5, 72.2, 71.1, 70.5, 68.3, 67.5, 61.8, 60.8, 58.3, 58.1, 53.9, 45.0, 43.7, 38.9, 37.0, 29.0, 27.0. **HRMS** (ESI+) req. for $C_{48}H_{56}N_7O_8S$ $[M+H]^+$ 890.3906, found 890.4007.

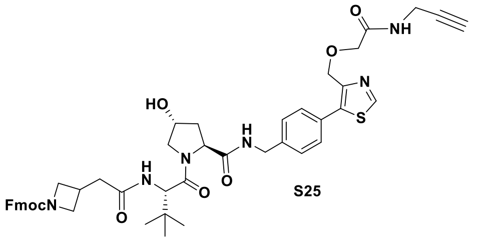

**S25**

**$^{1}$H NMR** (400 MHz, MeOD) δ = 8.96 (s, 1H), 7.77 (d, *J* = 7.6 Hz, 2H), 7.57 (d, *J* = 7.4 Hz, 2H), 7.50 – 7.41 (m, 4H), 7.37 (t, *J* = 7.4 Hz, 2H), 7.29 (t, *J* = 7.4 Hz, 2H), 4.65 – 4.53 (m, 5H), 4.50 (t, *J* = 2.2 Hz, 1H), 4.37 – 4.30 (m, 3H), 4.16 (t, *J* = 6.6 Hz, 1H), 4.07 – 3.95 (m, 6H), 3.89 (d, *J* = 11.2 Hz, 1H), 3.79 (dd, *J* = 11.1, 3.9 Hz, 1H), 3.64 (dt, *J* = 8.7, 4.6 Hz, 2H), 2.90 (dqd, *J* = 13.6, 7.8, 5.4 Hz, 1H), 2.67 – 2.48 (m, 3H), 2.22 (tdd, *J* = 12.4, 5.4, 3.5 Hz, 1H), 2.08 (ddd, *J* = 13.3, 9.2, 4.5 Hz, 1H), 1.04 (s, 9H). **$^{13}$C NMR** (101 MHz, MeOD) δ = 174.5, 173.4, 172.2, 171.8, 158.2, 153.9, 148.6, 145.2, 142.6, 141.1, 139.0, 130.6, 130.4, 129.1, 128.8, 128.2, 126.1, 121.0, 80.5, 72.2, 71.1, 70.5, 68.1, 67.5, 60.8, 59.0, 58.0, 43.6, 40.2, 38.9, 36.6, 29.0, 27.6, 27.1. **HRMS** (ESI+) req. for $C_{47}H_{53}N_6O_8S$ $[M+H]^+$ 861.3640, found 861.3694.

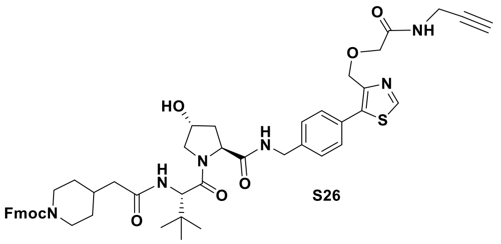

**S26**

**$^{1}$H NMR** (400 MHz, MeOD) δ = 8.97 (s, 1H), 7.85 (d, *J* = 9.0 Hz, 1H), 7.79 (d, *J* = 7.5 Hz, 2H), 7.58 (d, *J* = 7.5 Hz, 2H), 7.52 – 7.44 (m, 4H), 7.38 (t, *J* = 7.4 Hz, 2H), 7.31 (t, *J* = 7.5 Hz, 2H), 4.69 – 4.47 (m, 8H), 4.42 (s, 1H), 4.35 (d, *J* = 15.5 Hz, 1H), 4.21 (t, *J* = 6.2 Hz, 1H), 4.00 (d, *J* = 10.4 Hz, 5H), 3.92 (d, *J* = 11.1 Hz, 1H), 3.80 (dd, *J* = 10.8, 4.0 Hz, 2H), 2.72 (td, *J* = 13.0, 3.1 Hz, 2H), 2.58 (t, *J* = 2.9 Hz, 1H), 2.27 – 2.15 (m, 3H), 2.08 (ddd, *J* = 13.3, 9.1, 4.5 Hz, 1H), 1.89 (s, 1H), 1.59 (s, 2H), 1.05 (s, 9H). **$^{13}$C NMR** (101 MHz, MeOD) δ = 174.5, 174.4, 172.3, 171.8, 156.8, 154.0, 148.6, 145.3, 142.7, 141.2, 139.1, 130.7, 130.4, 129.1, 128.8, 128.2, 125.9, 125.9, 121.0, 80.5, 72.2, 71.1, 70.5, 68.2, 67.5, 60.8, 59.1, 58.0, 54.8, 45.1, 45.0, 43.7, 43.2, 38.9, 36.4, 34.7, 32.7, 29.0, 27.1. **HRMS** (ESI+) req. for $C_{49}H_{57}N_6O_8S$ $[M+H]^+$ 889.3953, found 889.3951.

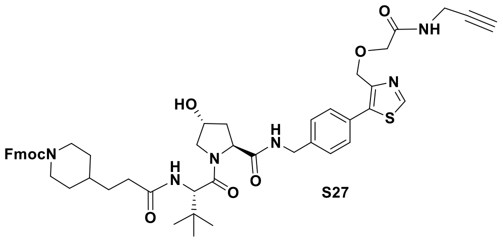

**S27**

**$^{1}$H NMR** (400 MHz, $CDCl_3$) δ = 8.91 (s, 1H), 7.74 (d, *J* = 7.3 Hz, 2H), 7.55 (d, *J* = 7.6 Hz, 2H), 7.49 (t, *J* = 5.6 Hz, 1H), 7.41 – 7.32 (m, 7H), 7.29 (td, *J* = 7.6, 1.3 Hz, 2H), 6.37 (d, *J* = 8.8 Hz, 1H), 4.71 – 4.64 (m, 3H), 4.59 (dd, *J* = 15.3, 6.6 Hz, 1H), 4.56 – 4.51 (m, 2H), 4.41 – 4.28 (m, 3H), 4.21 (t, *J* = 6.9 Hz, 1H), 4.06 (p, *J* = 2.9 Hz, 6H), 3.65 (dd, *J* = 11.2, 3.7 Hz, 1H), 2.73 (d, *J* = 14.6 Hz, 2H), 2.44 (ddd, *J* = 12.9, 8.1, 4.6 Hz, 1H), 2.37 – 2.19 (m, 3H), 2.19 – 2.10 (m, 1H), 1.69 – 1.52 (m, 4H), 1.40 (dqd, *J* = 14.6, 7.0, 4.8 Hz, 1H), 0.96 (s, 9H). **$^{13}$C NMR** (101 MHz, $CDCl_3$) δ = 173.6, 171.9, 171.2, 169.3, 155.3, 152.3, 146.4, 144.2, 141.4, 139.8, 137.8, 129.7, 128.8, 128.4, 127.8, 127.1, 125.1, 120.1, 116.5, 79.5, 77.4, 71.5, 70.1, 67.3, 66.5, 58.8, 57.6, 56.9, 47.5, 44.2, 43.2, 36.4, 35.6, 35.1, 33.5, 32.0, 31.8, 28.6, 26.6. **HRMS** (ESI+) req. for $C_{50}H_{59}N_6O_8S$ $[M+H]^+$ 903.4110, found 903.4135.

#### 5. Synthesis of modified VHL ligand – connector conjugates for VHL TR-FRET binding assay.

**Scheme S4.** Conversion of VHL ligands with Fmoc-protected connector amines to acetylated amines.

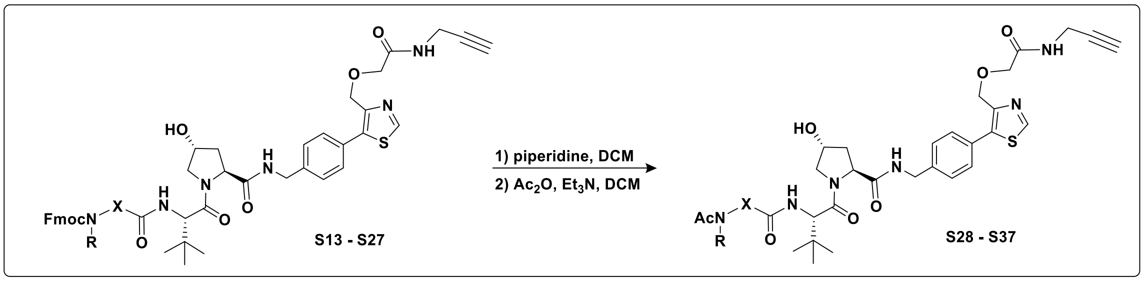


***Synthesis of S28 – S37:*** Conversion of Fmoc-protected connector amines to acetylated amines was performed according to **GP4** to generate the following compounds:

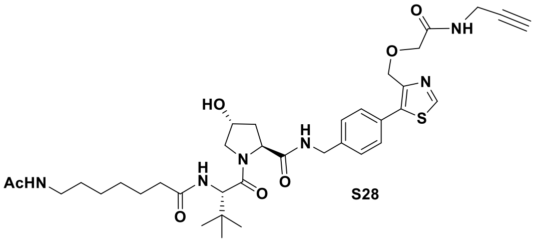

S28

**$^1$H NMR** (400 MHz, MeOD) δ = 9.00 (s, 1H), 7.48 (d, *J* = 1.7 Hz, 4H), 4.68 (s, 2H), 4.63 (s, 1H), 4.58 (t, *J* = 4.4 Hz, 1H), 4.54 (d, *J* = 2.8 Hz, 1H), 4.50 (s, 1H), 4.36 (d, *J* = 15.5 Hz, 1H), 4.03 (s, 2H), 4.00 (d, *J* = 2.5 Hz, 2H), 3.91 (d, *J* = 11.1 Hz, 1H), 3.80 (dd, *J* = 10.8, 3.9 Hz, 1H), 3.14 (t, *J* = 7.0 Hz, 2H), 2.59 (t, *J* = 2.6 Hz, 1H), 2.36 – 2.17 (m, 3H), 2.08 (ddd, *J* = 13.3, 9.2, 4.5 Hz, 1H), 1.91 (s, 3H), 1.67 – 1.56 (m, 2H), 1.49 (p, *J* = 6.7 Hz, 2H), 1.34 (p, *J* = 3.7 Hz, 4H), 1.04 (s, 9H). **$^{13}$C NMR** (101 MHz, MeOD) δ = 176.0, 174.5, 173.2, 172.3, 171.9, 154.0, 148.6, 141.2, 139.1, 130.7, 130.4, 129.1, 80.5, 72.2, 71.1, 70.5, 67.5, 60.8, 59.0, 58.0, 43.7, 40.4, 38.9, 36.5, 36.5, 30.2, 29.9, 29.0, 27.7, 27.0, 27.0, 26.9, 22.5. **HRMS** (ESI+) req. for $C_{36}H_{51}N_6O_7S$ $[M+H]^+$ 711.3534, found 711.3589.

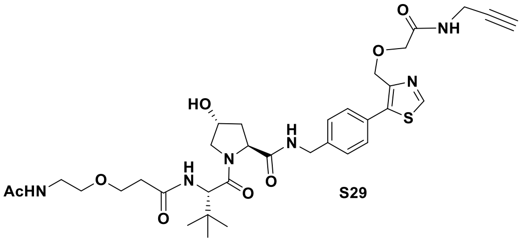

S29

**$^1$H NMR** (400 MHz, MeOD) δ = 9.00 (s, 1H), 7.48 (s, 4H), 4.70 – 4.65 (m, 3H), 4.61 – 4.48 (m, 3H), 4.37 (d, *J* = 15.6 Hz, 1H), 4.03 (s, 2H), 4.00 (d, *J* = 2.4 Hz, 2H), 3.90 (d, *J* = 10.9 Hz, 1H), 3.81 (dd, *J* = 10.9, 3.9 Hz, 1H), 3.77 – 3.66 (m, 2H), 3.55 – 3.49 (m, 2H), 3.35 (td, *J* = 5.3, 2.4 Hz, 2H), 2.59 (t, *J* = 2.5 Hz, 1H), 2.53 (ddd, *J* = 9.1, 7.2, 4.1 Hz, 2H), 2.23 (ddt, *J* = 13.2, 7.8, 1.9 Hz, 1H), 2.08 (ddd, *J* = 13.4, 9.2, 4.4 Hz, 1H), 1.92 (s, 3H), 1.05 (s, 9H). **$^{13}$C NMR** (101 MHz, MeOD) δ = 174.4, 173.8, 173.5, 172.2, 171.9, 154.0, 148.7, 141.2, 139.0, 130.7, 130.5, 129.2, 80.5, 72.2, 71.1, 70.5, 70.5, 68.0, 67.5, 60.8, 58.9, 58.1, 43.7, 40.5, 39.0, 37.2, 36.9, 29.0, 27.0, 27.0, 22.7. **HRMS** (ESI+) req. for $C_{34}H_{47}N_6O_8S$ $[M+H]^+$ 699.3171, found 699.3213.

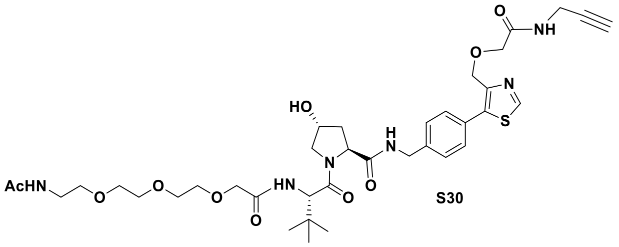

S30

**$^1$H NMR** (400 MHz, MeOD) δ = 9.00 (s, 1H), 7.49 (s, 4H), 4.69 (d, *J* = 7.8 Hz, 3H), 4.56 (dd, *J* = 16.1, 6.4 Hz, 2H), 4.50 (s, 1H), 4.37 (d, *J* = 15.6 Hz, 1H), 4.06 (d, *J* = 3.7 Hz, 2H), 4.03 (s, 2H), 4.00 (d, *J* = 2.4 Hz, 2H), 3.88 (d, *J* = 11.2 Hz, 1H), 3.81 (dd, *J* = 11.0, 3.7 Hz, 1H), 3.74 – 3.68 (m, 4H), 3.68 – 3.63 (m, 2H), 3.63 – 3.59 (m, 2H), 3.50 (t, *J* = 5.4 Hz, 2H), 3.36 – 3.31 (m, 2H), 2.59 (t, *J* = 2.5 Hz, 1H), 2.24 (ddt, *J* = 13.2, 7.4, 1.9 Hz, 1H), 2.09 (ddd, *J* = 13.2, 9.2, 4.4 Hz, 1H), 1.92 (s, 3H), 1.05 (s, 9H). **$^{13}$C NMR** (101 MHz, MeOD) δ = 174.4, 173.4, 172.1, 171.9, 171.7, 154.0, 148.6, 141.1, 139.1, 130.7, 130.4, 129.1, 80.5, 72.2, 72.2, 71.7, 71.5, 71.3, 71.1, 71.1, 70.5, 70.5, 67.5, 60.8, 58.1, 43.7, 40.5, 39.0, 37.1, 29.0, 27.0, 22.6. **HRMS** (ESI+) req. for $C_{37}H_{53}N_6O_{10}S$ $[M+H]^+$ 773.3538, found 773.3561.

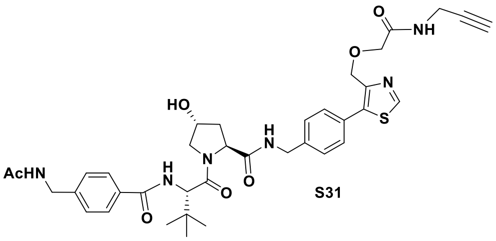

S31

**$^1$H NMR** (400 MHz, MeOD) δ = 8.99 (s, 1H), 7.81 (d, *J* = 8.0 Hz, 2H), 7.49 (s, 4H), 7.39 (d, *J* = 8.0 Hz, 2H), 4.68 (s, 2H), 4.64 – 4.56 (m, 2H), 4.53 (s, 1H), 4.41 (s, 2H), 4.35 (d, *J* = 15.6 Hz, 1H), 4.04 (s, 2H), 4.02 – 3.94 (m, 3H), 3.87 (dd, *J* = 10.9, 3.9 Hz, 1H), 2.58 (t, *J* = 2.5 Hz, 1H), 2.24 (dd, *J* = 13.1, 7.7 Hz, 1H), 2.12 (dddd, *J* = 13.3, 9.3, 8.7, 3.7 Hz, 1H), 2.00 (s, 3H), 1.11 (s, 9H). **$^{13}$C NMR** (101 MHz, MeOD) δ = 174.4, 173.3, 172.3, 171.9, 169.6, 154.0, 148.6, 144.3, 141.2, 139.1, 134.0, 130.7, 130.4, 129.1, 128.7, 128.6, 80.5, 72.2, 71.1, 70.5, 67.5, 60.9, 59.4, 58.2, 43.8, 43.7, 39.0, 37.3, 29.0, 27.1, 22.5. **HRMS** (ESI+) req. for $C_{37}H_{45}N_6O_7S$ $[M+H]^+$ 717.3065, found 717.3137.

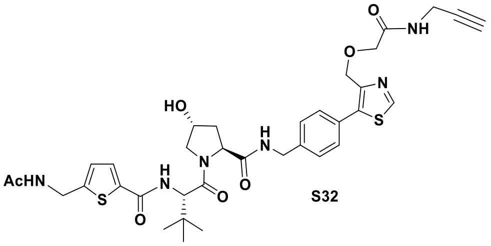

S32

**$^1$H NMR** (400 MHz, MeOD) δ = 9.00 (s, 1H), 7.65 (d, *J* = 3.6 Hz, 1H), 7.49 (s, 4H), 7.00 (d, *J* = 3.8 Hz, 1H), 4.68 (s, 2H), 4.59 (dd, *J* = 15.4, 7.8 Hz, 2H), 4.52 (s, 3H), 4.34 (d, *J* = 15.6 Hz, 1H), 4.04 (s, 2H), 4.00 (d, *J* = 2.5 Hz, 2H), 3.94 (d, *J* = 11.2 Hz, 1H), 3.84 (dd, *J* = 11.1, 3.8 Hz, 1H), 2.58 (t, *J* = 2.6 Hz, 1H), 2.24 (ddt, *J* = 13.3, 7.6, 1.6 Hz, 1H), 2.10 (ddd, *J* = 13.2, 9.4, 4.3 Hz, 1H), 1.97 (s, 3H), 1.08 (s, 9H). **$^{13}$C NMR** (101 MHz, MeOD) δ = 174.4, 173.1, 172.2, 171.9, 163.9, 154.0, 149.5, 148.6, 141.2, 139.1, 138.3, 130.7, 130.4, 130.1, 129.1, 127.4, 80.5, 72.2, 71.1, 70.5, 67.5, 60.9, 59.2, 58.2, 43.7, 39.1, 38.9, 37.3, 29.0, 27.1, 22.4. **HRMS** (ESI+) req. for $C_{35}H_{43}N_6O_7S$ $[M+H]^+$ 723.2629, found 723.2713.

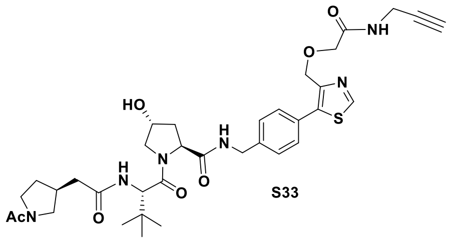

S33

**$^1$H NMR** (400 MHz, MeOD) δ = 9.00 (s, 1H), 7.95 (dd, *J* = 18.9, 8.9 Hz, 1H), 7.48 (d, *J* = 2.7 Hz, 5H), 4.67 (s, 2H), 4.64 (dd, *J* = 8.8, 2.5 Hz, 1H), 4.56 (dt, *J* = 14.0, 3.4 Hz, 2H), 4.50 (t, *J* = 2.2 Hz, 1H), 4.36 (d, *J* = 15.6 Hz, 1H), 4.03 (s, 2H), 4.00 (d, *J* = 2.6 Hz, 2H), 3.95 – 3.87 (m, 1H), 3.80 (dd, *J* = 10.8, 3.9 Hz, 1H), 3.67 (dd, *J* = 10.6, 6.8 Hz, 1H), 3.64 – 3.43 (m, 2H), 3.21 – 2.95 (m, 1H), 2.69 – 2.51 (m, 2H), 2.49 – 2.33 (m, 2H), 2.22 (ddd, *J* = 13.2, 7.4, 2.0 Hz, 1H), 2.18 – 2.04 (m, 2H), 2.03 (s, 3H), 1.78 – 1.56 (m, 1H), 1.05 (d, *J* = 2.5 Hz, 9H). **$^{13}$C NMR** (101 MHz, MeOD) δ = 174.5, 174.1, 174.1, 172.3, 172.2, 172.0, 171.9, 154.0, 148.6, 141.2, 139.1, 130.7, 130.4, 129.1, 80.5, 72.2, 71.1, 70.5, 67.5, 60.8, 59.1, 59.0, 58.0, 53.5, 51.8, 48.0, 46.1, 43.7, 39.5, 39.3, 39.0, 38.9, 37.7, 36.5, 36.4, 36.2, 32.6, 31.3, 29.0, 27.0, 22.2, 21.9. **HRMS** (ESI+) req. for $C_{35}H_{47}N_6O_7S$ $[M+H]^+$ 695.3221, found 695.3259.

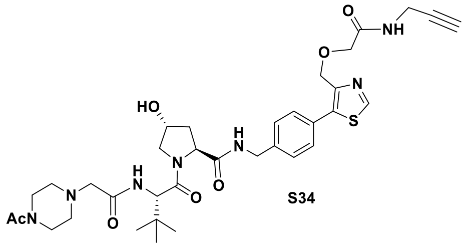

S34

**$^1$H NMR** (400 MHz, MeOD) δ = 8.99 (s, 1H), 7.49 (s, 4H), 4.68 (s, 2H), 4.65 (s, 1H), 4.58 (d, *J* = 7.3 Hz, 1H), 4.54 (d, *J* = 5.9 Hz, 1H), 4.50 (d, *J* = 3.3 Hz, 1H), 4.38 (d, *J* = 15.5 Hz, 1H), 4.03 (s, 2H), 3.99 (d, *J* = 2.7 Hz, 2H), 3.89 (dt, *J* = 11.2, 1.9 Hz, 1H), 3.80 (dd, *J* = 11.0, 3.7 Hz, 1H), 3.67 – 3.56 (m, 4H), 3.11 (d, *J* = 7.4 Hz, 2H), 2.63 – 2.55 (m, 3H), 2.52 (t, *J* = 5.2 Hz, 2H), 2.24 (ddt, *J* = 13.2, 7.9, 1.9 Hz, 1H), 2.09 (s, 3H), 1.05 (s, 9H). **$^{13}$C NMR** (101 MHz, MeOD) δ = 174.4, 172.0, 171.9, 171.8, 171.7, 154.0, 148.6, 141.1, 139.1, 130.7, 130.5, 129.6, 129.2, 80.5, 72.2, 71.1, 70.5, 67.5, 61.7, 60.8, 58.4, 58.2, 54.4, 54.0, 47.1, 43.7, 42.8, 39.0, 37.0, 29.0, 27.0, 27.0, 21.1. **HRMS** (ESI+) req. for $C_{35}H_{48}N_7O_7S$ $[M+H]^+$ 710.3330, found 710.3374.

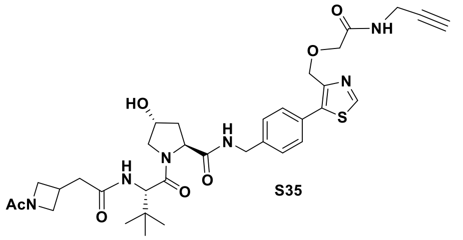

S35

**$^1$H NMR** (400 MHz, MeOD) δ = 9.00 (s, 1H), 7.48 (d, *J* = 2.4 Hz, 4H), 4.68 (s, 2H), 4.64 – 4.53 (m, 3H), 4.49 (s, 1H), 4.39 – 4.28 (m, 2H), 4.13 – 4.05 (m, 1H), 4.03 (s, 2H), 4.00 (d, *J* = 2.5 Hz, 2H), 3.96 – 3.85 (m, 2H), 3.79 (dd, *J* = 10.9, 3.9 Hz, 1H), 3.68 (dd, *J* = 10.2, 5.6 Hz, 1H), 2.96 (p, *J* = 5.8 Hz, 1H), 2.67 – 2.57 (m, 3H), 2.22 (ddt, *J* = 13.2, 7.7, 2.0 Hz, 1H), 2.08 (ddd, *J* = 13.3, 9.1, 4.5 Hz, 1H), 1.83 (s, 3H), 1.04 (s, 9H). **$^{13}$C NMR** (101 MHz, MeOD) δ = 174.5, 173.4, 173.2, 172.2, 172.2, 171.9, 154.0, 148.6, 141.2, 139.1, 130.7, 130.4, 129.1, 80.5, 72.2, 71.1, 70.5, 67.5, 60.8, 59.1, 58.0, 56.8, 56.7, 54.1, 54.0, 43.7, 40.1,

39.0, 36.5, 29.0, 27.0, 26.8, 26.7, 18.4. **HRMS** (ESI+) req. for $C_{34}H_{45}N_6O_7S$ $[M+H]^+$ 681.3065, found 681.3093.

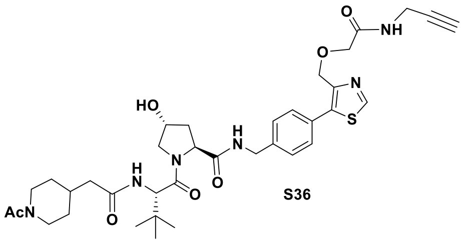

S36

**$^{1}$H NMR** (400 MHz, MeOD) δ = 9.00 (s, 1H), 7.48 (d, *J* = 1.9 Hz, 5H), 4.68 (s, 2H), 4.64 (d, *J* = 2.1 Hz, 1H), 4.60 – 4.53 (m, 2H), 4.48 (ddd, *J* = 14.7, 4.0, 2.1 Hz, 2H), 4.36 (dd, *J* = 15.6, 1.9 Hz, 1H), 4.03 (s, 2H), 4.00 (d, *J* = 2.5 Hz, 2H), 3.94 – 3.85 (m, 2H), 3.80 (dd, *J* = 10.9, 3.9 Hz, 1H), 3.16 – 3.04 (m, 1H), 2.67 – 2.56 (m, 2H), 2.28 – 2.18 (m, 3H), 2.13 – 1.95 (m, 6H), 1.81 – 1.67 (m, 2H), 1.30 – 1.07 (m, 3H), 1.04 (s, 9H). **$^{13}$C NMR** (101 MHz, MeOD) δ = 174.5, 174.5, 172.3, 172.3, 171.9, 171.4, 154.0, 148.6, 141.2, 139.1, 130.7, 130.4, 129.1, 80.5, 72.2, 71.1, 70.5, 67.5, 60.8, 59.1, 59.1, 58.0, 47.7, 43.7, 43.1, 42.9, 39.0, 36.4, 36.4, 34.9, 33.4, 33.3, 32.7, 32.6, 29.0, 27.1, 21.2. **HRMS** (ESI+) req. for $C_{36}H_{49}N_6O_7S$ $[M+H]^+$ 709.3378, found 709.3448.

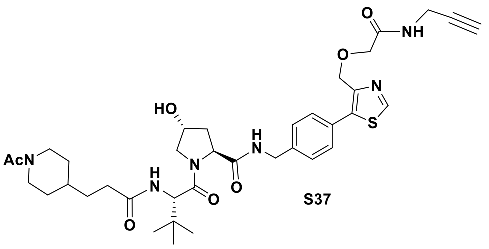

S37

**$^{1}$H NMR** (400 MHz, MeOD) δ = 9.00 (s, 1H), 7.48 (d, *J* = 2.4 Hz, 4H), 4.68 (s, 2H), 4.65 (d, *J* = 2.8 Hz, 1H), 4.59 – 4.53 (m, 2H), 4.48 (dt, *J* = 13.1, 2.1 Hz, 2H), 4.36 (d, *J* = 15.5 Hz, 1H), 4.03 (s, 2H), 4.00 (d, *J* = 2.5 Hz, 2H), 3.90 (tt, *J* = 9.3, 2.0 Hz, 2H), 3.80 (dd, *J* = 11.0, 3.8 Hz, 1H), 3.06 (tt, *J* = 13.2, 3.1 Hz, 1H), 2.63 – 2.52 (m, 2H), 2.34 (td, *J* = 7.3, 4.1 Hz, 2H), 2.22 (ddt, *J* = 13.2, 7.9, 2.0 Hz, 1H), 2.12 – 2.04 (m, 4H), 1.85 – 1.70 (m, 2H), 1.61 – 1.48 (m, 3H), 1.21 – 1.06 (m, 2H), 1.04 (s, 9H). **$^{13}$C NMR** (101 MHz, MeOD) δ = 175.9, 174.5, 172.3, 171.9, 171.3, 154.0, 148.6, 141.2, 139.1, 130.7, 130.4, 129.1, 80.5, 72.2, 71.1, 70.5, 67.5, 60.8, 59.0, 59.0, 58.1, 47.9, 43.7, 43.1, 43.0, 39.0, 36.7, 36.7, 36.5, 33.8, 33.8, 33.5, 33.3, 32.8, 32.8, 29.0, 27.0, 21.2. **HRMS** (ESI+) req. for $C_{37}H_{51}N_6O_7S$ $[M+H]^+$ 723.3534, found 723.3588.

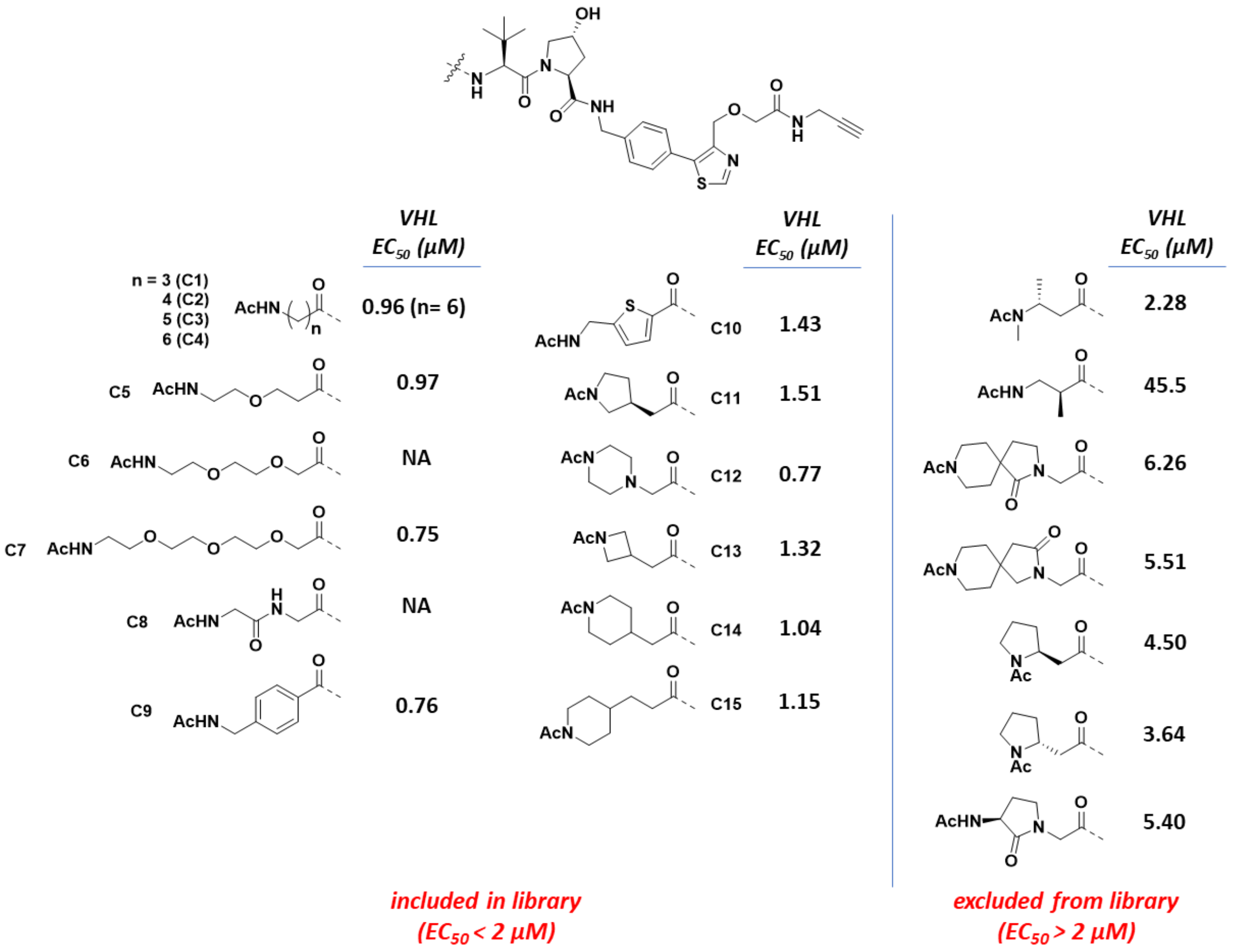


**Figure S2. Assessment of VHL ligand – connector conjugate binding to VHL for library inclusion.** Acetylated connectors conjugated to the modified VHL ligand were tested for binding to VHL using a TR-FRET peptide displacement assay (mean; n = 2). Connector conjugates that displayed $EC_{50}$ values less than 2 µM were included in the CIP-DEL library. NA = not available; compounds not tested.

#### 6. Synthesis of CIP-DEL screening hits and controls for validation experiments.

**Scheme S5.** Off-DNA synthesis of CIP-DEL screening hits.

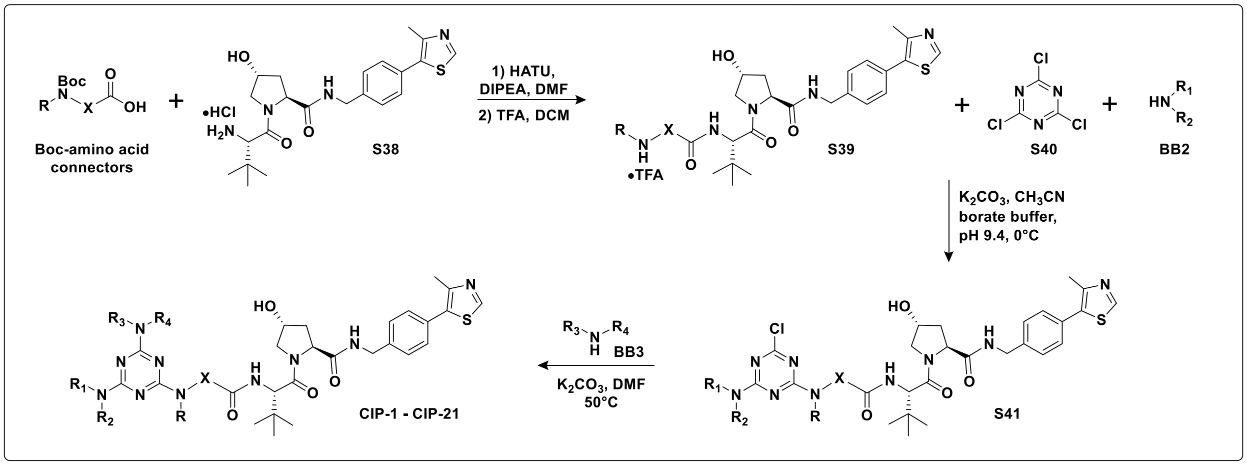


***Synthesis of CIP-1 – CIP-21:*** Connectors (1.2 equiv) were coupled to the VHL ligand amine **S38** (1.0 equiv) following **GP1** and products were purified by silica gel chromatography ($CH_2Cl_2$ – MeOH gradient). The coupled products were deprotected following **GP3** to provide **S39** which were taken on crude. Cyanuric chloride **S40** and **BB2** were then appended to the connector amines according to **GP5** to provide **S41**. The final triazine chloride was substituted with **BB3** (4 equiv) according to **GP6** to provide products **CIP-1 – CIP-S21** and the final products were purified by reverse-phase HPLC using gradients of water – acetonitrile both modified with 0.1% formic acid to generate the following compounds:

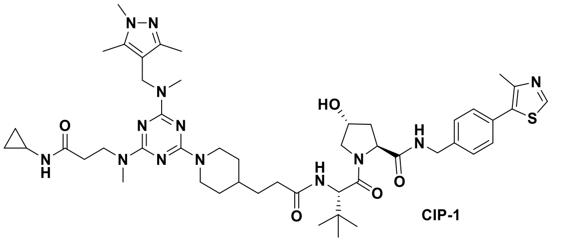

CIP-1

**$^1$H NMR** (400 MHz, MeOD) δ = 8.86 (s, 1H), 7.87 (d, *J* = 9.0 Hz, 1H), 7.49 – 7.38 (m, 4H), 4.77 (d, *J* = 12.8 Hz, 2H), 4.69 – 4.60 (m, 3H), 4.60 – 4.47 (m, 3H), 4.35 (d, *J* = 15.4 Hz, 1H), 3.91 (d, *J* = 11.0 Hz, 1H), 3.88 – 3.76 (m, 3H), 3.67 (s, 3H), 3.07 (s, 3H), 2.86 (s, 3H), 2.75 (t, *J* = 12.7 Hz, 2H), 2.59 (hept, *J* = 3.7 Hz, 1H), 2.47 (s, 3H), 2.44 (t, *J* = 6.6 Hz, 2H), 2.33 (hept, *J* = 6.9 Hz, 2H), 2.21 (s, 4H), 2.12 (s, 3H), 2.07 (td, *J* = 9.0, 4.5 Hz, 1H), 1.74 (d, *J* = 12.6 Hz, 2H), 1.62 – 1.45 (m, 3H), 1.15 – 1.07 (m, 2H), 1.04 (s, 9H), 0.65 (d, *J* = 5.3 Hz, 2H), 0.42 – 0.31 (m, 2H). **$^{13}$C NMR** (101 MHz, MeOD) δ = 176.0, 176.0, 174.5, 172.3, 166.8, 166.1, 152.8, 149.0, 147.6, 140.3, 140.1, 133.4, 131.5, 130.4, 129.0, 114.2, 71.1, 60.8, 59.0, 58.0, 46.2, 44.6, 43.7, 40.9, 38.9, 37.3, 36.5, 35.7, 35.5, 34.8, 33.9, 33.7, 33.2, 33.2, 32.4, 27.0, 23.3, 15.8, 12.0, 9.6, 6.4. **HRMS** (ESI+) req. for $C_{48}H_{70}N_{13}O_5S$ $[M+H]^+$ 940.5338, found 940.5406.

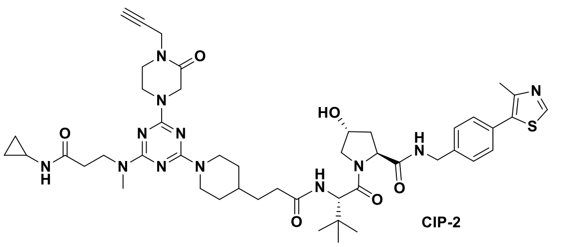

CIP-2

**$^{1}$H NMR** (400 MHz, DMSO) δ = 8.98 (s, 1H), 8.56 (t, *J* = 6.1 Hz, 1H), 7.90 (t, *J* = 4.5 Hz, 2H), 7.40 (q, *J* = 8.3 Hz, 4H), 5.12 (d, *J* = 3.5 Hz, 1H), 4.61 (d, *J* = 12.7 Hz, 2H), 4.55 (d, *J* = 9.4 Hz, 1H), 4.48 – 4.38 (m, 2H), 4.35 (s, 1H), 4.26 – 4.16 (m, 5H), 3.93 (t, *J* = 5.4 Hz, 2H), 3.73 – 3.60 (m, 4H), 3.41 (t, *J* = 5.1 Hz, 2H), 3.23 (t, *J* = 2.4 Hz, 1H), 2.98 (s, 3H), 2.75 – 2.63 (m, 2H), 2.57 (ddd, *J* = 12.0, 7.8, 4.1 Hz, 1H), 2.44 (s, 3H), 2.27 (t, *J* = 6.6 Hz, 3H), 2.21 – 2.12 (m, 1H), 2.08 – 1.98 (m, 1H), 1.90 (ddd, *J* = 12.9, 8.6, 4.6 Hz, 1H), 1.68 (t, *J* = 12.4 Hz, 2H), 1.51 – 1.35 (m, 3H), 0.94 (s, 11H), 0.56 (td, *J* = 7.1, 4.8 Hz, 2H), 0.36 – 0.28 (m, 2H). **$^{13}$C NMR** (101 MHz, DMSO) δ = 172.2, 172.0, 171.8, 169.7, 165.2, 164.7, 151.5, 147.7, 139.5, 131.2, 129.6, 128.7, 127.4, 79.0, 74.5, 68.9, 58.7, 56.3, 47.0, 45.0, 42.9, 41.7, 38.0, 35.3, 35.2, 34.6, 34.2, 32.2, 31.4, 26.4, 22.2, 16.0, 5.6. **HRMS** (ESI+) req. for $C_{47}H_{65}N_{12}O_6S$ $[M+H]^+$ 925.4865, found 925.4897.

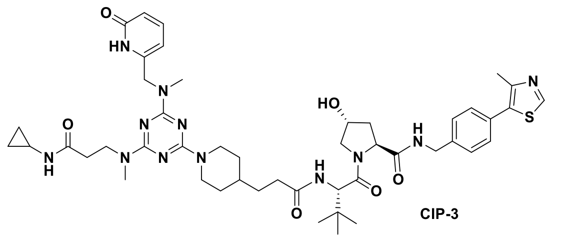

CIP-3

**$^{1}$H NMR** (400 MHz, MeOD) δ = 8.87 (s, 1H), 8.55 (s, 1H), 7.52 (dd, *J* = 9.2, 6.8 Hz, 1H), 7.49 – 7.37 (m, 4H), 6.40 (d, *J* = 9.2 Hz, 1H), 6.28 (d, *J* = 6.8 Hz, 1H), 4.74 (d, *J* = 11.3 Hz, 2H), 4.65 (s, 1H), 4.62 – 4.46 (m, 5H), 4.36 (d, *J* = 15.5 Hz, 1H), 3.91 (d, *J* = 10.9 Hz, 1H), 3.87 – 3.76 (m, 3H), 3.11 (d, *J* = 19.5 Hz, 5H), 2.77 (tt, *J* = 13.4, 3.5 Hz, 2H), 2.60 (tq, *J* = 7.4, 3.7 Hz, 1H), 2.47 (s, 3H), 2.42 (t, *J* = 6.6 Hz, 2H), 2.34 (td, *J* = 7.2, 5.4 Hz, 2H), 2.21 (dd, *J* = 13.3, 7.7 Hz, 1H), 2.07 (ddd, *J* = 13.3, 9.2, 4.5 Hz, 1H), 1.78 – 1.68 (m, 2H), 1.61 – 1.46 (m, 3H), 1.18 – 1.07 (m, 2H), 1.04 (s, 9H), 0.65 (td, *J* = 7.0, 5.1 Hz, 2H), 0.41 – 0.33 (m, 2H). **$^{13}$C NMR** (101 MHz, MeOD) δ = 176.0, 176.0, 175.0, 174.5, 172.3, 166.0, 152.8, 149.0, 148.3, 143.7, 140.3, 133.4, 131.5, 130.4, 129.0, 119.1, 106.7, 71.1, 60.8, 59.0, 58.1, 50.4, 46.5, 44.6, 43.7, 38.9, 37.2, 36.5, 35.5, 35.1, 35.0, 33.9, 33.7, 33.1, 27.0, 23.3, 15.8, 6.4. **HRMS** (ESI+) req. for $C_{47}H_{65}N_{12}O_6S$ $[M+H]^+$ 925.4865, found 925.4930.

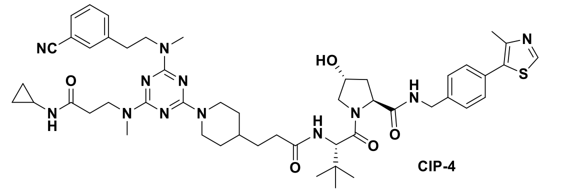

CIP-4

**$^{1}$H NMR** (400 MHz, MeOD) δ = 8.87 (s, 1H), 8.55 (s, 1H), 7.59 (t, *J* = 1.8 Hz, 1H), 7.53 (dd, *J* = 8.1, 1.5 Hz, 2H), 7.50 – 7.38 (m, 4H), 4.72 (d, *J* = 13.2 Hz, 2H), 4.65 (s, 1H), 4.62 – 4.46 (m, 3H), 4.35 (d, *J* = 15.5 Hz, 1H), 3.91 (d, *J* = 11.0 Hz, 1H), 3.85 – 3.76 (m, 4H), 3.09 – 3.00 (m, 5H), 2.95 (t, *J* = 7.2 Hz, 2H), 2.73 (tt, *J* = 12.8, 3.4 Hz, 2H), 2.60 (hept, *J* = 3.7 Hz, 1H), 2.47 (s, 2H), 2.43 (t, *J* = 6.7 Hz, 2H), 2.35 (h, *J* = 6.7 Hz, 2H), 2.22 (ddt, *J* = 13.1, 7.7, 1.8 Hz, 1H), 2.08 (ddd, *J* = 13.3, 9.1, 4.5 Hz, 1H), 1.74 (d, *J* = 12.7 Hz, 2H), 1.57 (q, *J* = 6.8 Hz, 2H), 1.04 (s, 10H), 0.66 (td, *J* = 8.8, 7.2 Hz, 2H), 0.42 – 0.34 (m, 2H). **$^{13}$C NMR** (101 MHz, MeOD) δ = 176.1, 174.5, 172.3, 166.8, 152.8, 150.4, 149.0, 143.1, 140.3, 135.0, 133.7, 131.5, 130.9, 130.4, 130.4, 129.0, 119.9, 113.2, 71.1, 60.8, 59.0, 58.0, 46.2, 44.6, 43.7, 38.9, 37.3, 36.6, 35.5, 34.8, 34.4,

33.9, 33.7, 33.2, 33.1, 27.0, 23.3, 15.8, 6.4. **HRMS** (ESI+) req. for $C_{50}H_{67}N_{12}O_5S$ $[M+H]^+$ 947.5073, found 947.5146.

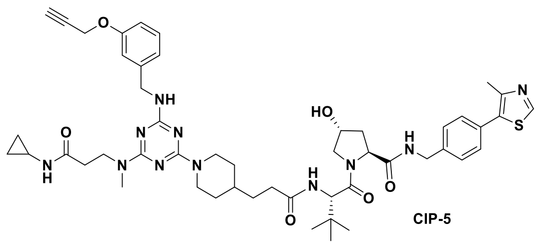

CIP-5

**$^1$H NMR** (400 MHz, DMSO) δ = 8.98 (s, 1H), 8.56 (t, *J* = 6.1 Hz, 1H), 7.89 (d, *J* = 9.3 Hz, 2H), 7.40 (q, *J* = 8.3 Hz, 4H), 7.20 (t, *J* = 8.0 Hz, 2H), 6.95 – 6.87 (m, 2H), 6.82 (dd, *J* = 8.0, 2.4 Hz, 1H), 5.12 (d, *J* = 3.5 Hz, 1H), 4.74 (d, *J* = 2.4 Hz, 2H), 4.64 – 4.51 (m, 3H), 4.43 (q, *J* = 7.4 Hz, 2H), 4.38 – 4.31 (m, 3H), 4.21 (dd, *J* = 15.9, 5.4 Hz, 1H), 3.72 – 3.58 (m, 4H), 3.52 (t, *J* = 2.4 Hz, 1H), 2.94 (s, 3H), 2.70 – 2.53 (m, 3H), 2.44 (s, 3H), 2.34 – 2.21 (m, 3H), 2.19 – 2.11 (m, 1H), 2.07 – 1.98 (m, 1H), 1.90 (ddd, *J* = 12.9, 8.7, 4.6 Hz, 1H), 1.64 (t, *J* = 12.7 Hz, 2H), 1.48 – 1.33 (m, 3H), 0.94 (s, 11H), 0.57 (td, *J* = 7.0, 4.8 Hz, 2H), 0.37 – 0.29 (m, 2H). **$^{13}$C NMR** (101 MHz, DMSO) δ = 172.7, 172.4, 172.4, 170.2, 157.6, 151.9, 148.2, 143.2, 140.0, 131.6, 130.1, 129.5, 129.1, 127.9, 120.9, 114.8, 112.8, 79.8, 78.5, 69.3, 59.2, 56.8, 55.7, 43.9, 43.2, 42.1, 38.4, 35.8, 35.7, 34.3, 32.7, 32.1, 31.9, 26.9, 22.7, 16.4, 6.0. **HRMS** (ESI+) req. for $C_{50}H_{66}N_{11}O_6S$ $[M+H]^+$ 948.4913, found 948.4943.

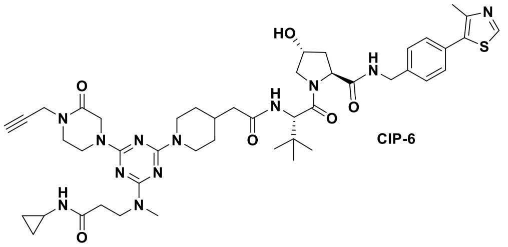

CIP-6

**$^1$H NMR** (400 MHz, DMSO) δ = 8.98 (s, 1H), 8.56 (t, *J* = 6.1 Hz, 1H), 7.94 – 7.83 (m, 2H), 7.40 (q, *J* = 8.4 Hz, 4H), 5.12 (d, *J* = 3.6 Hz, 1H), 4.62 – 4.51 (m, 3H), 4.49 – 4.39 (m, 2H), 4.35 (s, 1H), 4.27 – 4.17 (m, 5H), 3.93 (t, *J* = 5.4 Hz, 2H), 3.74 – 3.61 (m, 4H), 3.41 (t, *J* = 5.5 Hz, 2H), 3.23 (t, *J* = 2.5 Hz, 1H), 2.98 (s, 3H), 2.76 (t, *J* = 12.4 Hz, 2H), 2.57 (td, *J* = 7.3, 3.7 Hz, 1H), 2.44 (s, 3H), 2.27 (t, *J* = 7.2 Hz, 2H), 2.24 – 2.17 (m, 1H), 2.13 – 1.98 (m, 2H), 1.90 (ddd, *J* = 12.9, 8.5, 4.5 Hz, 2H), 1.63 (d, *J* = 12.5 Hz, 2H), 1.11 – 0.98 (m, 2H), 0.94 (s, 9H), 0.57 (td, *J* = 7.0, 4.7 Hz, 2H), 0.32 (dt, *J* = 6.7, 4.4 Hz, 2H). **$^{13}$C NMR** (101 MHz, DMSO) δ = 172.0, 171.8, 171.0, 169.7, 165.2, 164.7, 151.5, 147.7, 139.5, 131.2, 129.6, 128.7, 127.4, 79.0, 74.5, 68.9, 58.7, 56.4, 47.1, 45.0, 42.7, 41.7, 38.0, 35.1, 34.6, 34.2, 33.4, 31.5, 26.4, 22.2, 16.0, 5.6. **HRMS** (ESI+) req. for $C_{46}H_{63}N_{12}O_6S$ $[M+H]^+$ 911.4709, found 911.4735.

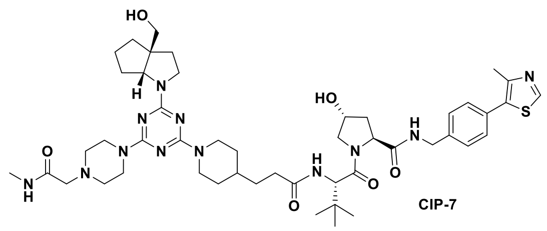

CIP-7

**$^1$H NMR** (400 MHz, DMSO) δ = 8.98 (s, 1H), 8.56 (t, *J* = 6.1 Hz, 1H), 7.89 (d, *J* = 9.3 Hz, 1H), 7.71 (q, *J* = 4.2 Hz, 1H), 7.40 (q, *J* = 8.1 Hz, 4H), 5.12 (d, *J* = 3.5 Hz, 1H), 4.79 (t, *J* = 5.2 Hz, 1H), 4.60 (d, *J* = 12.7 Hz,

2H), 4.54 (d, *J* = 9.4 Hz, 1H), 4.47 – 4.38 (m, 2H), 4.35 (s, 1H), 4.21 (dd, *J* = 15.8, 5.4 Hz, 1H), 3.96 (dd, *J* = 6.5, 3.2 Hz, 1H), 3.71 – 3.60 (m, 7H), 3.45 – 3.34 (m, 1H), 2.91 (s, 2H), 2.66 (s, 2H), 2.62 (d, *J* = 4.7 Hz, 3H), 2.44 (s, 3H), 2.39 (s, 4H), 2.34 – 2.26 (m, 1H), 2.21 – 2.11 (m, 1H), 2.03 (t, *J* = 10.4 Hz, 1H), 1.95 – 1.79 (m, 3H), 1.75 (s, 1H), 1.60 (ddd, *J* = 25.0, 13.0, 6.7 Hz, 7H), 1.51 – 1.31 (m, 5H), 0.94 (s, 11H). **$^{13}$C NMR** (101 MHz, DMSO) δ = 172.2, 172.0, 169.7, 163.1, 151.5, 147.7, 139.5, 131.2, 129.6, 128.7, 127.4, 68.9, 66.2, 64.4, 58.7, 56.4, 56.3, 55.1, 45.8, 42.8, 41.7, 38.0, 35.4, 35.2, 35.0, 33.4, 33.0, 32.2, 31.5, 26.4, 25.5, 24.5, 16.0. **HRMS** (ESI+) req. for $C_{48}H_{71}N_{12}O_6S$ $[M+H]^+$ 943.5335, found 943.5369.

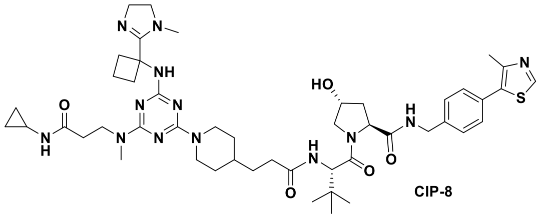

CIP-8

**$^{1}$H NMR** (400 MHz, DMSO) δ = 9.37 (s, 1H), 8.98 (s, 1H), 8.60 (d, *J* = 30.4 Hz, 1H), 7.89 (s, 1H), 7.48 (s, 1H), 7.40 (q, *J* = 8.3 Hz, 4H), 5.12 (s, 1H), 4.64 – 4.52 (m, 2H), 4.48 – 4.38 (m, 2H), 4.35 (s, 1H), 4.21 (dd, *J* = 15.9, 5.5 Hz, 1H), 3.70 – 3.61 (m, 2H), 3.51 – 3.38 (m, 3H), 3.25 – 3.19 (m, 1H), 3.18 – 3.08 (m, 1H), 2.94 (s, 3H), 2.91 – 2.82 (m, 1H), 2.72 (s, 1H), 2.66 (s, 3H), 2.63 – 2.55 (m, 3H), 2.54 (s, 2H), 2.44 (s, 3H), 2.34 – 2.13 (m, 6H), 2.06 – 1.99 (m, 1H), 1.90 (ddd, *J* = 12.9, 8.5, 4.5 Hz, 1H), 1.71 – 1.54 (m, 3H), 1.52 – 1.35 (m, 4H), 0.94 (s, 9H), 0.87 – 0.81 (m, 2H), 0.65 – 0.54 (m, 2H), 0.44 (s, 1H), 0.33 (s, 1H). **$^{13}$C NMR** (101 MHz, DMSO) δ = 172.2, 172.0, 172.0, 169.7, 164.4, 164.2, 151.5, 147.7, 139.5, 131.2, 129.6, 128.7, 127.4, 68.9, 58.7, 56.4, 56.3, 54.6, 42.7, 41.7, 40.4, 38.0, 37.0, 35.4, 35.2, 34.5, 33.8, 32.2, 31.9, 31.7, 31.5, 31.3, 29.0, 28.7, 26.4, 22.4, 16.0, 5.6, 5.3. **HRMS** (ESI+) req. for $C_{48}H_{70}N_{13}O_5S$ $[M+H]^+$ 940.5338, found 940.5388.

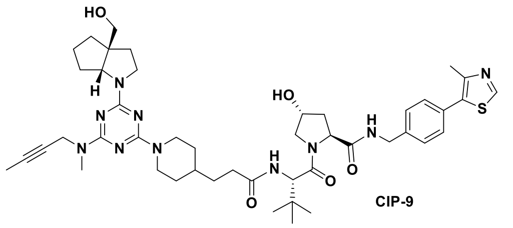

CIP-9

**$^{1}$H NMR** (400 MHz, DMSO) δ = 8.98 (s, 1H), 8.56 (t, *J* = 6.0 Hz, 1H), 7.89 (d, *J* = 9.4 Hz, 1H), 7.40 (q, *J* = 8.3 Hz, 4H), 5.12 (d, *J* = 3.5 Hz, 1H), 4.80 (t, *J* = 5.3 Hz, 1H), 4.62 (d, *J* = 12.8 Hz, 2H), 4.54 (d, *J* = 9.3 Hz, 1H), 4.43 (q, *J* = 7.3 Hz, 2H), 4.33 (d, *J* = 13.4 Hz, 3H), 4.21 (dd, *J* = 15.9, 5.4 Hz, 1H), 3.97 (s, 1H), 3.71 – 3.61 (m, 3H), 3.45 – 3.38 (m, 1H), 2.99 (s, 3H), 2.73 – 2.61 (m, 2H), 2.44 (s, 3H), 2.35 – 2.25 (m, 1H), 2.16 (dd, *J* = 13.9, 7.5 Hz, 1H), 2.07 – 1.99 (m, 1H), 1.95 – 1.80 (m, 3H), 1.76 (t, *J* = 2.4 Hz, 4H), 1.72 – 1.51 (m, 7H), 1.50 – 1.31 (m, 5H), 0.94 (s, 11H). **$^{13}$C NMR** (101 MHz, DMSO) δ = 172.2, 172.0, 169.7, 151.5, 147.7, 139.5, 131.2, 129.6, 128.7, 127.4, 78.5, 75.7, 68.9, 66.3, 64.4, 58.7, 56.4, 56.3, 55.1, 45.8, 42.8, 41.7, 38.0, 36.8, 35.4, 35.2, 35.0, 33.4, 33.3, 33.0, 32.8, 32.2, 31.7, 31.5, 26.4, 24.5, 16.0, 3.1. **HRMS** (ESI+) req. for $C_{46}H_{65}N_{10}O_5S$ $[M+H]^+$ 869.4855, found 869.4907.

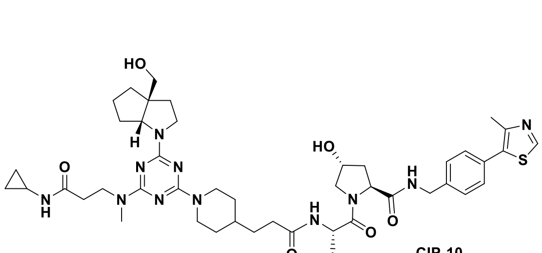

CIP-10

**$^{1}$H NMR** (400 MHz, DMSO) δ = 8.98 (s, 1H), 8.56 (t, *J* = 6.1 Hz, 1H), 7.93 – 7.85 (m, 2H), 7.40 (q, *J* = 8.2 Hz, 4H), 5.12 (d, *J* = 3.5 Hz, 1H), 4.79 (t, *J* = 4.8 Hz, 1H), 4.61 (d, *J* = 12.8 Hz, 2H), 4.54 (d, *J* = 9.4 Hz, 1H), 4.43 (q, *J* = 7.5 Hz, 2H), 4.38 – 4.32 (m, 1H), 4.21 (dd, *J* = 15.9, 5.5 Hz, 1H), 3.98 – 3.92 (m, 1H), 3.71 – 3.60 (m, 4H), 3.44 – 3.38 (m, 1H), 3.38 – 3.34 (m, 2H), 2.95 (s, 3H), 2.71 – 2.53 (m, 4H), 2.44 (s, 3H), 2.35 – 2.22 (m, 3H), 2.21 – 2.12 (m, 1H), 2.07 – 1.99 (m, 1H), 1.94 – 1.72 (m, 4H), 1.71 – 1.51 (m, 6H), 1.50 – 1.33 (m, 4H), 0.94 (s, 11H), 0.62 – 0.51 (m, 2H), 0.38 – 0.29 (m, 2H). **$^{13}$C NMR** (101 MHz, DMSO) δ = 214.6, 172.2, 172.0, 171.3, 169.7, 166.6, 164.2, 164.0, 151.5, 147.7, 139.5, 131.2, 129.7, 128.7, 127.4, 68.9, 66.3, 64.4, 58.7, 56.4, 56.3, 55.1, 45.8, 42.9, 41.7, 38.0, 35.4, 35.2, 35.0, 34.1, 33.8, 33.0, 32.2, 31.7, 31.5, 26.4, 24.5, 22.2, 16.0, 5.6. **HRMS** (ESI+) req. for $C_{48}H_{70}N_{11}O_6S$ $[M+H]^+$ 928.5226, found 928.5268.

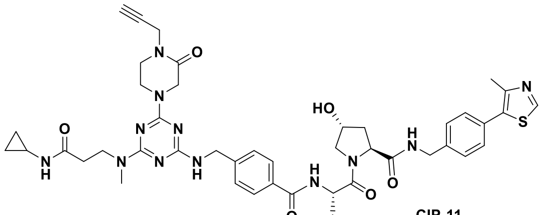

CIP-11

**$^{1}$H NMR** (400 MHz, DMSO) δ = 8.98 (s, 1H), 8.58 (t, *J* = 6.1 Hz, 1H), 7.95 – 7.82 (m, 2H), 7.79 (d, *J* = 7.9 Hz, 2H), 7.50 (t, *J* = 6.4 Hz, 1H), 7.44 – 7.34 (m, 6H), 5.14 (d, *J* = 3.6 Hz, 1H), 4.75 (d, *J* = 9.1 Hz, 1H), 4.49 – 4.34 (m, 5H), 4.28 – 4.15 (m, 5H), 3.95 – 3.86 (m, 2H), 3.72 (s, 2H), 3.69 – 3.59 (m, 2H), 3.43 – 3.38 (m, 1H), 3.24 – 3.18 (m, 1H), 3.02 – 2.91 (m, 3H), 2.68 – 2.53 (m, 2H), 2.44 (s, 3H), 2.30 – 2.19 (m, 2H), 2.04 (ddd, *J* = 12.2, 7.1, 2.1 Hz, 1H), 1.91 (ddd, *J* = 13.0, 8.7, 4.5 Hz, 1H), 1.02 (s, 9H), 0.60 – 0.52 (m, 2H), 0.37 – 0.27 (m, 2H). **$^{13}$C NMR** (101 MHz, DMSO) δ = 171.9, 171.8, 169.5, 166.3, 165.2, 151.5, 147.8, 144.4, 139.5, 134.4, 132.3, 131.2, 129.7, 128.7, 127.5, 127.5, 127.0, 79.0, 74.5, 68.9, 58.8, 57.2, 56.4, 47.0, 45.0, 41.7, 37.9, 35.6, 34.6, 33.7, 26.5, 22.2, 16.0, 5.6. **HRMS** (ESI+) req. for $C_{47}H_{59}N_{12}O_6S$ $[M+H]^+$ 919.4396, found 919.4417.

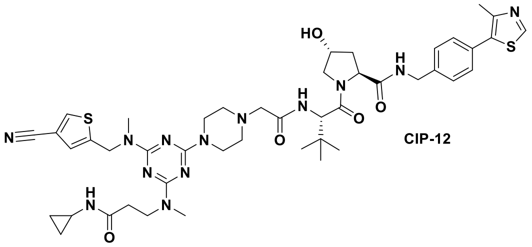

CIP-12

**$^{1}$H NMR** (400 MHz, DMSO) δ = 8.94 (s, 1H), 8.59 (t, *J* = 6.1 Hz, 1H), 8.38 (s, 1H), 7.90 (s, 1H), 7.82 (d, *J* = 9.6 Hz, 1H), 7.45 (s, 1H), 7.44 – 7.36 (m, 4H), 5.14 (d, *J* = 3.4 Hz, 1H), 4.83 (s, 2H), 4.52 (d, *J* = 9.6 Hz, 1H), 4.47 – 4.33 (m, 3H), 4.26 (dd, *J* = 15.5, 5.8 Hz, 1H), 3.81 – 3.58 (m, 8H), 3.13 – 2.94 (m, 9H), 2.59 – 2.52 (m, 2H), 2.49 – 2.44 (m, 2H), 2.42 (s, 3H), 2.28 (t, *J* = 5.8 Hz, 2H), 2.05 (dd, *J* = 12.5, 8.1 Hz, 1H), 1.90 (ddd, *J* = 13.1, 9.0, 4.5 Hz, 1H), 0.95 (s, 9H), 0.55 (d, *J* = 7.0 Hz, 2H), 0.33 – 0.26 (m, 2H). **$^{13}$C NMR** (101 MHz, DMSO) δ 171.8, 171.8, 169.3, 168.3, 164.7, 151.4, 147.7, 139.5, 137.8, 131.2, 129.7, 128.9, 128.7, 127.5, 127.5, 115.5, 68.9, 60.7, 58.8, 56.6, 55.9, 52.8, 52.8, 52.7, 45.7, 45.6, 44.9, 42.9, 42.9, 42.8, 41.7, 37.9, 35.8, 33.8, 33.3, 26.3, 22.2, 15.9, 5.6. **HRMS** (ESI+) req. for $C_{45}H_{60}N_{13}O_5S_2$ $[M+H]^+$ 926.4276, found 926.4310.

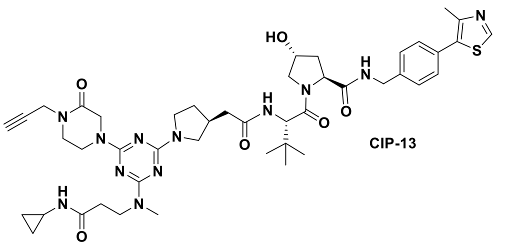

CIP-13

**[1]H NMR** (400 MHz, DMSO) δ = 8.98 (s, 1H), 8.56 (t, *J* = 6.1 Hz, 1H), 7.98 (d, *J* = 9.2 Hz, 1H), 7.90 (s, 1H), 7.40 (q, *J* = 8.3 Hz, 4H), 5.13 (d, *J* = 3.6 Hz, 1H), 4.57 (d, *J* = 9.3 Hz, 1H), 4.44 (dd, *J* = 15.0, 6.8 Hz, 2H), 4.35 (s, 1H), 4.27 – 4.17 (m, 5H), 3.93 (s, 2H), 3.73 – 3.62 (m, 5H), 3.57 (td, *J* = 7.9, 3.8 Hz, 1H), 3.41 (t, *J* = 5.6 Hz, 2H), 3.23 (t, *J* = 2.5 Hz, 1H), 3.04 (dd, *J* = 10.2, 7.4 Hz, 1H), 2.98 (s, 3H), 2.58 (td, *J* = 7.4, 3.8 Hz, 1H), 2.46 – 2.39 (m, 5H), 2.31 – 2.22 (m, 3H), 2.10 – 1.85 (m, 4H), 1.60 – 1.49 (m, 1H), 0.95 (s, 9H), 0.57 (td, *J* = 7.0, 4.7 Hz, 2H), 0.37 – 0.28 (m, 2H). **[13]C NMR** (101 MHz, DMSO) δ = 172.0, 171.9, 171.0, 169.6, 165.9, 164.4, 151.5, 147.7, 139.5, 131.2, 129.7, 128.7, 127.4, 79.0, 74.5, 68.9, 58.7, 56.4, 47.0, 45.0, 44.9, 44.8, 41.7, 38.4, 38.0, 35.3, 34.6, 34.2, 33.8, 30.4, 26.4, 22.2, 16.0, 5.6. **HRMS** (ESI+) req. for $C_{45}H_{61}N_{12}O_6S$ $[M+H]^+$ 897.4552, found 897.4587.

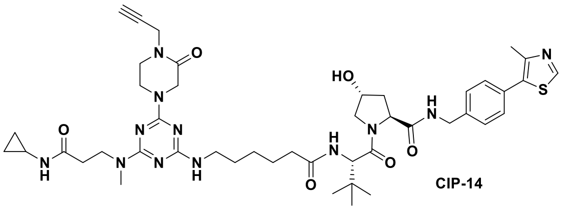

CIP-14

**[1]H NMR** (400 MHz, DMSO) δ = 8.98 (s, 1H), 8.55 (t, *J* = 6.0 Hz, 1H), 7.91 (s, 1H), 7.83 (d, *J* = 9.3 Hz, 1H), 7.40 (q, *J* = 8.3 Hz, 4H), 6.78 (d, *J* = 6.5 Hz, 1H), 5.12 (d, *J* = 3.5 Hz, 1H), 4.53 (d, *J* = 9.4 Hz, 1H), 4.43 (dd, *J* = 15.8, 7.4 Hz, 2H), 4.35 (s, 1H), 4.26 – 4.17 (m, 5H), 3.96 – 3.87 (m, 2H), 3.71 – 3.61 (m, 4H), 3.40 (s, 2H), 3.22 (t, *J* = 2.5 Hz, 1H), 3.19 (d, *J* = 6.7 Hz, 2H), 2.96 (d, *J* = 10.2 Hz, 3H), 2.57 (ddd, *J* = 9.7, 6.5, 2.7 Hz, 1H), 2.44 (s, 3H), 2.30 – 2.21 (m, 3H), 2.11 (q, *J* = 7.0 Hz, 1H), 2.07 – 1.99 (m, 1H), 1.90 (ddd, *J* = 12.8, 8.5, 4.5 Hz, 1H), 1.47 (dq, *J* = 14.4, 8.2, 7.1 Hz, 4H), 1.29 – 1.21 (m, 2H), 0.92 (s, 9H), 0.57 (d, *J* = 6.6 Hz, 2H), 0.33 (s, 2H). **[13]C NMR** (101 MHz, DMSO) δ = 172.1, 172.0, 171.9, 169.7, 165.3, 151.5, 147.7, 139.5, 131.2, 129.6, 128.7, 127.4, 79.0, 74.5, 68.9, 58.7, 56.4, 56.3, 47.0, 45.0, 44.8, 41.7, 38.0, 35.2, 35.0, 34.6, 33.8, 29.1, 26.4, 26.2, 25.3, 22.2, 16.0, 5.6. **HRMS** (ESI+) req. for $C_{45}H_{63}N_{12}O_6S$ $[M+H]^+$ 899.4709, found 899.4736.

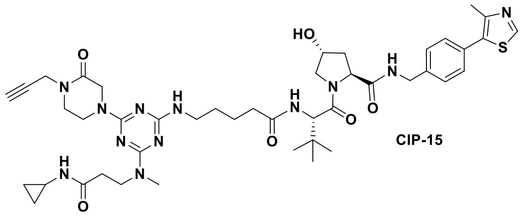

CIP-15

**[1]H NMR** (400 MHz, DMSO) δ = 8.98 (s, 1H), 8.56 (t, *J* = 6.0 Hz, 1H), 7.91 (s, 1H), 7.83 (d, *J* = 9.3 Hz, 1H), 7.40 (q, *J* = 8.2 Hz, 4H), 6.81 (d, *J* = 5.7 Hz, 1H), 5.12 (d, *J* = 3.6 Hz, 1H), 4.53 (d, *J* = 9.4 Hz, 1H), 4.48 – 4.39 (m, 2H), 4.35 (s, 1H), 4.26 – 4.17 (m, 5H), 3.97 – 3.87 (m, 2H), 3.72 – 3.60 (m, 4H), 3.41 (s, 2H), 3.24 – 3.17 (m, 3H), 2.97 (d, *J* = 13.7 Hz, 3H), 2.58 (s, 1H), 2.44 (s, 3H), 2.31 – 2.22 (m, 3H), 2.11 (dd, *J* = 14.3, 7.1 Hz, 1H), 2.03 (dd, *J* = 11.9, 7.7 Hz, 1H), 1.90 (ddd, *J* = 12.8, 8.5, 4.6 Hz, 1H), 1.54 – 1.42 (m, 4H), 0.93 (s, 9H), 0.60 – 0.54 (m, 2H), 0.33 (s, 2H). **[13]C NMR** (101 MHz, DMSO) δ = 172.1, 172.0, 171.9, 169.7, 165.3, 151.5, 147.7, 139.5, 131.2, 129.7, 128.7, 127.4, 79.0, 74.5, 68.9, 58.7, 56.4, 56.3, 47.0, 45.0, 44.9, 41.7, 38.0, 35.2, 34.8, 34.6, 33.8, 29.0, 26.4, 23.1, 22.2, 16.0, 5.6. **HRMS** (ESI+) req. for $C_{44}H_{61}N_{12}O_6S$ $[M+H]^+$ 885.4552, found 885.4582.

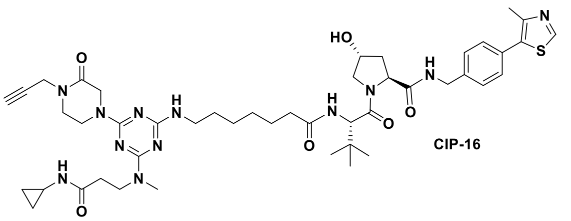

CIP-16

**$^{1}$H NMR** (400 MHz, DMSO) δ = 8.98 (s, 1H), 8.55 (t, *J* = 6.1 Hz, 1H), 7.90 (s, 1H), 7.83 (d, *J* = 9.3 Hz, 1H), 7.40 (dd, *J* = 8.3, 7.0 Hz, 4H), 6.78 (s, 1H), 5.12 (d, *J* = 3.6 Hz, 1H), 4.54 (d, *J* = 9.3 Hz, 1H), 4.47 – 4.38 (m, 2H), 4.34 (s, 1H), 4.26 – 4.17 (m, 5H), 3.91 (dd, *J* = 10.8, 5.6 Hz, 2H), 3.72 – 3.60 (m, 4H), 3.40 (s, 2H), 3.22 (t, *J* = 2.5 Hz, 1H), 3.18 (t, *J* = 6.7 Hz, 2H), 2.97 (d, *J* = 11.2 Hz, 3H), 2.62 – 2.53 (m, 1H), 2.44 (s, 3H), 2.31 – 2.21 (m, 3H), 2.14 – 1.99 (m, 2H), 1.90 (ddd, *J* = 13.0, 8.6, 4.6 Hz, 1H), 1.52 – 1.39 (m, 4H), 1.30 – 1.19 (m, 4H), 0.93 (s, 9H), 0.60 – 0.52 (m, 2H), 0.33 (s, 2H). **$^{13}$C NMR** (101 MHz, DMSO) δ = 172.1, 172.0, 171.9, 169.7, 165.3, 151.5, 147.7, 139.5, 131.2, 129.6, 128.7, 127.4, 79.0, 74.5, 68.9, 58.7, 56.4, 56.3, 47.0, 45.0, 44.8, 41.7, 38.0, 35.2, 34.9, 34.6, 33.8, 29.1, 28.5, 26.4, 26.3, 25.5, 22.2, 16.0, 5.6. **HRMS** (ESI+) req. for $C_{46}H_{65}N_{12}O_6S$ $[M+H]^+$ 913.4865, found 913.4912.

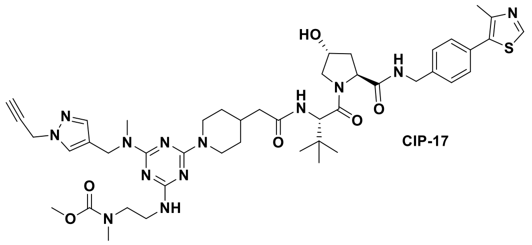

CIP-17

**$^{1}$H NMR** (400 MHz, DMSO) δ = 8.98 (s, 1H), 8.56 (t, *J* = 6.0 Hz, 1H), 7.87 (d, *J* = 9.3 Hz, 1H), 7.67 (d, *J* = 4.3 Hz, 1H), 7.47 – 7.32 (m, 6H), 6.73 (s, 1H), 5.11 (d, *J* = 3.5 Hz, 1H), 4.96 (s, 2H), 4.67 – 4.47 (m, 6H), 4.47 – 4.39 (m, 2H), 4.35 (s, 1H), 4.22 (dd, *J* = 15.9, 5.5 Hz, 1H), 3.66 (s, 2H), 3.58 – 3.41 (m, 5H), 3.01 – 2.90 (m, 3H), 2.86 – 2.66 (m, 6H), 2.44 (s, 3H), 2.27 – 2.17 (m, 1H), 2.12 – 1.99 (m, 2H), 1.90 (tt, *J* = 8.3, 4.6 Hz, 2H), 1.62 (s, 2H), 1.12 – 0.99 (m, 2H), 0.94 (s, 9H). **$^{13}$C NMR** (101 MHz, DMSO) δ = 172.0, 171.0, 169.7, 156.1, 151.5, 147.7, 139.5, 139.1, 131.2, 129.7, 128.7, 127.4, 118.3, 78.6, 76.2, 68.9, 58.7, 56.4, 52.2, 42.6, 41.8, 41.7, 40.7, 38.0, 35.1, 33.5, 33.2, 31.5, 31.4, 26.4, 16.0. **HRMS** (ESI+) req. for $C_{45}H_{62}N_{13}O_6S$ $[M+H]^+$ 912.4661, found 912.4724.

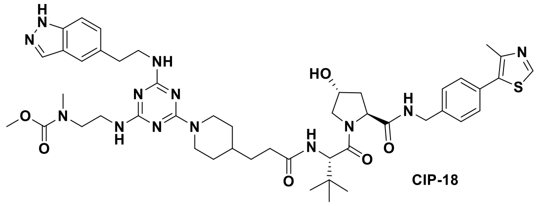

CIP-18

**$^{1}$H NMR** (400 MHz, DMSO) δ = 12.93 (s, 1H), 8.98 (s, 1H), 8.56 (t, *J* = 6.1 Hz, 1H), 7.97 (s, 1H), 7.89 (d, *J* = 9.3 Hz, 1H), 7.54 (s, 1H), 7.47 – 7.35 (m, 5H), 7.21 (d, *J* = 8.6 Hz, 1H), 6.70 (s, 1H), 6.53 (s, 1H), 5.12 (d, *J* = 3.5 Hz, 1H), 4.71 – 4.52 (m, 3H), 4.43 (q, *J* = 7.3 Hz, 2H), 4.35 (s, 1H), 4.21 (dd, *J* = 15.9, 5.5 Hz, 1H), 3.70 – 3.62 (m, 2H), 3.58 – 3.49 (m, 3H), 3.46 – 3.38 (m, 2H), 3.37 – 3.33 (m, 2H), 3.31 – 3.28 (m, 3H), 2.95 – 2.75 (m, 6H), 2.44 (s, 3H), 2.35 – 2.25 (m, 1H), 2.22 – 2.12 (m, 1H), 2.03 (t, *J* = 10.3 Hz, 1H), 1.90 (ddd, *J* = 12.9, 8.6, 4.5 Hz, 1H), 1.73 – 1.57 (m, 2H), 1.50 – 1.35 (m, 3H), 0.93 (s, 11H). **$^{13}$C NMR** (101 MHz, DMSO) δ = 172.2, 172.0, 169.7, 156.1, 151.5, 147.7, 139.5, 138.8, 133.0, 131.6, 131.2, 129.7, 128.7, 127.4, 123.1, 119.4, 109.9, 68.9, 58.7, 56.4, 56.3, 52.2, 52.1, 41.7, 38.0, 35.4, 35.2, 32.2, 31.7, 31.5, 26.4, 16.0. **HRMS** (ESI+) req. for $C_{47}H_{64}N_{13}O_6S$ $[M+H]^+$ 938.4818, found 938.4854.

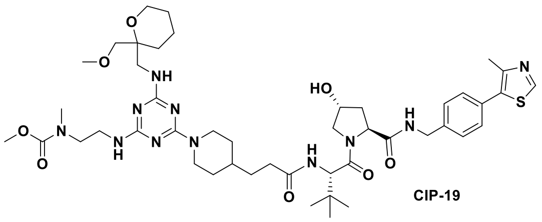

CIP-19

**$^{1}$H NMR** (400 MHz, DMSO) δ = 8.98 (s, 1H), 8.56 (t, *J* = 6.0 Hz, 1H), 7.89 (d, *J* = 9.3 Hz, 1H), 7.40 (q, *J* = 8.2 Hz, 4H), 6.70 (s, 1H), 6.51 (s, 1H), 6.00 – 5.78 (m, 1H), 5.12 (d, *J* = 3.5 Hz, 1H), 4.61 (s, 2H), 4.54 (d, *J* = 9.3 Hz, 1H), 4.43 (q, *J* = 7.8 Hz, 2H), 4.35 (s, 1H), 4.21 (dd, *J* = 15.9, 5.5 Hz, 1H), 3.70 – 3.63 (m, 2H), 3.63 – 3.50 (m, 6H), 3.31 – 3.28 (m, 4H), 3.24 (s, 3H), 2.83 (s, 3H), 2.71 – 2.57 (m, 2H), 2.44 (s, 3H), 2.35 – 2.25 (m, 1H), 2.21 – 2.11 (m, 1H), 2.07 – 1.98 (m, 1H), 1.90 (ddd, *J* = 12.9, 8.6, 4.6 Hz, 1H), 1.73 – 1.51 (m, 5H), 1.51 – 1.33 (m, 8H), 0.93 (s, 11H). **$^{13}$C NMR** (101 MHz, DMSO) δ = 172.2, 172.0, 169.7, 156.1, 151.5, 147.7, 139.5, 131.2, 129.6, 128.7, 127.4, 68.9, 61.0, 58.8, 58.7, 56.4, 56.3, 52.2, 41.7, 38.0, 35.4, 35.2, 32.2, 31.7, 31.5, 27.9, 26.4, 25.2, 18.5, 16.0. **HRMS** (ESI+) req. for $C_{46}H_{70}N_{11}O_8S$ $[M+H]^+$ 936.5124, found 936.5161.

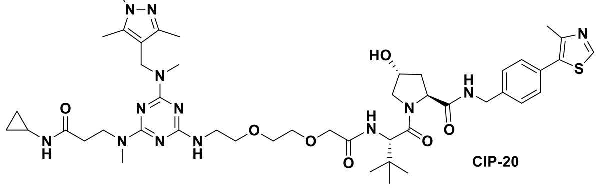

CIP-20

**$^{1}$H NMR** (400 MHz, MeOD) δ = 9.01 (s, 1H), 8.86 (t, *J* = 5.0 Hz, 1H), 8.34 (s, 1H), 7.90 – 7.76 (m, 1H), 7.45 (t, *J* = 4.9 Hz, 1H), 7.35 (t, *J* = 5.0 Hz, 4H), 4.80 – 4.73 (m, 1H), 4.72 – 4.56 (m, 3H), 4.55 – 4.43 (m, 2H), 4.22 (d, *J* = 15.2 Hz, 1H), 4.08 (dt, *J* = 16.0, 4.5 Hz, 1H), 3.96 (dt, *J* = 16.3, 4.4 Hz, 1H), 3.89 – 3.77 (m, 3H), 3.75 – 3.55 (m, 10H), 3.36 (s, 1H), 3.05 (s, 3H), 2.81 (s, 3H), 2.65 – 2.54 (m, 1H), 2.51 – 2.37 (m, 5H), 2.24 – 2.15 (m, 4H), 2.15 – 2.02 (m, 4H), 1.29 (s, 1H), 1.04 (s, 9H), 0.65 (q, *J* = 5.6 Hz, 2H), 0.39 (q, *J* = 5.3 Hz, 2H). **$^{13}$C NMR** (101 MHz, MeOD) δ = 175.9, 174.2, 172.3, 171.5, 152.8, 149.0, 147.6, 140.1, 139.8, 133.4, 131.5, 130.3, 129.3, 113.9, 72.7, 71.7, 71.2, 71.0, 70.7, 61.0, 58.1, 58.0, 44.0, 41.7, 38.8, 37.6, 35.8, 35.5, 27.0, 23.3, 16.0, 12.0, 9.7, 6.4, 6.4. **HRMS** (ESI+) req. for $C_{46}H_{68}N_{13}O_7S$ $[M+H]^+$ 946.5080, found 946.5140.

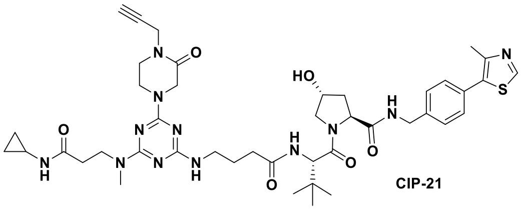

CIP-21

**$^{1}$H NMR** (400 MHz, DMSO) δ = 8.98 (s, 1H), 8.56 (t, *J* = 6.0 Hz, 1H), 7.90 (d, *J* = 9.0 Hz, 2H), 7.40 (dd, *J* = 8.4, 7.0 Hz, 4H), 6.82 (s, 1H), 5.12 (d, *J* = 3.6 Hz, 1H), 4.54 (d, *J* = 9.3 Hz, 1H), 4.47 – 4.39 (m, 2H), 4.35 (s, 1H), 4.27 – 4.21 (m, 3H), 4.21 – 4.17 (m, 3H), 3.97 – 3.89 (m, 2H), 3.73 – 3.63 (m, 4H), 3.44 – 3.38 (m, 2H), 3.22 (t, *J* = 2.6 Hz, 1H), 3.19 – 3.13 (m, 1H), 3.03 (s, 1H), 2.99 (s, 2H), 2.95 (s, 1H), 2.61 – 2.55 (m, 1H), 2.44 (s, 3H), 2.32 – 2.24 (m, 3H), 2.14 (dt, *J* = 14.6, 7.4 Hz, 1H), 2.03 (dd, *J* = 11.8, 8.4 Hz, 1H), 1.90 (ddd, *J* = 12.9, 8.6, 4.6 Hz, 1H), 1.69 (dq, *J* = 14.5, 7.1 Hz, 2H), 0.94 (s, 9H), 0.57 (td, *J* = 7.0, 4.7 Hz, 2H), 0.36 – 0.30 (m, 2H). **$^{13}$C NMR** (101 MHz, DMSO) δ = 214.5, 212.8, 205.4, 202.7, 172.0, 171.8, 171.8, 169.6, 165.3, 151.5, 147.7, 139.5, 131.2, 129.6, 128.7, 127.4, 79.0, 74.5, 68.9, 58.7, 56.4, 56.4, 47.0, 45.0, 45.0, 41.7, 38.0, 35.5, 35.3, 34.6, 34.6, 32.8, 26.4, 22.2, 16.0, 5.6. **HRMS** (ESI+) req. for $C_{43}H_{59}N_{12}O_6S$ $[M+H]^+$ 871.4396, found 871.4407.

**Scheme S6.** Synthesis of ***cis*-CIP-1** negative control.

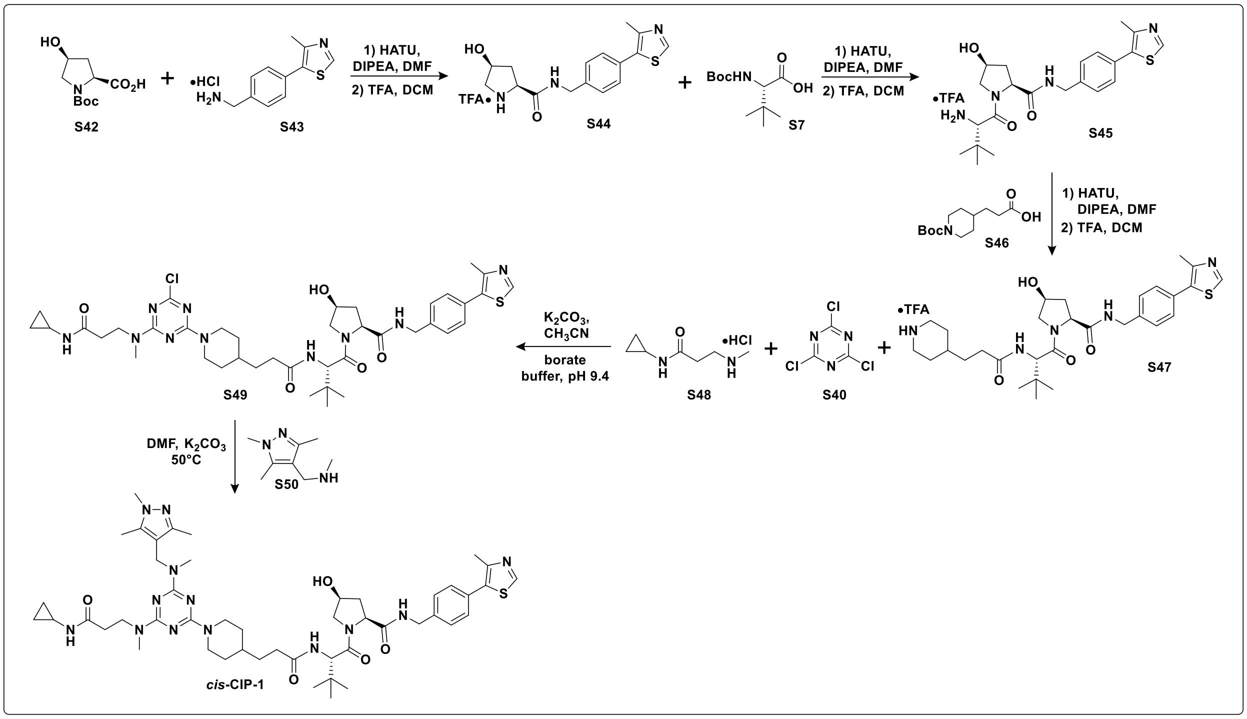


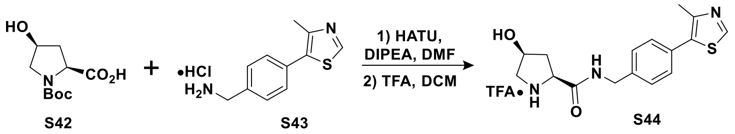


***Synthesis of S44:*** *N*-Boc-*cis*-hydroxyproline **S42** (300 mg, 1.30 mmol, 1.2 equiv) was coupled to amine **S43** (260 mg, 1.08 mmol, 1.0 equiv) according to **GP1** and purified by flash chromatography on silica ($CH_2Cl_2$ – MeOH gradient) to provide coupled product containing residual DIPEA. The coupled product was deprotected according to **GP3** to provide product **S44** mixed with DIPEA·TFA (204.8 mg, 0.475 mmol, ~37% yield over two-steps).

**$^1$H NMR** (400 MHz, MeOD) δ = 9.34 (s, 1H), 7.47 (s, 4H), 5.07 (s, 1H), 4.61 – 4.48 (m, 3H), 4.48 – 4.43 (m, 1H), 3.47 – 3.34 (m, 2H), 2.63 (ddt, *J* = 14.6, 10.2, 4.7 Hz, 1H), 2.51 (s, 3H), 2.21 (ddt, *J* = 14.2, 3.9, 1.9 Hz, 1H). **$^{13}$C NMR** (101 MHz, MeOD) δ = 170.0, 154.6, 146.1, 140.7, 135.0, 130.6, 130.5, 130.2, 129.3, 70.4, 60.0, 54.6, 44.2, 39.4, 14.5. **HRMS** (ESI+) req. for $C_{16}H_{20}N_3O_2S$ $[M+H]^+$ 318.1271, found 318.1288.

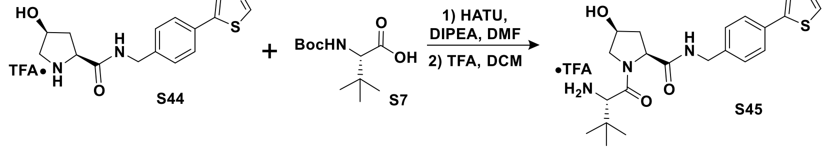


***Synthesis of S45:*** Boc-Tle-OH **S7** (113 mg, 0.487 mmol, 1.1 equiv) was coupled to amine **S44** (141 mg, 0.443 mmol, 1.0 equiv) according to **GP1** and purified by flash chromatography on silica ($CH_2Cl_2$ – MeOH gradient) to provide coupled product containing DIPEA. The coupled product was deprotected according to **GP3** to provide product **S45** mixed with DIPEA·TFA (241 mg, 0.443 mmol, ~quantitative yield over two-steps).

**$^1$H NMR** (400 MHz, MeOD) δ = 9.52 (s, 1H), 7.51 (d, *J* = 8.2 Hz, 2H), 7.46 (d, *J* = 8.4 Hz, 2H), 4.61 (dd, *J* = 9.0, 5.0 Hz, 1H), 4.57 (d, *J* = 15.6 Hz, 1H), 4.49 – 4.35 (m, 2H), 4.03 (s, 1H), 3.93 (dd, *J* = 10.7, 5.0 Hz, 1H), 3.67 – 3.59 (m, 1H), 2.54 (s, 3H), 2.49 (td, *J* = 8.7, 4.5 Hz, 1H), 1.98 (dt, *J* = 13.5, 4.9 Hz, 1H), 1.12 (s, 9H). **$^{13}$C NMR** (101 MHz, MeOD) δ = 174.5, 168.7, 164.9, 155.1, 144.8, 141.3, 135.9, 130.4, 129.2, 71.3, 61.1, 60.1, 57.3, 43.7, 37.8, 35.5, 26.6, 14.0. **HRMS** (ESI+) req. for $C_{22}H_{31}N_4O_3S$ $[M+H]^+$ 431.2111, found 431.2132.

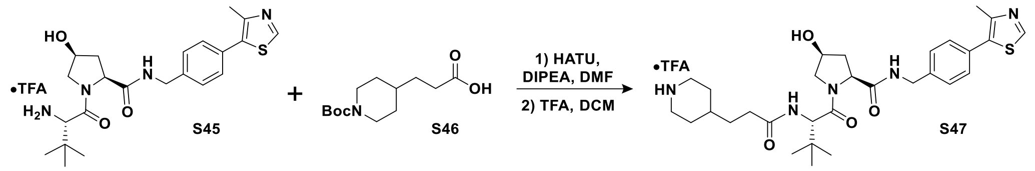


***Synthesis of S47:*** Acid **S46** (125 mg, 0.487 mmol, 1.1 equiv) was coupled to amine **S45** (141 mg, 0.443 mmol, 1.0 equiv) according to **GP1** and was purified by flash chromatography on silica ($CH_2Cl_2$ – MeOH gradient) to provide coupled product with residual DIPEA. The coupled product was deprotected according to **GP3** to provide product **S47** containing DIPEA·TFA (150 mg, 0.219 mmol, ~49% yield over two-steps).

**$^1$H NMR** (400 MHz, MeOD) δ = 9.27 (s, 1H), 7.49 (d, *J* = 8.2 Hz, 2H), 7.44 (d, *J* = 8.2 Hz, 2H), 4.58 – 4.49 (m, 2H), 4.47 – 4.36 (m, 2H), 4.04 (dd, *J* = 10.6, 5.1 Hz, 1H), 3.77 – 3.70 (m, 1H), 3.37 (dt, *J* = 12.2, 3.6 Hz, 2H), 3.01 – 2.89 (m, 2H), 2.51 (s, 3H), 2.43 (ddt, *J* = 12.7, 8.8, 4.4 Hz, 1H), 2.38 – 2.26 (m, 2H), 2.03 – 1.89 (m, 3H), 1.67 – 1.51 (m, 4H), 1.46 – 1.38 (m, 2H), 1.04 (s, 9H). **$^{13}$C NMR** (101 MHz, MeOD) δ = 175.8, 175.0, 172.6, 154.2, 146.6, 140.9, 134.9, 130.4, 130.3, 129.2, 71.5, 61.0, 59.5, 57.6, 45.2, 43.8, 37.9, 35.8, 34.5, 33.3, 32.8, 29.7, 27.0, 14.8. **HRMS** (ESI+) req. for $C_{30}H_{44}N_5O_4S$ $[M+H]^+$ 570.3109, found 570.3167.

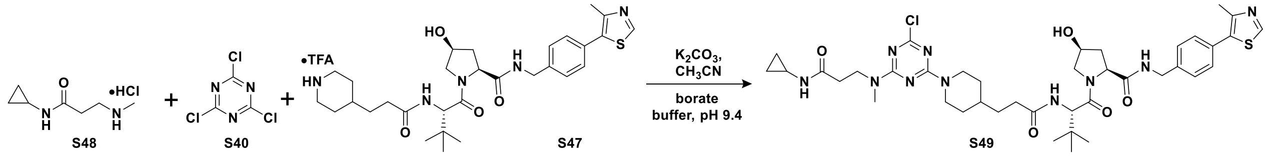


***Synthesis of S49:*** The triazine conjugate was synthesized with amine **S47** (150 mg, 0.219 mmol, 1.0 equiv) and building block **S48** (39.2 mg, 0.219 mmol, 1.0 equiv) according to **GP5** to give **S49** (113 mg, 0.137 mmol, 62% yield).

**$^1$H NMR** (400 MHz, $CDCl_3$) δ = 8.64 (s, 1H), 7.74 – 7.57 (m, 1H), 7.30 (dd, *J* = 12.3, 8.6 Hz, 4H), 6.17 (t, *J* = 10.3 Hz, 1H), 5.57 (d, *J* = 9.5 Hz, 1H), 4.69 – 4.55 (m, 4H), 4.50 (d, *J* = 9.1 Hz, 1H), 4.42 (dt, *J* = 9.3, 4.3 Hz, 1H), 4.27 (dd, *J* = 15.1, 5.0 Hz, 1H), 3.88 (dd, *J* = 11.1, 4.2 Hz, 1H), 3.83 – 3.70 (m, 3H), 3.03 (d, *J* = 11.3 Hz, 3H), 2.76 – 2.68 (m, 2H), 2.65 – 2.57 (m, 1H), 2.45 (s, 3H), 2.44 – 2.35 (m, 2H), 2.27 (d, *J* = 14.2 Hz, 1H), 2.23 – 2.12 (m, 3H), 1.67 (d, *J* = 12.7 Hz, 2H), 1.50 (qd, *J* = 16.5, 14.9, 7.3 Hz, 4H), 1.06 (q, *J* = 12.2, 11.4 Hz, 2H), 0.89 (s, 9H), 0.67 (dd, *J* = 6.2, 5.7 Hz, 2H), 0.47 – 0.41 (m, 1H), 0.40 – 0.32 (m, 1H). **$^{13}$C NMR** (101 MHz, $CDCl_3$) δ = 172.8, 172.7, 172.4, 172.1, 164.8, 163.8, 150.4, 148.5, 137.4, 131.5, 131.1, 129.6, 128.1, 71.1, 59.9, 58.5, 57.0, 53.5, 45.2, 43.7, 43.4, 35.7, 35.4, 35.1, 35.0, 33.4, 31.8, 31.7, 26.6, 26.4, 22.6, 16.1, 6.4, 6.3. **HRMS** (ESI+) req. for $C_{40}H_{56}ClN_{10}O_5S$ $[M+H]^+$ 823.3839, found 823.3923.

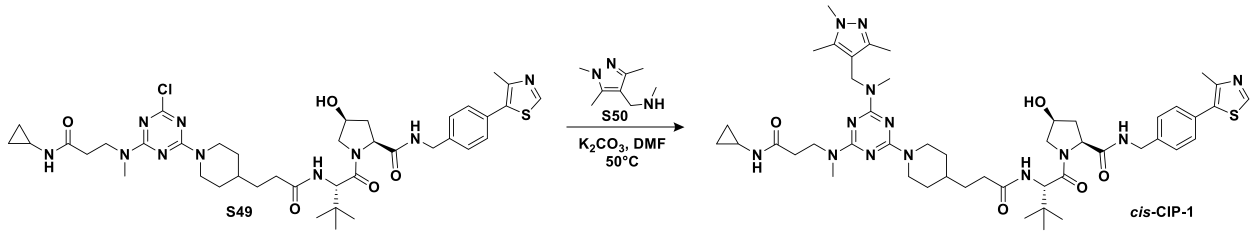


***Synthesis of cis-CIP-1:*** Chlorotriazine **S49** (56.1 mg, 0.068 mmol, 1.0 equiv) was substituted with amine **S50** (41.8 mg, 0.273 mmol, 4 equiv) according to **GP6** to provide product ***cis*-CIP-1** after prep HPLC purification (41.7 mg, 0.044 mmol, 64% yield).

$^{1}$**H NMR** (400 MHz, MeOD) δ = 8.86 (s, 1H), 7.90 (d, *J* = 8.1 Hz, 1H), 7.45 (d, *J* = 8.3 Hz, 2H), 7.40 (d, *J* = 8.3 Hz, 2H), 4.77 (d, *J* = 12.4 Hz, 2H), 4.66 (s, 2H), 4.58 – 4.48 (m, 3H), 4.42 – 4.33 (m, 2H), 4.05 (dd, *J* = 10.5, 5.1 Hz, 1H), 3.84 (t, *J* = 6.7 Hz, 2H), 3.70 (dd, *J* = 10.7, 3.9 Hz, 1H), 3.67 (s, 3H), 3.07 (s, 3H), 2.85 (s, 3H), 2.79 – 2.69 (m, 2H), 2.63 – 2.55 (m, 1H), 2.47 (s, 3H), 2.46 – 2.39 (m, 3H), 2.37 – 2.27 (m, 2H), 2.21 (s, 3H), 2.12 (s, 3H), 1.98 (dt, *J* = 13.3, 4.5 Hz, 1H), 1.73 (d, *J* = 12.1 Hz, 2H), 1.60 – 1.45 (m, 3H), 1.15 – 1.01 (m, 11H), 0.65 (td, *J* = 7.2, 5.0 Hz, 2H), 0.37 (tq, *J* = 6.9, 4.5 Hz, 2H). $^{13}$**C NMR** (101 MHz, MeOD) δ = 176.4, 176.3, 176.0, 174.9, 172.6, 172.6, 166.8, 166.1, 152.8, 149.1, 147.6, 140.1, 140.0, 133.4, 131.6, 130.4, 129.0, 114.2, 71.5, 61.0, 59.4, 59.3, 57.6, 46.2, 44.6, 43.8, 40.9, 37.9, 37.4, 35.9, 35.7, 35.5, 34.8, 33.7, 33.7, 33.2, 33.1, 32.4, 27.0, 23.3, 15.8, 12.0, 9.6, 6.4. **HRMS** (ESI+) req. for $C_{48}H_{70}N_{13}O_5S$ $[M+H]^+$ 940.5338, found 940.5430.

**Scheme S7.** Synthesis of **BL-1** BRD4$^{BD1}$ ligand with connector.

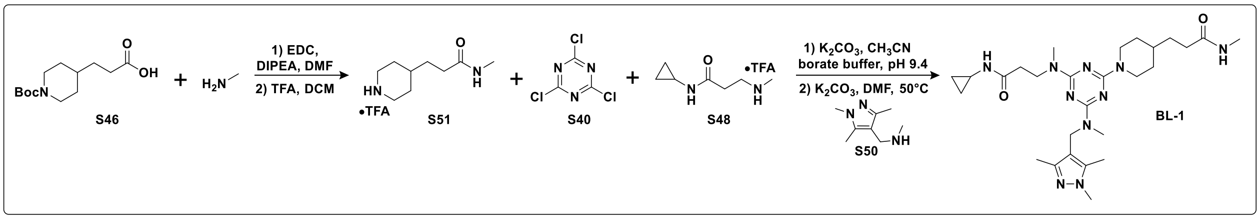


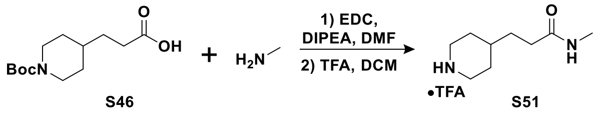


***Synthesis of S51:*** To a sample vial was added acid **S46** (300 mg, 1.17 mmol, 1.0 equiv), DMF (2 mL), EDC (246 mg, 1.28 mmol, 1.1 equiv), and DIPEA (0.611 mL, 3.50 mmol, 3 equiv). After stirring at room temperature for 5 mins, methylamine hydrochloride (94 mg, 1.40 mmol, 1.2 equiv) was added. After 1 h, UPLC-MS showed incomplete consumption of starting material and additional EDC (224 mg, 1.17 mmol, 1.0 equiv) and methylamine hydrochloride (79 mg, 1.17 mmol, 1.0 equiv) was added. After stirring for 1 h, the reaction was partitioned between $CH_2Cl_2$ and saturated aqueous $NH_4Cl$ and the organic phase was isolated. The aqueous phase was further extracted with $CH_2Cl_2$ (3x) and the organics were concentrated and purified by flash chromatography on silica ($CH_2Cl_2$ – MeOH gradient). The isolated product was further purified on a 50 g reverse-phase ISCO C18 column and eluted with a gradient of water and acetonitrile, both modified with 0.1% $NH_4OH$, to provide the desired coupling product (386 mg, 1.43 mmol, 73% yield). The isolated product was deprotected according to **GP3** to provide product **S51** (407 mg, 1.43 mmol, quantitative yield).

**$^{1}$H NMR** (400 MHz, MeOD) δ = 3.37 (dt, *J* = 13.0, 3.4 Hz, 2H), 2.94 (td, *J* = 12.9, 3.0 Hz, 2H), 2.70 (s, 3H), 2.24 (t, *J* = 7.3 Hz, 2H), 1.93 (d, *J* = 14.0 Hz, 2H), 1.59 (t, *J* = 6.7 Hz, 3H), 1.48 – 1.31 (m, 2H). **$^{13}$C NMR** (101 MHz, MeOD) δ = 176.4, 45.2, 45.1, 34.5, 33.8, 32.9, 29.7, 29.7, 26.4. **LRMS** (ESI+) req. for $C_9H_{19}N_2O$ $[M+H]^+$ 171.1, found 171.1.

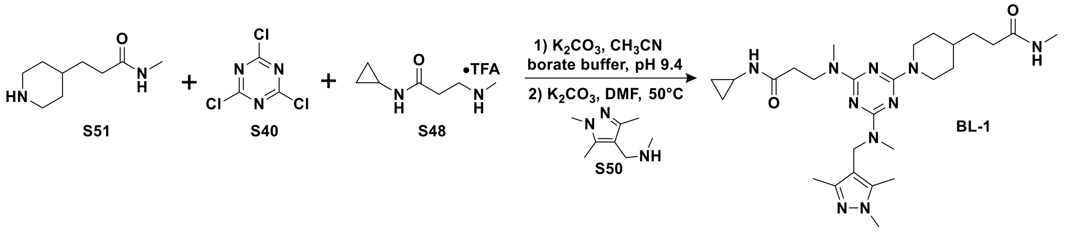


***Synthesis of BL-1:*** The disubstituted triazine intermediate was synthesized from amine **S51** (69 mg, 0.245 mmol, 1.0 equiv) and amine **S48** (63 mg, 0.245 mmol, 1.0 equiv) according to **GP5** to obtain the disubstituted chlorotriazine intermediate (23 mg, 0.055 mmol, 22% yield).

**$^{1}$H NMR** (400 MHz, $CDCl_3$) δ = 5.52 (s, 1H), 4.70 (d, *J* = 13.4 Hz, 2H), 3.82 (t, *J* = 6.6 Hz, 2H), 3.24 – 3.04 (m, 3H), 2.81 (d, *J* = 4.4 Hz, 6H), 2.74 – 2.42 (m, 3H), 2.21 (t, *J* = 7.5 Hz, 2H), 1.77 (d, *J* = 13.0 Hz, 2H), 1.71 – 1.50 (m, 3H), 1.22 – 1.05 (m, 2H), 0.75 (q, *J* = 6.9 Hz, 2H), 0.62 – 0.39 (m, 2H). **HRMS** (ESI+) req. for $C_{19}H_{31}ClN_7O_2$ $[M+H]^+$ 424.2222, found 424.2246.

The chlorotriazine intermediate (11.6 mg, 0.027 mmol, 1.0 equiv) was then substituted with amine **S50** (21.0 mg, 0.137 mmol, 5 equiv) according to **GP6** to provide product **BL-1** after preparative HPLC purification (9.5 mg, 0.018 mmol, 64% yield).

**$^{1}$H NMR** (400 MHz, MeOD) δ = 4.77 (d, *J* = 12.3 Hz, 2H), 4.67 (s, 2H), 3.84 (t, *J* = 6.7 Hz, 2H), 3.68 (s, 3H), 3.07 (s, 3H), 2.86 (s, 3H), 2.76 (td, *J* = 12.6, 2.3 Hz, 5H), 2.71 (d, *J* = 4.2 Hz, 3H), 2.59 (tt, *J* = 7.4, 3.8 Hz, 1H), 2.44 (t, *J* = 6.7 Hz, 2H), 2.22 (s, 5H), 2.12 (s, 3H), 1.75 (d, *J* = 3.4 Hz, 1H), 1.72 (d, *J* = 3.0 Hz, 1H), 1.61 – 1.45 (m, 3H), 1.10 (qd, *J* = 12.5, 4.1 Hz, 2H), 0.70 – 0.62 (m, 2H), 0.42 – 0.34 (m, 2H). **$^{13}$C NMR** (101 MHz, MeOD) δ = 176.9, 176.1, 166.9, 166.8, 166.1, 147.7, 140.1, 114.2, 46.2, 44.5, 40.9, 37.4, 35.7, 35.5, 34.8, 34.3, 33.8, 33.2, 32.4, 26.3, 23.3, 11.9, 9.6, 6.4. **HRMS** (ESI+) req. for $C_{27}H_{45}N_{10}O_2$ $[M+H]^+$ 541.3721, found 541.3771.

#### 7. Building block validation for the CIP-DEL library.

To select amines for inclusion in the triazine substitution reactions, two building block profiling campaigns were performed using library-like reaction conditions. First, we profiled the one-pot triazine addition and cycle 2 amine substitution reaction with a collection of 264 primary amines and 169 secondary amines (**Scheme S8**). Next, we profiled the cycle 3 amine substitution reaction with the same collection of building blocks and a benzylamine-substituted model triazine substrate (**Scheme S9**). We used UPLC-MS analysis of each substitution reaction to determine the percentage of product formed by integration of the UV signal at 260 nm. For library inclusion, we selected building blocks that generated 75% or more product peak area, excluding remaining starting material peak area (which was general very minor) and non-DNA peaks lacking a total ion chromatogram (TIC) signal.

**Scheme S8.** Experimental profiling of cycle 2 amines.

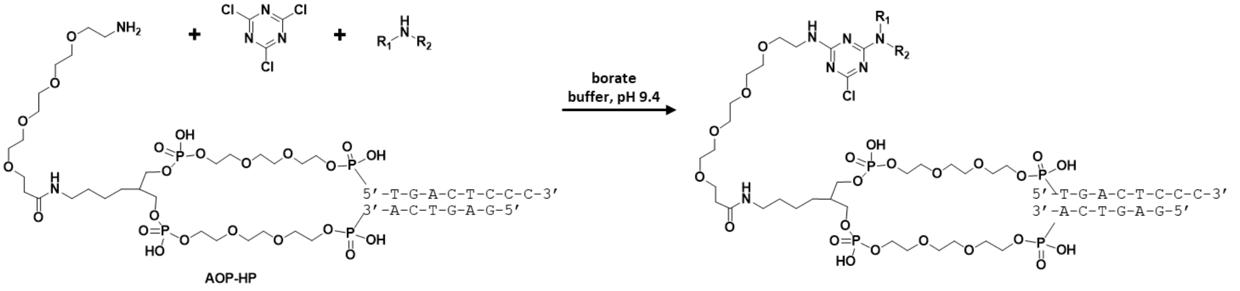

***Screening of cycle 2 amine building blocks:*** using 96-well plates, each building block was profiled by adding 5 µL of 150 mM borate buffer pH 9.4 and 1 µL of AOP-HP (1 mM solution in borate buffer, 1 nmol per well, 1 equiv). The plates were cooled to 4°C on a thermocycler and 5 µL of cyanuric chloride (3 mM in $CH_3CN$, 15 nmol, 15 equiv) was added. The reactions were incubated at 4°C for 1 h, then 1 µL of each test amine (200 mM in 1:1 $CH_3CN$ : $dH_2O$, 200 nmol, 200 equiv) was added. The solutions were incubated at 4°C for 18 h and ethanol precipitated with the addition of 1 µL of 5 M NaCl and 80 µL ethanol. After incubating the 96-well plates on dry ice for 10 min, the pellets were isolated by centrifugation and further washed with 100 µL of ice cold 90% ethanol. Final pellets were dried under vacuum for 60 min. The dried pellets were dissolved in 150 µL $dH_2O$ and analyzed directly by UPLC-MS. Using a cutoff of 75% product peak area, we selected 163 primary amines and 127 secondary amines for library inclusion.

**Scheme S9.** Experimental profiling of cycle 3 amines.

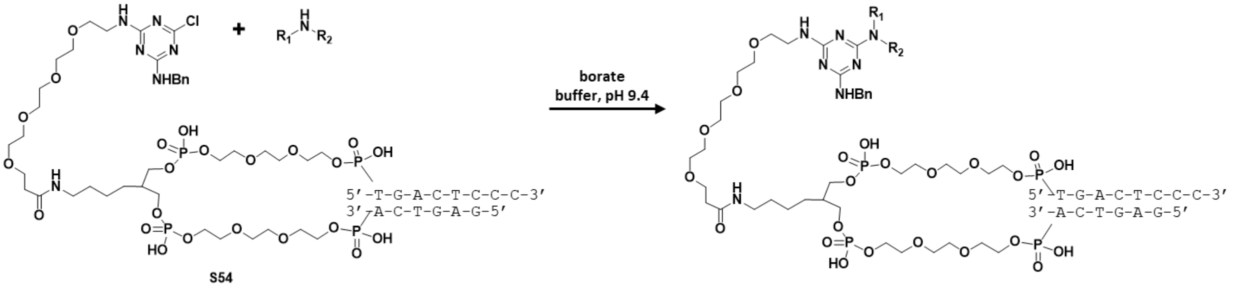

***Screening of cycle 3 amine building blocks:*** using a benzylamine substituted triazine **S54** as the model substrate, we profiled our collection of primary and secondary amines for cycle 3 substitutions. In 96-well plates, 13.8 µL of **S54** (72.5 µM solution in borate buffer pH 9.4, 1 nmol per well, 1 equiv) was added to each well followed by 1.2 µL of each amine (200 mM solutions in 1:1 $CH_3CN$ : $dH_2O$, 240 nmol, 240 equiv). The plates were sealed and heated to 80°C in a thermocycler for 21 h. Each reaction was ethanol precipitated with the addition of 1 µL of 5 M NaCl and 80 µL ice cold ethanol. Plates were incubated on dry ice for 10 min and pellets were isolated by centrifugation and further washed with 100 µL of ice cold 90% ethanol. Final pellets were dried under vacuum for 30 min, then dissolved in 100 µL of $dH_2O$ and analyzed directly by UPLC-MS. Using a cutoff of 75% product peak area, we selected 146 primary amines and 103 secondary amines for cycle 3 substitutions.

#### 8. CIP-DEL library production.

**Scheme S10.** Synthesis of azido-AOP-HP.

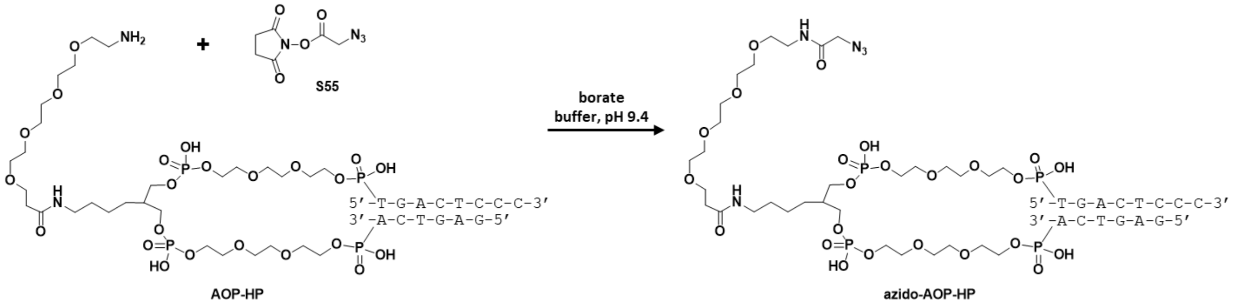


***Synthesis of Azido-AOP-HP:*** A 400 mM solution of activated azido-acetic acid **S55** was generated *in situ* by combining EDC·HCl (24 mg, 0.125 mmol, 1.0 equiv) and 2-azidoacetic acid (11 µL, 0.150 mmol, 1.2 equiv) in DMF (0.2 mL), followed by the addition of N-hydroxysuccinimide (16 mg, 0.138 mmol, 1.1 equiv) in DMF (0.113 mL). The reaction mixture was stirred at room temperature for 1 h, then used as is.

A solution of AOP-HP (1.0 mL, 1.0 µmol, 1.0 equiv) in 150 mM borate buffer, pH 9.4 was combined with the DMF solution of NHS-azido-acetic acid prepared above (0.10 mL, 40 µmol, 40 equiv). The resulting mixture was vortexed, spun down, and incubated at room temperature. After 1 h, UPLC-MS analysis showed complete consumption of AOP-HP starting material. The DNA was ethanol precipitated with the addition of 5 M NaCl (0.10 mL) and cold EtOH (2.75 mL), and then incubated on dry ice for 30 mins. The suspension was centrifuged and the supernatant decanted. The isolated pellet was washed with cold EtOH (2 x 3 mL, 90% EtOH) and the final pellet was dried under vacuum. The pellet was dissolved in 100 mM PBS, pH 8.0 (0.65 mL) and quantified by NanoDrop: 1.00 µmol, 1.54 mM, quantitative yield.

**MS** (ESI-) expected 5265 Da, found 5268 Da.

***UV:***

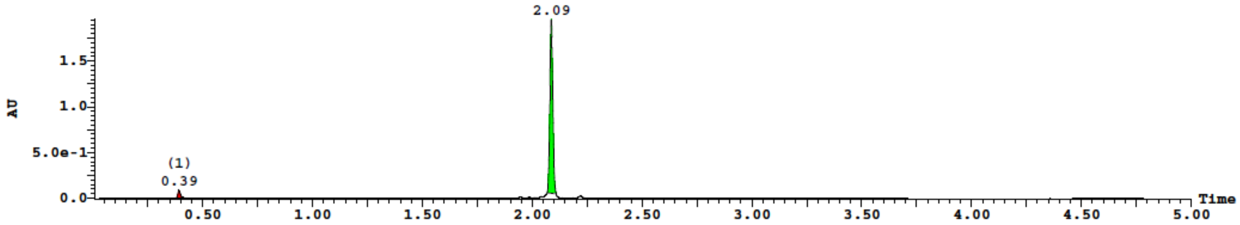


***TIC:***

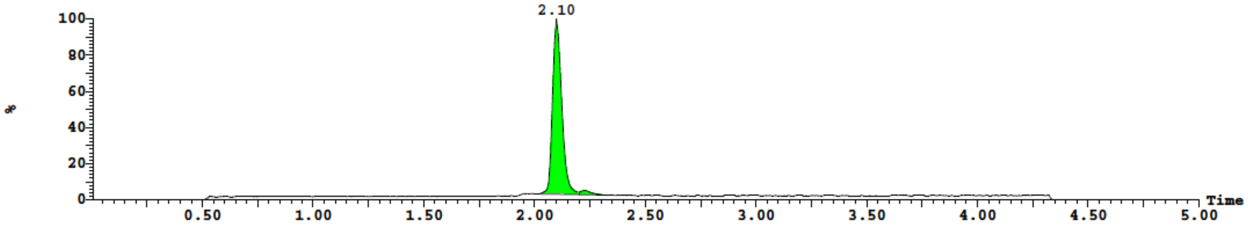

**Scheme S11.** Attachment of VHL-connector conjugates to azido-AOP-HP.

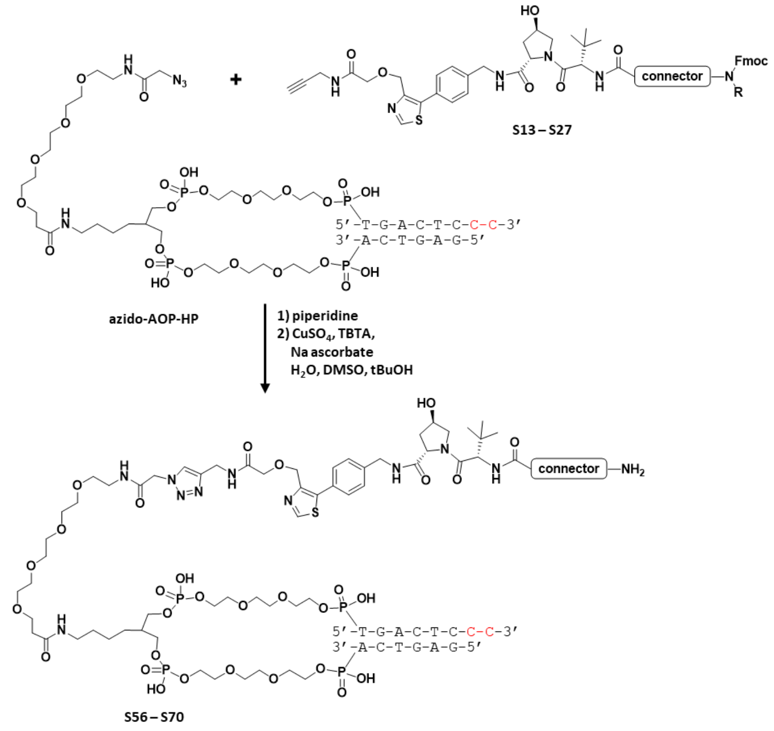


***Synthesis of S56 – S70:*** VHL-connector conjugates **S13** – **S27** were dissolved in DMSO (200 mM) and coupled to the azido-AOP-HP as follows. First, the Fmoc group was deprotected *in situ* by combining DMSO (30 µL), VHL-connector conjugate (200 mM in DMSO, 40 µL, 8.00 µmol, 40 equiv), and piperidine (0.8 µL, 8.00 µmol, 40 equiv). The mixture was vortexed briefly, spun down, and left standing at room temperature for 0.5 h. To the resulting solution was added azido-AOP-HP (1.13 mM solution in 100 mM phosphate buffer pH 8.0, 177 µL, 0.2 µmol, 1.0 equiv), sodium ascorbate (10 mM solution in d$H_2O$, 40 µL, 0.40 µmol, 2 equiv), TBTA (2 mM in 4:1 *t*BuOH : DMSO, 20 µL, 0.04 µmol, 0.2 equiv), and $CuSO_4$ (50 mM solution in d$H_2O$, 8 µL, 0.40 µmol, 2 equiv). The final reaction mixture was incubated at room temperature with shaking until UPLC-MS analysis showed complete consumption of starting material. A solution of sodium diethyldithiocarbamate (100 mM in d$H_2O$, 15 µL, 3.75 equiv based on Cu) was added to scavenge copper from the reaction. Upon scavenger addition, a brown precipitate formed which was removed by centrifugation and the isolated supernatant was ethanol precipitated with the addition of 5 M NaCl (30 µL) and cold EtOH (825 µL). The mixture was incubated at -20°C for 1 h and the precipitate was isolated by centrifugation. The pellet was washed with an additional 1 mL cold EtOH and allowed to dry for 1 h. The crude material was purified by preparative HPLC and product-containing fractions were lyophilized. The purified material was dissolved in 150 µL of 100 mM phosphate buffer pH 8.0 and quantified by NanoDrop and analyzed by UPLC-MS (gradient: 10 - 45% B).

***Synthesis of S56:*** isolated 0.118 µmol, 59% yield. **MS** (ESI-) expected 5891 Da, found 5894 Da.

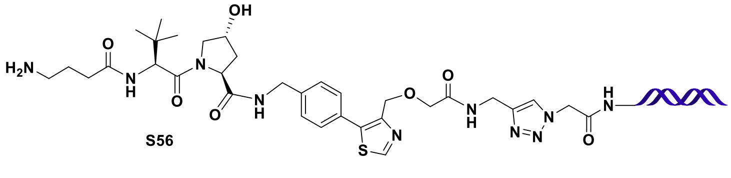

S56

***UV:***

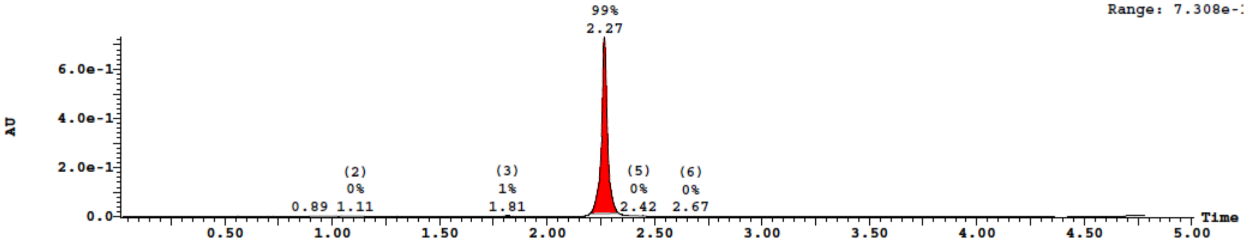


***TIC:***

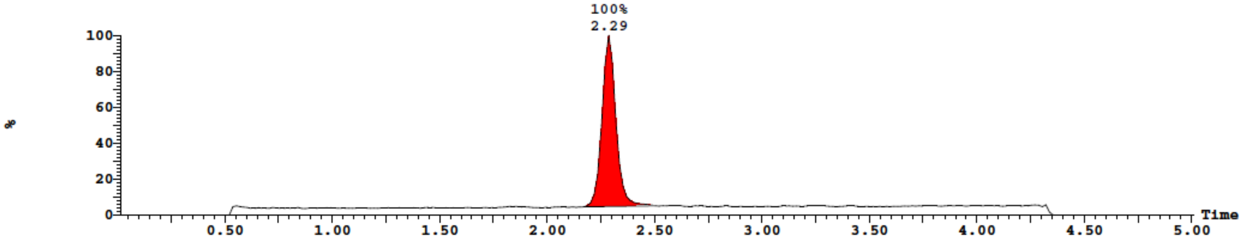


***Synthesis of S57:*** isolated 0.125 µmol, 62% yield. **MS** (ESI-) expected 5905 Da, found 5908 Da.

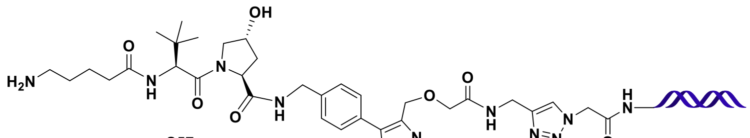


S57

***UV:***

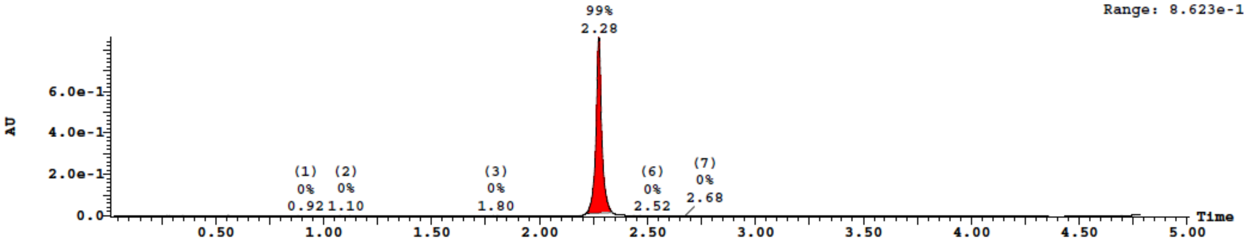


***TIC:***

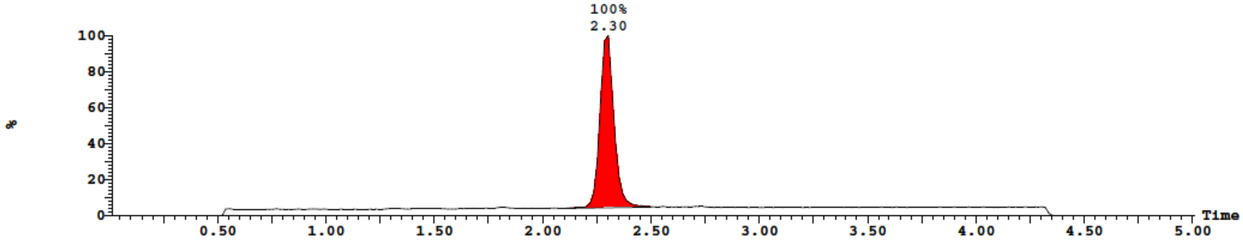

***Synthesis of S58:*** isolated 0.124 µmol, 62% yield. **MS** (ESI-) expected 5919 Da, found 5922 Da.

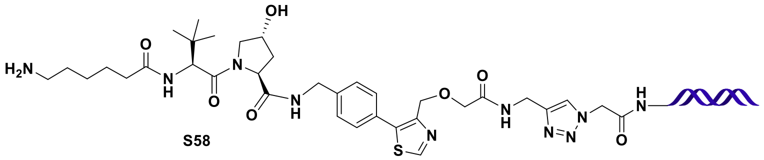


**S58**

***UV:***

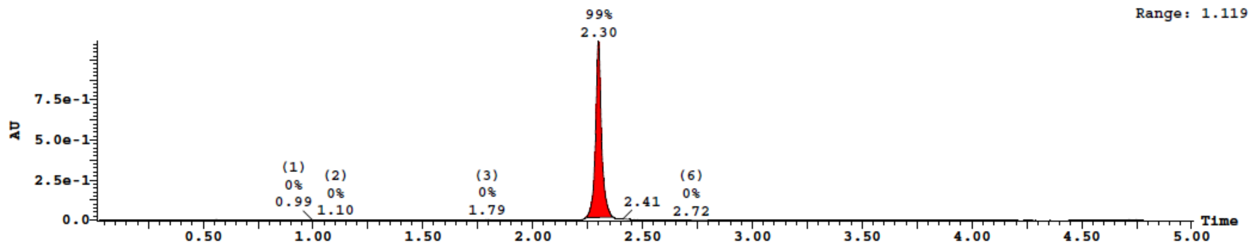


***TIC:***

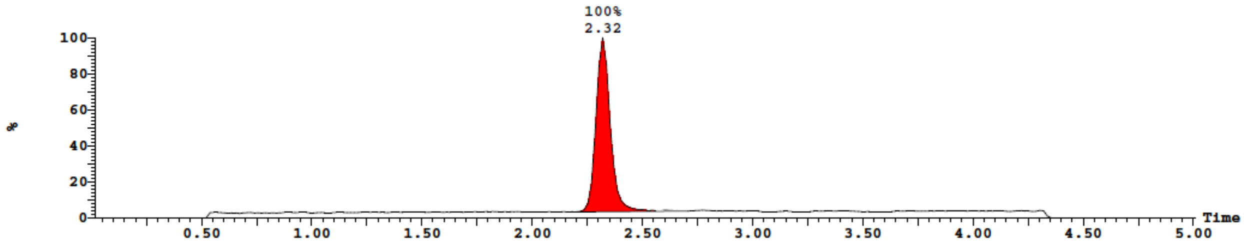


***Synthesis of S59:*** isolated 0.116 µmol, 58% yield. **MS** (ESI-) expected 5933 Da, found 5936 Da.

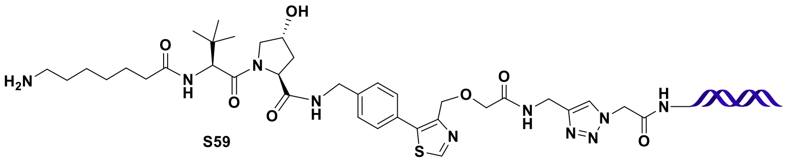


**S59**

***UV:***

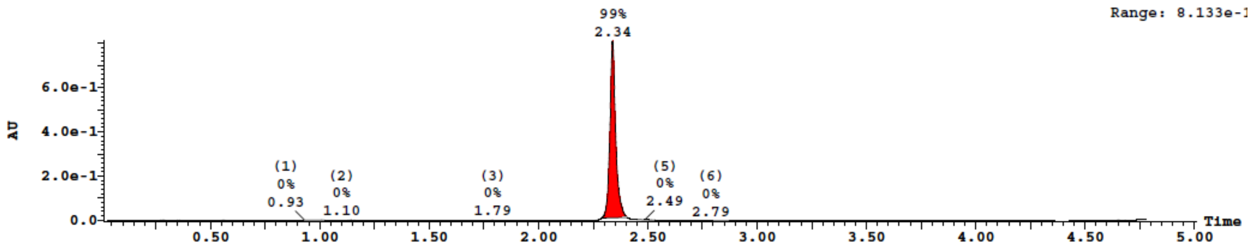

*TIC:*

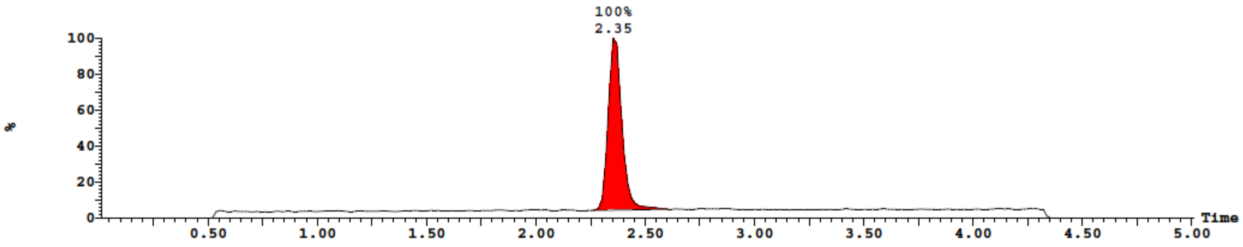


***Synthesis of S60:*** isolated 0.108 µmol, 54% yield. **MS** (ESI-) expected 5921 Da, found 5924 Da.

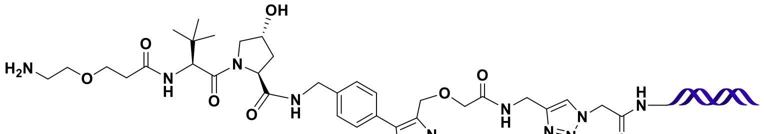

S60

*UV:*

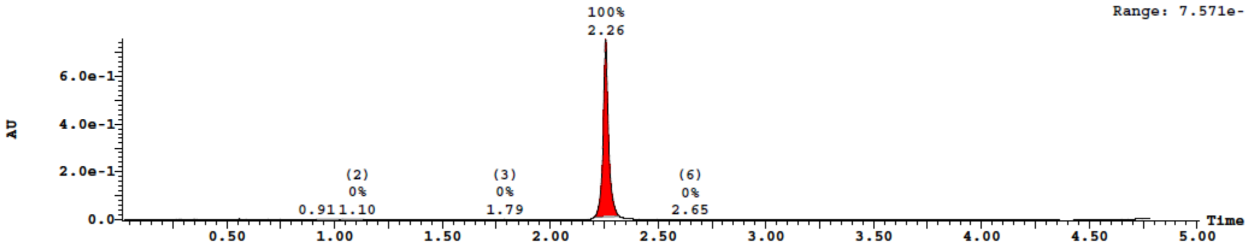


*TIC:*

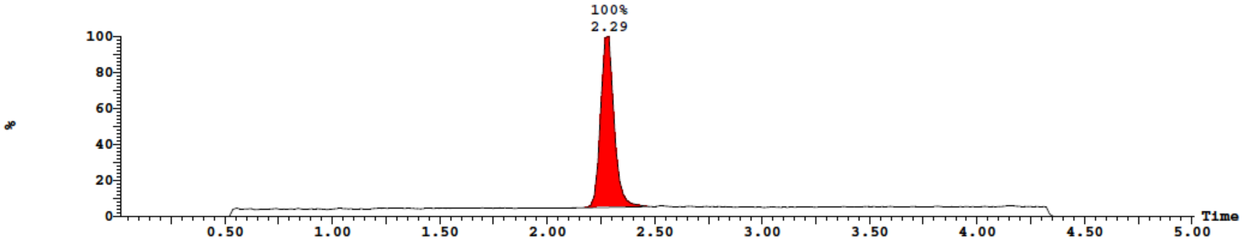


***Synthesis of S61:*** isolated 0.124 µmol, 62% yield. **MS** (ESI-) expected 5951 Da, found 5954 Da.

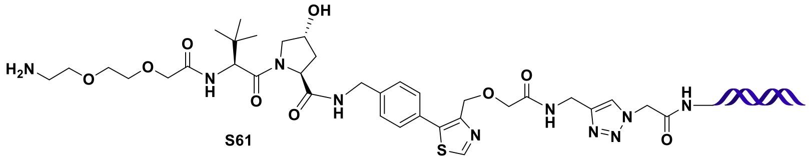

S61

*UV:*

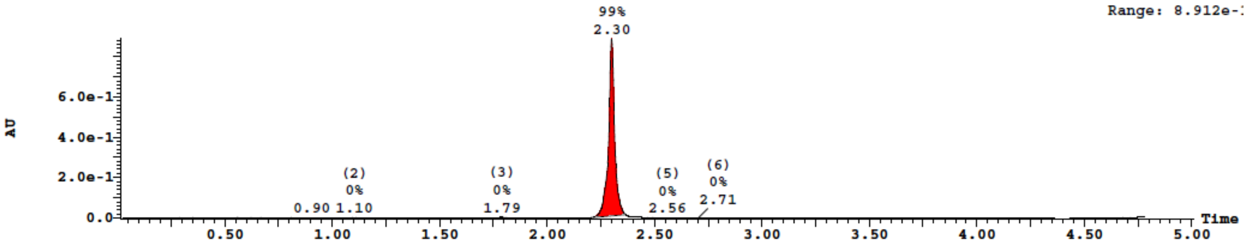


*TIC:*

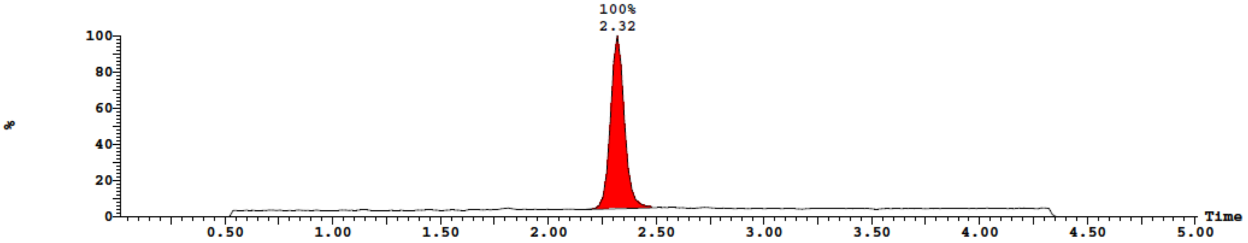


***Synthesis of S62:*** isolated 0.113 µmol, 56% yield. **MS** (ESI-) expected 5995 Da, found 5998 Da.

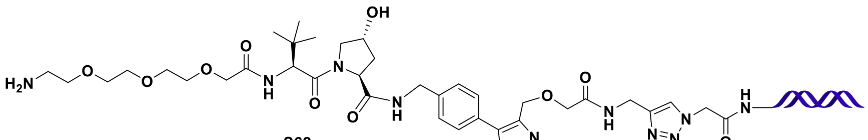


S62

*UV:*

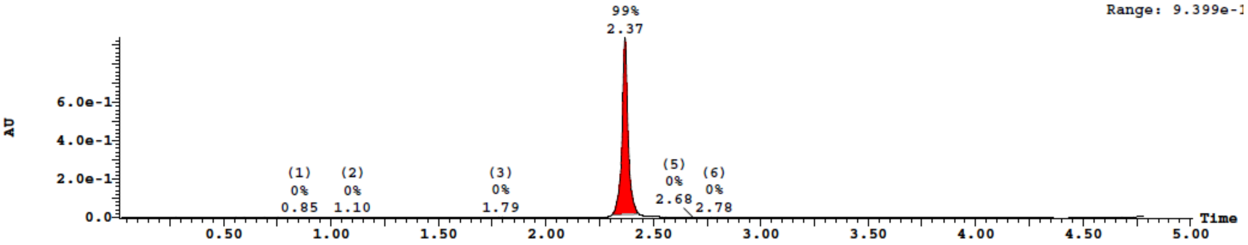


*TIC:*

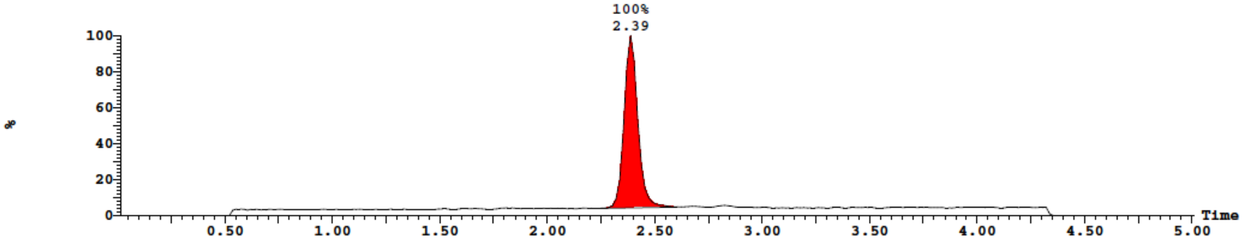

***Synthesis of S63:*** isolated 0.117 µmol, 58% yield. **MS** (ESI-) expected 5920 Da, found 5923 Da.

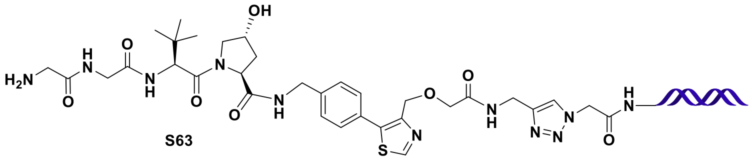

S63

*UV:*

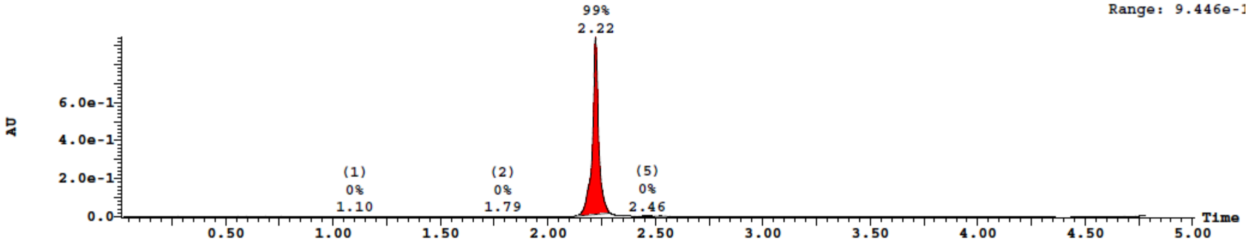


*TIC:*

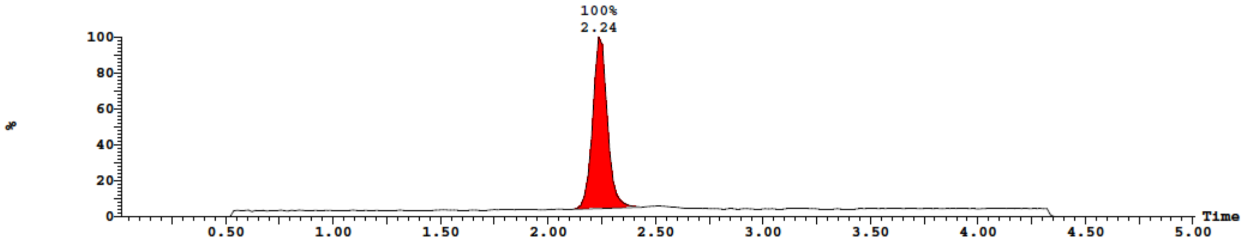


***Synthesis of S64:*** isolated 0.123 µmol, 62% yield. **MS** (ESI-) expected 5939 Da, found 5942 Da.

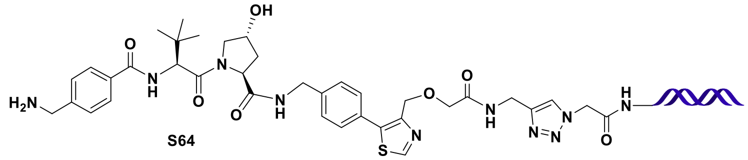

S64

*UV:*

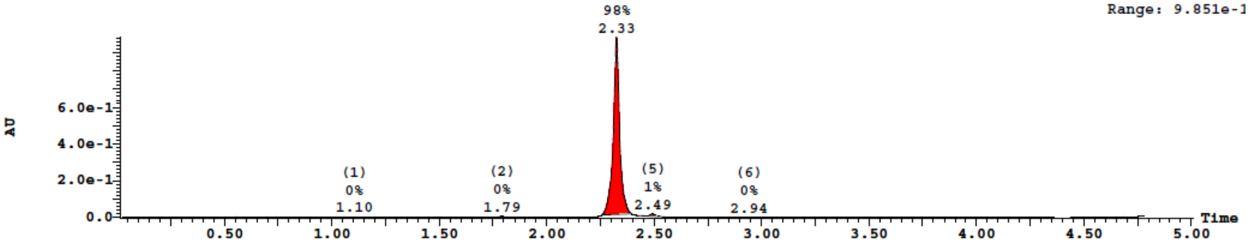

*TIC:*

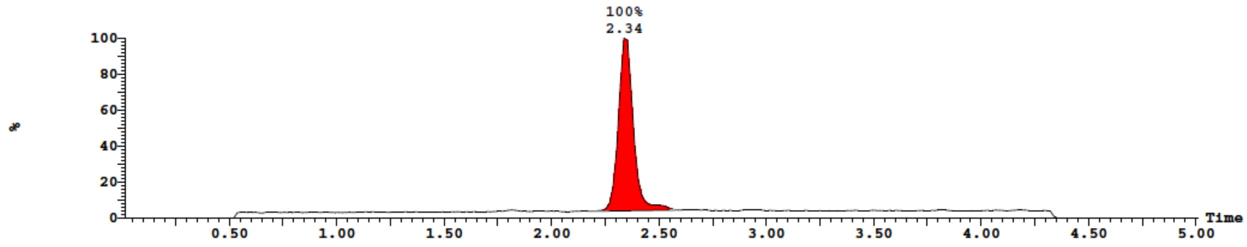


***Synthesis of S65:*** isolated 0.129 µmol, 64% yield. **MS** (ESI-) expected 5945 Da, found 5948 Da.

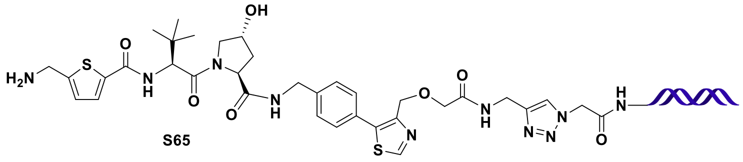

S65

*UV:*

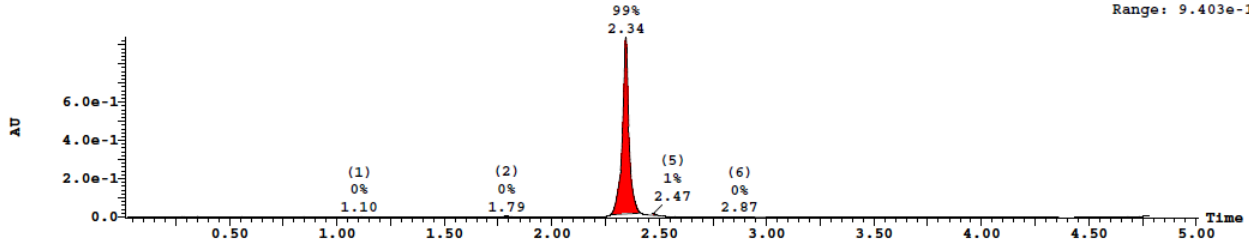


*TIC:*

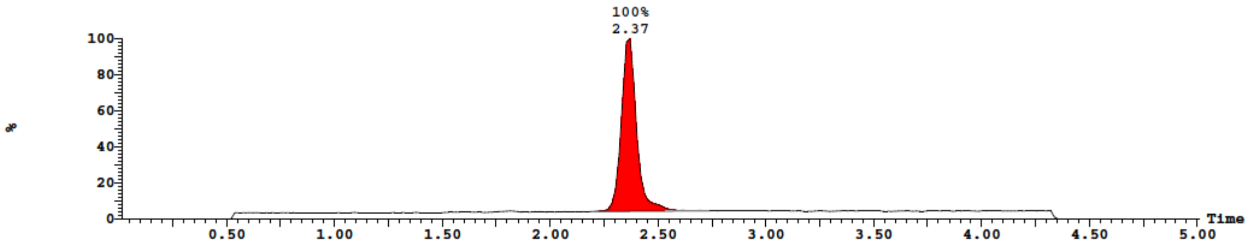


***Synthesis of S66:*** isolated 0.133 µmol, 67% yield. **MS** (ESI-) expected 5917 Da, found 5920 Da.

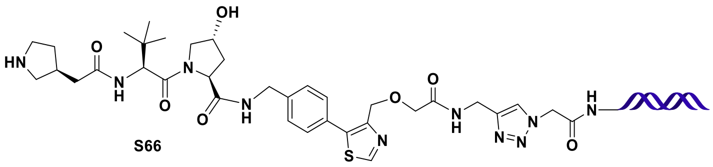

S66

*UV:*

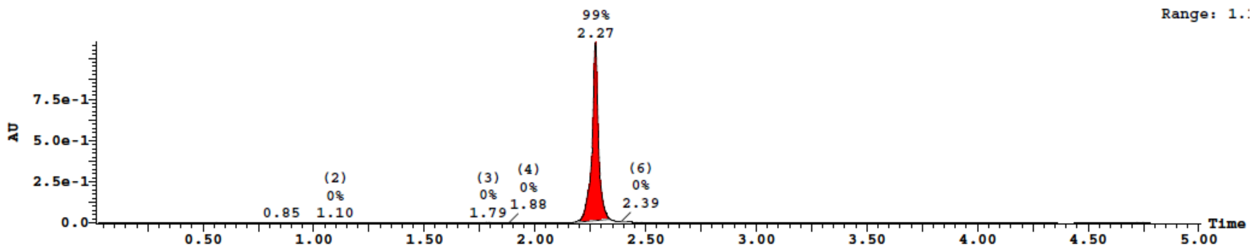


*TIC:*

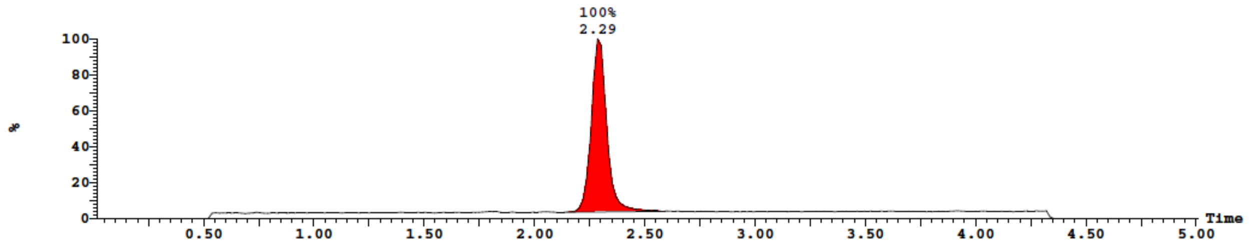


***Synthesis of S67:*** isolated 0.074 µmol, 37% yield. **MS** (ESI-) expected 5932 Da, found 5935 Da.

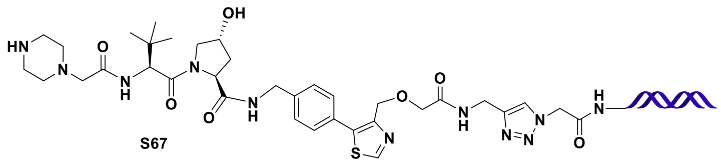

S67

*UV:*

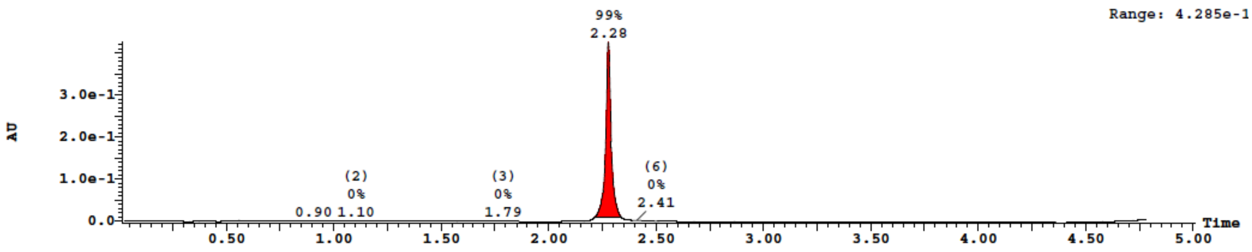


*TIC:*

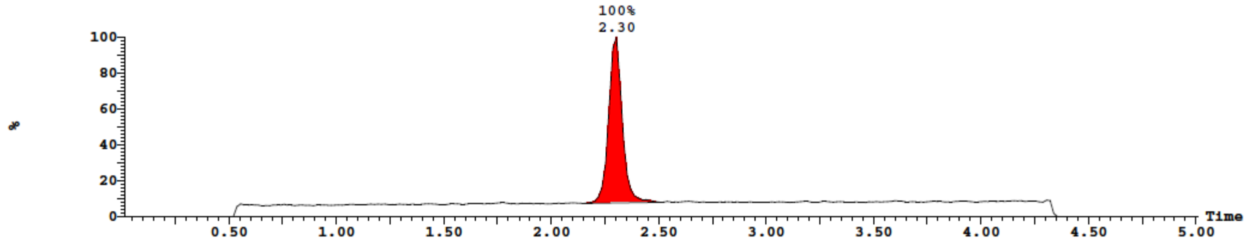

***Synthesis of S68:*** isolated 0.079 µmol, 40% yield. **MS** (ESI-) expected 5903 Da, found 5906 Da.

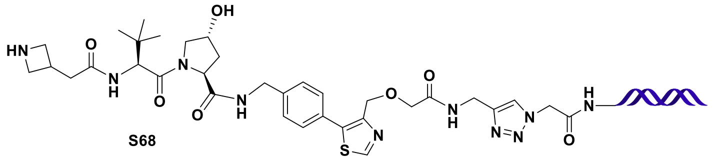


**S68**

***UV:***

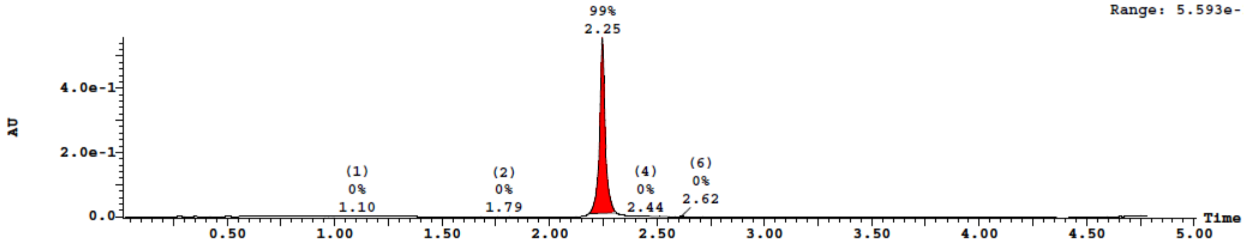


***TIC:***

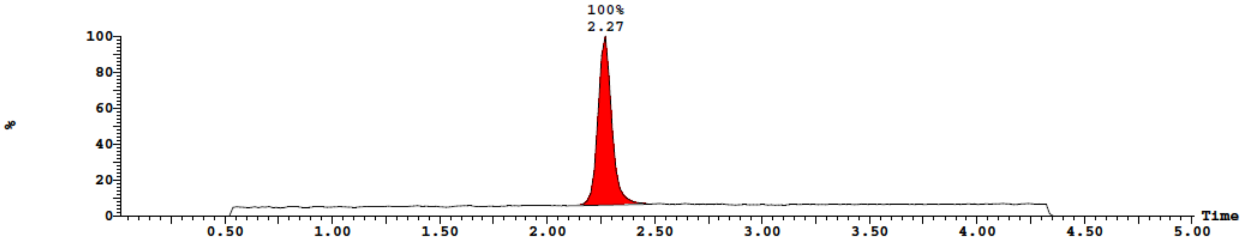


***Synthesis of S69:*** isolated 0.074 µmol, 37% yield. **MS** (ESI-) expected 5931 Da, found 5933 Da.

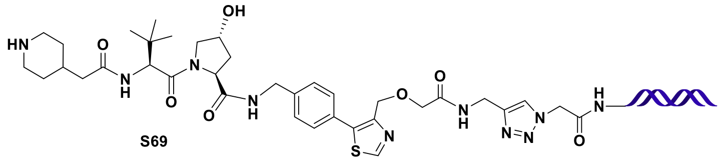


**S69**

***UV:***

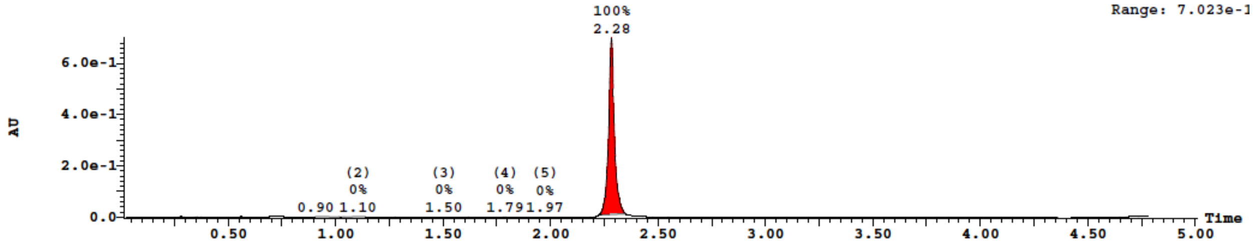

*TIC:*

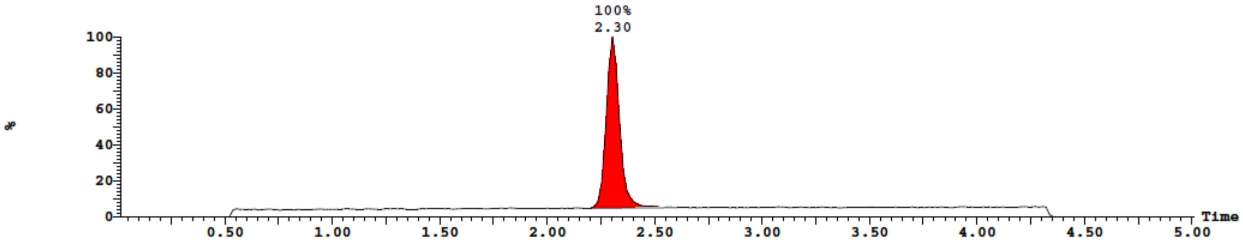


***Synthesis of S70:*** isolated 0.065 µmol, 32% yield. **MS** (ESI-) expected 5945 Da, found 5948 Da.

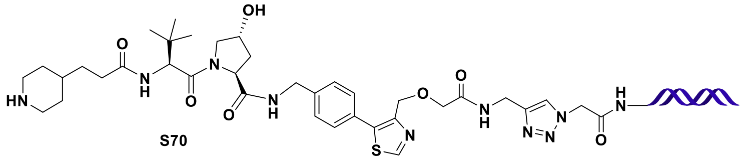


**S70**

*UV:*

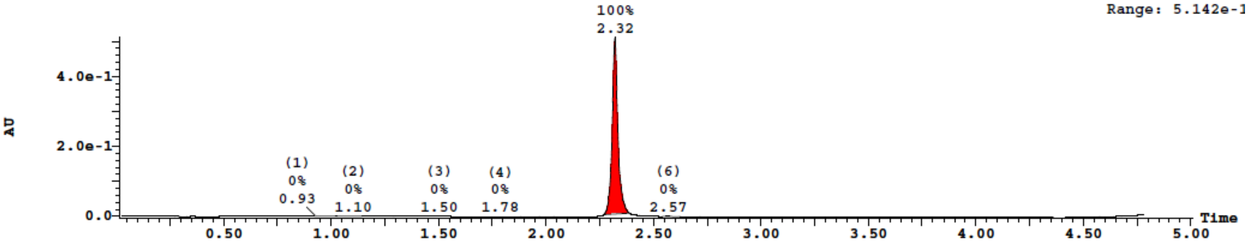


*TIC:*

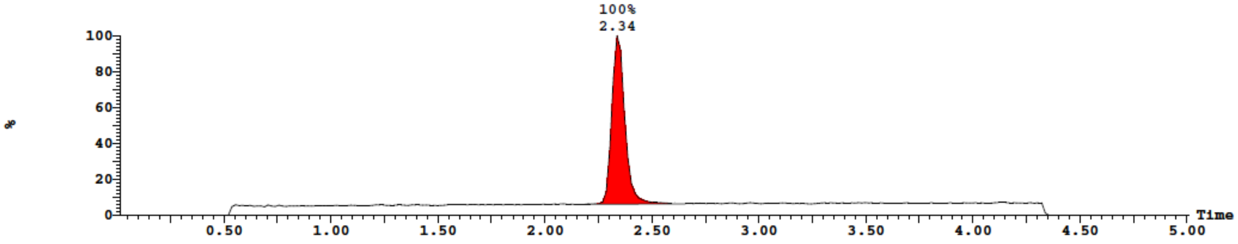

**Scheme S12.** Double ligation for addition of forward primer binding site and cycle 1 connector encoding tags.

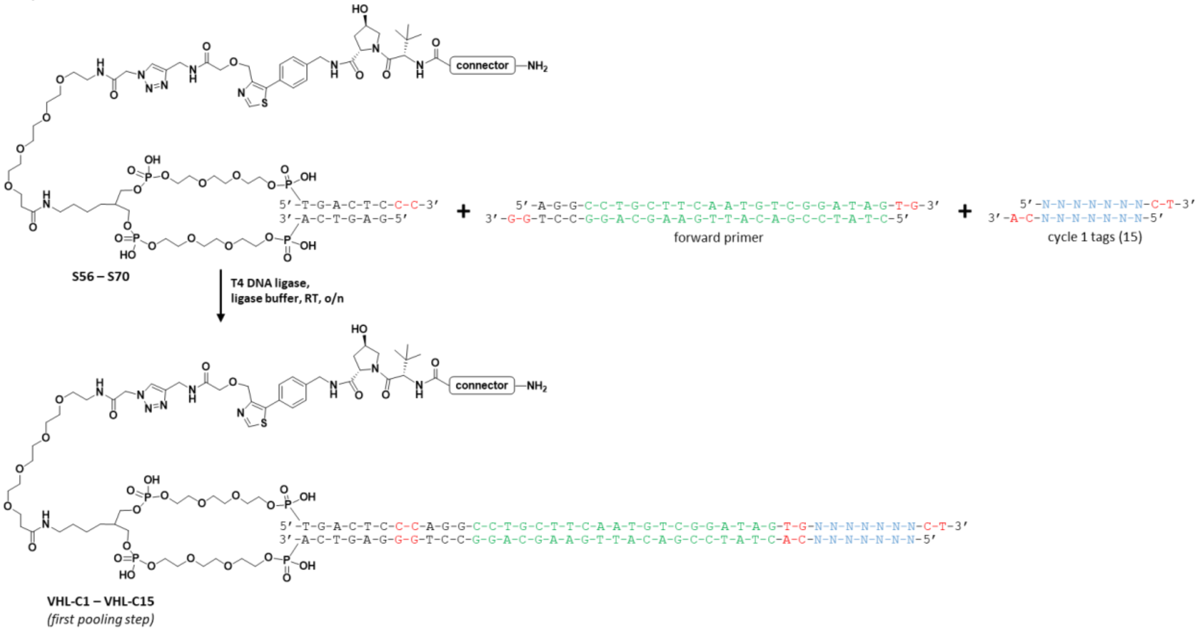


***Addition of forward primer binding site and cycle 1 tags:*** To a solution of each VHL ligand-connector DNA conjugate **S56** – **S70** (32 nmol, 66 µL, 1.6 equiv) was added 10x T4 ligase buffer (Lucigen, 20 µL), pre-annealed forward primer binding site oligo (1 mM in $dH_2O$, 20 µL, 20 nmol, 1 equiv), a unique cycle 1 tag (1 mM in $dH_2O$, 44 µL, 44 nmol, 2.2 equiv), and T4 DNA Ligase (Lucigen, 30 Weiss U / µL, 0.8 µL, 24 Weiss U / ligation). Each ligation was gently shaken overnight at room temperature. Reaction completion was assessed by running an analytical sample of each ligation reaction on a 4% agarose gel (E-gel EX 4%), which indicated complete consumption of forward primer (**Figure S3**). Each ligation was ethanol precipitated with the addition of 25 µL of a 5 M NaCl solution and 600 µL of ice-cold ethanol. The reactions were incubated at -20°C for 1 h and the precipitate was isolated by centrifugation. The supernatant was carefully decanted and the pellets were air-dried overnight. Each pellet was dissolved in dH2O and pooled into a Amicon Ultra 3 kDa centrifugal filter and buffer exchanged with ~5 volumes of $dH_2O$ to remove excess salt. The final combined library material was analyzed by UPLC-MS (gradient: 10 - 45% B) and showed minor peaks for residual forward primer and cycle 1 tags and a large broad peak containing the product mixture (**Figure S4**). The pooled material was quantified spectrophotometrically (NanoDrop): 21.24 µg/µL (A260/280= 1.42; A260/230= 1.60); volume = 430 µL; ~340 nmol (~quantitative yield; theoretical maximum: 300 nmol, plus excess forward primer and cycle 1 tag DNA).

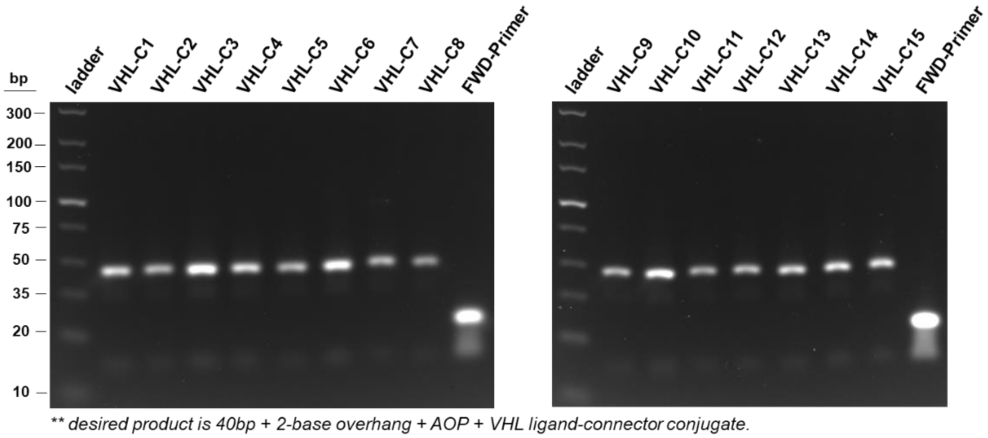


**Figure S3.** Agarose gel analysis of double ligations between **S56** – **S70**, the forward primer oligo, and the cycle 1 tags for connector encoding.

*UV:*

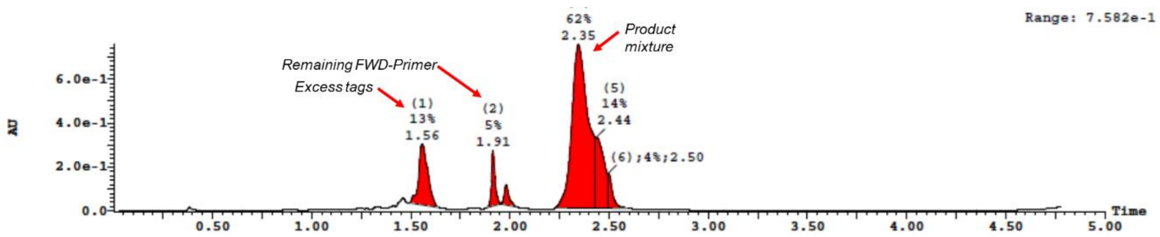


*TIC:*

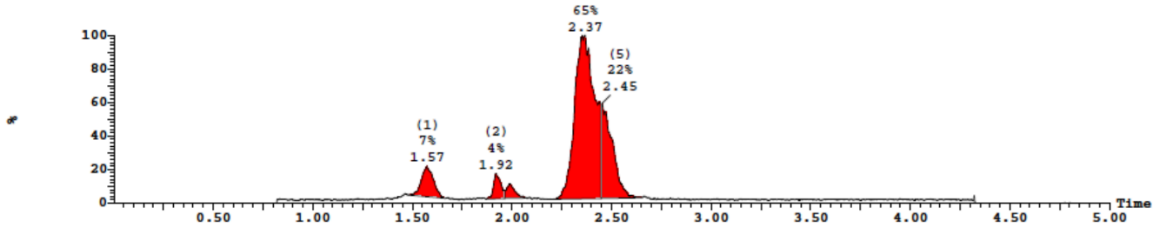


**Figure S4.** UPLC-MS analysis of pooled and buffer-exchanged library material after double ligation of the forward primer oligo and cycle 1 tags.

**Scheme S13.** Cycle 2 tag ligations for encoding of triazine amine substituents.

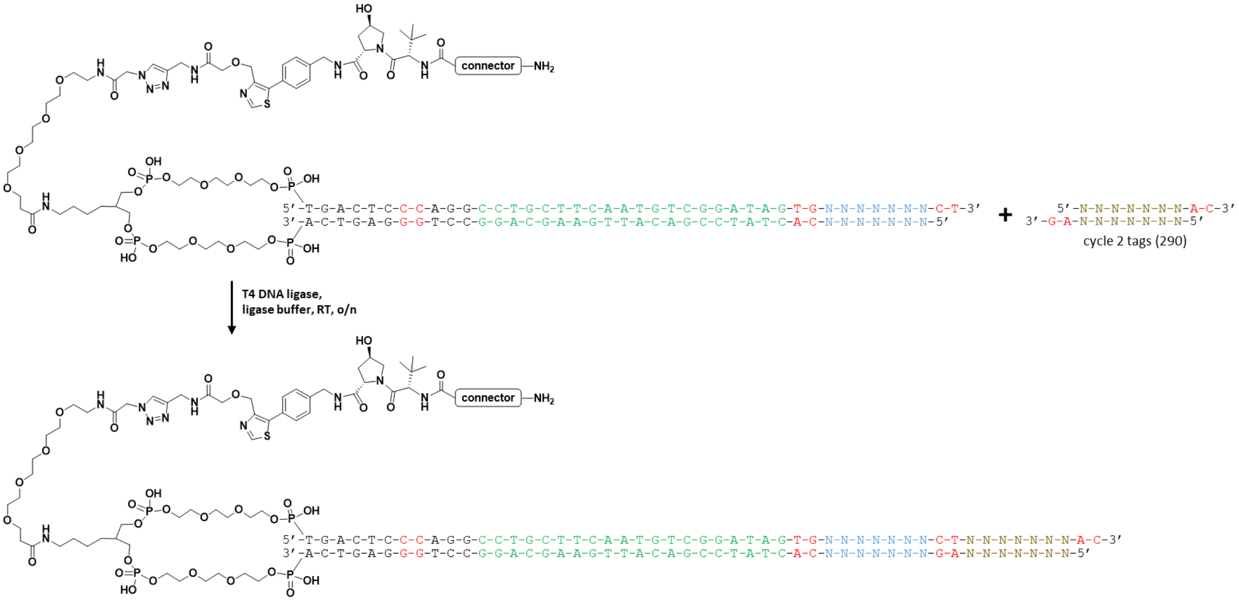


***Addition of cycle 2 tags for amine encoding:*** The pooled material from the forward primer and cycle 1 ligations (300 nmol) was split across 290 wells using 96-well plates (1.03 nmol / well in 5 µL $dH_2O$, 1 equiv). To each well was added a unique cycle 2 tag (2.3 µL, 1 mM solution in $dH_2O$, 2.3 nmol, 2.2 equiv) followed by a mixture of T4 DNA ligase and 5x ligase buffer (2.7 µL, 1.2 Weiss U / well) for a final volume of 10 µL per ligation. The ligations were incubated at RT overnight and an analytical sample from the diagonal wells across each plate was analyzed on a 4% agarose gel to assess reaction completion (**Figure S5**). All ligations appeared complete, and the reactions were precipitated with the addition of 2 µL of 5 M NaCl and 40 µL of ice-cold ethanol. Plates were incubated at -20°C for 1 h and the pellets were isolated by centrifugation. The supernatant was carefully decanted and the pellets were washed with an additional 100 µL of ice-cold ethanol and then air-dried for 1 h. Each pellet was dissolved in 10 µL of 150 mM borate buffer, pH 9.4 and used directly for $S_NAr$ reactions.

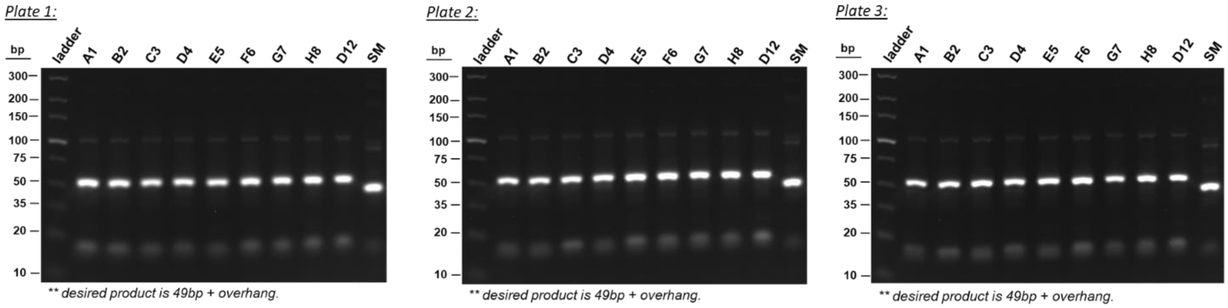


**Figure S5.** Agarose gel analysis of cycle 2 tag ligations across diagonal wells of each 96-well plate.

**Scheme S14.** Triazine addition and $S_NAr$ substitution of connector amines.

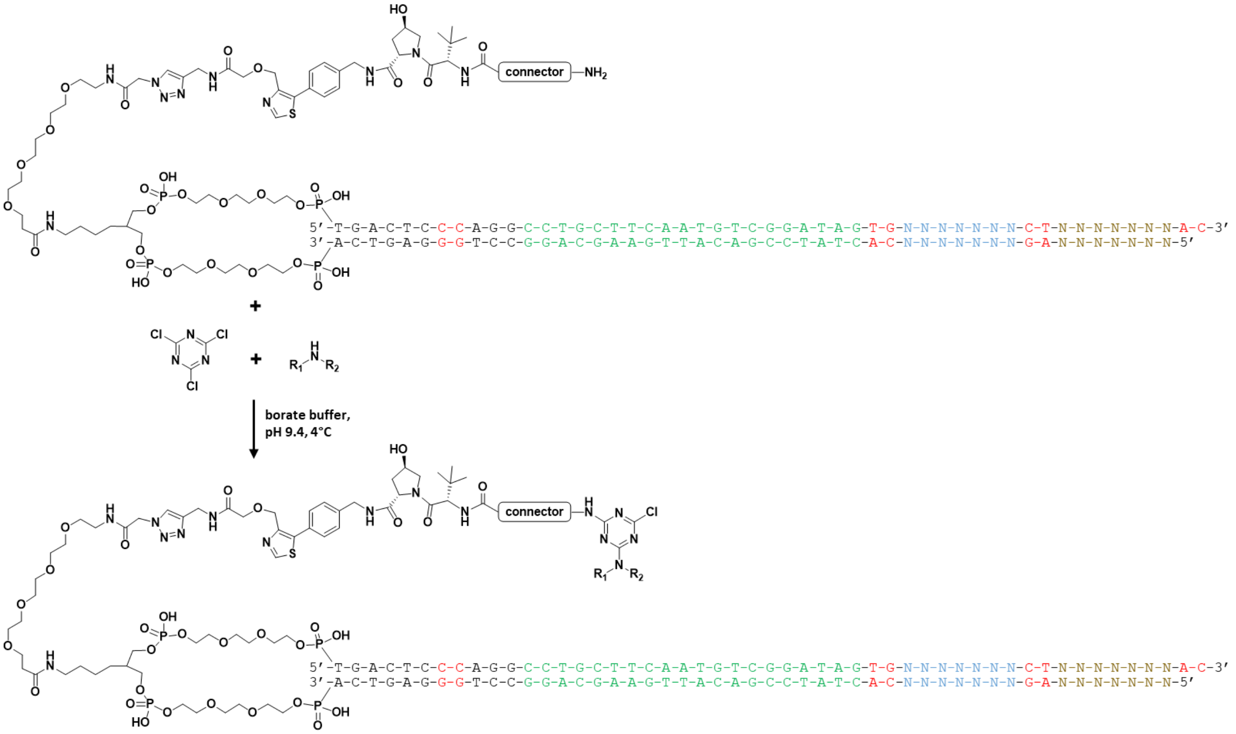


***Connector amine substitution with triazine and further $S_NAr$ with 290 cycle 2 amines:*** Each of the 290 wells from the above cycle 2 ligations (0.1 mM in 150 mM borate buffer pH 9.4, 10 µL) was cooled to 4°C in a thermocycler and cyanuric chloride (5 µL of 3 mM solution in $CH_3CN$, 15 equiv) was added. The plates were kept at 4°C for 1 h, then each well received one of 290 amines (200 mM in 1:1 $CH_3CN$:$dH_2O$, 1 µL, 200 equiv). The reactions were maintained at 4°C for 9 h and then precipitated with the addition of 2.5 µL of 5 M NaCl and 80 µL of ice-cold ethanol. The plates were incubated at -20°C for 1.25 h and the precipitate was isolated by centrifugation. The supernatant was carefully decanted and the pellets were washed with an additional 100 µL of ice-cold ethanol and air-dried for 2 h. The air-dried pellets were dissolved in $dH_2O$ (50 µL), pooled, and transferred to an Amicon Ultra 10 kDa spin column. The pooled material was concentrated and buffer exchanged with ~10 volumes of $dH_2O$ to remove excess salt and reagents. The pooled material was analyzed by UPLC-MS (**Figure S6**) and quantified by spectrophotometrically (NanoDrop): 13.66 µg/µL (A260/280= 1.83; A260/230= 2.02); volume = 900 µL.

***UV:***

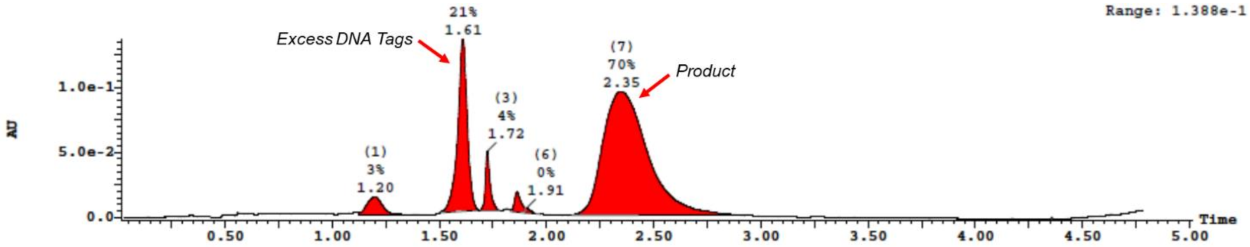


**Figure S6.** UPLC-MS analysis of pooled and buffer-exchanged library material after triazine addition and cycle 2 amine substitutions.

**Scheme S15.** Cycle 3 tag ligations for encoding of final triazine substitutions.

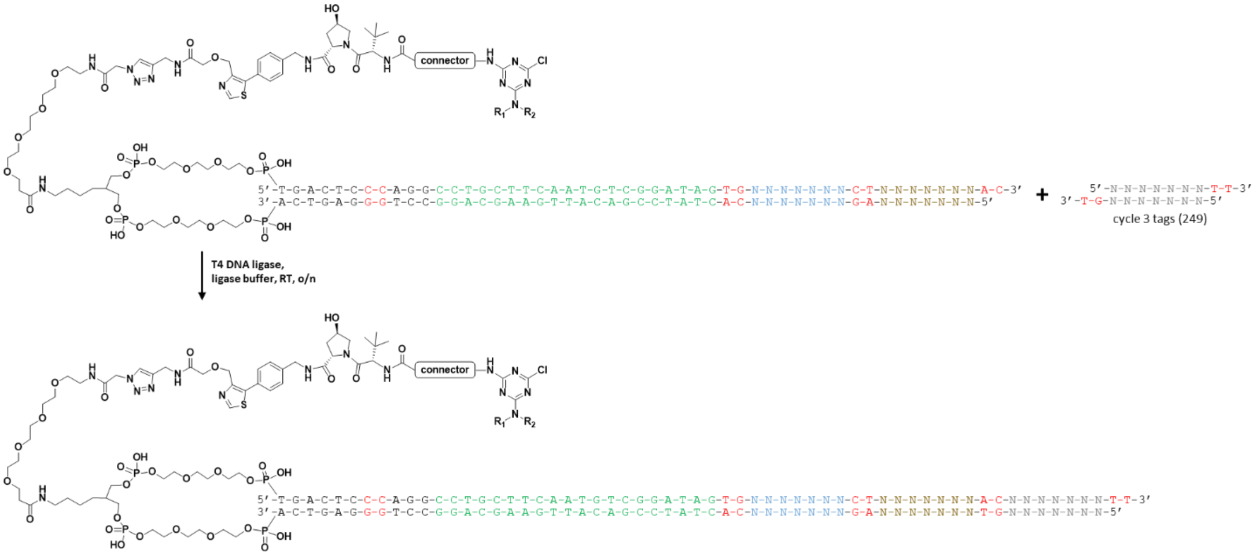

***Ligation of cycle 3 tags for encoding of 249 final triazine substitutions:*** The pooled material from cycle 2 was split across 249 wells (1.2 nmol / well in 5 µL in d$H_2O$, 1 equiv). A unique cycle 3 tag (2.6 µL, 1 mM solution in d$H_2O$, 2.6 nmol, 2.2 equiv) was added to each well, followed by a freshly prepared solution of T4 DNA ligase in 5x ligase buffer (2.4 µL, 1.4 Weiss U / well) to give a final volume of 10 µL per well. The plates were briefly spun down and incubated at RT overnight. Reaction completion was assessed by 4% agarose gel analysis of analytical samples from the diagonal wells of each plate (**Figure S7**). All ligations were deemed complete and were ethanol precipitated with the addition of 2.2 µL of 5 M NaCl and 45 µL of ice-cold ethanol. The plates were incubated at -20°C for 10 h and the precipitate was isolated by centrifugation. The supernatant was carefully decanted and the pellets were washed with an additional 100 µL of ice-cold ethanol and air-dried for 1 h. Each pellet was dissolved in 150 mM borate buffer pH 9.4 (12.6 µL) and taken on directly for the final $S_NAr$ step.

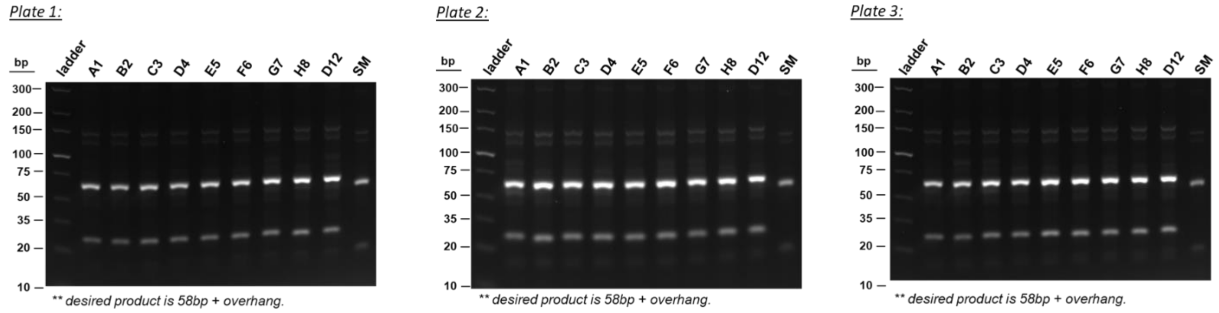


**Figure S7.** Agarose gel analysis of cycle 3 tag ligations across diagonal wells of each 96-well plate.

**Scheme S16.** Final triazine substitution with 249 amines.

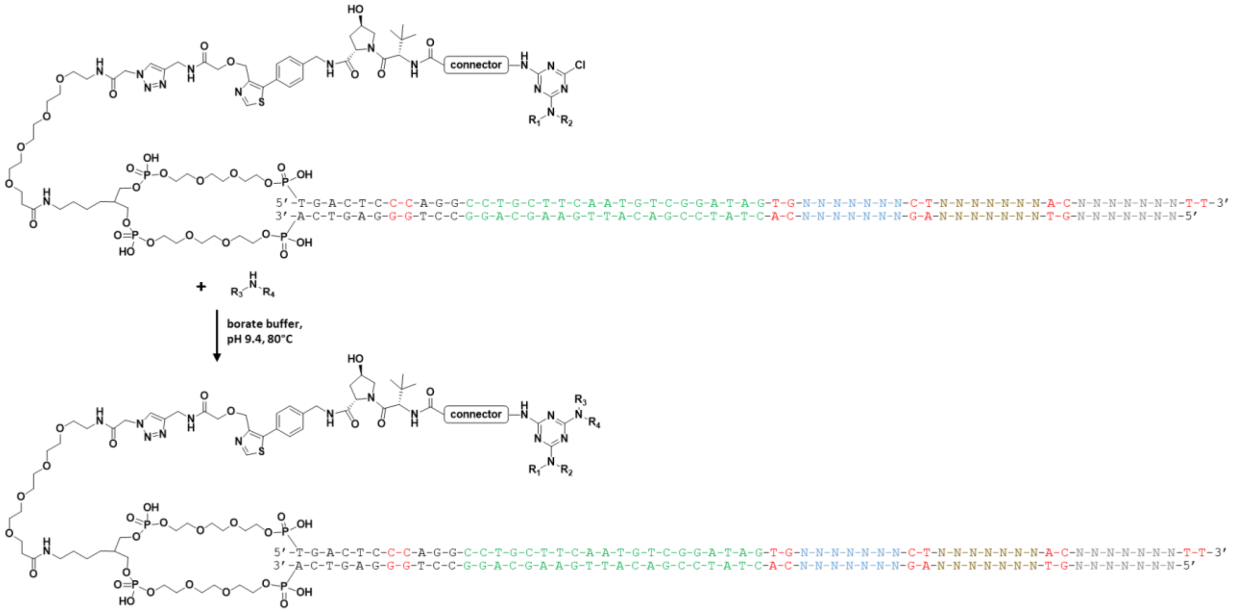


***Final substitution of triazine with 249 cycle 3 amines:*** To each of the 249 wells from the above cycle 3 ligations (12.6 µL in 150 mM borate buffer pH 9.4, 1.2 nmol, 1 equiv) was added one of 249 amines (200 mM in 1:1 $CH_3CN$:$dH_2O$, 2.4 µL, 480 nmol, 400 equiv). The reaction plates were sealed, heated to 80°C for 16 h, and then precipitated with the addition of 2.5 µL of 5 M NaCl and 100 µL of ice-cold ethanol. The plates were incubated at -20°C for 2 h and the precipitate was isolated by centrifugation. The supernatant was carefully decanted and the pellets were washed with an additional 100 µL of ice-cold ethanol and air-dried for 45 mins. The pellets were dissolved in $dH_2O$, pooled, and transferred to an Amicon Ultra 10 kDa spin column and buffer exchanged with ~10 volumes of $dH_2O$. The pooled library material was analyzed by UPLC-MS (**Figure S8**) and quantified spectrophotometrically (NanoDrop): 17.08 µg/µL (A260/280= 1.83; A260/230= 1.97); volume = 840 µL.

***UV:***

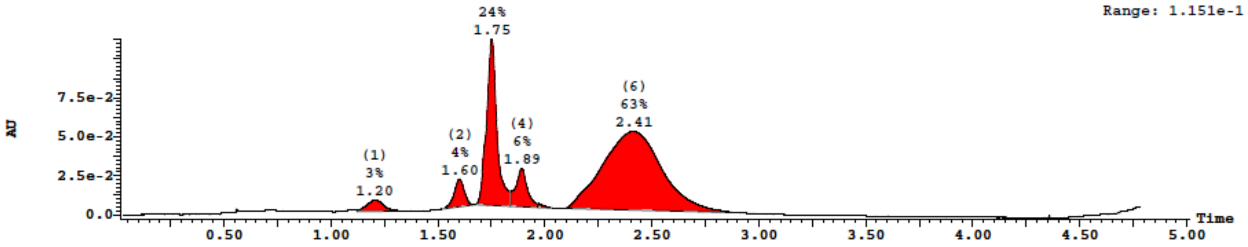


**Figure S8.** UPLC-MS analysis of pooled and buffer-exchanged library material after cycle 3 amine substitutions of triazine.

**Scheme S17.** CIP-DEL closing tag ligation.

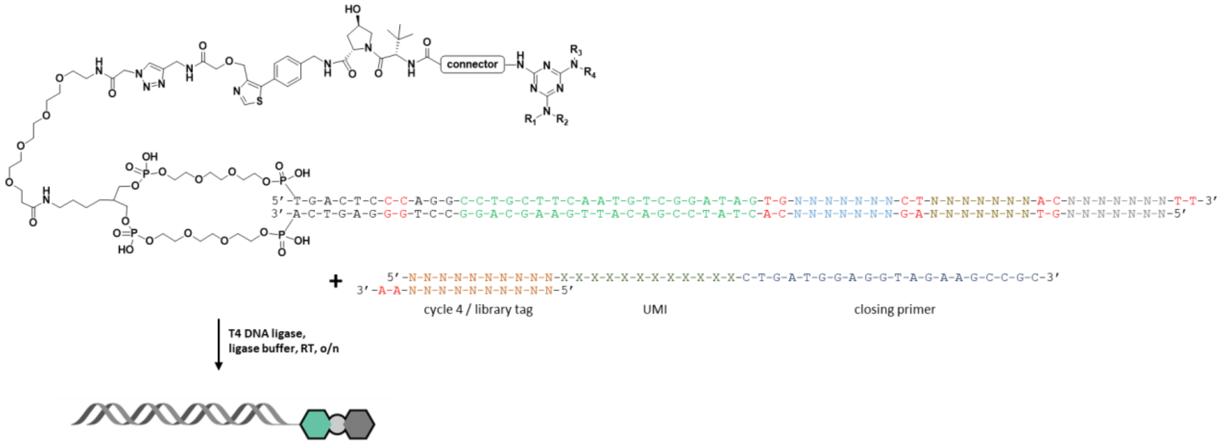


***Closing tag ligation:*** The CIP-DEL library synthesis was completed with the addition of a closing DNA segment containing a double-stranded 10 bp library tag, a single-stranded 12 nucleotide unique molecular identifier (UMI), and a single-stranded 20 nucleotide closing primer binding site. Approximately 65 nmol (182 µL, ~0.357 mM in $dH_2O$) of the crude library was closed by adding the closing tag (library tag: top 5' – CGTCAACGGT – 3') (118 µL, 118 nmol, 1 mM solution in $dH_2O$, 1.8 equiv), 10x ligase buffer (97.4 µL, Lucigen), and T4 DNA ligase (2.6 µL, 30 Weiss U / µL, Lucigen). The ligation mixture was gently vortexed, spun down, and incubated at room temperature. After 16 h, ligation progress was analyzed by 4% agarose gel electrophoresis (**Figure S9.A**) and judged to be mostly complete. The ligated product band was purified by preparative agarose gel electrophoresis using a BioRad 491 Prep Cell (**Figure S10**) per the manufacturer's protocol. Briefly, a 4% agarose gel solution was prepared by heating 8 g agarose and 200 mL 1x TBE buffer. The partially cooled solution was then poured into the large gel casting tube in the Prep Cell assembly while water was circulated through the cooling core. After 1 h, the cooling water was turned off and the gel was allowed to set overnight. The gel casting tube was then installed in the Prep Cell assembly. The cooling buffer reservoir, elution buffer reservoir, and recirculation reservoir were filled with 1x TBE and the cooling buffer circulation was started. The ligation mixture (400 µL) was combined with 80 µL of 5x sample loading dye (Invitrogen, NOVEX Hi-Density TBE Sample Buffer 5x, Cat #: LC6678) and loaded on to the gel. Electrophoresis was initiated with 12 W constant power. After running for 1.25 h, the first dye band from the loading buffer had reached the bottom of the gel and an automatic fraction collector was used to collect 1.5 mL fractions at a flow rate of 0.9 mL/min. The isolated fractions were analyzed on a 4% agarose gel (**Figure S9.B**) and the product-containing fractions were pooled. The pooled solution was transferred to a 10 kDa Amicon Ultra spin column, buffer exchanged with ~10 volumes of $dH_2O$, and concentrated. The final purified CIP-DEL library was analyzed by UPLC-MS (**Figure S11**) and quantified by QuBit dsDNA BR assay (Invitrogen) per the manufacturer's instructions: 1.060 µg/µL.

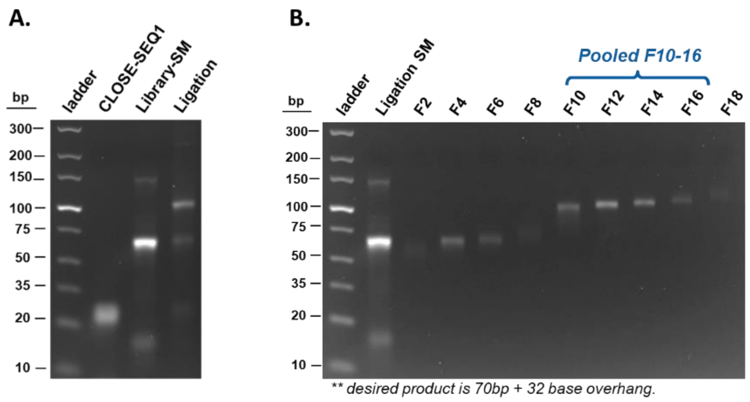


**Figure S9. (A)** Agarose gel analysis of closing tag ligation. **(B)** Analysis of purified library fractions from preparative gel electrophoresis.

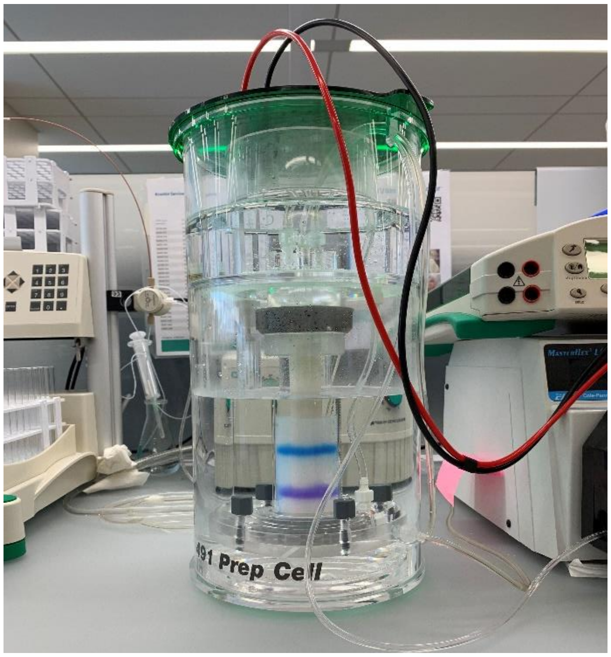


**Figure S10.** Preparative agarose gel electrophoresis of final closed CIP-DEL library using a BioRad 491 Prep Cell.

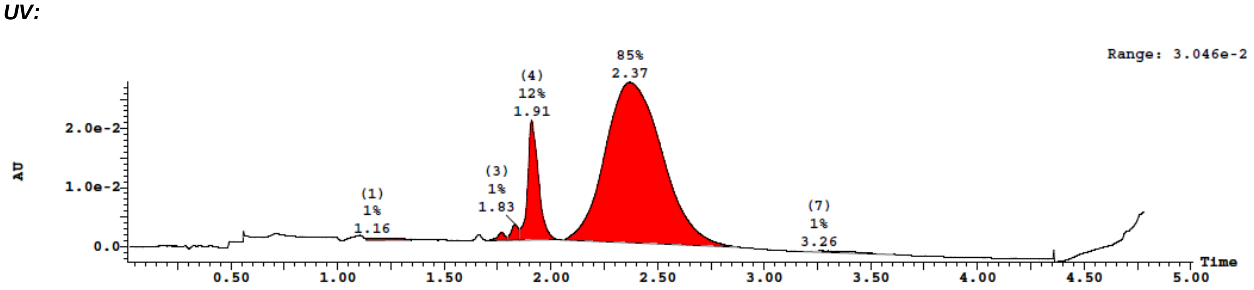


**Figure S11.** UPLC-MS analysis of final closed and purified CIP-DEL library.

#### 9. Synthesis of MZ1-DEL positive control.

**Scheme S18.** Coupling of (+)-JQ1-acid to AOP-HP with VHL ligand and **MZ1** connector.

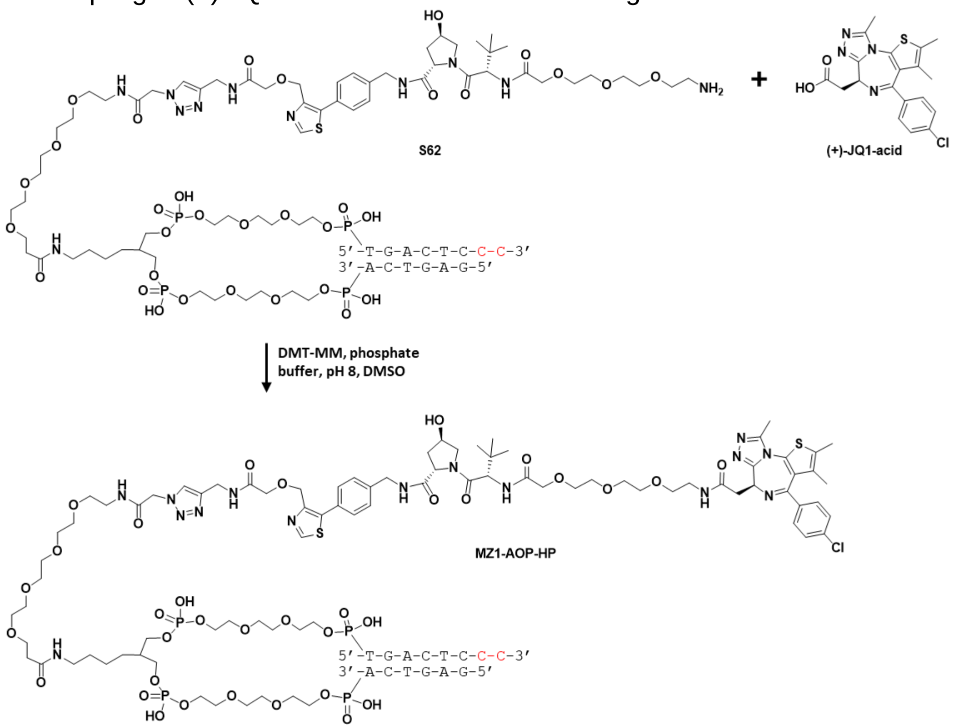


***Synthesis of MZ1-AOP-HP:*** (+)-JQ1-acid was pre-activated by combining the acid (16.0 µl, 1.60 µmol, 100 mM solution in DMSO) and DMT-MM (11.7 µl, 1.40 µmol, 120 mM solution in $dH_2O$) in a 1.5 mL microcentrifuge tube. The solution was vortexed briefly, spun down, and incubated at room temperature for 10 mins. To the activated acid was added **S62** (50.0 µl, 0.04 µmol, 0.80 mM solution in phosphate buffer pH 8) and the mixture was incubated at room temperature with shaking overnight. Analysis of the reaction by UPLC-MS showed a single major peak with product mass and the reaction mixture was precipitated with the addition of 5 M NaCl (8 µL) and ice cold EtOH (250 µL). The suspension was incubated on dry ice for 15 mins, and then spun down. The supernatant was carefully removed with a pipette and the pellet was washed with an additional 0.5 mL cold 90% EtOH. The pellet was dissolved in 80 µL $dH_2O$ and purified by preparative HPLC to obtain 16.6 nmol (NanoDrop, 42% yield) of product.

**MS** (ESI-) expected 6377 Da, found 6383 Da.

***UV:***

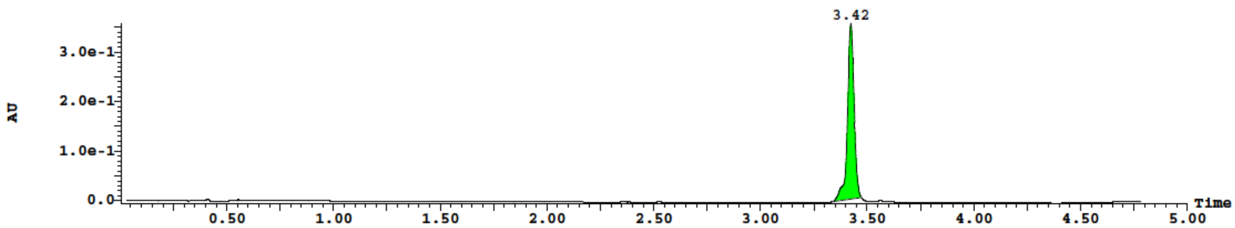

***TIC:***

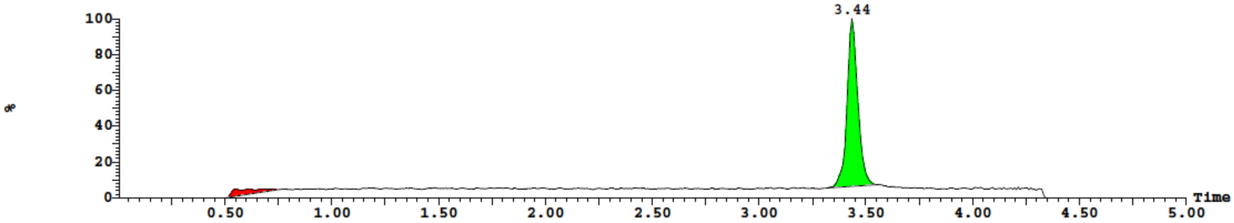


**Scheme S19.** Encoding of **MZ1** control with a single ligation of forward primer and cycle 1 to 3 DNA tags.

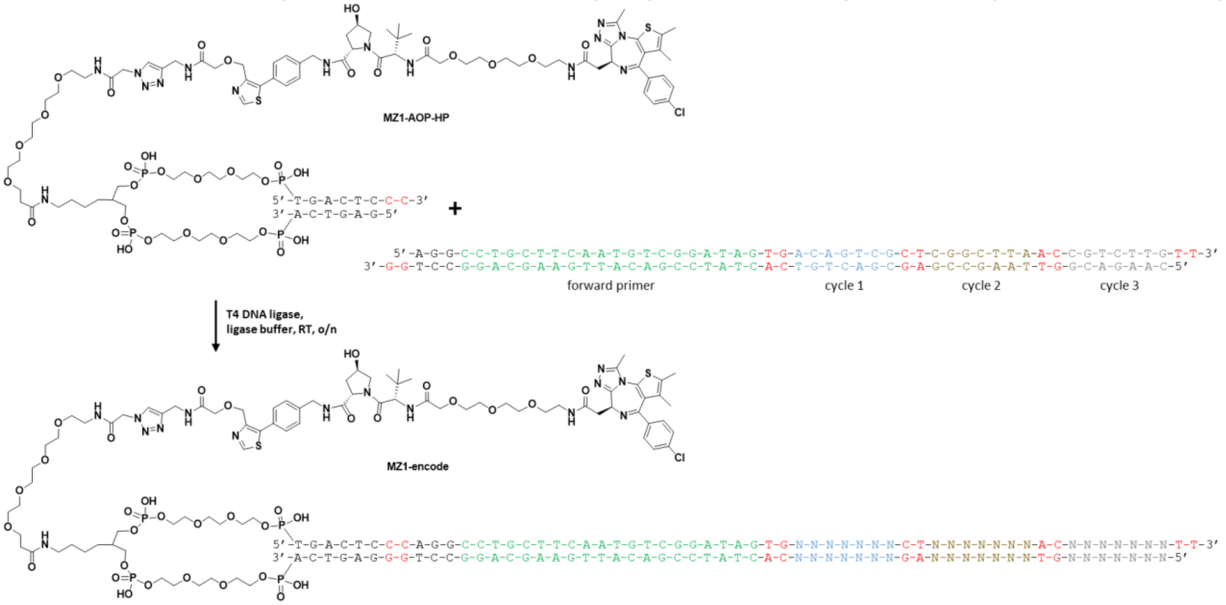


***Synthesis of MZ1-encode:*** The **MZ1-AOP-HP** conjugate was ligated to an oligo containing the entire forward primer and cycle 1, 2, and 3 encoding regions with unique sequences for the positive control. The ligation was performed by combining 25 µL of **MZ1-AOP-HP** (15.7 nmol, 0.627 mM solution in $dH_2O$, 1 equiv) with 31.4 µL **MZ1-TAG** (31.4 nmol, 2 equiv, 1 mM solution in $dH_2O$), 13 µL ligase buffer (10x), and 0.6 µL T4 DNA Ligase (30 Weiss U/µL). The reaction was gently mixed, spun down, and incubated at room temperature for 16 h. An analytical sample was analyzed by 4% agarose gel electrophoresis and indicated the reaction was complete (**Figure S12.A**). The ligase was inactivated by heating the solution to 70°C for 5 mins and the sample was purified by preparative HPLC (**Figure S12.B, C**). Purified product was analyzed by UPLC-MS (**Figure S12.D**) and quantified by NanoDrop to provide 9.3 mol (59% yield, 0.31 mM solution in $dH_2O$) of product.

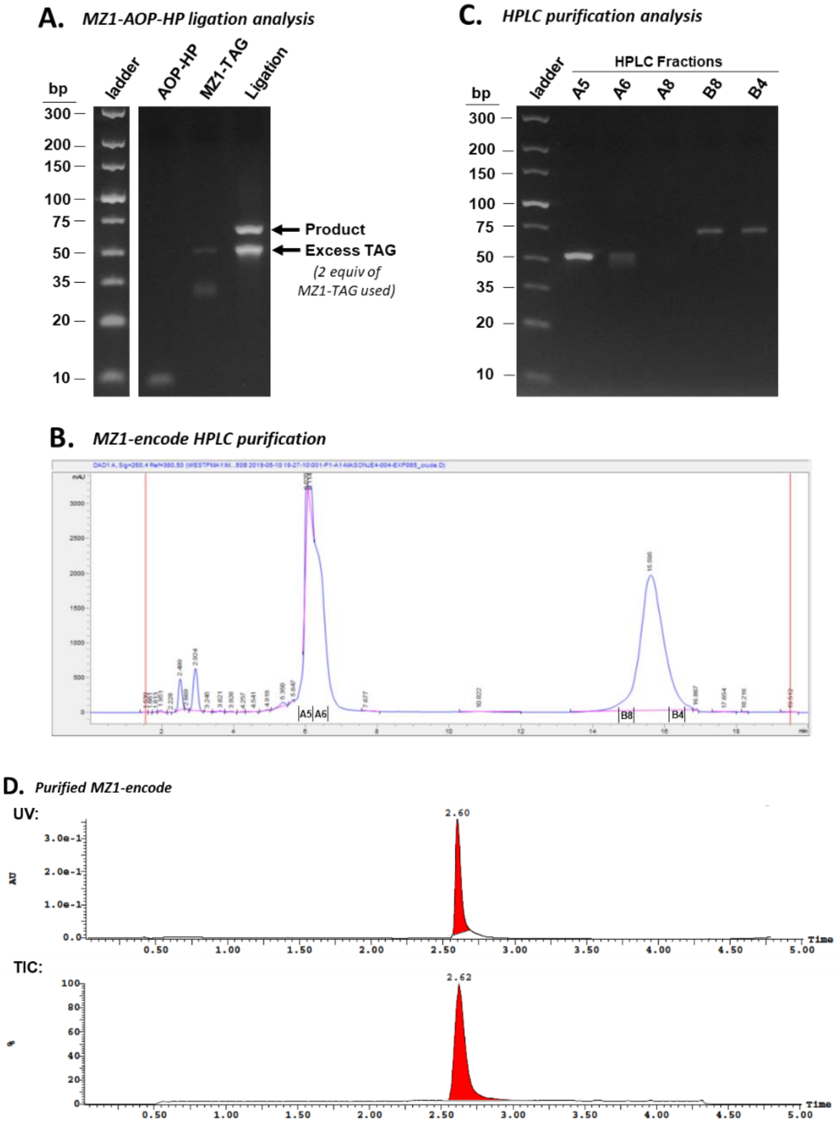


**Figure S12.** Ligation of **MZ1-TAG** encoding for positive control. **(A)** Analysis of crude ligation reaction by 4% agarose gel electrophoresis. **(B)** Preparative HPLC UV-260nm trace of sample purification. **(C)** Agarose gel analysis of HPLC fractions. **(D)** Purified **MZ1-encode** sample analysis by UPLC-MS.

**Scheme S20.** Ligation of closing tag to provide **MZ1-DEL** control.

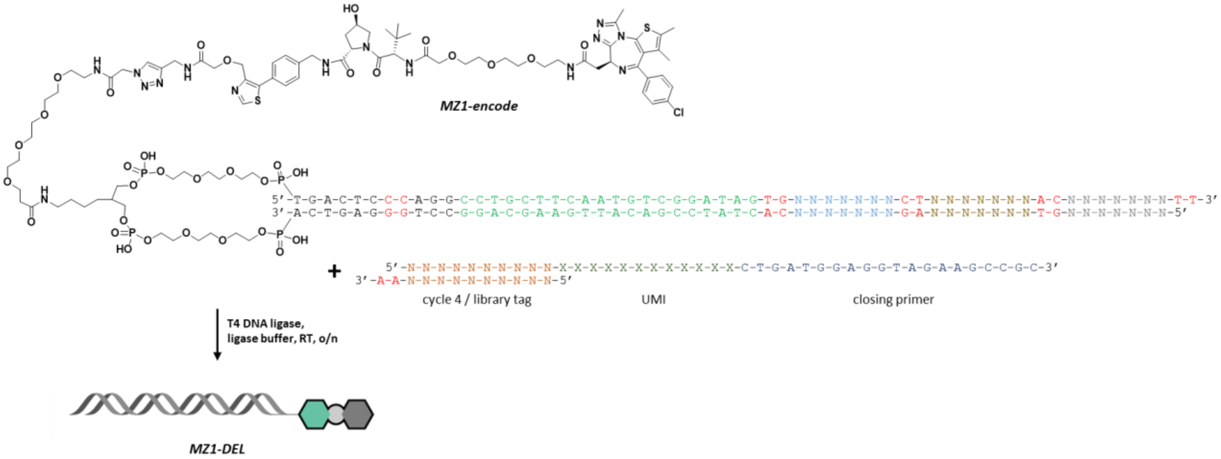


***Synthesis of MZ1-DEL:*** To a 0.5 nmol sample of purified **MZ1-encode** was added the final closing DNA sequence containing the library tag (top 5' – GCTCAACAGC – 3'; dsDNA), the UMI (ssDNA), and the reverse primer-binding site (ssDNA). In a 1.5 mL microcentrifuge tube was combined 5.3 µL $dH_2O$, 1.6 µL **MZ1-encode** (0.5 nmol, 0.31 mM in $dH_2O$, 1 equiv), 0.9 µL closing tag (0.9 nmol, 1 mM solution in dH2O, 1.8 equiv), 2.0 µL ligase buffer (10x), and 0.2 µL T4 DNA ligase (30 Weiss U/µL). The ligation mixture was gently mixed, spun down, and incubated at room temperature for 16 h. Analysis of the reaction was done by 4% agarose gel electrophoresis and the ligation was judged complete (**Figure S13.A**). The ligation mixture was diluted to 220 µL and heated to 70°C for 5 mins to inactivate the ligase. After cooling to room temperature, the crude mixture was loaded across all wells of a 4% agarose gel and the product band was separated by electrophoresis. The gel band containing product was excised and DNA was isolated by gel extraction (Qiagen). Purified **MZ1-DEL** was analyzed by 4% agarose gel (**Figure S13.B**) and quantified by QuBit dsDNA BR assay (Invitrogen) per the manufacturer's instructions: 65.6 ng/µL, 0.124 nmol, 1.19 µM solution in dH2O (25% yield).

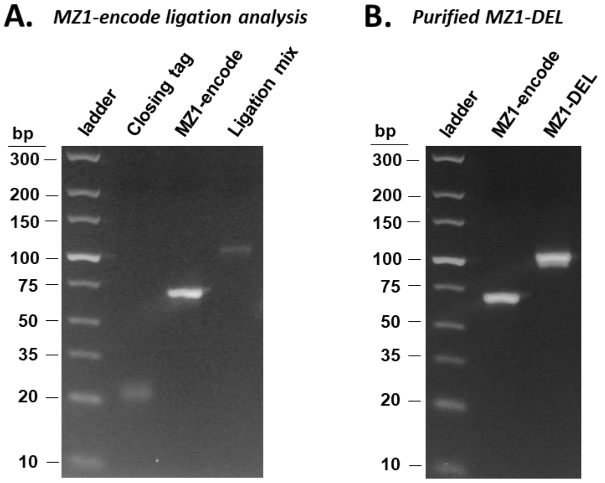


**Figure S13.** Addition of closing tag to **MZ1-encode** to give **MZ1-DEL**. **(A)** Analysis of ligation reaction after 16 h. **(B)** Final **MZ1-DEL** after agarose gel purification.

#### 10. CIP-DEL selections.

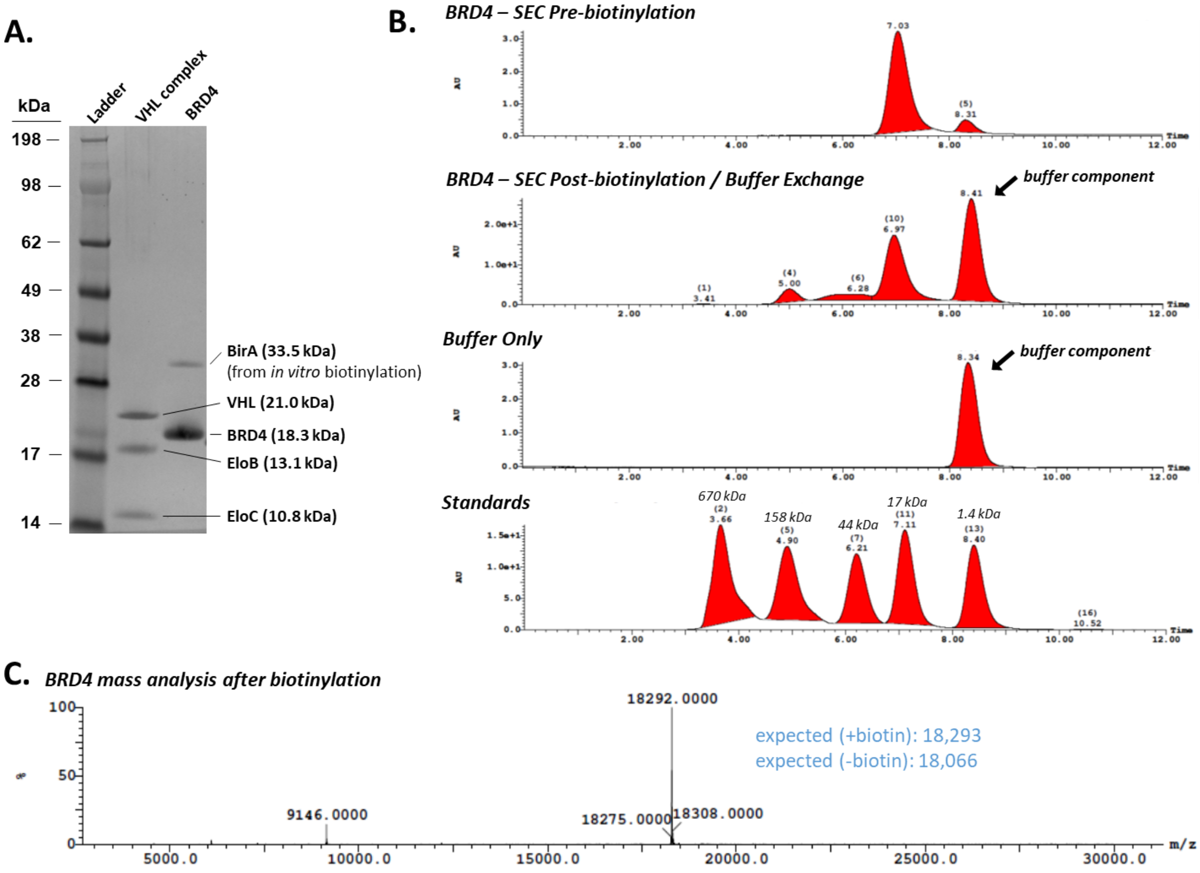


**Figure S14. (A)** SDS-PAGE analysis of VHL complex (VHL + Elongin B + Elongin C) and BRD4$^{BD1}$ protein used in CIP-DEL selection experiments. **(B)** Analytical size-exclusion analysis of BRD4$^{BD1}$ protein before and after *in vitro* biotinylation. **(C)** TOF-MS analysis of biotinylated BRD4$^{BD1}$ protein showing expected m/z for biotinylated protein and no visible m/z for non-biotinylated protein.

**Proteins**. The first bromodomain of BRD4 (N44 – E168) was synthesized in codon optimized form by GeneArt synthesis services (Thermo Fisher) with an N-terminal His-tag and C-terminal Avi-tag and cloned in to a pDEST expression plasmid. The protein was expressed in *E. coli* BL21(DE3) as previously described,[35] concentrated to 5.3 mg/mL in 25 mM HEPES pH 7.5, 150 mM NaCl, 10% glycerol, and stored at -80°C prior to use. For use in CIP-DEL selections, the BRD4 protein was first biotinylated *in vitro* using the BirA500 biotinylation kit (Avidity LLC) per the manufacturer's instructions. The biotinylation reaction was incubated at room temperature for 30 mins, then 4°C overnight. The reaction mixture was transferred to an Amicon Ultra 4 kDa centrifugal filter and buffer exchanged with ~10 volumes of selection buffer (25 mM HEPES pH 7.5, 150 mM NaCl, 0.05% Tween-20). Final protein quantification was performed using the Pierce 660nm protein assay with Pierce bovine gamma globulin standard set (Thermo Fisher). BRD4 biotinylation was confirmed by mass spectrometry using a Xevo G2-SX TOF LC-MS instrument (Water) (**Figure S14.C**) and monodispersed protein was confirmed by analytical size-exclusion chromatography (SEC) pre- and post-biotinylation (**Figure S14.B**). Final protein purity was assessed by SDS-PAGE, which showed a primary band of expected mass for BRD4 and a minor band corresponding to the BirA enzyme used for *in vitro* biotinylation (**Figure S14.A**). Although the BirA enzyme was not removed prior to performing selections, the first step of the selection experiment was to immobilize BRD4 to streptavidin-coated beads and wash away unbound protein (ie. BirA, along with excess BRD4).

VHL (M54 – D213), elonginB (M1 – Q118), and elonginC (M17 – C112) (VBC complex) were co-expressed in *E. coli* BL21(DE3) with an N-terminal 8x-His tag on VHL and an HRV 3C protease site for tag removal. Protein expression was performed as described previously[36] and the final protein with His-tag removed was concentrated to 3.22 mg/mL in 25 mM Tris pH 7.5, 150 mM NaCl, 1 mM TCEP and stored at -80°C until use.

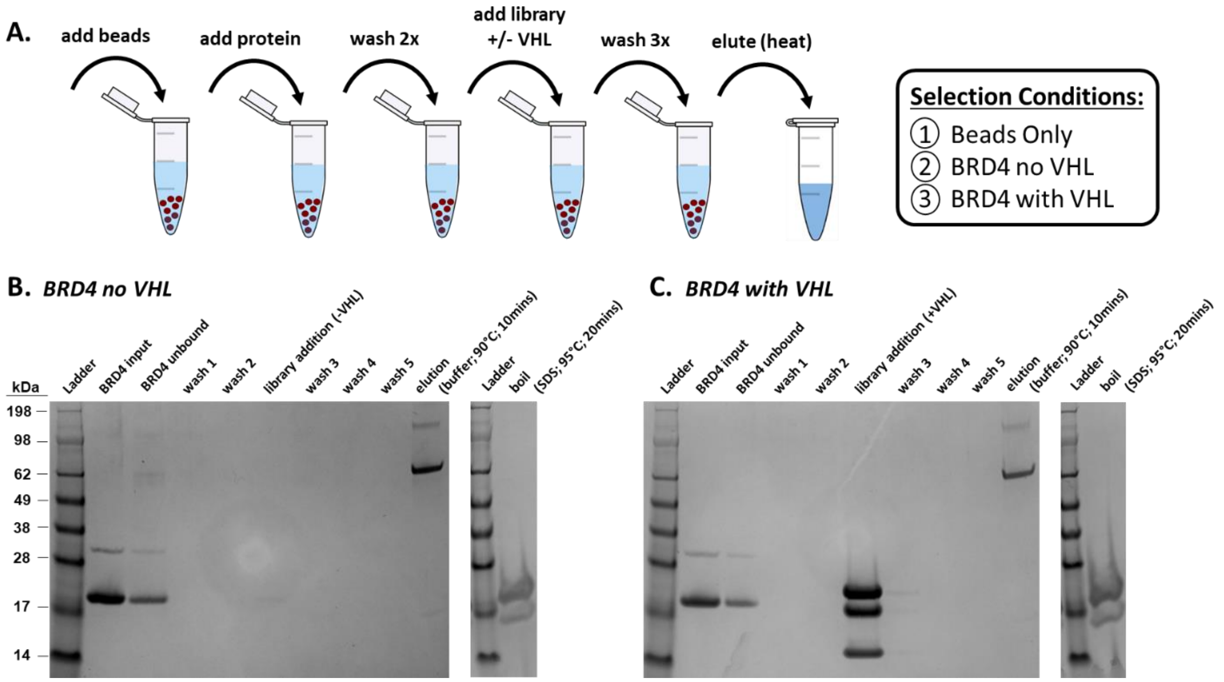


**Figure S15. (A)** Overview of the selection workflow. Three selections were performed in parallel: beads only control, $BRD4^{BD1}$ without VHL, and $BRD4^{BD1}$ with VHL pre-incubated with the CIP-DEL library. **(B) & (C)** The presence of BRD4 and VHL protein complex was monitored after each step of the selection experiment by SDS-PAGE. Beads were first incubated with excess BRD4 (lanes labelled “BRD4 input”) and unbound protein was removed (lanes labelled “BRD4 unbound”). Beads were then washed twice (lanes labelled “wash 1 and 2”) and the CIP-DEL library was added to the beads in selection buffer with and without pre-incubation with the VHL protein complex. The library was incubated with the beads for 1 h at room temperature and the solution was removed (lanes labelled “library addition”). The beads were then washed three times (lanes labelled “wash 3, 4, and 5”). The beads were resuspended in BE buffer and library members retained during the selection were eluted with heat (“elution; buffer; 90°C; 10 mins”). To ensure BRD4 was retained on the beads throughout the selection, the beads were resuspended in SDS loading buffer and boiled to remove biotinylation $BRD4^{BD1}$ (“boil; SDS; 95°C; 20 mins”).

**Buffers.**

***BE Buffer (Bind-Elute):***
25mM HEPES pH 7.4, 150mM NaCl, 0.05% Tween-20

***S Buffer (Selection):***
25mM HEPES pH 7.4, 150mM NaCl, 0.05% Tween-20, 0.1mg/mL ssDNA (Aldrich: D1626-1G), 20mM imidazole

**Selections.** Three aliquots of the CIP-DEL library were prepared for each of the selections performed in parallel: 1) beads only control, 2) BRD4 (-) VHL, and 3) BRD4 (+) VHL. We targeted the addition of ~7 million copies of each library compound per selection. Given the library size of 1,083,150 compounds, we added 12.7 pmol of CIP-DEL library to each selection experiment. In addition, our positive control compound MZ1-DEL was spiked into each library aliquot with equivalent representation (i.e. ~7 million copies of this single control compound). A master stock of CIP-DEL library with MZ1-DEL was diluted in S Buffer by combining 3.2 µL of CIP-DEL library (19.9 µM solution in dH2O), 7.0 µL of MZ1-DEL control compound (11.9 pM solution in dH2O, obtained by serial dilution), and 240 µL of S buffer. Three 50 µL aliquots of this pool were transferred to 1.5 mL microcentrifuge tubes for each selection. To the BRD4 (+) VHL aliquot was added 14.5 µL of VHL protein complex (75 µM solution in S buffer) and the beads only and BRD4 (-) VHL aliquots received an equal volume of S buffer without protein. Each sample was incubated at room temperature for 30 mins.

Meanwhile, 50 µL of beads (DynaBeads MyOne Streptavidin C1, Invitrogen, P/N: 65001) was transferred to three separate 1.5 mL microcentrifuge tubes for each selection experiment. The samples were placed on a magnetic rack and the solution was removed from the beads. To the beads only control sample was added 50 µL BE buffer, whereas the BRD4 (-) VHL and BRD4 (+) VHL samples received 50 µL of biotinylated $BRD4^{BD1}$ protein stock solution prepared by combining 98.1 µL BE buffer and 11.9 µL of $BRD4^{BD1}$ (1.86 mg/mL solution in BE buffer, 10 µg $BRD4^{BD1}$ protein per sample). The samples were incubated at room temperature for 20 mins with gentle mixing and then placed on a magnet. The supernatant was removed and the resin was washed once with 200 µL BE buffer (wash 1) and once with 200 µL S buffer (wash 2). Following removal of the wash buffer, the library solutions prepared above (64.5 µL) were added to their respective bead sample and incubated at room temperature for 1 h with gentle mixing to prevent the beads from settling.

Each sample was returned to the magnetic rack and the resin was washed with 200 µL S buffer (wash 3) then 2 x 200 µL BE buffer (wash 4 & 5), switching microcentrifuge tubes after the final wash to avoid contamination. Finally, 50 µL of BE buffer was added to each tube and the solutions were heated to 90℃ for 10 min to elute bound library members. Immediately after heating, the tubes were placed on a magnet and the supernatants were transferred to new 1.5 mL microcentrifuge tubes. To confirm that BRD4 protein was successfully immobilized on the streptavidin beads, the BRD4 bead samples were resuspended in 20 µL of SDS loading buffer and 1 µL of reducing agent and the samples were heated to 95°C for 20 min. The tubes were transferred to a magnetic rack and the supernatants were collected in new 1.5 mL microcentrifuge tubes. All selection supernatants were saved and stored at -80°C for further analysis. Later, an aliquot of each supernatant was run on a 4 – 12% NuPAGE SDS gel to track protein loadings throughout the selection experiments (**Figure S15.B & C**).

**Post Selection DNA Processing.** DNA isolated from the three selection experiments was cleaved from the AOP-HP and small molecule components by restriction digest with StuI. To the 50 µL sample of screening output from each selection experiment was added 6 µL of 10x CutSmart buffer (New England Biolabs) and 1 µL of StuI (NEB, P/N: R0187) and each reaction mixture was heated to 37°C with gentle mixing for 3 h. The cleaved DNA was then PCR amplified directly by combining 15 µL of the restriction digest reaction mixture with 8 µL nuclease-free water, 1 µL of i5 index primer (10 µM, 5'-AATGATACGG CGACCACCGA GATCTACACX XXXXXXXCTT AGCTCCCAGC GACCTGCTTC AATGTCGGAT AGTG-3'), 1 µL i7 index primer (10 µM, 5'-CAAGCAGAAG ACGGCATACG AGATXXXXXX XXACCATGCT CACTGCGGCT TCTACCTCCA TCAG-3'), and 25 µl of 2x master mix (Invitrogen Platinum™ Hot Start PCR Master Mix, P/N: 13000012). The indexing sequences (indicated with X's above) were as follows: beads only - i5: GTAAGGAG; i7: AGGAGTCC; BRD4 (-) VHL- i5: ACTGCATA; i7: AGGAGTCC; BRD4 (+) VHL- i5: GTAAGGAG; i7: CATGCCTA. Following PCR amplification (95°C for 2 min; then [22 cycles of 95°C (15 sec), 55°C (15 sec), 72°C (30 sec)]; then 72°C for 7 min; then hold at 4°C), products were purified on a 4% agarose gel and the library band was isolated by gel extraction (Qiagen, P/N: 28704). Purified DNA was quantified by qPCR using Illumina P5 and P7 flowcell adapters as primers and Power Sybr Green PCR Master Mix (Applied Biosystems, P/N: 4367660) on an Applied Biosystems ViiA 7 system. The three selection outputs were evenly pooled (along with additional unrelated samples) based on the $C_T$ values obtained from qPCR and the resulting pool was submitted for Illumina NGS sequencing using a MiSeq v2 50 cycle reagent kit (P/N: MS-102-2001). Custom primers were used for CIP-DEL barcode sequencing (5'-CTTAGCTCCC AGCGACCTGC TTCAATGTCG GATAGTG-3') and i7 indexing (5'-CTGATGGAGG TAGAAGCCGC AGTGAGCATG GT-3').

#### 11. NGS data processing and analysis.

Reads observed during sequencing were processed into counts for each barcoded small molecule through a custom Python script. FastQ files were read using the BioPython package and used to construct a sparse mapping describing the number of reads for each unique sequence. Fifty nucleotide-long reads were split into five regions corresponding to the cycle 1 tag, cycle 2 tag, cycle 3 tag, library tag, and unique molecular identifier (UMI). The barcode region (comprising the three cycle tags and the library tag) was used to aggregate the reads corresponding to each small molecule. To account for errors during PCR amplification and sequencing, up to two errors in the library tag and up to one error in the cycle 1 tag were allowed as measured by the Hamming distance between the expected and observed tags (no errors were allowed in cycle 2 and 3 tags). This error tolerance provided a marginal increase in the number of matched reads while still allowing unambiguous assignment of an observed sequence to a single expected barcode at that position. UMIs were used to deduplicate reads by constructing a directional graph that accounts for errors during PCR and sequencing, using the umi_tools package by Smith et al.[37] The number of unique UMI clusters corresponding to each barcode corresponds to that barcode's (integer) count.

Enrichment values were estimated by comparing count values across two distinct experimental conditions (i.e., a protein-containing selection versus the beads-only control), using a recently proposed Poisson enrichment metric.[38] We assume that each barcoded small molecule exists at a fixed but unknown concentration before PCR and NGS, $\lambda$. We also make the simplifying assumption that PCR amplification does not change the relative abundance of barcodes and that the stochastic process of NGS can be modeled using Poisson statistics. Poisson distributions are characterized by a single parameter $\lambda$ that in this setting describes the relative abundance of a particular barcode compared to the total abundance of all other barcodes in the sample. If the (unknown) relative abundance of one compound following one experimental condition is $\lambda_1$ and following a second experimental condition is $\lambda_2$ (e.g., where the second experiment is a beads-only control), we define the enrichment ratio as $R = \lambda_1/\lambda_2$. Defining $k_1$ and $k_2$ as the integer numbers of counts for the particular barcode and $n_1$ and $n_2$ as the total integer counts for that experimental condition, naive estimates of $\lambda_1$ and $\lambda_2$ are simply $k_1/n_1$ and $k_2/n_2$, and hence a naïve estimate of $R$ is $(k_1/n_1)/(k_2/n_2)$. However, this ratio does not consider uncertainty in the observed values of $k$, which is necessarily an integer and typically close to zero using standard sequencing depths; further, it fails to produce a finite estimate of the enrichment ratio when $k_2 = 0$.

As an uncertainty-aware enrichment metric, we directly estimate the value of $R = \lambda_1/\lambda_2$ and its confidence interval rather than the values of $\lambda_1$ and $\lambda_2$ separately. We therefore perform a Poisson ratio test that formally seeks to evaluate a hypothesis test about the value of the ratio of two Poisson metrics based on the observed integer counts and total exposures (total counts)[39]:

$$H_0: \frac{\lambda_1}{\lambda_2} \leq R_0 \qquad H_1: \frac{\lambda_1}{\lambda_2} > R_0$$

Using a variance stabilizing square root transformation, we calculate a z-score as:

$$z = 2\frac{\left(k_1 + \frac{3}{8}\right)^{1/2} - \left(k_2 + \frac{3}{8}\right)^{\frac{1}{2}}\left(\frac{n_1}{n_2}R_0\right)^{1/2}}{\left(1 + \frac{n_1}{n_2}R_0\right)^{1/2}}$$

As we expect this score to follow a standard normal distribution, we can calculate a p-value for different values of $R_0$. To estimate the lower and upper bounds for the enrichment ratio, we therefore invert this hypothesis test and calculate the values of $R_0$ corresponding to $z = \pm 2$. This procedure can be applied to the count data for individual compounds (single barcodes), as well as groups of compounds (sets of barcodes, e.g., those sharing a particular building block) as done for the linker analysis to identify **C15** as strongly enriched.

For analysis of our screening data, we use the lower bound (lb) of the 95% confidence interval as our enrichment metric since this provides a more conservative estimate of enrichment that may reduce the number of false positives. In effect, this approach accounts for the uncertainty present when barcode counts are low: if a barcode is observed three times in a screen and once in the beads-only control, we obtain a 3-fold enrichment but with low certainty; if a barcode is observed 150 times in the screen and 50 times in the beads-only control, we obtain a 3-fold enrichment with higher certainty.

***Note regarding DNA barcode analysis:***

We profiled a common set of amine building blocks for both cycle 2 and cycle 3 triazine substitutions using the model reaction systems described in section 7 (scheme S8 and S9). A cutoff of 75% UPLC-MS product yield was used for including amine building blocks in the CIP-DEL library. Since the cutoff was applied independently for cycle 2 and cycle 3 profiling, some amines met the inclusion criteria for cycle 2, some amines met the inclusion criteria for cycle 3, and some amines met the inclusion criteria for both cycles 2 and 3. Including a subset of identical amines in cycles 2 and 3 generated redundant library members due to symmetry of the triazine scaffold (e.g. BBx and BBy added in cycles 2 and 3 generates the same molecule as BBx and BBy added in cycles 3 and 2, respectively).

A total of 218 amines (121 primary amines and 97 secondary amines) were included in both cycle 2 and 3 substitutions, producing 354,795 duplicate library members (with distinct DNA barcodes) and 728,355 unique library members. For analysis of duplicate library members, we selected the component with the greatest enrichment value in the BRD4 (+) VHL selection condition. Enrichment values for the BRD4 (-) VHL selection were then matched with the selected library member. This analysis is based on the assumption that higher enrichment values correspond to library members with greater synthetic yields and therefore, the more representative enrichment values. We note however that variability was observed in the enrichment values between identical library members encoded with distinct DNA barcodes and further studies on the implications of redundant DNA encoding are needed.

#### 12. VHL TR-FRET binding assay.

Binding competition reactions were created with a mixture of 10 nM purified recombinant VHL protein complex consisting of N-terminally Avi-tagged and biotinylated VHL, untagged Elongin-B and untagged Elongin-C, 2 nM terbium-conjugated Streptavidin (ThermoFisher P/N: PV3966) and 24 nM FITC-peptide probe: (bA)-DEALA(HyP)YIPD-OH (New England Peptide) in binding assay buffer (100 mM NaCl, 50 mM Tris pH 7.5, 0.1% Pluronic F127, 1 mM DTT). The mixture was dispensed into 1536-well plates (Greiner P/N: 782075) with 5 µL / well using a tip-based dispenser (GNF Systems One Tip Dispenser). 50 nL of CIP compounds in DMSO was added to target wells using an acoustic liquid handler (Labcyte Echo 500). The plates were incubated at RT for 60 minutes, and then read on a plate reader (Perkin Elmer Envision) using a TR-FRET protocol (terbium donor exc: 340 / 80 nm, ems: 492 / 7.5 nm; FITC acceptor ems: 520 / 7.5 nm). The TR-FRET signal was calculated as the ratio: signal 520 nm / signal 492 nm. $AC_{50}$ values were calculated from curve fits generated by Spotfire analysis software (Tibco).

#### 13. SPR experimental details.

*Binary complexes*: His-BRD4(44-168)-avi was immobilized on a streptavidin chip using a Biacore 8K (GE Healthcare). To determine the kinetic affinity of the small molecules, compounds were diluted in DMSO such that final concentration gradients in DMSO were 50x. 3.1 µL of the small molecules were then added to 150 µL of buffer (50 mM Tris pH 8.0, 150 mM NaCl, 0.01% Tween 20, 1 mM EDTA, 1 mM DTT) in a 384-deep well plate (Greiner Bio-One P/N: 78127) and mixed using a Biomek FX. The small molecule solution gradients (6 concentrations / 2-fold dilutions) were then injected at 45 µL / min for 120 seconds of contact time followed by 1200 seconds of dissociation time in running buffer (50 mM Tris pH 8.0, 150 mM NaCl, 0.01% Tween 20, 1 mM EDTA, 1 mM DTT, 2% DMSO). The single-cycle kinetics data was fit to a 1:1 binding model or steady-state model to measure the association rate $k_a$ (1 / M·s), the dissociation rate $k_d$ (1 / s), and the affinity $K_D$ (M).

*Ternary complexes*: His-BRD4(44-168)-avi was immobilized on a streptavidin chip using a Biacore 8K (GE Healthcare). Compounds were diluted in DMSO to give 50x final concentration gradients. To determine the apparent kinetic affinity for ternary complex formation for small molecules with immobilized BRD4 and soluble untagged VHL complex (VHL, Elongin-B, and Elongin-C), 3.1 µL of the small molecules were added to 150 µL of buffer (50 mM Tris pH 8.0, 150 mM NaCl, 0.01% Tween 20, 1 mM EDTA, 1 mM DTT) containing 2 or 4 µM VHL complex in a 384-deep well plate (Greiner Bio-One P/N: 78127) and mixed using a Biomek FX. The protein / small molecule solution gradients (6 compound concentrations / 2-fold dilutions) were then injected at 45 µL / min for 120 seconds of contact time followed by a dissociation time of 1200 seconds in running buffer (50 mM Tris pH 8.0, 150 mM NaCl, 0.01% Tween 20, 1 mM EDTA, 1 mM DTT, 2% DMSO). The single-cycle kinetics data was fit to a 1:1 binding model or steady-state model to measure the association rate $k_a$ (1 / M·s), the dissociation rate $k_d$ (1 / s), and the affinity $K_D$ (M).

*Data Analysis:* A detailed report of all kinetic and steady-state binding parameters for CIP compounds, along with calculated α-values, is provided in **Extended Data Table 1**. To simplify subsequent analyses, we averaged all experimentally determined $K_D$ values for each compound and used this data (shown in **Table 1**) to look for correlations across our data sets.

#### 14. NanoBit cellular ternary complex complementation assay.

HEK293 cells were transfected by plating 15 mL of cells at 0.625 x $10^6$ cells / mL in complete media (DMEM with 10% FBS) into a 75 $cm^2$ flask. 90 µL of Fugene HD (Promega P/N: E2312) was combined with 1500 µL of OptiMEM I (Gibco P/N: 31985-062) and added to 30 µg of total DNA (15 µg of HSVTK-driven VHL-LgBiT plasmid and 15 µg of CMV-driven SmBiT-$BRD4^{BD1}$ plasmid). The transfection mixture was incubated for 15 minutes at RT before being added dropwise to the plated cells. After 24 h of incubation in a cell culture incubator (37˚C, 5% $CO_2$), the transfected cells were trypsinized, normalized to 0.5 x $10^6$ cells / mL in complete media and plated into 1536-well plates, 5 µL / well using a tip-based dispenser (GNF Systems One Tip Dispenser). The plates were incubated in a cell culture incubator (37˚C, 5% $CO_2$) for 5 h. Next, 5 nL of 1 mM MLN4924 in DMSO was added to every well using an acoustic liquid handler (Labcyte Echo 500), and the plates were returned to the incubator for an additional 17 h. Next, 10 nL of CIP compounds in 100% DMSO, or DMSO alone as a control, were dispensed into target wells using the Echo, and the plates were returned to the incubator for another 2 h. The plates were allowed to cool to RT for 10 minutes, and 5 µL of NanoBiT NanoGlo reagent, consisting of NanoGlo subtrate (Promega P/N: N2058) diluted 1:25 in a mixture of 50% NanoGlo LCS buffer (Promega P/N: N206C) and 50% PBS, was dispensed into each well using the GNF dispenser. After a final incubation at RT for 10 minutes, the plates were read on a luminescence plate reader (Perkin Elmer ViewLux). Luminescence for each data point was normalized using the equation 100*(well value-NC)/NC, where NC is the neutral control well (DMSO). Curves were fitted using GraphPad Prism v9.0.0.

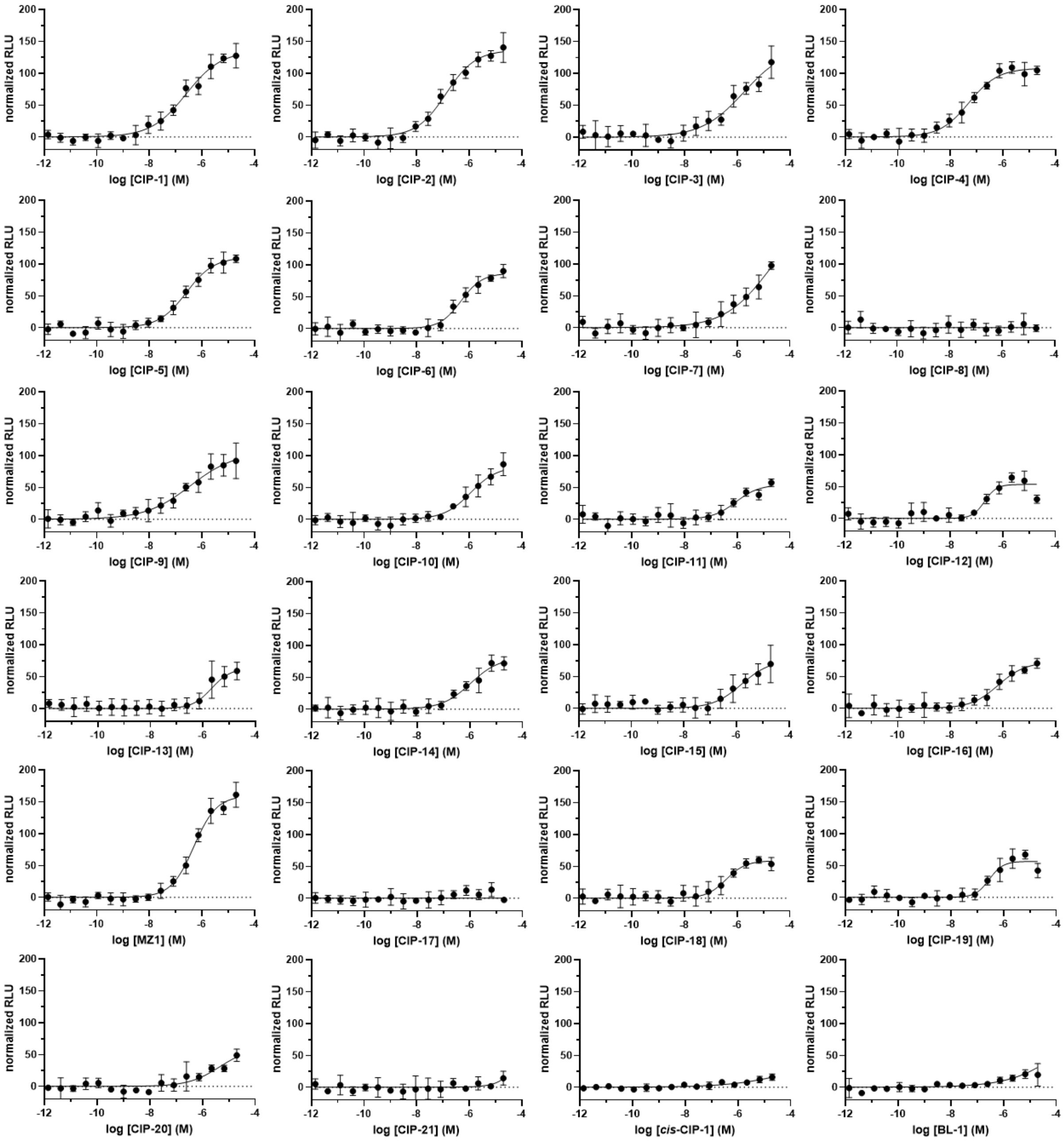


**Figure S16.** Representative curves for cell-based ternary complex formation using the NanoBit assay (n = 2 independent experiments with 2 and 4 technical replicates per experiment; 4 technical replicates shown).

#### 15. HiBit cellular protein degradation assay.

HEK293 cells were transfected by plating 15 mL of cells at 0.625 x $10^6$ cells / mL in complete media (DMEM with 10% FBS) into a 75 $cm^2$ flask. 90 µL of Fugene HD (Promega P/N: E2312) was combined with 500 µL of OptiMEM I (Gibco P/N: 31985-062) and added to 30 µg of total DNA (15 µg of CMV-driven VHL plasmid and 15 µg of HSVTK-driven $BRD4^{BD1}$-HiBiT plasmid). The transfection mixture was incubated for 15 minutes at RT before being added dropwise to the plated cells. After 24 h of incubation in a cell culture incubator (37˚C, 5% $CO_2$), the transfected cells were trypsinized, normalized to 0.5 x $10^6$ cells / mL in complete media and plated into 1536-well plates, 5 µL / well using a tip-based dispenser (GNF Systems One Tip Dispenser). Immediately after, 5 nL of either 1 mM MLN4924 in DMSO, or DMSO alone, was added to wells using an acoustic liquid handler (Labcyte Echo 500) and the plates were incubated in a cell culture incubator (37˚C, 5% $CO_2$) for 5 h. Next, 10 nL of CIP compounds in 100% DMSO, or DMSO alone as a control, were dispensed into target wells using the Echo, and the plates were returned to the incubator for another 17 h. The plates were allowed to cool to RT for 10 minutes, and 5 µL of HiBiT lytic reagent consisting of LgBiT protein (Promega P/N: N4018) diluted 1:100 and lytic substrate (Promega P/N: N246B) diluted 1:50 in lytic buffer (Promega P/N: N247B), was dispensed into each well using the GNF dispenser. After a final incubation at RT for 10 minutes, the plates were read on a luminescence plate reader (Perkin Elmer ViewLux). Luminescence for each data point was normalized using the equation -100*((well value-NC)/(AC-NC)), where NC is the neutral control well (DMSO) and AC is the active control well (mock-transfected with no DNA). $DC_{50}$ values were calculated from curve fits generated by GraphPad Prism v9.0.0.

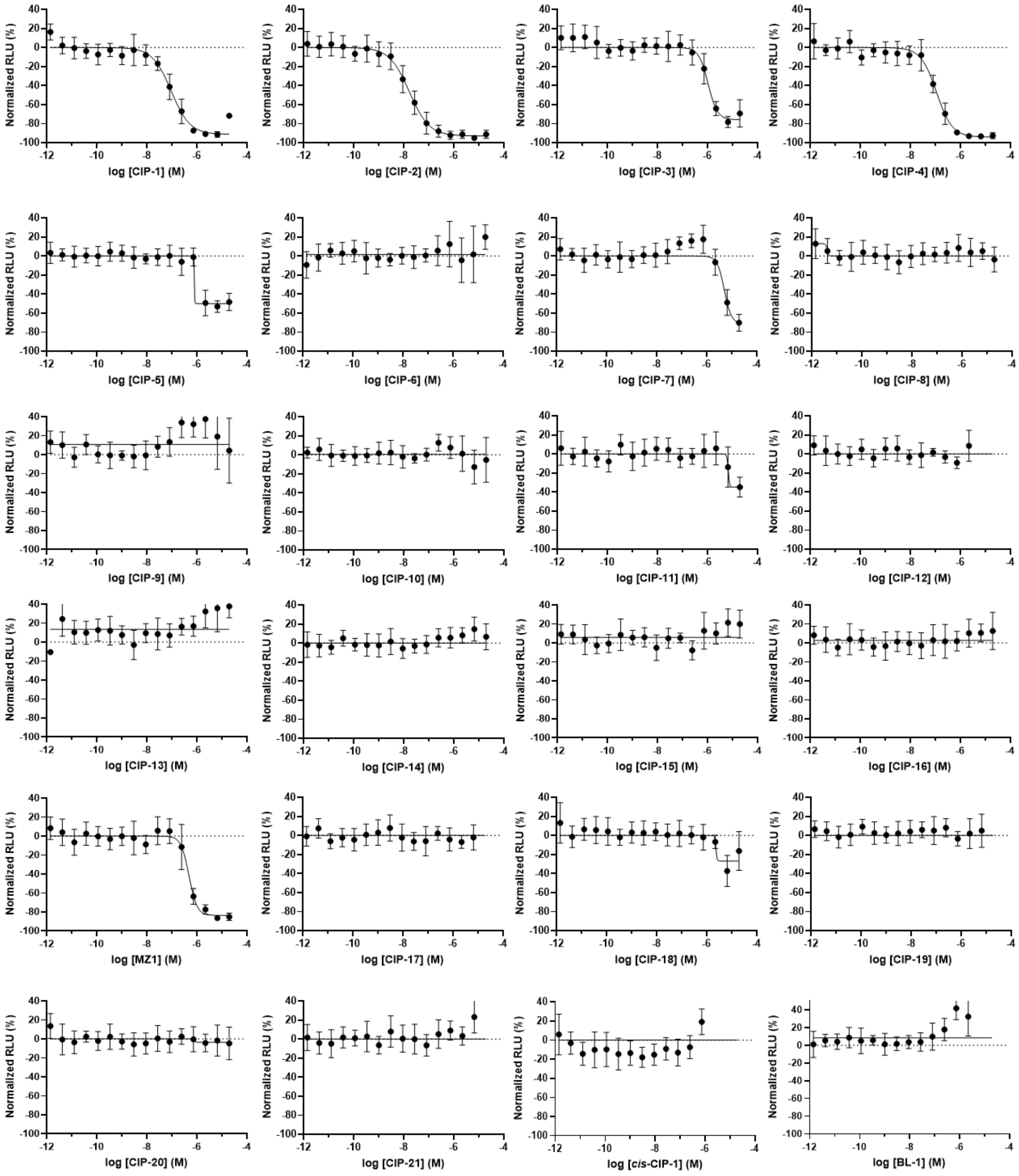


**Figure S17.** Individual curves for cell-based HiBit protein degradation assay (n = 2 independent experiments with 4 technical replicates per experiment).

**Table S1.** Summary of off-DNA validation data with error values for replicates.

| ID # | DEL Enrichment, lb | | | BRD4 Binary SPR | | | Ternary SPR | | | SPR | VHL Binding, TR-FRET | | | | | NanoBit | | | | HiBit | | | |
|---|---|---|---|---|---|---|---|---|---|---|---|---|---|---|---|---|---|---|---|---|---|---|---|
| | (-) VHL | (+) VHL | (+) / (-) ratio | $K_D$ (nM) | ± SD | N | $K_D$ (nM) | ± SD | N | BRD4 alpha | (-) BRD4 (nM) | ± SD | (+) BRD4 (nM) | ± SD | VHL alpha | $pEC_{50}$ | SEM | Emax (%) | SEM | $pDC_{50}$ | SEM | Dmax (%) | SEM |
| CIP-1 | 81 | 389 | 4.8 | 45 | 13 | 4 | 158 | 38 | 9 | 0.29 | 31 | 1 | 42 | 4 | 0.74 | 6.79 | 0.10 | 133 | 5.9 | 7.02 | 0.05 | 91 | 2.7 |
| CIP-2 | 120 | 246 | 2.0 | 13 | 1 | 2 | 34 | 7 | 9 | 0.39 | 27 | 3 | 26 | 3 | 1.05 | 6.99 | 0.07 | 139 | 4.3 | 7.78 | 0.05 | 93 | 1.8 |
| CIP-3 | 20 | 162 | 8.2 | 114 | 28 | 4 | 636 | 26 | 9 | 0.18 | 38 | 3 | 60 | 1 | 0.63 | 6.20 | 0.14 | 122 | 8.9 | 5.97 | 0.05 | 76 | 4.0 |
| CIP-4 | 15 | 150 | 10.2 | 250 | 80 | 7 | 276 | 24 | 7 | 0.91 | 19 | 1 | 24 | 5 | 0.79 | 7.37 | 0.09 | 113 | 4.1 | 6.95 | 0.04 | 94 | 2.2 |
| CIP-5 | 0.9 | 144 | 168.6 | 566 | 210 | 5 | 726 | 31 | 4 | 0.78 | 16 | 2 | 16 | 3 | 0.95 | 6.64 | 0.07 | 115 | 4.3 | 6.10 | > 1 | 50 | 2.4 |
| CIP-6 | 44 | 106 | 2.4 | 26 | NA | 1 | 410 | 81 | 7 | 0.06 | 23 | 1 | 178 | 12 | 0.13 | 6.20 | 0.09 | 101 | 5.5 | < 5 | -- | -- | -- |
| CIP-7 | 0.2 | 100 | 545.2 | 17350 | 3465 | 4 | 2680 | 432 | 8 | 6.47 | 39 | 1 | 15 | 15 | 2.63 | 5.65 | 0.20 | 109 | 14.1 | 5.29 | 0.05 | 71 | 5.2 |
| CIP-8 | 4 | 88 | 22.1 | 3909 | 205 | 3 | -- | -- | -- | -- | 63 | 0.5 | 66 | 5 | 0.96 | < 5 | -- | -- | -- | < 5 | -- | -- | -- |
| CIP-9 | 0.1 | 72 | 806.4 | -- | -- | -- | 2799 | 356 | 8 | > 7.14[‡] | 54 | 2 | 28 | 5 | 1.91 | 6.64 | 0.14 | 100 | 6.4 | < 5 | -- | -- | -- |
| CIP-10 | 0.02 | 53 | 2322 | -- | -- | -- | 3542 | 310 | 4 | > 5.65[‡] | 24 | 1 | 19 | 6 | 1.24 | 5.71 | 0.21 | 100 | 12.4 | < 5 | -- | -- | -- |
| CIP-11 | 5 | 49 | 10.1 | 896 | 301 | 8 | -- | -- | -- | -- | 31 | 2 | 29 | 2 | 1.10 | 5.98 | 0.15 | 61 | 6.0 | 5.16 | > 1 | 34.8 | 12.4 |
| CIP-12 | 34 | 33 | 1.0 | 70 | 3 | 2 | 969 | 242 | 4 | 0.07 | 6 | 0.2 | 283 | 24 | 0.02 | 6.88 | 0.10 | 50 | 3.1 | < 5 | -- | -- | -- |
| CIP-13 | 33 | 32 | 1.0 | 145 | 7 | 2 | -- | -- | -- | -- | 52 | 1 | 372 | 23 | 0.14 | 5.99 | 0.10 | 50 | 4.1 | < 5 | -- | -- | -- |
| CIP-14 | 13 | 31 | 2.4 | 527 | 6 | 2 | 4342 | 857 | 4 | 0.12 | 31 | 1 | 41 | 0.2 | 0.75 | 6.05 | 0.13 | 74 | 6.2 | < 5 | -- | -- | -- |
| CIP-15 | 5 | 20 | 4.0 | 318 | 4 | 2 | -- | -- | -- | -- | 39 | 4 | 68 | 8 | 0.58 | 5.84 | 0.22 | 71 | 10.0 | < 5 | -- | -- | -- |
| CIP-16 | 12 | 19 | 1.6 | 324 | 22 | 2 | -- | -- | -- | -- | 27 | 0.1 | 55 | 1 | 0.49 | 6.28 | 0.08 | 72 | 3.7 | < 5 | -- | -- | -- |
| MZ1 | 22 | 16 | 0.7 | 14 | 8 | 9 | 49 | 6 | 9 | 0.29 | 51 | 6 | 49 | 9 | 1.03 | 6.31 | 0.06 | 160 | 6.1 | 6.31 | 0.04 | 84 | 2.7 |
| CIP-17 | 12 | 12 | 1.0 | -- | -- | -- | -- | -- | -- | -- | 9 | 2 | 11 | 4 | 0.80 | < 5 | -- | -- | -- | < 5 | -- | -- | -- |
| CIP-18 | 13 | 12 | 0.9 | 921 | NA | 1 | -- | -- | -- | -- | 11 | 1 | 26 | 1 | 0.43 | 6.55 | 0.12 | 57 | 3.9 | 5.63 | > 1 | 27 | 4.7 |
| CIP-19 | 6 | 6 | 1.1 | 845 | 328 | 2 | -- | -- | -- | -- | 38 | 2 | 76 | 4 | 0.50 | 6.52 | 0.08 | 60 | 3.1 | < 5 | -- | -- | -- |
| CIP-20 | 0.4 | 5 | 12.4 | 3870 | 833 | 4 | 3323 | 312 | 7 | 1.16 | 153 | 1 | 88 | 1 | 1.74 | < 5 | -- | -- | -- | < 5 | -- | -- | -- |
| CIP-21 | 8 | 2 | 0.2 | 791 | NA | 1 | -- | -- | -- | -- | 27 | 4 | 57 | 0.2 | 0.47 | < 5 | -- | -- | -- | < 5 | -- | -- | -- |

*See* ***Table 1*** *of main text for additional details.*

#### 16. Cell-Titer Glo cell viability assay.

The Cell-Titer Glo assay was performed identically to the HiBiT cellular protein degradation assay, except 5 µL of Cell-Titer Glo substrate (Promega P/N: G7571) was added instead of HiBiT lytic reagent. Luminescence was not normalized.

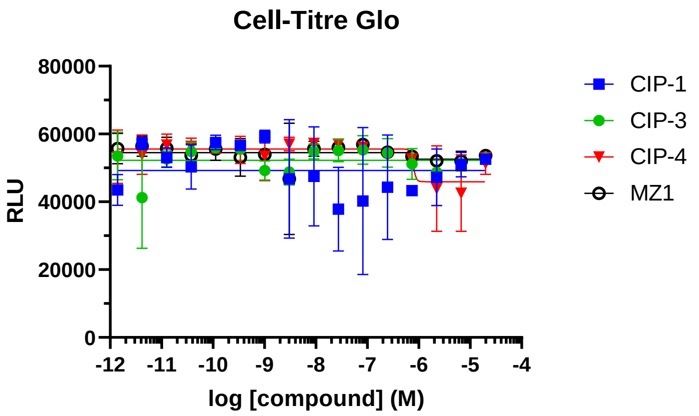


**Figure S18.** Cell viability assessment using three degraders identified from CIP-DEL library and MZ1 control (n = 1 experiment with 3 technical replicates).

#### 17. Western blotting.

Degradation of endogenous BRD4 was validated by western blotting in HEK293 cells cultured in DMEM supplemented with 10% FBS and 100 U / mL penicillin-streptomycin. For each treatment condition, 350k cells in 2 mL of media were plated in 6-well culture plates and incubated at 37°C with 5% $CO_2$. After 4 h, 200 µL of media was removed and replaced with 200 µL of 10x drug treatment solution diluted in media. Final DMSO concentrations were 1%. Cells were treated with compounds for 18 h, then placed on ice and washed twice with ice cold PBS. The cells were lysed with 100 µL RIPA buffer (20 mM Tris pH 7.4, 150 mM NaCl, 2 mM EDTA, 0.1% SDS, 1% NP-40, 0.25% deoxycholate, 1 mM $Na_2VO_4$, 1 mM PMSF, 1 mM NaF, and Roche protease inhibitor cocktail) and the lysates were clarified by centrifugation at 20,000 x g for 30 min at 4°C. Protein concentrations were determined for each lysate using the Pierce Detergent Compatible Bradford Assay (P/N: 1863028) with Bio-Rad BGG protein standards (P/N: 5000209) and 30 µg of each protein lysate was separated via electrophoresis on polyacrylamide gels. Proteins were transferred to nitrocellulose membranes using the Trans-Blot Turbo Transfer System (Bio-Rad) and the membranes were blocked with 5% BSA in TBST for 30 min. The blots were probed with an anti-BRD4 rabbit monoclonal antibody (Abcam, P/N: ab128874, 1:1000) at 4°C with gentle shaking overnight, then stained with IRDye 800CW donkey anti-rabbit secondary antibody (LiCOR, P/N: 926-32213, 1:10,000) and imaged using a LiCOR Odyssey infrared imaging system. Finally, blots were probed with an anti-β-Actin primary antibody (Cell Signaling Technology, P/N: 4970S, 1:2000) and IRDye 800CW secondary (as above) and imaged to confirm uniform protein loading.

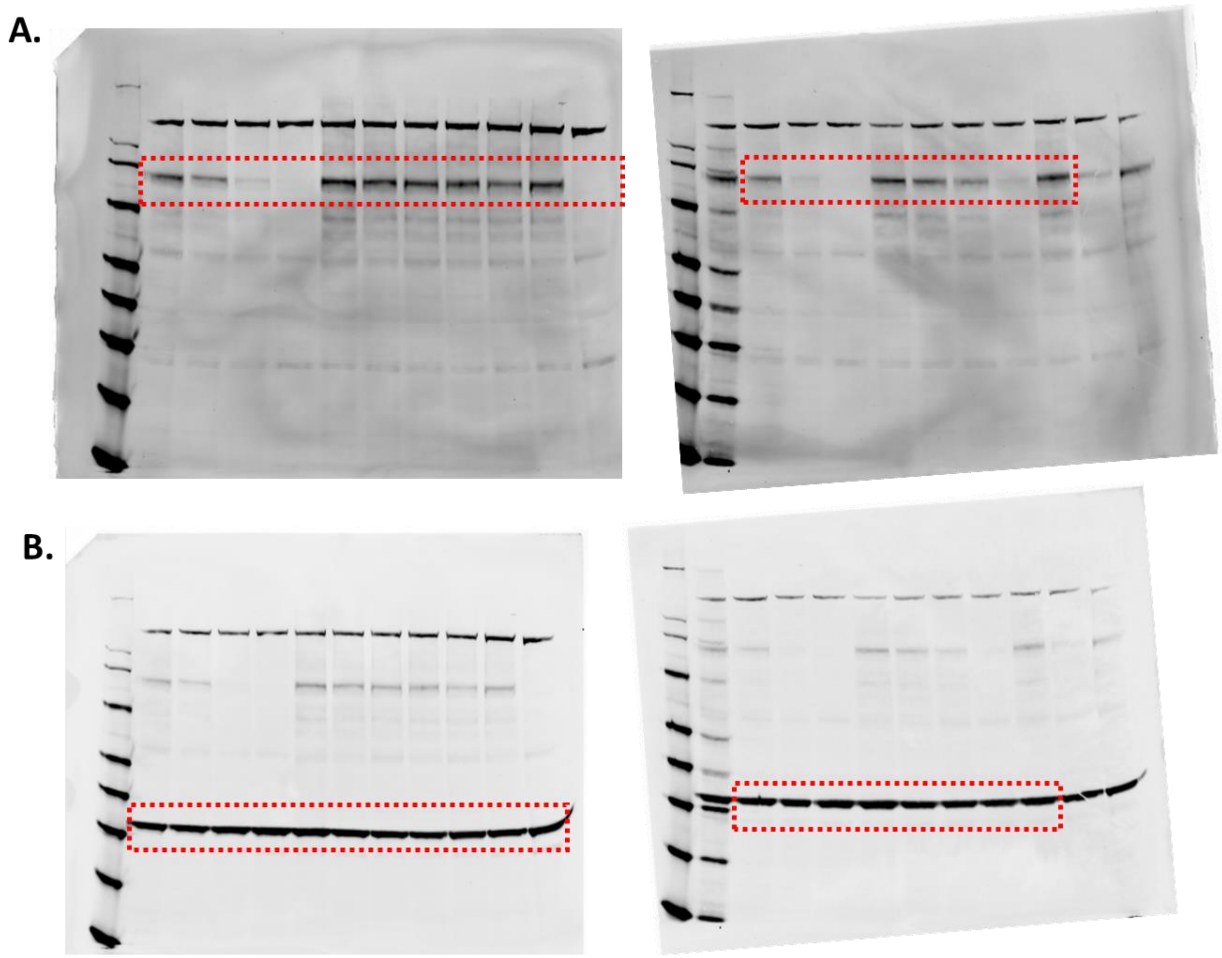


**Figure S19.** Uncropped western blot gels. (**A**) Gel imaging after staining with primary anti-BRD4 antibody. (**B**) Gel imaging after re-straining with secondary anti-β-actin antibody.

#### 18. X-ray crystallography.

VCB complex (233 μM) and BRD4$^{BD1}$ (1280 μM) were dialyzed separately into buffer containing 5 mM Hepes pH 7.5, 100 mM NaCl, and 0.5 mM TCEP. After dialysis overnight, VCB complex and BRD4$^{BD1}$ were mixed at 1:1 molar ratio in the presence of 200 μM **CIP-1**. The VCB / BRD4$^{BD1}$ / **CIP-1** complex was concentrated to 10 mg / ml (180 μM) prior to crystallization screens. Multiple crystal hits were identified from related conditions such as 0.2 M sodium malonate, 20% w/v PEG 3350 (Hampton research PEG/Ion E8) and 0.2 M sodium citrate, 20% w/v PEG 3350 (Hampton research PEG/Ion D10), after 2 – 4 days of incubation. Single crystals suitable for data collection were obtained after several rounds of optimization with streak seeding in 0.2 M sodium malonate, 16 – 20% w/v PEG 3350. Crystals were cryo-protected in reservoir solution supplemented with 20% glycerol and flash-cooled in liquid nitrogen for data collection. Datasets were collected under cryogenic conditions (100 K) at the Advanced Light Source (ALS) beamline 5.0.2. All data were processed using CCP4[40], employing XDS[41] for data integration and AIMLESS[42] for scaling. Molecular replacement was carried out using PHASER[43] with coordinates from human VHL-ElonginC-ElonginB complex (PDB ID: 1VCB)[44] and the first bromodomain of human BRD4 (PDB ID: 2OSS).[45] Structure refinement was carried out in PHENIX[46] alternated with manual fitting in Coot.[47] Data collection and structure refinement statistics are included in **Extended Data Table 2**.
