## Supplementary material for "DNA-encoded library (DEL)-enabled discovery of proximity-inducing small molecules": Spectral Data

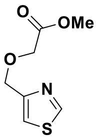
O
OMe
O
N
S

**S2**
($CDCl_3$)

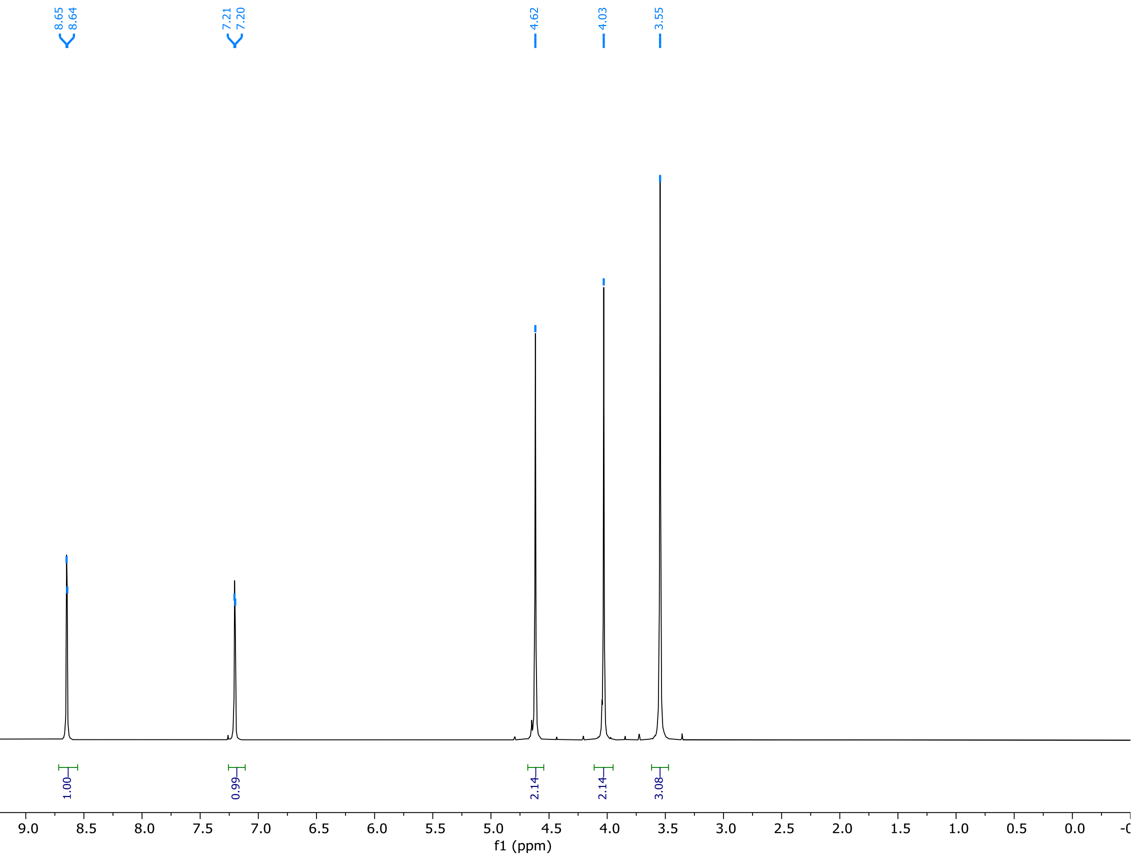
8.65
8.64
7.21
7.20
4.62
4.03
3.55
1.00
0.99
2.14
2.14
3.08
f1 (ppm)

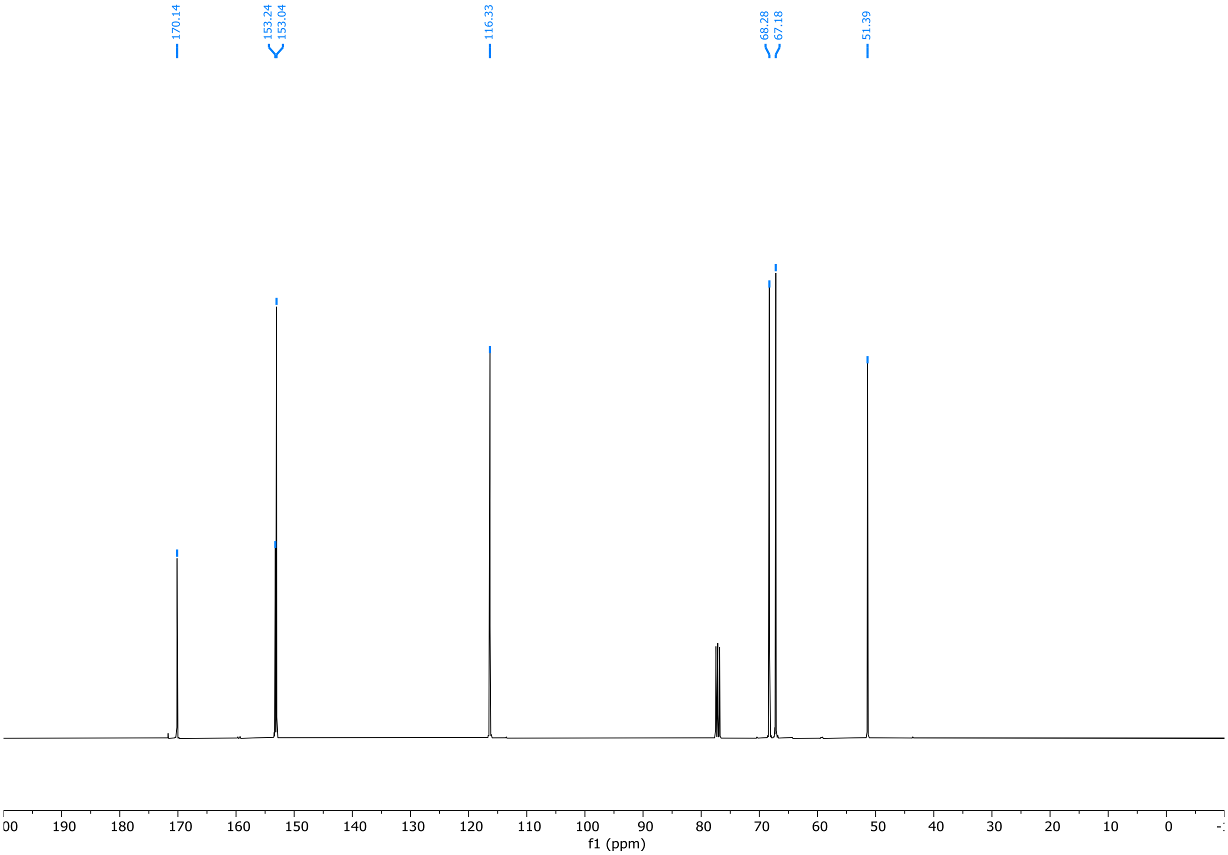
170.14
153.24
153.04
116.33
68.28
67.18
51.39
f1 (ppm)

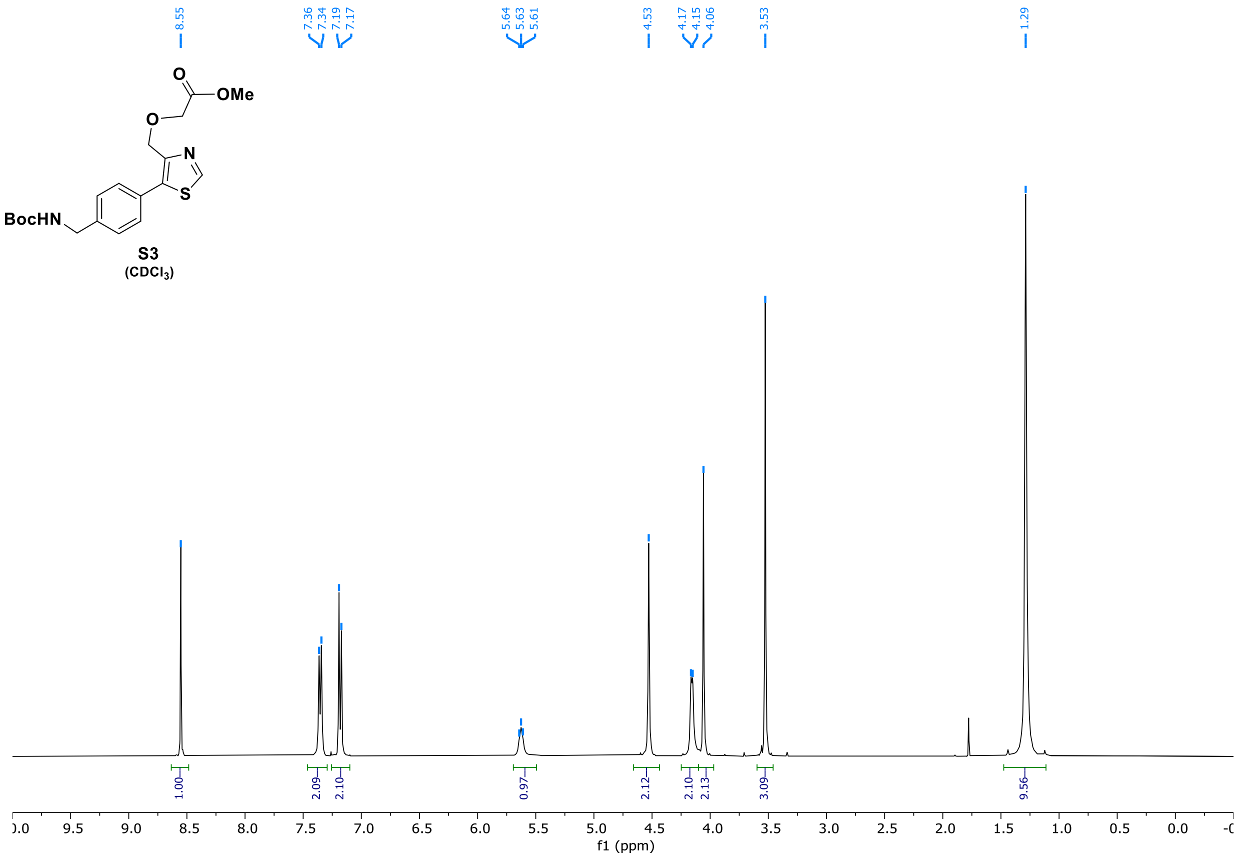
O
OMe
O
N
S
BocHN
S3
(CDCl3)
8.55
7.36
7.34
7.19
7.17
5.64
5.63
5.61
4.53
4.17
4.15
4.06
3.53
1.29
1.00
2.09
2.10
0.97
2.12
2.10
2.13
3.09
9.56
f1 (ppm)

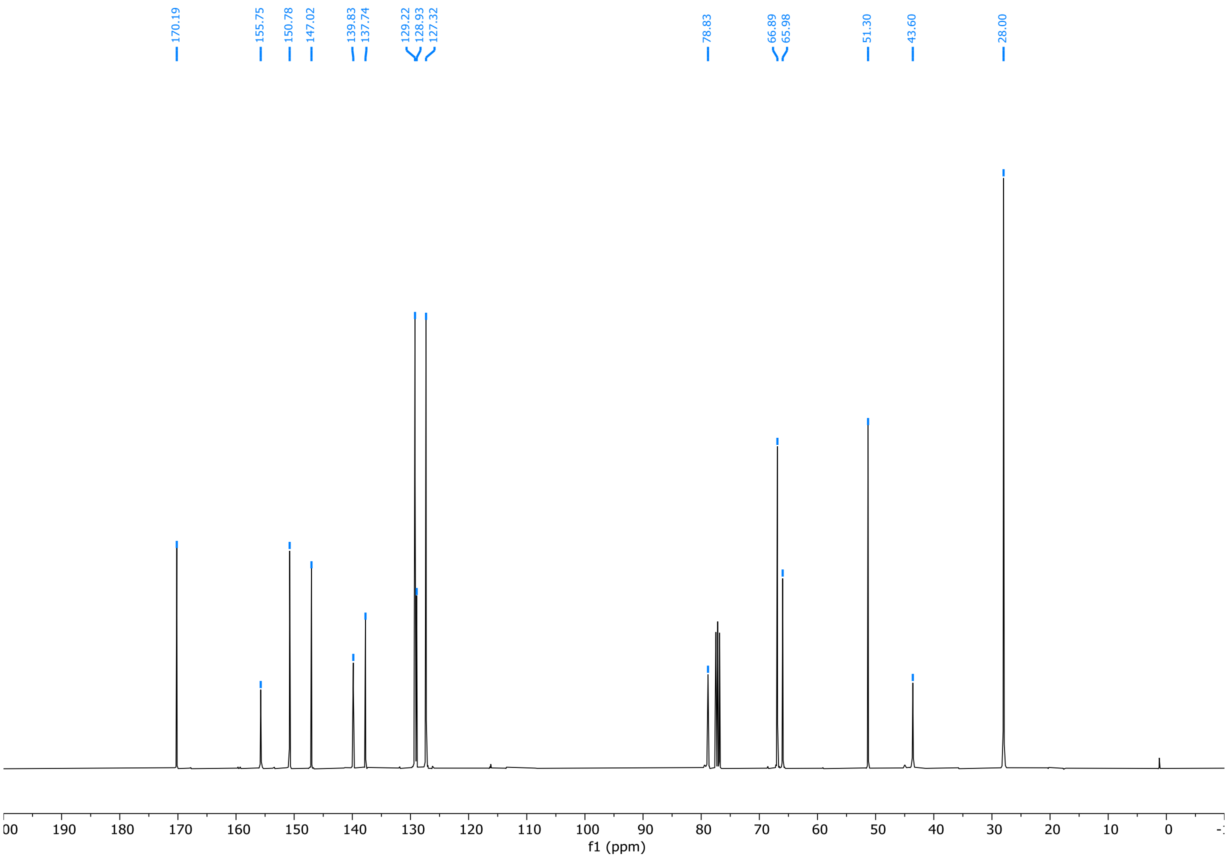
170.19
155.75
150.78
147.02
139.83
137.74
129.22
128.93
127.32
78.83
66.89
65.98
51.30
43.60
28.00
f1 (ppm)

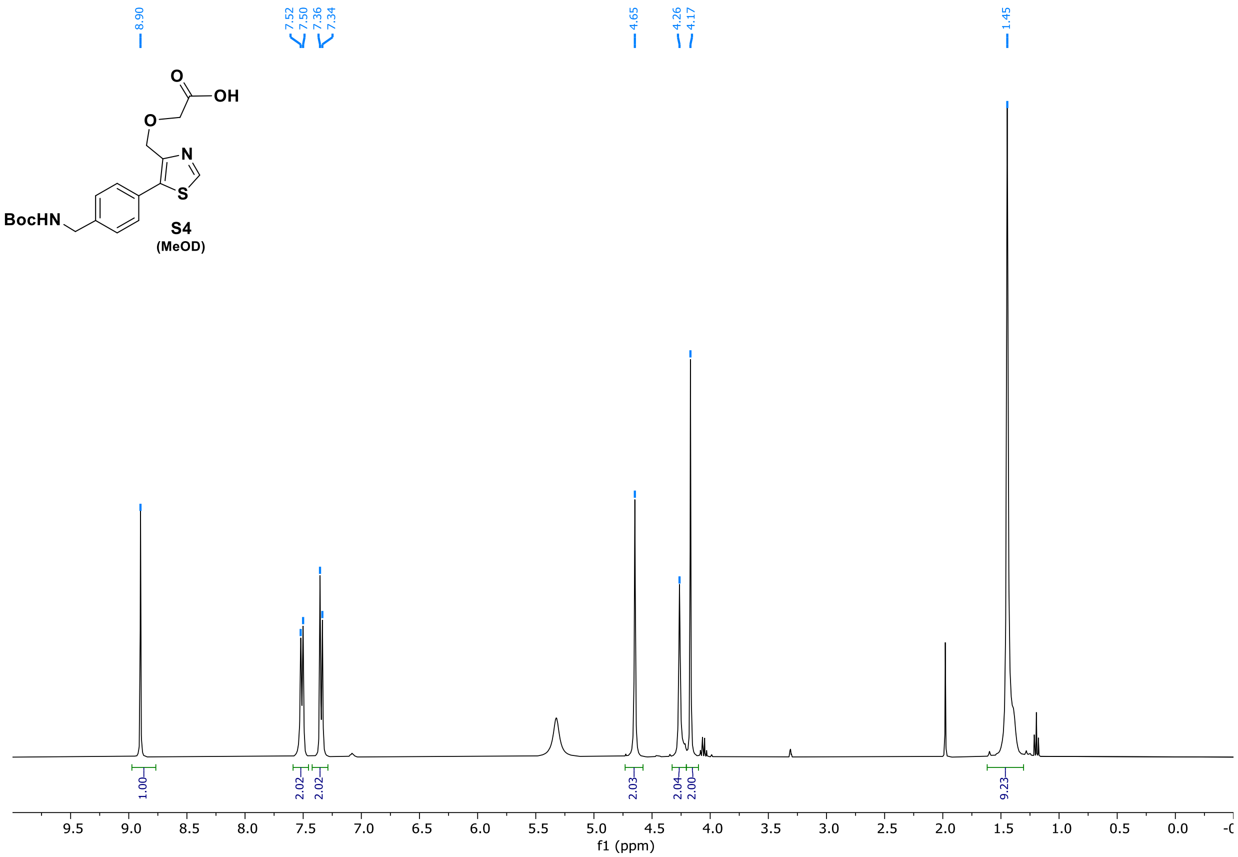
8.90
7.52
7.50
7.36
7.34
4.65
4.26
4.17
1.45
O
OH
O
N
S
BocHN
S4
(MeOD)
1.00
2.02
2.02
2.03
2.04
2.00
9.23
9.5
9.0
8.5
8.0
7.5
7.0
6.5
6.0
5.5
5.0
4.5
4.0
3.5
3.0
2.5
2.0
1.5
1.0
0.5
0.0
f1 (ppm)

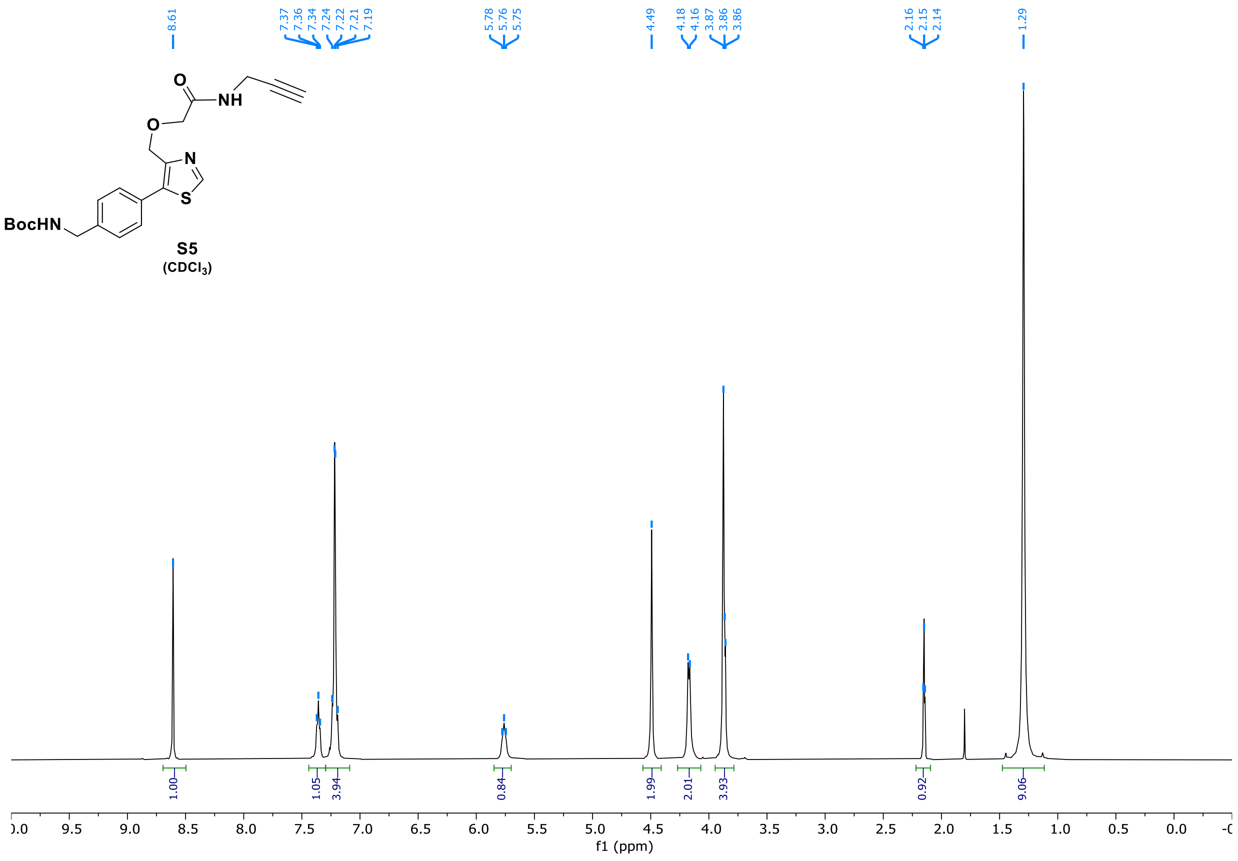
8.61
7.37
7.36
7.34
7.24
7.22
7.21
7.19
5.78
5.76
5.75
4.49
4.18
4.16
3.87
3.86
3.86
2.16
2.15
2.14
1.29
O
NH
O
N
S
BocHN
S5
(CDCl3)
1.00
1.05
3.94
0.84
1.99
2.01
3.93
0.92
9.06
9.5
9.0
8.5
8.0
7.5
7.0
6.5
6.0
5.5
5.0
4.5
4.0
3.5
3.0
2.5
2.0
1.5
1.0
0.5
0.0
f1 (ppm)

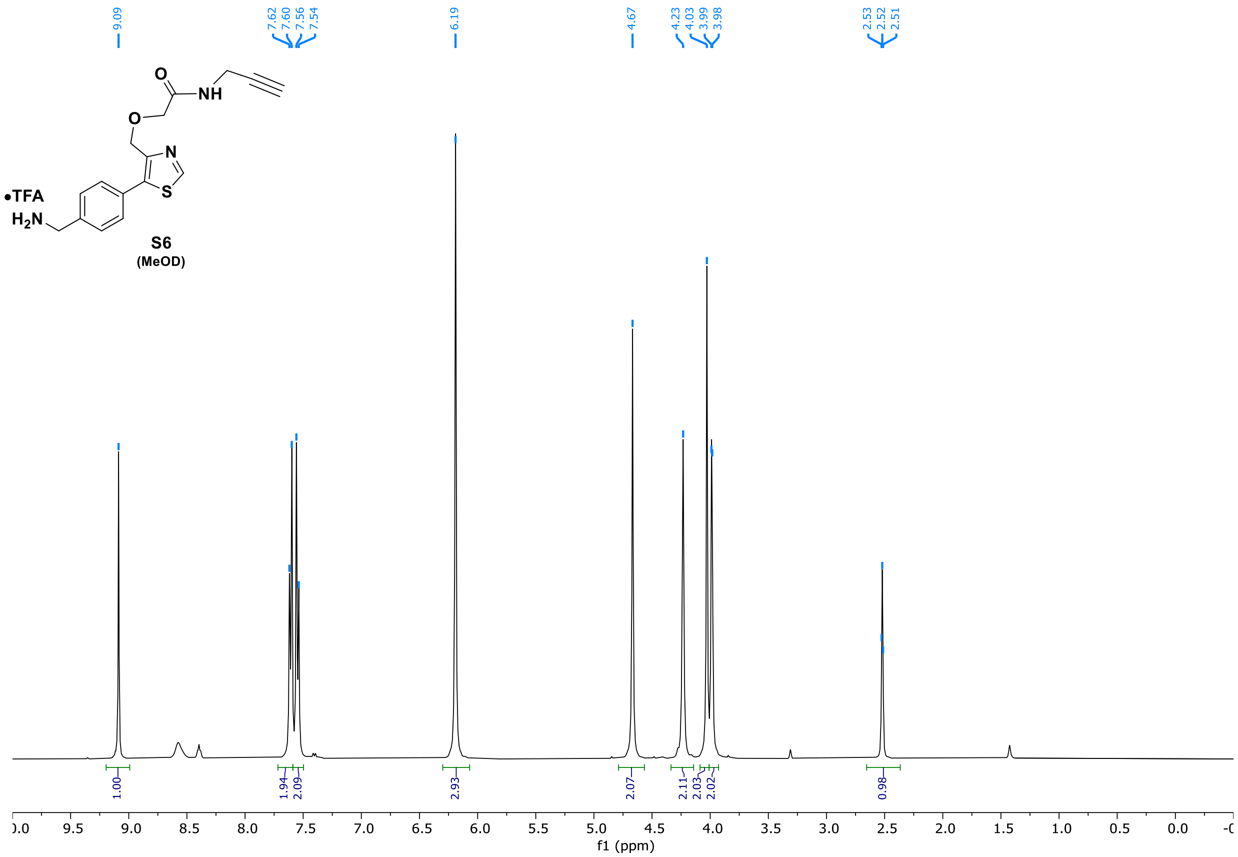
9.09
7.62
7.60
7.56
7.54
6.19
4.67
4.23
4.03
3.99
3.98
2.53
2.52
2.51
O
NH
O
N
S
•TFA
H2N
S6
(MeOD)
1.00
1.94
2.09
2.93
2.07
2.11
2.03
2.02
0.98
f1 (ppm)

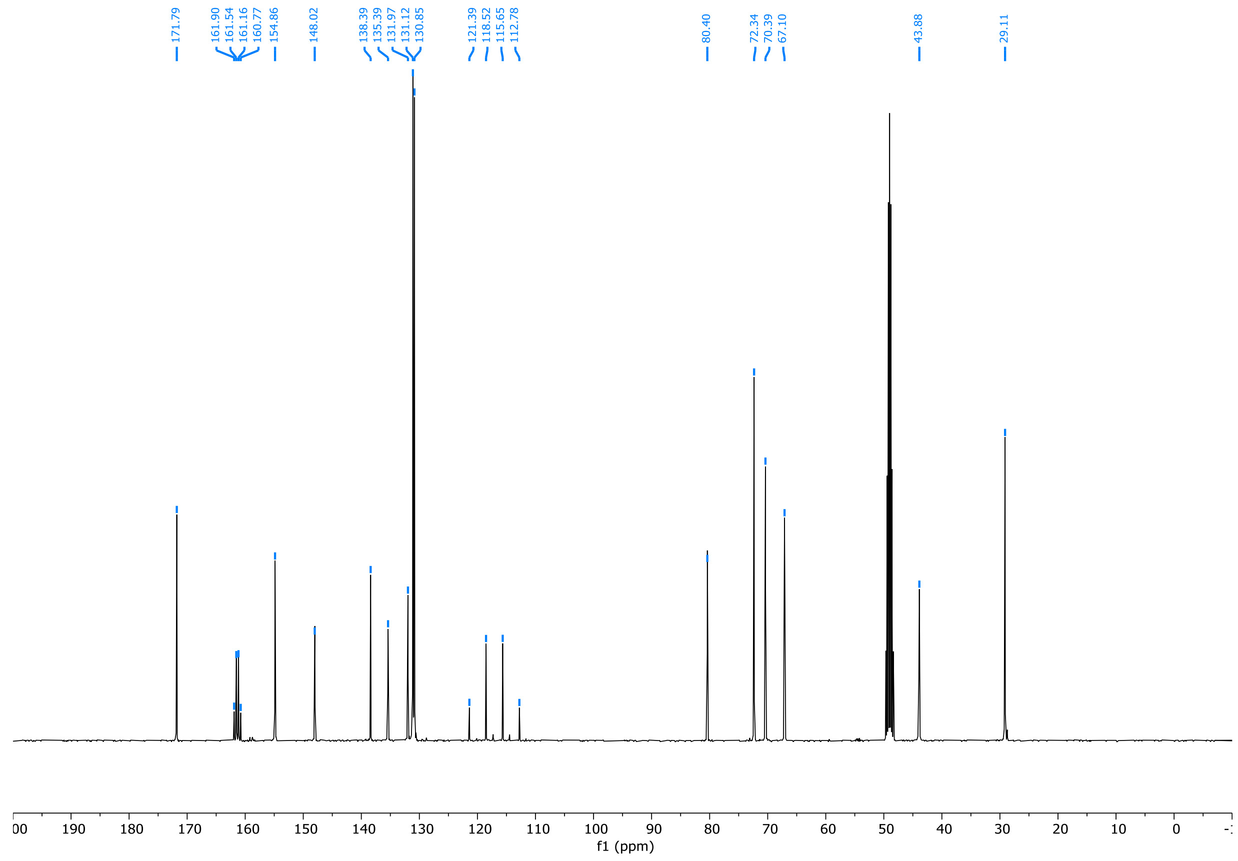
171.79
161.90
161.54
161.16
160.77
154.86
148.02
138.39
135.39
131.97
131.12
130.85
121.39
118.52
115.65
112.78
80.40
72.34
70.39
67.10
43.88
29.11
f1 (ppm)

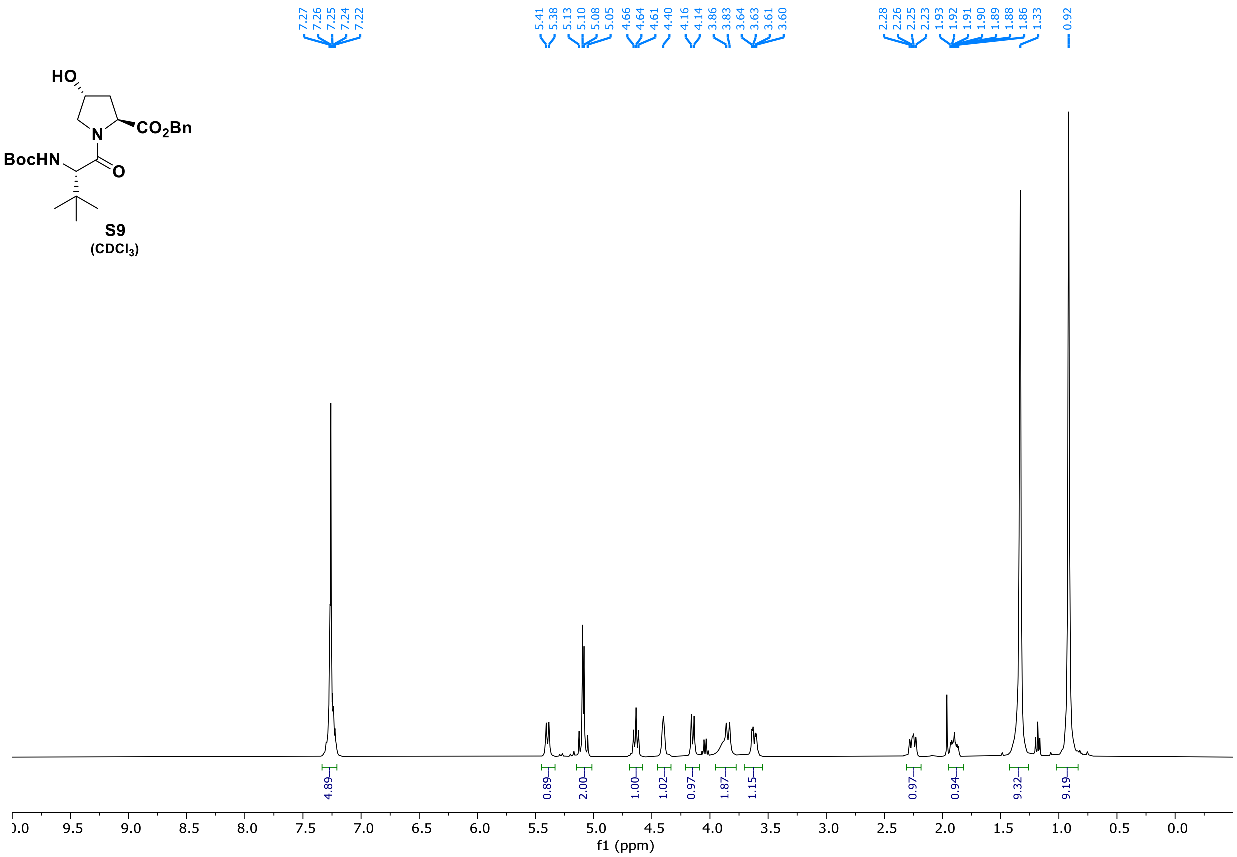
HO
$CO_2Bn$
BocHN
N
O
S9
($CDCl_3$)
7.27
7.26
7.25
7.24
7.22
5.41
5.38
5.13
5.10
5.08
5.05
4.66
4.64
4.61
4.40
4.16
4.14
3.86
3.83
3.64
3.63
3.61
3.60
2.28
2.26
2.25
2.23
1.93
1.92
1.91
1.90
1.89
1.88
1.86
1.33
0.92
4.89
0.89
2.00
1.00
1.02
0.97
1.87
1.15
0.97
0.94
9.32
9.19
f1 (ppm)

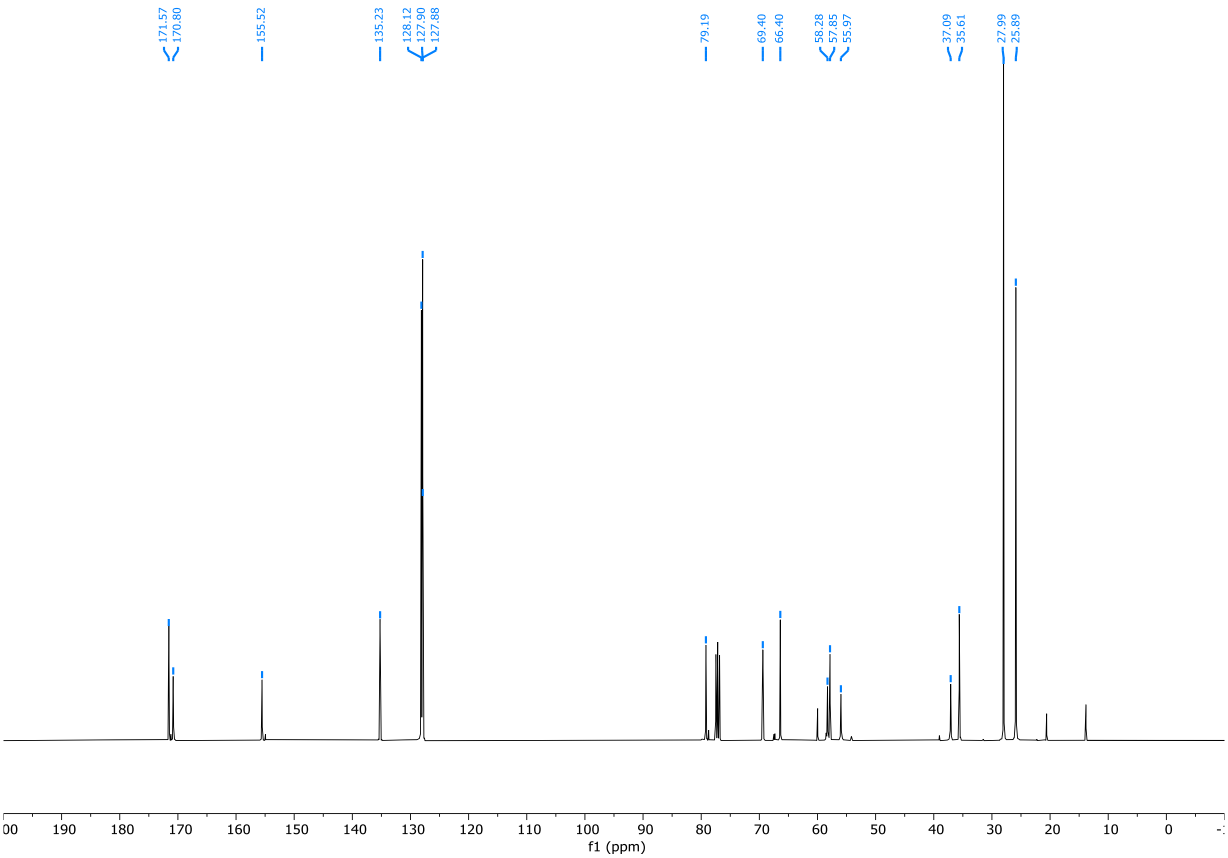
171.57
170.80
155.52
135.23
128.12
127.90
127.88
79.19
69.40
66.40
58.28
57.85
55.97
37.09
35.61
27.99
25.89
f1 (ppm)

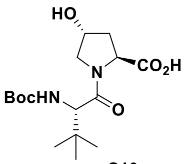
HO
CO2H
N
BocHN
O

**S10**
**(MeOD)**

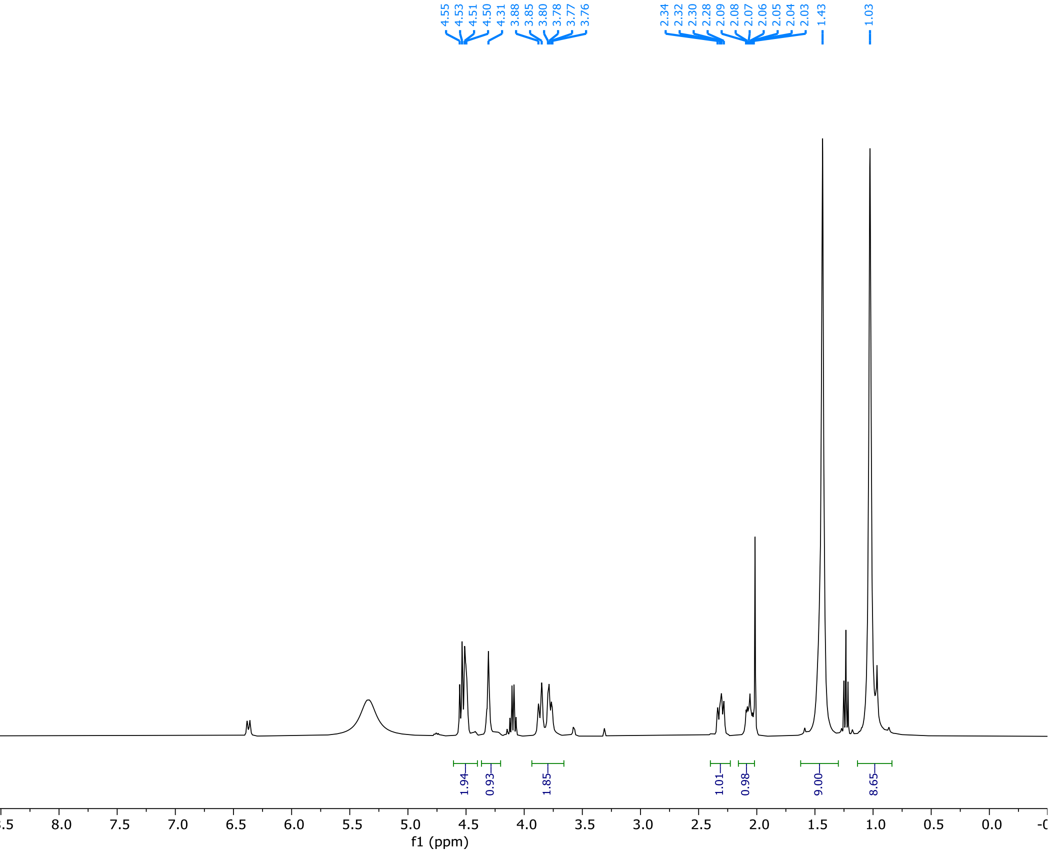
4.55
4.53
4.51
4.50
4.31
3.88
3.85
3.80
3.78
3.77
3.76
2.34
2.32
2.30
2.28
2.09
2.08
2.07
2.06
2.05
2.04
2.03
1.43
1.03
1.94
0.93
1.85
1.01
0.98
9.00
8.65
9.5 9.0 8.5 8.0 7.5 7.0 6.5 6.0 5.5 5.0 4.5 4.0 3.5 3.0 2.5 2.0 1.5 1.0 0.5 0.0
f1 (ppm)

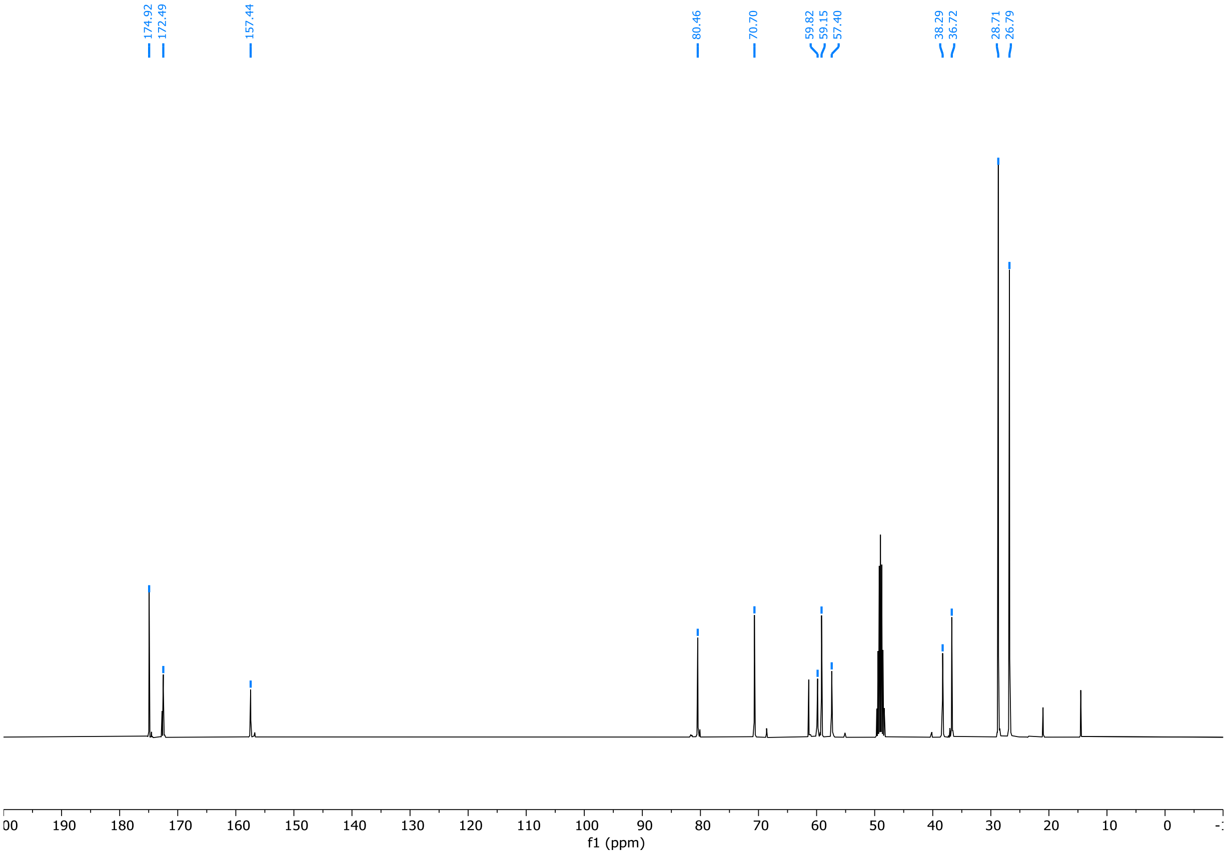
174.92
172.49
157.44
80.46
70.70
59.82
59.15
57.40
38.29
36.72
28.71
26.79
190 180 170 160 150 140 130 120 110 100 90 80 70 60 50 40 30 20 10 0
f1 (ppm)

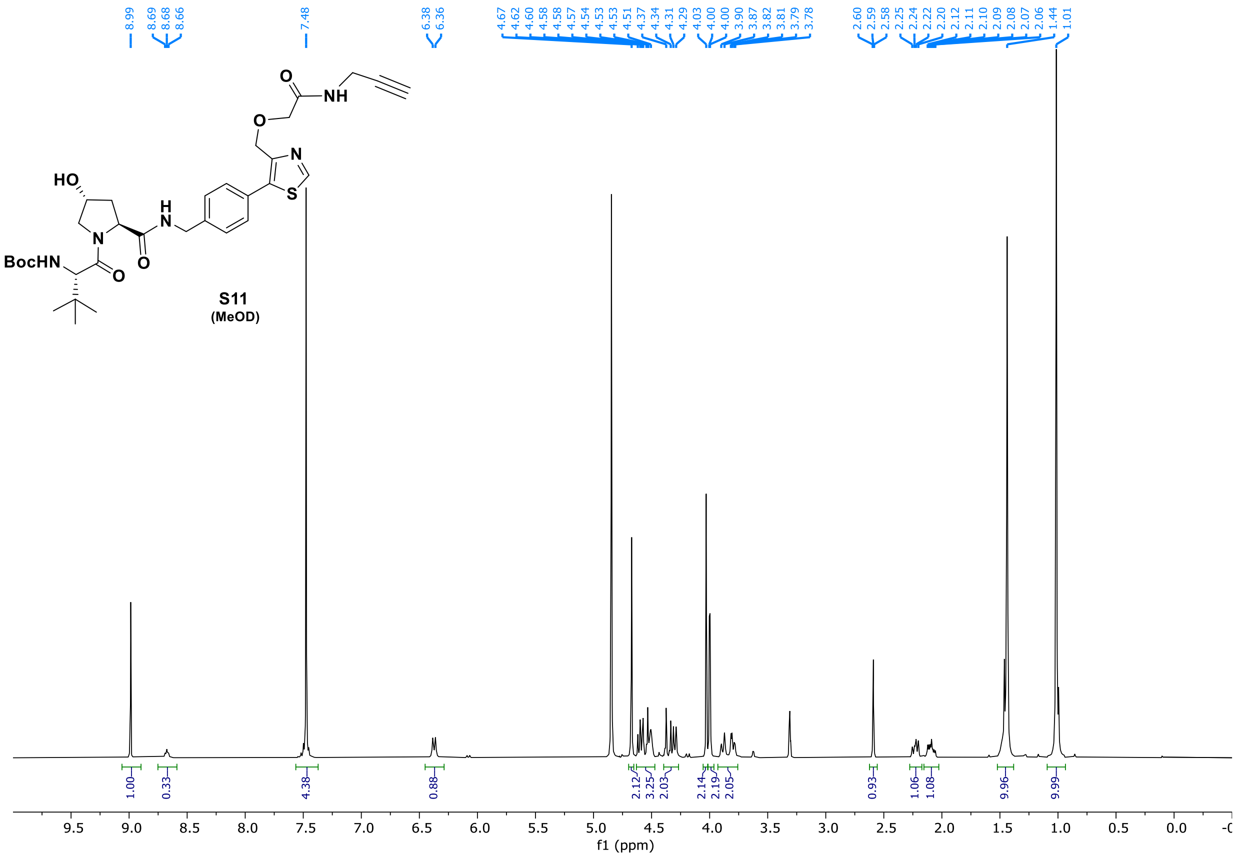
8.99
8.69
8.68
8.66
7.48
6.38
6.36
4.67
4.62
4.60
4.58
4.58
4.57
4.54
4.53
4.53
4.51
4.37
4.34
4.31
4.29
4.03
4.00
4.00
3.90
3.87
3.82
3.81
3.79
3.78
2.60
2.59
2.58
2.25
2.24
2.22
2.20
2.12
2.11
2.10
2.09
2.08
2.07
2.06
1.44
1.01
HO
BocHN
N
H
N
O
O
O
O
NH
N
S
S11
(MeOD)
1.00
0.33
4.38
0.88
2.12
3.25
2.03
2.14
2.19
2.05
0.93
1.06
1.08
9.96
9.99
f1 (ppm)

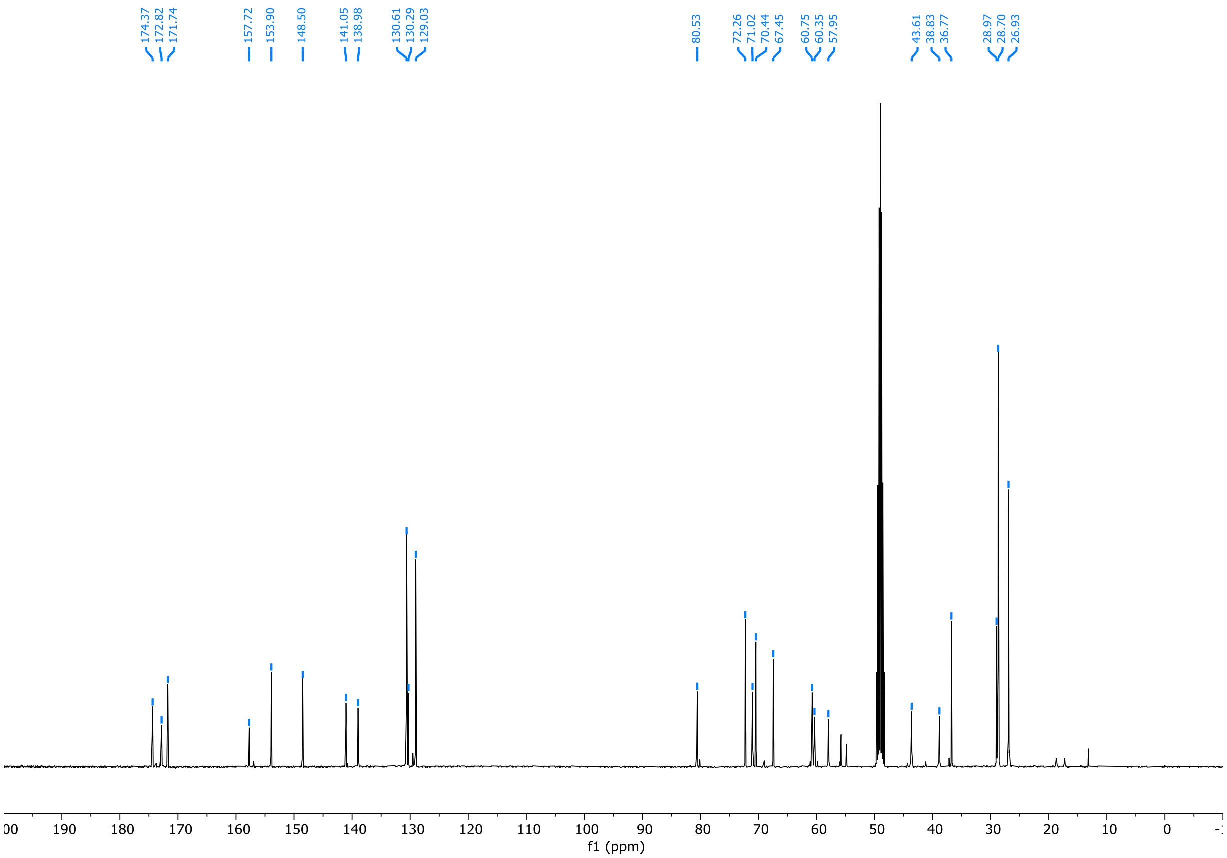
174.37
172.82
171.74
157.72
153.90
148.50
141.05
138.98
130.61
130.29
129.03
80.53
72.26
71.02
70.44
67.45
60.75
60.35
57.95
43.61
38.83
36.77
28.97
28.70
26.93
f1 (ppm)

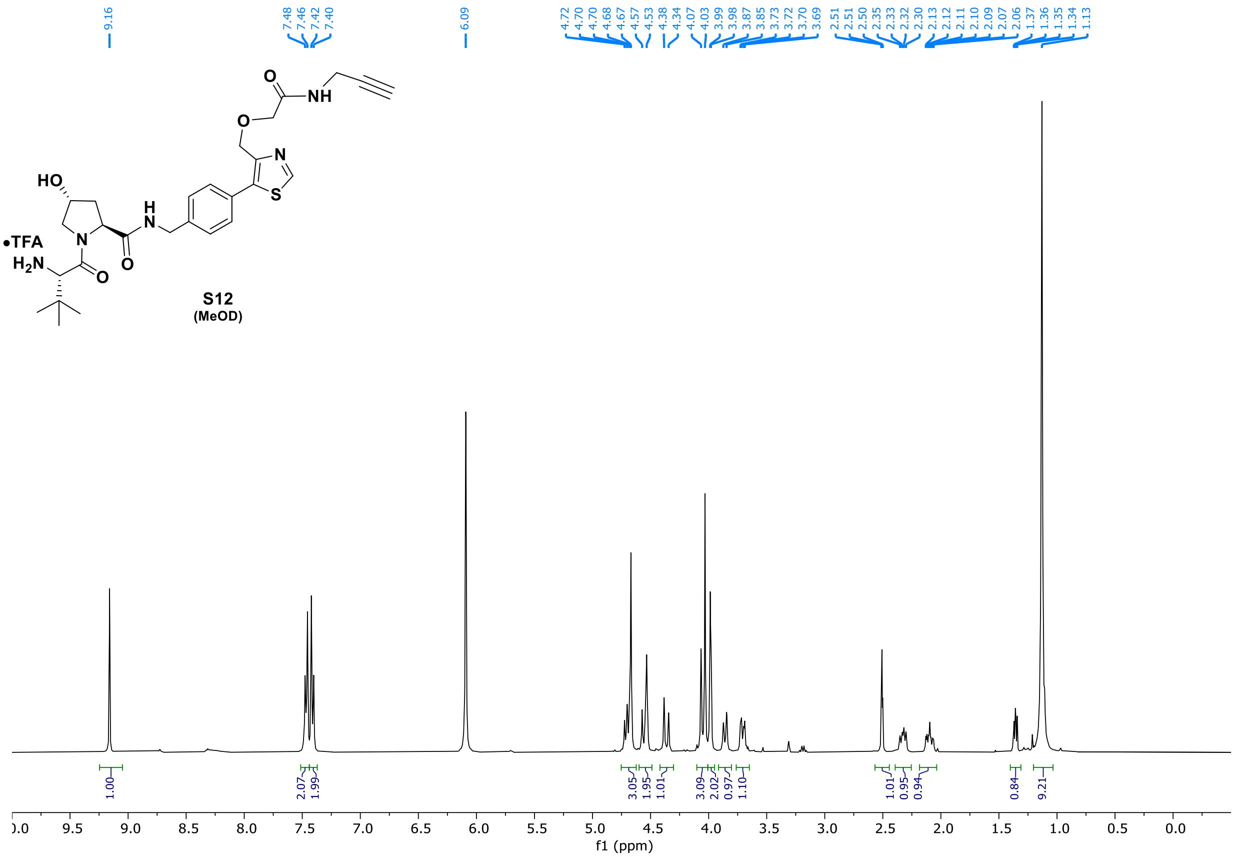
9.16
7.48 7.46 7.42 7.40
6.09
4.72 4.70 4.70 4.68 4.67 4.57 4.53 4.38 4.34 4.07 4.03 3.99 3.98 3.87 3.85 3.73 3.72 3.70 3.69
2.51 2.51 2.50 2.35 2.33 2.32 2.30 2.13 2.12 2.11 2.10 2.09 2.07 2.06 1.37 1.36 1.35 1.34 1.13
•TFA
S12
(MeOD)
1.00
2.07 1.99
3.05 1.95 1.01
3.09 2.02 0.97 1.10
1.01 0.95 0.94
0.84
9.21
f1 (ppm)

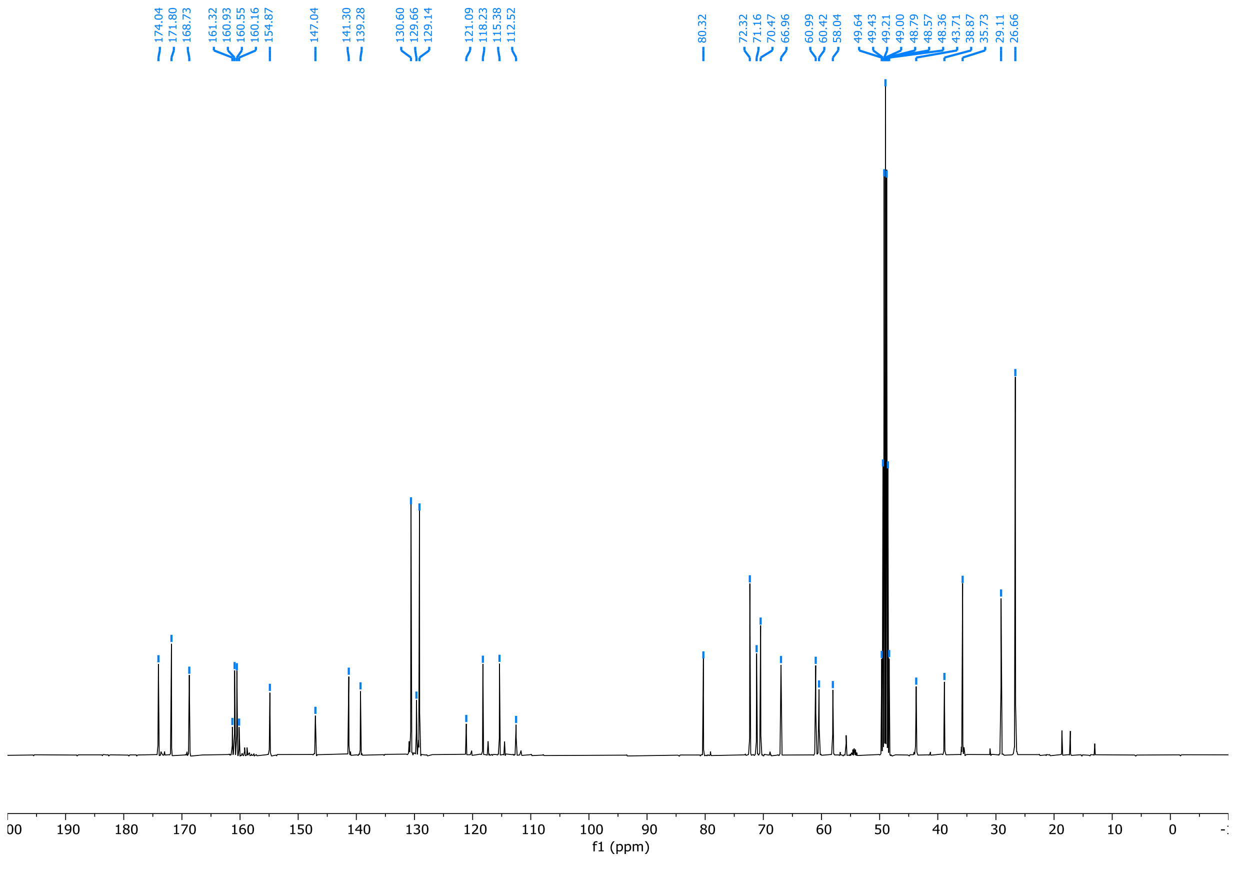
174.04 171.80 168.73 161.32 160.93 160.55 160.16 154.87
147.04 141.30 139.28
130.60 129.66 129.14
121.09 118.23 115.38 112.52
80.32
72.32 71.16 70.47 66.96 60.99 60.42 58.04
49.64 49.43 49.21 49.00 48.79 48.57 48.36 43.71 38.87 35.73 29.11 26.66
f1 (ppm)

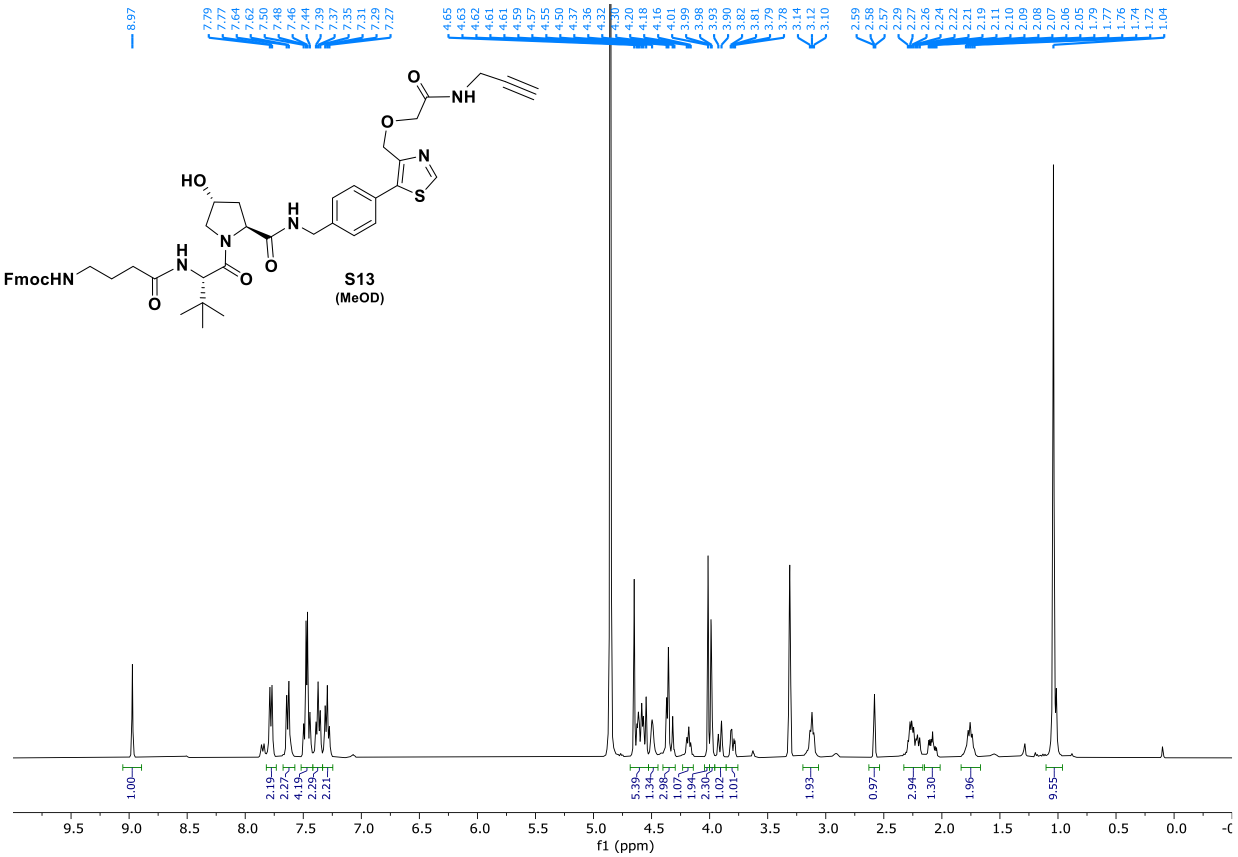

8.97
7.79 7.77 7.64 7.62 7.50 7.48 7.46 7.44 7.39 7.37 7.35 7.31 7.29 7.27
4.65 4.63 4.62 4.61 4.61 4.59 4.57 4.55 4.50 4.37 4.36 4.32 4.30 4.20 4.18 4.16 4.01 3.99 3.98 3.93 3.90 3.82 3.81 3.79 3.78
3.14 3.12 3.10
2.59 2.58 2.57 2.29 2.27 2.26 2.24 2.22 2.21 2.19 2.11 2.10 2.09 2.08 2.07 2.06 2.05 1.79 1.77 1.76 1.74 1.72 1.04
FmocHN
HO
N
H
O
NH
S
1.00 2.19 2.27 4.19 2.29 2.21 5.39 1.34 2.98 1.07 1.94 2.30 1.02 1.01 1.93 0.97 2.94 1.30 1.96 9.55
f1 (ppm)


S13
(MeOD)

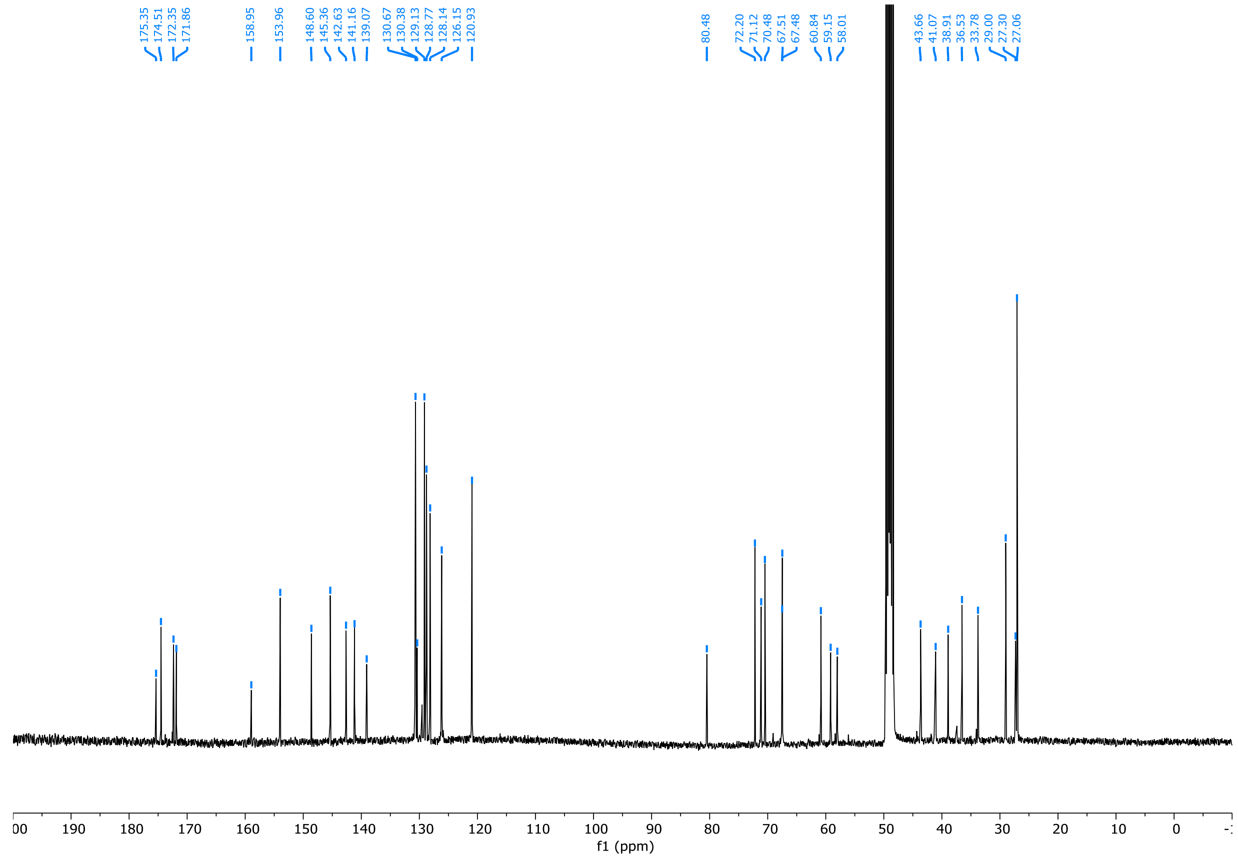

175.35 174.51 172.35 171.86 158.95 153.96 148.60 145.36 142.63 141.16 139.07 130.67 130.38 129.13 128.77 128.14 126.15 120.93
80.48 72.20 71.12 70.48 67.51 67.48 60.84 59.15 58.01
43.66 41.07 38.91 36.53 33.78 29.00 27.30 27.06
f1 (ppm)

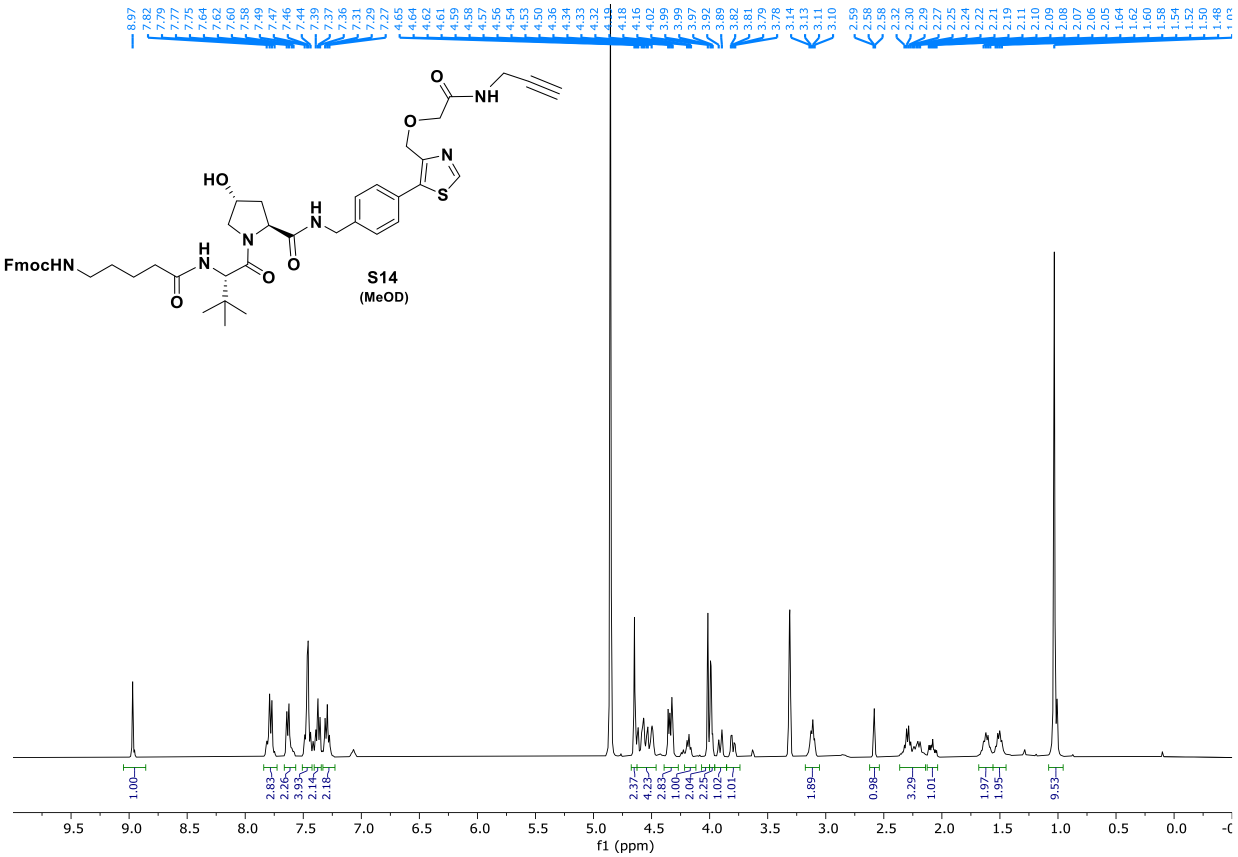
8.97
7.82
7.79
7.77
7.75
7.64
7.62
7.60
7.58
7.49
7.47
7.46
7.44
7.39
7.37
7.36
7.31
7.29
7.27
4.65
4.64
4.62
4.61
4.59
4.58
4.57
4.56
4.54
4.53
4.50
4.36
4.34
4.33
4.32
4.19
4.18
4.16
4.02
3.99
3.99
3.97
3.92
3.89
3.82
3.81
3.79
3.78
3.14
3.13
3.11
3.10
2.59
2.58
2.58
2.32
2.30
2.29
2.27
2.25
2.24
2.22
2.21
2.19
2.11
2.10
2.09
2.08
2.07
2.06
2.05
1.64
1.62
1.60
1.58
1.54
1.52
1.50
1.48
FmocHN
HO
S14
(MeOD)
1.00
2.83
2.26
3.93
2.14
2.18
2.37
4.23
2.83
1.00
2.04
2.25
1.02
1.01
1.89
0.98
3.29
1.01
1.97
1.95
9.53
f1 (ppm)

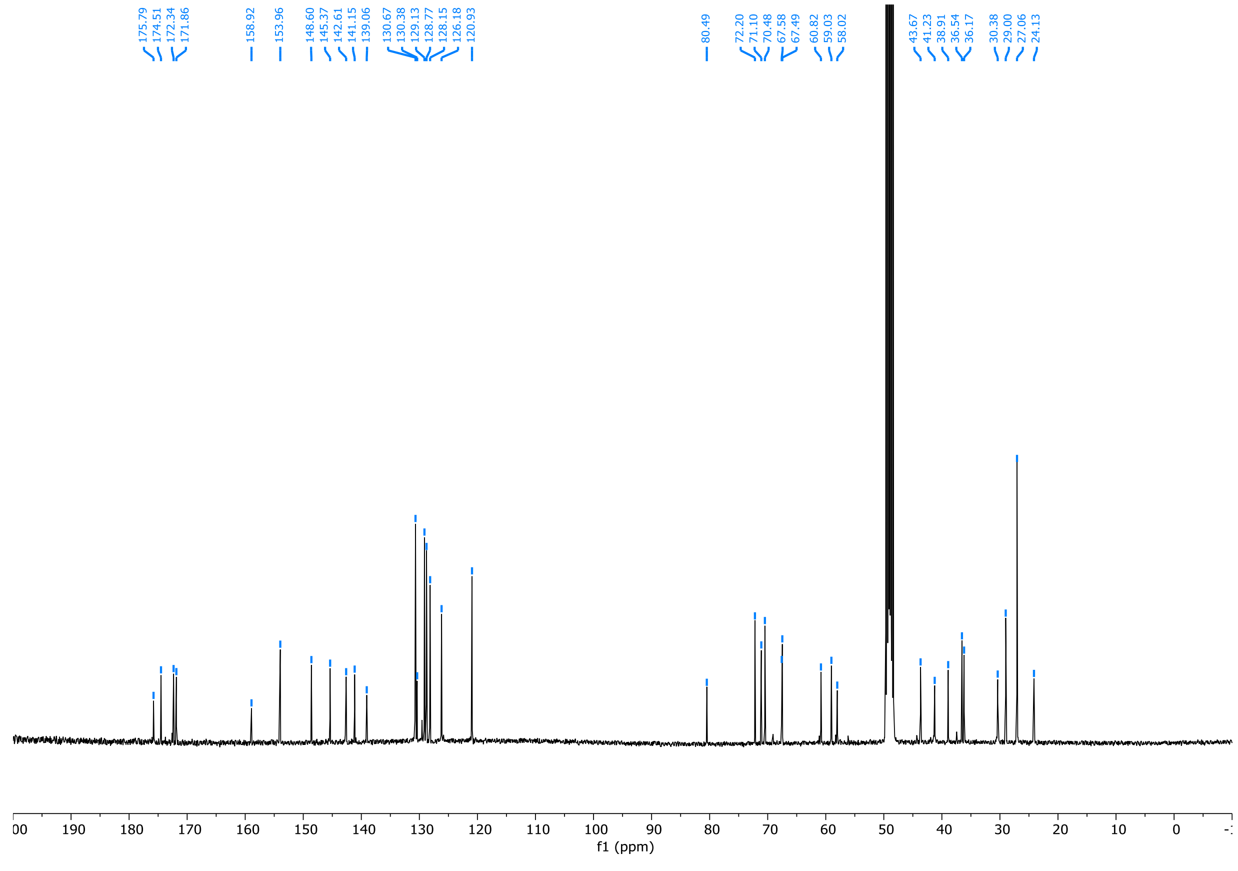
175.79
174.51
172.34
171.86
158.92
153.96
148.60
145.37
142.61
141.15
139.06
130.67
130.38
129.13
128.77
128.15
126.18
120.93
80.49
72.20
71.10
70.48
67.58
67.49
60.82
59.03
58.02
43.67
41.23
38.91
36.54
36.17
30.38
29.00
27.06
24.13
f1 (ppm)

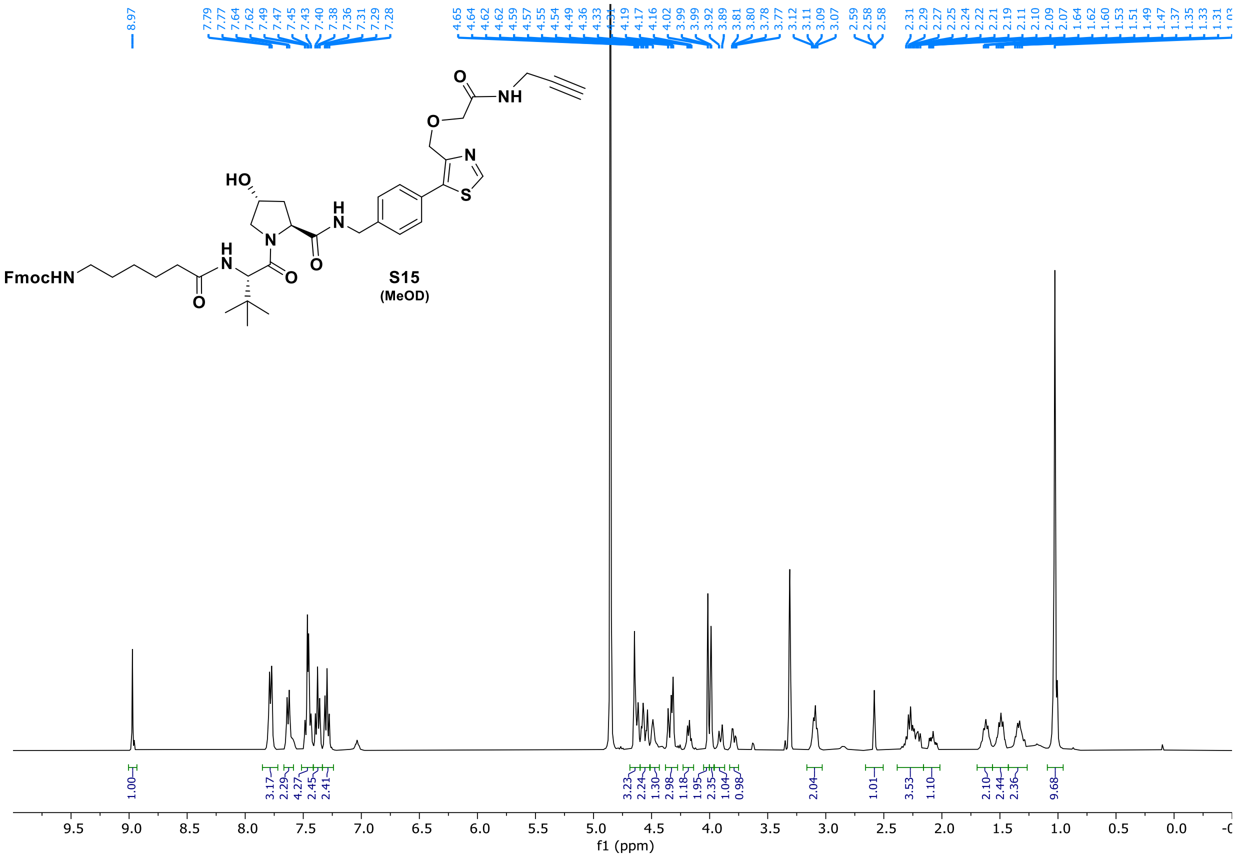

S15
(MeOD)

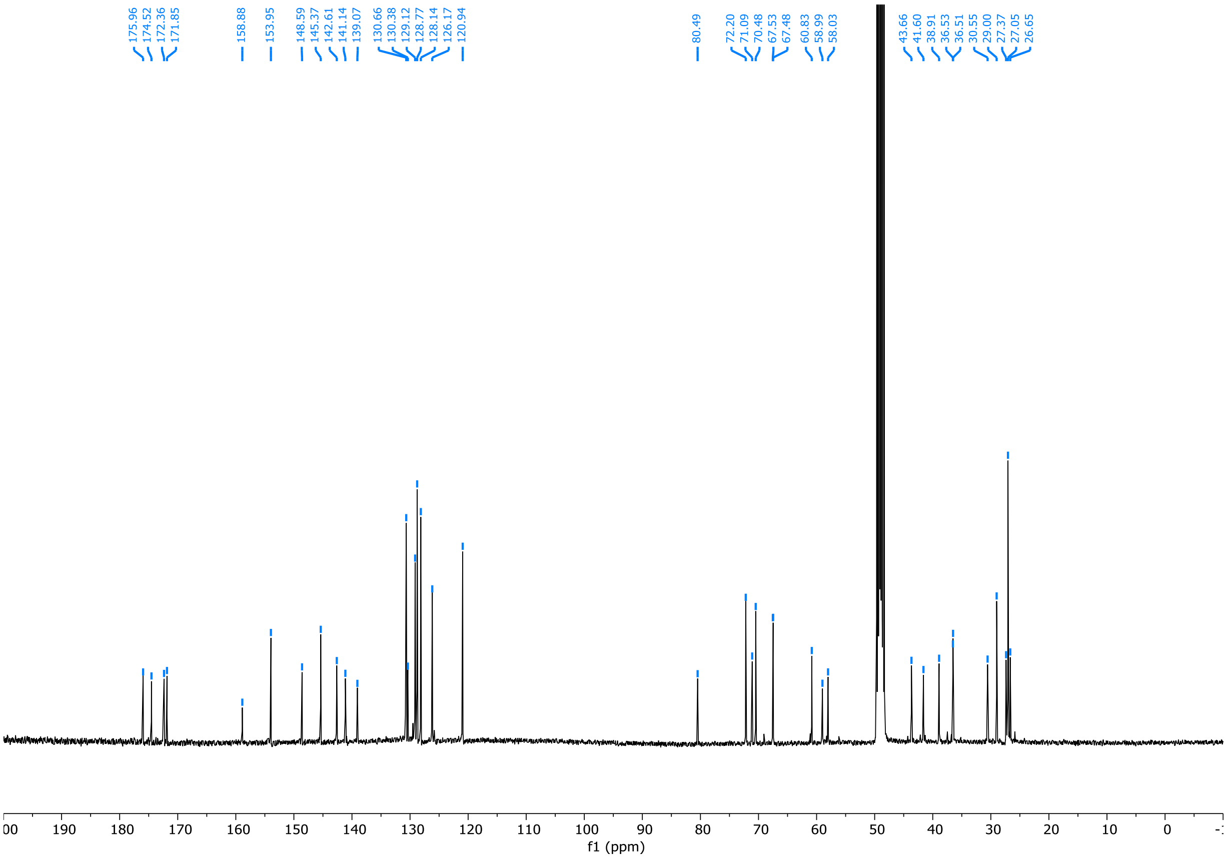

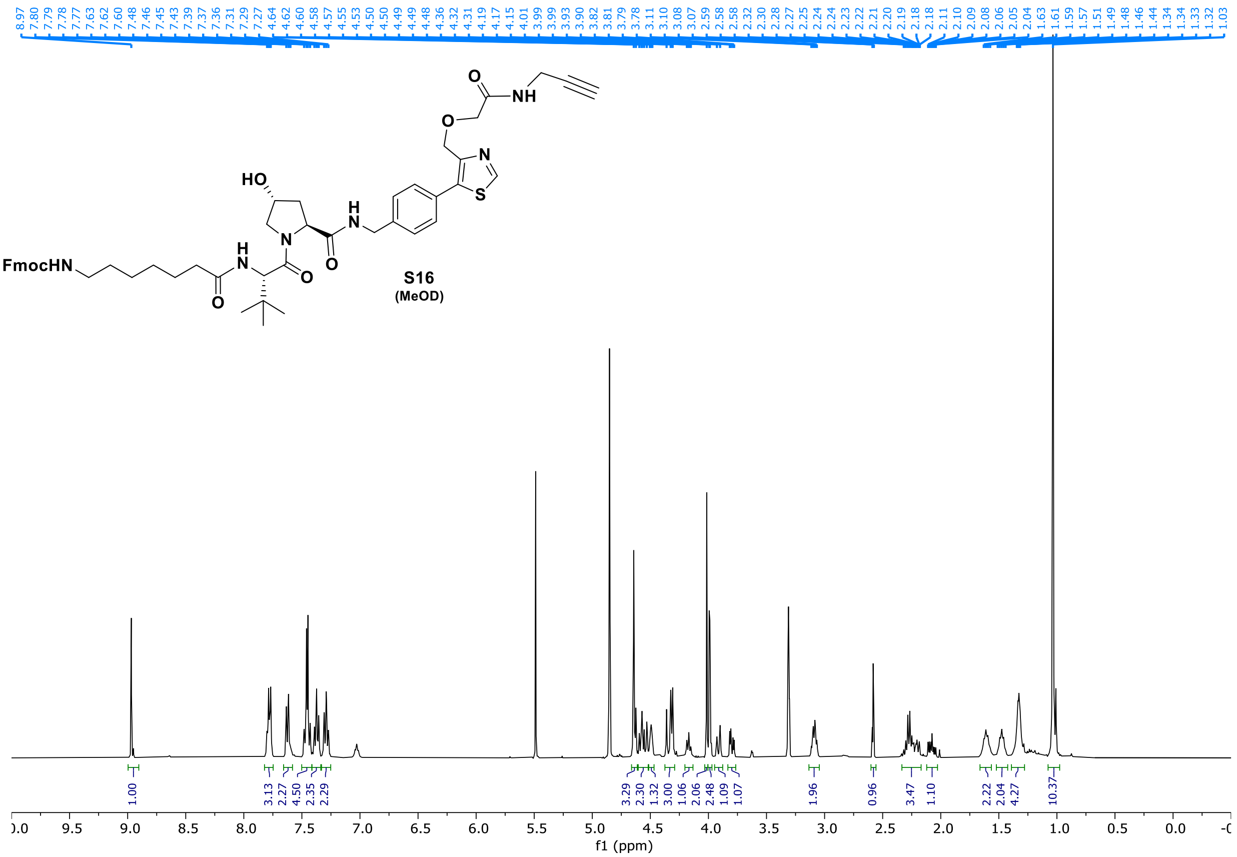
8.97 7.80 7.79 7.78 7.77 7.63 7.62 7.60 7.48 7.46 7.45 7.43 7.39 7.37 7.36 7.31 7.29 7.27 4.64 4.62 4.60 4.58 4.57 4.55 4.53 4.50 4.50 4.49 4.49 4.48 4.36 4.32 4.31 4.19 4.17 4.15 4.01 3.99 3.99 3.93 3.90 3.82 3.81 3.79 3.78 3.11 3.10 3.08 3.07 2.59 2.58 2.58 2.32 2.30 2.28 2.27 2.25 2.24 2.24 2.23 2.22 2.21 2.20 2.19 2.18 2.18 2.11 2.10 2.09 2.08 2.06 2.05 2.04 1.63 1.61 1.59 1.57 1.51 1.49 1.48 1.46 1.44 1.34 1.34 1.33 1.32 1.03
FmocHN
HO
N
H
NH
O
S
S16
(MeOD)
1.00 3.13 2.27 4.50 2.35 2.29 3.29 2.30 1.32 3.00 1.06 2.06 2.48 1.09 1.07 1.96 0.96 3.47 1.10 2.22 2.04 4.27 10.37
f1 (ppm)

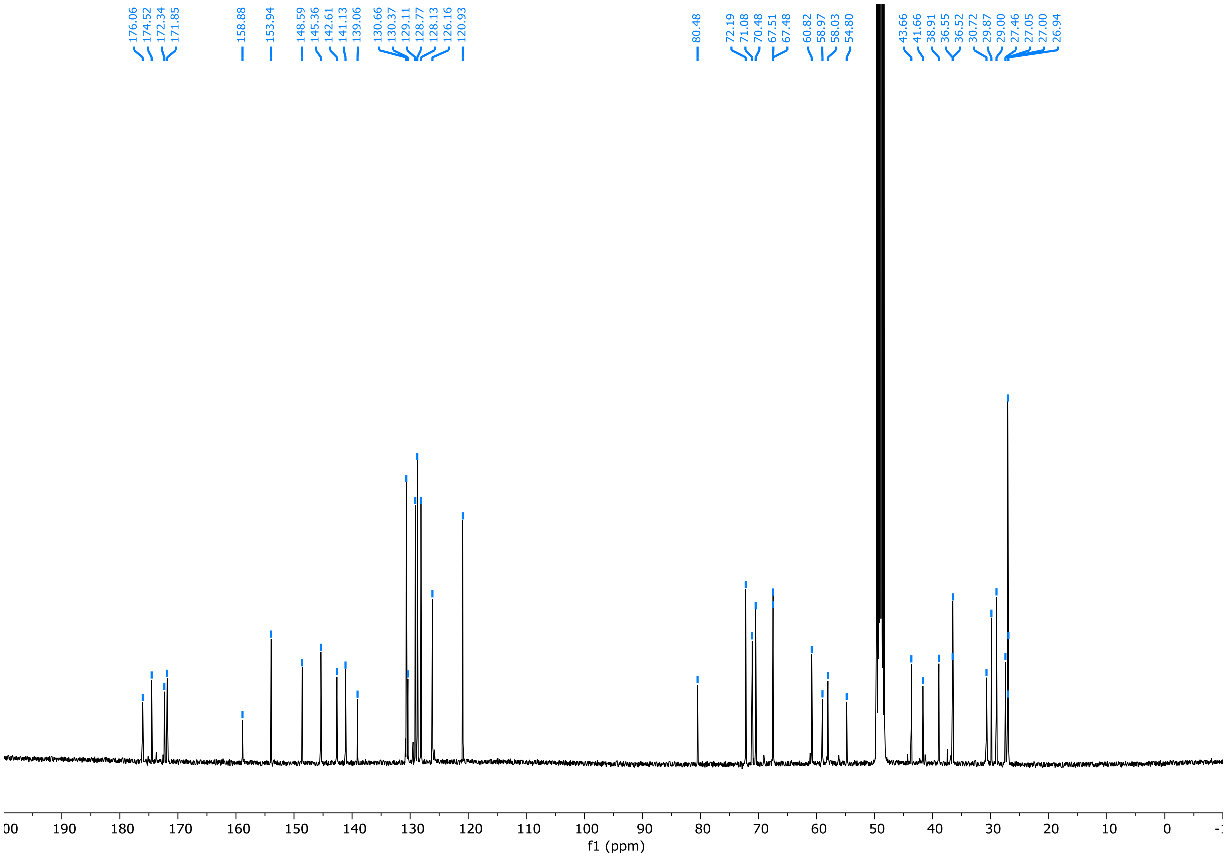
176.06 174.52 172.34 171.85 158.88 153.94 148.59 145.36 142.61 141.13 139.06 130.66 130.37 129.11 128.77 128.13 126.16 120.93 80.48 72.19 71.08 70.48 67.51 67.48 60.82 58.97 58.03 54.80 43.66 41.66 38.91 36.55 36.52 30.72 29.87 29.00 27.46 27.05 27.00 26.94
f1 (ppm)

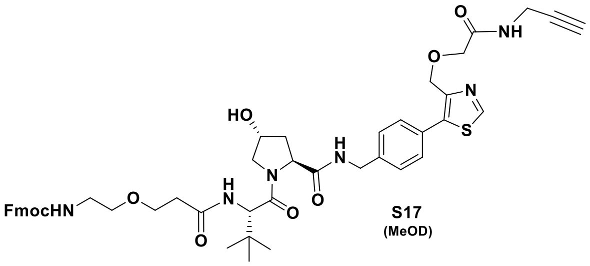
FmocHN
HO
N
H
N
O
H
N
O
N
S
O
NH

**S17**
(MeOD)

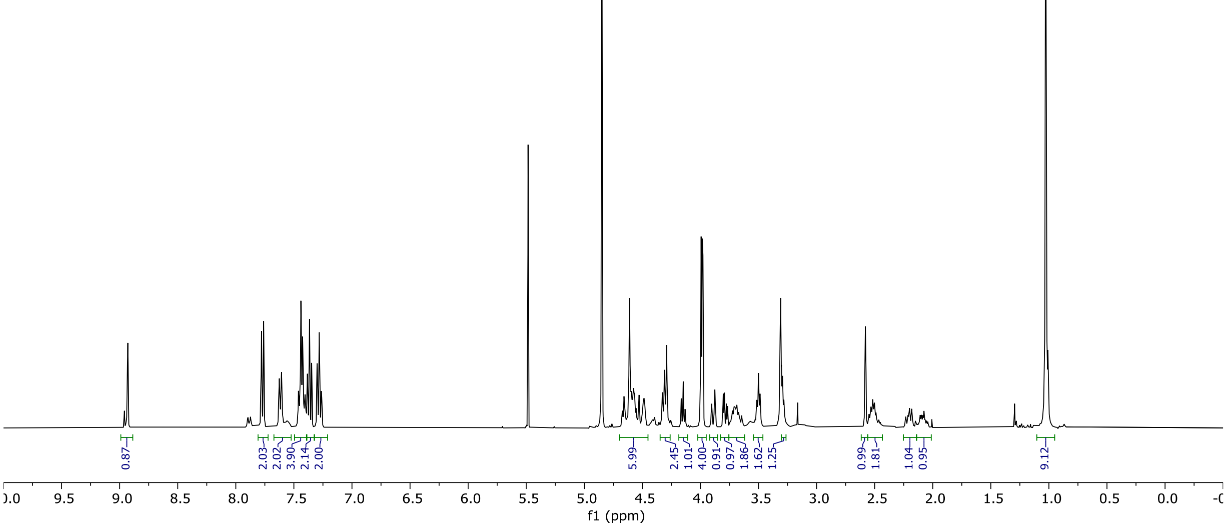
8.93 7.78 7.76 7.63 7.61 7.46 7.44 7.42 7.40 7.38 7.37 7.35 7.30 7.28 7.26 4.67 4.66 4.65 4.61 4.60 4.58 4.57 4.55 4.53 4.49 4.33 4.31 4.29 4.17 4.15 4.13 3.99 3.98 3.98 3.90 3.88 3.80 3.79 3.78 3.77 3.72 3.71 3.70 3.69 3.67 3.66 3.65 3.64 3.51 3.50 3.49 3.29 3.28
2.59 2.58 2.57 2.56 2.55 2.54 2.53 2.52 2.50 2.49 2.48 2.46 2.24 2.23 2.23 2.22 2.21 2.20 2.19 2.18 2.18 2.11 2.10 2.09 2.08 2.06
0.87 2.03 2.02 3.90 2.14 2.00 5.99 2.45 1.01 4.00 0.91 0.97 1.86 1.62 1.25 0.99 1.81 1.04 0.95 9.12
9.5 9.0 8.5 8.0 7.5 7.0 6.5 6.0 5.5 5.0 4.5 4.0 3.5 3.0 2.5 2.0 1.5 1.0 0.5 0.0
f1 (ppm)

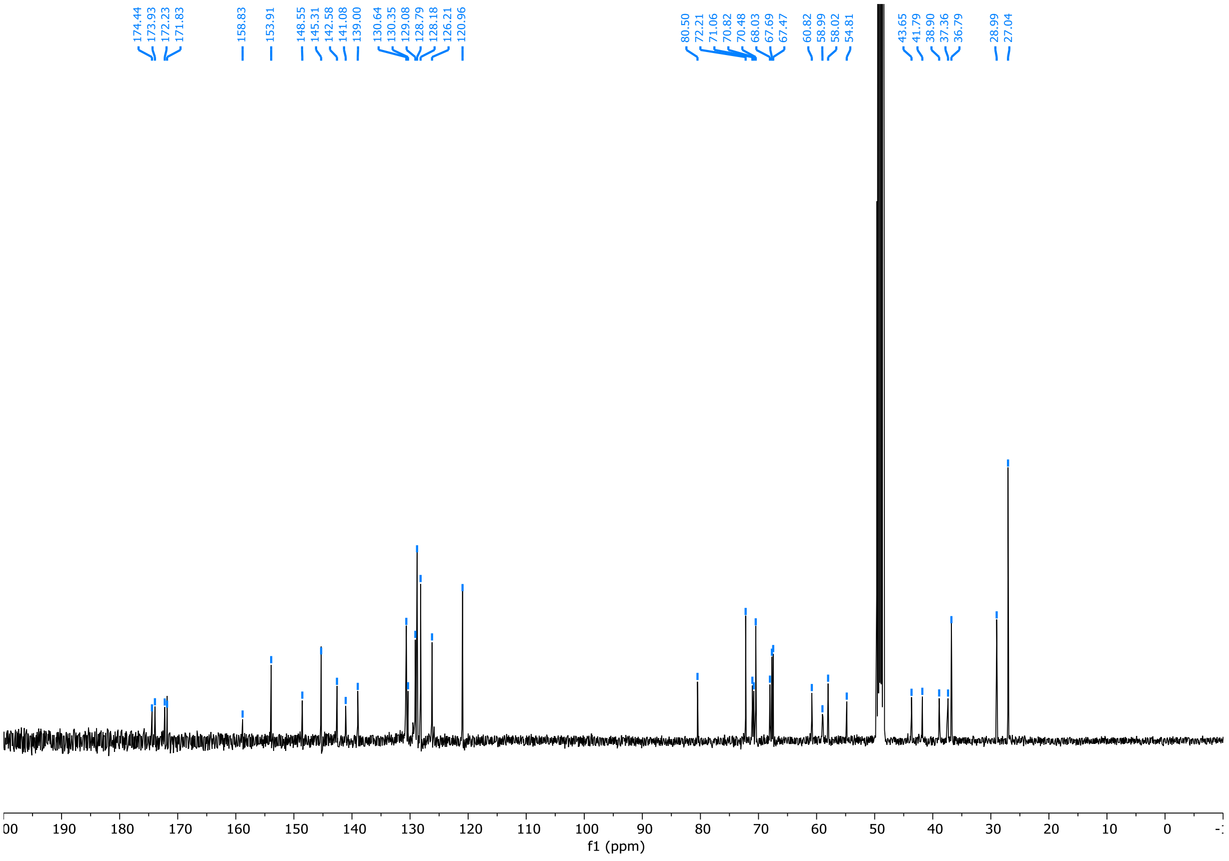
174.44 173.93 172.23 171.83 158.83 153.91 148.55 145.31 142.58 141.08 139.00 130.64 130.35 129.08 128.79 128.18 126.21 120.96
80.50 72.21 71.06 70.82 70.48 68.03 67.69 67.47 60.82 58.99 58.02 54.81 43.65 41.79 38.90 37.36 36.79 28.99 27.04
190 180 170 160 150 140 130 120 110 100 90 80 70 60 50 40 30 20 10 0
f1 (ppm)

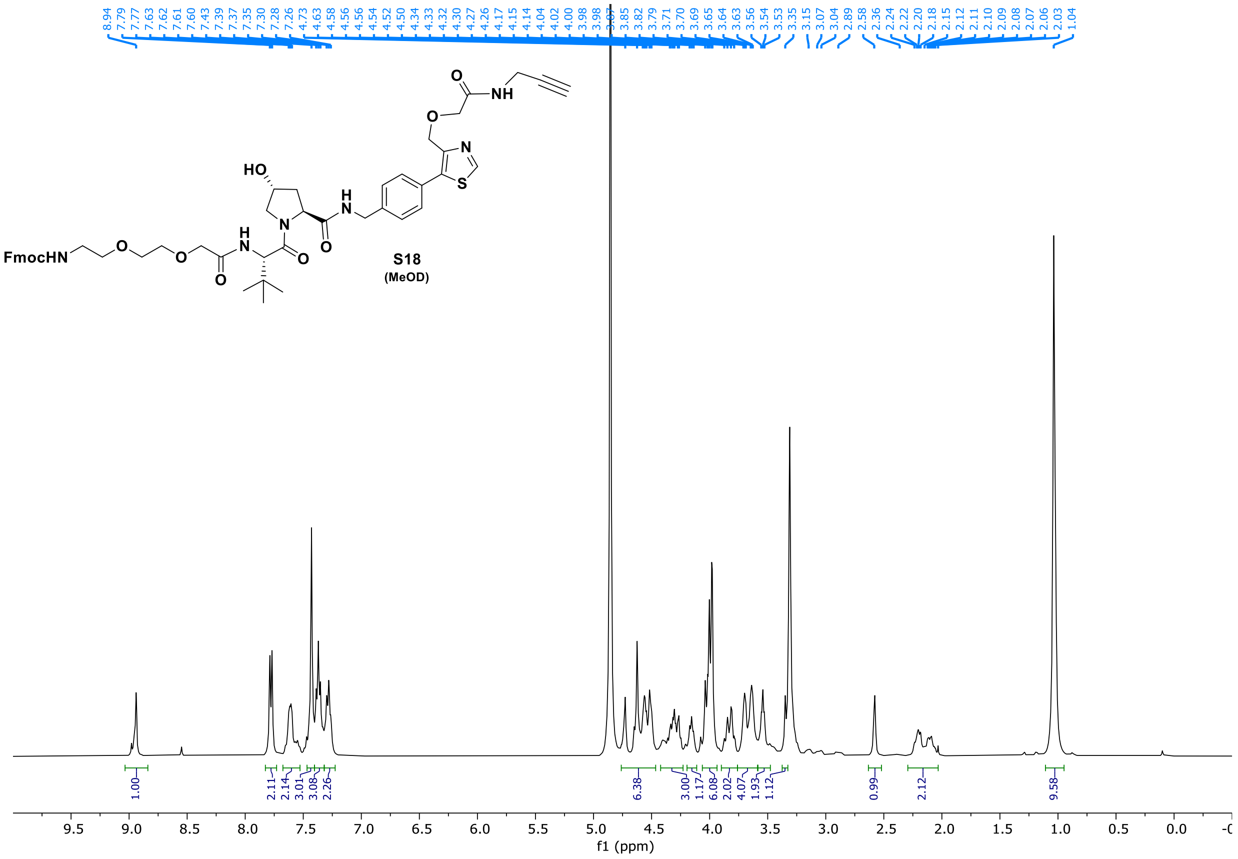

8.94 7.79 7.77 7.63 7.62 7.61 7.60 7.43 7.39 7.37 7.35 7.30 7.28 7.26 4.73 4.63 4.58 4.56 4.56 4.54 4.52 4.50 4.34 4.33 4.32 4.30 4.27 4.26 4.17 4.15 4.14 4.04 4.02 4.00 3.98 3.98 3.87 3.85 3.82 3.79 3.71 3.70 3.69 3.65 3.64 3.63 3.56 3.54 3.53 3.35 3.15 3.07 3.04 2.89 2.58 2.36 2.24 2.22 2.20 2.18 2.15 2.12 2.11 2.10 2.09 2.08 2.07 2.06 2.03 1.04
FmocHN
HO
S18
(MeOD)
1.00 2.11 2.14 3.01 3.08 2.26 6.38 3.00 1.17 6.08 2.02 4.07 1.93 1.12 0.99 2.12 9.58
f1 (ppm)


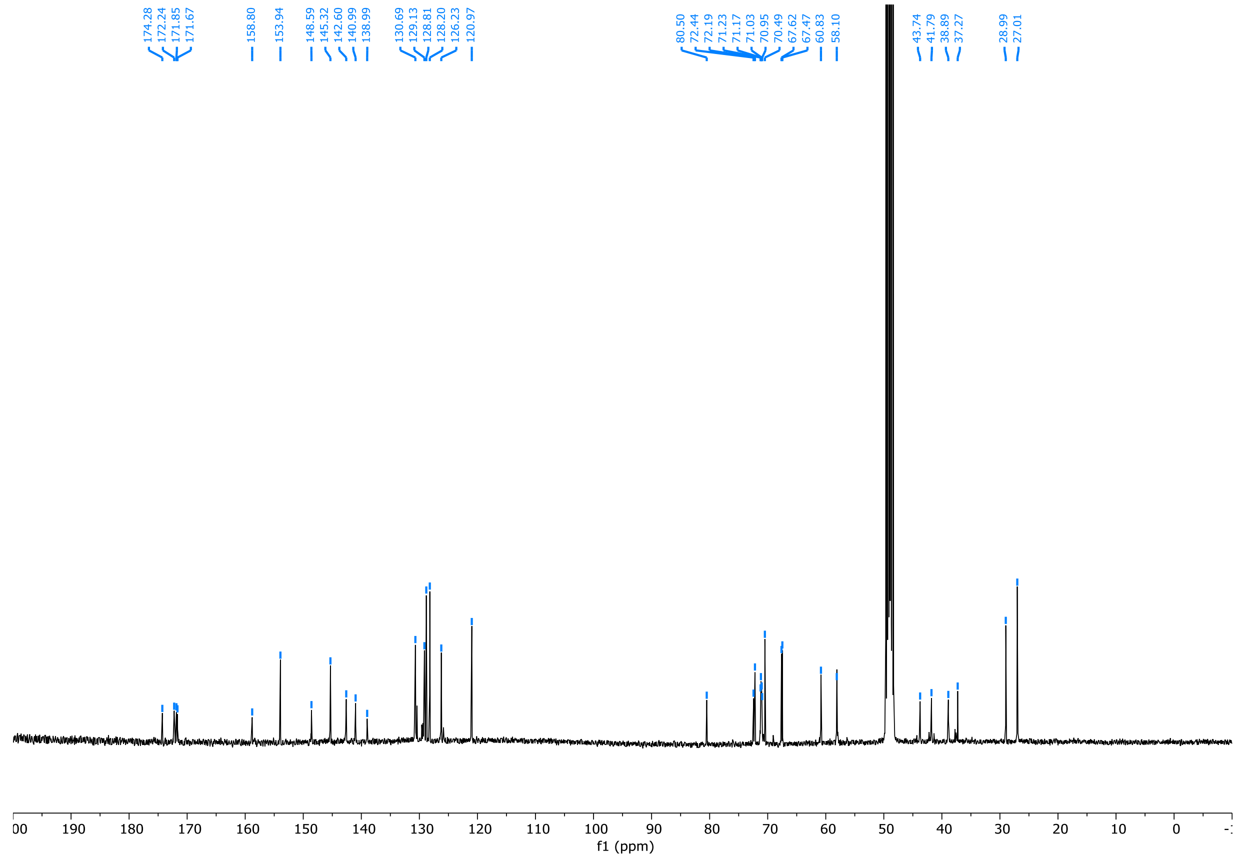

174.28 172.24 171.85 171.67 158.80 153.94 148.59 145.32 142.60 140.99 138.99 130.69 129.13 128.81 128.20 126.23 120.97 80.50 72.44 72.19 71.23 71.17 71.03 70.95 70.49 67.62 67.47 60.83 58.10 43.74 41.79 38.89 37.27 28.99 27.01
f1 (ppm)

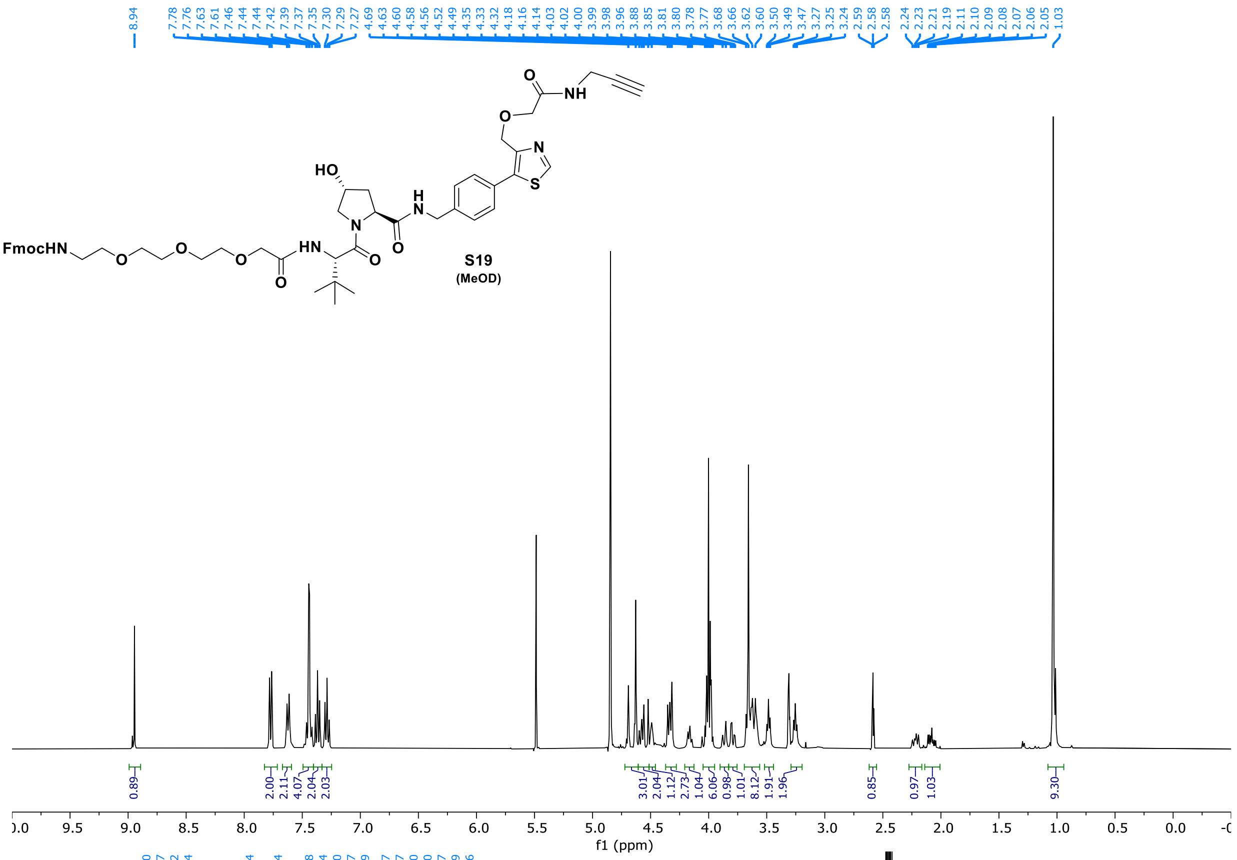
8.94
7.78 7.76 7.63 7.61 7.46 7.44 7.44 7.42 7.39 7.37 7.35 7.30 7.29 7.27
4.69 4.63 4.60 4.58 4.56 4.52 4.49 4.35 4.33 4.32 4.18 4.16 4.14 4.03 4.02 4.00 3.99 3.98 3.96 3.88 3.85 3.81 3.80 3.78 3.77 3.68 3.66 3.62 3.60 3.50 3.49 3.47 3.27 3.25 3.24
2.59 2.58 2.58
2.24 2.23 2.21 2.19 2.11 2.10 2.09 2.08 2.07 2.06 2.05
1.03
HO
O
NH
O
N
S
N
H
N
O
FmocHN
O
O
O
HN
O
O
S19
(MeOD)
0.89 2.00 2.11 4.07 2.04 2.03 3.01 2.04 1.12 2.73 1.04 6.06 0.98 1.01 8.12 1.91 1.96 0.85 0.97 1.03 9.30
f1 (ppm)

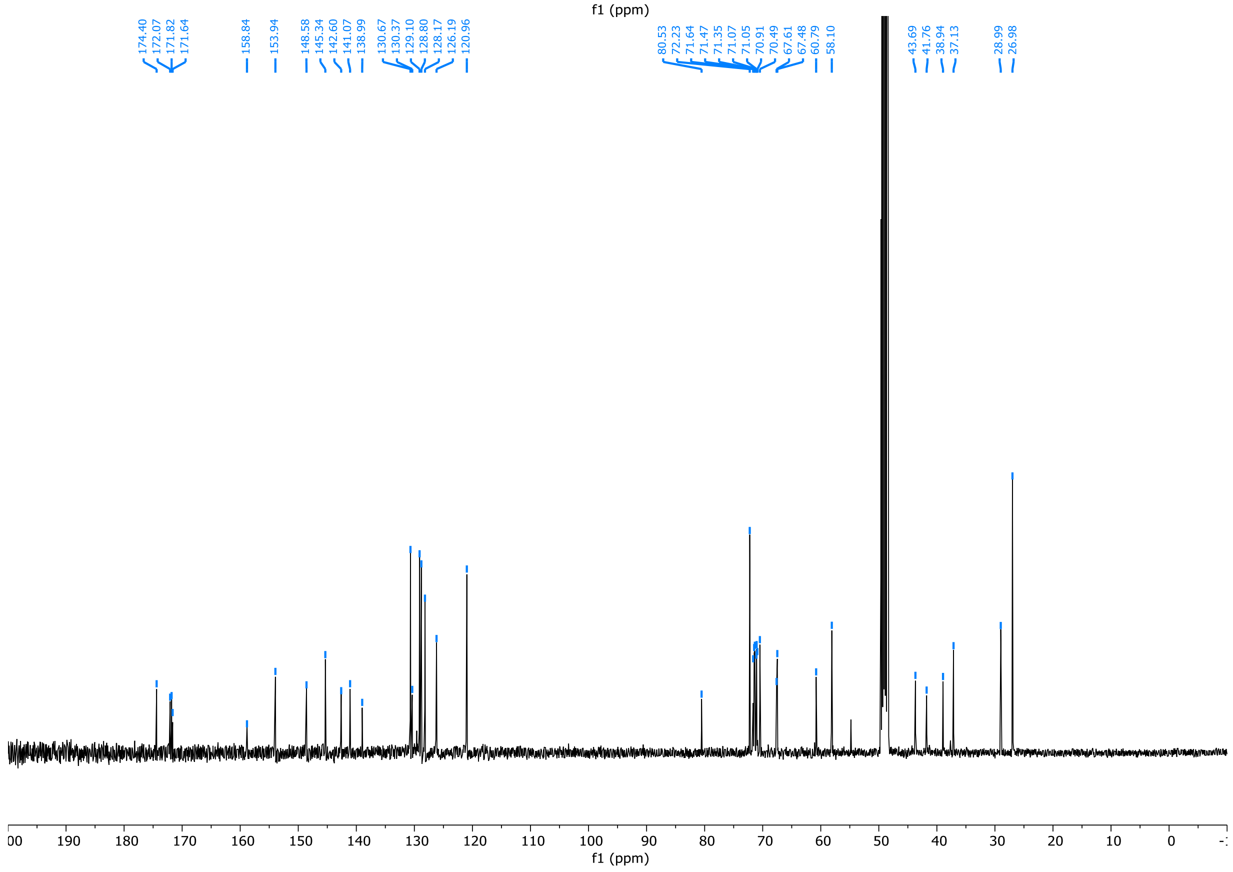
174.40 172.07 171.82 171.64
158.84
153.94
148.58 145.34 142.60 141.07 138.99
130.67 130.37 129.10 128.80 128.17 126.19 120.96
80.53 72.23 71.64 71.47 71.35 71.07 71.05 70.91 70.49 67.61 67.48 60.79 58.10
43.69 41.76 38.94 37.13
28.99 26.98
f1 (ppm)

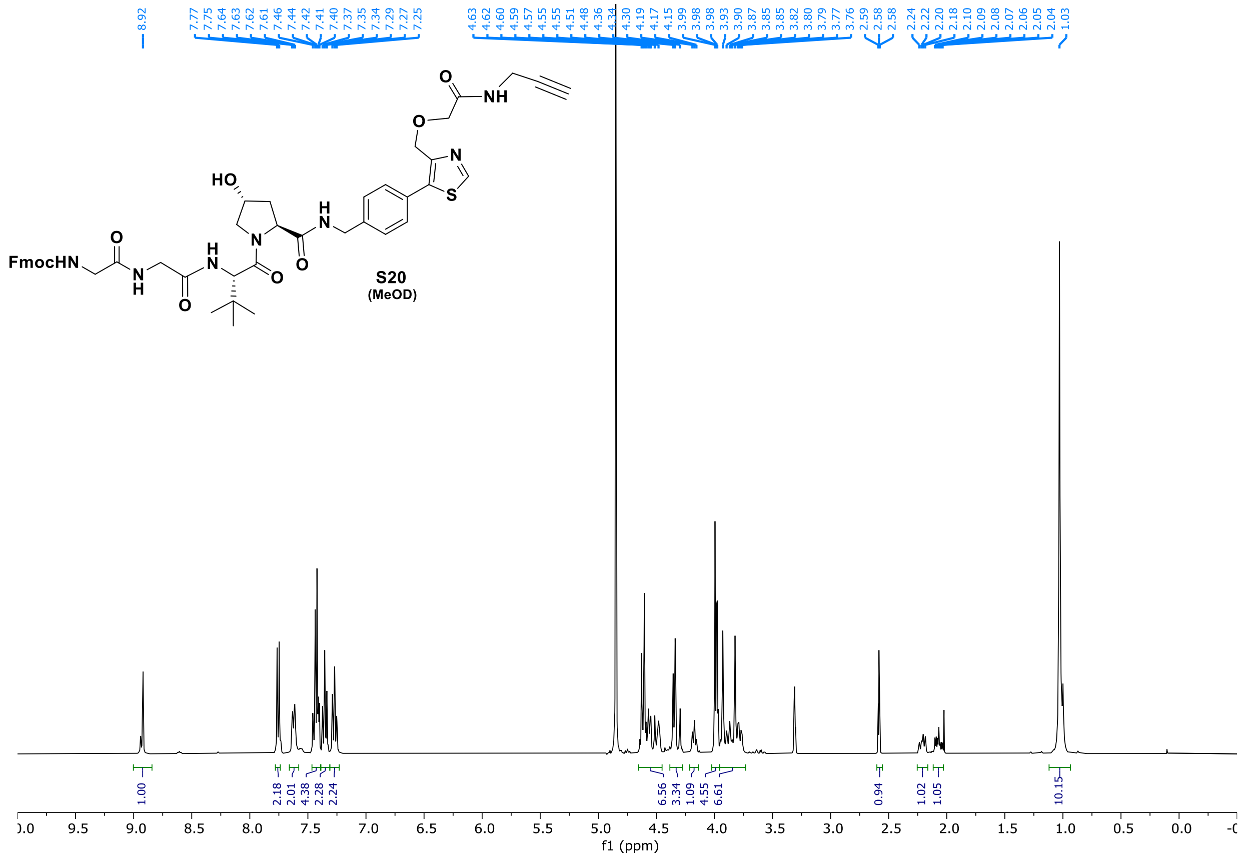
S20
(MeOD)
f1 (ppm)

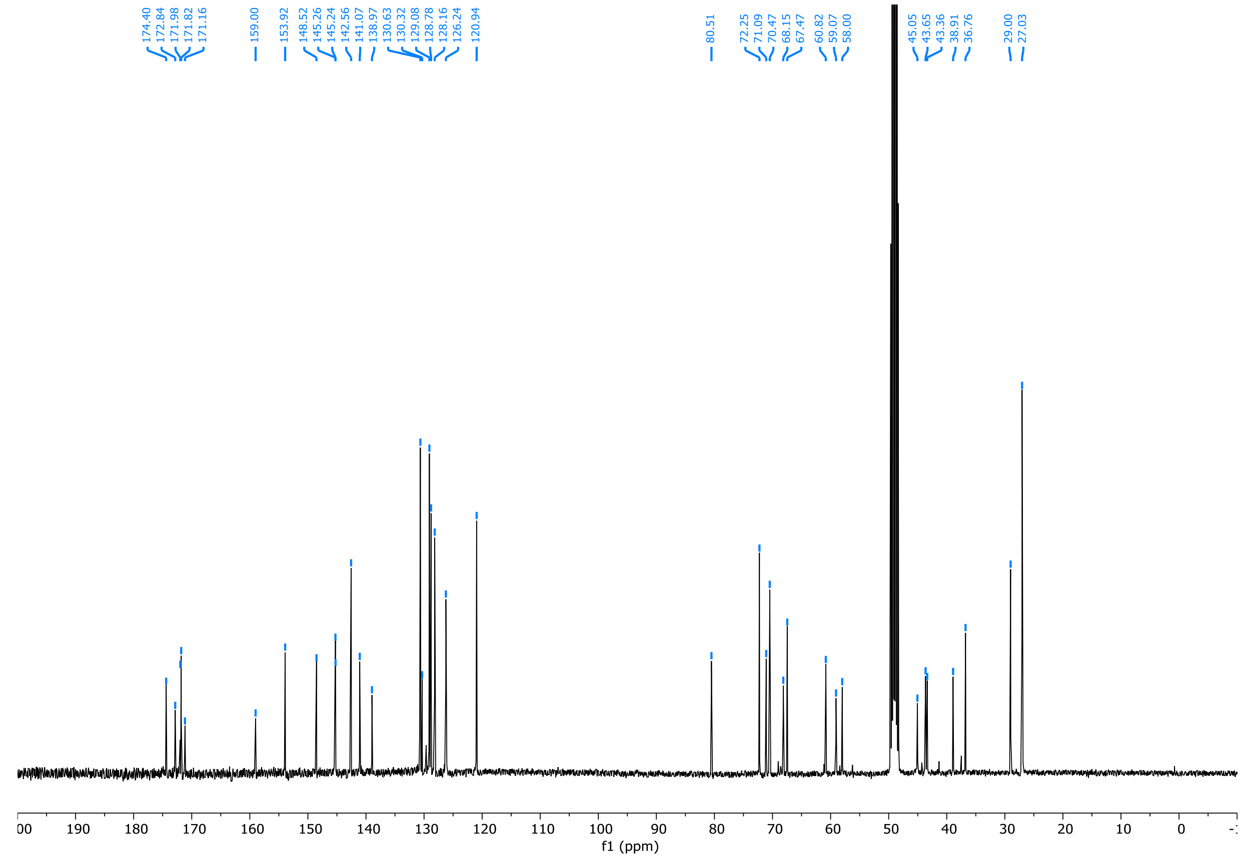
f1 (ppm)

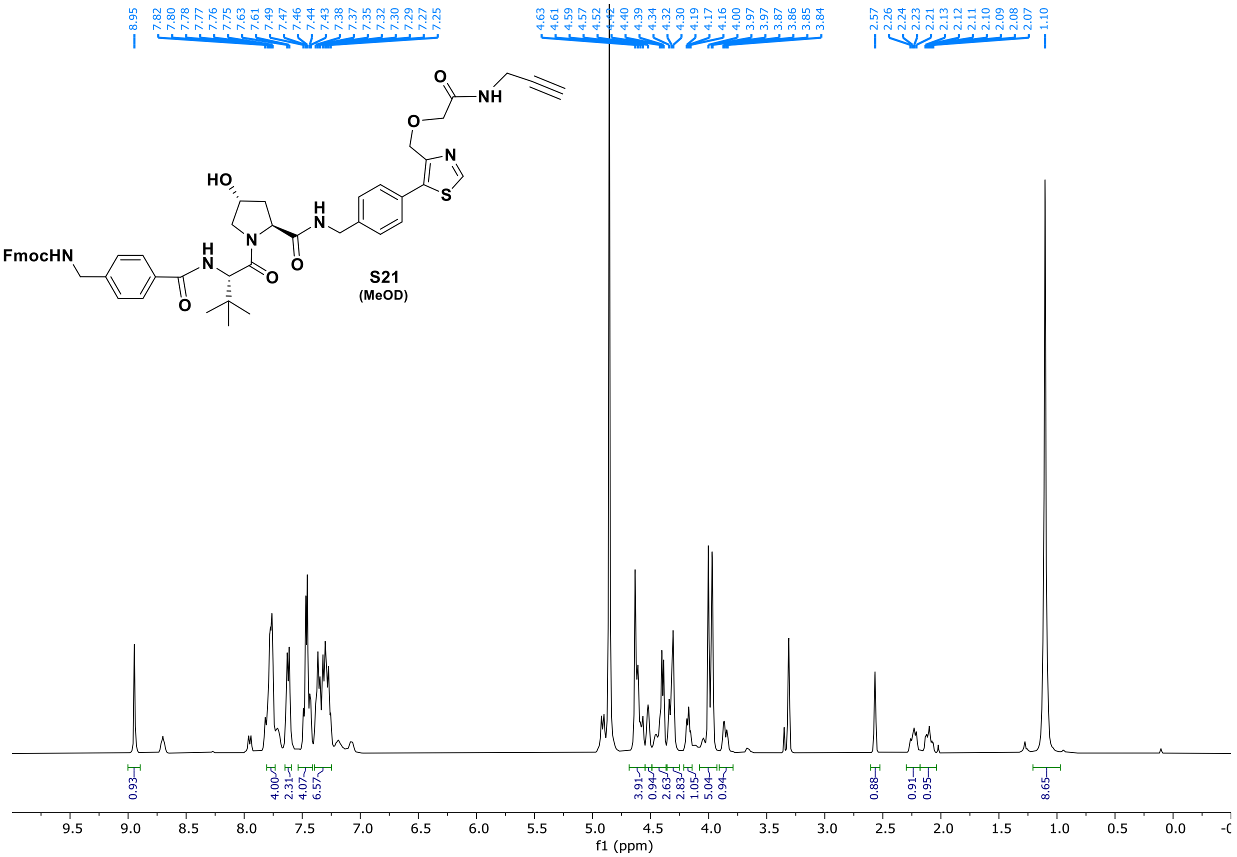
FmocHN
HO
S21
(MeOD)
8.95 7.82 7.80 7.78 7.77 7.76 7.75 7.63 7.61 7.49 7.47 7.46 7.44 7.43 7.38 7.37 7.35 7.32 7.30 7.29 7.27 7.25
4.63 4.61 4.59 4.57 4.52 4.42 4.40 4.39 4.34 4.32 4.30 4.19 4.17 4.16 4.00 3.97 3.97 3.87 3.86 3.85 3.84
2.57 2.26 2.24 2.23 2.21 2.13 2.12 2.11 2.10 2.09 2.08 2.07 1.10
0.93 4.00 2.31 4.07 6.57 3.91 0.94 2.63 2.83 1.05 5.04 0.94 0.88 0.91 0.95 8.65
f1 (ppm)

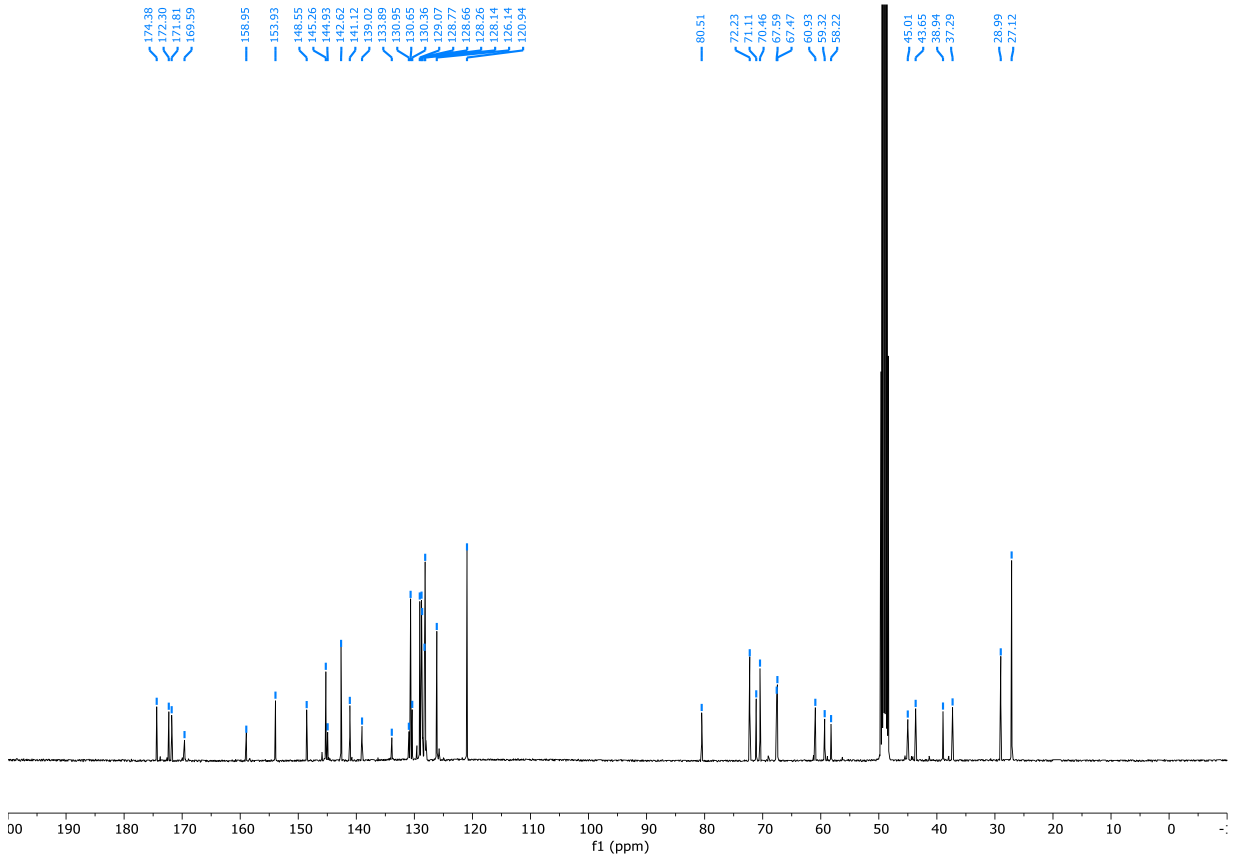
174.38 172.30 171.81 169.59 158.95 153.93 148.55 145.26 144.93 142.62 141.12 139.02 133.89 130.95 130.65 130.36 129.07 128.77 128.66 128.26 128.14 126.14 120.94
80.51 72.23 71.11 70.46 67.59 67.47 60.93 59.32 58.22 45.01 43.65 38.94 37.29 28.99 27.12
f1 (ppm)

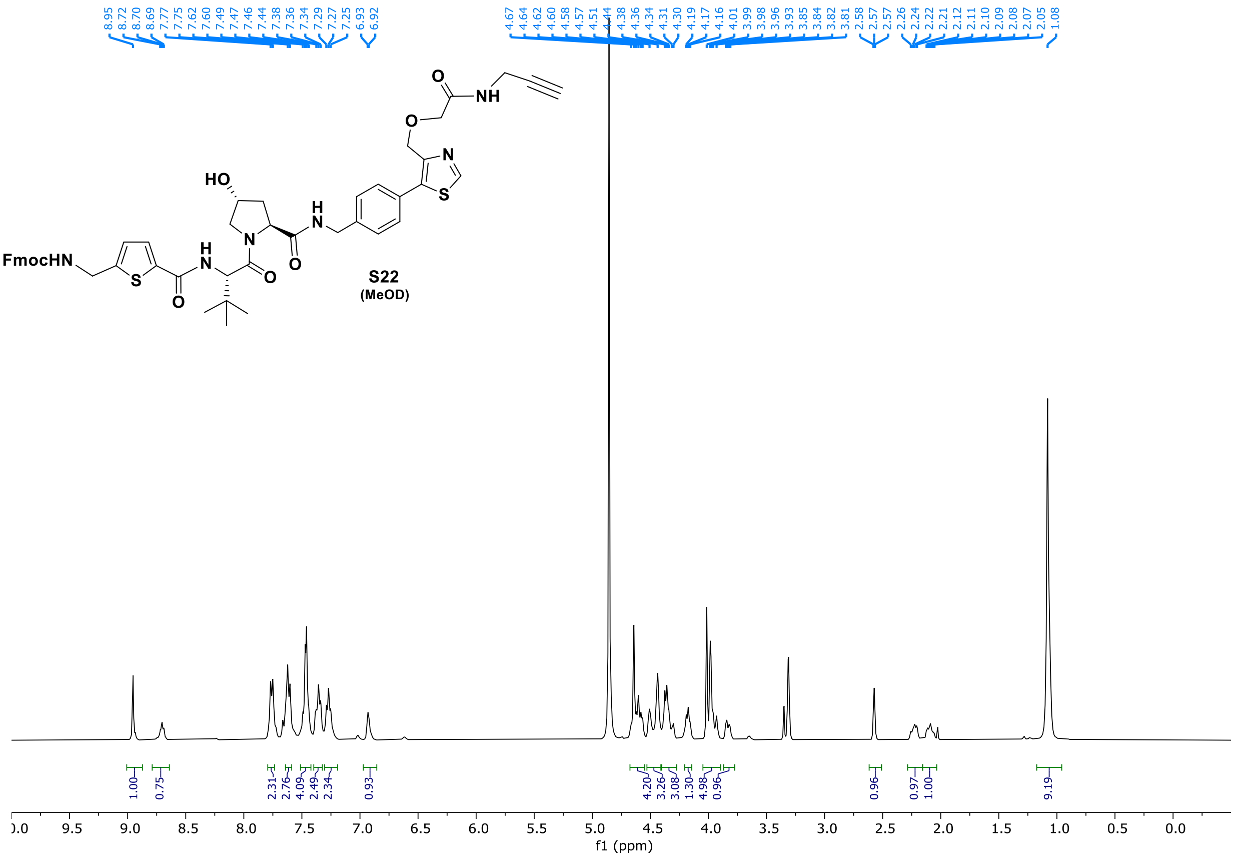
FmocHN
HO
S22
(MeOD)
f1 (ppm)

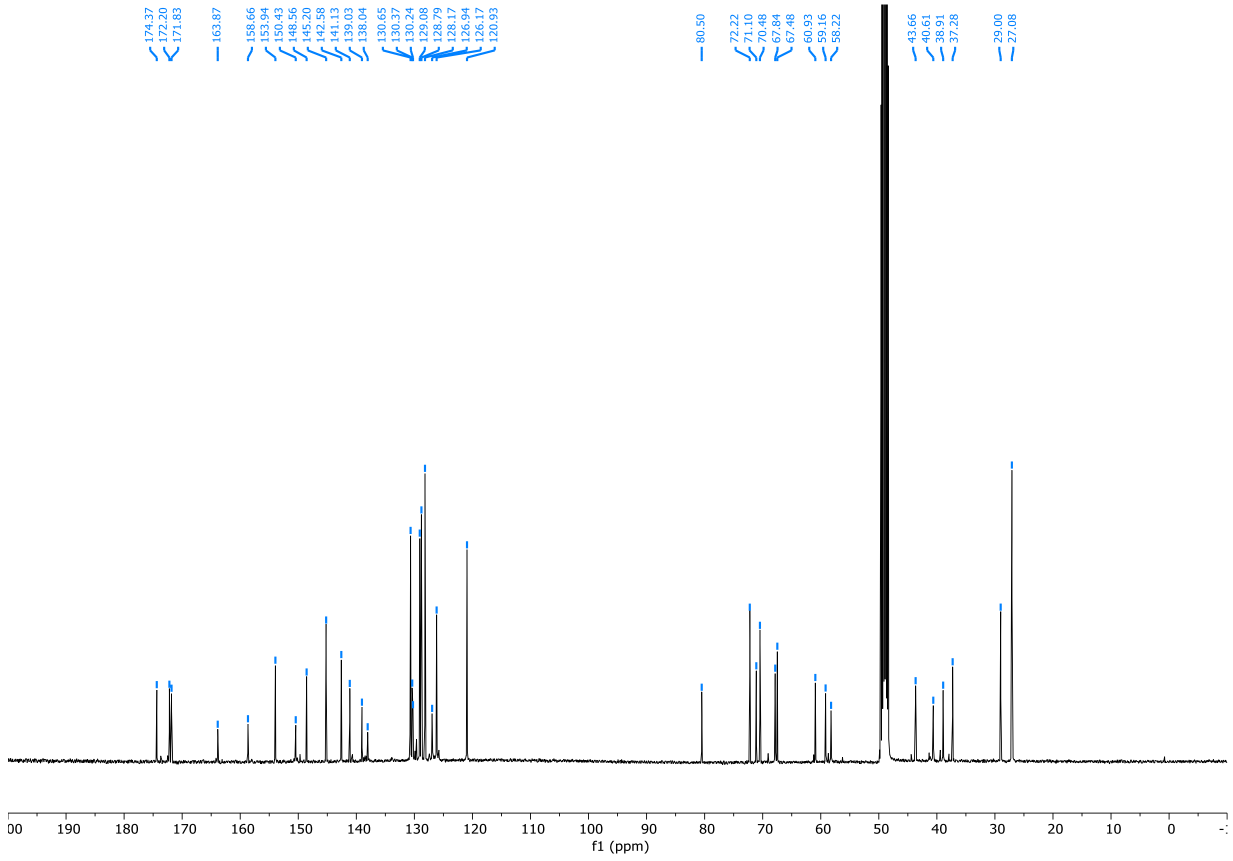
f1 (ppm)

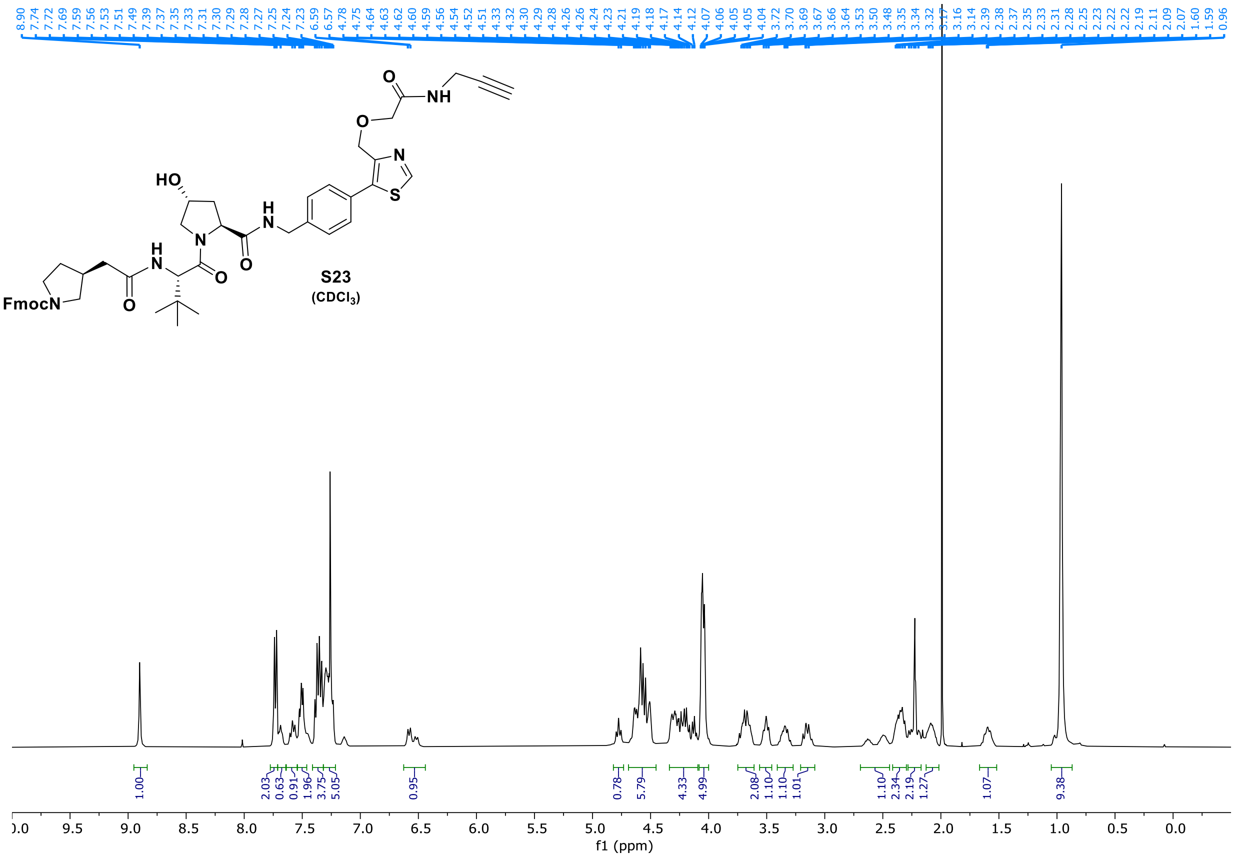
HO
FmocN
S23
(CDCl3)
f1 (ppm)

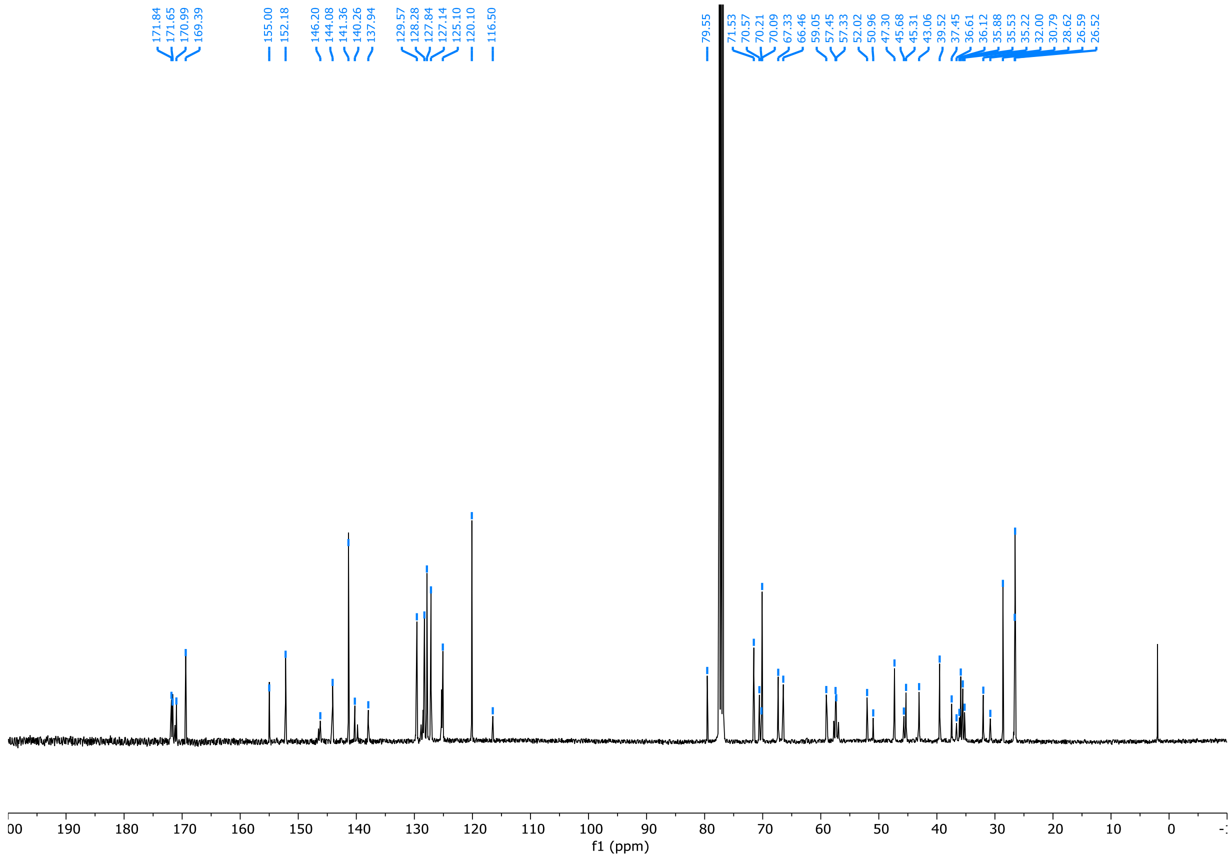
f1 (ppm)

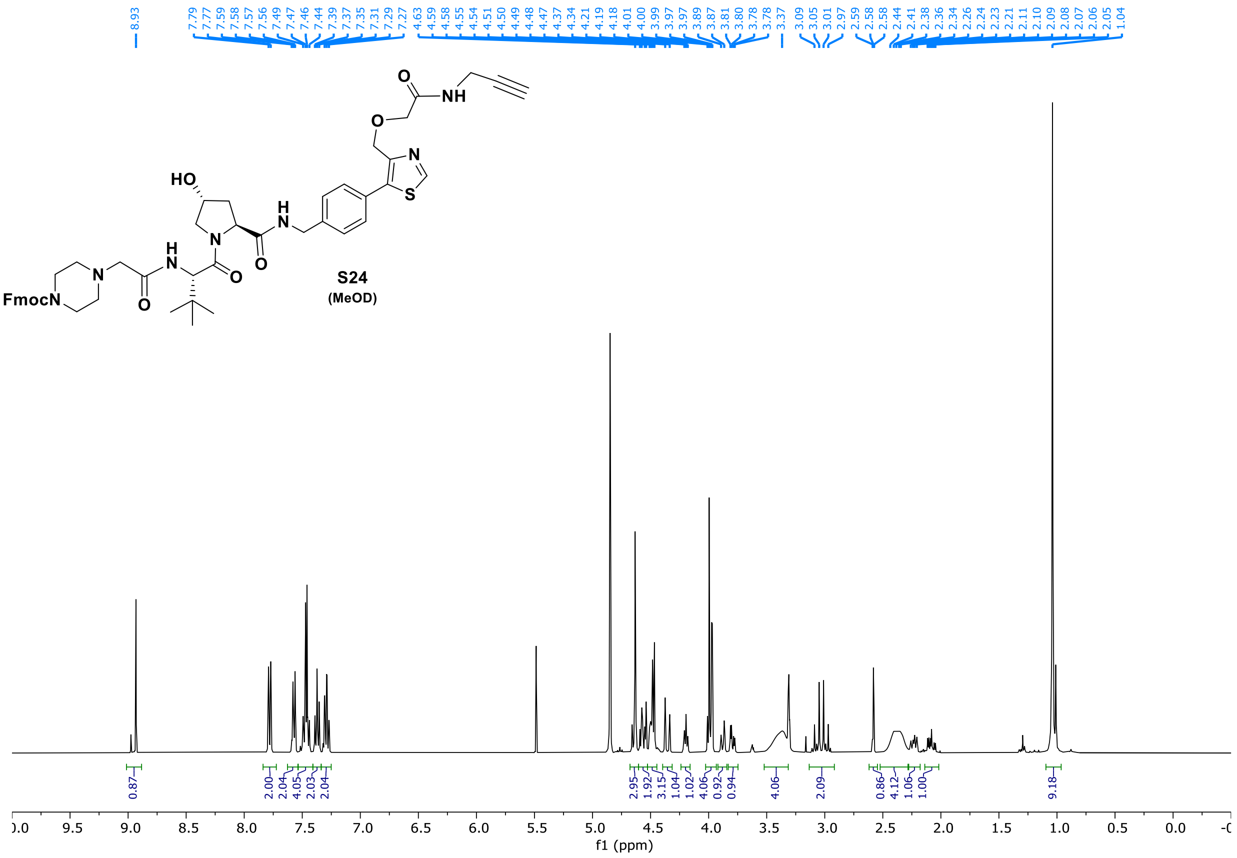
8.93
7.79 7.77 7.59 7.58 7.57 7.56 7.49 7.47 7.46 7.44 7.39 7.37 7.35 7.31 7.29 7.27
4.63 4.59 4.58 4.55 4.54 4.51 4.50 4.49 4.48 4.47 4.37 4.34 4.21 4.19 4.18 4.01 4.00 3.99 3.97 3.97 3.89 3.87 3.81 3.80 3.78 3.78
3.37
3.09 3.05 3.01 2.97 2.59 2.58 2.58 2.44 2.41 2.38 2.36 2.34 2.26 2.24 2.23 2.21 2.11 2.10 2.09 2.08 2.07 2.06 2.05 1.04
HO
FmocN
S24
(MeOD)
0.87 2.00 2.04 4.05 2.03 2.04 2.95 1.92 3.15 1.04 1.02 4.06 0.92 0.94 4.06 2.09 0.86 4.12 1.06 1.00 9.18
f1 (ppm)

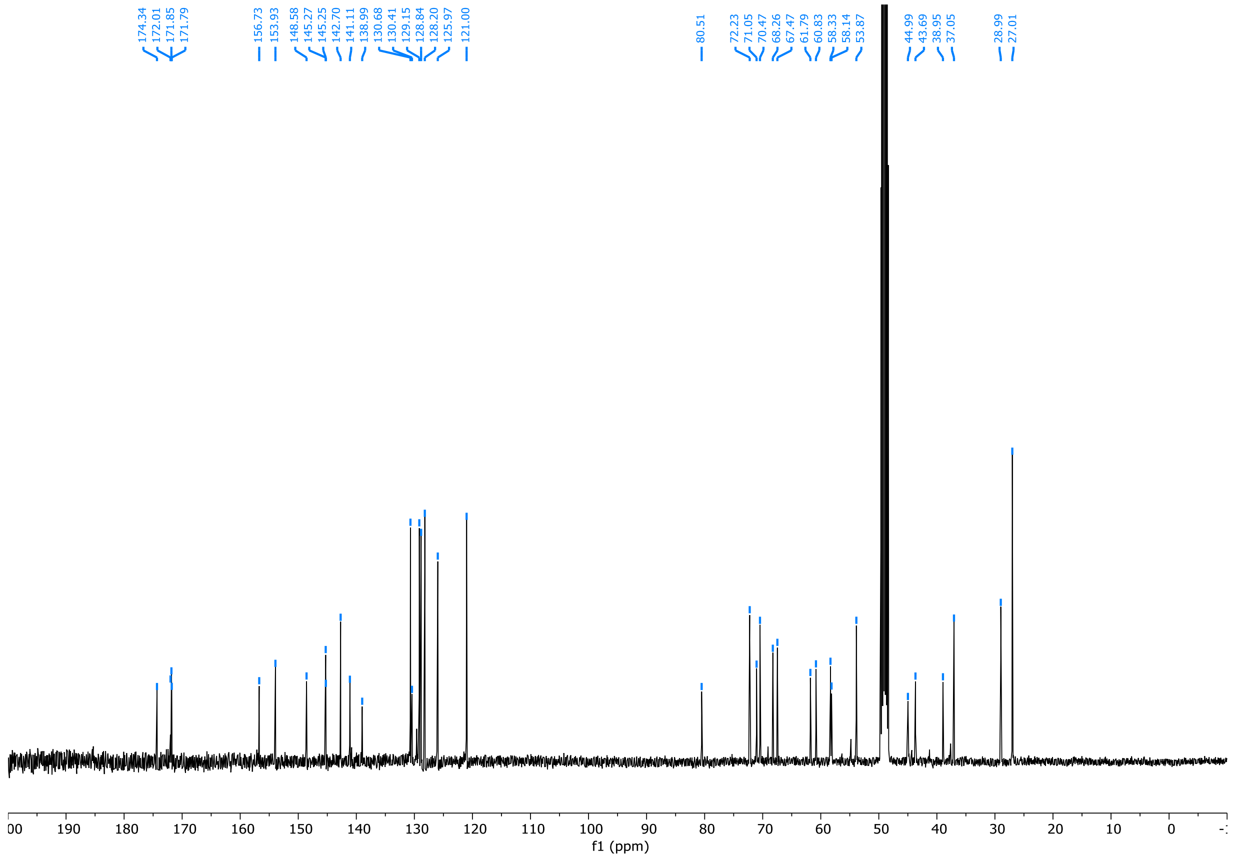
174.34 172.01 171.85 171.79
156.73 153.93 148.58 145.27 145.25 142.70 141.11 138.99 130.68 130.41 129.15 128.84 128.20 125.97 121.00
80.51
72.23 71.05 70.47 68.26 67.47 61.79 60.83 58.33 58.14 53.87
44.99 43.69 38.95 37.05
28.99 27.01
f1 (ppm)

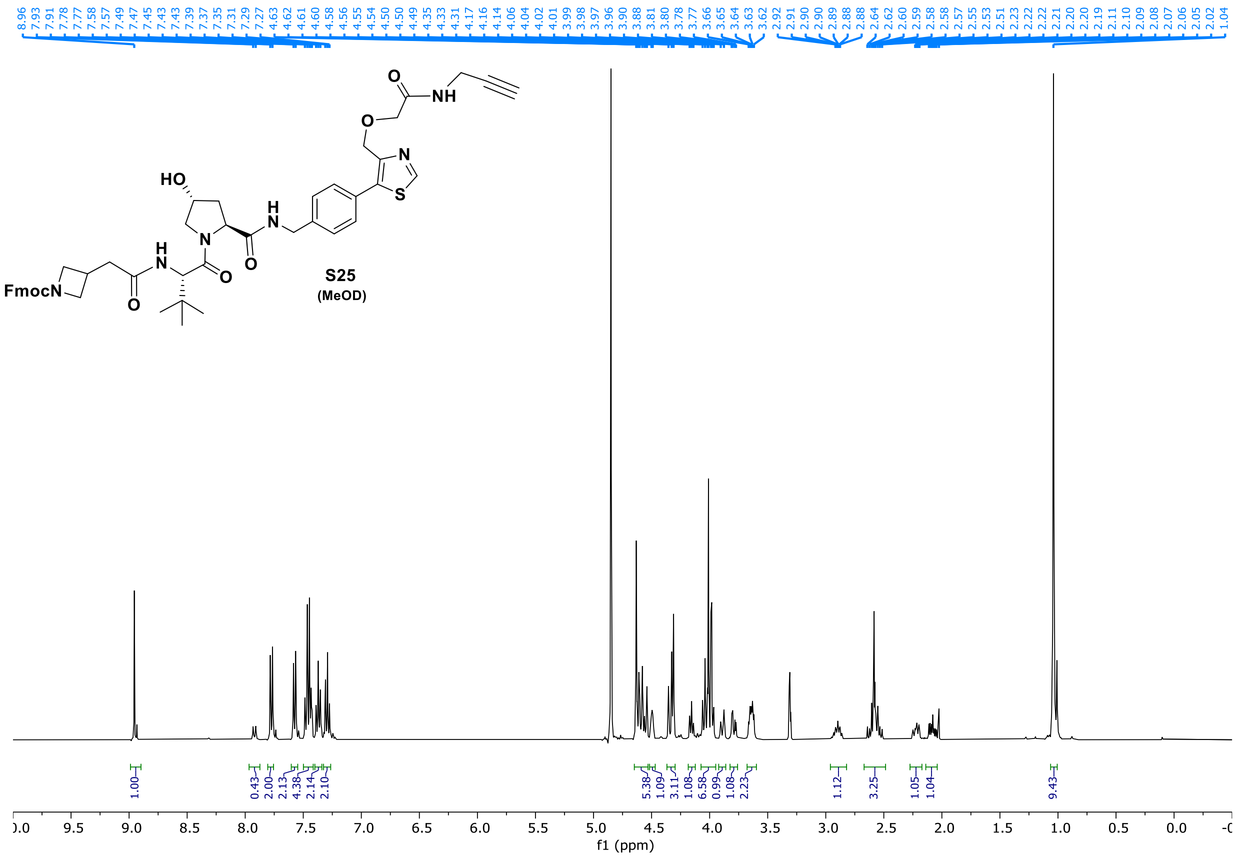

8.96 7.93 7.91 7.78 7.77 7.58 7.57 7.49 7.47 7.45 7.43 7.43 7.39 7.37 7.35 7.31 7.29 7.27 4.63 4.62 4.61 4.60 4.58 4.56 4.55 4.54 4.50 4.50 4.49 4.35 4.33 4.31 4.17 4.16 4.14 4.06 4.04 4.02 4.01 3.99 3.98 3.97 3.96 3.90 3.88 3.81 3.80 3.78 3.77 3.66 3.65 3.64 3.63 3.62 2.92 2.91 2.90 2.90 2.89 2.88 2.88 2.64 2.62 2.60 2.59 2.58 2.58 2.57 2.55 2.53 2.51 2.23 2.22 2.22 2.21 2.20 2.20 2.19 2.11 2.10 2.09 2.08 2.07 2.06 2.05 2.02 1.04
HO
H
N
O
NH
O
N
S
FmocN
S25
(MeOD)
1.00
0.43 2.00 2.13 4.38 2.14 2.10
5.38 1.09 3.11 1.08 6.58 0.99 1.08 2.23
1.12
3.25
1.05 1.04
9.43
f1 (ppm)


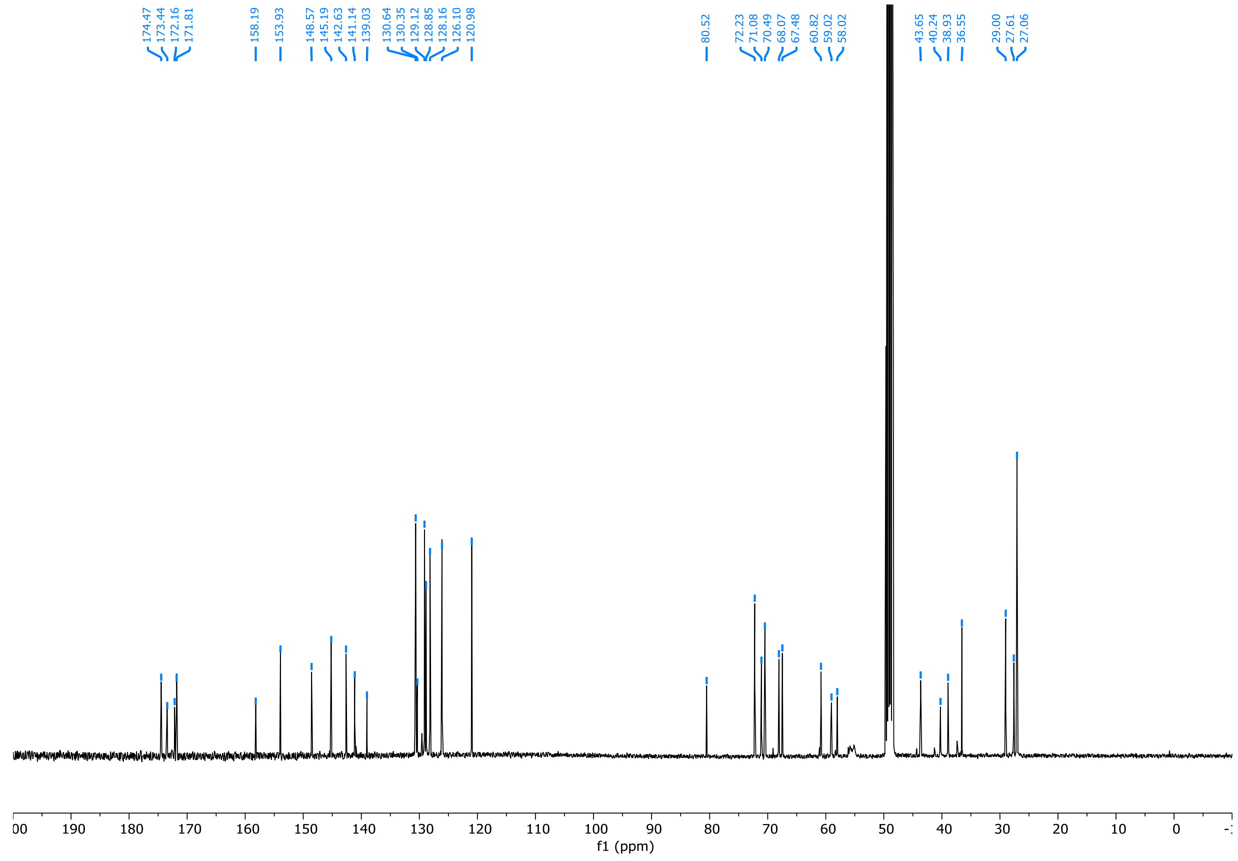

174.47 173.44 172.16 171.81
158.19 153.93
148.57 145.19 142.63 141.14 139.03
130.64 130.35 129.12 128.85 128.16 126.10 120.98
80.52
72.23 71.08 70.49 68.07 67.48
60.82 59.02 58.02
43.65 40.24 38.93 36.55
29.00 27.61 27.06
f1 (ppm)

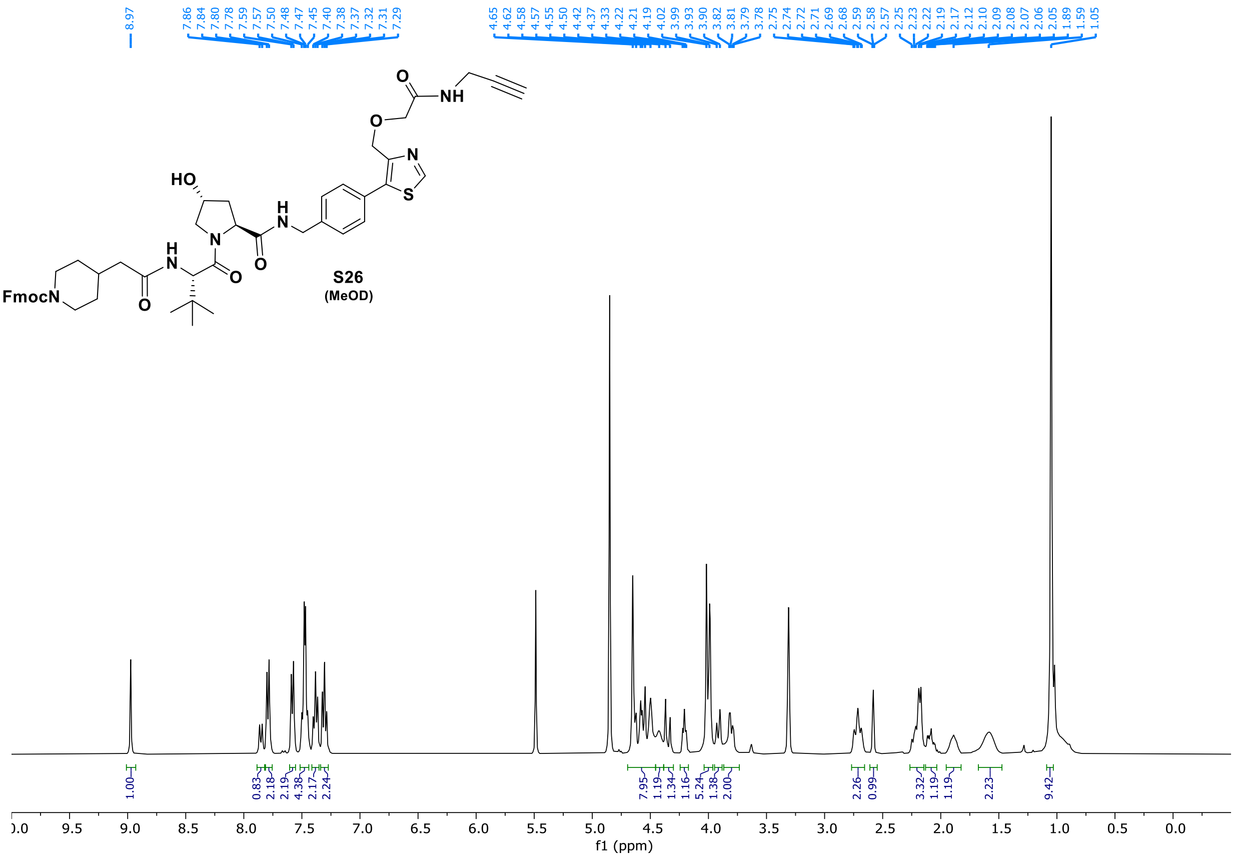
8.97
7.86 7.84 7.80 7.78 7.59 7.57 7.50 7.48 7.47 7.45 7.40 7.38 7.37 7.32 7.31 7.29
4.65 4.62 4.58 4.57 4.55 4.50 4.42 4.37 4.33 4.22 4.21 4.19 4.02 3.99 3.93 3.90 3.82 3.81 3.79 3.78 2.75 2.74 2.72 2.71 2.69 2.68 2.59 2.58 2.57 2.25 2.23 2.22 2.19 2.17 2.12 2.10 2.09 2.08 2.07 2.06 2.05 1.89 1.59 1.05
HO
H
N
O
NH
N
S
FmocN
S26
(MeOD)
1.00 0.83 2.18 2.19 4.38 2.17 2.24 7.95 1.19 1.34 1.16 5.24 1.38 2.00 2.26 0.99 3.32 1.19 1.19 2.23 9.42
f1 (ppm)

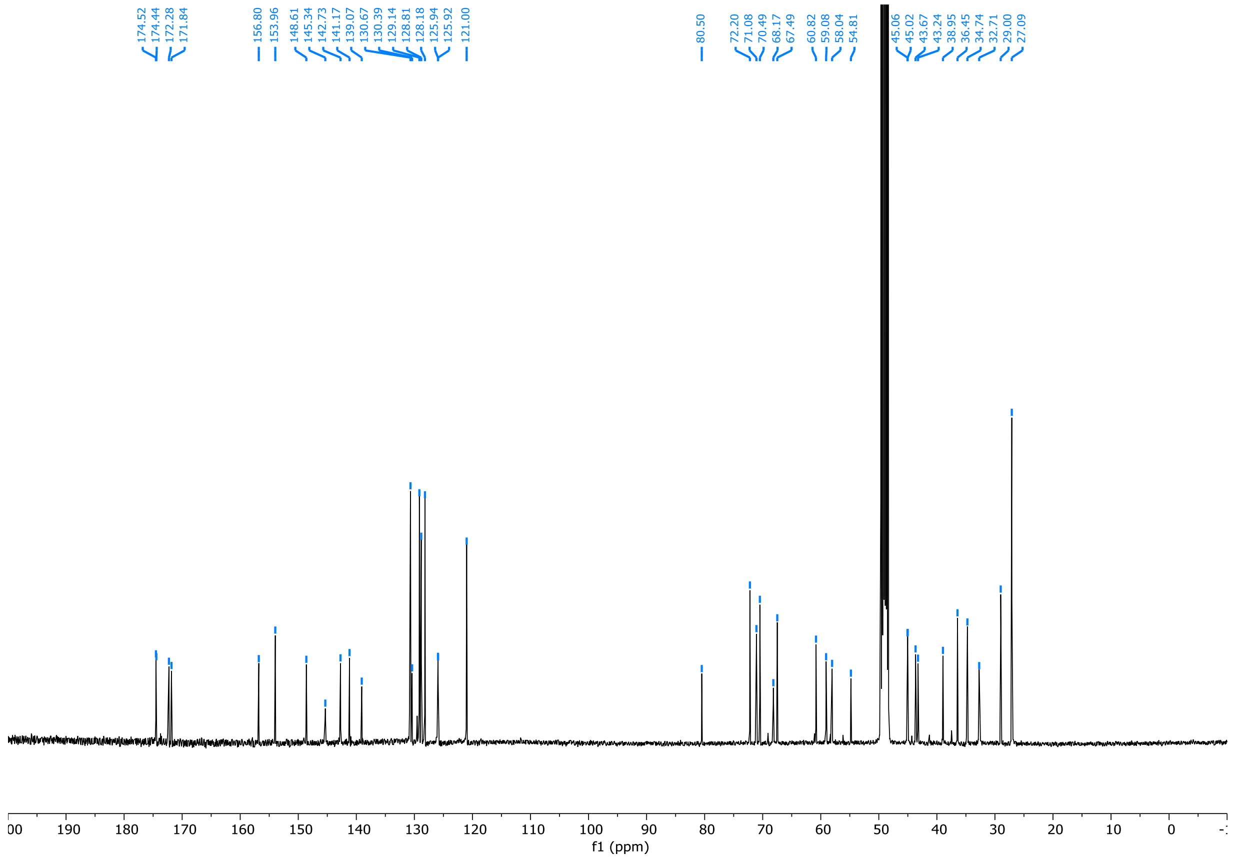
174.52 174.44 172.28 171.84 156.80 153.96 148.61 145.34 142.73 141.17 139.07 130.67 130.39 129.14 128.81 128.18 125.94 125.92 121.00
80.50 72.20 71.08 70.49 68.17 67.49 60.82 59.08 58.04 54.81 45.06 45.02 43.67 43.24 38.95 36.45 34.74 32.71 29.00 27.09
f1 (ppm)

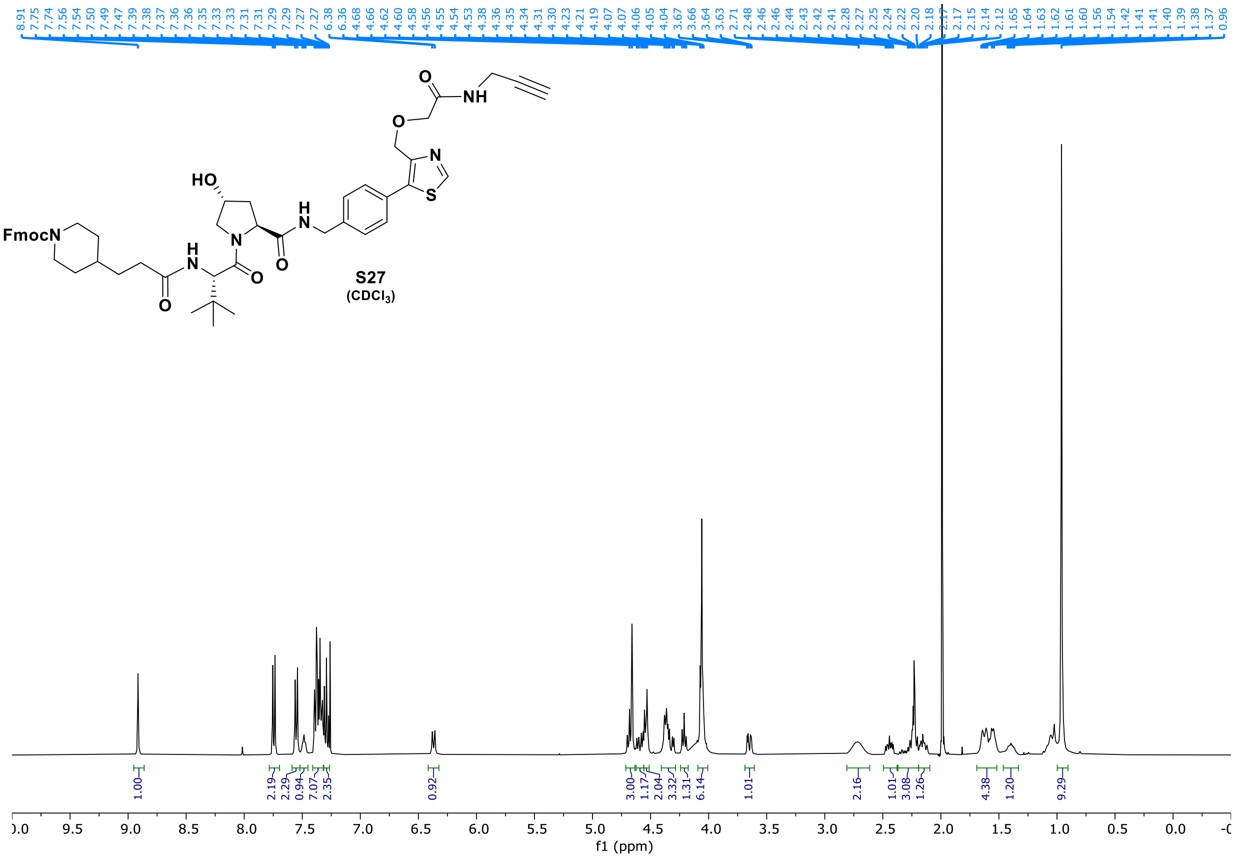
8.91 7.75 7.74 7.56 7.54 7.50 7.49 7.47 7.39 7.38 7.37 7.36 7.36 7.35 7.33 7.33 7.31 7.31 7.29 7.29 7.27 7.27 6.38 6.36 4.68 4.66 4.62 4.60 4.58 4.56 4.55 4.54 4.53 4.38 4.36 4.35 4.34 4.31 4.30 4.23 4.21 4.19 4.07 4.06 4.05 4.04 3.67 3.66 3.64 3.63 2.71 2.48 2.46 2.44 2.43 2.42 2.41 2.28 2.27 2.25 2.24 2.22 2.20 2.18 2.17 2.17 2.15 2.14 2.12 1.65 1.64 1.63 1.62 1.61 1.60 1.56 1.54 1.42 1.41 1.40 1.39 1.38 1.37 0.96
O
NH
O
N
S
HO
H
N
FmocN
H
N
N
O
O
O
S27
(CDCl3)
1.00 2.19 2.29 0.94 7.07 2.35 0.92 3.00 1.17 2.04 3.32 1.31 6.14 1.01 2.16 1.01 3.08 1.26 4.38 1.20 9.29
9.5 9.0 8.5 8.0 7.5 7.0 6.5 6.0 5.5 5.0 4.5 4.0 3.5 3.0 2.5 2.0 1.5 1.0 0.5 0.0
f1 (ppm)

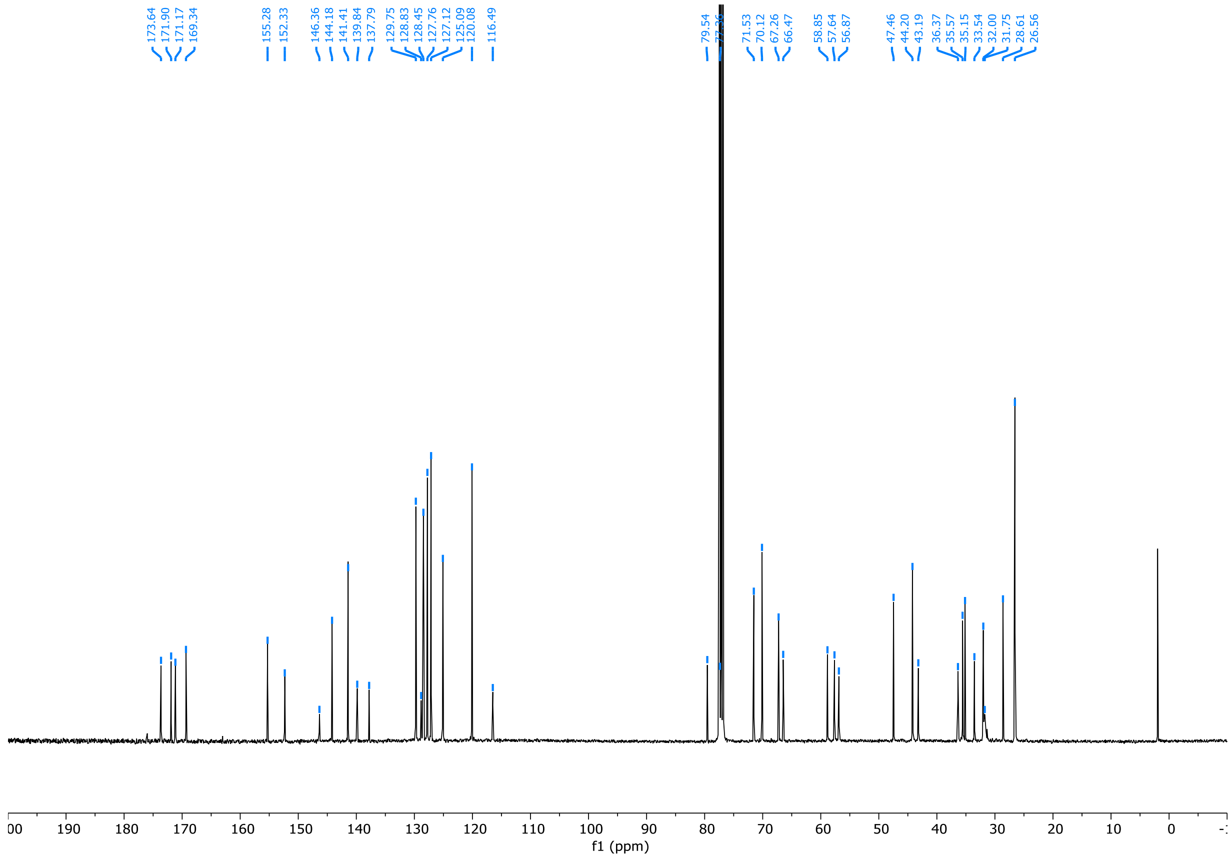
173.64 171.90 171.17 169.34 155.28 152.33 146.36 144.18 141.41 139.84 137.79 129.75 128.83 128.45 127.76 127.12 125.09 120.08 116.49 79.54 71.53 70.12 67.26 66.47 58.85 57.64 56.87 47.46 44.20 43.19 36.37 35.57 35.15 33.54 32.00 31.75 28.61 26.56
190 180 170 160 150 140 130 120 110 100 90 80 70 60 50 40 30 20 10 0
f1 (ppm)

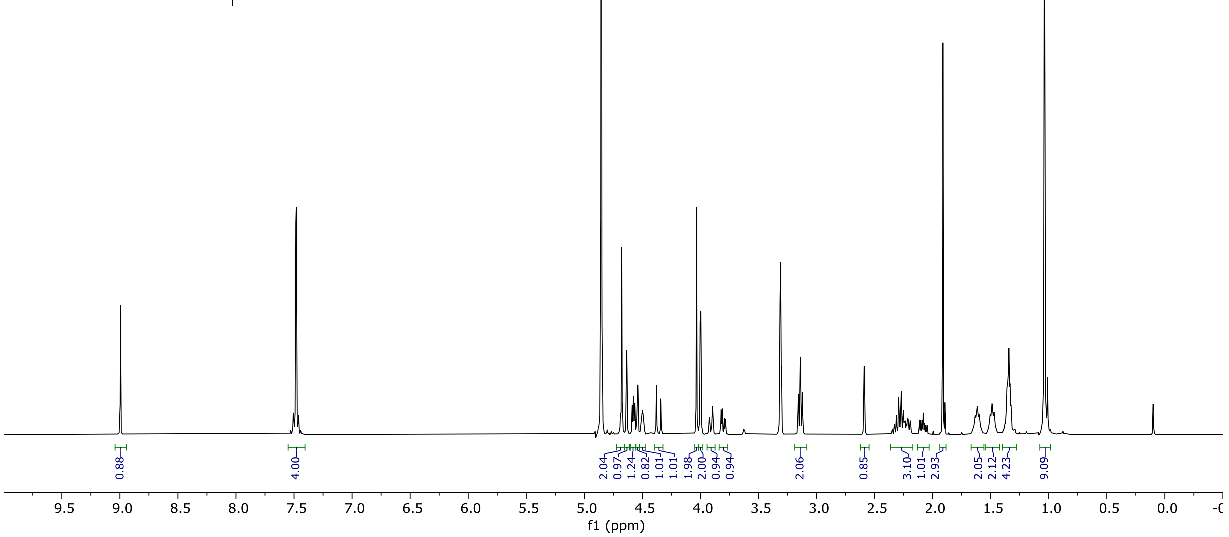
9.00
7.51 7.49 7.48 7.46 4.68 4.63 4.59 4.58 4.57 4.55 4.54 4.50 4.38 4.34 4.03 4.00 4.00 3.92 3.89 3.82 3.81 3.79 3.78 3.16 3.14 3.12 2.60 2.59 2.58 2.35 2.33 2.31 2.29 2.29 2.28 2.27 2.25 2.24 2.24 2.23 2.22 2.21 2.21 2.20 2.19 2.19 2.11 2.10 2.09 2.08 2.07 2.06 2.05 1.91 1.65 1.64 1.63 1.62 1.62 1.60 1.60 1.58 1.52 1.51 1.49 1.47 1.45 1.36 1.35 1.34 1.33 1.32 1.04
0.88 4.00 2.04 0.97 1.24 0.82 1.01 1.01 1.98 2.00 0.94 0.94 2.06 0.85 3.10 1.01 2.93 2.05 2.12 4.23 9.09
f1 (ppm)

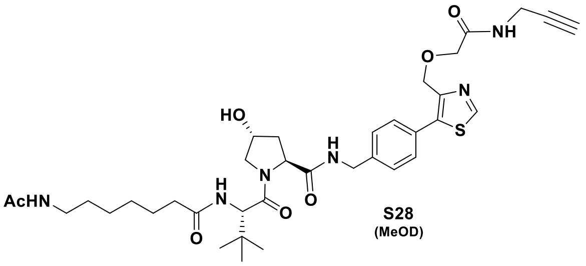

**S28**
(MeOD)

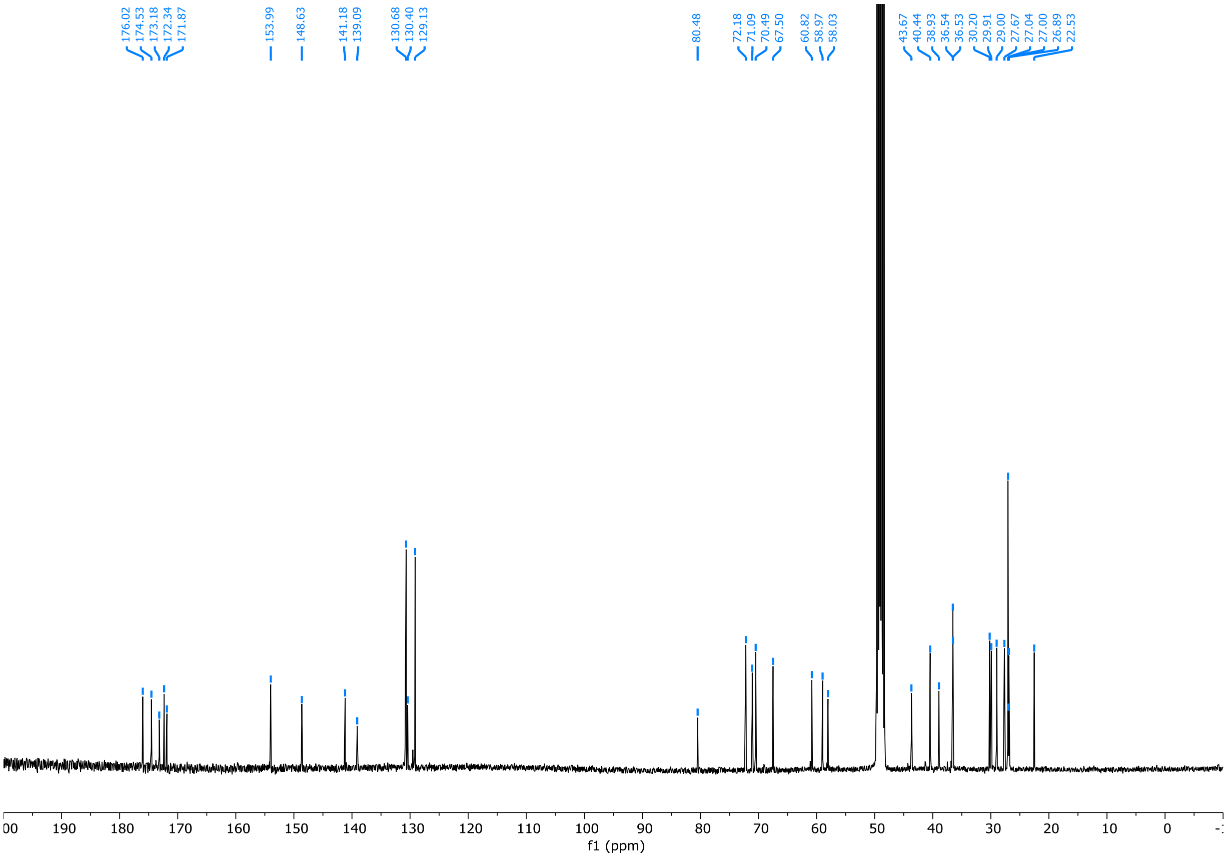
176.02 174.53 173.18 172.34 171.87 153.99 148.63 141.18 139.09 130.68 130.40 129.13 80.48 72.18 71.09 70.49 67.50 60.82 58.97 58.03 43.67 40.44 38.93 36.54 36.53 30.20 29.91 29.00 27.67 27.04 27.00 26.89 22.53
f1 (ppm)

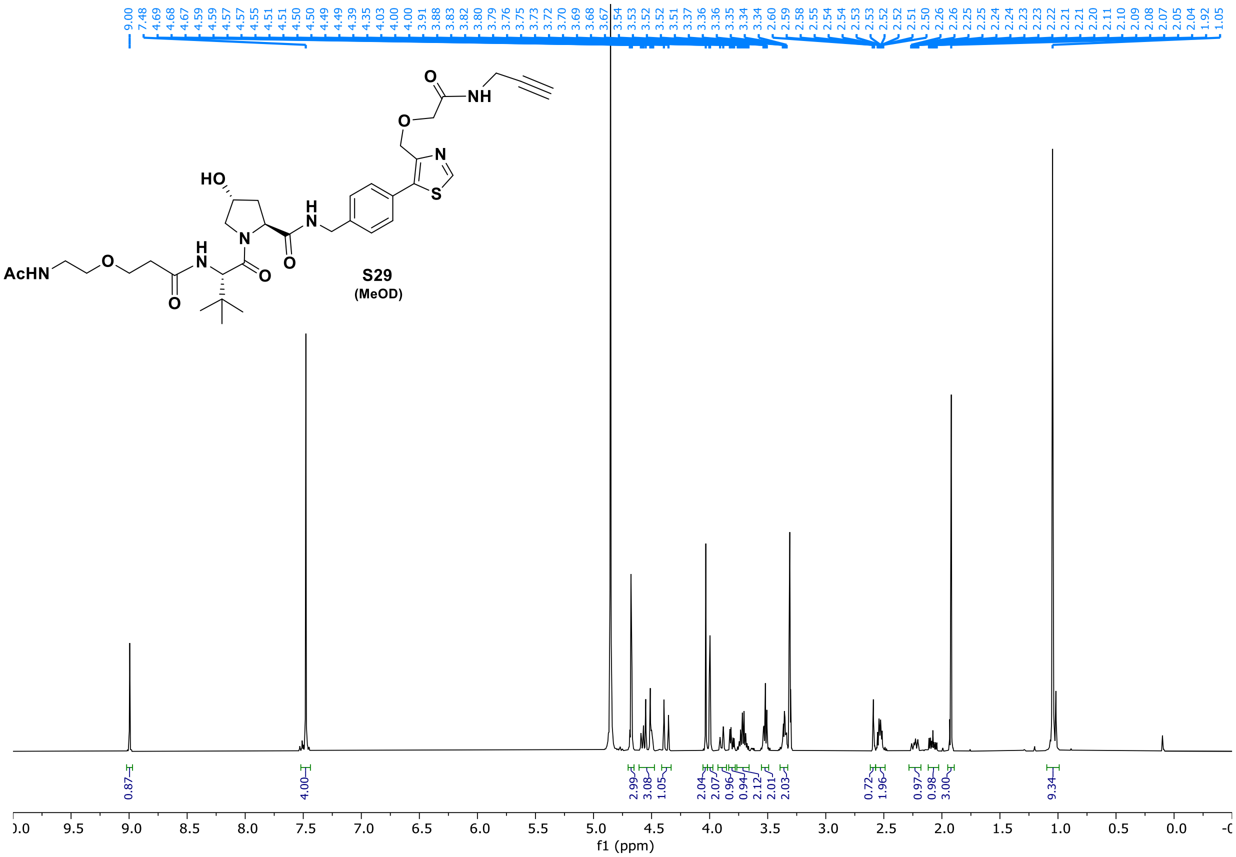

S29
(MeOD)

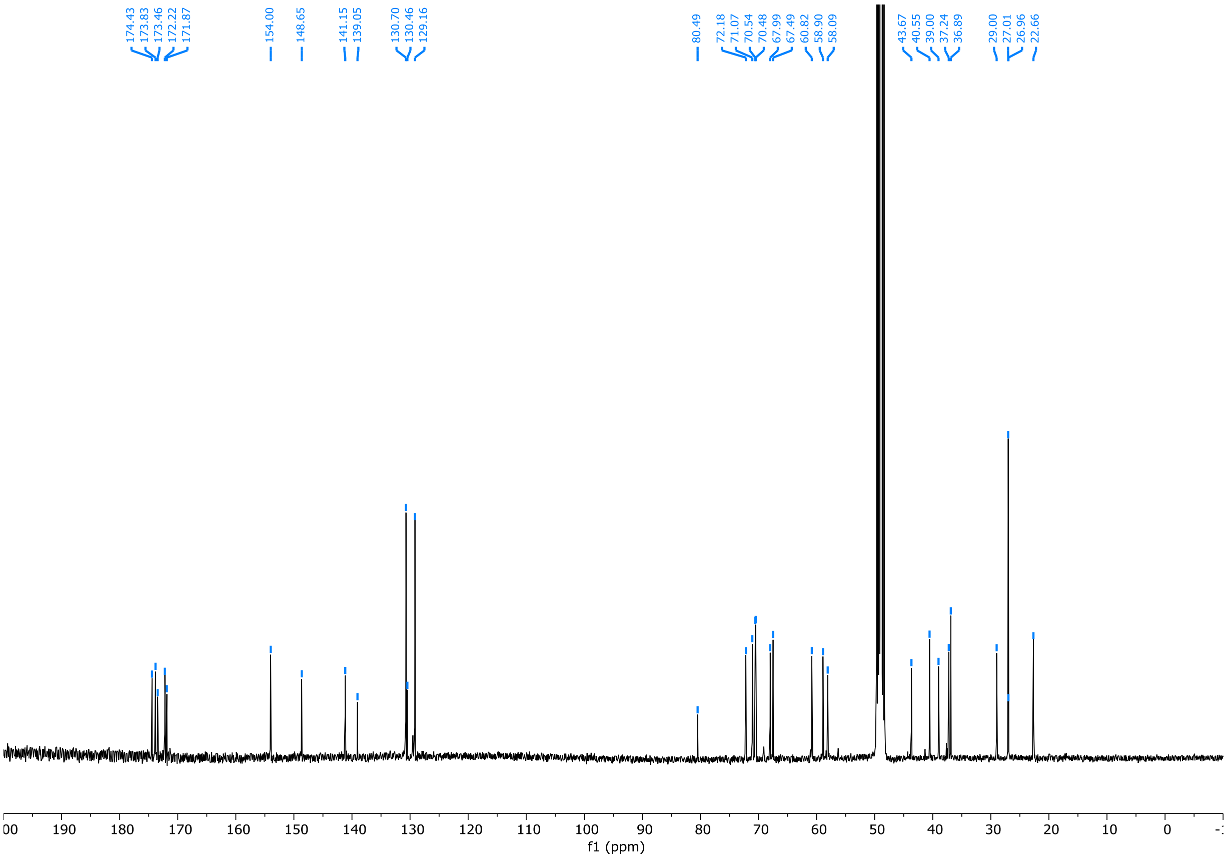

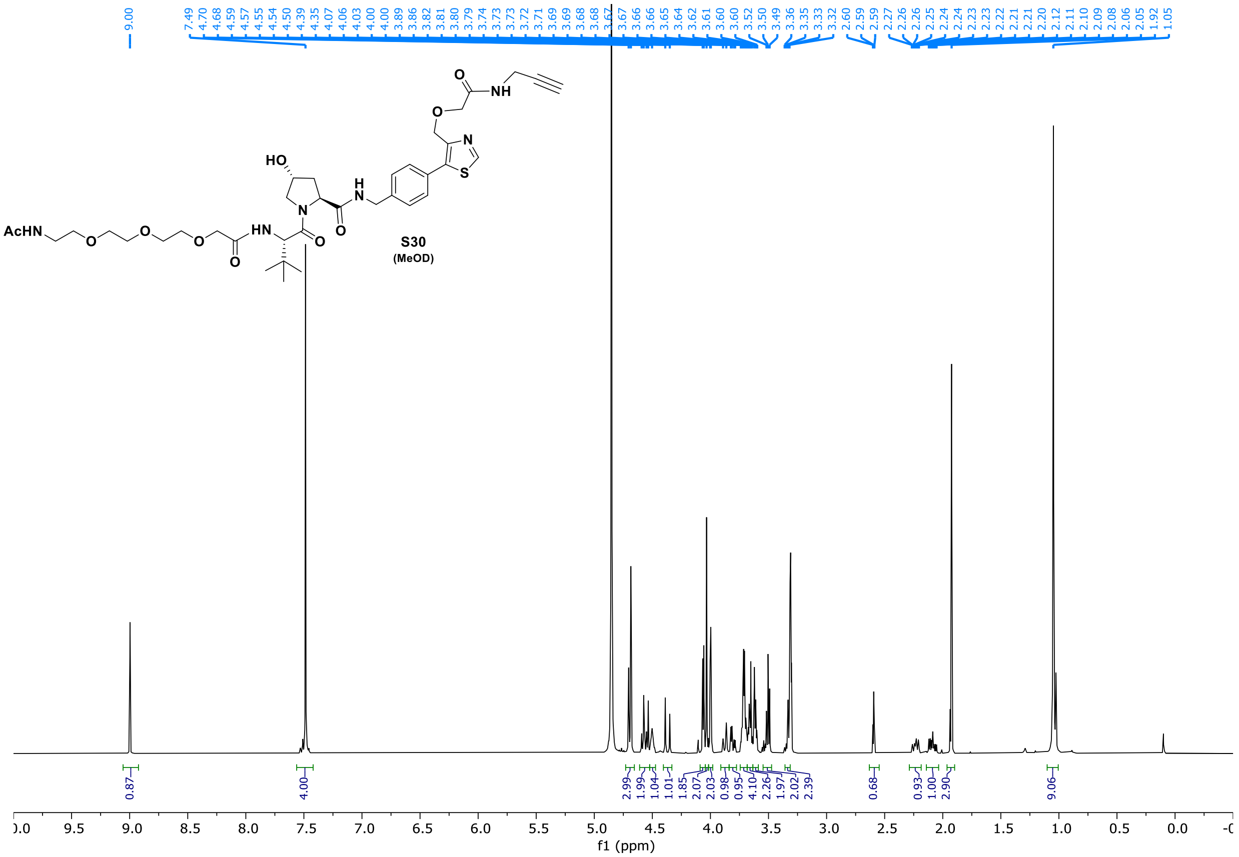
9.00
7.49 4.70 4.68 4.59 4.57 4.55 4.54 4.50 4.39 4.35 4.07 4.06 4.03 4.00 4.00 3.89 3.86 3.82 3.81 3.80 3.79 3.74 3.73 3.73 3.72 3.71 3.69 3.69 3.68 3.67 3.67 3.66 3.66 3.65 3.64 3.62 3.61 3.60 3.60 3.52 3.50 3.49 3.36 3.35 3.33 3.32 2.60 2.59 2.59 2.27 2.26 2.26 2.25 2.24 2.24 2.23 2.23 2.22 2.21 2.21 2.20 2.12 2.11 2.10 2.09 2.08 2.06 2.05 1.92 1.05
AcHN
HO
HN
N
H
NH
O
N
S
S30
(MeOD)
0.87 4.00 2.99 1.99 1.04 1.01 1.85 2.07 2.03 0.98 0.95 4.10 2.26 1.97 2.02 2.39 0.68 0.93 1.00 2.90 9.06
f1 (ppm)

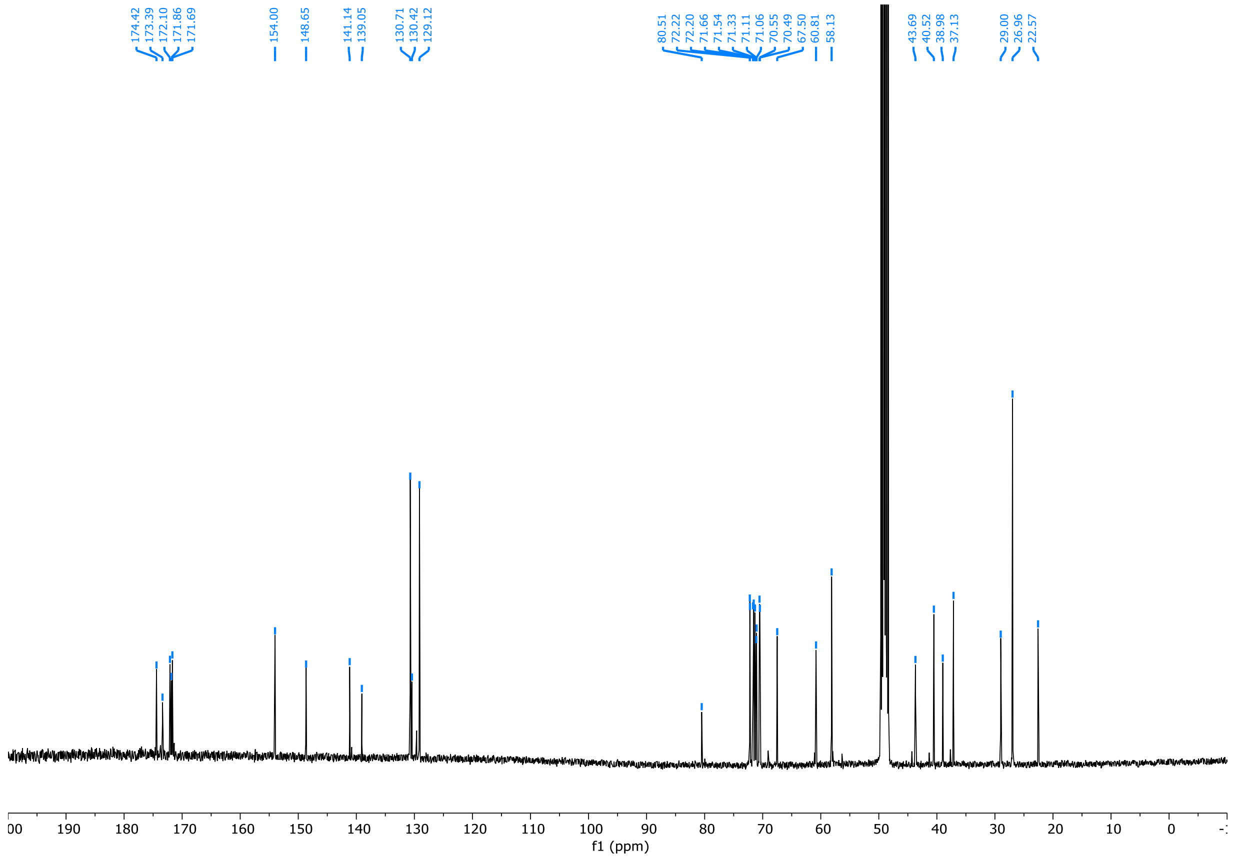
174.42 173.39 172.10 171.86 171.69
154.00
148.65
141.14 139.05
130.71 130.42 129.12
80.51 72.22 72.20 71.66 71.54 71.33 71.11 71.06 70.55 70.49 67.50 60.81 58.13
43.69 40.52 38.98 37.13
29.00 26.96 22.57
f1 (ppm)

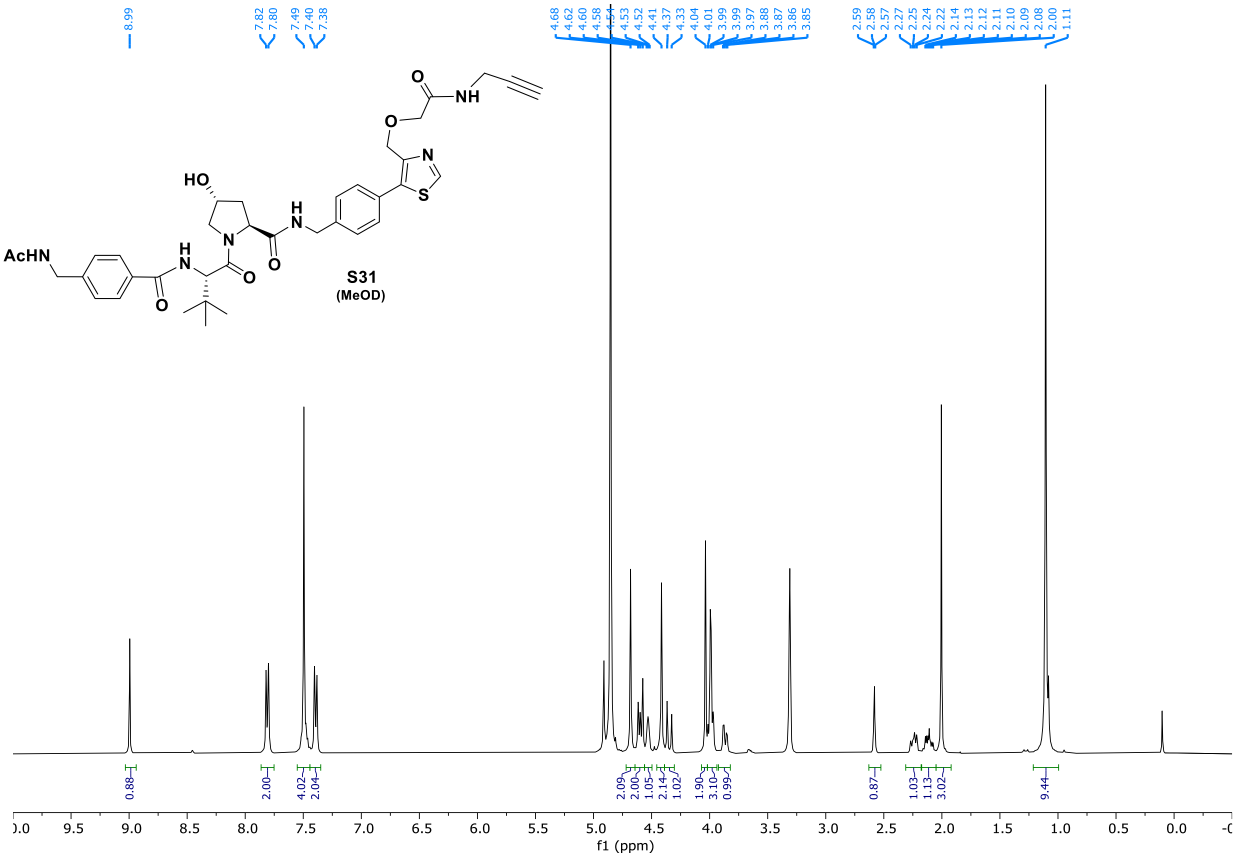
S31
(MeOD)
8.99
7.82
7.80
7.49
7.40
7.38
4.68
4.62
4.60
4.58
4.54
4.53
4.52
4.41
4.37
4.33
4.04
4.01
3.99
3.99
3.97
3.88
3.87
3.86
3.85
2.59
2.58
2.57
2.27
2.25
2.24
2.22
2.14
2.13
2.12
2.11
2.10
2.09
2.08
2.00
1.11
0.88
2.00
4.02
2.04
2.09
2.00
1.05
2.14
1.02
1.90
3.10
0.99
0.87
1.03
1.13
3.02
9.44
f1 (ppm)

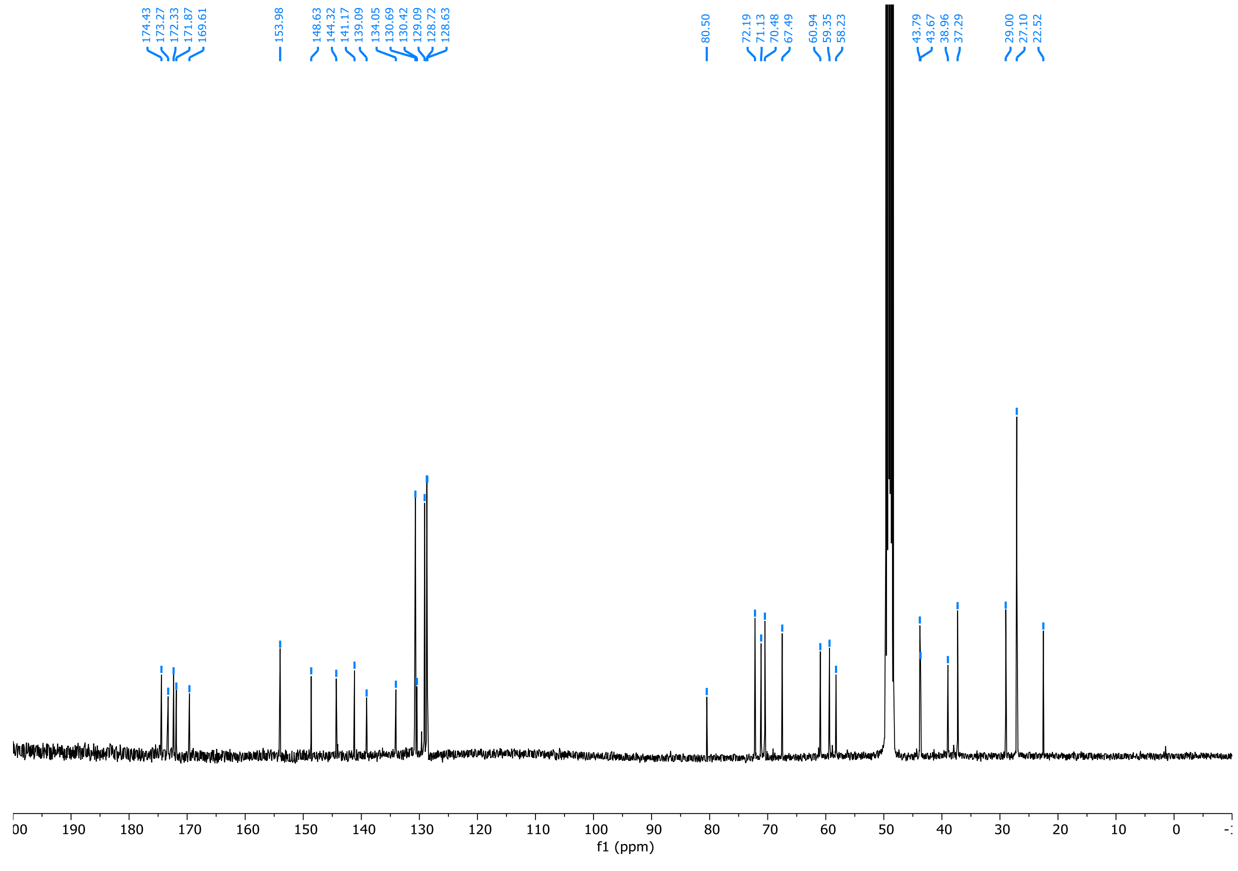
174.43
173.27
172.33
171.87
169.61
153.98
148.63
144.32
141.17
139.09
134.05
130.69
130.42
129.09
128.72
128.63
80.50
72.19
71.13
70.48
67.49
60.94
59.35
58.23
43.79
43.67
38.96
37.29
29.00
27.10
22.52
f1 (ppm)

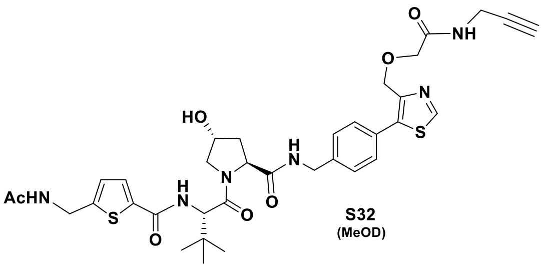
AcHN
S
O
H
N
HO
N
O
H
N
O
N
S
O
O
NH

S32
(MeOD)

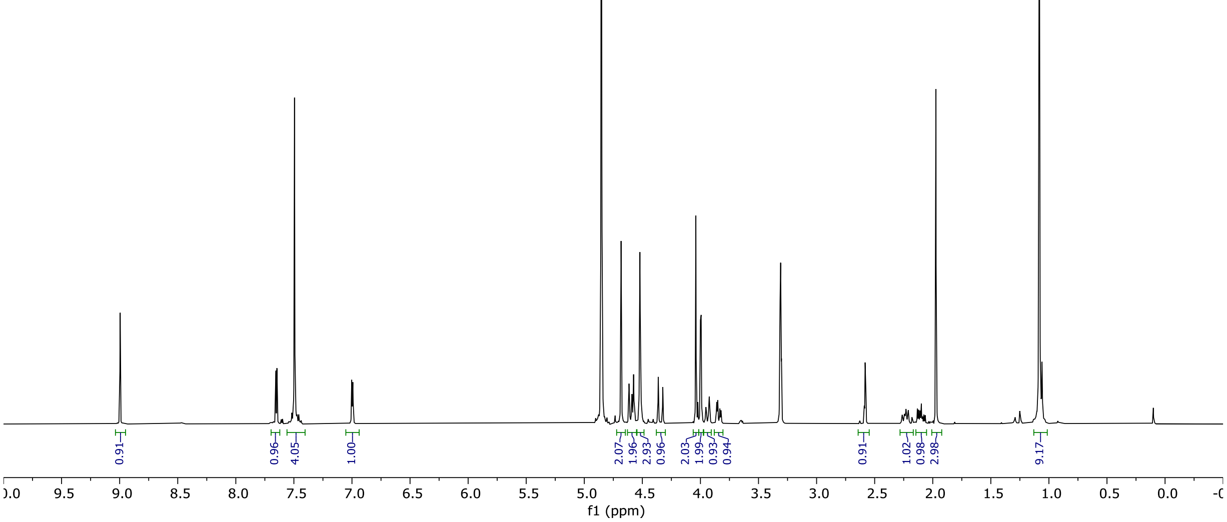
9.00
7.66
7.65
7.49
7.00
6.99
4.68
4.61
4.59
4.58
4.57
4.52
4.36
4.32
4.04
4.00
3.99
3.95
3.92
3.86
3.85
3.83
3.82
2.59
2.58
2.58
2.27
2.26
2.26
2.25
2.24
2.24
2.23
2.23
2.22
2.21
2.21
2.13
2.12
2.11
2.10
2.09
2.08
2.07
1.97
1.08
0.91
0.96
4.05
1.00
2.07
1.96
2.93
0.96
2.03
1.99
0.93
0.94
0.91
1.02
0.98
2.98
9.17
f1 (ppm)

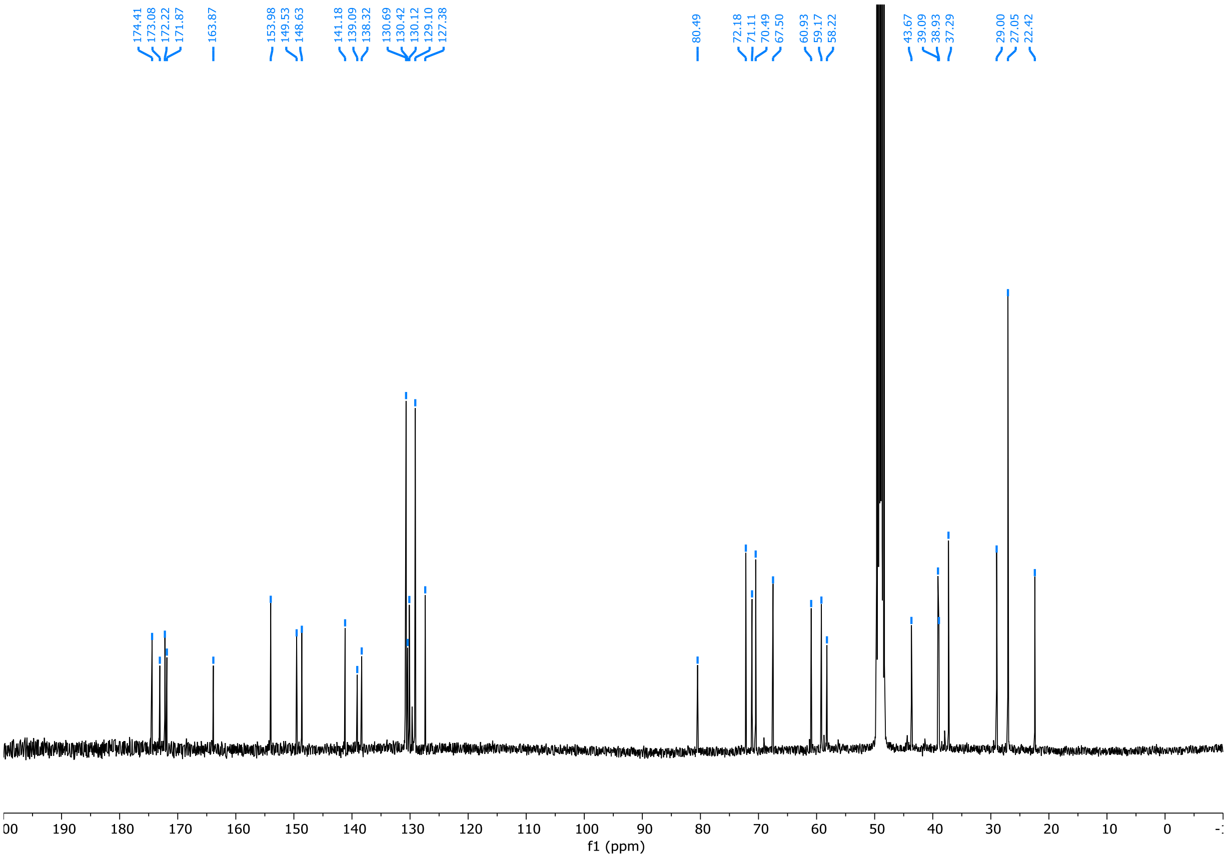
174.41
173.08
172.22
171.87
163.87
153.98
149.53
148.63
141.18
139.09
138.32
130.69
130.42
130.12
129.10
127.38
80.49
72.18
71.11
70.49
67.50
60.93
59.17
58.22
43.67
39.09
38.93
37.29
29.00
27.05
22.42
f1 (ppm)

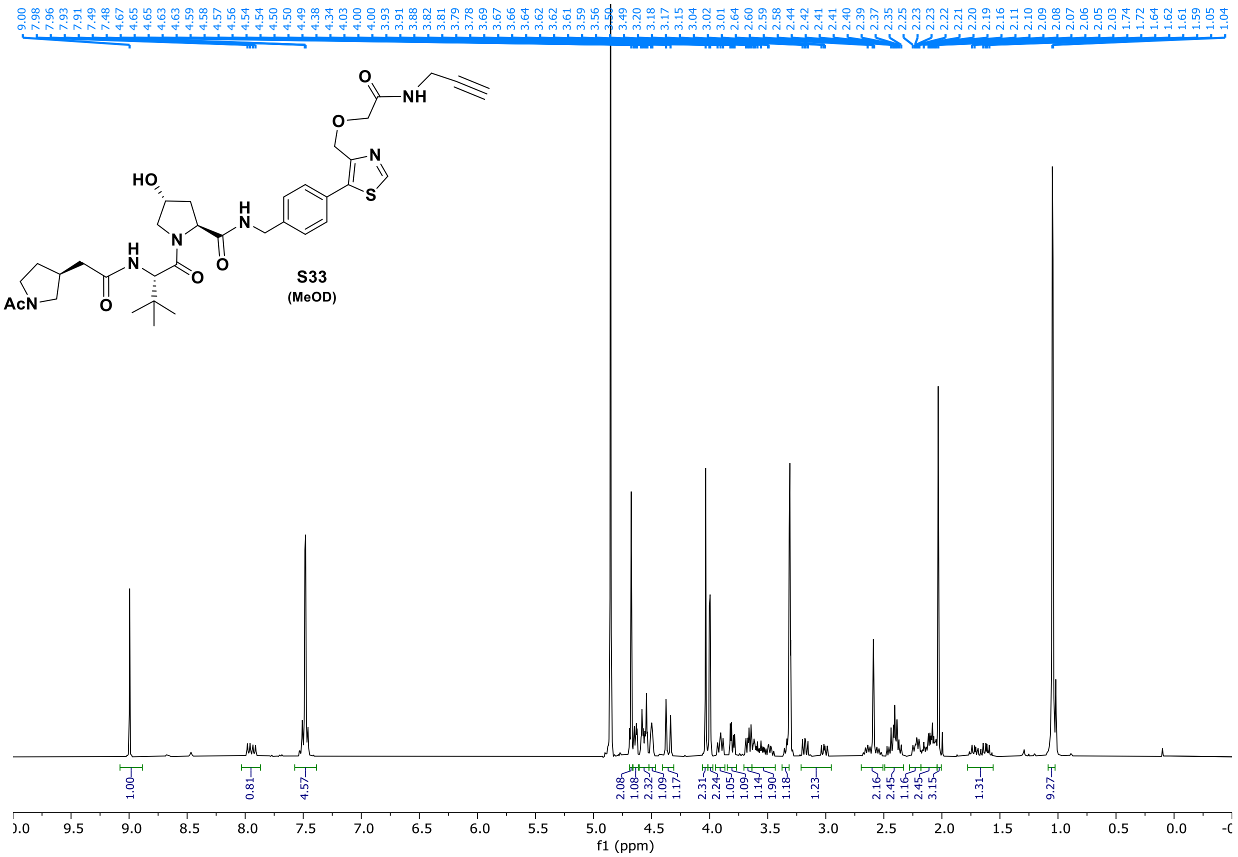

S33
(MeOD)


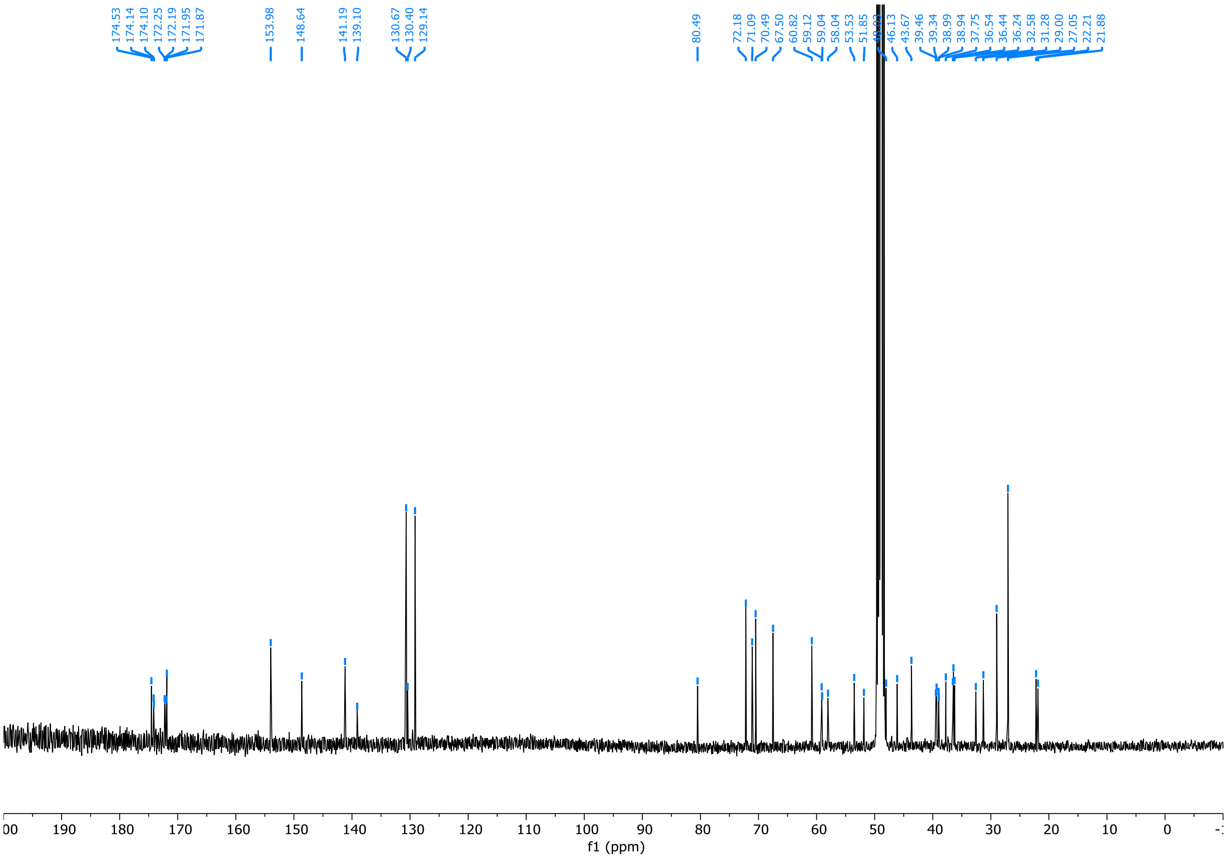

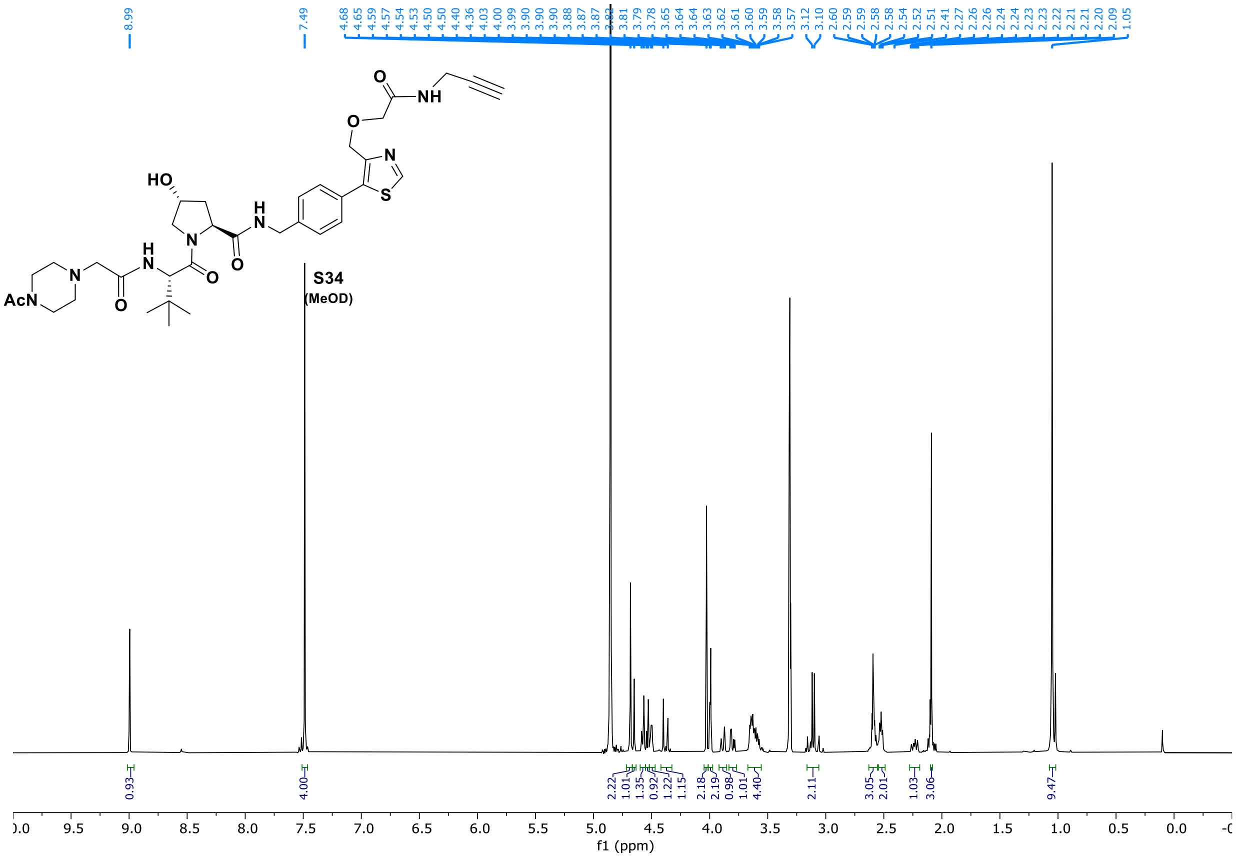
S34
(MeOD)
f1 (ppm)

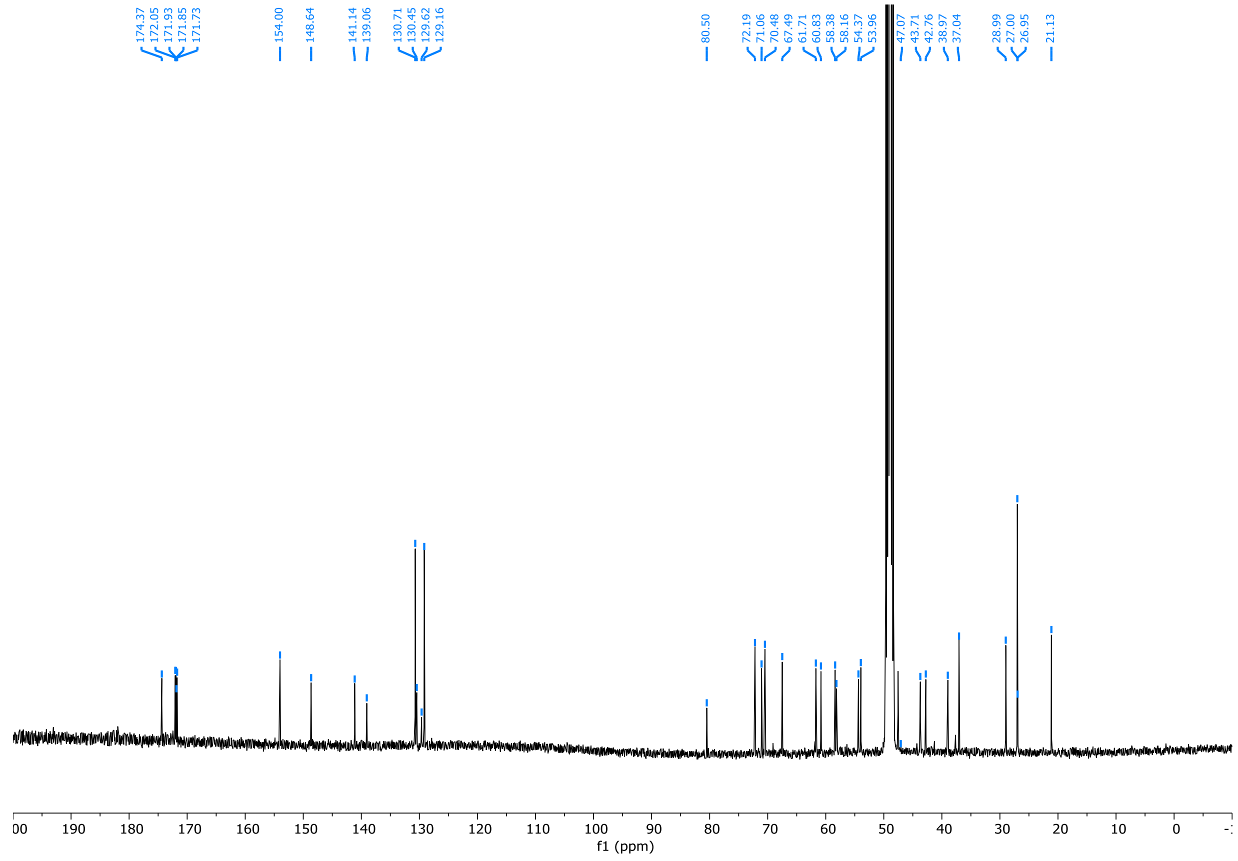
f1 (ppm)

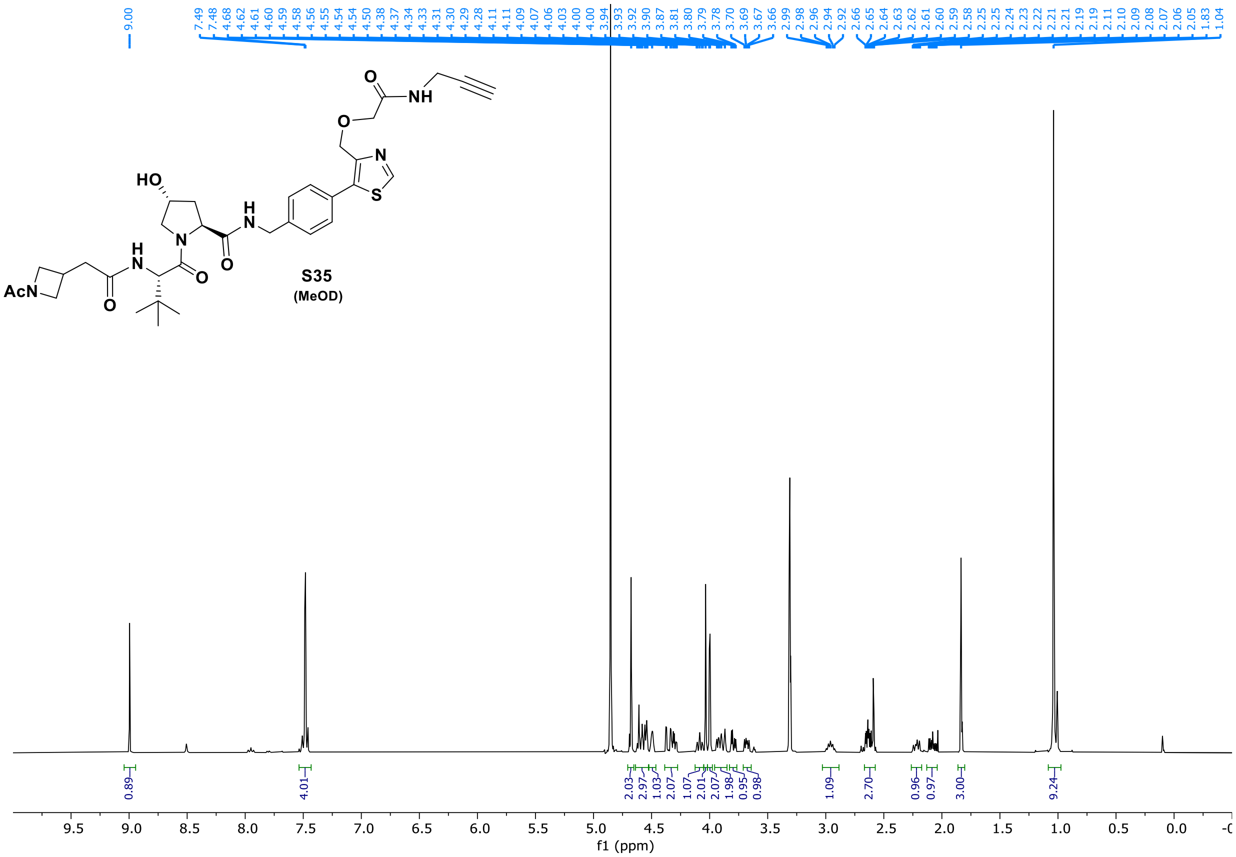
9.00
7.49 7.48 4.68 4.62 4.61 4.60 4.59 4.58 4.56 4.55 4.54 4.54 4.50 4.38 4.37 4.34 4.33 4.31 4.30 4.29 4.28 4.11 4.11 4.09 4.07 4.06 4.03 4.00 4.00 3.94 3.93 3.92 3.90 3.87 3.81 3.80 3.79 3.78 3.70 3.69 3.67 3.66 2.99 2.98 2.96 2.94 2.92 2.66 2.65 2.64 2.63 2.62 2.61 2.60 2.59 2.58 2.25 2.25 2.24 2.23 2.22 2.21 2.21 2.19 2.19 2.11 2.10 2.09 2.08 2.07 2.06 2.05 1.83 1.04
O
NH
O
N
S
HO
H
N
N
H
N
O
O
AcN
O
S35
(MeOD)
0.89
4.01
2.03 2.97 1.03 2.07 1.07 2.01 2.07 1.98 0.95 0.98
1.09
2.70
0.96 0.97
3.00
9.24
9.5 9.0 8.5 8.0 7.5 7.0 6.5 6.0 5.5 5.0 4.5 4.0 3.5 3.0 2.5 2.0 1.5 1.0 0.5 0.0
f1 (ppm)

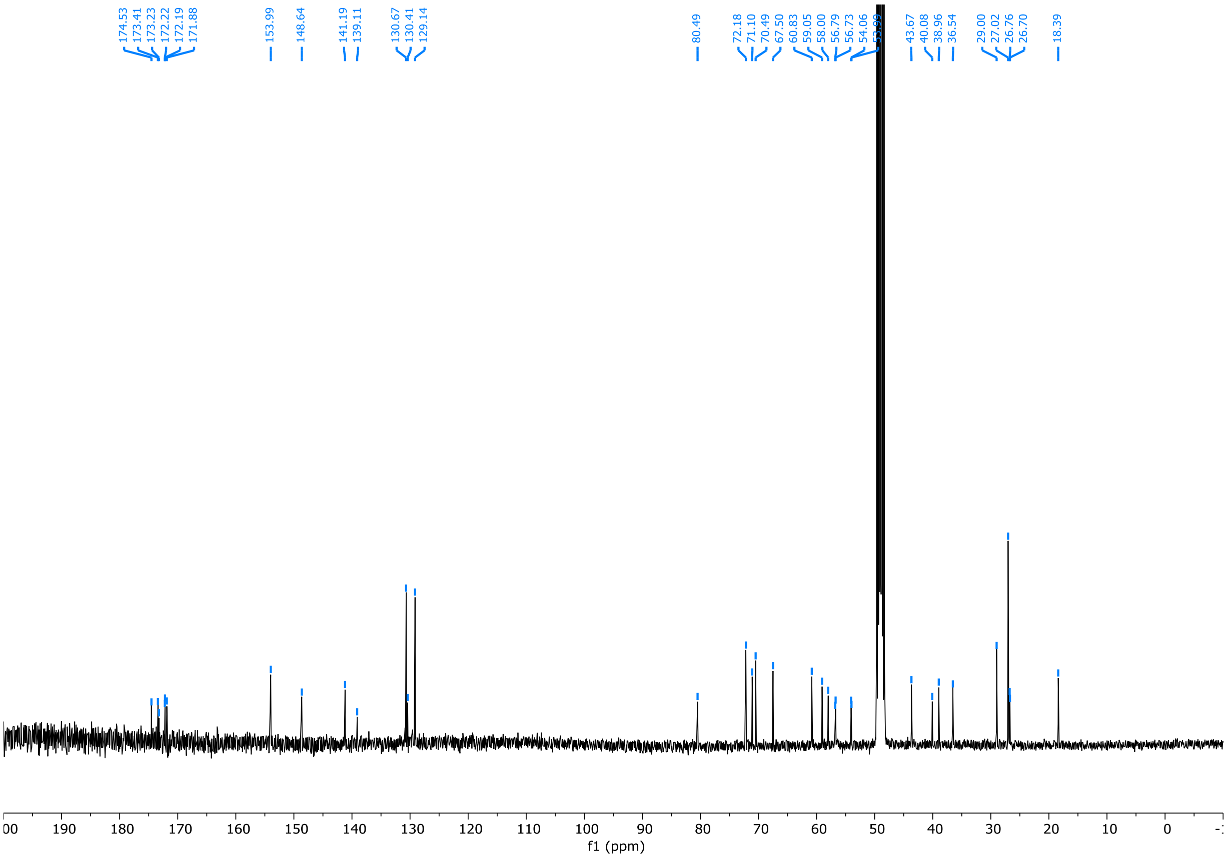
174.53 173.41 173.23 172.22 172.19 171.88
153.99
148.64
141.19 139.11
130.67 130.41 129.14
80.49
72.18 71.10 70.49 67.50 60.83 59.05 58.00 56.79 56.73 54.06
43.67 40.08 38.96 36.54
29.00 27.02 26.76 26.70
18.39
190 180 170 160 150 140 130 120 110 100 90 80 70 60 50 40 30 20 10 0
f1 (ppm)

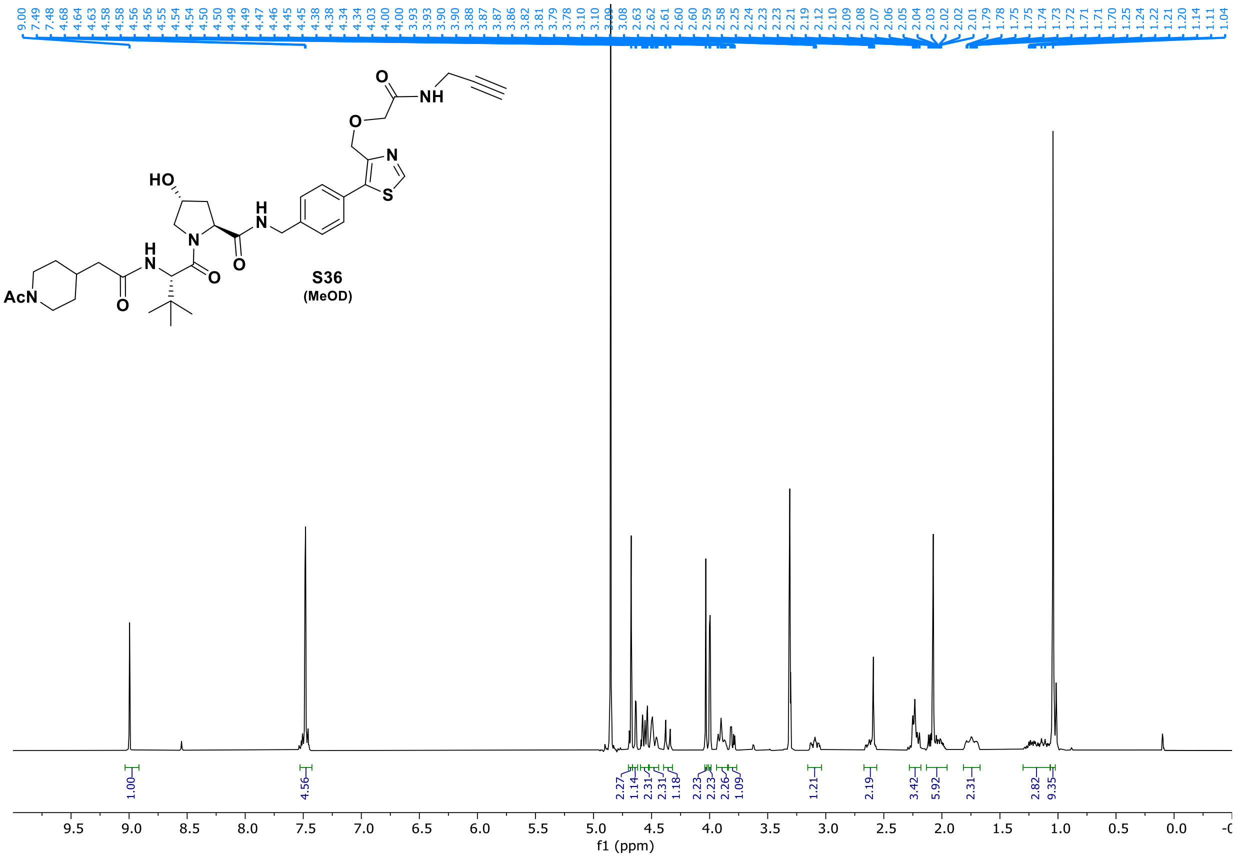
HO
AcN
S36
(MeOD)
9.00 7.49 7.48 4.68 4.64 4.63 4.58 4.58 4.56 4.56 4.55 4.54 4.54 4.50 4.50 4.49 4.49 4.47 4.46 4.45 4.38 4.38 4.34 4.34 4.03 4.00 3.93 3.93 3.90 3.88 3.87 3.87 3.86 3.82 3.81 3.79 3.78 3.10 3.10 3.09 3.08 2.63 2.62 2.61 2.60 2.60 2.59 2.58 2.25 2.24 2.23 2.23 2.21 2.19 2.12 2.10 2.09 2.08 2.07 2.06 2.05 2.04 2.03 2.02 2.02 2.01 1.79 1.78 1.75 1.75 1.74 1.73 1.72 1.71 1.71 1.70 1.25 1.24 1.22 1.21 1.20 1.14 1.11 1.04
1.00 4.56 2.27 1.14 2.31 2.31 1.18 2.23 2.23 2.26 1.09 1.21 2.19 3.42 5.92 2.31 2.82 9.35
9.5 9.0 8.5 8.0 7.5 7.0 6.5 6.0 5.5 5.0 4.5 4.0 3.5 3.0 2.5 2.0 1.5 1.0 0.5 0.0
f1 (ppm)

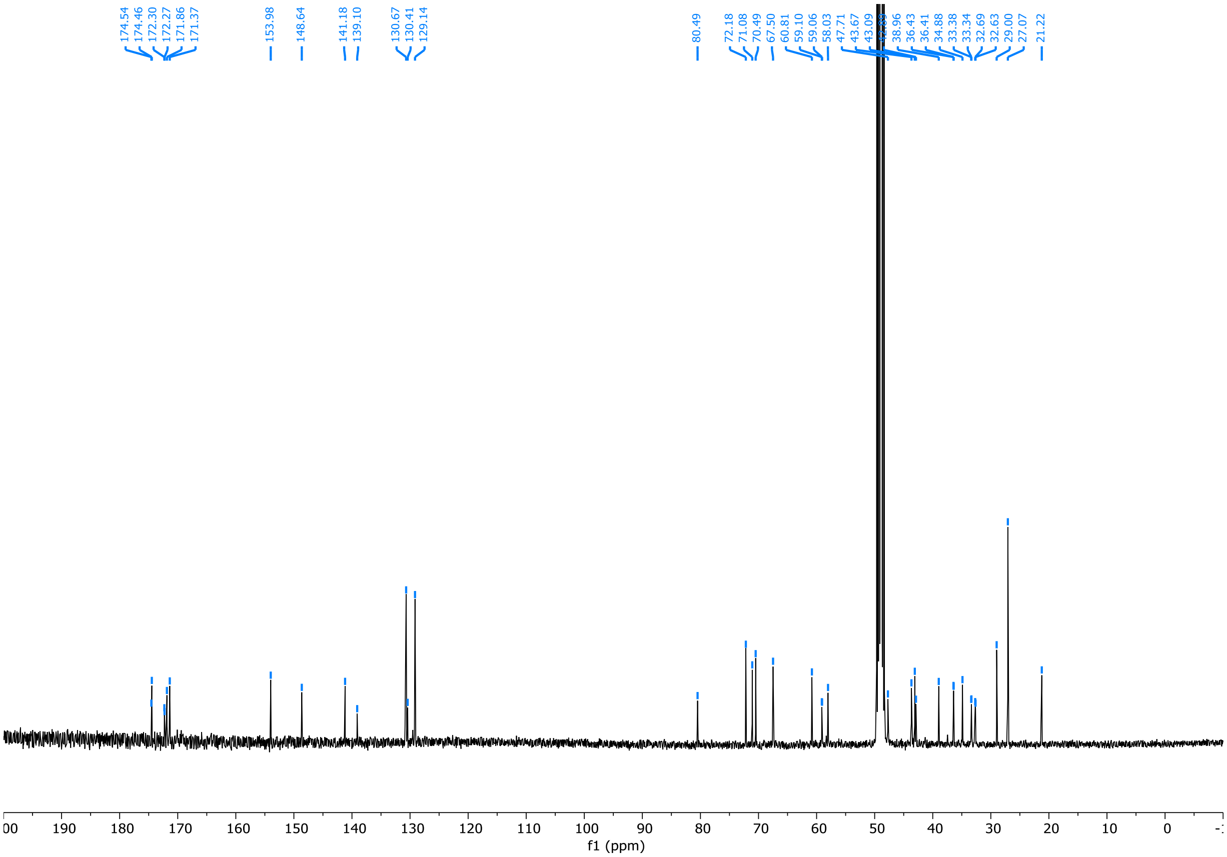
174.54 174.46 172.30 172.27 171.86 171.37 153.98 148.64 141.18 139.10 130.67 130.41 129.14 80.49 72.18 71.08 70.49 67.50 60.81 59.10 59.06 58.03 47.71 43.67 43.09 38.96 36.43 36.41 34.88 33.38 33.34 32.69 32.63 29.00 27.07 21.22
190 180 170 160 150 140 130 120 110 100 90 80 70 60 50 40 30 20 10 0
f1 (ppm)

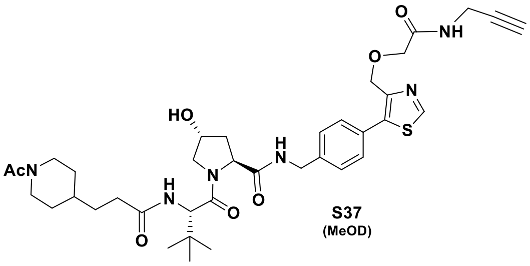
AcN
HO
N
H
N
O
O
H
N
O
N
S
O
O
NH

S37
(MeOD)

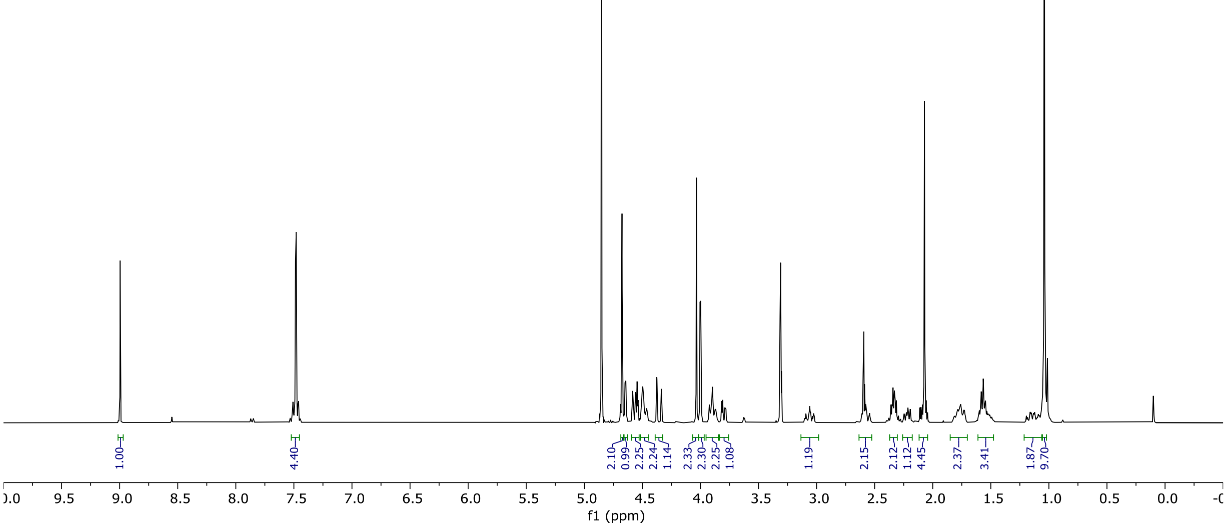
9.00 7.49 7.48 4.68 4.65 4.64 4.58 4.56 4.55 4.54 4.50 4.50 4.49 4.47 4.46 4.38 4.34 4.03 4.00 4.00 3.93 3.92 3.92 3.90 3.90 3.89 3.88 3.87 3.87 3.82 3.81 3.79 3.78 3.09 3.07 3.06 3.05 3.03 2.61 2.60 2.59 2.59 2.58 2.57 2.54 2.36 2.35 2.34 2.33 2.32 2.31 2.25 2.23 2.22 2.22 2.21 2.21 2.20 2.19 2.19 2.11 2.10 2.09 2.07 2.06 2.04 1.78 1.76 1.76 1.73 1.73 1.60 1.58 1.56 1.55 1.55 1.53 1.53 1.52 1.51 1.16 1.15 1.13 1.12 1.09 1.04
1.00
4.40
2.10
0.99
2.25
2.24
1.14
2.33
2.30
2.25
1.08
1.19
2.15
2.12
1.12
4.45
2.37
3.41
1.87
9.70
f1 (ppm)

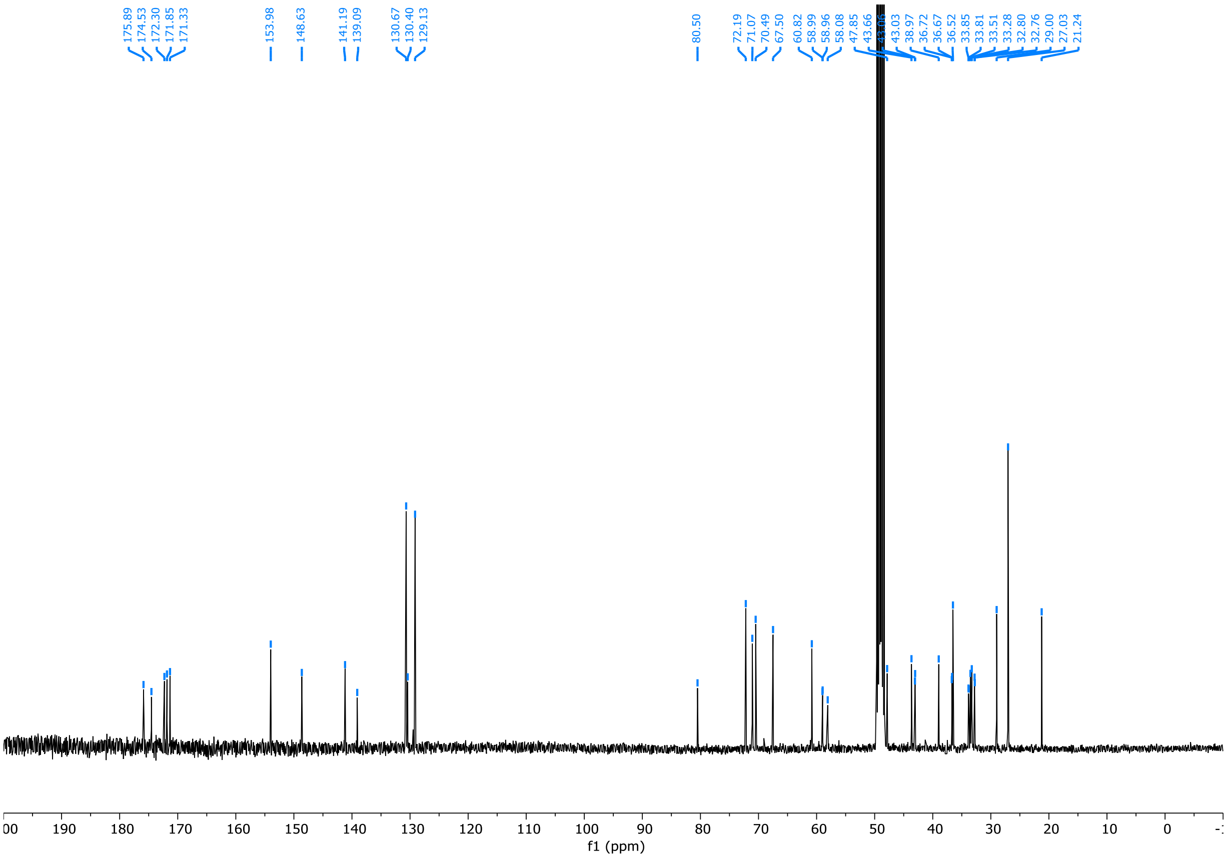
175.89 174.53 172.30 171.85 171.33
153.98
148.63
141.19 139.09
130.67 130.40 129.13
80.50
72.19 71.07 70.49 67.50 60.82 58.99 58.96 58.08 47.85 43.66 43.03 38.97 36.72 36.67 36.52 33.85 33.81 33.51 33.28 32.80 32.76 29.00 27.03 21.24
f1 (ppm)

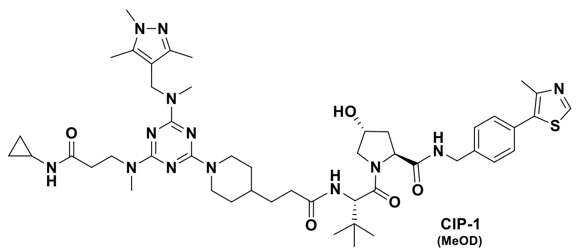
N-N
N
HO
N
S
O
H
N
N
N
H
N
N
O
N
N
H
N
N
O
O
O

CIP-1
(MeOD)

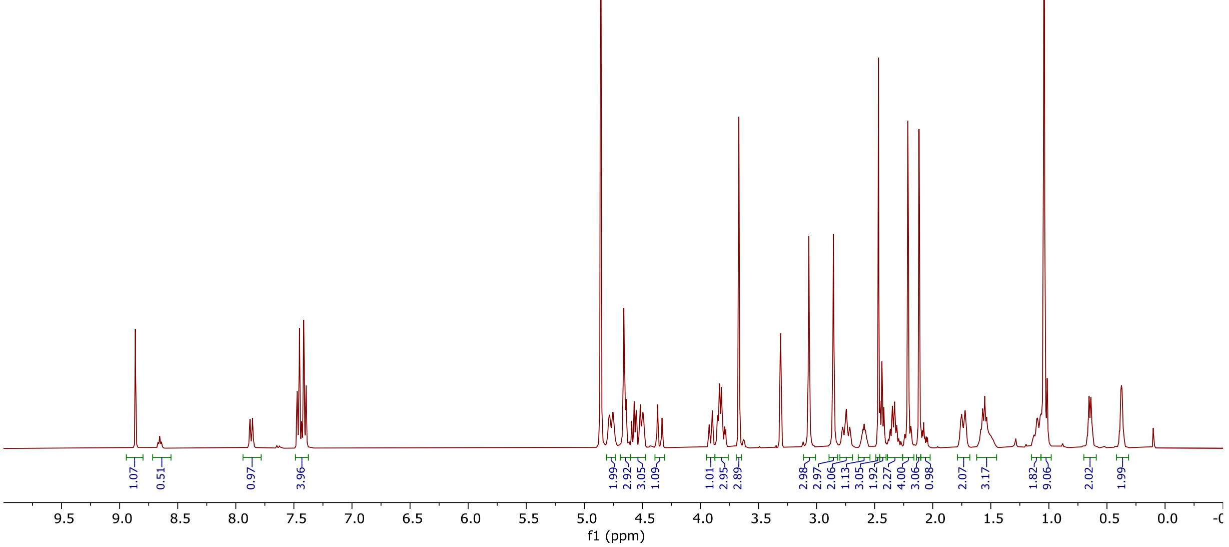
8.86
7.88
7.86
7.47
7.45
7.43
7.43
7.41
7.39
4.78
4.75
4.66
4.64
4.61
4.59
4.57
4.55
4.53
4.52
4.50
4.49
4.49
4.37
4.33
3.92
3.90
3.85
3.84
3.82
3.81
3.79
3.78
3.67
3.07
2.86
2.78
2.74
2.71
2.62
2.61
2.60
2.59
2.58
2.57
2.56
2.47
2.45
2.44
2.42
2.38
2.37
2.35
2.33
2.31
2.29
2.27
2.21
2.12
2.10
2.09
2.08
2.07
2.06
2.04
1.75
1.72
1.59
1.57
1.55
1.54
1.52
1.50
1.13
1.10
1.07
1.04
0.65
0.64
0.38
0.38
0.37
0.37
1.07
0.51
0.97
3.96
1.99
2.92
3.05
1.09
1.01
2.95
2.89
2.98
2.97
2.06
1.13
3.05
1.92
2.27
4.00
3.06
0.98
2.07
3.17
1.82
9.06
2.02
1.99
9.5 9.0 8.5 8.0 7.5 7.0 6.5 6.0 5.5 5.0 4.5 4.0 3.5 3.0 2.5 2.0 1.5 1.0 0.5 0.0
f1 (ppm)

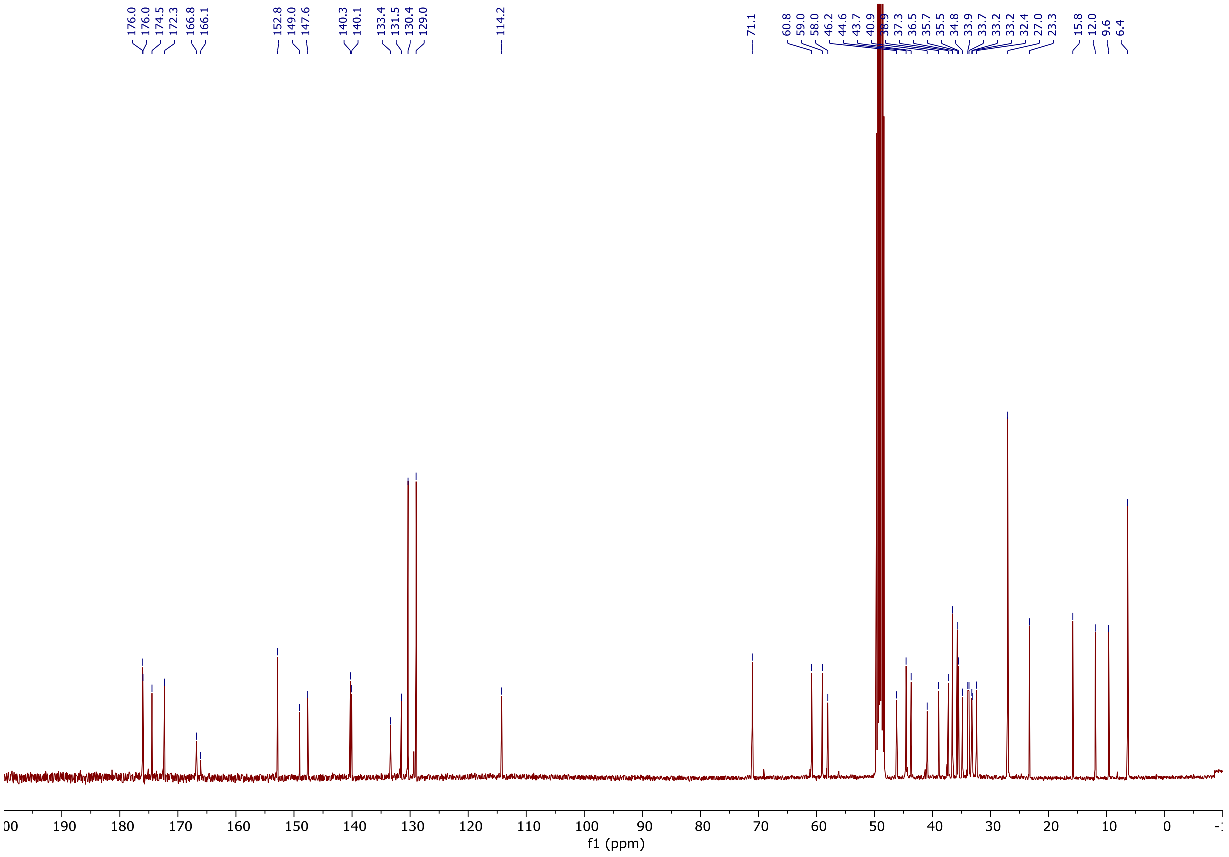
176.0
176.0
174.5
172.3
166.8
166.1
152.8
149.0
147.6
140.3
140.1
133.4
131.5
130.4
129.0
114.2
71.1
60.8
59.0
58.0
46.2
44.6
43.7
40.9
37.3
36.5
35.7
35.5
34.8
33.9
33.7
33.2
33.2
32.4
27.0
23.3
15.8
12.0
9.6
6.4
190 180 170 160 150 140 130 120 110 100 90 80 70 60 50 40 30 20 10 0
f1 (ppm)

**COSY:**

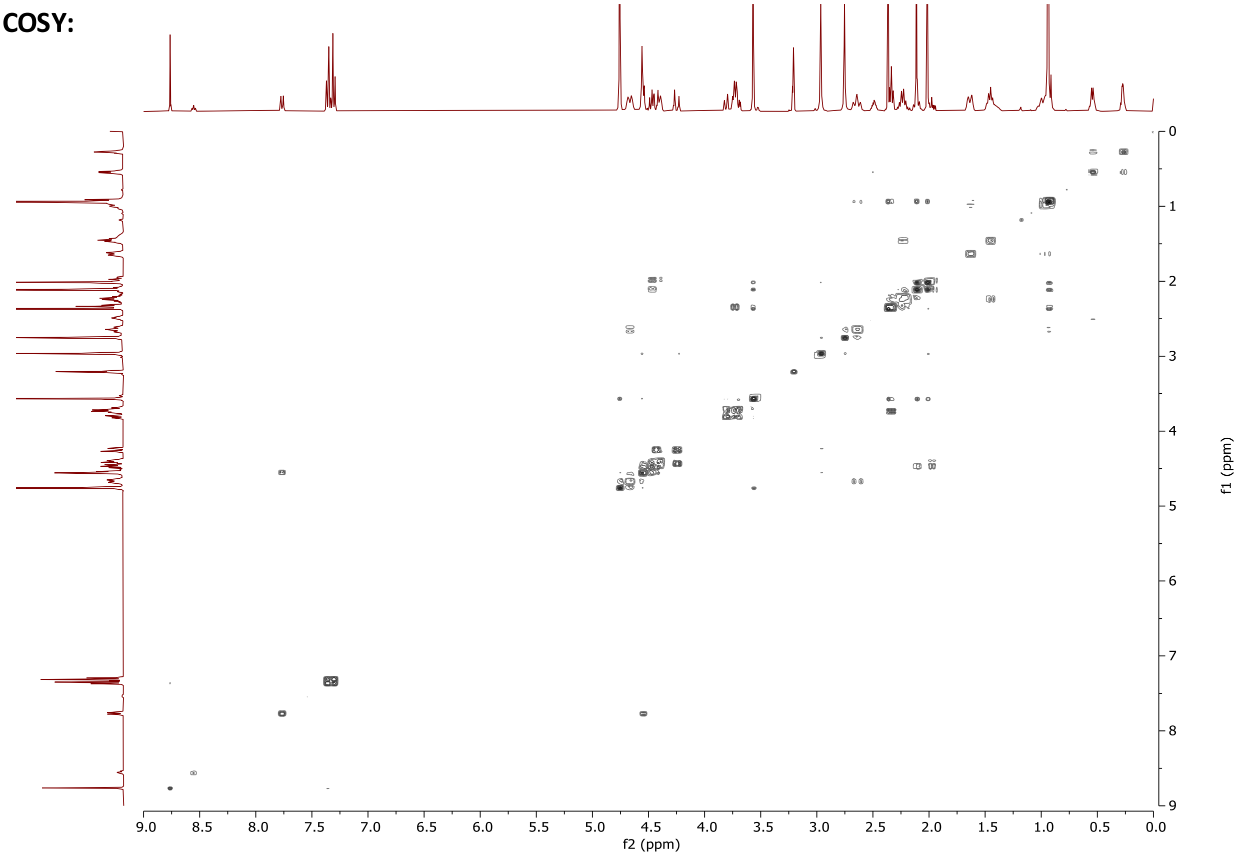


**HSQC:**

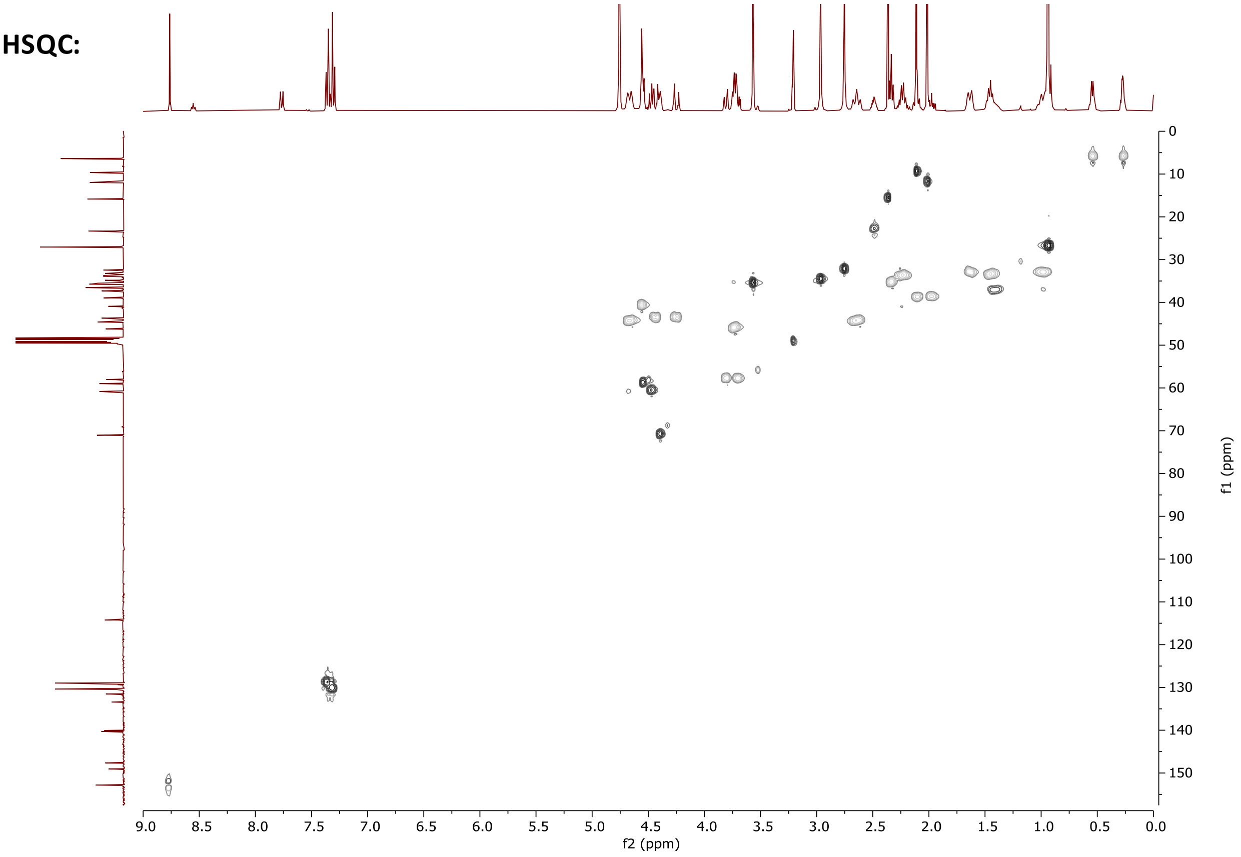

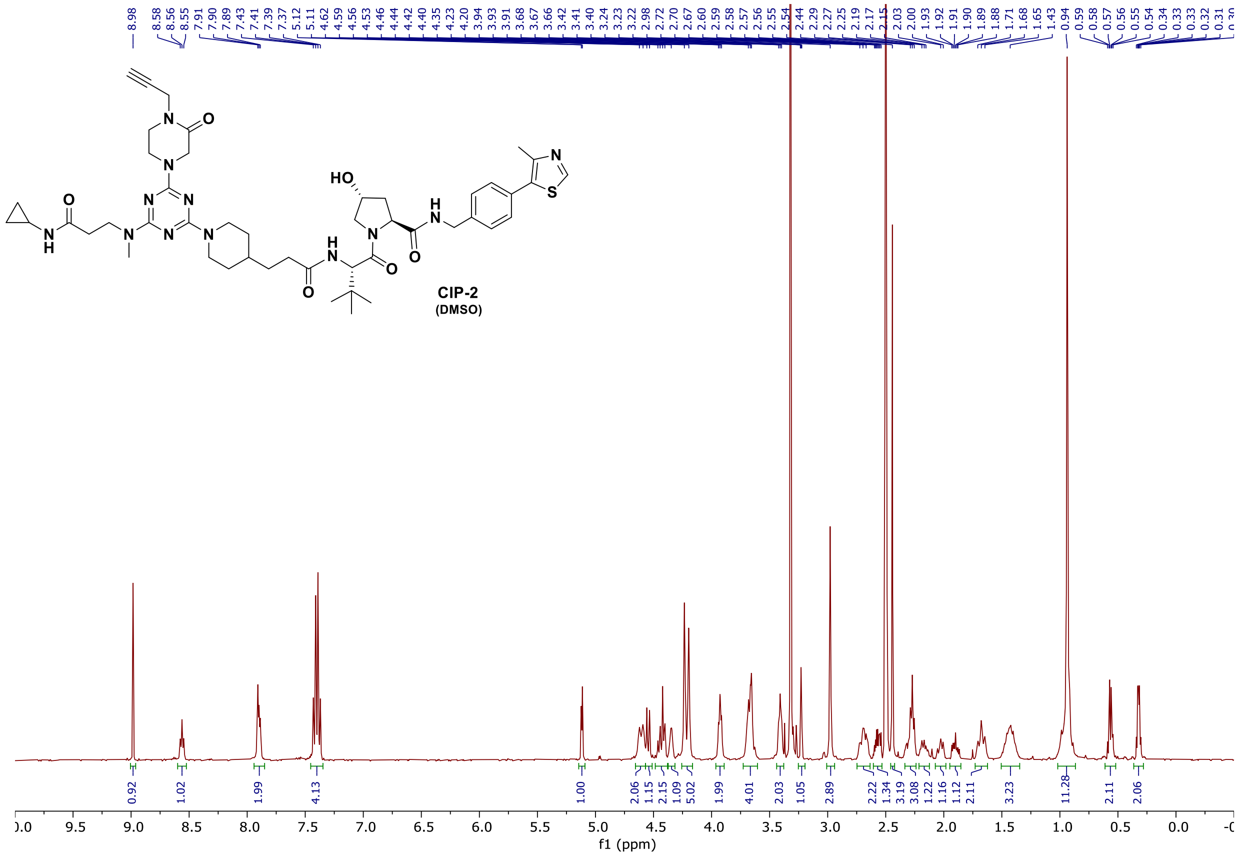

**CIP-2**
**(DMSO)**



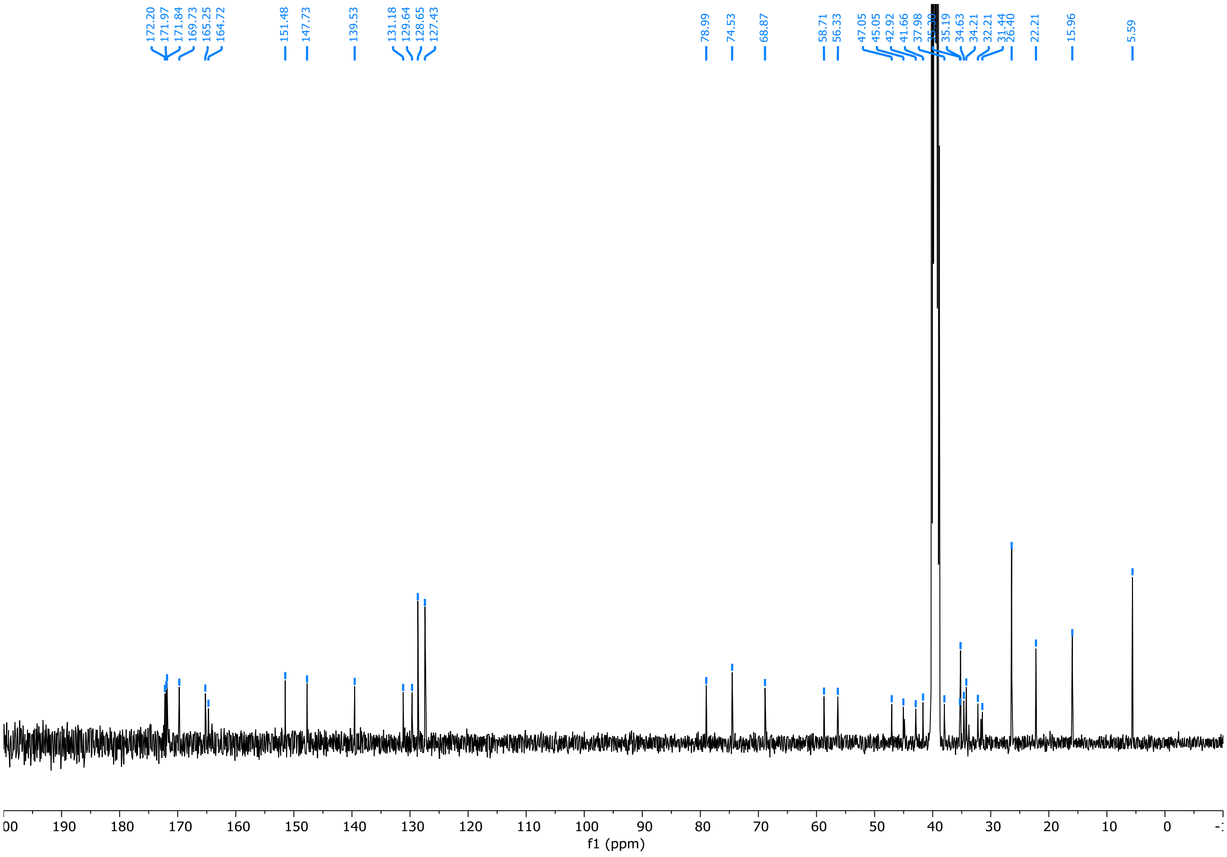

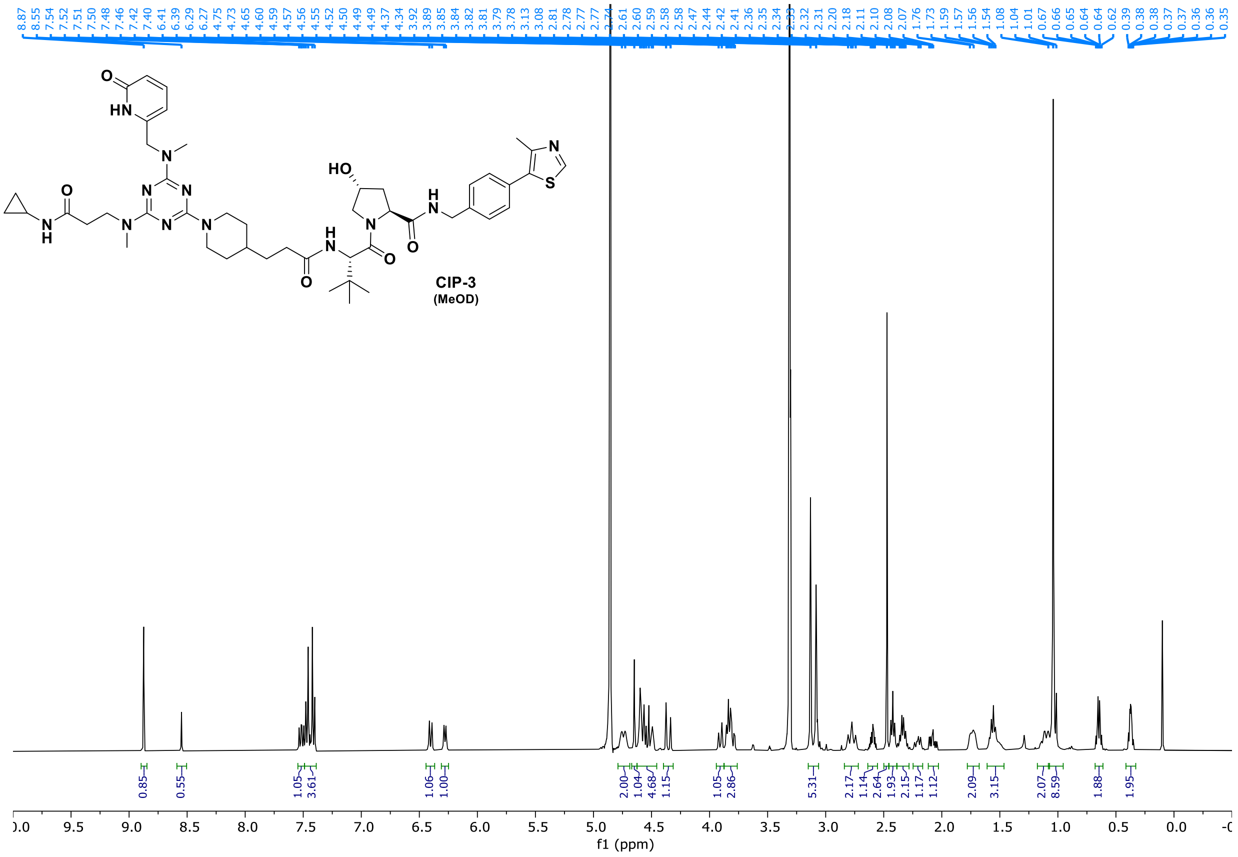
CIP-3
(MeOD)
f1 (ppm)

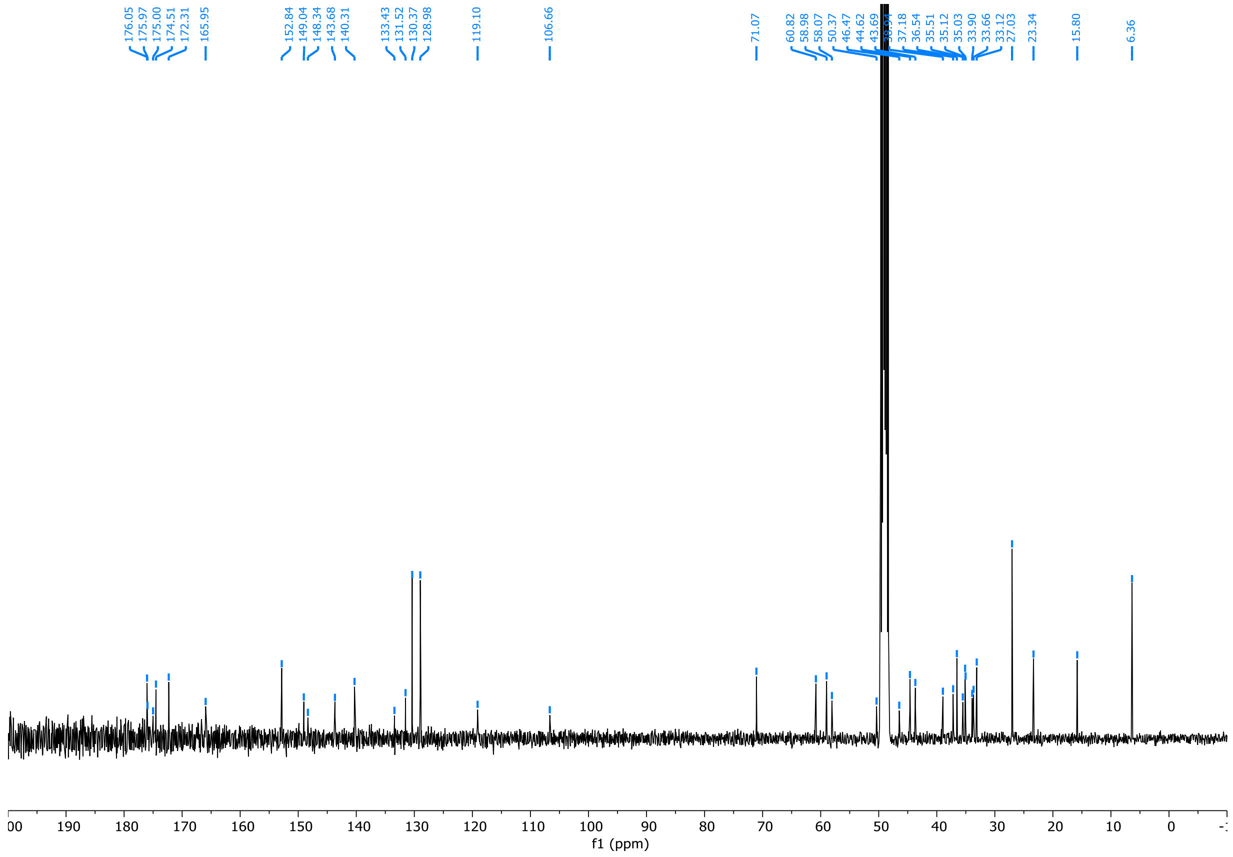
f1 (ppm)

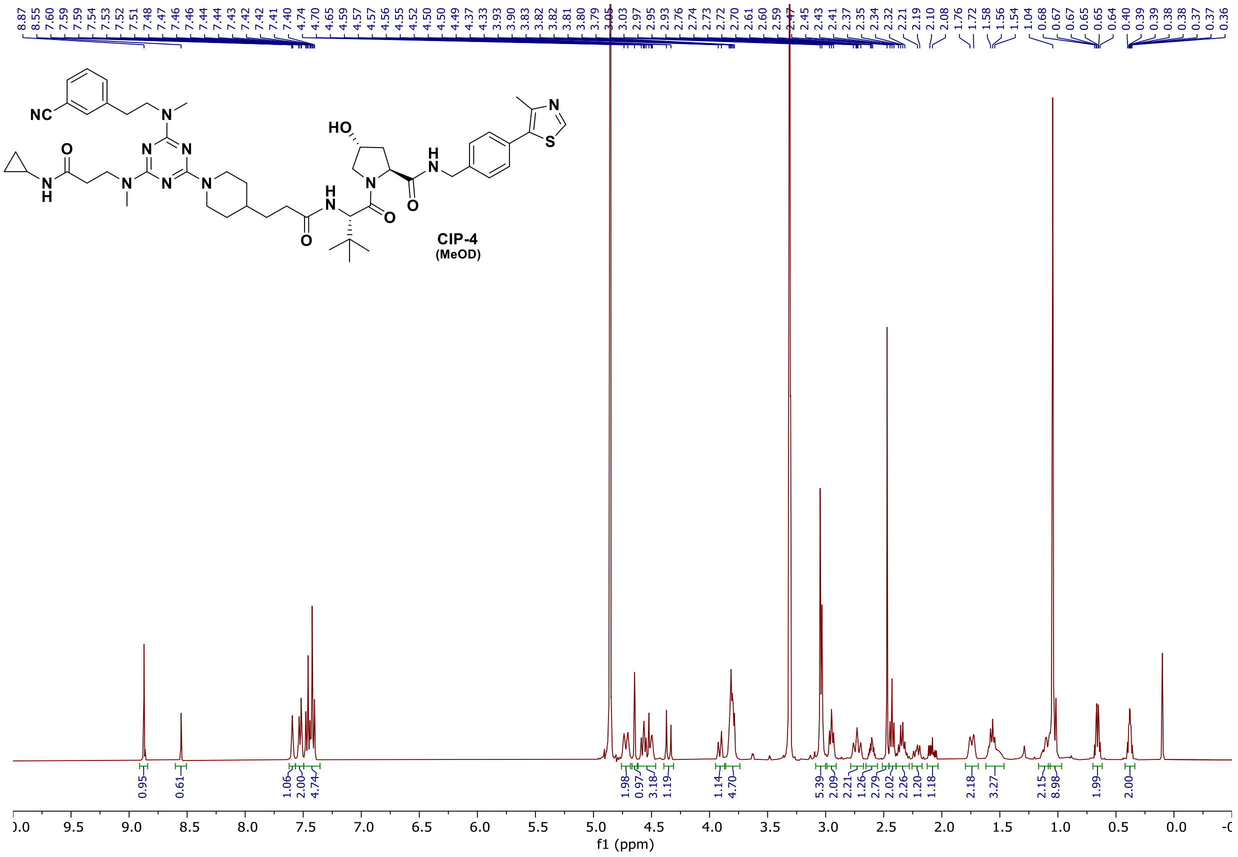

8.87 8.55 7.60 7.59 7.59 7.54 7.53 7.52 7.51 7.48 7.47 7.46 7.46 7.44 7.44 7.43 7.42 7.42 7.41 7.40 4.74 4.70 4.65 4.59 4.57 4.56 4.55 4.52 4.50 4.49 4.37 4.33 3.93 3.90 3.83 3.82 3.82 3.81 3.80 3.79 3.05 3.03 2.97 2.95 2.93 2.76 2.74 2.73 2.72 2.70 2.61 2.60 2.59 2.47 2.45 2.43 2.41 2.37 2.35 2.34 2.32 2.21 2.19 2.10 2.08 1.76 1.72 1.58 1.56 1.54 1.04 0.68 0.67 0.65 0.65 0.64 0.40 0.39 0.39 0.38 0.37 0.36
CIP-4
(MeOD)
0.95 0.61 1.06 2.00 4.74 1.98 0.97 3.18 1.19 1.14 4.70 5.39 2.09 2.21 1.26 2.79 2.02 2.26 1.20 1.18 2.18 3.27 2.15 8.98 1.99 2.00
f1 (ppm)


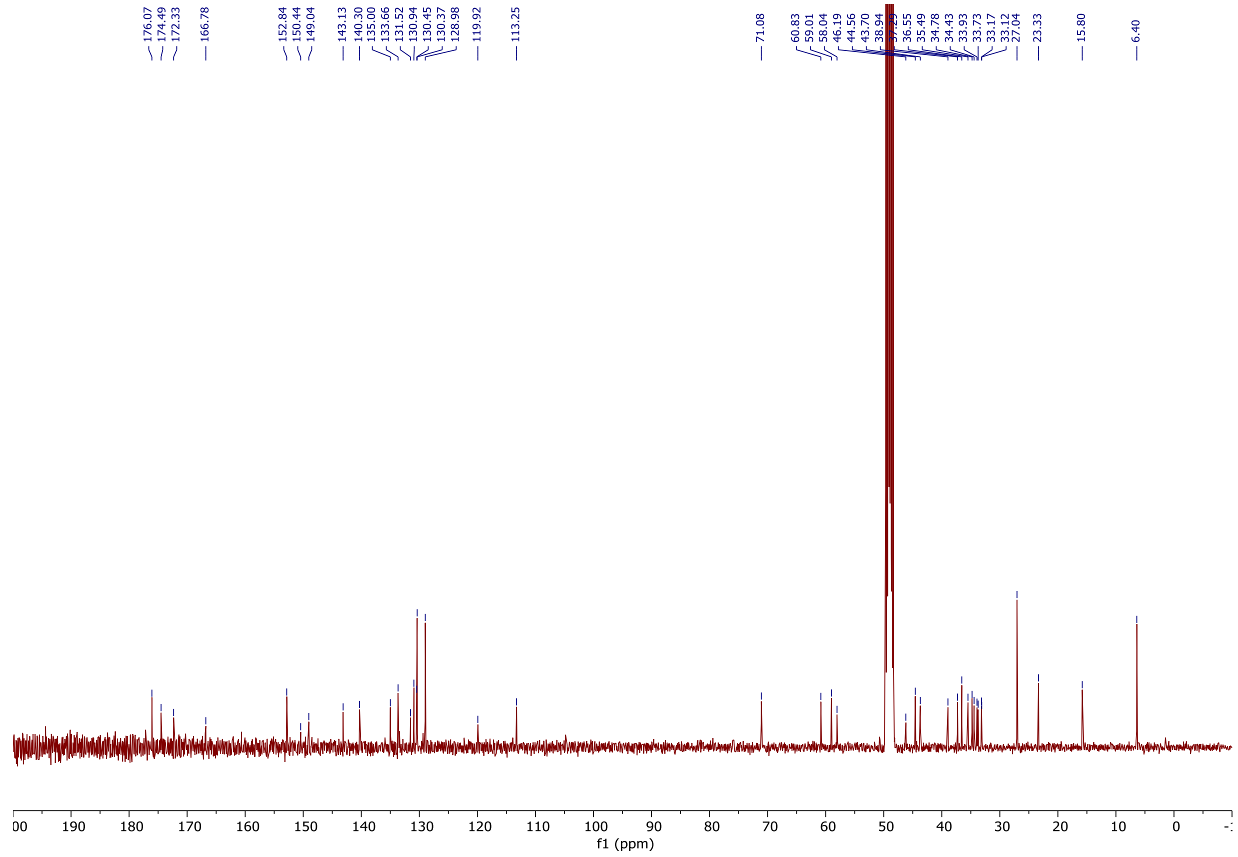

176.07 174.49 172.33 166.78 152.84 150.44 149.04 143.13 140.30 135.00 133.66 131.52 130.94 130.45 130.37 128.98 119.92 113.25 71.08 60.83 59.01 58.04 46.19 44.56 43.70 38.94 36.55 35.49 34.78 34.43 33.93 33.73 33.17 33.12 27.04 23.33 15.80 6.40
f1 (ppm)

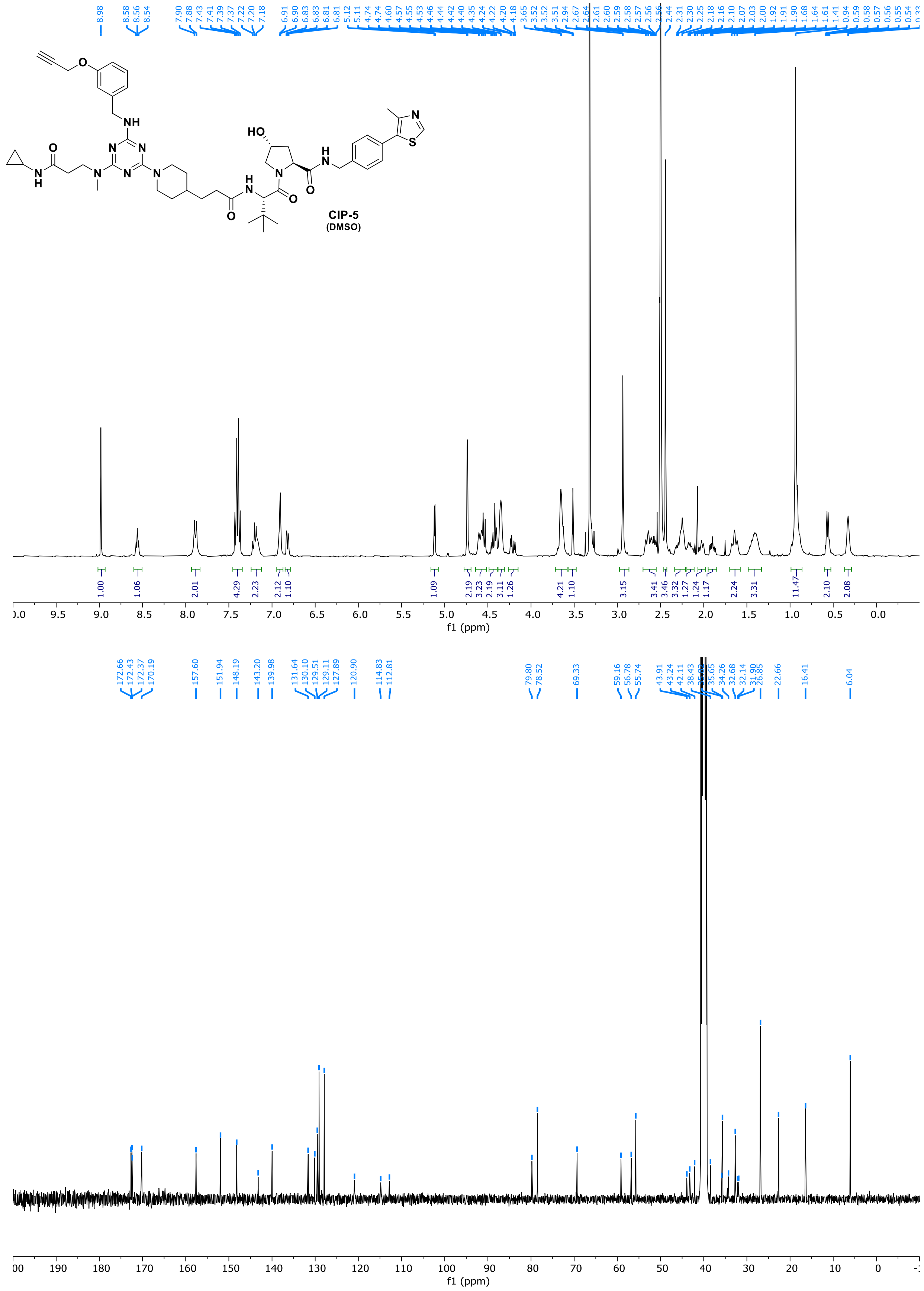
CIP-5
(DMSO)
f1 (ppm)
f1 (ppm)

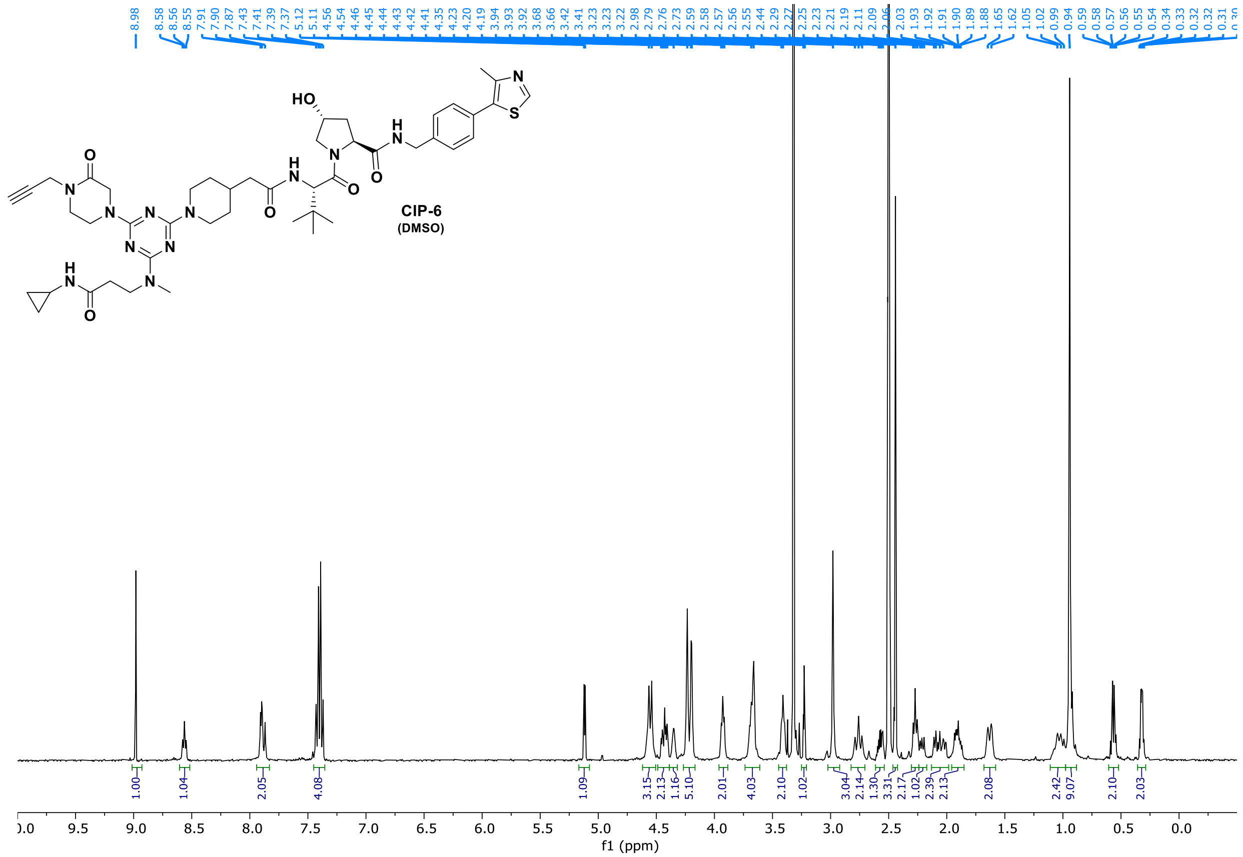
CIP-6
(DMSO)
f1 (ppm)

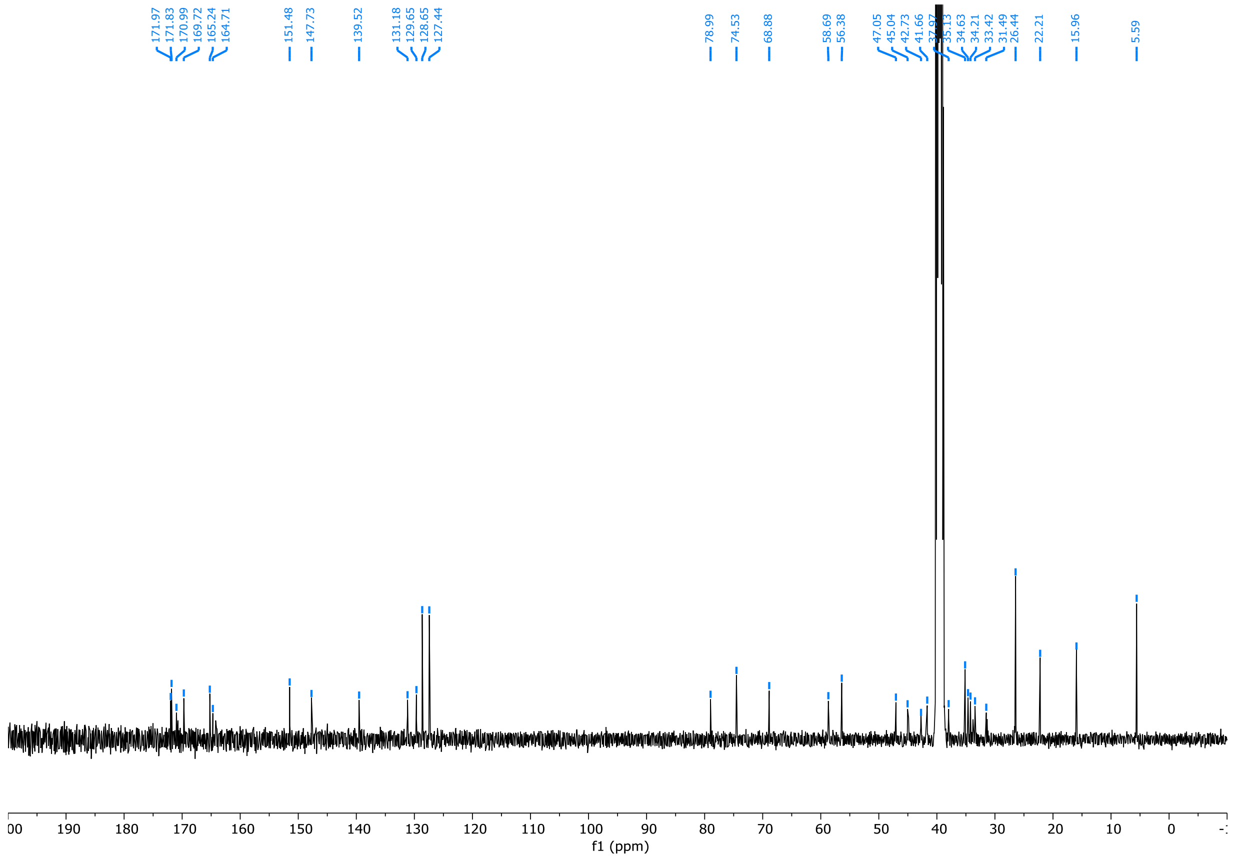
f1 (ppm)

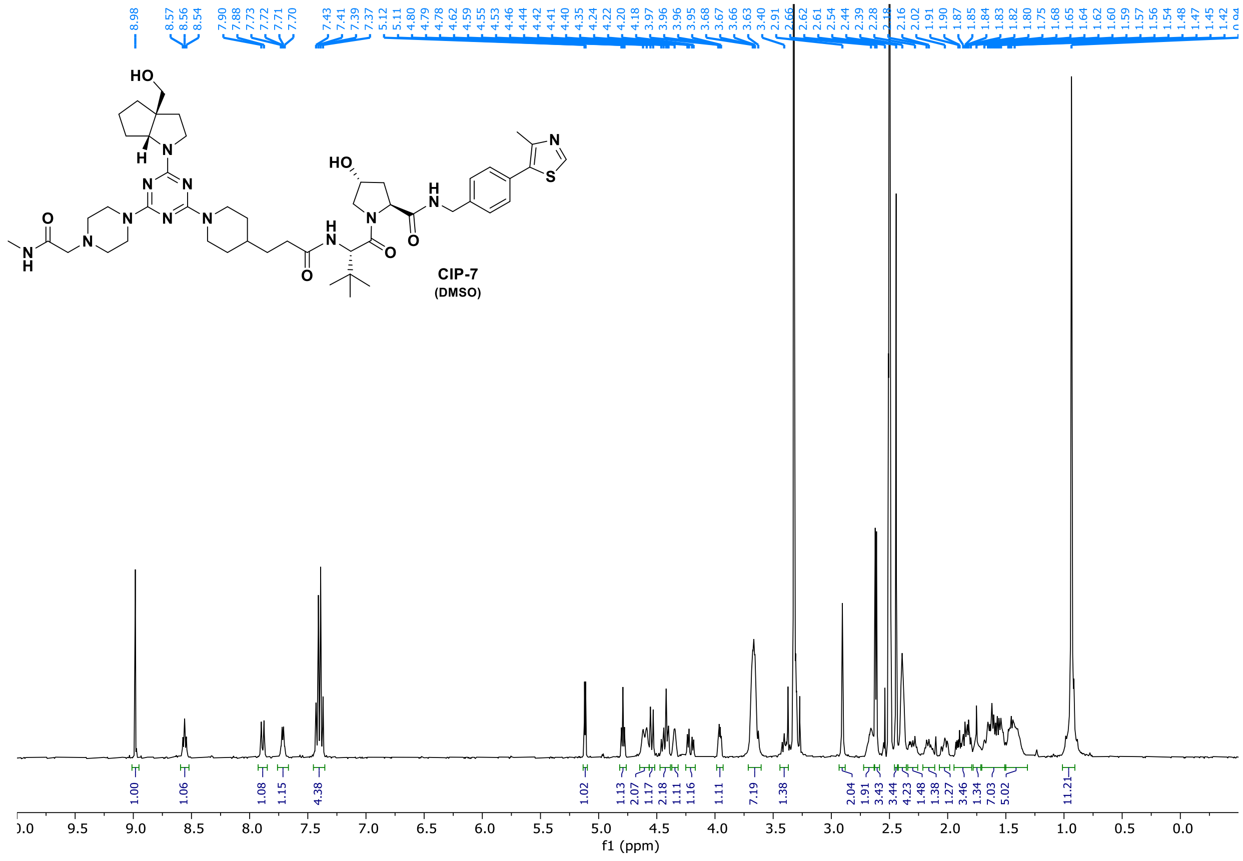
HO
H
N
HO
NH
O
CIP-7
(DMSO)
f1 (ppm)

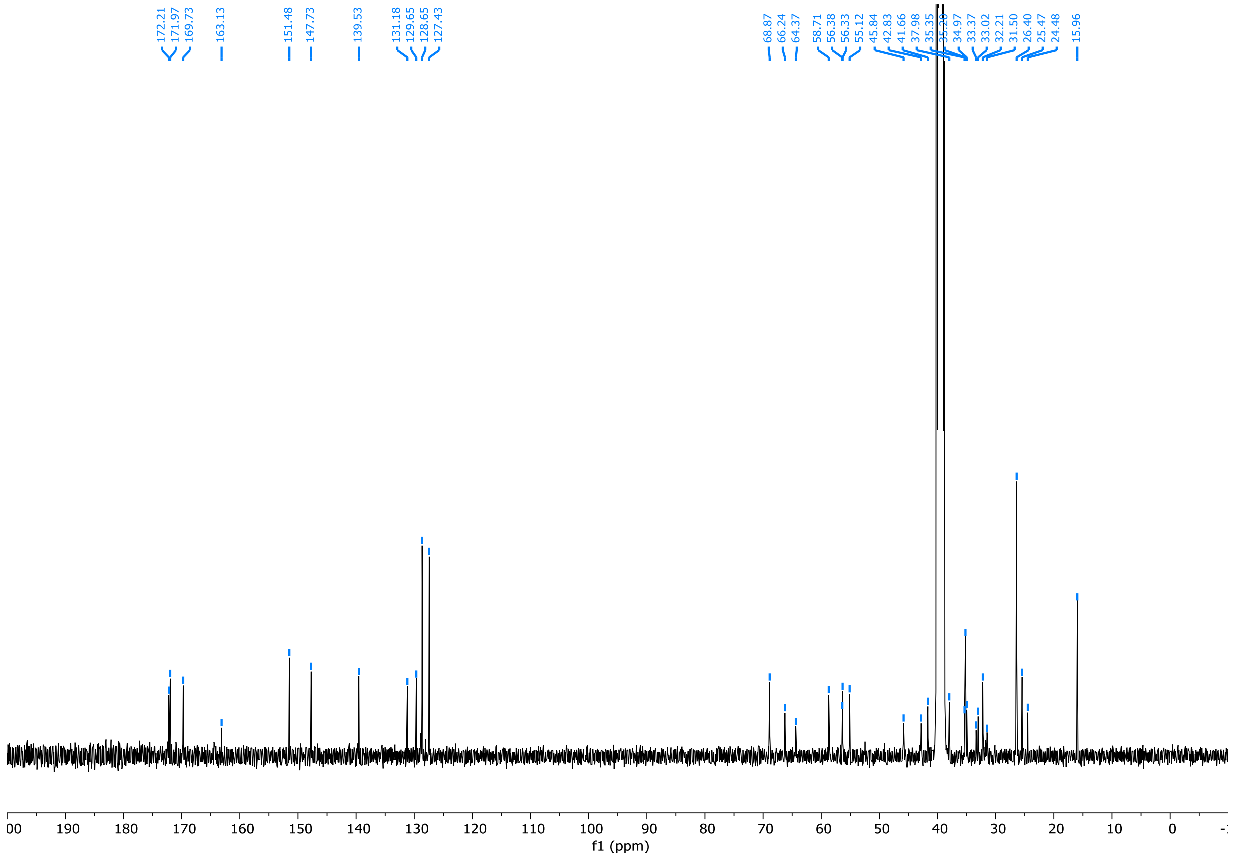
f1 (ppm)

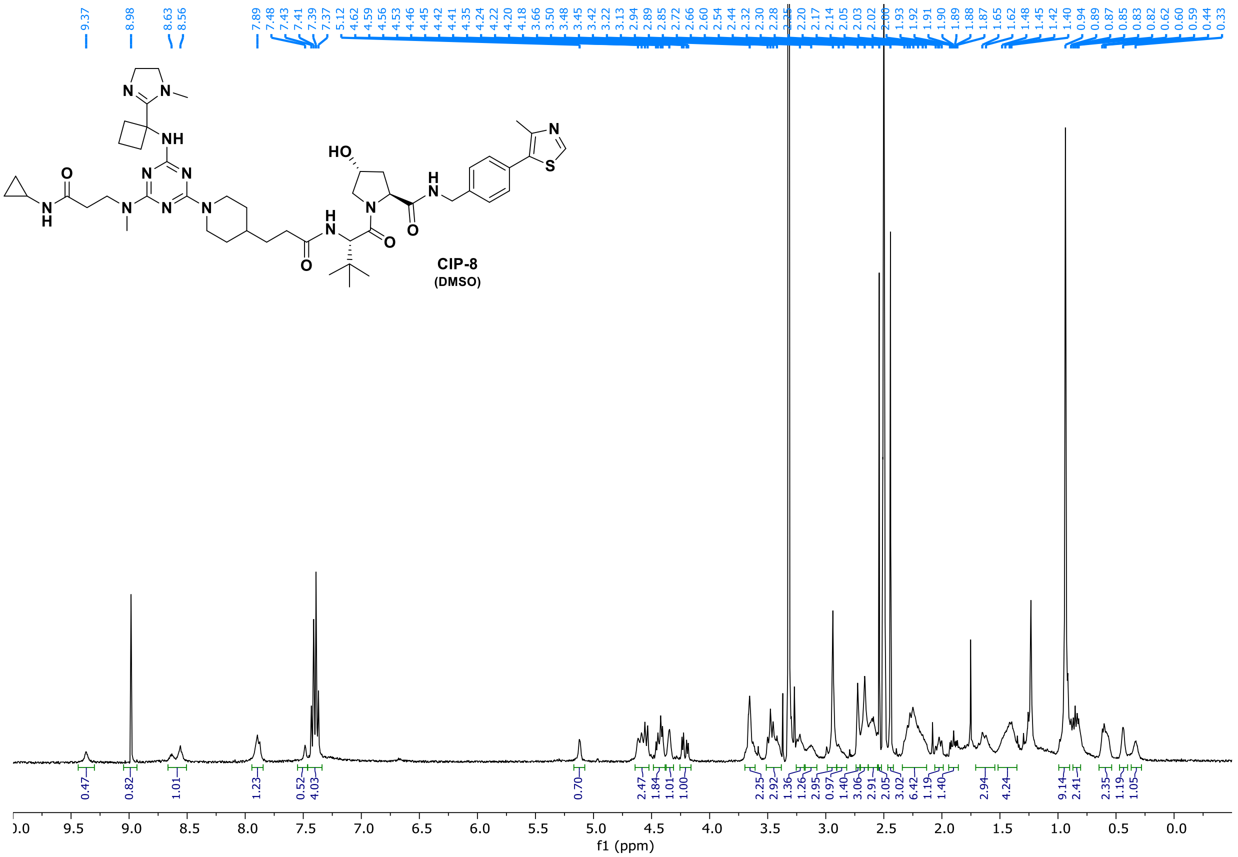
CIP-8
(DMSO)
f1 (ppm)

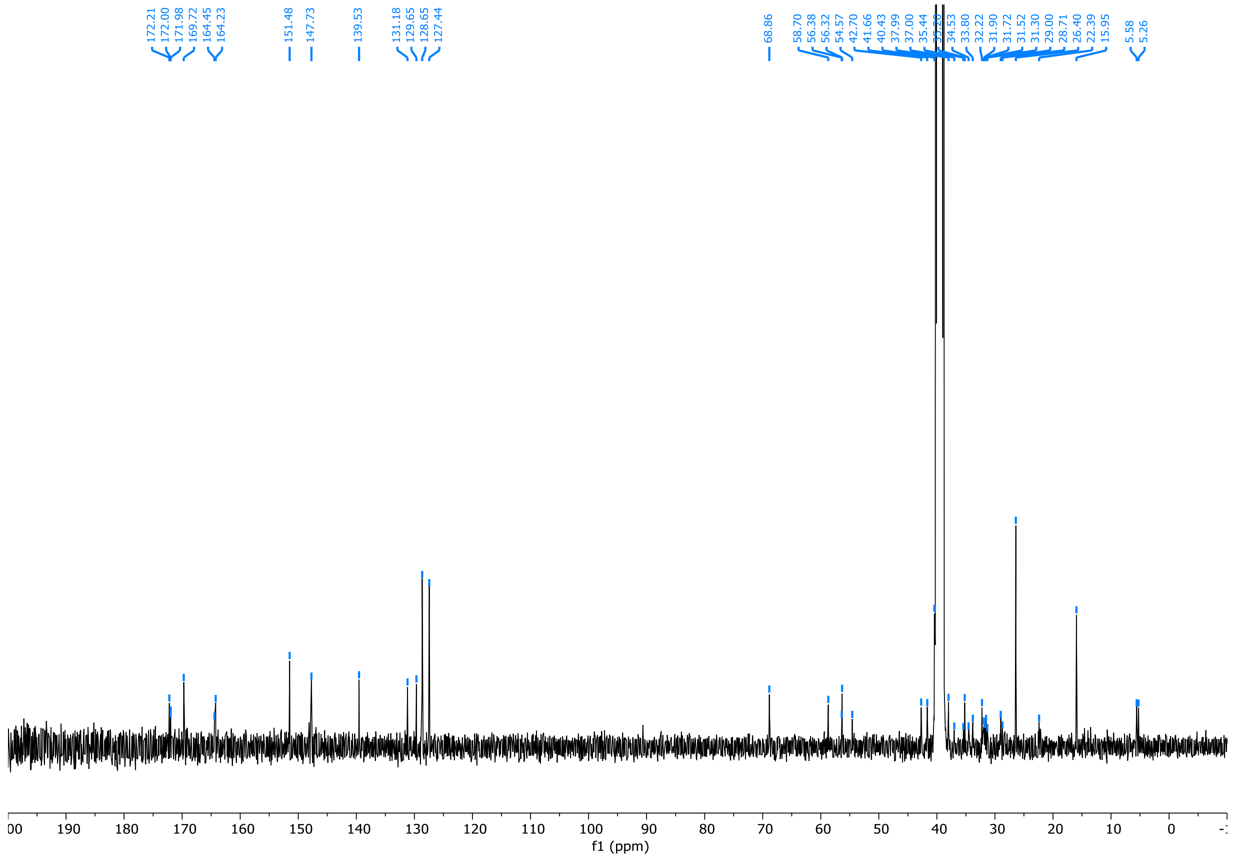
f1 (ppm)

**CIP-9**
**(DMSO)**

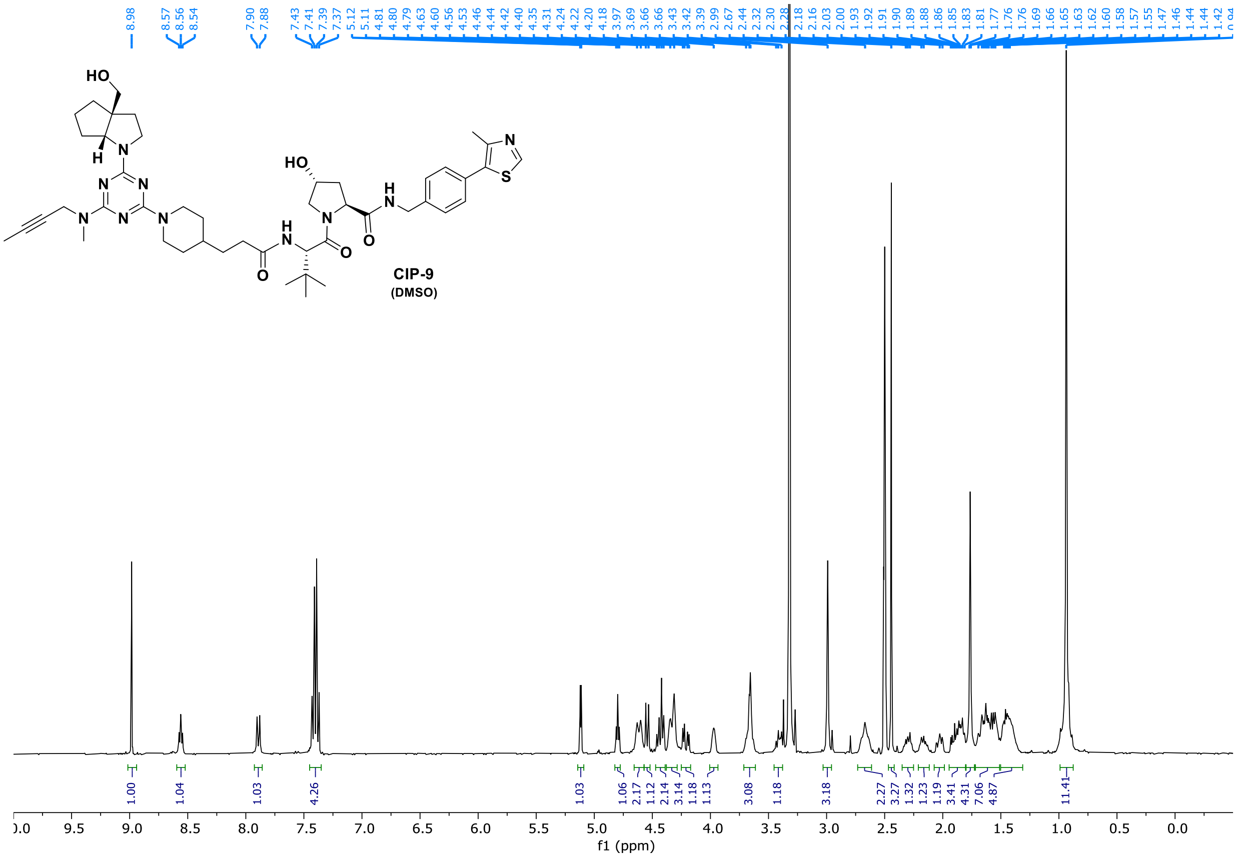

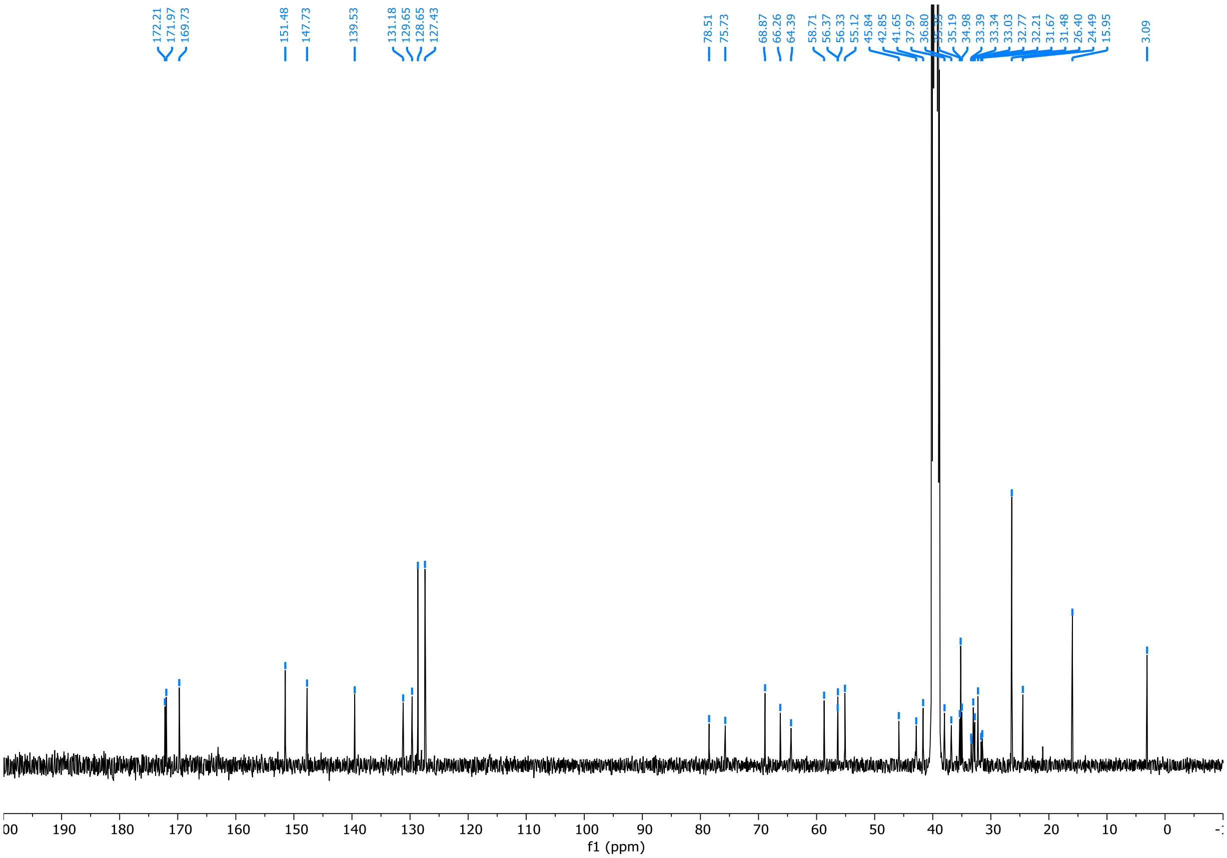

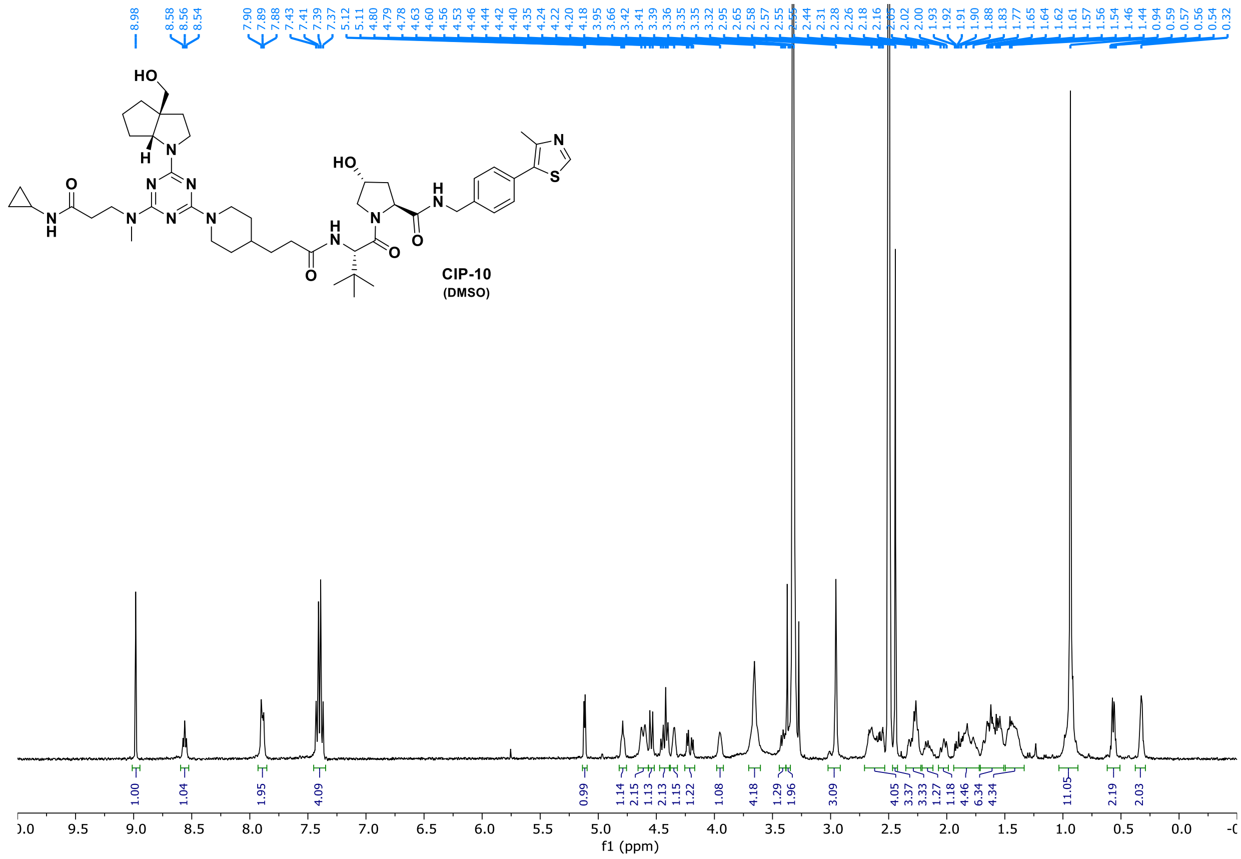

CIP-10
(DMSO)

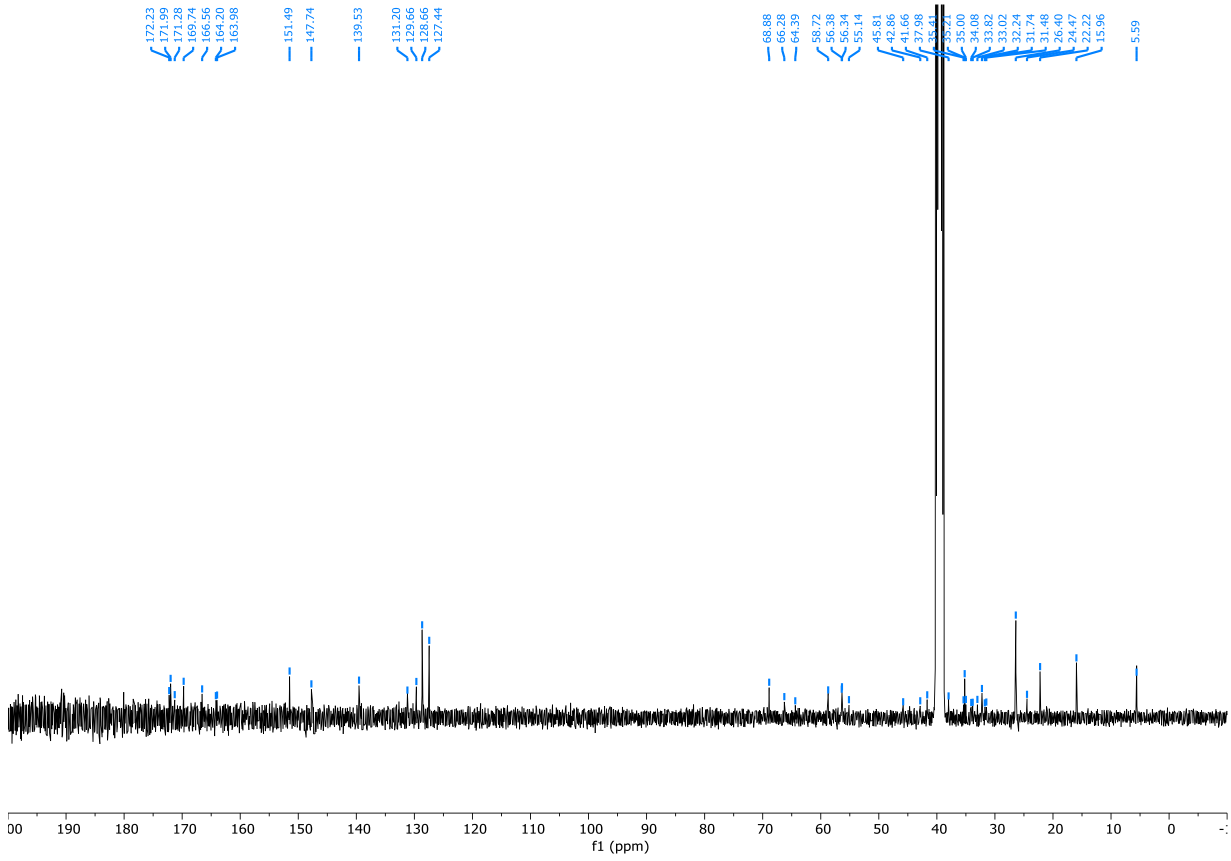

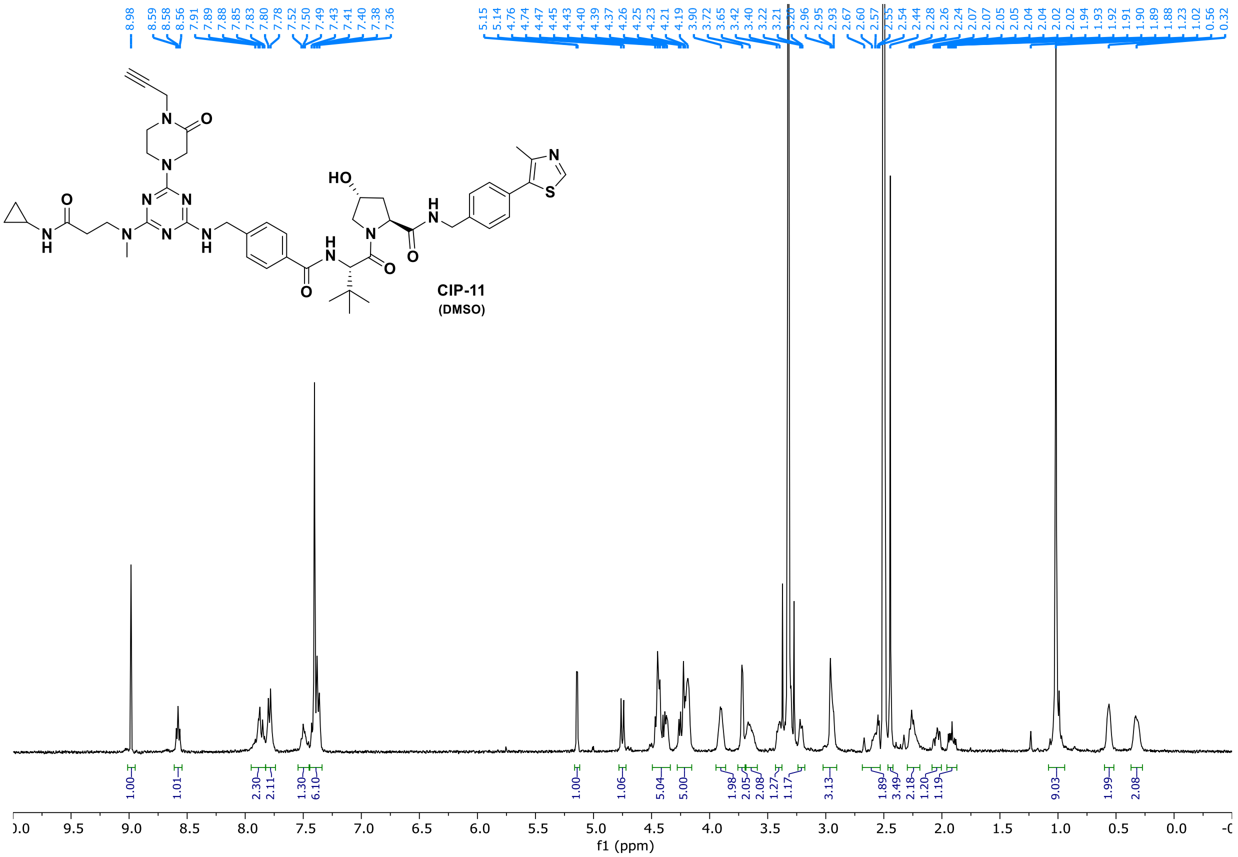

**CIP-11**
**(DMSO)**


8.98 8.59 8.58 8.56 7.91 7.89 7.88 7.85 7.83 7.80 7.78 7.52 7.50 7.49 7.43 7.41 7.40 7.38 7.36
5.15 5.14 4.76 4.74 4.47 4.45 4.43 4.40 4.39 4.37 4.26 4.25 4.23 4.21 4.19 3.90 3.72 3.65 3.42 3.40 3.22 3.21 2.96 2.95 2.93 2.67 2.60 2.57 2.55 2.54 2.44 2.28 2.26 2.24 2.07 2.07 2.05 2.05 2.04 2.04 2.02 2.02 1.94 1.93 1.92 1.91 1.90 1.89 1.88 1.23 1.02 0.56 0.32
1.00 1.01 2.30 2.11 1.30 6.10 1.00 1.06 5.04 5.00 1.98 2.05 2.08 1.27 1.17 3.13 1.89 3.49 2.18 1.20 1.19 9.03 1.99 2.08
f1 (ppm)


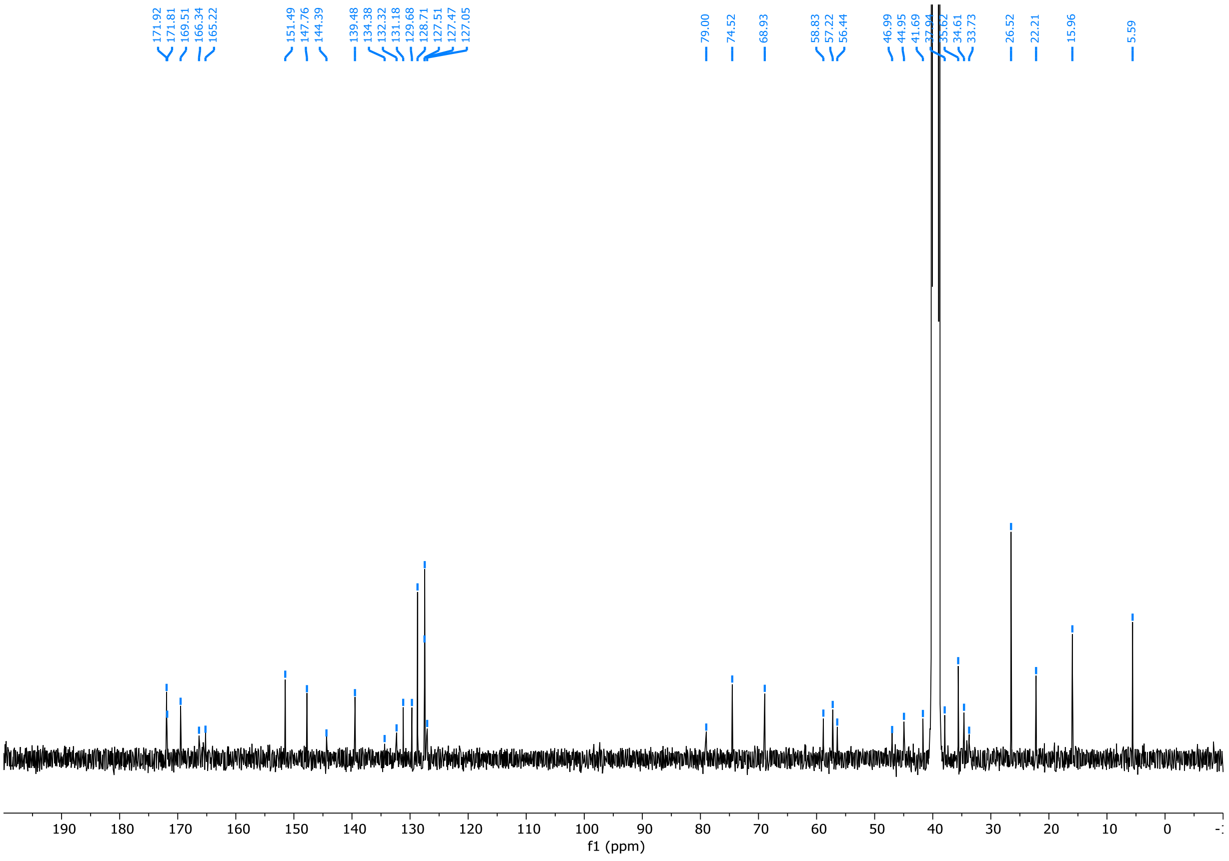

171.92 171.81 169.51 166.34 165.22 151.49 147.76 144.39 139.48 134.38 132.32 131.18 129.68 128.71 127.51 127.47 127.05
79.00 74.52 68.93 58.83 57.22 56.44 46.99 44.95 41.69 35.62 34.61 33.73 26.52 22.21 15.96 5.59
f1 (ppm)

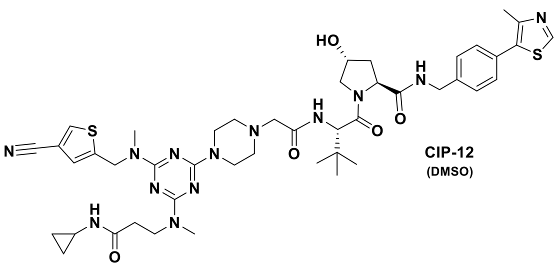
HO
N
H
N
S
N
N
H
O
O
N
N
N
N
N
N
S
N
H
N
O
N
O

**CIP-12**
(DMSO)

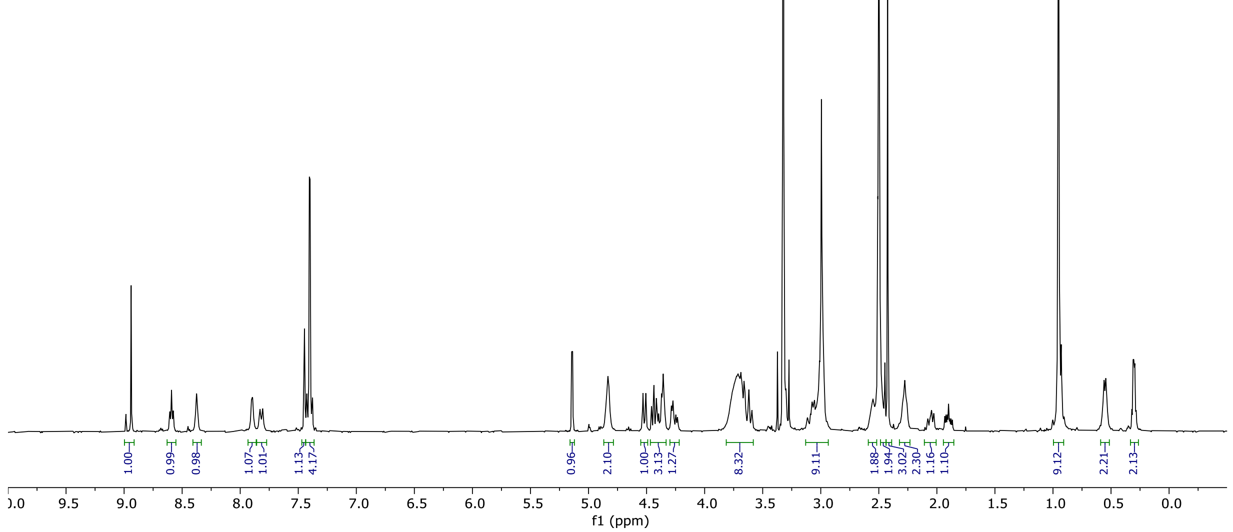
8.94
8.61
8.59
8.58
8.38
7.90
7.83
7.81
7.45
7.43
7.40
7.40
7.38
5.15
5.14
4.83
4.53
4.51
4.46
4.44
4.41
4.37
4.36
4.29
4.27
4.25
4.23
3.72
3.69
3.68
3.66
3.65
3.62
3.59
3.11
3.07
3.05
3.01
2.99
2.55
2.54
2.48
2.45
2.42
2.29
2.28
2.26
2.08
2.06
2.06
2.05
2.03
1.93
1.92
1.91
1.90
1.89
1.88
1.87
0.95
0.56
0.54
0.32
0.31
0.30
0.30
0.28
1.00
0.99
0.98
1.07
1.01
1.13
4.17
0.96
2.10
1.00
3.13
1.27
8.32
9.11
1.88
1.94
3.02
2.30
1.16
1.10
9.12
2.21
2.13
9.5
9.0
8.5
8.0
7.5
7.0
6.5
6.0
5.5
5.0
4.5
4.0
3.5
3.0
2.5
2.0
1.5
1.0
0.5
0.0
f1 (ppm)

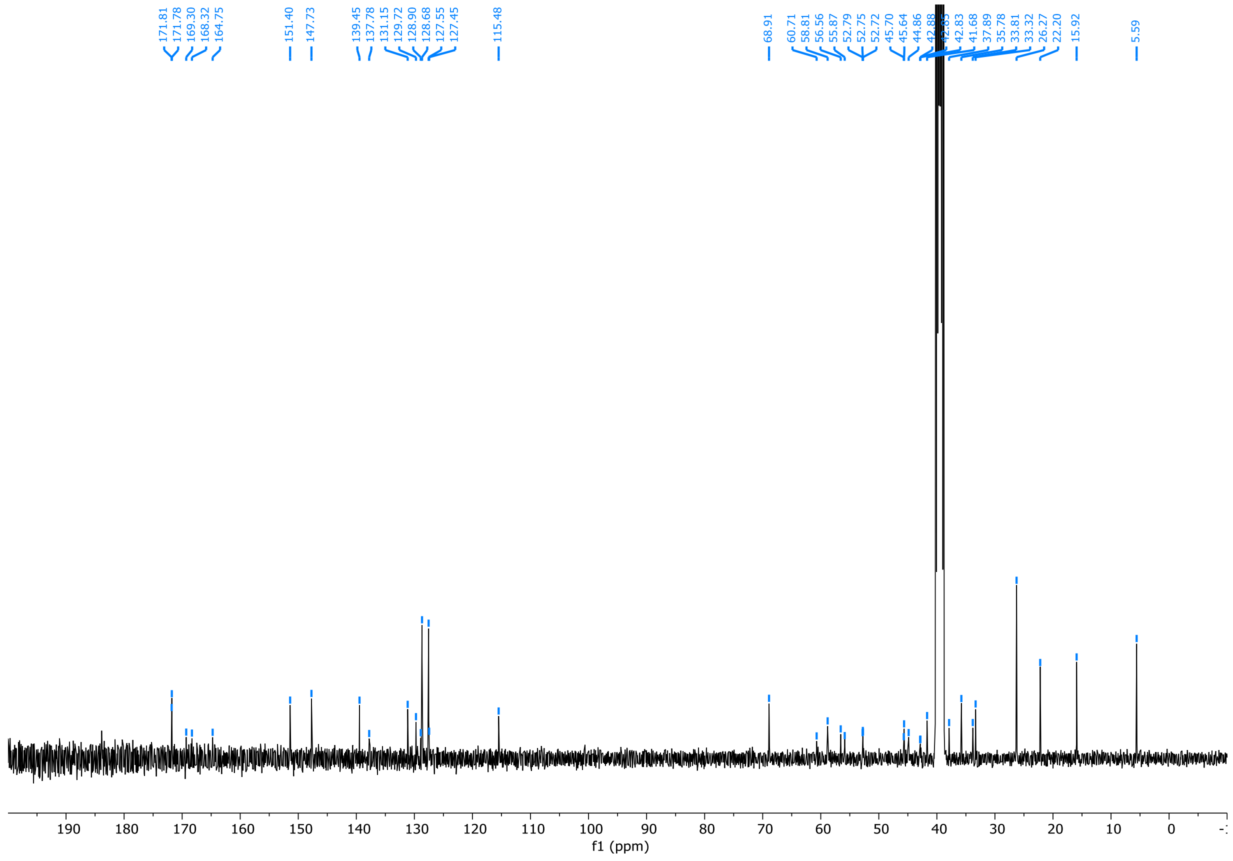
171.81
171.78
169.30
168.32
164.75
151.40
147.73
139.45
137.78
131.15
129.72
128.90
128.68
127.55
127.45
115.48
68.91
60.71
58.81
56.56
55.87
52.79
52.75
52.72
45.70
45.64
44.86
42.83
41.68
37.89
35.78
33.81
33.32
26.27
22.20
15.92
5.59
190
180
170
160
150
140
130
120
110
100
90
80
70
60
50
40
30
20
10
0
f1 (ppm)

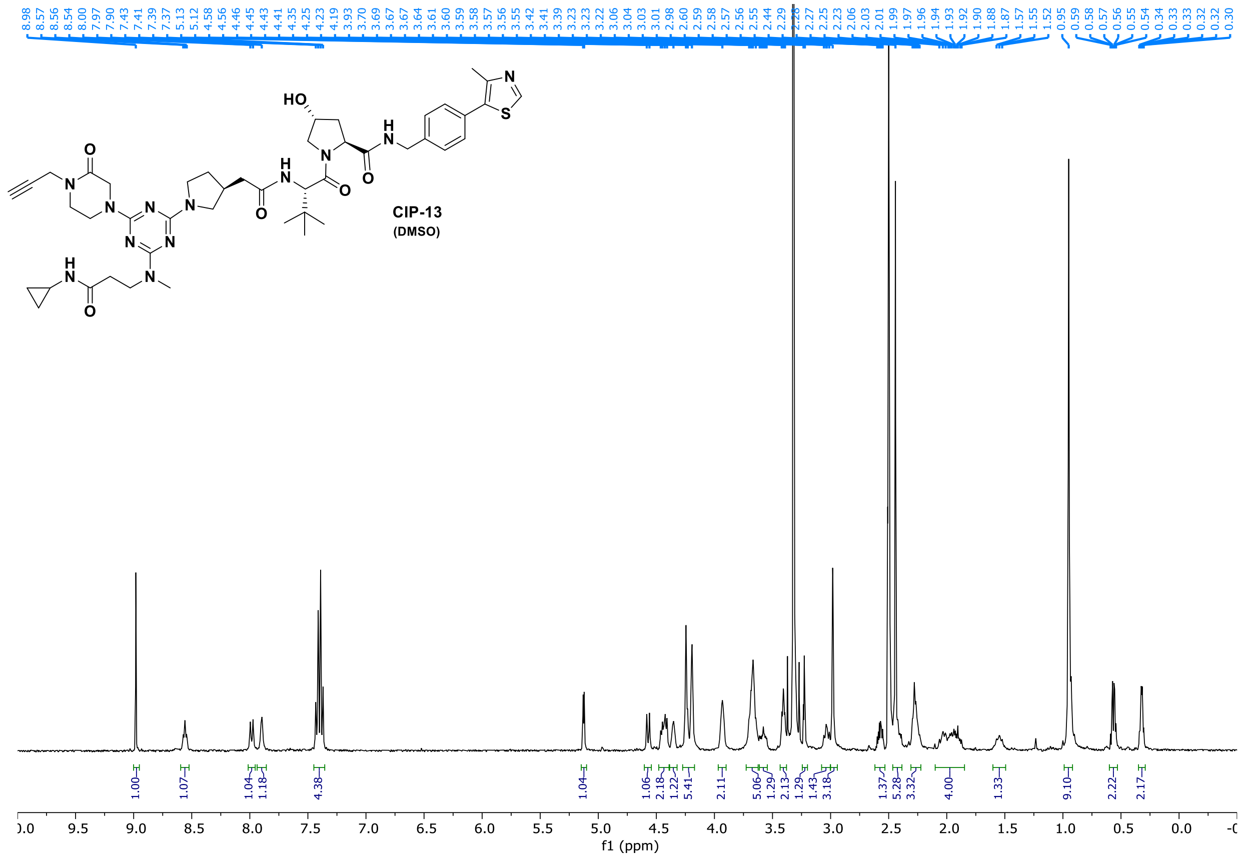

HO
N
S
H
N
O
N
N
N
N
N
N
N
N
H
N
H
O
O
O
8.98 8.57 8.56 8.54 8.00 7.97 7.90 7.43 7.41 7.39 7.37 5.13 5.12 4.58 4.56 4.46 4.45 4.43 4.41 4.35 4.25 4.23 4.19 3.93 3.70 3.69 3.67 3.67 3.64 3.61 3.60 3.59 3.58 3.57 3.56 3.55 3.42 3.41 3.39 3.23 3.22 3.06 3.04 3.03 3.01 2.98 2.60 2.59 2.58 2.57 2.56 2.55 2.44 2.29 2.27 2.25 2.23 2.06 2.03 2.01 1.99 1.97 1.96 1.94 1.93 1.92 1.90 1.88 1.87 1.57 1.55 1.52 0.95 0.59 0.58 0.57 0.56 0.55 0.54 0.34 0.33 0.32 0.32 0.30
1.00 1.07 1.04 1.18 4.38 1.04 1.06 2.18 1.22 5.41 2.11 5.06 1.29 2.13 1.29 1.43 3.18 1.37 5.28 3.32 4.00 1.33 9.10 2.22 2.17
9.5 9.0 8.5 8.0 7.5 7.0 6.5 6.0 5.5 5.0 4.5 4.0 3.5 3.0 2.5 2.0 1.5 1.0 0.5 0.0
f1 (ppm)


CIP-13
(DMSO)

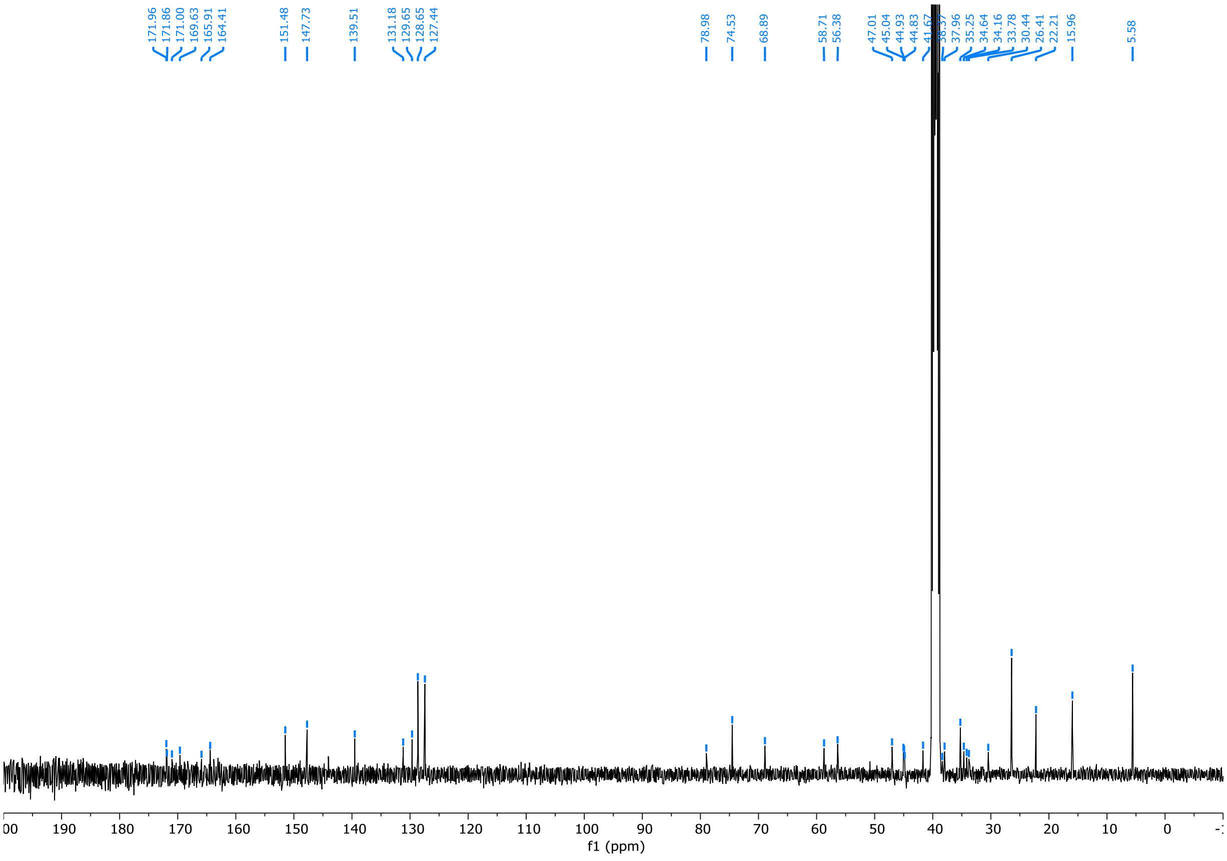

171.96 171.86 171.00 169.63 165.91 164.41 151.48 147.73 139.51 131.18 129.65 128.65 127.44 78.98 74.53 68.89 58.71 56.38 47.01 45.04 44.93 44.83 41.67 37.96 35.25 34.64 34.16 33.78 30.44 26.41 22.21 15.96 5.58
190 180 170 160 150 140 130 120 110 100 90 80 70 60 50 40 30 20 10 0
f1 (ppm)

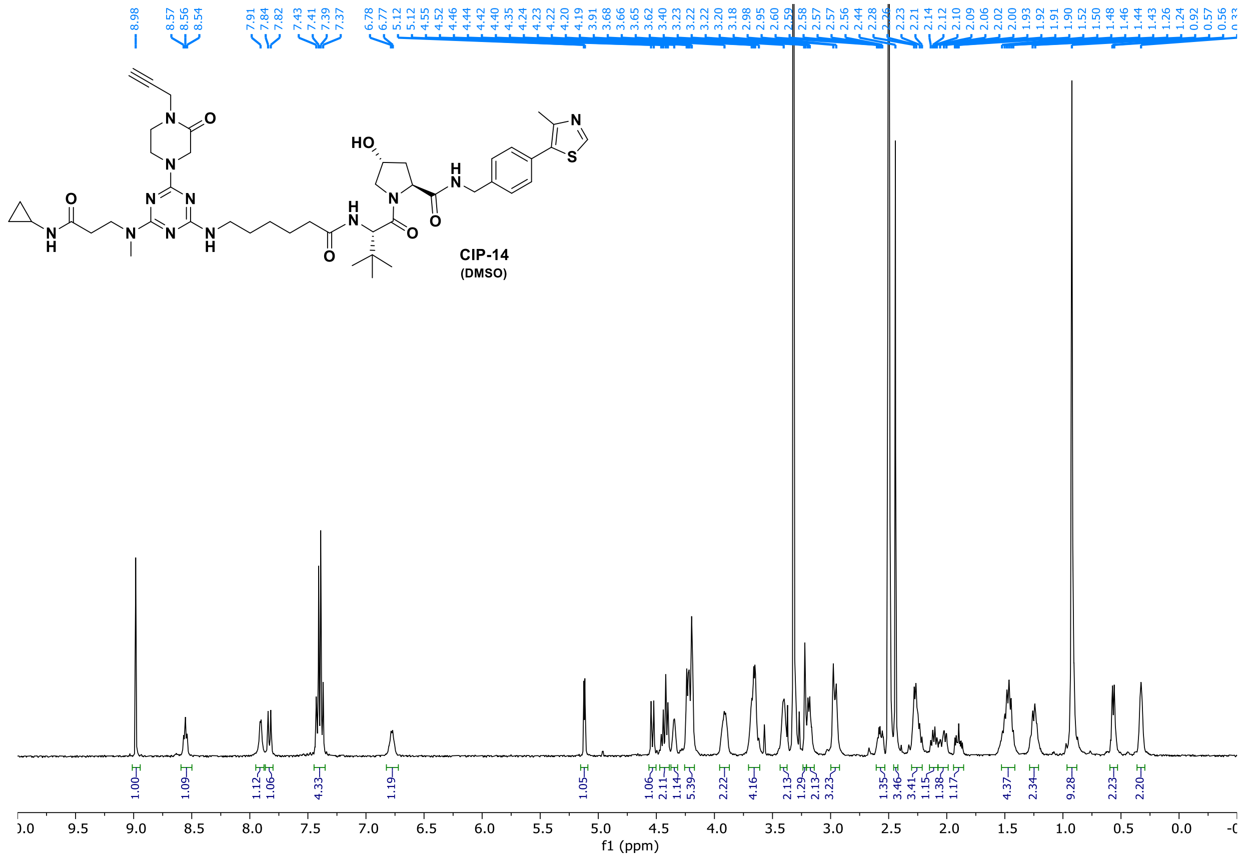
CIP-14
(DMSO)
8.98
8.57 8.56 8.54
7.91 7.84 7.82
7.43 7.41 7.39 7.37
6.78 6.77
5.12 5.12
f1 (ppm)

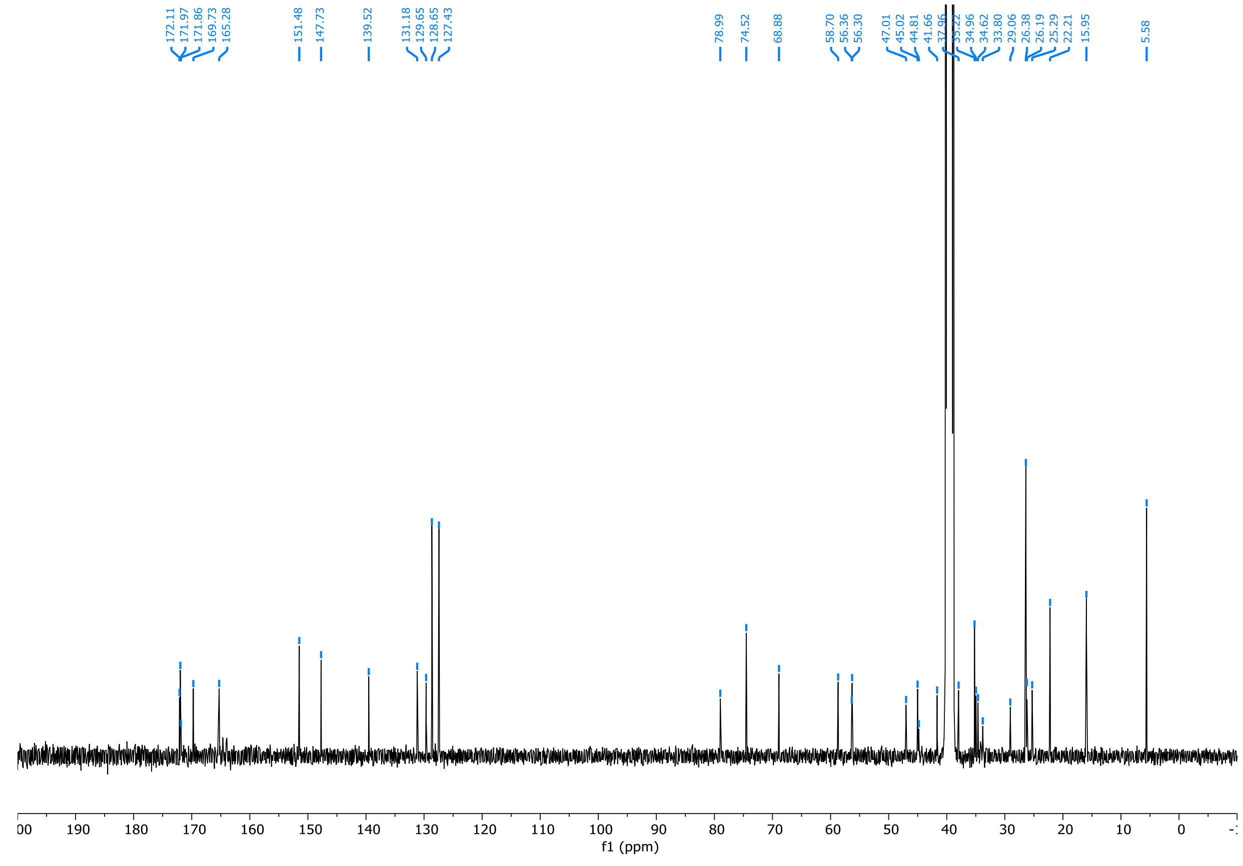
172.11 171.97 171.86 169.73 165.28
151.48 147.73
139.52
131.18 129.65 128.65 127.43
78.99 74.52 68.88
58.70 56.36 56.30
47.01 45.02 44.81 41.66
34.96 34.62 33.80 29.06 26.38 26.19 25.29 22.21 15.95
5.58
f1 (ppm)

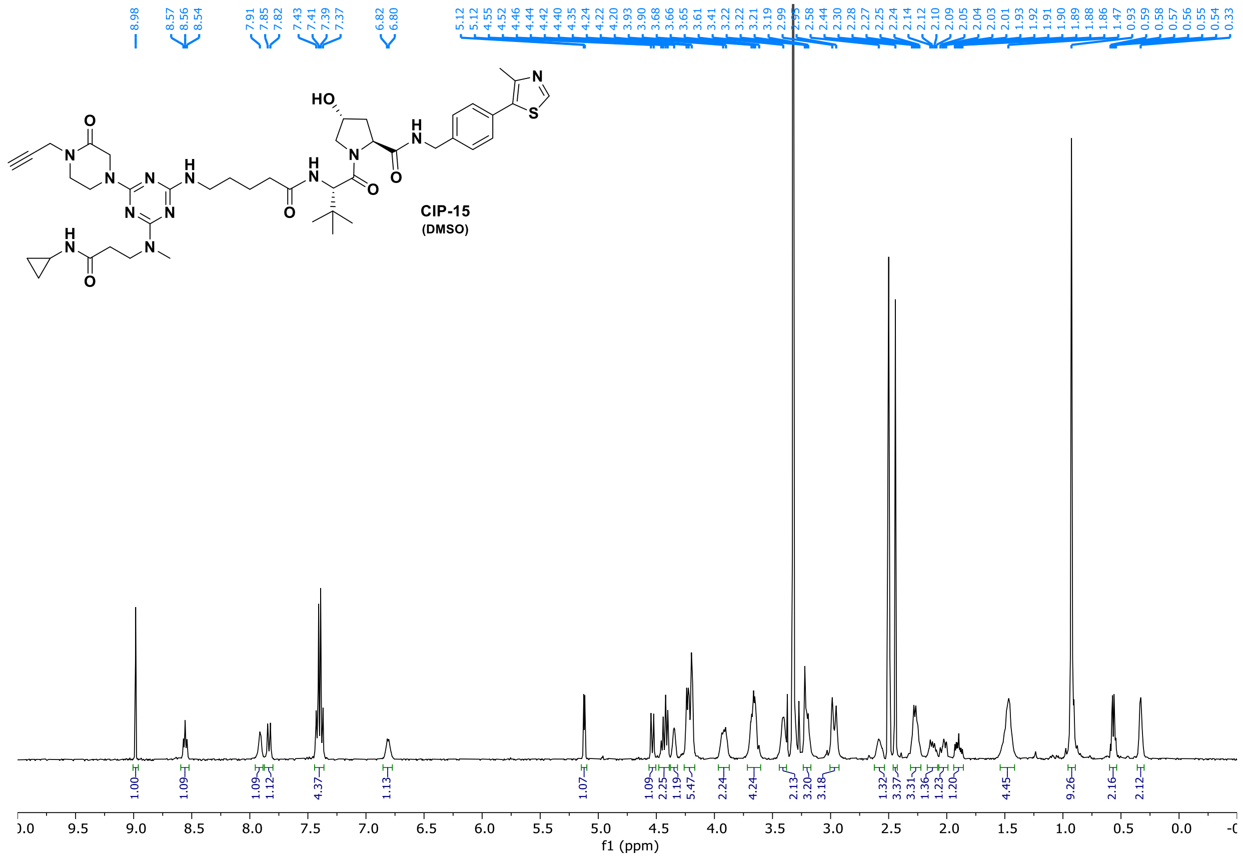


CIP-15
(DMSO)

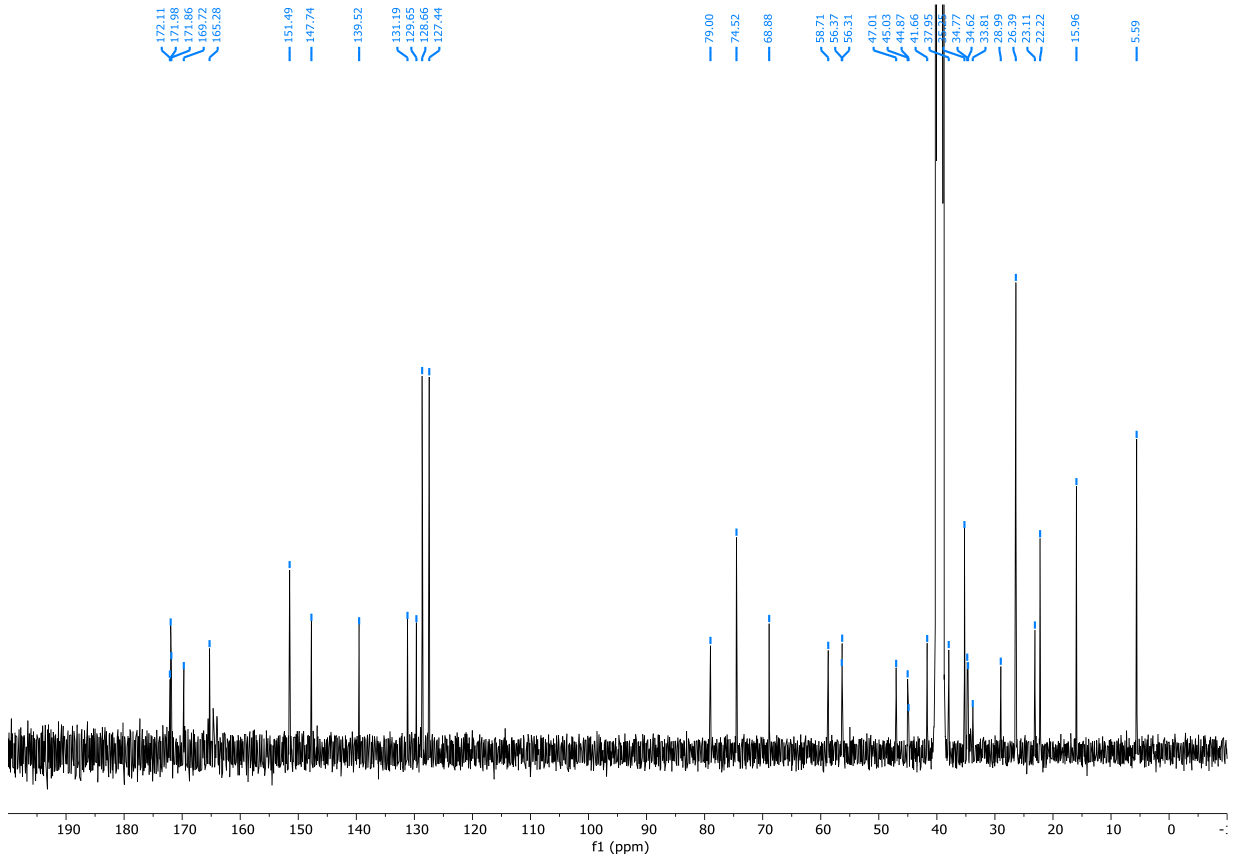

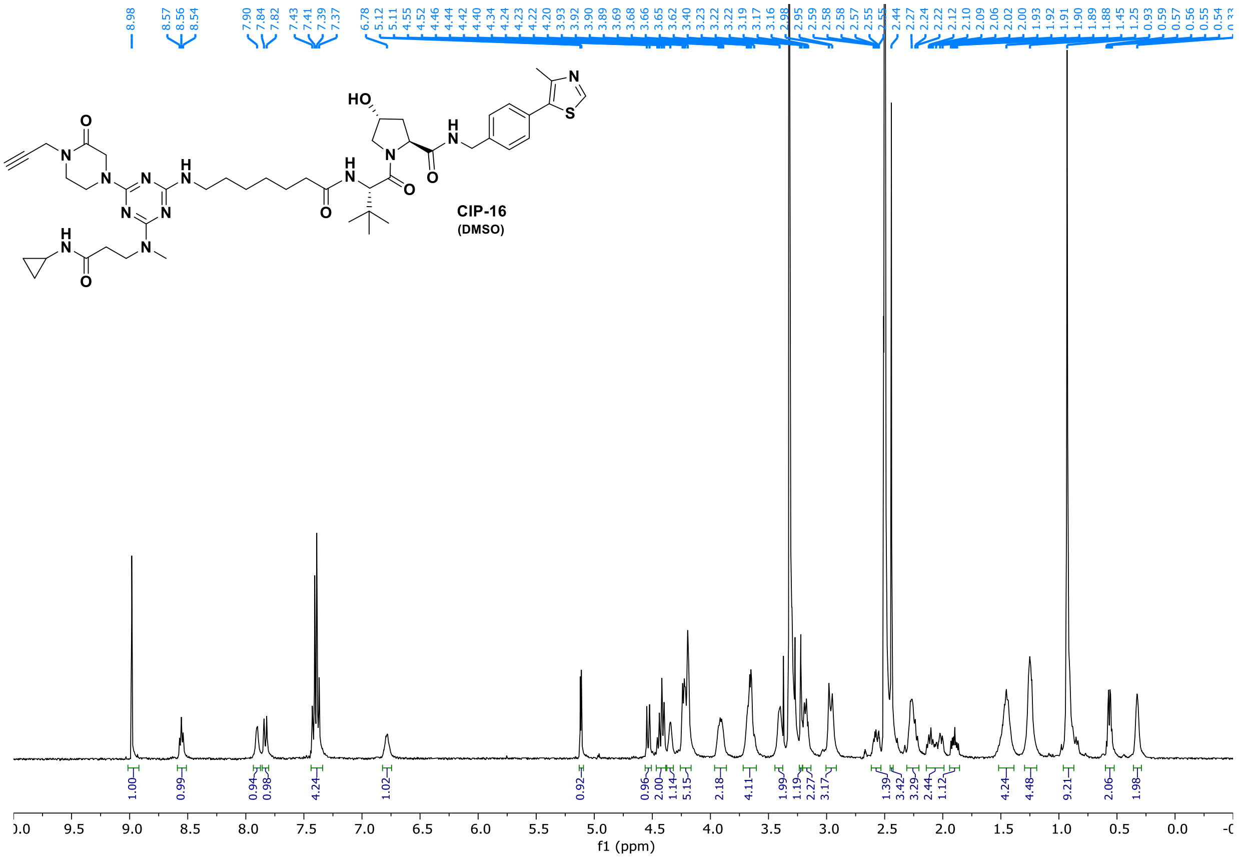
CIP-16
(DMSO)
f1 (ppm)

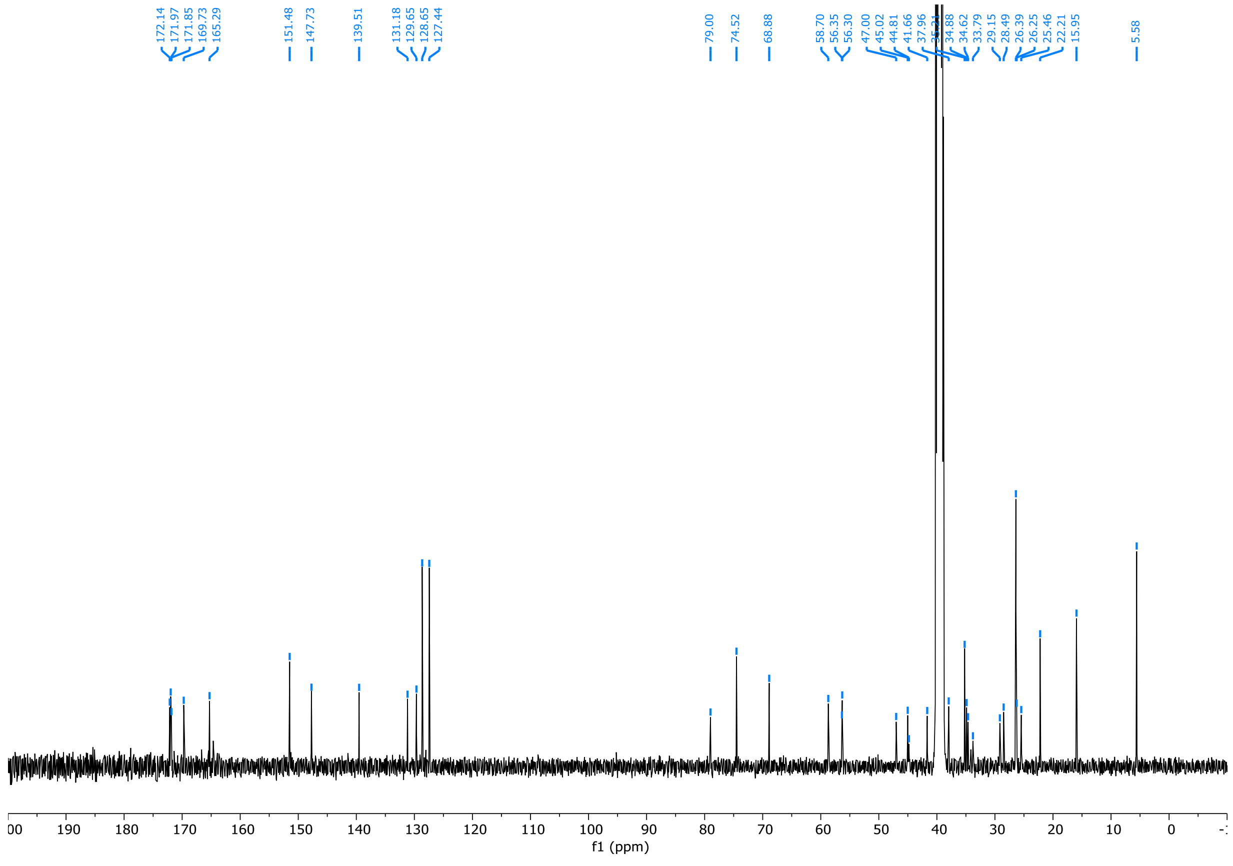
f1 (ppm)

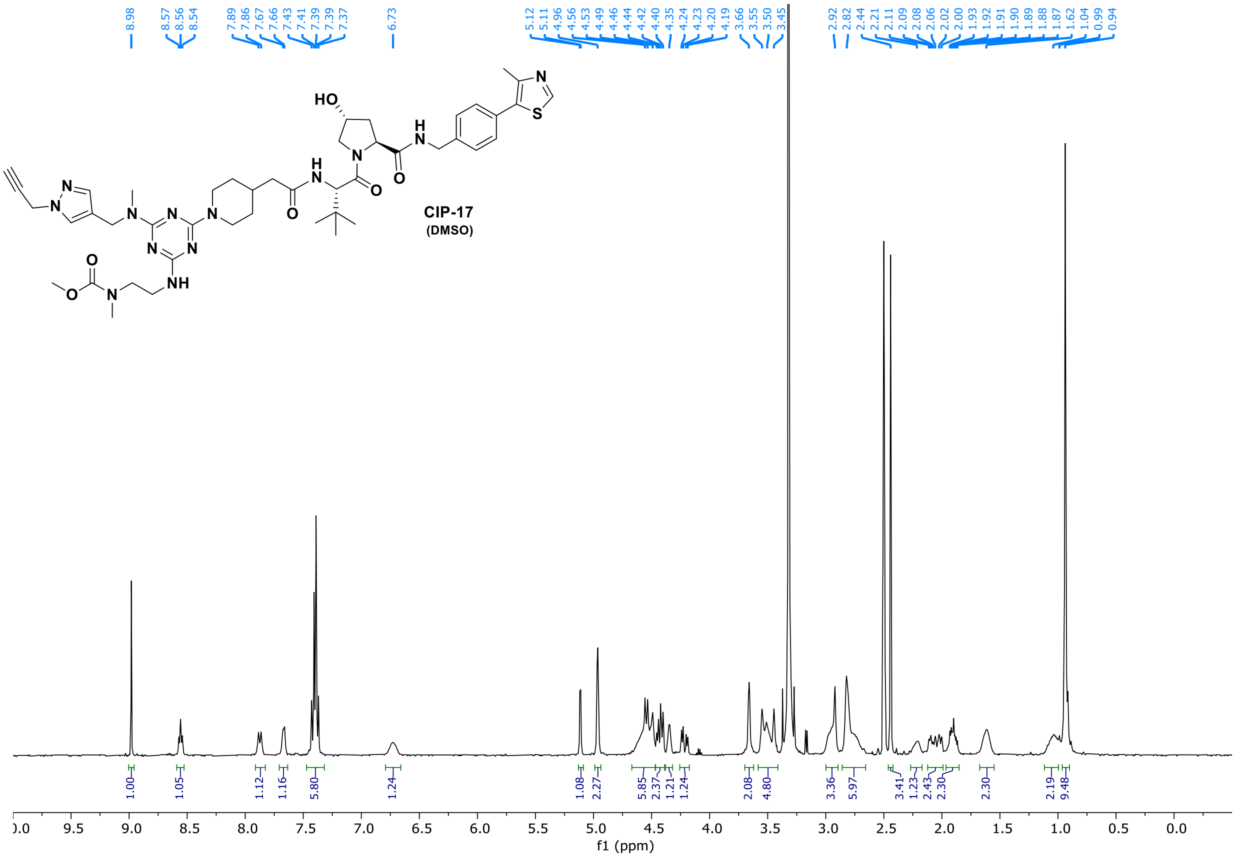
8.98
8.57
8.56
8.54
7.89
7.86
7.67
7.66
7.43
7.41
7.39
7.39
7.37
6.73
5.12
5.11
4.96
4.56
4.53
4.49
4.46
4.44
4.42
4.40
4.35
4.24
4.23
4.20
4.19
3.66
3.55
3.50
3.45
2.92
2.82
2.44
2.21
2.11
2.09
2.08
2.06
2.02
2.00
1.93
1.92
1.91
1.90
1.89
1.88
1.87
1.62
1.04
0.99
0.94
HO
N
H
N
S
N
N
N
N
N
N
N
N
H
NH
O
O
O
O
O
N
CIP-17
(DMSO)
1.00
1.05
1.12
1.16
5.80
1.24
1.08
2.27
5.85
2.37
1.21
1.24
2.08
4.80
3.36
5.97
3.41
1.23
2.43
2.30
2.30
2.19
9.48
).0
9.5
9.0
8.5
8.0
7.5
7.0
6.5
6.0
5.5
5.0
4.5
4.0
3.5
3.0
2.5
2.0
1.5
1.0
0.5
0.0
f1 (ppm)

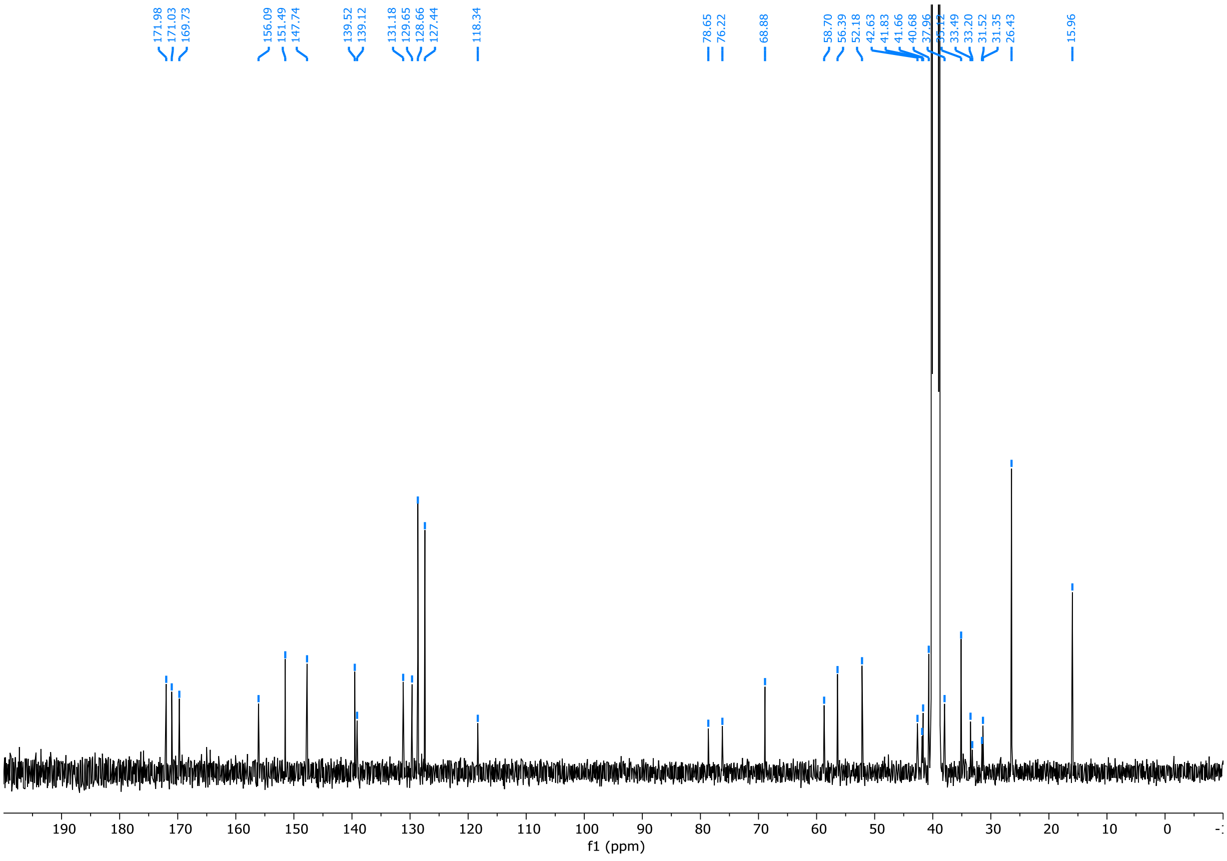
171.98
171.03
169.73
156.09
151.49
147.74
139.52
139.12
131.18
129.65
128.66
127.44
118.34
78.65
76.22
68.88
58.70
56.39
52.18
42.63
41.83
41.66
40.68
37.96
35.12
33.49
33.20
31.52
31.35
26.43
15.96
190
180
170
160
150
140
130
120
110
100
90
80
70
60
50
40
30
20
10
0
f1 (ppm)

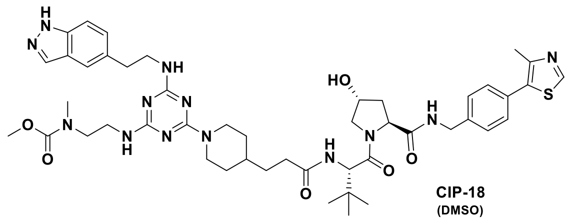
HO
NH
N
S
O
N
H


**CIP-18**
(DMSO)

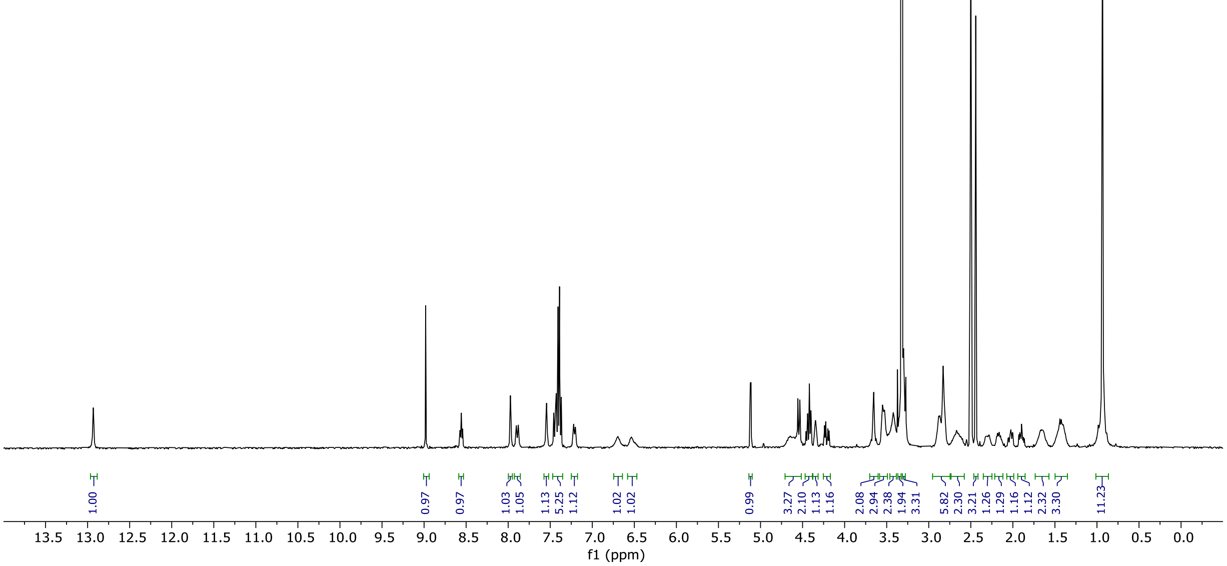
12.93
8.98
8.57
8.56
8.54
7.97
7.90
7.88
7.54
7.46
7.44
7.43
7.41
7.39
7.37
7.22
7.20
6.70
6.53
5.12
5.11
4.56
4.53
4.46
4.44
4.42
4.40
4.35
4.24
4.22
4.20
4.19
3.66
3.55
3.53
3.42
3.35
3.34
3.31
3.30
2.88
2.83
2.44
2.30
2.16
2.03
2.00
1.93
1.92
1.91
1.90
1.88
1.66
1.44
1.42
1.00
0.97
0.97
1.03
1.05
1.13
5.25
1.12
1.02
1.02
0.99
3.27
2.10
1.13
1.16
2.08
2.94
2.38
1.94
3.31
5.82
2.30
3.21
1.26
1.29
1.16
1.12
2.32
3.30
11.23
13.5 13.0 12.5 12.0 11.5 11.0 10.5 10.0 9.5 9.0 8.5 8.0 7.5 7.0 6.5 6.0 5.5 5.0 4.5 4.0 3.5 3.0 2.5 2.0 1.5 1.0 0.5 0.0
f1 (ppm)

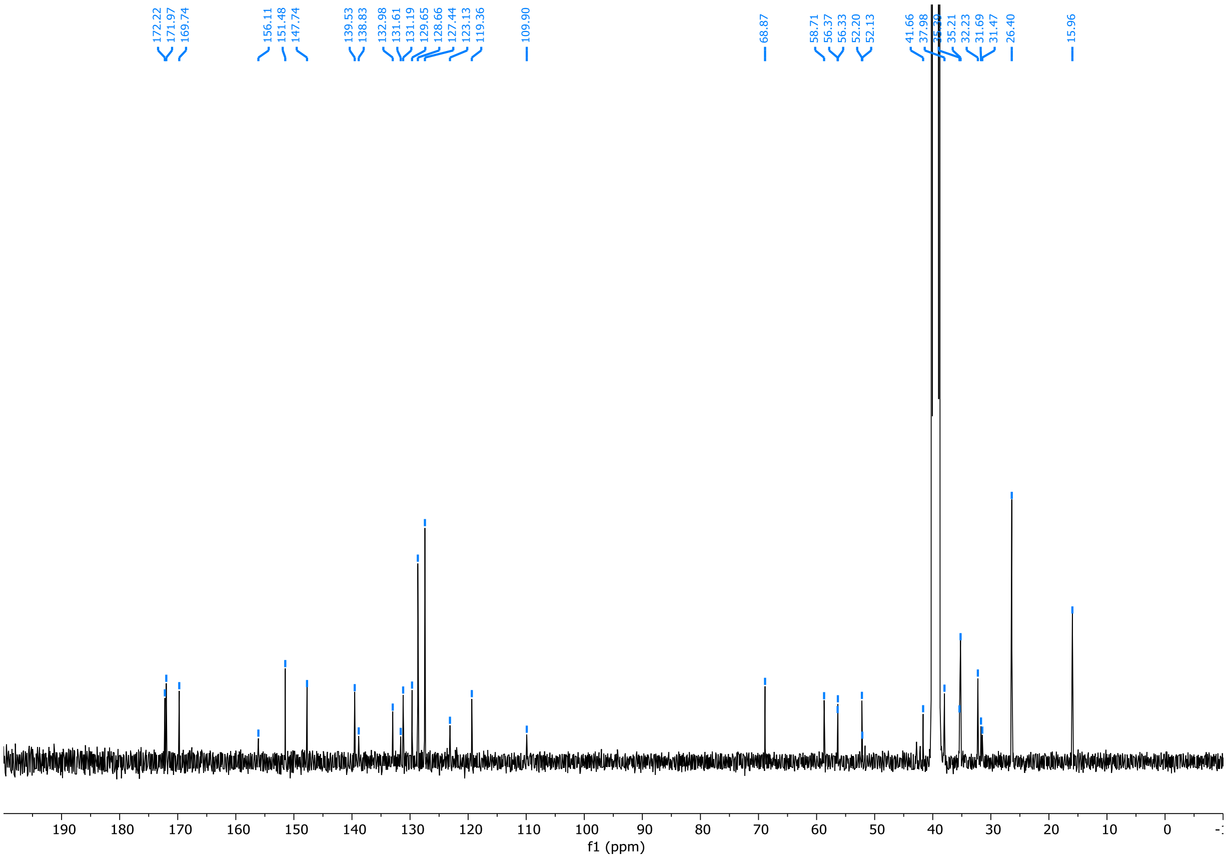
172.22
171.97
169.74
156.11
151.48
147.74
139.53
138.83
132.98
131.61
131.19
129.65
128.66
127.44
123.13
119.36
109.90
68.87
58.71
56.37
56.33
52.20
52.13
41.66
37.98
35.21
32.23
31.69
31.47
26.40
15.96
190 180 170 160 150 140 130 120 110 100 90 80 70 60 50 40 30 20 10 0
f1 (ppm)

**CIP-19**
**(DMSO)**

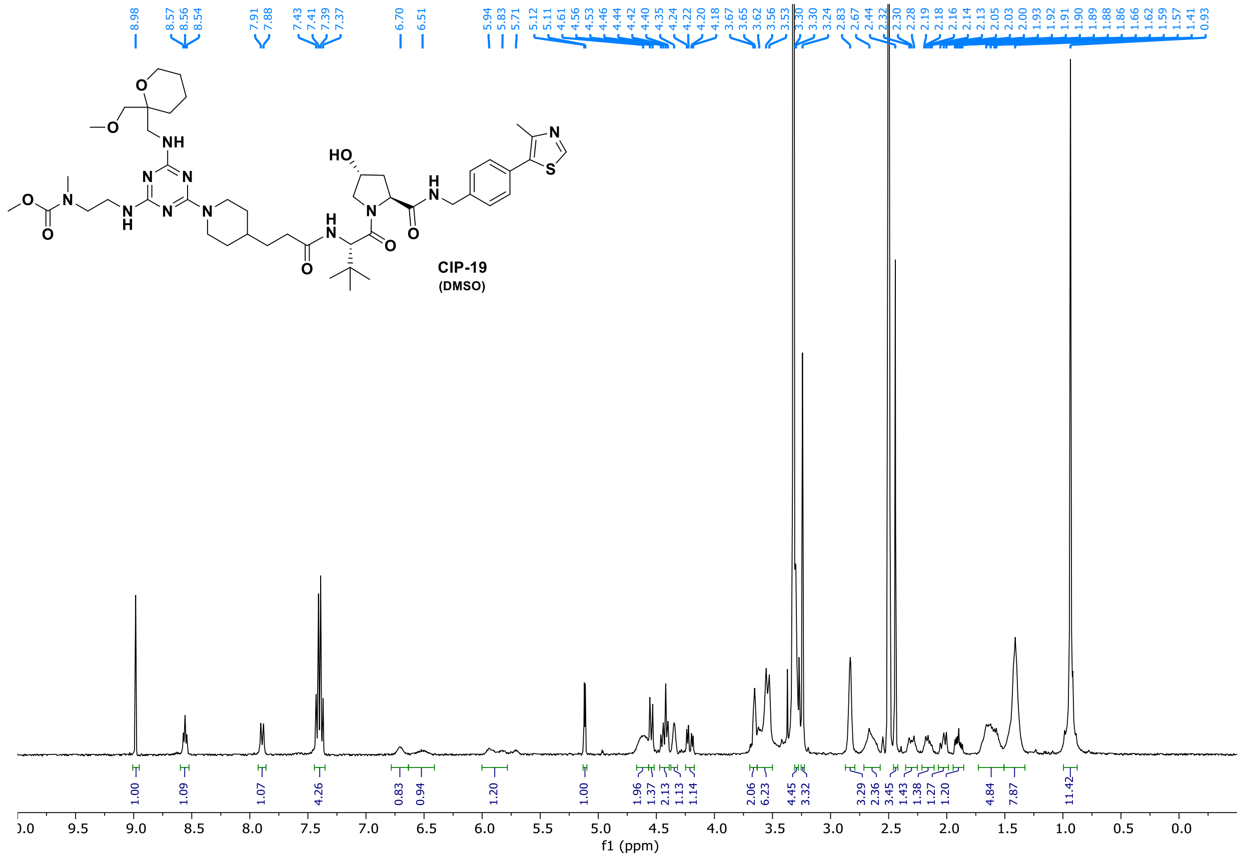
8.98
8.57
8.56
8.54
7.91
7.88
7.43
7.41
7.39
7.37
6.70
6.51
5.94
5.83
5.71
5.12
5.11
4.61
4.56
4.53
4.46
4.44
4.42
4.40
4.35
4.24
4.22
4.20
4.18
3.67
3.65
3.62
3.56
3.53
3.30
3.30
3.24
2.83
2.67
2.44
2.32
2.30
2.28
2.19
2.18
2.16
2.14
2.13
2.05
2.03
2.00
1.93
1.92
1.91
1.90
1.89
1.86
1.66
1.62
1.59
1.57
1.41
0.93
1.00
1.09
1.07
4.26
0.83
0.94
1.20
1.00
1.96
1.37
2.13
1.13
1.14
2.06
6.23
4.45
3.32
3.29
2.36
3.45
1.43
1.38
1.27
1.20
4.84
7.87
11.42
f1 (ppm)

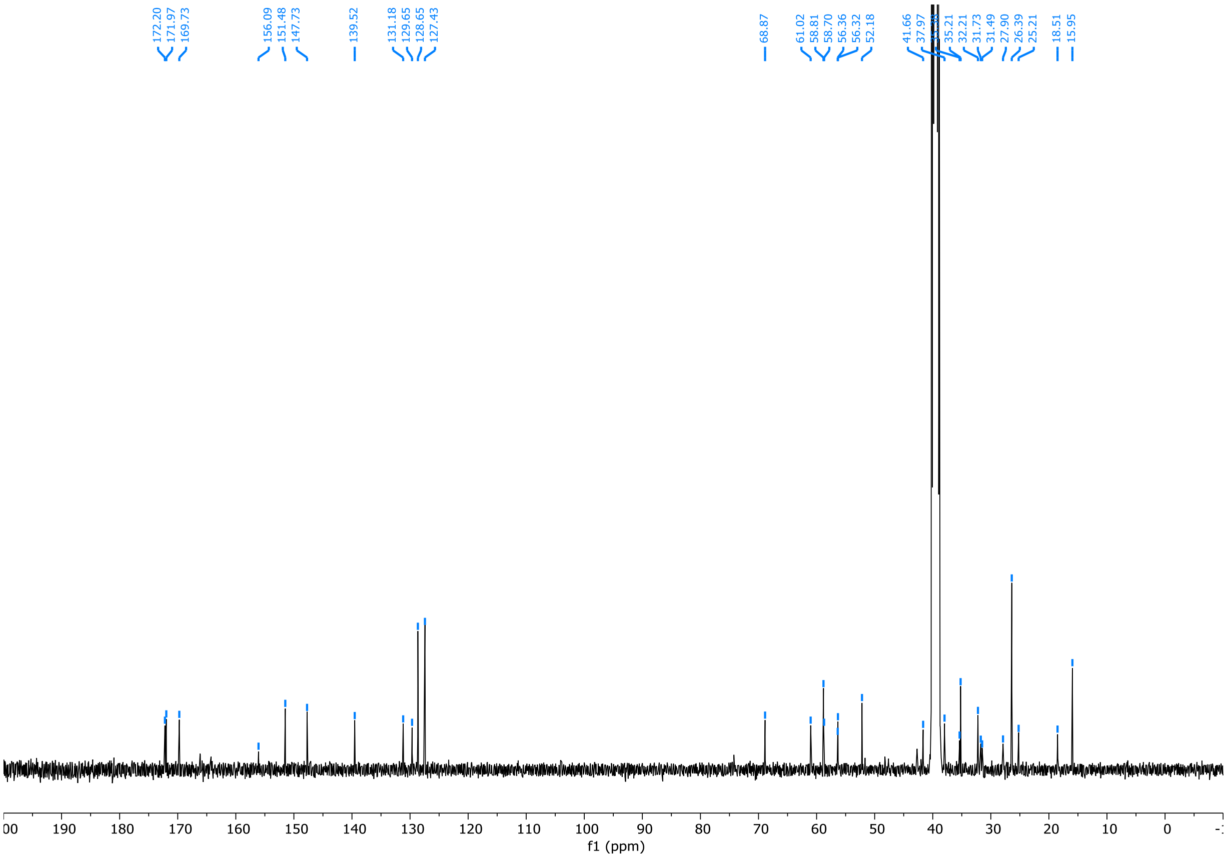
172.20
171.97
169.73
156.09
151.48
147.73
139.52
131.18
129.65
128.65
127.43
68.87
61.02
58.81
58.70
56.36
56.32
52.18
41.66
37.97
35.21
32.21
31.73
31.49
27.90
26.39
25.21
18.51
15.95
f1 (ppm)

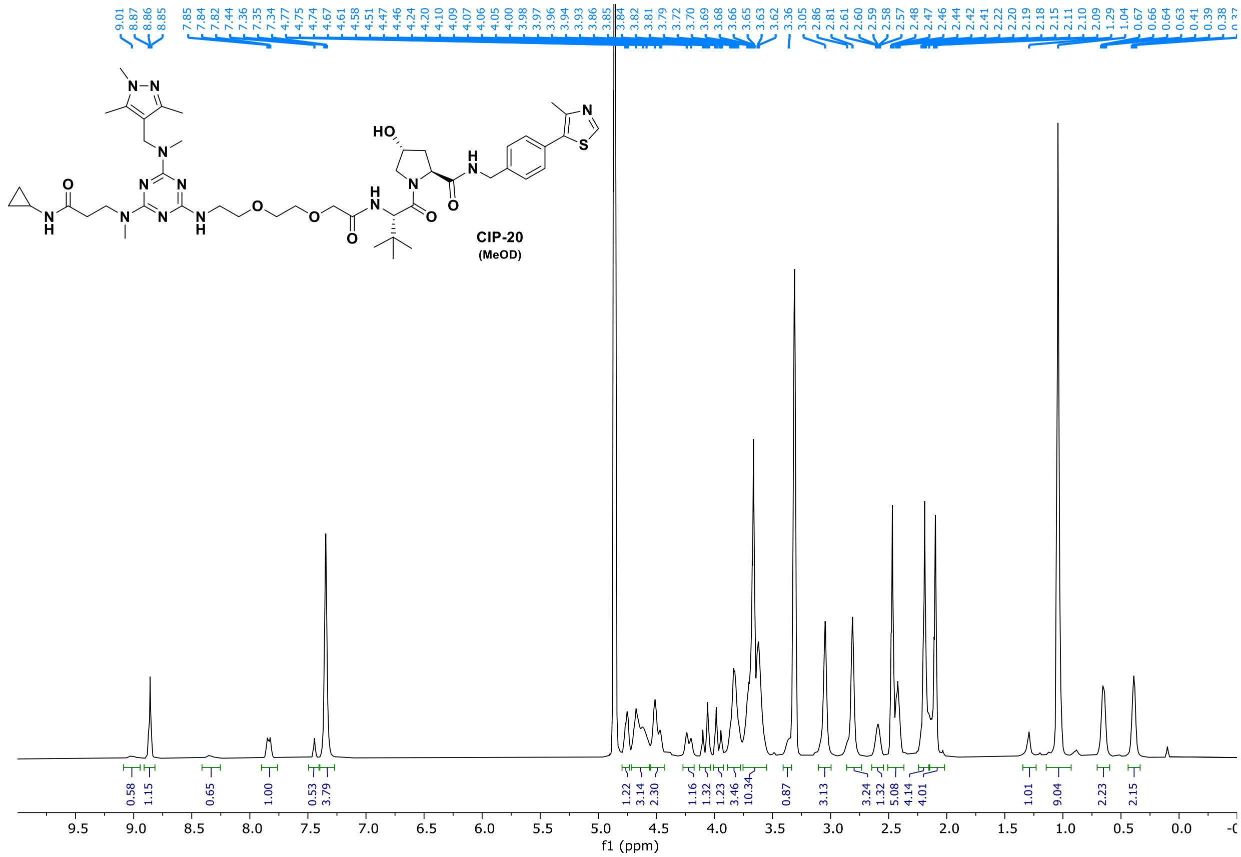

0.58 1.15 0.65 1.00 0.53 3.79 1.22 3.14 2.30 1.16 1.32 1.23 3.46 10.34 0.87 3.13 3.24 1.32 5.08 4.14 4.01 1.01 9.04 2.23 2.15
f1 (ppm)


**CIP-20**
**(MeOD)**

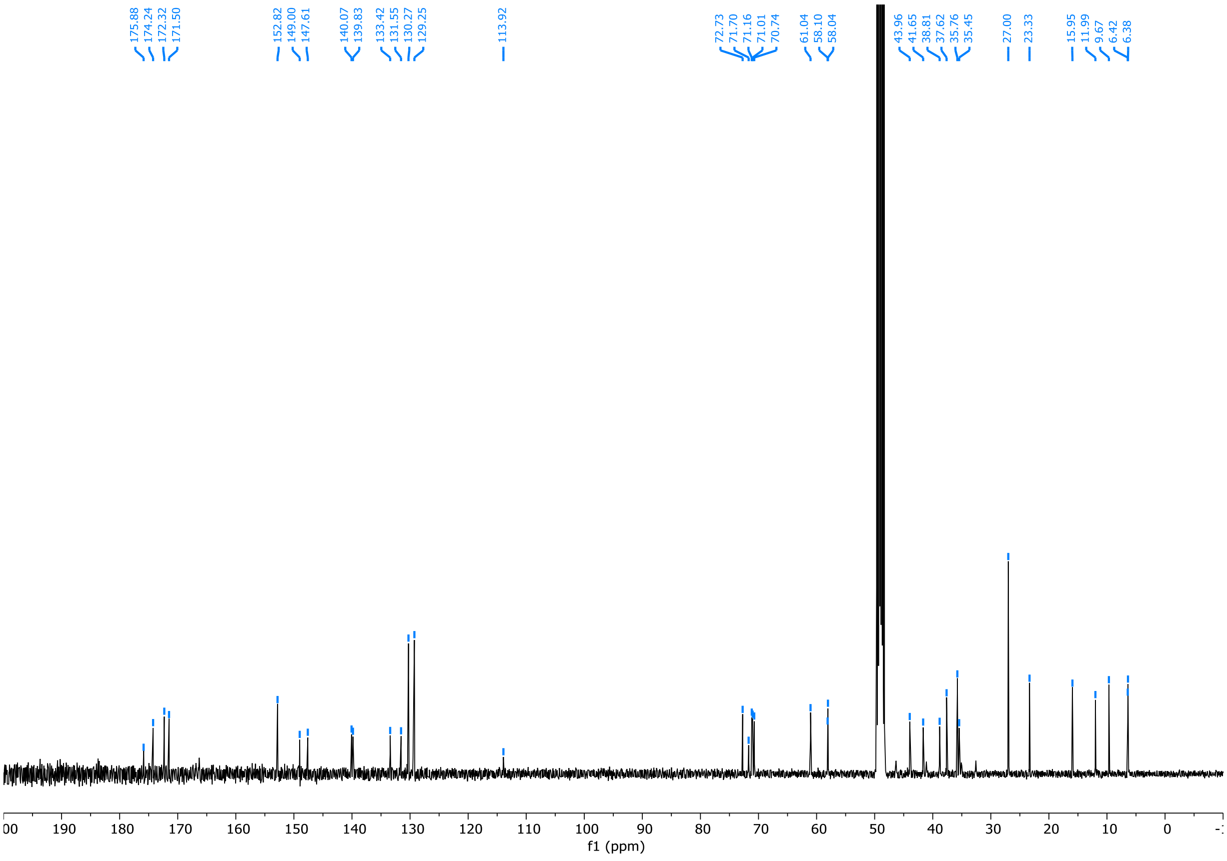

175.88 174.24 172.32 171.50 152.82 149.00 147.61 140.07 139.83 133.42 131.55 130.27 129.25 113.92 72.73 71.70 71.16 71.01 70.74 61.04 58.10 58.04 43.96 41.65 38.81 37.62 35.76 35.45 27.00 23.33 15.95 11.99 9.67 6.42 6.38
f1 (ppm)

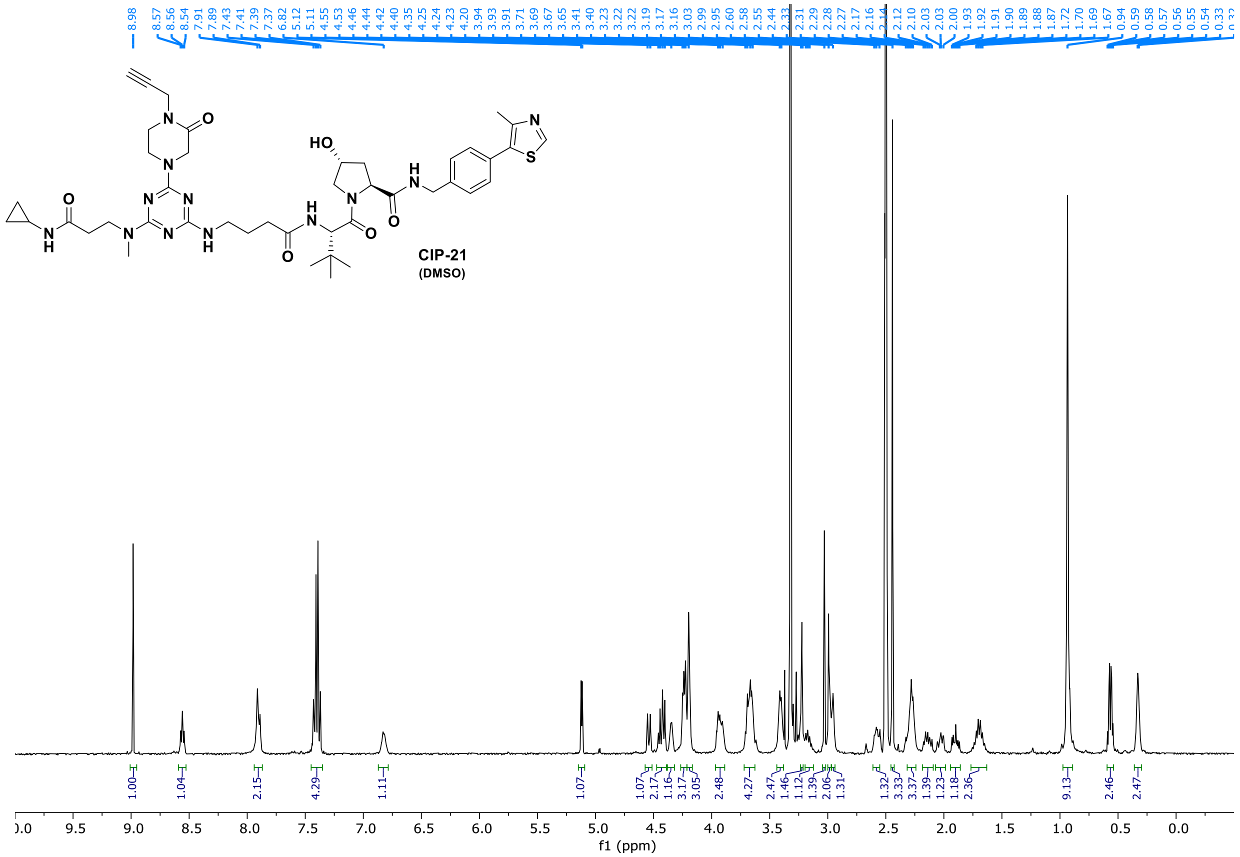
CIP-21
(DMSO)
f1 (ppm)

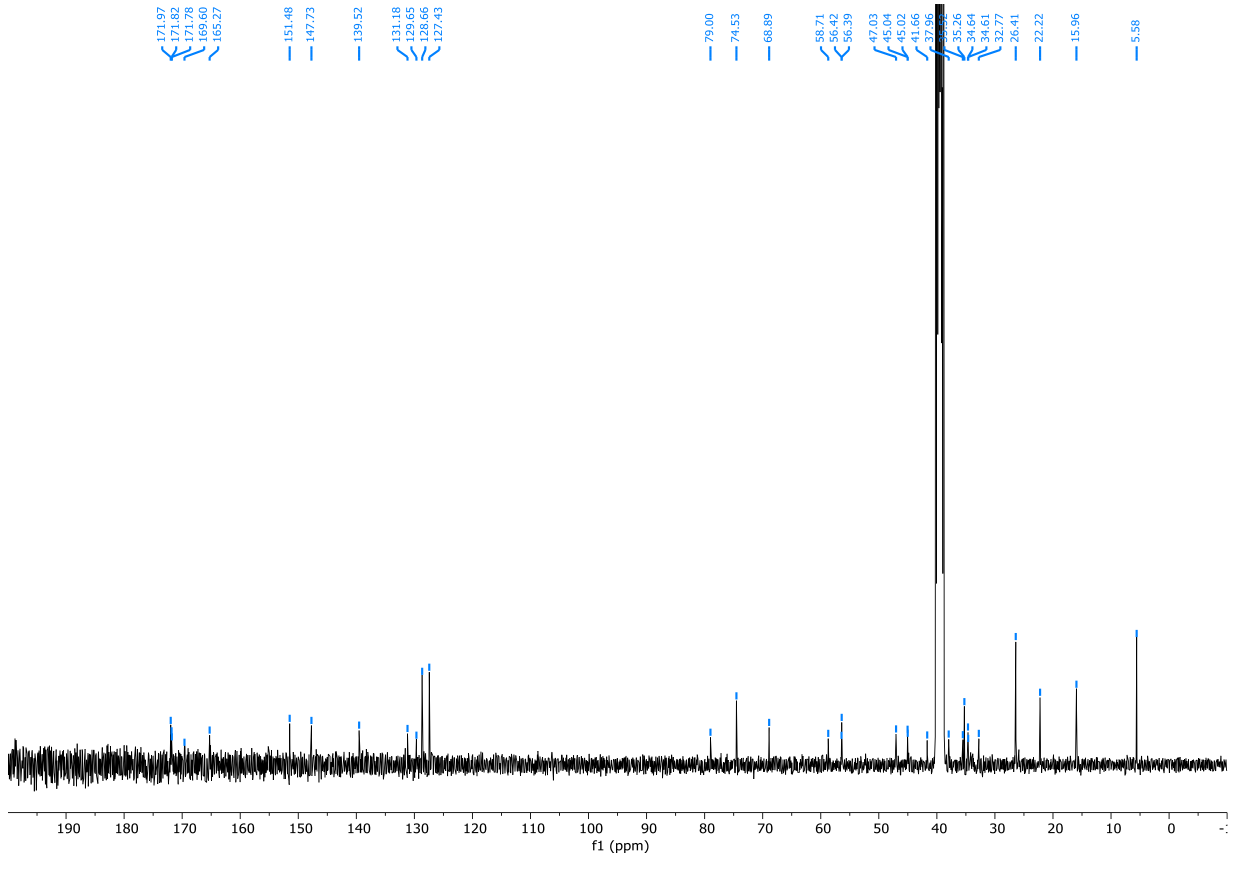
f1 (ppm)

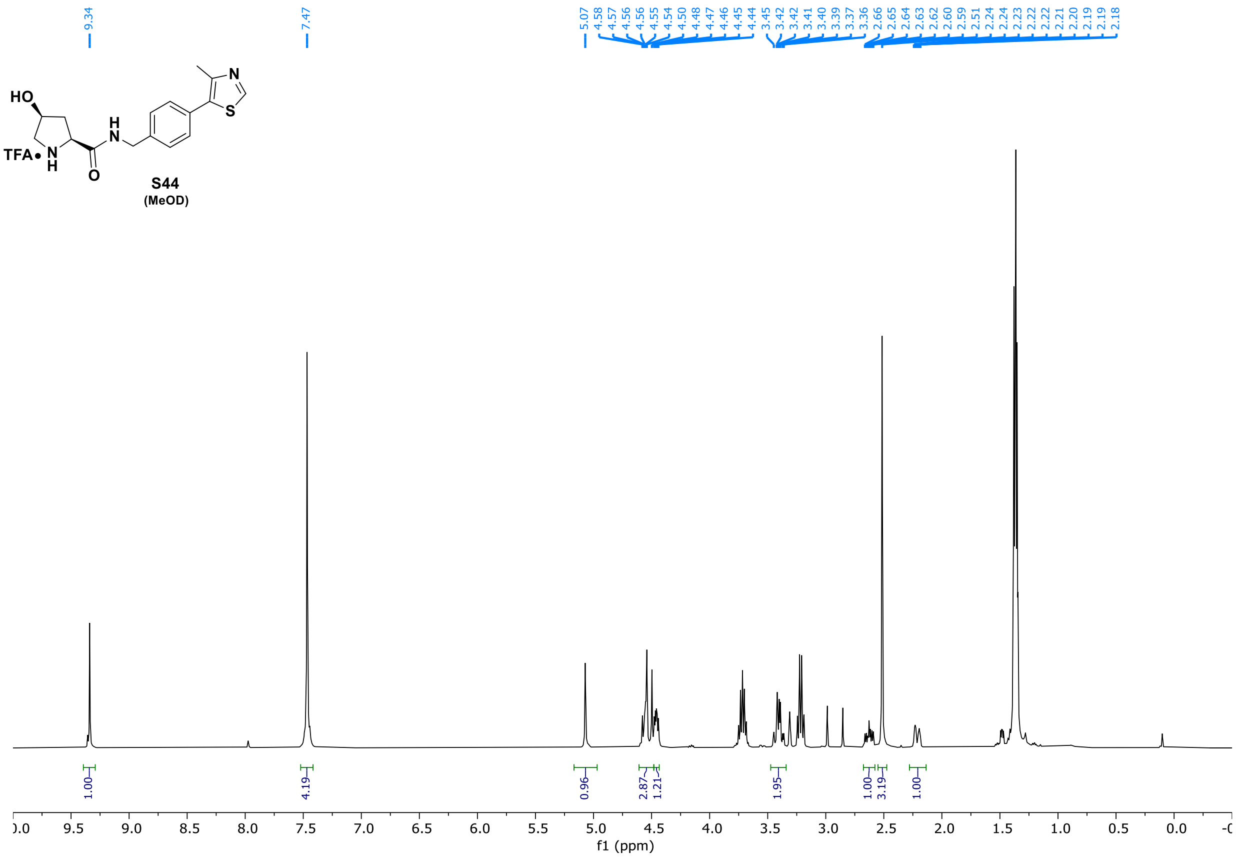
9.34
7.47
5.07
4.58
4.57
4.56
4.56
4.55
4.54
4.50
4.48
4.47
4.46
4.45
4.44
3.45
3.42
3.42
3.41
3.40
3.39
3.37
3.36
2.66
2.65
2.64
2.63
2.62
2.60
2.59
2.51
2.24
2.24
2.23
2.22
2.22
2.21
2.20
2.19
2.19
2.18
HO
N
S
TFA•
H
N
H
O
1.00
4.19
0.96
2.87
1.21
1.95
1.00
3.19
1.00
f1 (ppm)

**S44**
(MeOD)

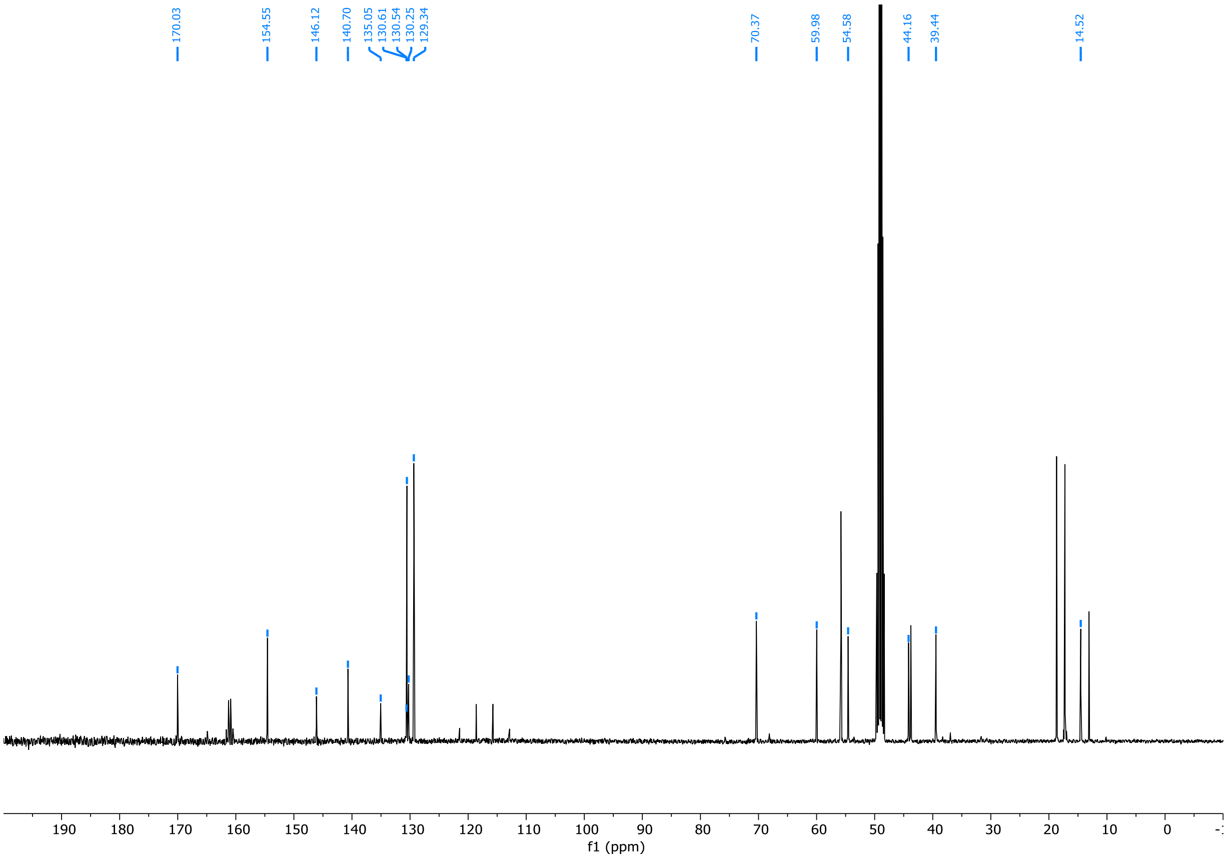
170.03
154.55
146.12
140.70
135.05
130.61
130.54
130.25
129.34
70.37
59.98
54.58
44.16
39.44
14.52
f1 (ppm)

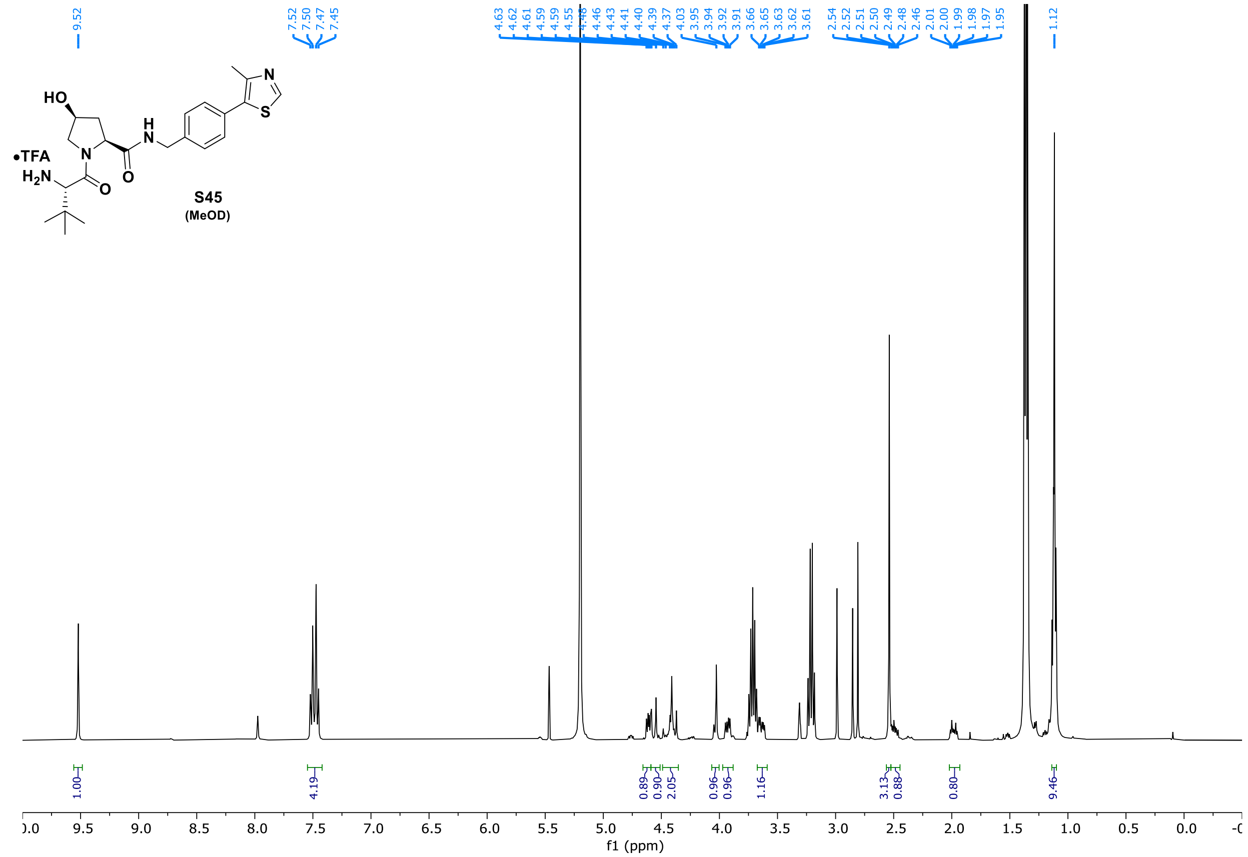
HO
N
H
N
N
S
•TFA
$H_2N$
O
O
9.52
7.52
7.50
7.47
7.45
4.63
4.62
4.61
4.59
4.59
4.55
4.48
4.46
4.43
4.41
4.40
4.39
4.37
4.03
3.95
3.94
3.92
3.91
3.66
3.65
3.63
3.62
3.61
2.54
2.52
2.51
2.50
2.49
2.48
2.46
2.01
2.00
1.99
1.98
1.97
1.95
1.12
1.00
4.19
0.89
0.90
2.05
0.96
0.96
1.16
3.13
0.88
0.80
9.46
f1 (ppm)

**S45**
(MeOD)

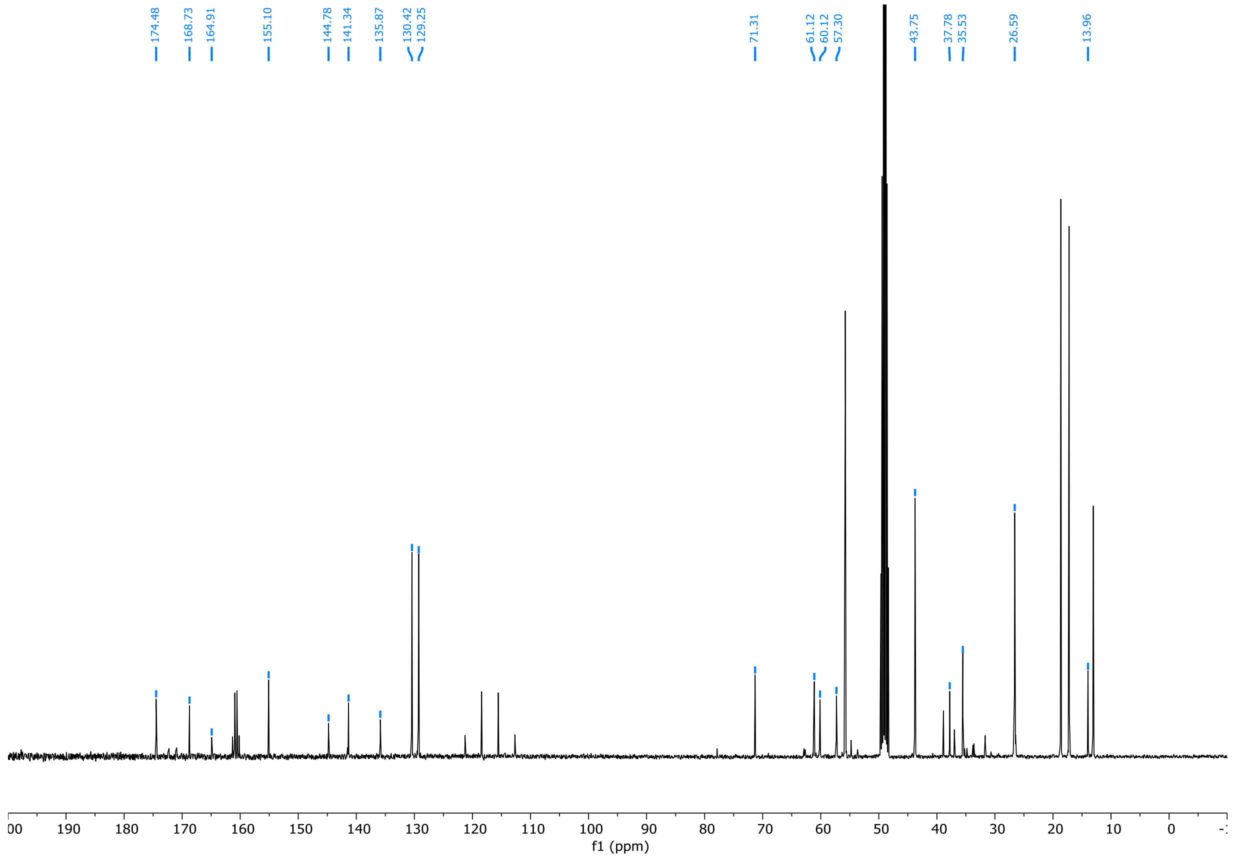
174.48
168.73
164.91
155.10
144.78
141.34
135.87
130.42
129.25
71.31
61.12
60.12
57.30
43.75
37.78
35.53
26.59
13.96
f1 (ppm)

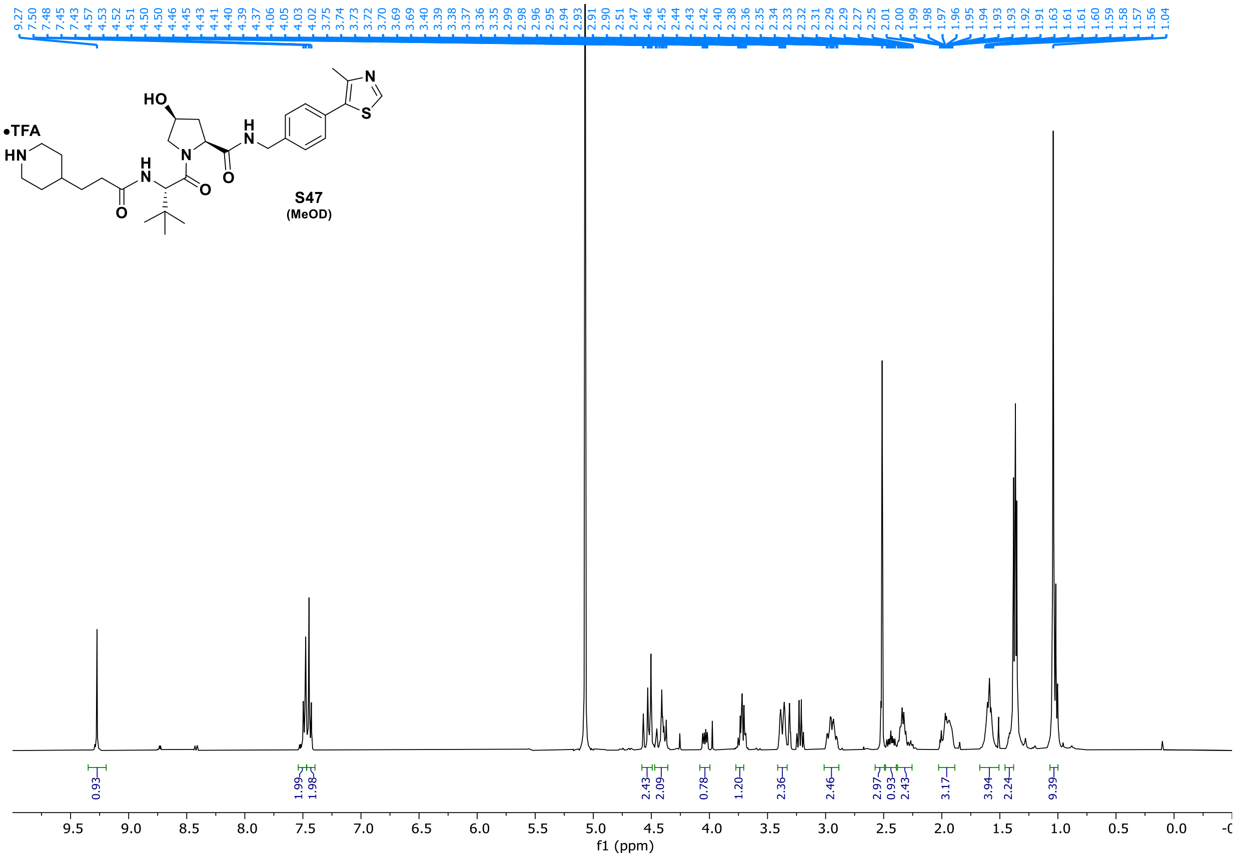
•TFA
HN
HO
H
N
N
H
N
O
O
O
N
S
S47
(MeOD)
9.27 7.50 7.48 7.45 7.43 4.57 4.53 4.52 4.51 4.50 4.50 4.46 4.45 4.43 4.41 4.40 4.39 4.37 4.06 4.05 4.03 4.02 3.75 3.74 3.73 3.72 3.70 3.69 3.69 3.40 3.39 3.38 3.37 3.36 3.35 2.99 2.98 2.96 2.95 2.94 2.93 2.91 2.90 2.51 2.47 2.46 2.45 2.44 2.43 2.42 2.40 2.38 2.36 2.35 2.34 2.33 2.32 2.31 2.29 2.29 2.27 2.25 2.01 2.00 1.99 1.98 1.97 1.96 1.95 1.94 1.93 1.93 1.92 1.91 1.63 1.61 1.61 1.60 1.59 1.58 1.57 1.56 1.04
0.93 1.99 1.98 2.43 2.09 0.78 1.20 2.36 2.46 2.97 0.93 2.43 3.17 3.94 2.24 9.39
9.5 9.0 8.5 8.0 7.5 7.0 6.5 6.0 5.5 5.0 4.5 4.0 3.5 3.0 2.5 2.0 1.5 1.0 0.5 0.0
f1 (ppm)

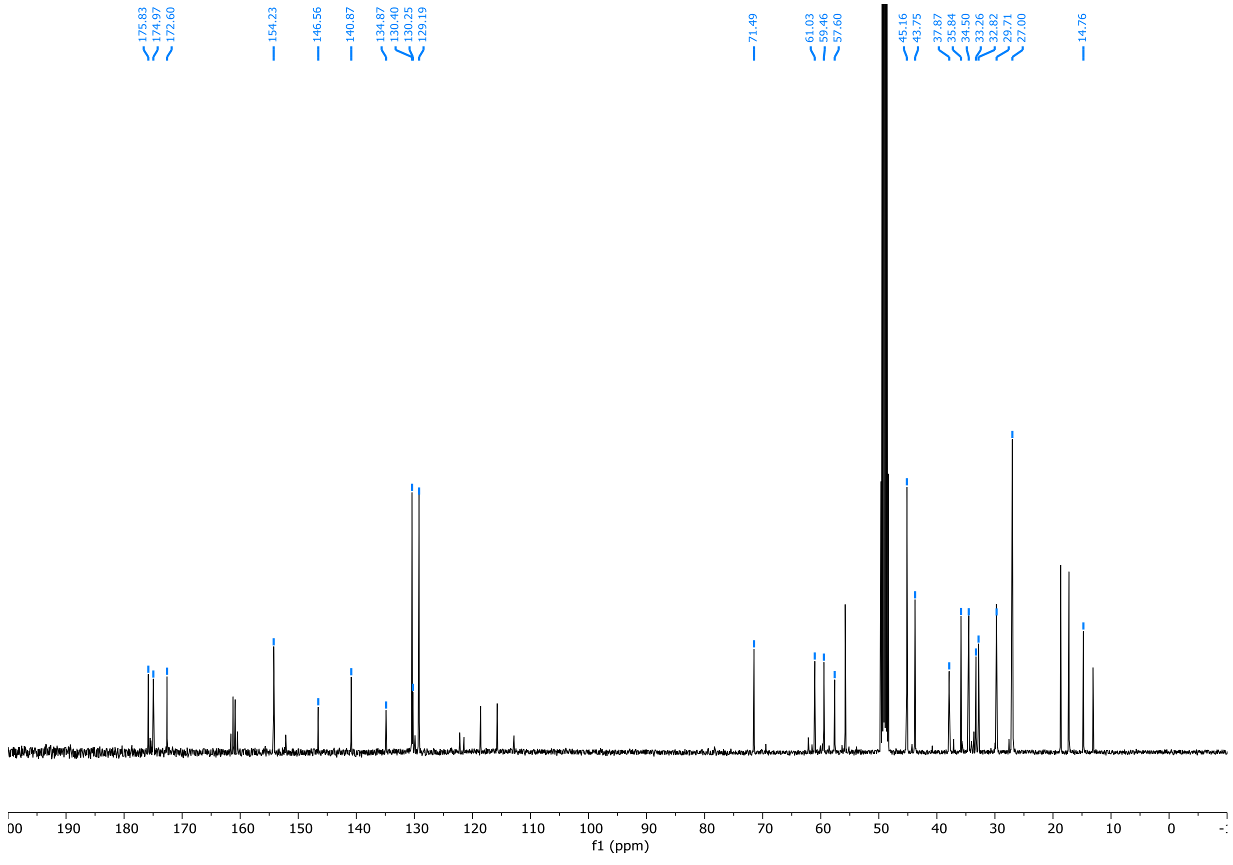
175.83 174.97 172.60 154.23 146.56 140.87 134.87 130.40 130.25 129.19 71.49 61.03 59.46 57.60 45.16 43.75 37.87 35.84 34.50 33.26 32.82 29.71 27.00 14.76
190 180 170 160 150 140 130 120 110 100 90 80 70 60 50 40 30 20 10 0
f1 (ppm)

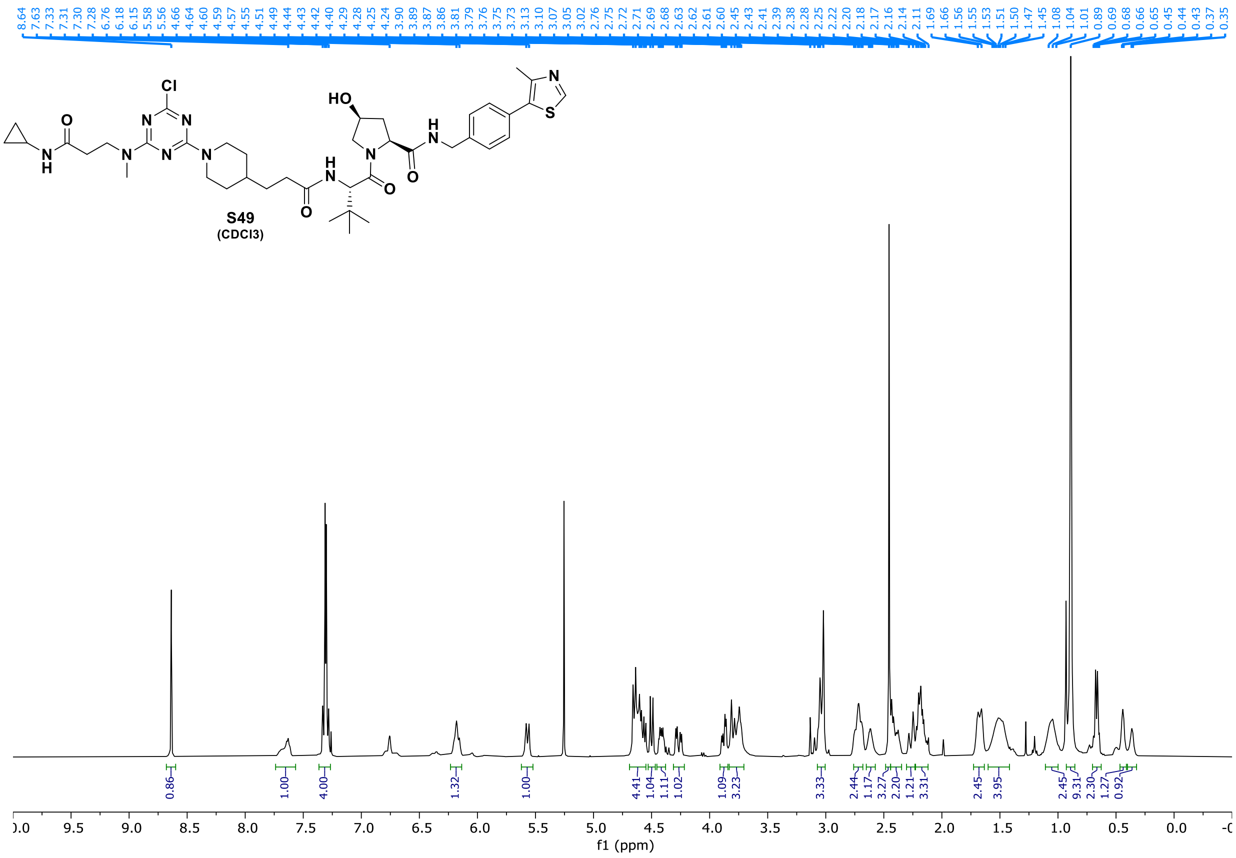
8.64 7.63 7.33 7.31 7.30 7.28 6.76 6.18 6.15 5.58 5.56 4.66 4.64 4.60 4.59 4.57 4.55 4.51 4.49 4.44 4.43 4.42 4.40 4.29 4.28 4.25 4.24 3.90 3.89 3.87 3.86 3.81 3.79 3.76 3.75 3.73 3.13 3.10 3.07 3.05 3.02 2.76 2.75 2.72 2.71 2.69 2.68 2.63 2.62 2.61 2.60 2.45 2.43 2.41 2.39 2.38 2.28 2.25 2.22 2.20 2.18 2.17 2.16 2.14 2.11 1.69 1.66 1.56 1.55 1.53 1.51 1.50 1.47 1.45 1.08 1.04 1.01 0.89 0.69 0.68 0.66 0.65 0.45 0.44 0.43 0.37 0.35
Cl
N
HO
N
N
N
N
H
N
N
O
N
H
S
O
N
H
O
N
O
S49
(CDCl3)
0.86 1.00 4.00 1.32 1.00 4.41 1.04 1.11 1.02 1.09 3.23 3.33 2.44 1.17 3.27 2.20 1.21 3.31 2.45 3.95 2.45 9.31 2.30 1.27 0.92
9.5 9.0 8.5 8.0 7.5 7.0 6.5 6.0 5.5 5.0 4.5 4.0 3.5 3.0 2.5 2.0 1.5 1.0 0.5 0.0
f1 (ppm)

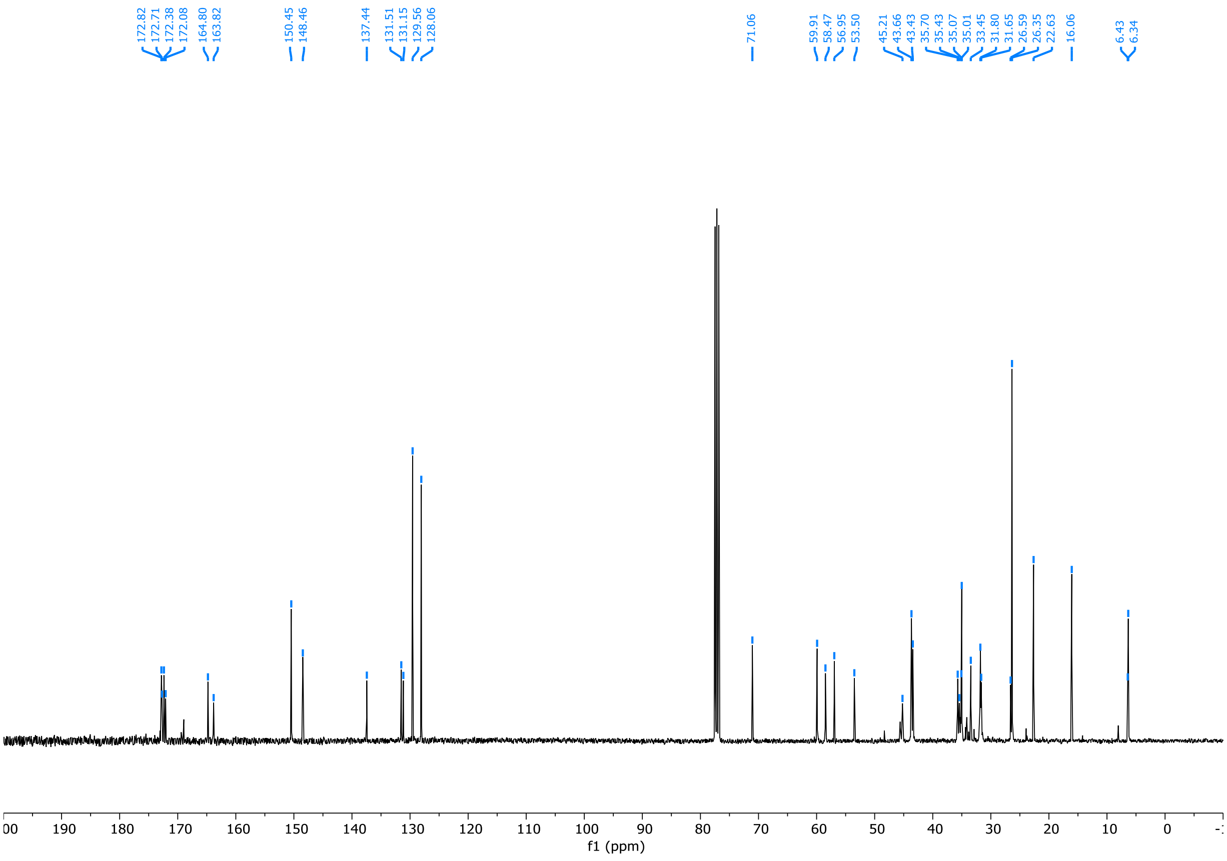
172.82 172.71 172.38 172.08 164.80 163.82 150.45 148.46 137.44 131.51 131.15 129.56 128.06 71.06 59.91 58.47 56.95 53.50 45.21 43.66 43.43 35.70 35.43 35.07 35.01 33.45 31.80 31.65 26.59 26.35 22.63 16.06 6.43 6.34
190 180 170 160 150 140 130 120 110 100 90 80 70 60 50 40 30 20 10 0
f1 (ppm)

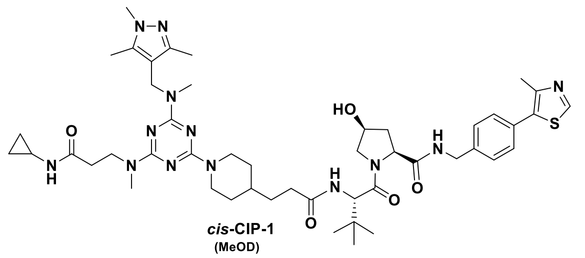


*cis*-CIP-1
(MeOD)

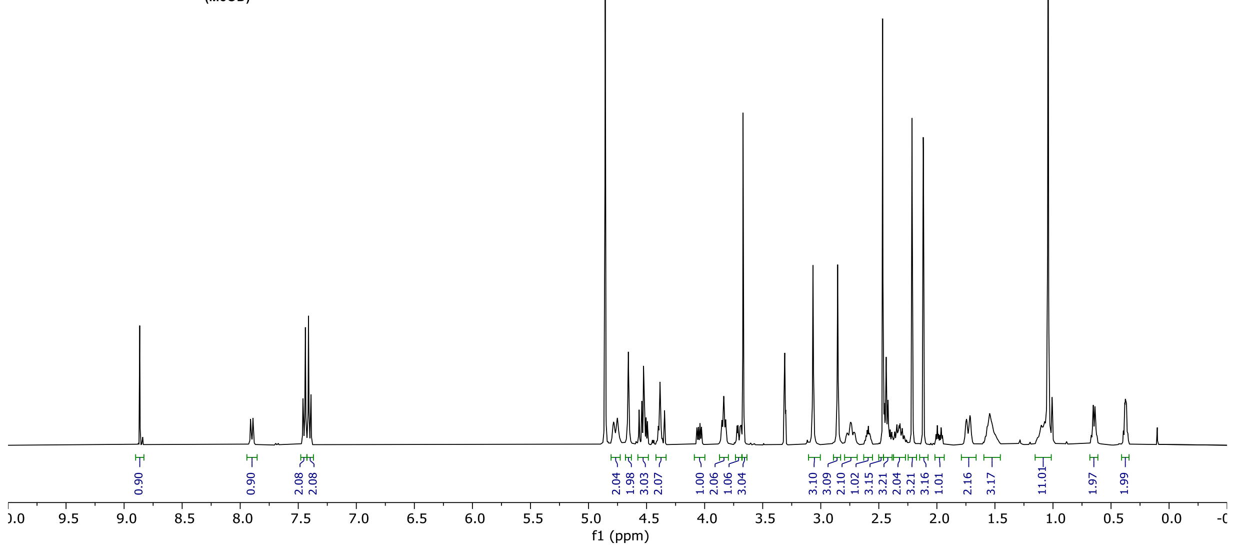


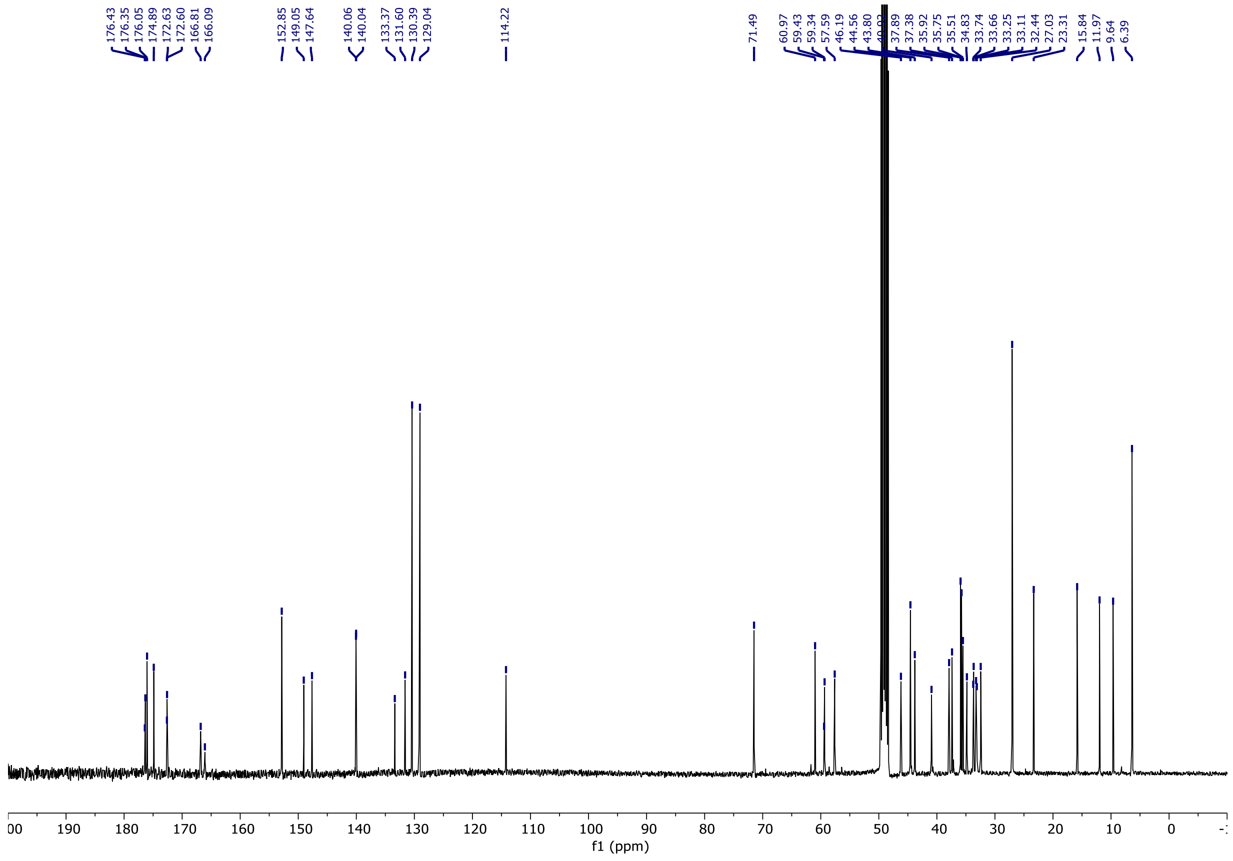

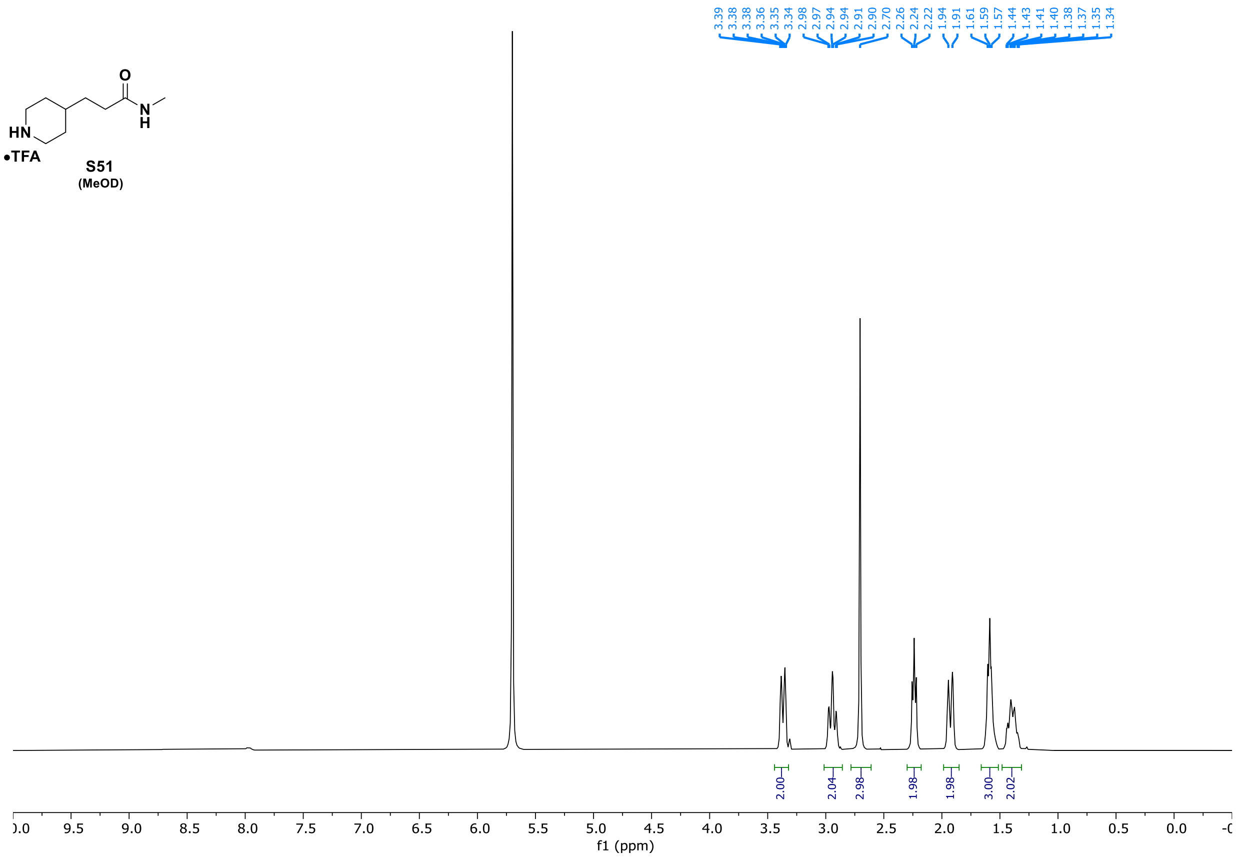
O
HN
N
H
•TFA
S51
(MeOD)
3.39 3.38 3.38 3.36 3.35 3.34 2.98 2.97 2.94 2.94 2.91 2.90 2.70 2.26 2.24 2.22 1.94 1.91 1.61 1.59 1.57 1.44 1.43 1.41 1.40 1.38 1.37 1.35 1.34
2.00 2.04 2.98 1.98 1.98 3.00 2.02
f1 (ppm)

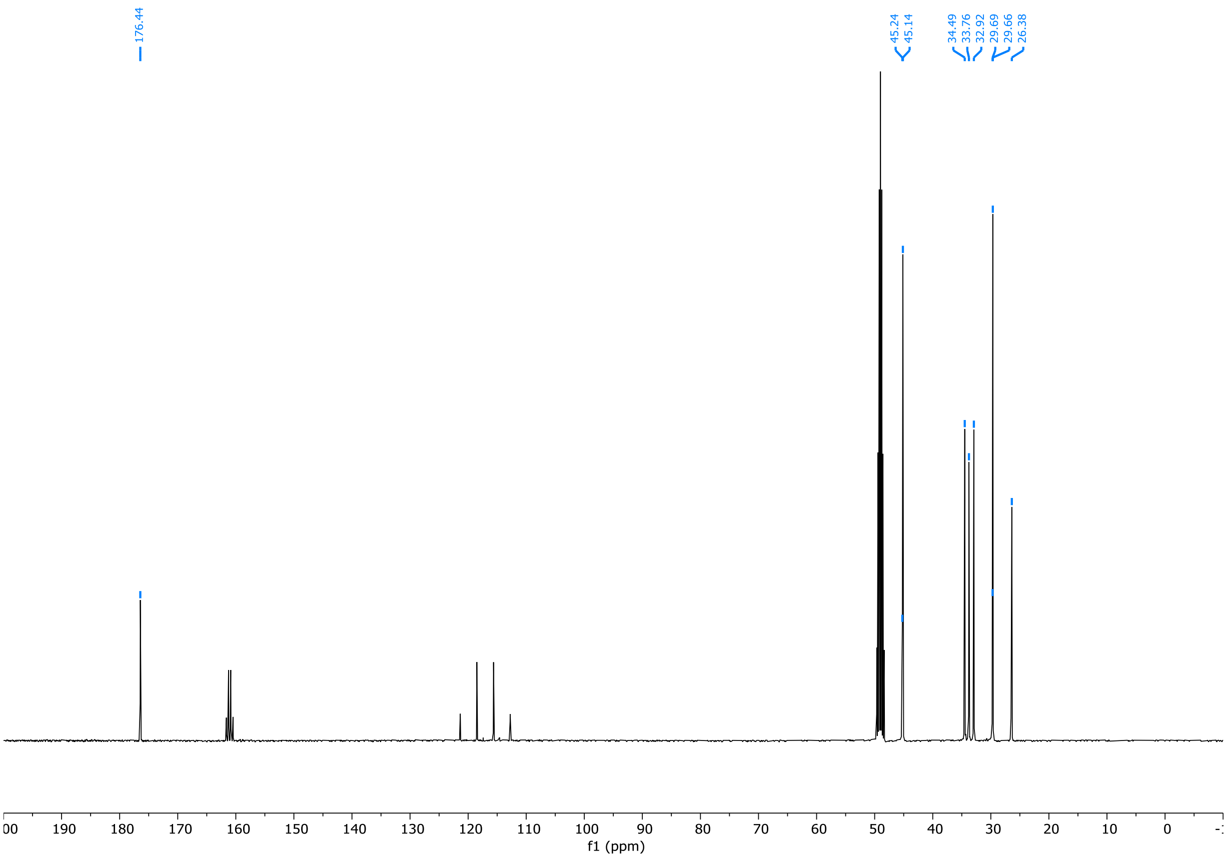
176.44
45.24 45.14
34.49 33.76 32.92 29.69 29.66 26.38
f1 (ppm)

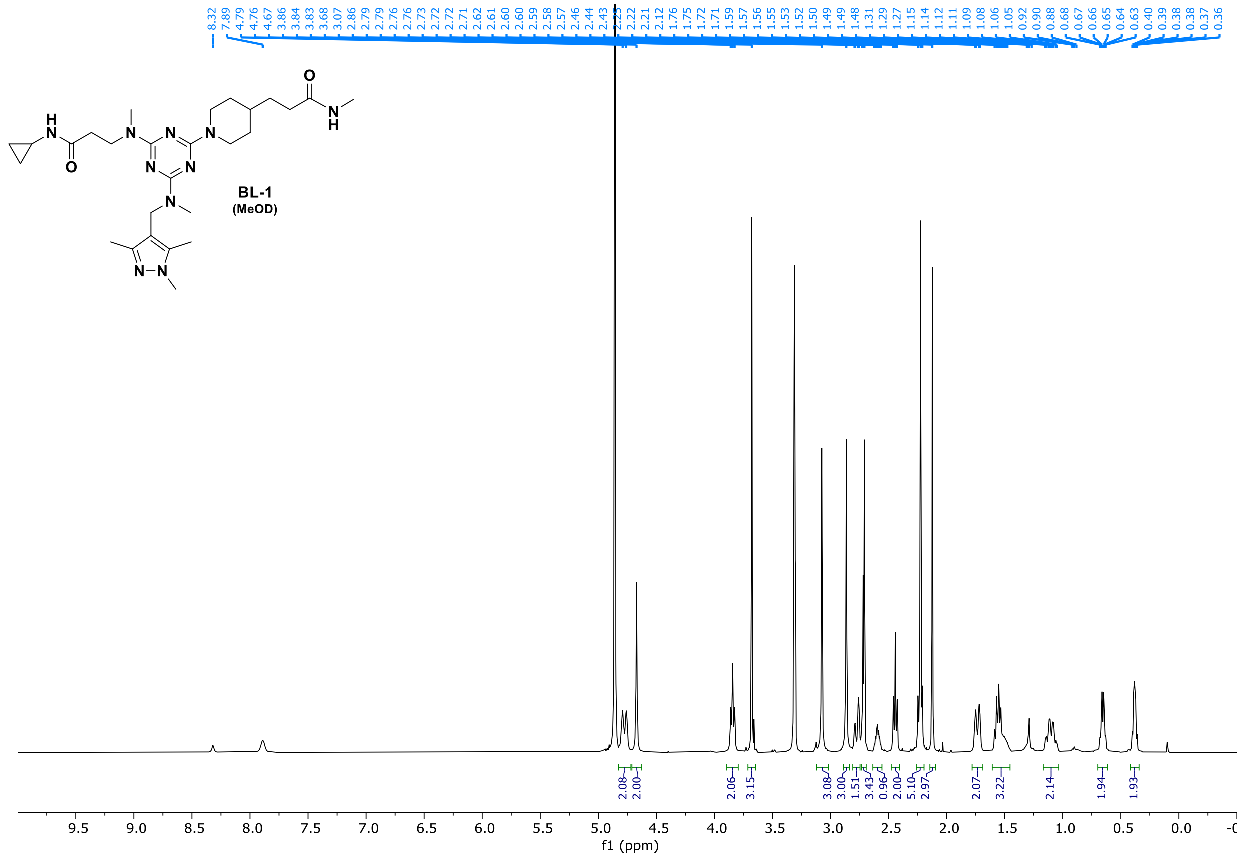


**BL-1**
(MeOD)

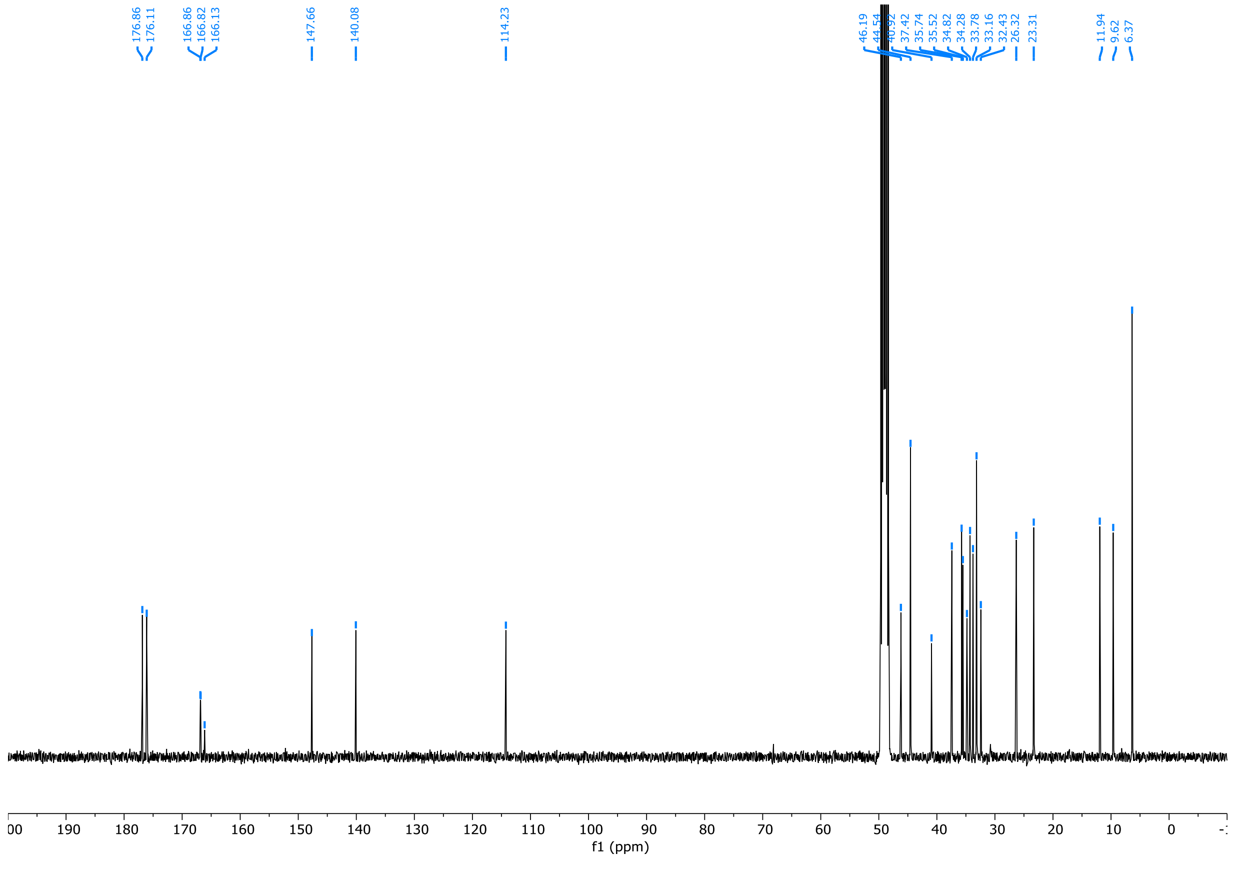
